# Interlaboratory comparison of standardised metabolomics and lipidomics analyses in human and rodent blood using the MxP® Quant 500 kit

**DOI:** 10.1101/2024.11.13.619447

**Authors:** Gözde Ertürk Zararsiz, Jutta Lintelmann, Alexander Cecil, Jennifer Kirwan, Gernot Poschet, Hagen M. Gegner, Sven Schuchardt, Xue Li Guan, Daisuke Saigusa, David Wishart, Jiamin Zheng, Rupasri Mandal, Kendra Adams, J. Will Thompson, Michael P. Snyder, Kevin Contrepois, Songjie Chen, Nadia Ashrafi, Sumeyya Akyol, Ali Yilmaz, Stewart F. Graham, Thomas M. O’Connell, Karel Kalecký, Teodoro Bottiglieri, Alice Limonciel, Hai Tuan Pham, Therese Koal, Jerzy Adamski, Gabi Kastenmüller

## Abstract

Metabolomics and lipidomics are pivotal in understanding phenotypic variations beyond genomics. However, quantification and comparability of mass spectrometry (MS)-derived data are challenging. Standardised assays can enhance data comparability, enabling applications in multi-center epidemiological and clinical studies. Here we evaluated the performance and reproducibility of the MxP® Quant 500 kit across 14 laboratories. The kit allows quantification of 634 different metabolites from 26 compound classes using triple quadrupole MS. Each laboratory analysed twelve samples, including human plasma and serum, lipaemic plasma, NIST SRM 1950, and mouse and rat plasma, in triplicates. 505 out of the 634 metabolites were measurable above the limit of detection in all laboratories, while eight metabolites were undetectable in our study. Out of the 505 metabolites, 412 were observed in both human and rodent samples. Overall, the kit exhibited high reproducibility with a median coefficient of variation (CV) of 14.3 %. CVs in NIST SRM 1950 reference plasma were below 25 % and 10 % for 494 and 138 metabolites, respectively. To facilitate further inspection of reproducibility for any compound, we provide detailed results from the in-depth evaluation of reproducibility across concentration ranges using Deming regression. Interlaboratory reproducibility was similar across sample types, with some species-, matrix-, and phenotype-specific differences due to variations in concentration ranges. Comparisons with previous studies on the performance of MS-based kits (including the AbsoluteIDQ p180 and the Lipidyzer) revealed good concordance of reproducibility results and measured absolute concentrations in NIST SRM 1950 for most metabolites, making the MxP® Quant 500 kit a relevant tool to apply metabolomics and lipidomics in multi-center studies.

## INTRODUCTION

Metabolomics plays a crucial role in unraveling the molecular basis of phenotypes, complementing information from other omics layers. For instance, while genomics explains only 3 % of the variance in a population’s body mass index (BMI), metabolomics can account for 50 % of the observed variance (1), capturing environmental factors such as nutrition (2, 3), lifestyle (4–7), medication (8, 9), as well as intrinsic factors like aging and hormonal status (10–13). To enable large-scale studies for identifying reliable biomarkers of diseases and their progression and ultimately for developing potential companion diagnostics, metabolomics approaches need to be accessible (technology availability), comprehensive (high metabolite coverage), reproducible (robust analytical performance), sustainable (facilitating reanalysis), and scalable (parallel assays in independent units). Despite the implementation of stable isotopically labeled metabolites as valuable internal standards throughout the community, accurate quantification and comparability of mass spectrometry-derived metabolomics data remain major challenges. One approach to mitigate these challenges is the use of standardised kits that enable scientists to compare their data independently of their analytical platforms (14, 15).

To test the reproducibility and scalability of metabolomics analyses using such kits, various ring trials (16, 17) or interlaboratory studies (18) have been conducted worldwide. These initiatives have provided crucial insights into the accessibility of metabolomics analytics, thereby supporting epidemiological study design (19), clinical translational projects (20), and the development of diagnostic clinical procedures (21).

In this study, we aimed to assess the metabolite coverage and performance of the MxP® Quant 500 kit, specifically applied to human and rodent blood plasma samples. We sought to investigate the reproducibility of quantitative measurements across 14 distinct laboratories worldwide utilizing different instrument setups to ultimately determine the feasibility of multi-center studies using the MxP® Quant 500 kit.

## MATERIALS AND METHODS

### MxP® Quant 500 kit

The MxP® Quant 500 kit (biocrates life sciences ag, Innsbruck, Austria; later abbreviated as biocrates) was utilised as a ready-to-use method according to the manufacturer’s instructions for targeted metabolite quantification in 10 µl liquid biopsy samples (e.g., plasma or serum).

#### Metabolite coverage

Depending on the mass spectrometer used, the method targets up to 628 metabolites (Waters instruments) and 630 metabolites (Sciex instruments), thereof 624 metabolites are detectable on both instruments. The metabolites DG-O(16:0_20:4), PC aa C30:2, PC aa C38:1, SM C22:3, TG(14:0_39:3), and TG(20:1_31:0) are exclusively detected using Sciex mass spectrometers while DG-O(18:2_18:2) and HexCer(d16:1/20:0), TG(14:0_40:5), and TG(20:1_32:0) are only detected using Waters mass spectrometers. This sums up to 634 different metabolites from 26 compound classes covered by the kit. Detection and quantification of metabolites are carried out by triple quadrupole mass spectrometry (QqQ-MS or MS/MS). Out of the 634 metabolites, 106 metabolites (alkaloids, amine oxides, amino acids, amino acids related, bile acids, biogenic amines, carboxylic acids, cresols, fatty acids, hormones and related, indoles and derivatives, nucleobases and related, vitamins and cofactors) are quantified via separation by ultra-high-performance chromatography (UHPLC) before the MS (LC-MS/MS) analysis. The remaining 528 metabolites (carbohydrates and related, acylcarnitines, lysophosphatidylcholines, phosphatidylcholines, sphingomyelins, ceramides, dihydroceramides, hexosylceramides, dihexosylceramides, trihexosylceramides, cholesteryl esters, diacylglycerols, and triacylglycerols) are analysed via flow injection analysis (FIA-MS/MS). Analyte-specific multiple reaction monitoring (MRM) is used for quantification. The principal workflow is based on a patented 96-well plate format and consists of a derivatisation step, the extraction of the analytes, and the LC- and FIA-MS/MS determination. More detailed information on covered metabolites is given in **Supplementary Table S1**.

#### Measurement standardisation and kit software

All MxP® Quant 500 kits are delivered together with a standard operating procedure (SOP) containing detailed information for standardised sample preparation, instrument set-up, MS measurement (including system suitability testing), and data processing. The biocrates proprietary software (MetIDQ at the time of this study) is also an integral part of the kit and must be installed before beginning with kit preparation. The software is used for controlling the standardised kit workflow, performing the quantitation, and checking the technical validity of the kit analysis. The kit includes a reagent set including human plasma quality control (QC) samples (anticoagulant: EDTA) representing low, medium, and high, partially spiked concentration levels, standard solutions for system suitability tests for LC and FIA, a 7-point calibrator set samples for LC, as well as mobile phase additives for FIA solvents, and two deep well plates with silicone mats for extract dilution.

#### Metabolite nomenclature

In this manuscript, we adopted the same metabolite nomenclature and abbreviations as used by the manufacturer at the time of measurement. Thereby, the short names for ceramides, cholesteryl esters, diacylglycerols, and triacylglycerols largely follow the standard lipid nomenclature (22). Phosphatidylcholines are denoted as lysoPC a Cx:y (lysophosphatidylcholines), PC aa Cx:y and PC ae Cx:y (diacyl- and acyl-alkyl-phosphatidylcholines, respectively), where x represents the total number of carbon atoms and y the number of double bonds in the fatty acid side chains. Note: In contrast to the LC-MS/MS part of the analysis, which allows for the separation of isobaric and, to some extent, of constitutional and cis-trans stereoisomers, the FIA of hexoses and lipids does not use pre-MS/MS UHPLC separation of metabolites. In combination with the unit resolution of triple quadrupole mass spectrometers, the selectivity for the metabolites determined by FIA is decreased. As a consequence, FIA analysis cannot provide specific information regarding the bond types (acyl (ester) or alkyl (ether)) or positions or chain lengths of the fatty acid residues linked to each lipid’s backbone. For all FIA lipids, the detected signal is thus the sum of several isobaric/isomeric lipids, and the lipid name assigned is only one potential representative of an - in some cases - large number of possible isobars and isomers. As an example, in contrast to what PC aa and PC ae labels suggest, the measurement cannot differentiate between acyl- and alkyl bonds (or any other possible isobar); these labels were chosen under the assumption that fatty acids with even chain lengths are more common/abundant (23); similarly, sphingomyelins are labeled assuming d18:1 for the fatty acid moiety in the amide bond (i.e., SM C18:0 corresponds to SM 36:1). A further elucidation of the potential mixtures and/or an improved assignment of the lipids was not a goal of this study. For phosphatidylcholines, more details on the constituents of the measured sums in human plasma can be found in (23). To come closer to recent developments in the nomenclature of lipid species by the LIPID MAPS community (22), a new metabolite nomenclature was released by biocrates and is now applied in the latest version of the kit software (WebIDQ). Both nomenclatures are available in **Supplementary Table S1**. For the present study, the 26 compound classes as listed by the manufacturer were further aggregated into 11 classes to facilitate visualisations in the figures. Specifically, the classes alkaloids, carbohydrates and related, carboxylic acids, cresols, hormones and related, indoles and derivatives, nucleobases and related, vitamins and cofactors, of which some contained only one metabolite, were summarised in the group “Others”, comprising in total 21 metabolites. Also, all ceramides (ceramides, dihydroceramides, hexosylceramides, dihexosylceramides, trihexosylceramides) were assigned to a single class “Ceramides” (n = 71); phosphatidylcholines and lysophosphatidylcholines were summarised as “Glycerophospholipids” (n = 90); amino acids, amino acids related, amine oxides, and biogenic amines were grouped into a single class termed “Amino Acids and Amines” (n = 60).

### Samples for interlaboratory comparison

A set of twelve blinded plasma and serum samples (3 x 60 µl per sample) was provided by biocrates to each laboratory as an interlaboratory project sample set (**Figure 1**). The sample set contained 1 x NIST SRM 1950, 3 x individual male human plasma samples, 3 x individual female human plasma samples, 1 x individual male human serum sample, 1 x individual female human serum sample, 1 x lipaemic human plasma pool, 1 x mouse plasma pool, 1 x rat plasma pool. Human plasma/serum was collected and aliquoted by in.vent Diagnostika GmbH (Berlin, Germany). Informed consent for blood collection was requested and obtained from each individual, and the documents are available on request. The NIST SRM 1950 sample was purchased from the National Institute of Standards and Technology (NIST, Gaithersburg, USA) and aliquoted at biocrates. The pooled mouse and rat plasma samples were bought from Sera Laboratories International Ltd (West Sussex, United Kingdom) and aliquoted at biocrates. All samples were blinded for the measurements, meaning that the samples were numbered from 1 to 12 and the underlying identification was not disclosed to the participants of the interlaboratory comparison.

**Figure 1.**
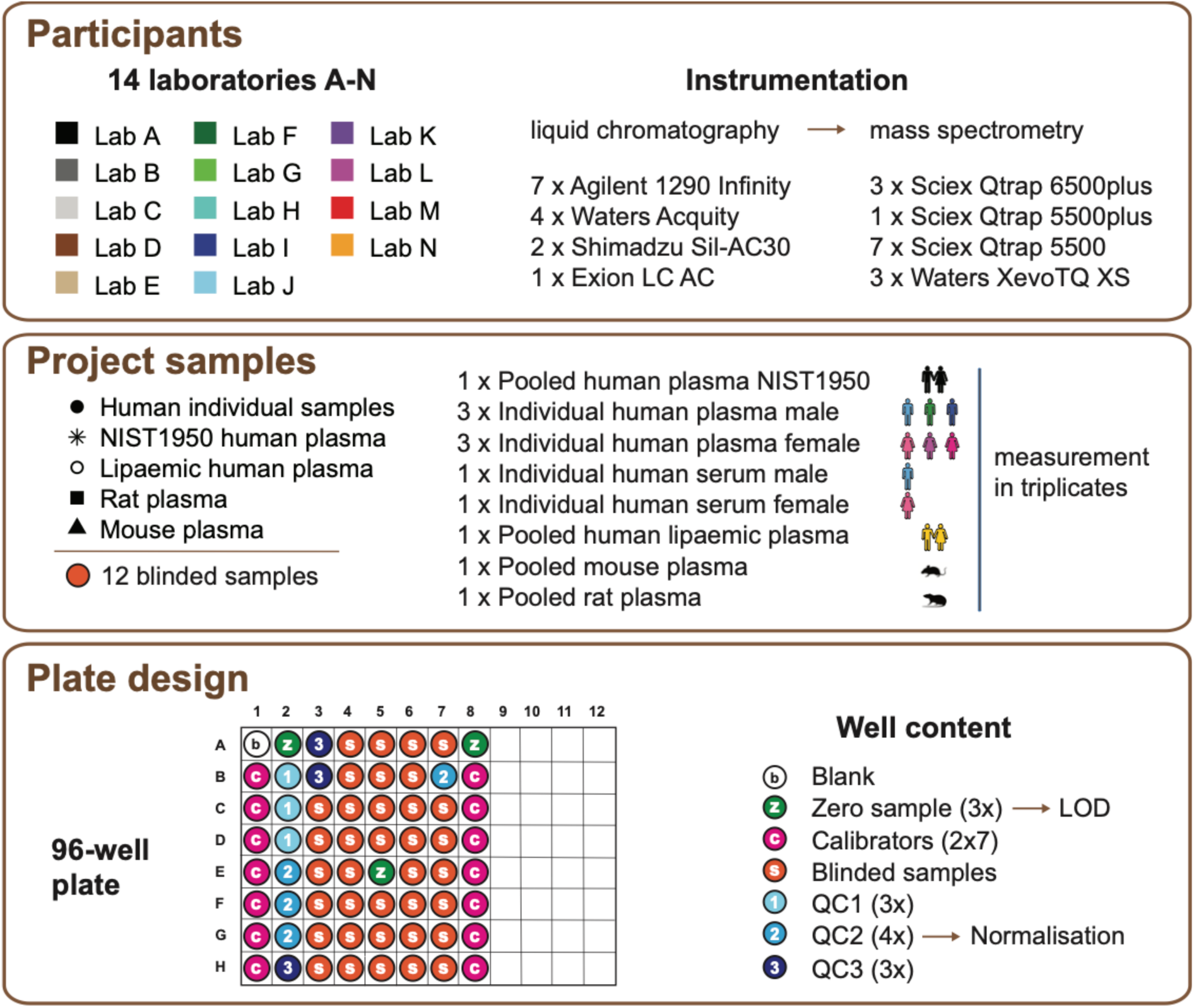
Graphical overview of participating laboratories, project samples, and plate design. Each participating laboratory is denoted by a unique letter and color code used throughout this manuscript. Analogously, project sample types are denoted by different symbols.

### Instrumentation for interlaboratory comparison

The participating laboratories used different UHPLC-MS/MS setups (**Figure 1**); for UHPLC, setups included Agilent 1290 Infinity, Waters Acquity, Shimadzu Sil-AC30, or Exion LC AC systems; for MS/MS analysis, Sciex Qtrap 5500/5500plus/6500plus or Waters XevoTQ XS instruments were used.

### Analytical procedure within the present study

An overview of the complete workflow of the interlaboratory comparison is given in Supplementary Figure S1.

#### Materials

Biological (project) samples and MxP® Quant 500 kits were sent to the participants on dry ice shortly before the scheduled measurements. In addition to the SOPs, each laboratory received a detailed technical guide containing further information on instrument requirements, cleaning instructions, plate layout of project samples, data processing of raw data, and data upload. The project sample set was analysed in each laboratory following a predefined 96-well plate layout (1 x blank, 3 x zero samples (only internal standards), 2 sets of 7 calibrators each (beginning and end of the plate), 3 x QC1, QC3, 4 x QC2, 3 x 12 project samples) (**Figure 1**). Required materials not included in the kit, namely methanol, ethanol, acetonitrile, 2-propanol, ammonium acetate, formic acid, phosphate-buffered saline (PBS), pyridine, phenylisothiocyanate (PITC), water, and gases, were purchased independently by each laboratory and had at least LC-MS grade purity.

#### System suitability testing

The performances of the individual UHPLC-MS/MS systems in the laboratories were checked by system suitability tests (SST), which had to be performed by injecting and controlling test samples for FIA- and LC-MS/MS in each laboratory before starting with kit preparation and measurement of the project samples. SST raw data files were submitted to biocrates for review and approval.

#### Sample preparation

After SST approval, sample preparation was carried out in strict accordance with the SOP delivered with the kit. Briefly, calibrators and QC samples were prepared according to protocol, and 10 µl of all samples (including QC samples, calibrators, zero samples, and the project samples) were pipetted onto the filter inserts of the 96-well plate following the predefined plate layout. The filters of the kit plate contained all internal standards and thus no further pipetting step was needed. The samples were dried under a gentle stream of nitrogen for 30 min and subsequently derivatised with 50 µl of a PITC solution at room temperature for 60 min. Then, the derivatised samples were dried for 60 min, again under a gentle stream of nitrogen. Metabolites were extracted by applying 300 µl of a solution of 5 mM ammonium acetate in methanol to each well and shaking the plate for 30 min at 450 rpm. The plate was centrifuged to collect the extracts through the filter inserts. Extracts were diluted with either running solvent (1:50, methanol with mobile phase additive provided by biocrates) for FIA, or with water (1:1) for LC-MS/MS analysis.

#### FIA and LC-MS/MS

Instrument-specific ready-to-use and validated analytical methods for FIA- and UHPLC-MS/MS data acquisition were provided with the kit. The diluted sample extracts were analysed twice in positive ion FIA-MS/MS and twice in UHPLC-MS/MS mode (one in positive ion mode, one in negative ion mode) by (overall four) different instrument-specific methods. UHPLC-separations were performed on a kit-specific UHPLC column applying gradients from 100 % water with 0.2 % formic acid to 100 % acetonitrile with 0.2 % formic acid at 0.8 ml/min and 50 °C. The injection volume was 5 µl, the autosampler temperature was 10 °C and the total LC-MS/MS analysis time was approximately 6 min per injection. FIA-MS/MS analysis was performed with an injection volume of 20 µl. The diluted sample extracts were injected into the mass spectrometer with methanol including the FIA mobile phase additive at 0.03 ml/min. The total FIA-MS/MS run time was approximately 3 min per injection. Mass spectrometric detection and quantification for FIA- and LC-MS/MS was carried out based on analyte-specific mass transitions in the MRM mode.

#### Raw data processing

Raw data files were imported into the kit software (Version “Oxygen-DB110-3005”) and processed by following the protocol provided in the user manual (UM-MetIDQ-Oxygen-11). Firstly, retention times of all LC-peaks were adjusted using QC2 runs in positive and negative ion mode. Then, the integration of every peak in every sample was checked manually and corrected, if necessary. Afterward, calibration curves of metabolites with 7-point calibration were inspected. Quadratic regression with 1/x weighting was used for the calculation of the calibration curves according to the technical guide provided by the manufacturer. The laboratories were advised to remove calibration points if they were significantly out of range. Compliance of the curve fits (regression coefficient R^2^) ≥ 0.98 was checked. After plate validation and approval, the LC-MS/MS data were ready for export (csv- and xlsx-files). FIA data was automatically processed, quantified, and validated in the kit software. The concentration unit of the results was µmol/l. Quantification was achieved on different quality levels (7-point calibration using calibrators or 1-point calibration using internal standard concentrations). The median value of the zero samples (PBS in three wells of the plate) was calculated for each metabolite as an approximation of the background noise. The limit of detection (LOD) was defined as three times this median for each analyte. In case of missing signal intensities in all zero samples, LOD values calculated during the method validation were automatically implemented by the kit software.

#### Data collection and preparation

Participating laboratories were assigned random identities depicted by letters A-N. The letter coding was disclosed to corresponding laboratories but not to other participants of this project. Each laboratory put together a package containing the following data and files: the kit software export *.metidq project file and *.xlsx- and *.csv-result files as QC2-normalised data and result files without normalisation of any type; *xlsx or *csv files with retention time data. Additionally, the raw data files of system suitability test samples and all samples measured on the plate were included. These packages were pseudonymised and collected at the secure data repository of Helmholtz Zentrum München. Individual *.metidq projects were imported into a single MetIDQ repository, inspected for data consistency and verified for quality of calibration curves, peak integration and QC-matching, and the data were exported without normalisation of any type. One laboratory could repeat the measurements with an identical set of samples due to technical issues discovered in this step. The *.csv files were harmonised concerning layout, number of rows and columns (columns for metabolites not measured by Waters/Sciex instruments were added, respectively) to facilitate further data evaluation.

### Data analysis

All submitted data sets from 14 laboratories were collected and statistically analysed at the Helmholtz Zentrum München. All measurements below the laboratory-specific LODs were ignored, i.e., set to “missing” in the data set for all described analyses unless explicitly stated otherwise.

#### Metabolite coverage

For the analysis of metabolite coverage, the sources of “missingness” (e.g., LOD, not having passed laboratory-specific analytical QC) were not differentiated, i.e., we use the term “missingness” for every metabolite and its concentration value that could not be identified during the analyses. We considered a metabolite as detected in a laboratory if at least one aliquot of at least one project sample showed a value in the data table of this laboratory. To investigate whether missingness was linked to LOD levels, i.e., level of noise and impurities in zero samples, we ranked laboratories by their calculated LODs for each metabolite. Thereby, rank 1 represents the laboratory with the lowest LOD level and rank 14 the laboratory with the highest LOD level. A Spearman rank correlation test was performed between the median rank of a laboratory across all metabolites and the median percentage of missingness of each laboratory across all 12 x 3 measured project sample aliquots.

#### Comparison of measured values of QC2 with reference (target) values used by biocrates for normalisation

For each metabolite with less than 25% missing values (n=512), we calculated the mean absolute percentage error (MAPE) of its concentrations in QC2 aliquots measured by each laboratory compared to the “target” value which was determined by the manufacturer and is used within the kit software for normalisation of experimental sample concentrations to QC2 sample measurements. In other words, the MAPE represents a measure of how close a laboratory’s observed concentrations for a particular metabolite in the QC2 sample are compared to the concentrations in QC2 specified by biocrates.

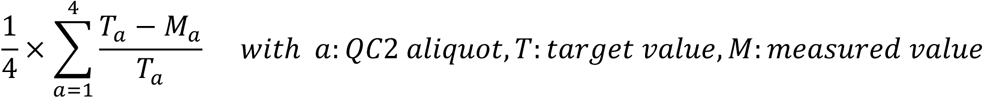

The median and mean of MAPEs across laboratories were calculated for each metabolite.

#### QC2 normalisation

Normalisation of metabolite concentrations measured for the project samples was performed separately for each dataset from each participating laboratory. For this purpose, we used the concentrations measured for QC2 samples in each laboratory and the QC2 target concentration values provided by biocrates (i.e., as used in the kit software). For each metabolite and laboratory data set, we calculated the median concentration of the four QC2 samples and determined the metabolite & laboratory-specific normalisation factor as the ratio of the metabolite target value divided by the QC2 median value. Metabolite concentrations measured for the project samples were then multiplied with their corresponding normalisation factors. If QC2 normalisation was not possible for a metabolite in a laboratory because no concentration was available for any of the four QC2 sample aliquots for this metabolite, the concentrations measured for the project samples in this laboratory were omitted (i.e., they were set “missing” in the QC2-normalised data files). For 19 metabolites, this was the case in all 14 participating laboratories. A list of all affected metabolites is provided in **Supplementary Table S2**.

#### Principal component analysis

We performed a principal component analysis (PCA) based on the 268 metabolites without any missing value using the *prcomp* function in R. To provide an overview of sources of variations, we overlaid the score plots for principal components 1 and 2 with information on the sample type and the laboratory, in which the sample was measured. To ensure that the variation of the subset of 268 metabolites is representative of the entire data set, we repeated the PCA including all 480 metabolites with less than 25 % missing values in the project samples (missing values were imputed with 0.0 for PCA calculation). Comparison of the score plots for principal components 1 and 2 showed the same pattern of variation in the score plots for both subsets of the data.

#### Interlaboratory comparison of metabolite concentrations

All analyses related to the concordance of metabolite concentrations between participating laboratories were based on values of the 561 metabolites for which at least one concentration value was available in more than three laboratories after QC2-normalisation. First, we calculated the coefficient of variation (CV = standard deviation/mean) across laboratories for each metabolite based on its mean concentration in the three aliquots of each project sample type within a laboratory. CV calculations were performed for the data sets before and after QC2-normalisation. Second, method comparison analyses were applied to evaluate whether there was a systematic difference in metabolite concentrations among the 14 laboratories. These analyses were applied separately for each metabolite according to the CLSI EP09-A3 guidelines (24) and involved three steps: *Step 1 - Pairwise laboratory comparisons using Deming regression:* the analytical measurement error ratios were assumed to be constant between laboratories and thus Deming regression was applied to implement the pairwise interlaboratory comparisons. The measurement errors were estimated from the triplicate measurements of each metabolite according to formulas stated in (25). The regression coefficients were estimated using the least-squares approach of Deming regression. The 95 % confidence intervals for the fitted regression coefficients were estimated using the Jackknife method. In addition, the presence of systematic error was determined by evaluating the confidence intervals of the regression coefficients. If the confidence interval for the intercept excluded 0, it was assumed that a constant error was present, and if the regression slope excluded 1, it was assumed that a proportional error was present. In the presence of at least one of the constant or proportional errors between two laboratories, a systematic error was considered to be present for the measured metabolite. The results for each metabolite were summarised in matrices showing the type of systematic error (N: No difference, C: Constant difference, P: Proportional difference, CP: Both constant and proportional difference) (**Supplementary Data S1**). *Step 2 - Calculation of the pairwise interlaboratory relative differences:* Absolute and relative differences (AD, RD) were calculated using the fitted regression coefficients to estimate the quantity of the systematic differences. These statistics were calculated separately for the minimum, median and maximum statistics of each metabolite using the formula

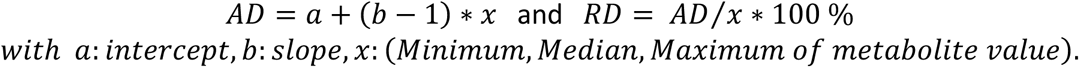

Again, the results for each metabolite were summarised in matrices showing the pairwise relative differences between laboratories (**Supplementary Data S2**). *Step 3 - Consolidation of differences:* After calculating the absolute and relative differences for each pair of laboratories for each metabolite, the obtained results were summarised. For this purpose, the median of pairwise relative differences was calculated for each laboratory and each metabolite. These median relative differences were log-transformed (base 2) and summarised using circular heatmap plots. The log-median relative differences of each laboratory were placed on each track of the graph, and the circle is split into the 11 metabolite classes as defined above. Circular heatmaps were generated using the R package *circlize* (26). Detailed results of the pairwise laboratory comparisons of measurements are additionally provided in an R/Shiny web-tool which can be accessed from https://biotools.erciyes.edu.tr/MethodCompRing/.

#### Comparison of measurements for NIST SRM 1950 to reference values determined by other studies

Reference values were collected from different sources: provided by NIST and supplied by different laboratories (18, 27, 28). To assess the concordance of concentrations measured for the NIST sample in this study with respective reference values from the literature, we calculated the MAPE for all metabolites, for which reference values were available in the above-mentioned sources. MAPE calculation was conducted analogous to the procedure used for comparing measured to target QC2 concentrations (see above).

Where not stated otherwise, statistical analyses and plot generation were performed using R (version 4.2.2).

## RESULTS

### Overview of the study

In this study, we assessed the performance and reproducibility of the MxP® Quant 500 kit in an interlaboratory comparative analysis. Fourteen laboratories (designated as A to N) located across different regions worldwide and utilizing various instrumentation setups participated in this study (**Figure 1**). Each laboratory received aliquots of twelve blinded samples, encompassing diverse sample types. These included human plasma samples obtained from three males and three females, along with serum samples from one male and one female from the same individuals. Additionally, a human lipaemic pooled plasma sample, a NIST SRM 1950 sample, as well as pooled mouse and rat plasma samples were included. To ensure consistency, all samples were analysed in triplicates by each participating laboratory, following a standardised plate design. The plate design incorporated additional samples for calibration, determination of the limit of detection (zero samples), and quality control samples for data normalisation (QC1, QC2, QC3) (**Figure 1**; for detailed procedures, see Methods).

Depending on the mass spectrometer used, the kit enabled the measurement of up to 628 metabolites on Waters instruments and 630 metabolites on Sciex instruments. In total, the kit covered 634 different metabolites, with 624 metabolites overlapping between the two systems (see Methods). These metabolites encompassed 26 compound classes, comprising a total of 107 small molecules (e.g., amino acids, fatty acids) and 527 complex lipids (e.g., triacylglycerols) (**Figure 2**, **Supplementary Table S1**). For the present study, we assigned metabolites to one of 10 metabolite classes to group structurally related metabolites (e.g., “Acylcarnitines”, “Amino Acids and Amines”, “Bile Acids”, “Ceramides”, “Diacylglycerols”, “Fatty Acids”, “Glycerophospholipids”, “Sphingomyelins”, and “Triacylglycerols”) or to the group “Others”, which comprises 21 measured metabolites from various functionally unrelated classes, namely alkaloids, carbohydrates and related, carboxylic acids, cresols, hormones and related, indoles and derivatives, nucleobases and related, vitamins and cofactors.

**Figure 2:**
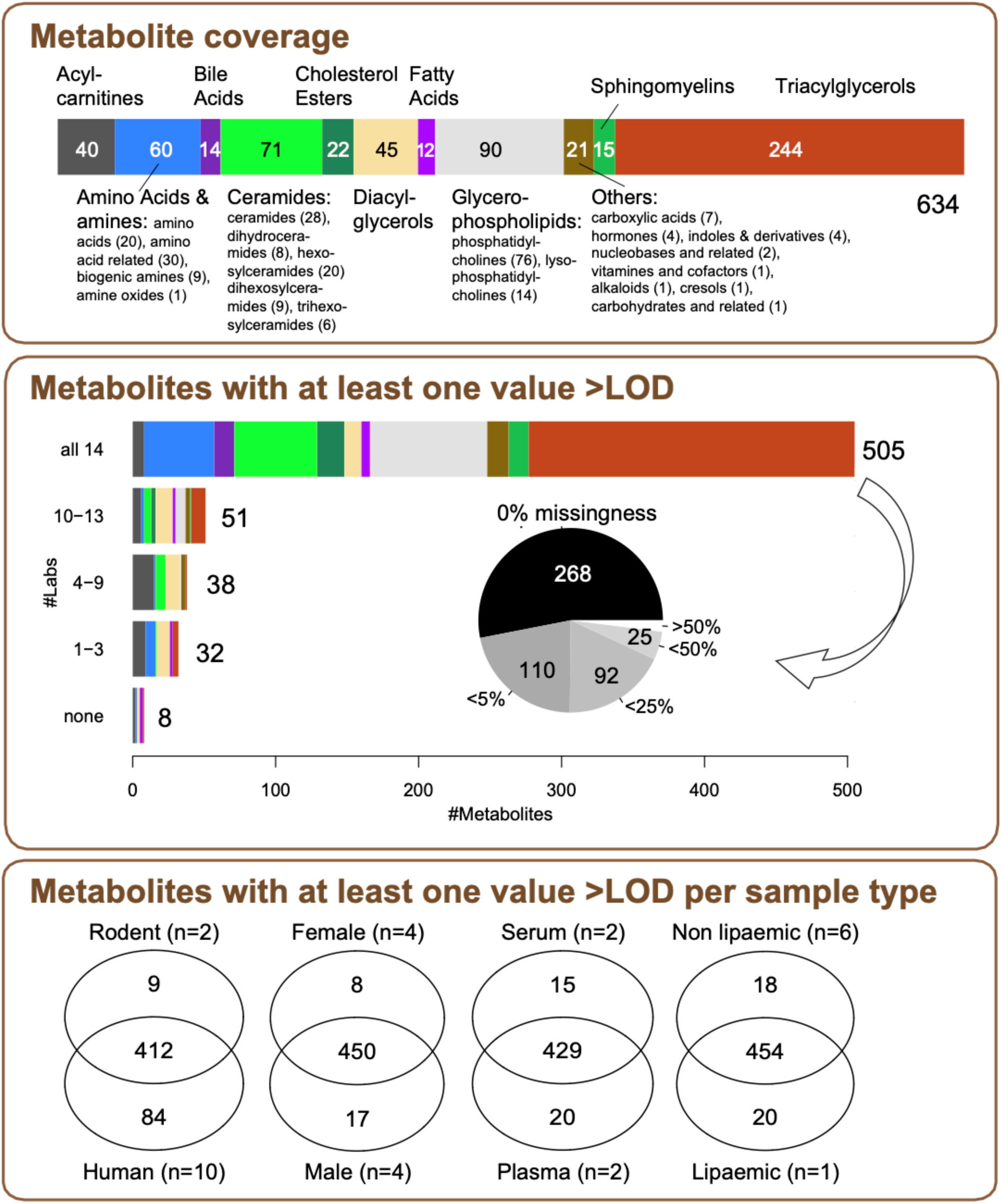
Metabolites measured in project samples across participating laboratories. The *top panel* provides an overview of the metabolite coverage of the MxP® Quant 500 kit. The *central panel* shows how many of these metabolites were detected in at least one measurement above LOD for the project samples within this study. The horizontal bars depict metabolite classes and numbers of metabolites that were detected in all 14 laboratories, in 10 to 13, in 4-9, or in 1-3 laboratories, respectively. Overall, 505 metabolites were measured in all 14 laboratories. The pie chart displays the proportion of these metabolites that were measured in all project samples (i.e., 0 % missingness), with less than 5 %, 25 %, and 50 %, or at least 50 % missingness. In the *bottom panel*, Venn diagrams depict comparisons of metabolite coverage between the different sample types for the 505 metabolites detectable in all laboratories.

### Metabolite coverage and overall performance

Out of the 634 targeted metabolites, 505 were detected by all 14 laboratories, i.e., a concentration above the calculated limit of detection (LOD) was reported for at least one project sample (**Figure 2, Supplementary Table S3**). On the other hand, eight metabolites were not detected in any of the twelve samples in any laboratory (**Table 1**). For 62.5 % of the 40 targeted acylcarnitines, at least 5 laboratories did not observe any value above the LOD in any project sample (**Table 1**).

**Table 1:**
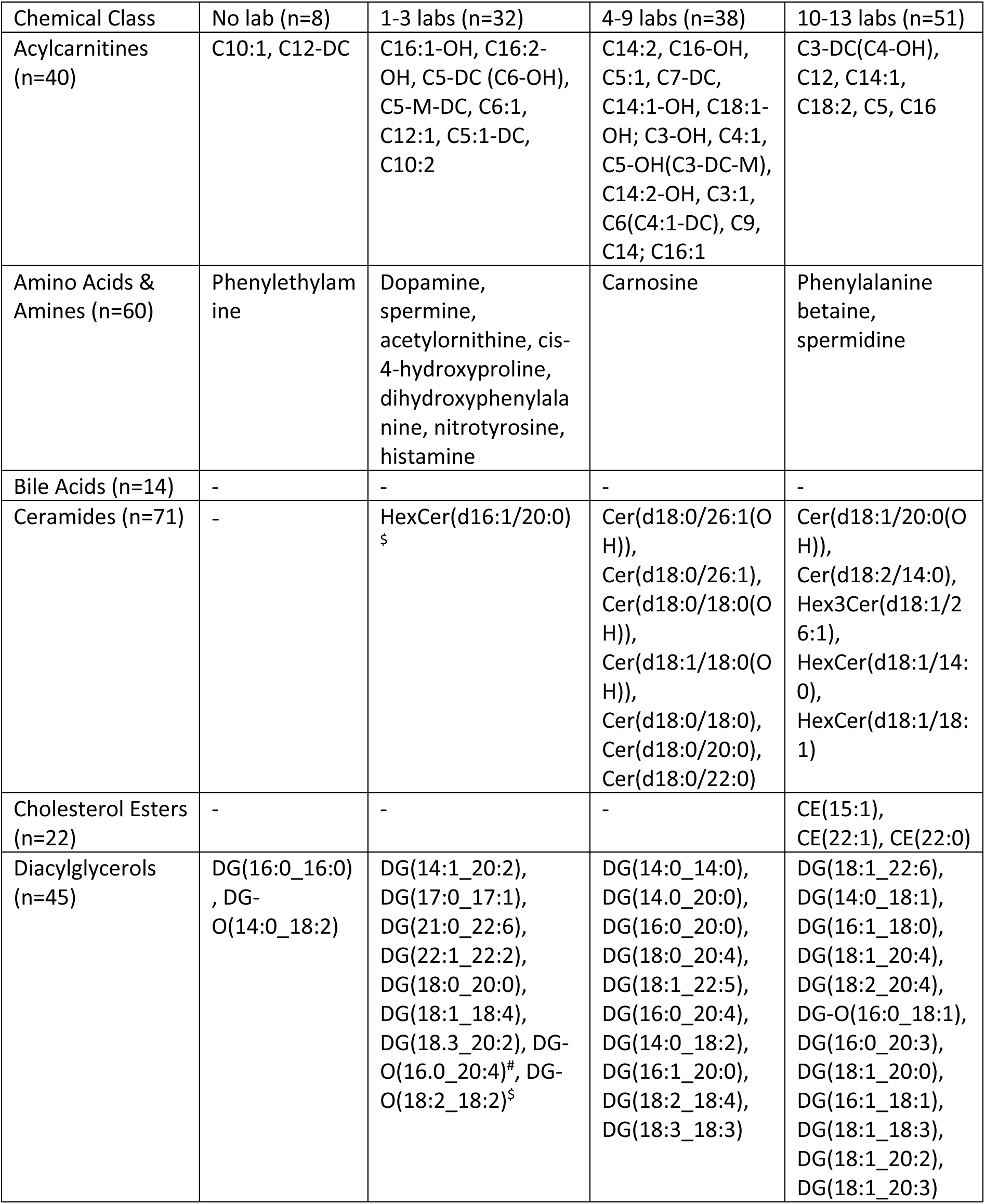
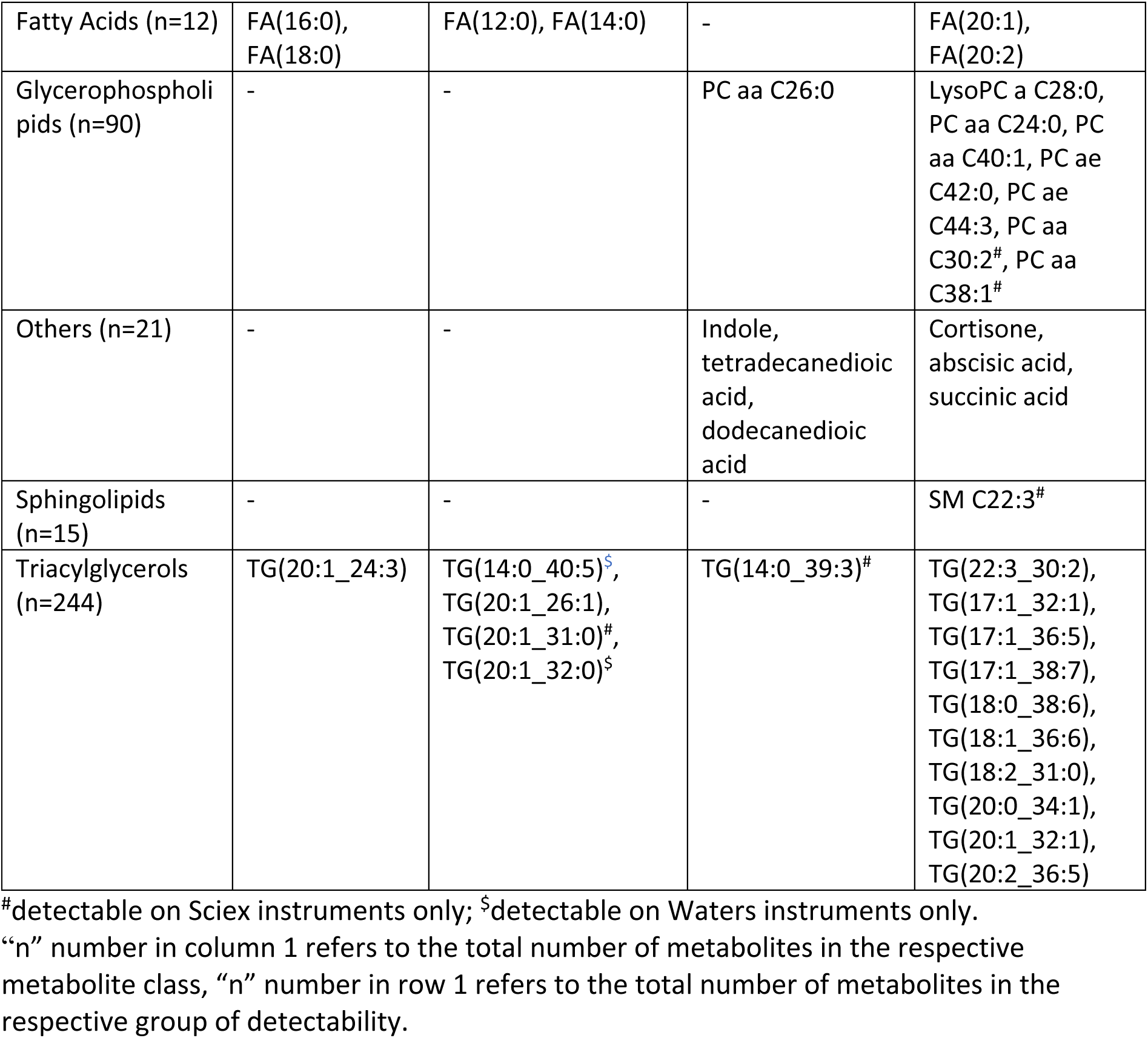
Metabolites which were only partially covered across laboratories.

| Chemical Class | No lab (n=8) | 1-3 labs (n=32) | 4-9 labs (n=38) | 10-13 labs (n=51) |
| --- | --- | --- | --- | --- |
| Acylcarnitines (n=40) | C10:1, C12-DC | C16:1-OH, C16:2-OH, C5-DC (C6-OH), C5-M-DC, C6:1, C12:1, C5:1-DC, C10:2 | C14:2, C16-OH, C5:1, C7-DC, C14:1-OH, C18:1-OH; C3-OH, C4:1, C5-OH(C3-DC-M), C14:2-OH, C3:1, C6(C4:1-DC), C9, C14; C16:1 | C3-DC(C4-OH), C12, C14:1, C18:2, C5, C16 |
| Amino Acids & Amines (n=60) | Phenylethylamine | Dopamine, spermine, acetylorntithine, cis-4-hydroxyproline, dihydroxyphenylalanine, nitrotyrosine, histamine | Carnosine | Phenylalanine betaine, spermidine |
| Bile Acids (n=14) | - | - | - | - |
| Ceramides (n=71) | - | HexCer(d16:1/20:0)<sup>s</sup> | Cer(d18:0/26:1(OH)), Cer(d18:0/26:1), Cer(d18:0/18:0(OH)), Cer(d18:1/18:0(OH)), Cer(d18:0/18:0), Cer(d18:0/20:0), Cer(d18:0/22:0) | Cer(d18:1/20:0(OH)), Cer(d18:2/14:0), Hex3Cer(d18:1/26:1), HexCer(d18:1/14:0), HexCer(d18:1/18:1) |
| Cholesterol Esters (n=22) | - | - | - | CE(15:1), CE(22:1), CE(22:0) |
| Diacylglycerols (n=45) | DG(16:0_16:0), DG-O(14:0_18:2) | DG(14:1_20:2), DG(17:0_17:1), DG(21:0_22:6), DG(22:1_22:2), DG(18:0_20:0), DG(18:1_18:4), DG(18:3_20:2), DG-O(16:0_20:4)<sup>#</sup>, DG-O(18:2_18:2)<sup>s</sup> | DG(14:0_14:0), DG(14:0_20:0), DG(16:0_20:0), DG(18:0_20:4), DG(18:1_22:5), DG(16:0_20:4), DG(14:0_18:2), DG(16:1_20:0), DG(18:2_18:4), DG(18:3_18:3) | DG(18:1_22:6), DG(14:0_18:1), DG(16:1_18:0), DG(18:1_20:4), DG(18:2_20:4), DG-O(16:0_18:1), DG(16:0_20:3), DG(18:1_20:0), DG(16:1_18:1), DG(18:1_18:3), DG(18:1_20:2), DG(18:1_20:3) |
| Fatty Acids (n=12) | FA(16:0),<br>FA(18:0) | FA(12:0), FA(14:0) | - | FA(20:1),<br>FA(20:2) |
| Glycerophospholipids (n=90) | - | - | PC aa C26:0 | LysoPC a C28:0, PC aa C24:0, PC aa C40:1, PC ae C42:0, PC ae C44:3, PC aa C30:2<sup>#</sup>, PC aa C38:1<sup>#</sup> |
| Others (n=21) | - | - | Indole, tetradecanedioic acid, dodecanedioic acid | Cortisone, abscisic acid, succinic acid |
| Sphingolipids (n=15) | - | - | - | SM C22:3<sup>#</sup> |
| Triacylglycerols (n=244) | TG(20:1_24:3) | TG(14:0_40:5)<sup>§</sup>, TG(20:1_26:1), TG(20:1_31:0)<sup>#</sup>, TG(20:1_32:0)<sup>§</sup> | TG(14:0_39:3)<sup>#</sup> | TG(22:3_30:2), TG(17:1_32:1), TG(17:1_36:5), TG(17:1_38:7), TG(18:0_38:6), TG(18:1_36:6), TG(18:2_31:0), TG(20:0_34:1), TG(20:1_32:1), TG(20:2_36:5) |

<sup>#</sup>detectable on Sciex instruments only; <sup>§</sup>detectable on Waters instruments only.
“n” number in column 1 refers to the total number of metabolites in the respective metabolite class, “n” number in row 1 refers to the total number of metabolites in the respective group of detectability.

Out of the 505 commonly detected metabolites, measurements for 268 metabolites were complete, i.e., their concentrations were above LOD in all aliquots of all project samples in all laboratories; 202 metabolites showed a missingness below 25 % (i.e., more than 378 out of the 504 (= 14*12*3) measurements were above LOD for these metabolites).

The majority (n = 412) of the 505 metabolites that were reliably measured across all participating laboratories were detected in human as well as rodent blood. Nine metabolites (5-aminovaleric acid, anserine, phenylacetylglycine, putrescine, hydroxyglutaric acid, Cer(d18:1/26:0), DG(18:2_18:3), DG(18:2_20:0), FA(20:3)) were only detected in rodent blood despite the smaller number of rodent samples (n = 2) compared to human samples (n = 10). We also observed differences in metabolite coverage between serum versus plasma, male versus female, and lipaemic versus non-lipaemic samples. Details are provided in **Supplementary Table S4**.

Regarding the proportion of missing values for a metabolite, we observed some differences among the 14 participating laboratories (**Figure 3**, **Supplementary Table S3**). We hypothesised that the number of missing values per sample associated with the LODs, which were calculated based on the zero samples in each laboratory. To test this hypothesis, we ranked laboratories by their calculated LODs for each metabolite and related the median rank of metabolites for each laboratory to the laboratories’ median proportion of missing values across all samples (36 measurements). As a result, we found that higher LOD values significantly correlated with more missingness in the reported data (Spearman rho = 0.695, *p* = 0.0057).

**Figure 3.**
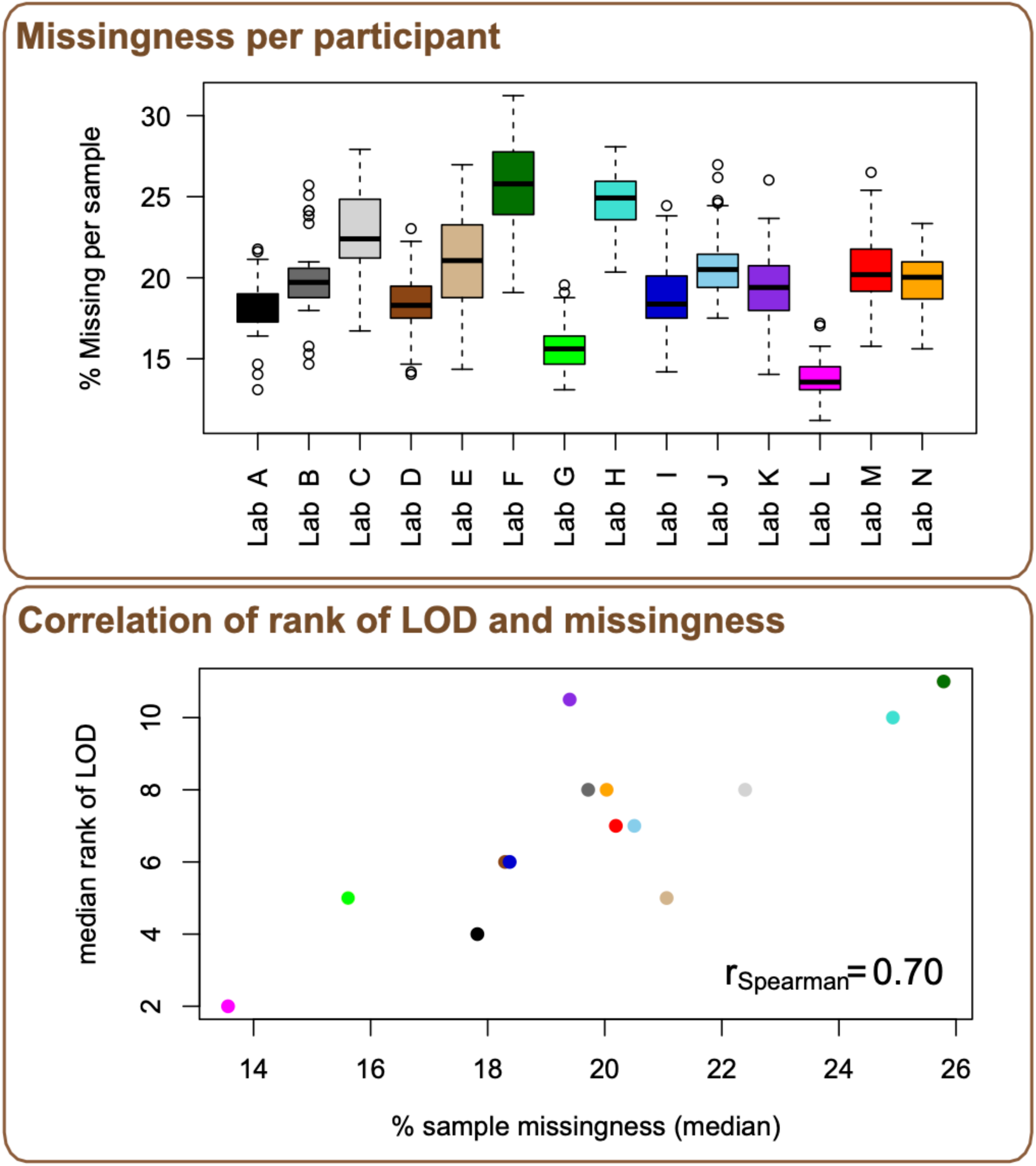
Laboratory-specific missingness. *Top panel:* Laboratories showed differences in the number of missing measurements. *Bottom panel:* Thereby, higher laboratory-specific LODs correlated with higher overall missingness. Color coding of laboratories as in Figure 1.

To provide an overview of the variation observed across all 504 measured project samples (14 laboratories x 12 project sample types in triplicates), we performed a principal component analysis (PCA) of the (unnormalised) measured concentrations and mapped information about sample type (denoted by symbol) and laboratory (denoted by color) onto the PCA scoring plot for the principal components (PC) 1 and 2 (**Figure 4** (center left)). This analysis showed that the largest variation in the data was linked to the lipid content of the samples, separating lipaemic samples from the rest on PC1. Also, differences between the laboratories contribute to the variation depicted in PC1 dimension. The variation attributed to the different species is captured in the PC2 dimension.

**Figure 4:**
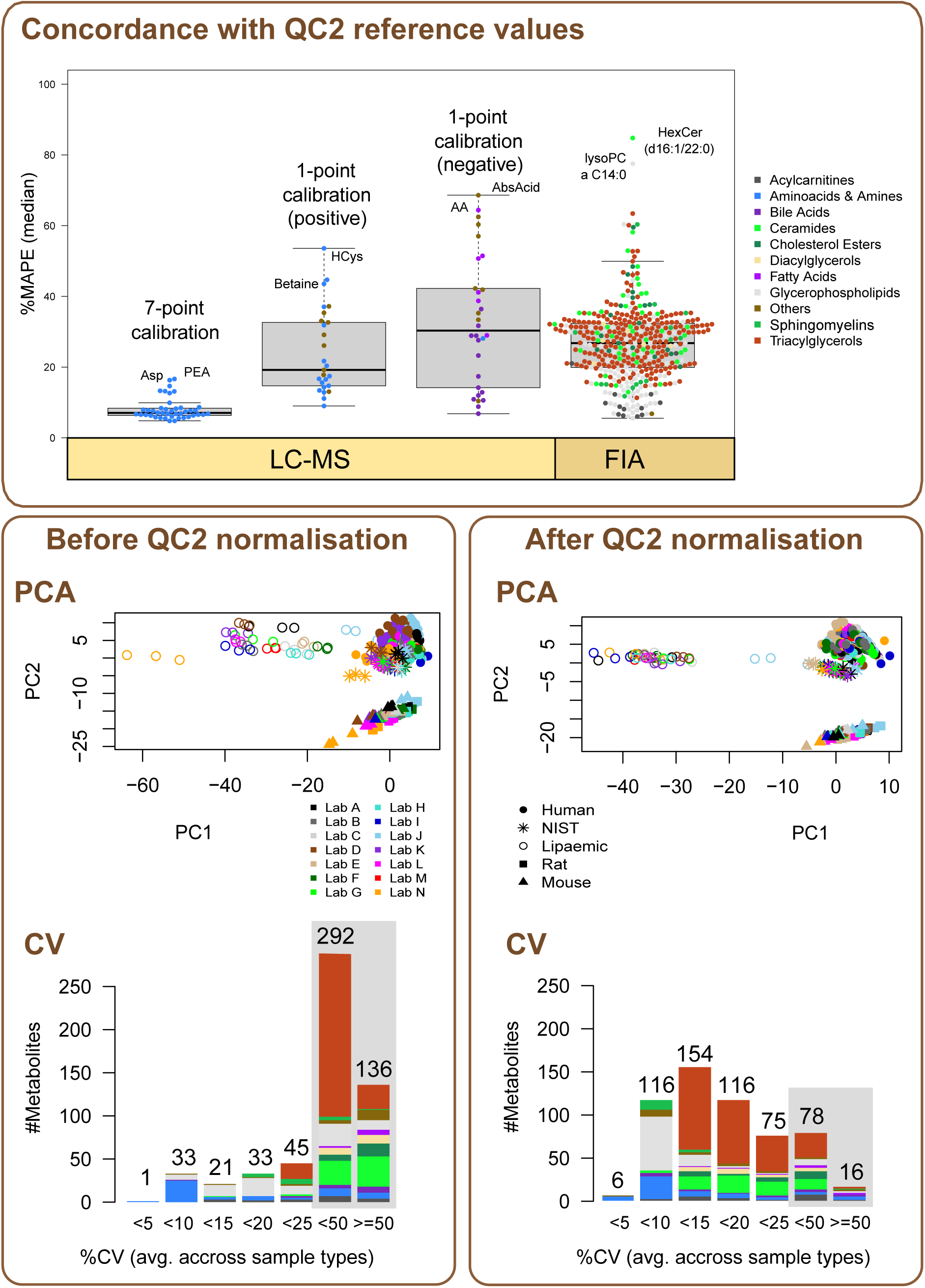
Normalisation of metabolites using QC2. *Top panel:* Mean absolute percentage error (MAPE) distribution of measured QC2 concentrations of all metabolites detected in QC2 samples compared to their reference target values provided by the kit manufacturer is shown separated by the measurement method used: (from left to right) after LC separation: (i) metabolites quantified with multi-point calibration curves (7-point calibration), (ii) metabolites quantified using 1-point calibration detected in positive or (iii) negative mode, and (iv) metabolites detected by flow injection analysis (FIA). (Metabolite abbreviations: Asp asparagine, PEA - phenylethylamine, HCys – homocysteine, AA – arachidonic acid, AbsAcid – abscisic acid.) *Central panel:* PCA score plot for PC1 and PC2 before (left) and after (right) QC2 normalisation (center). *Bottom panel:* Average interlaboratory CVs of all sample types for 561 metabolites before and after QC2 normalisation.

### Effects of normalisation

To obtain a reliable quantification and minimise batch effects, the kit manufacturer recommends normalizing the measured metabolite plasma concentrations to the concentrations in the quality control sample QC2, which largely resembles the physiological concentration ranges of metabolites in human plasma (except a few spiked metabolites). For this purpose, the kit software provides reference values for the concentration of each metabolite in QC2. To assess how close a laboratory’s observed value was to the target value specified by the manufacturer before normalisation, we calculated the mean absolute percentage error (MAPE) of QC2 concentrations measured by each laboratory compared to the reference values (**Supplementary Table S5)**. Nineteen metabolites (7 acylcarnitines, 5 diacylglycerols, 2 fatty acids, 2 glycerophospholipids, 3 triacylglycerols) did not have any value above LOD in QC2 in any laboratory in this study, i.e., QC2 could not be used as a reference for their normalisation. Those metabolites were omitted in further analyses. For 10 metabolites (FA(18:2), CE(14:1), DG(18:1_20:1), Hex3Cer(d18:1/20:0), HexCer(d16:1/24:0), HexCer(d18:2/16:0), TG(17:1_38:5), TG(20:2_36:5), TG(20:4_35:3), TG(22:2_32:4)), the median MAPE across the 14 laboratories was above 100 %, indicating very large differences between measured concentrations and reference values. The distribution of median MAPEs for the remaining metabolites is depicted in **Figure 4** (top) with the different methods used for their measurement and quantification. Considering metabolites determined by LC-MS, the 7-point calibration resulted in the lowest MAPE values followed by a 1-point calibration in positive mode. The weakest concordance with QC2 reference values was observed for 1-point calibration in negative mode. For metabolites measured by FIA, no 7-point calibration is available; nevertheless, acylcarnitines and glycerophospholipids showed MAPE values comparable to those achieved with that quantification mode. In contrast, the concordance of measured and reference values varied for the other metabolites analysed by FIA. In particular, ceramides, and triacylglycerols showed a lower agreement with QC2 reference values.

After QC2-normalisation, 561 metabolites with at least one (normalised) concentration value in more than three laboratories were available for further analyses. Overall, the normalisation reduced the variation in the data that is attributed to differences between the 14 laboratories as can be seen in the PCA score plot based on the QC2-normalised data (**Figure 4**, center right). This improvement in the reproducibility of measurement results is also reflected when comparing the coefficients of variation (CVs) for the 561 metabolites before and after normalisation (**Figure 4**, bottom; **Supplementary Table S6**). While, before QC2-normalisation, only 133 metabolites showed an average interlaboratory CV of all sample types below 25 %, 467 metabolites met this threshold after normalisation. Thereby, the effects of QC2-based normalisation were most prominent for those metabolites, for which the MAPE and its variance across laboratories were high (**Figure 4**, **Supplementary Table S6**). In particular, the reproducibility of measured concentrations increased for triacylglycerols and ceramides. Out of 239 triacylglycerols, only 18 (7.5 %) and out of 66 ceramides, only 3 (4.5 %) had a CV below 25 % before normalisation. After normalisation, 209 (87.4 %) triacylglycerols and 54 ceramides (81.8 %) showed a CV below 25 % (**Supplementary Table S7**). In addition, the reproducibility of metabolites with high MAPE that were detected using LC separation largely improved through the normalisation. As an example, MAPE for homocysteine was 53.6 %, indicating large interlaboratory differences in the measured concentrations before normalisation. Accordingly, the interlaboratory CV for homocysteine was 103.5 % before normalisation, dropping to 16.9 % after QC2 normalisation.

### Interlaboratory reproducibility

For analyses of interlaboratory reproducibility in our study, we used the QC2-normalised data set, comprising 561 metabolites, for which (normalised) data was available from more than three laboratories. The median interlaboratory CV of all 561 metabolites and all sample types was 14.3 %. Thereby, 122 (22 %) metabolites had a mean CV below 10 % and 467 (83 %) metabolites below 25 % (**Figure 4**, **Supplementary Table S7**). The lowest CVs were observed for the amino acids (7.7 % on average), most of which were quantified using 7-point calibration. In contrast, fatty acids measures were less reproducible, showing relatively high CVs (34.7 % on average). For other classes such as acylcarnitines, glycerophospholipids, or ceramides, CVs varied between the different analytes. For example, while C2 had a mean CV of 6.1 %, C6(C4:1-DC) was only measured with a mean CV of 66.5 % (**Supplementary Table S6**).

For a more detailed analysis of the systematic (constant and proportional) differences of measurements across the 14 laboratories, we performed pairwise laboratory comparisons of measurements using Deming regression analysis on each metabolite (see Methods) (**Supplementary Data S1**). Based on the fitted regression coefficients for the minimum, median, and maximum statistics of each metabolite, we calculated the absolute and relative differences for each pair of laboratories, representing estimates for the systematic differences in measurements (**Supplementary Data S2**). Detailed matrices of pairwise laboratory comparisons of measurements are provided at https://biotools.erciyes.edu.tr/MethodCompRing/. To summarise these results, we calculated the log-median relative differences (LMRD) of each laboratory and generated circular heatmaps (**Figure 5**, center) with saturated green color indicating minor differences (LMRD ≤ 2), and saturated red color indicating large differences (LMRD > 8); white color indicates missing information due to lack of measurements for the corresponding metabolites.

**Figure 5:**
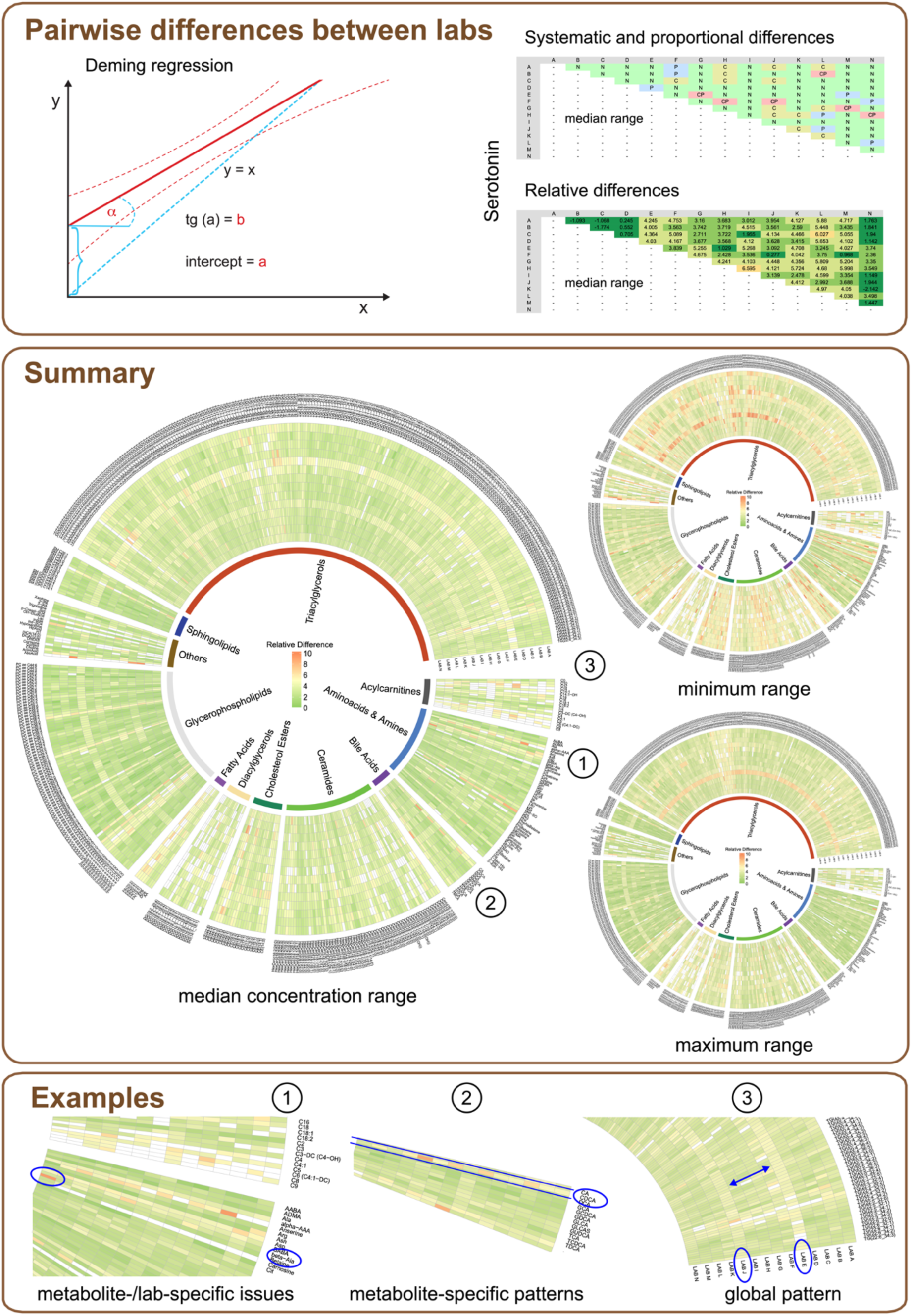
Pairwise laboratory comparisons of measurements using Deming regression. *Top panel:* Deming regression was used to assess pairwise systematic constant and proportional differences of measurements. Results for each metabolite are provided as matrices indicating whether concentrations between two laboratories were not significantly different (“N”), or showed significant constant (“C”), proportional (“P”) or constant and proportional differences (“CP”). In addition, we also provide matrices containing and color-coding the observed relative differences for each metabolite in the minimum, median, and maximum ranges. *Center panel:* Circular heatmaps visualising the log-median relative differences of each laboratory for each metabolite for the median, minimum, and maximum concentration ranges using a color gradient from saturated green (LMRD ≤ 2) to saturated red (LMRD > 8); white color indicates missing information due to lack of measurements for the metabolite in this laboratory. *Bottom panel:* Inlets of the circular heatmaps show selected examples of observed patterns of reproducibility.

These heatmaps allow a detailed inspection of results for each metabolite and pinpoint various patterns of differences (**Figure 5**, center and bottom): (i) Overall, laboratories produced concordant results for most of the metabolites with the smallest differences being observed for median concentration ranges (LMRD ≤ 4 for 76 % of metabolites). (ii) Measurements from laboratories E and J showed larger differences compared to the remaining laboratories, in particular for triacylglycerols in the minimum concentration range. (iii) In the classes “Acylcarnitines” and “Diacylglycerols”, many metabolites were not consistently detected across laboratories as indicated by the high number of white cells in the circular plots. (iv) Bile acid measurements showed overall good concordance across laboratories in the maximum and medium ranges except for CDCA (all laboratories) and TMCA (Laboratory H); larger differences were observed in the minimum range. (v) For asparagine, concentrations measured by Laboratory A differed considerably in higher concentration ranges while measurements of all remaining laboratories showed good concordance; similarly, beta-alanine concentrations measured by Laboratory N differed significantly from measurements of Laboratories A-M at all concentration ranges. (For the latter case, we could trace back the issue to an error in peak integration.)

### Comparison of reproducibility across different sample types

Comparing the number of metabolites with CVs below 25 %, the interlaboratory reproducibility was largely similar across sample types with numbers of metabolites ranging between 400 and 494 for rat pool plasma and NIST SRM 1950, respectively (**Figure 6, Supplementary Table S6**). While the NIST SRM 1950 samples showed the best reproducibility in terms of metabolites with a CV below 25 %, it showed the lowest number (n=138) of metabolites with a CV below 10 % among the human blood samples. In general, we observed four representative patterns of CV distributions related to sample types (**Figure 6**, bottom left): (i) metabolites with similarly low CVs for all sample types (example: acetylcarnitine (C2)), (ii) metabolites with similarly low CVs for all human sample types but higher CVs for rodent samples (example: HexCer(d16:1/22:0)), (iii) metabolites with similarly low CVs for the two rodent sample types but higher CVs for all human samples (example: PC ae C38:1), and (iv) metabolites with large variation across sample types (example: TG(18:0_30:1): higher CVs for rodent plasma and human serum samples). In many cases, variations in the CVs across sample types (and metabolites) can be explained by differences in the mean metabolite concentrations, with higher CVs generally being significantly associated with lower mean concentrations (Spearman rho ranging from r_lipaemic_ = −0.22 to r_plasma_female_1_ = −0.60) (**Supplementary Table S8**).

**Figure 6:**
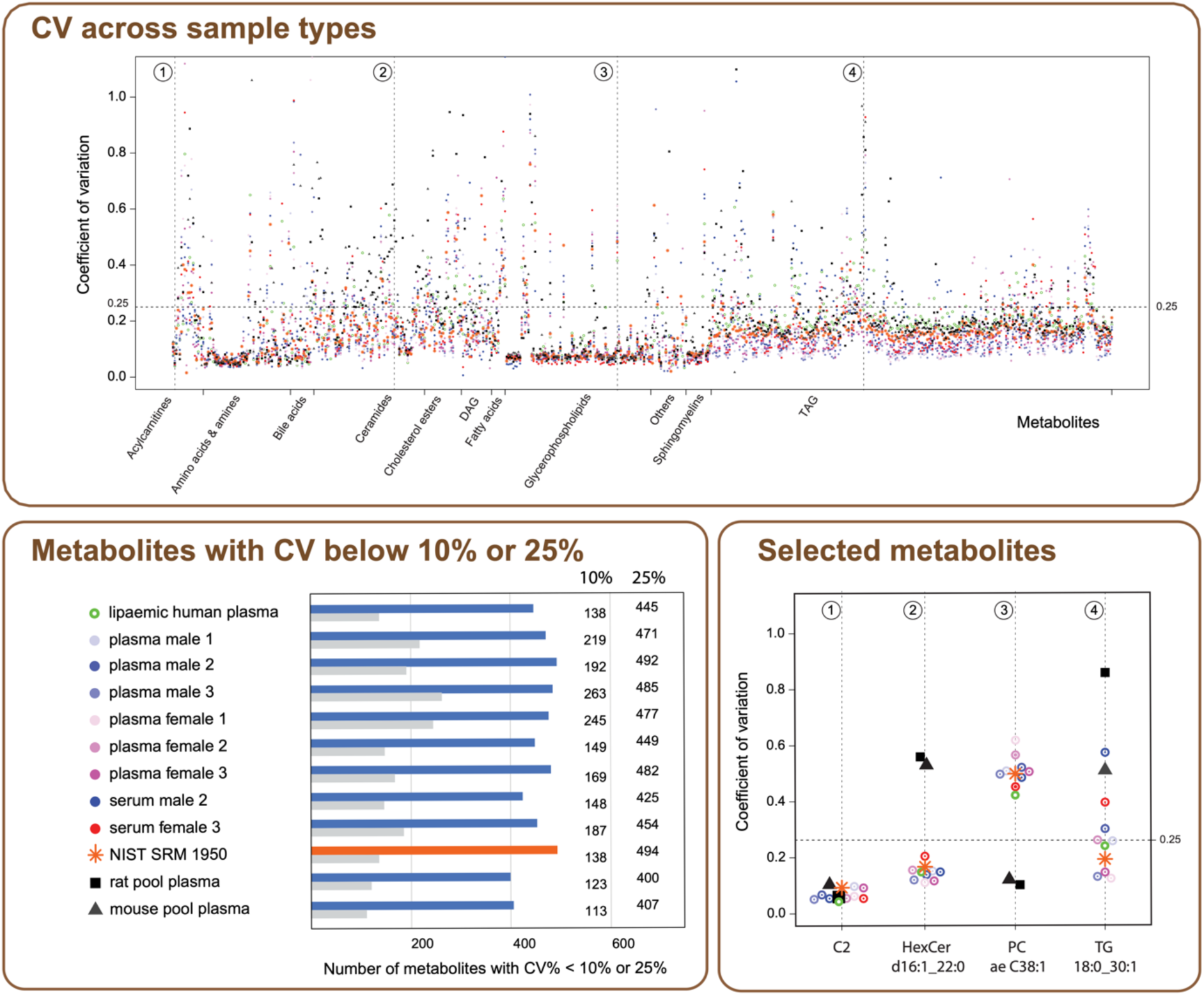
CV values in different sample types. The *top panel* shows all CVs as calculated for each metabolite and sample type. Vertical dotted lines indicate the overlay of CV values for specific metabolites that have been selected as examples in the lower right panel. Sample NIST SRM 1950 is highlighted in orange. The *bottom left panel* illustrates the number of metabolites with CVs below 25 % (color) or 10 % (grey) within each sample type. The *bottom right panel* shows the distribution of CVs across sample types for acetylcarnitine (C2), HexCer(d16:1/22:0), PC ae C38:1, and TG(18:0_30:1).

### Comparison of absolute quantification with other kits

The NIST SRM 1950 sample has been used in various ring trials and studies of interlaboratory reproducibility in the past (17, 18, 27, 29). In Table 2, we compare CVs for metabolites covered by the MxP® Quant 500 kit in the NIST SRM 1950 sample to those reported for the NIST sample using other assays, namely the AbsoluteIDQ p180 (29), the AbsoluteIDQ p400HR (17), and the Lipidyzer (18). In study 1, the AbsoluteIDQ p180 kit (using the same LC-MS/MS and FIA-MS/MS setup for triple quadrupole mass spectrometers as in our study for a subset of 188 metabolites) was applied, and the CVs were calculated for triplicate measurements performed in 6 laboratories. In study 2, the AbsoluteIDQ p400HR kit was used to perform LC-MS/MS and FIA-MS/MS measurements with the Q Exactive Orbitrap mass spectrometer (Thermo Fisher Scientific), quantifying up to 408 metabolites in triplicate measurements in 14 laboratories. In study 3, the Lipidyzer kit was used to quantify lipids with SelexION (differential ion mobility or DSM) metabolite fractionation before analyses on triple quadrupole mass spectrometers in 5 replicate measurements in 9 laboratories.

**Table 2.** Mean interlaboratory CVs of metabolite classes for measurements of NIST SRM 1950 by different kits.

| Metabolite classes* | Metabolite subclasses # | This study |  | Other studies |  |  |
| --- | --- | --- | --- | --- | --- | --- |
|  |  | Mean CV (%) in class * | Mean CV (%) in subclass # | CV in study 1 § | CV in study 2 & | CV in study 3 Δ |
| Amino Acids & Amines | Amino acids | 15.7 | 7.4 | 10.2 | **7.1** | - |
|  | Amino acids related |  | 26.1 | 24.0 | - | - |
|  | Biogenic amines |  | 24.6 |  | 27.0 | - |
|  | Amine oxide |  | 4.6 | - | - | - |
| Bile acids |  | 13.5 | 13.5 | - | - | - |
| Others | Sugars | 17.6 | **5.2** | 9.4 | 9.3 | - |
|  | Carboxylic acids |  | 24.6 | - | - | - |
|  | Cresols |  | 8.8 | - | - | - |
|  | Vitamines and cofactors |  | 21.2 | - | - | - |
|  | Alkaloids |  | 4.9 | - | - | - |
|  | Hormones and related |  | 16.5 | - | - | - |
|  | Indoles and derivatives |  | 41.1 | - | - | - |
|  | Nucleobases and related |  | 23.8 | - | - | - |
| Cholesterol esters |  | 23.4 | 23.4 | 15.5 | - | **3.6** |
| Fatty acids |  | 27.6 | 27.6 | - | - | **9.9** |
| Acylcarnitines |  | 20.7 | **20.7** | 38.2 | 34.0 | - |
| Ceramides | Ceramides | 16.9 | 17.4 | - | - | **5.7** |
|  | Dihydroceramides |  | **14.0** | - | - | 25.7 |
|  | Dihexosylceramides |  | 19.6 | - | - | - |
|  | Trihexosylceramides |  | 18.4 | - | - | - |
|  | Hexosylceramides |  | 15.3 | - | - | **4.6** |
| Glycerophospholipids | Lysophosphatidylcholines | 12.9 | 14.4 | 22.6 | 18.0 | **8.2** |
|  | Phosphatidylcholines |  | **11.5** |  | 16.0 | 12.3 |
| Sphingomyelins |  | 11.7 | **11.7** | 15.1 | 68.0 | 20.4 |
| Diacylglycerols |  | 21.4 | 21.4 | 24.8 | - | **7.8** |
| Triacylglycerols |  | 17.5 | 17.5 |  | - | **1.7** |

\* Metabolites assignment into groups as used in this study

Although these four targeted panels only share partial coverage, large overlaps in specific chemical classes could be compared. For most metabolite classes, the previously reported interlaboratory CVs were similar to the CVs observed in our study. This holds especially true for other biocrates kits (AbsoluteIDQ p180 and AbsoluteIDQ p400HR). The lipid-focused Lipidyzer approach revealed similar CVs for phosphatidylcholines, better CVs for cholesterol esters, fatty acids, ceramides, hexosylceramides, lysophosphatidylcholines, diglycerols and triglycerols, and worse CVs for dihydroceramides and sphingomyelins than those measured in our study. The median CV of all covered metabolites in our study and studies 1-3 were 17.2 %, 19.0 %, 18.0 %, and 10.0 %, respectively.

To further evaluate the reproducibility of metabolite concentrations measured for the NIST SRM 1950 across different studies and measurement platforms, we directly compared the absolute concentrations measured in this study with concentrations that were provided by the manufacturer (NIST) as certified or reference (concentrations determined applying various analytical methods) as well as with concentrations measured and provided in two previous studies (18, 27). As a measure of concordance, we calculated the MAPE, relating the concentrations measured by each of the laboratories participating in our study to the externally determined concentrations (**Figure 7**, top; **Supplementary Table S9**). Fatty acids (FAs) were among the metabolites where the MxP® Quant 500 kit-derived concentrations showed the largest differences to reference values. In contrast, measurements for most complex lipids showed a low MAPE except for several ceramides and diacylglycerols. Also, measured and certified concentrations for amino acids in the NIST (28) SRM 1950 sample showed low MAPE values except for glycine and alanine, for which most laboratories participating in our study found higher concentrations in the NIST sample (**Figure 7**, bottom). Specific metabolites with the highest MAPE (discordant measurements) and lowest MAPE (concordant measurement) are indicated in **Figure 7**. Among the metabolites in NIST SRM 1950 for which we found good concordance of measured and certified/reference concentrations included cortisol (0.231 μmol/l ± 0.005 (NIST) vs 0.181 μmol/l ± 0.019 (this study)), homocysteine (8.500 μmol/l ± 0.200 vs 8.695 μmol/l ± 1.877), and glucose (4560 μmol/l ± 56 vs sum of hexoses 4747 μmol/l ± 361, with glucose being the major hexose in human plasma (30)).

**Figure 7:**
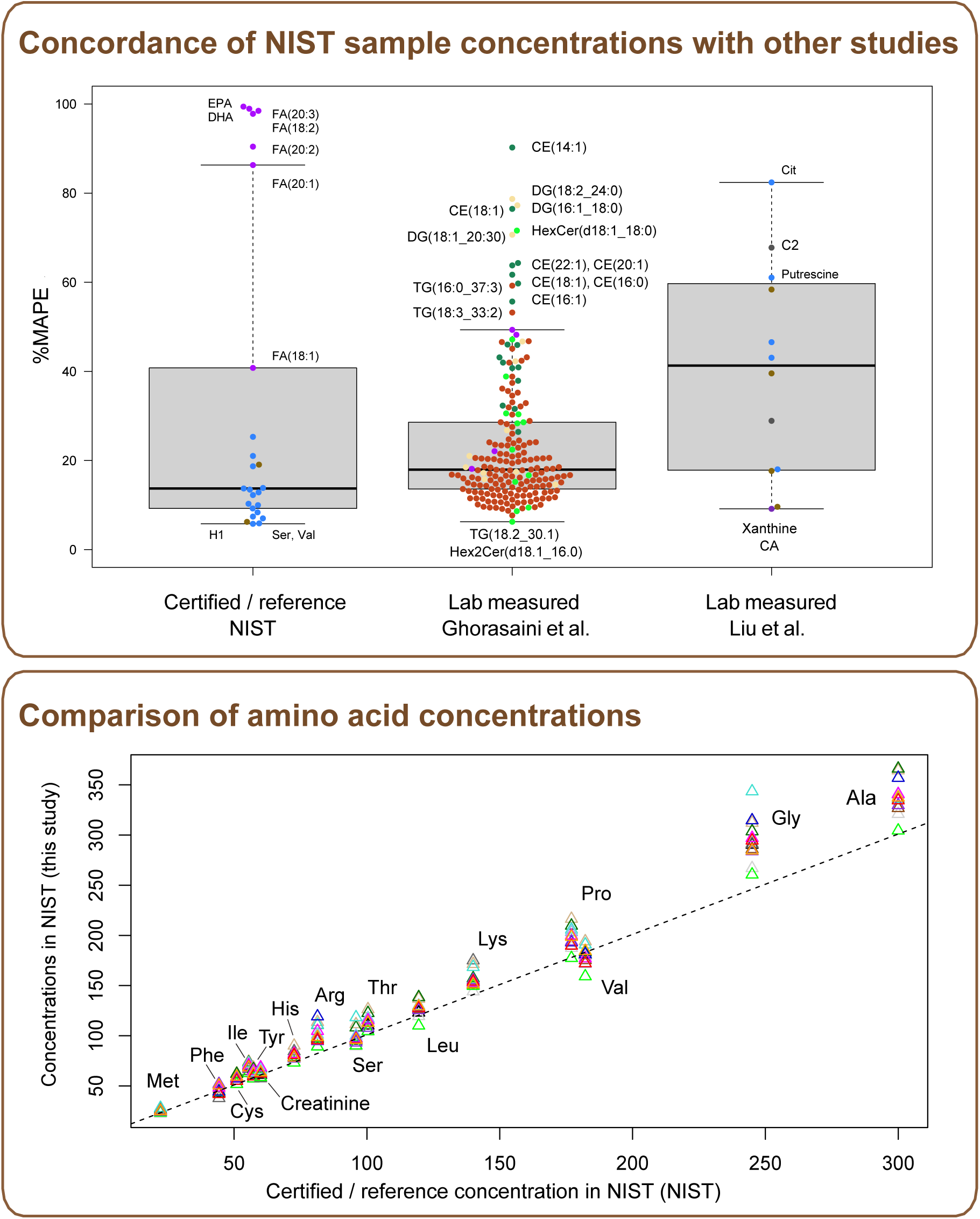
Comparison of metabolite concentrations in the NIST SRM 1950 sample between studies. *Top panel:* Mean absolute percentage error (% MAPE) of data from this project versus those (i) certified by NIST (or provided as reference) (28), (ii) measured by Ghorasaini et al. (18), and (iii) Liu et al. (27). Color coding for metabolites represents metabolite classes as in Figure 2. *Bottom panel:* A correlation plot of amino acid values measured in this study versus those reported for the SRM 1950 probe as certified or reference by NIST. For each amino acid, individual concentrations measured in the Laboratories A-N are shown. Color coding of the laboratories as in Figure 1. Abbreviations: EPA - eicosapentaenoic acid (FA(20:5)), DHA - docosahexaenoid acid (FA(22:6)), Cit - citrulline, C2 – acetylcarnitine, H1 – hexoses, CA – cholic acid.

## DISCUSSION

Contemporary omics studies of human phenotypes face the unmet need to explain tolerable variability of molecular mechanisms in health and intolerable deviations in disease. Beyond the importance of depicting adequate reference values for a broad range of molecular pathways, there is a growing need to demonstrate the diversity of referenced values in different ethnicities (31) or under distinct environmental conditions (32, 33). In genomics research, a so-called “pangenomics” approach addresses these aspects (34). To accomplish panmetabolomic studies, metabolomic data sets from large population-wide biobanks need to be generated and combined. To this end, it is crucial to explore the compatibility, and limits of available metabolomics methodologies, in particular concerning the reproducibility and concordance of measurements within and across laboratories.

Moreover, the increasing availability of large population-wide biobanks that also support longitudinal analyses of clinical and molecular omics phenotypes calls for a scaling up of sample throughput. One approach to increase the throughput of metabolomics is the distribution of measurements across multiple laboratories using standardised methods that provide a broad metabolite coverage while ensuring precise quantification. In this study, we investigated the reproducibility of quantifications for more than 630 metabolites using the MxP® Quant 500 kit across 14 laboratories.

### More than 70 % of kit metabolites were reliably detected and reproducibly quantified across laboratories

Despite the use of different instrumental setups, the coverage of the 624 metabolites that are targeted on both, Sciex and Waters instruments remained essentially consistent. The majority of metabolites (n = 505) were detected in all laboratories, with only 8 of the 624 theoretically possible metabolites remaining undetected across all laboratories. Likely, concentrations of some of these undetected metabolites (e.g., phenylethylamine, C10:1, C12-DC) were very low in the studied samples, which mostly represent blood levels in healthy, non-fasting conditions (35–37). Analogously, variations in the detectability of metabolites across laboratories were largely explained through laboratory-specific differences in the estimated limits of detection (LOD) in our study. However, low concentrations do not explain the undetectability in all cases as some lipids with relatively high abundance in (human) blood were also affected (e.g., FA(16:0), FA(18:0)) (38).

For all 561 metabolites with considerable coverage across laboratories, we observed high reproducibility of quantifications with a median coefficient of variation (CV) of 14.3 %. Thereby, 467 metabolites had CVs below 25 %. As expected, most amino acids which were analyzed using LC and 7-point calibration showed the lowest CVs. But notably, various glycerophospholipids and acylcarnitines, which were analyzed using FIA, also demonstrated excellent reproducibility (CV < 10 %). In general, low CVs correlated with low mean concentrations of the metabolites. However, in case of the lipaemic samples, many lipids showed higher CVs, suggesting that the upper limit for their reliable quantification was reached in these samples. It was beyond the scope of this study to ascertain whether this was due to reaching mass spectrometer detection limits, ion suppression effects, reaching the limit of solubility in the sample preparation stage, or a combination of all three.

Beyond CVs, we performed a detailed analysis of systematic and proportional differences in the measured metabolite concentrations between each pair of laboratories using Deming regression, which confirmed the overall high reproducibility of measurements. Visualisation of the detailed results in circular plots facilitated evaluating and comparing the reproducibility of measurements at low, medium, and high concentration ranges for each metabolite, and pinpointed specific patterns in the reproducibility across concentration ranges, metabolite classes, or laboratories (Figure 5). For instance, overall, the smallest relative differences between measurements occurred for the median range, while reproducibility was generally lower for low- and high-concentration ranges. The reproducibility patterns of laboratories E and J across metabolites differed slightly from those of the other laboratories for the median- and low-concentration ranges. While one of these two laboratories used a Waters instrument, the other two laboratories with Waters instruments did not show this pattern. Also, metabolites where measurements from one laboratory strongly deviated from those of the other 13 laboratories could be directly differentiated from metabolites for which quantifications generally differed across laboratories – a differentiation that is not possible based on CVs. For example, beta-alanine concentrations provided by Laboratory N differed from those of the other laboratories (red color) while the remaining laboratories agreed in the reported quantities (green color). Following up on this, we found that Laboratory N had an issue in peak integration for this metabolite. On the other hand, when it comes to chenodeoxycholic acid (CDCA), the quantifications showed considerable discrepancies between laboratories, as indicated by the red color representing multiple laboratories. To enable further examination of the comprehensive metabolite results, in addition to the examples discussed, we offer online access at https://biotools.erciyes.edu.tr/MethodCompRing/.

### Results consistent across species and sample types

The comparative analysis of metabolite coverage for the distinct sample types profiled in this study, which included human (female, male, lipaemic, NIST SRM 1950) plasma and serum (female, male) as well as rodent (mouse, rat) plasma, exhibited unique species-, sex-, matrix- and phenotype-specific differences. Nonetheless, the overall reproducibility of measurements in rodent plasma was comparable to that in human plasma. Similar transferability had been previously reported for the Lipidyzer kit, which also was originally validated for human plasma only, but performed well on mouse samples in cross-platform comparisons (39, 40). Thus, the high detectability of metabolites in these sample types and the good reproducibility of quantification in our study suggest that the MxP® Quant 500 kit can be applied beyond human plasma as an extension of the current recommended use by the manufacturer.

Evaluation of the matrix-specific CVs also indicated a high reproducibility for the quantification of at least 400 metabolites which showed CVs below 25 %. For various metabolites, CVs differed between sample types, in particular when comparing human and rodent samples. Considering the general relation of high CVs and low mean concentrations within sample types in our study, the observed differences in reproducibility of quantifications between human and rodent samples are most likely not caused by matrix effects but by species-specific differences in blood concentrations that are in the range of metabolite-specific LOD, lower limit of quantification (LLOQ; or upper limit of quantification (ULOQ) in the case of lipaemic plasma). As an example, the difference in the CVs was most striking for HexCer(d16:1/22:0) with low concentrations and a high CV in rodent plasma (mean concentration 0.009 µmol/l; CV > 53.0 %) versus higher concentrations and lower CVs in human plasma (mean concentrations 0.06 µmol/l; CV < 15.5 %). Conversely, PC ae C38:1 had low concentrations and high CVs in human plasma (mean concentrations 0.41 µmol/l; CV > 48.9 %) versus higher concentration and lower CVs in rodent plasma (mean concentration 1.97 µmol/l; CV < 11.4 %).

As concentrations can significantly vary depending on the conditions at the time of blood draw (e.g., fasting vs fed state) or with specific phenotypes, metabolites can be increased and thus more reproducibly quantified in certain circumstances. For example, PC ae C38:1 is elevated in breast cancer patients (41). Similarly, quantifications of metabolites, which were not detected by any laboratory in the samples of this study, could be reproducible in samples with higher concentrations. For instance, phenylethylamine has a very low concentration in human plasma, unless the concentration is significantly elevated e.g., in hyperactivity disorders (42, 43). The C10:1 (decenoylcarnitine) is elevated in human blood during fasting (36) or low-calorie diets (44) or in atrial fibrillation (45).

### Normalisation essential for reproducible quantification across laboratories

To ensure high reliability and reproducibility, all components of the MxP® Quant 500 kit are strongly controlled. Nevertheless, laboratory-specific differences, such as instrumental set-up, instrument condition, and variation in ambient parameters during sample preparation and measurement of each kit plate can significantly influence measurement results. This leads to batch effects when the measurement of a sample set is distributed over multiple kit plates and laboratories. To compensate for these batch effects and to generally improve the accuracy of quantifications, normalisation of the data is commonly advised (46, 47) and also strongly recommended by the manufacturer. In this study, we performed batch correction based on the measurements for four replicates of the QC2 quality control sample (provided in the kit) and the reference target values for the metabolite levels in the QC2 sample as also used in the kit software to perform normalisation for each plate. Comparing the reproducibility of the measured quantities with and without normalisation showed that this step is essential to ensure the comparability of results across laboratories.

Improvement of reproducibility after QC2-normalisation was not only observed for the profiled human plasma samples but also for the human serum and rodent plasma samples though QC2 is based on human plasma and, thus, does not necessarily meet the conditions for other sample types regarding concentration ranges and matrix effects. In general, for sample types other than human plasma, the use of study sample pools could further improve the outcome of data normalisation as they better match the conditions of the investigated sample and matrix type (48). Some normalisation approaches, such as the commonly applied QC-based robust LOESS (locally estimated scatterplot smoothing) signal correction (QC-RLSC) algorithm for intra and inter-batch correction, require that the QC samples are representative of the matrix they are modeling, otherwise, those signals are filtered out (49). However, for large studies, the preparation of a representative sample pool of sufficient volume to be used on all plates of the study is often challenging and commercially available material is used instead (50). In the present study, the reproducibility of measurements in human serum and rodent plasma samples increased considerably with the use of the plasma pool QC2, confirming that normalisation based on less representative sample pools is suitable to adjust for technical variation to some extent.

Beyond using QC2 or study sample pools, NIST SRM 1950 samples have proven useful for aligning measurements even between studies where different metabolomic kits and platforms were applied (27, 51). This specific reference material, consisting of pooled plasma samples from healthy individuals composed of equal numbers of women and men aged 40-50 years with a racial distribution reflecting the US population, can facilitate not only data harmonisation but also quality assurance (48, 52–55). However, in contrast to QC2 samples, carefully determined target values for normalisation are unavailable for most measured metabolites.

### Interlaboratory reproducibility and quantification consistent with previous studies

Two ring trials were previously conducted with biocrates kits AbsoluteIDQ p180 (29) and AbsoluteIDQ p400HR (17), with the latter being based on the same sample collection as in the present study. In those ring trials and our study, the LC-MS approach achieved a lower CV compared to FIA. This is not unexpected since matrix effects and incomplete ionisation as sources of variations likely play a larger role in FIA due to the lack of prior separation of molecules. For the AbsoluteIDQ p180 kit, using triple quadrupole mass spectrometry, the median CV between laboratories was 7.6 %, with 85 % of metabolites having a median CV between laboratories of <20 % when considering all sample types (29). While the median CV was higher in our study (14.3 %), this result is probably due to the inclusion of rodent samples and the larger set of metabolites. Focusing on the reproducibility of measurements in the NIST SRM 1950 samples, the mean interlaboratory CVs in our study were comparable to or better than those of both the AbsoluteIDQ p180 kit and the AbsoluteIDQ p400HR when identical metabolites were considered. With the AbsoluteIDQ p400HR kit, using an orbitrap mass spectrometer, interlaboratory precision was high for all analyte classes (mean CVs < 18 %), with the best performance observed for amino acids (7.1 % mean CV vs 7.4 % in our study) (17). Although the same mix of sample types including human and rodent plasma was used in the AbsoluteIDQ p400HR ring trial, we could not readily compare our results on the sample-specific CV differences as metabolites, for which we observed large differences, were not part of the overlapping analytes.

In the comparison of CVs between our study and a previous study (18) that used the lipidomics Lipidyzer kit, separating lipids using differential mobility (DMS) before MRM detection, CVs of dihydroceramides and sphingomyelins had lower CVs than that in the Lipidyzer study (14.0 % versus 25.7 %, and 11.7 % versus 20.4 %, respectively). In contrast, the Lipidyzer study had overall lower mean CVs compared to our study for the remaining lipid subclasses, including most ceramide classes, free fatty acids, diacyl- and triacylglycerols.

Beyond CVs, this study added a substantial amount of quantitative data for the NIST SRM 1950 sample to the metabolite concentrations, which have been measured and reported using various analytical methods for this reference material. Only a portion of the measured concentrations are provided as certified or reference values by NIST; additional metabolite concentrations were measured in different laboratories and have been provided by the research community (18, 27, 48, 53, 54, 56). A current study by the metabolomics quality assurance and control consortium (www.mQACC.org) is seeking to collate these results. In our study, the amino acid concentrations determined by the 14 participating laboratories generally corresponded well with the certified/reference values of the NIST (28) SRM 1950 sample. Nonetheless, for 6 out of the 15 amino acids (arginine, threonine, lysine, proline, glycine, and alanine), concentrations measured in our study were slightly but consistently higher. One of the possible explanations for this observation could be sample degeneration. A previous study reported that the concentrations of 8 out of 18 amino acids (including arginine, proline, and glycine) increased over a period of 5 years in plasma samples when stored at −80°C, while 10 did not show any significant change (57). We therefore hypothesise that the deviation of the metabolite certified/reference concentrations for the NIST SRM 1950 sample from the concentrations reported in this study is due to sample storage rather than a bias of the MxP® Quant 500 kit.

In contrast, large differences between certified/reference concentrations and concentrations measured in our study were seen for fatty acids. Those metabolites also showed very low reproducibility between the laboratories and did not pass the validation procedure by the manufacturer, suggesting that at least in blood samples representing “normal” conditions like this study’s project samples (including NIST SRM 1950), fatty acid measurements from the MxP® Quant 500 kit are not reliable. In general, free fatty acids pose problems in MS-based metabolomics workflows that build on protocols aiming at extraction of a wide range of different metabolites; fatty acids are aliphatic and do not always dissolve fully in aqueous solvents. Notably, the longest-chain fatty acids were more likely to be missing in this study. In addition, certain free fatty acids, especially C16:0 and C18:0 are ubiquitous contaminants of the environment, including common plastic lab consumables. Park et al. have previously suggested use of glass tubes, combined with extensive washing with both methanol and chloroform may be required for accurate quantification of these two species (58).

To assess the kit’s performance regarding lipids for which no certified/reference values were available from NIST, we compared the concentrations measured in our study with the ones determined in a previous ring trial which used the Lipidyzer kit (18). While the concordance of concentrations measured in our study with those reported previously was high for most triacylglycerols (MAPE <20 %), for the majority of the cholesterol esters compared, the measured concentrations differed considerably.

### Strengths and limitations

This study included a wide array of analytical instruments across multiple laboratories to assess the interlaboratory comparability of measurements under typical conditions of multi-center studies. For a thorough analysis of reproducibility, we used statistical tools recommended for method comparison in clinical applications thereby going beyond calculations of CVs. The provision and visualisation of the detailed statistical results for all tested sample types, including the widely used reference material NIST SRM 1950 as well as rodent plasma, will facilitate planning of future studies.

However, our study also comes with limitations. First, the kit manufacturer provided the materials used in this study, and the experimental conditions were strictly controlled, including the use of new chromatography columns. This tightly controlled environment might have improved consistency, but may not fully reflect the conditions encountered in routine analyses. Second, the limited sample set used in this study does not encompass the full range of metabolite concentrations observable in human and rodent blood. Due to the interrelations of concentration ranges and reproducibility of quantifications, our results might over-or underestimate the comparability of measurements. For example, in the tested samples acylcarnitine concentrations were not representative for plasma samples under fasted conditions, in which higher concentrations could have led to higher detectability and reproducibility of measurements. Third, in our study, we applied the same QC sample (QC2) to normalize metabolites, not taking specific concentration ranges into account. As a consequence, metabolites, for which levels were below LOD in the QC samples, were excluded from further analysis irrespective of their detectability in the project samples. While the impact of the choice of QC was not investigated in this study, it might be critical for specific applications of the MxP® Quant 500 kit. For example, the concentrations of various acylcarnitines in QC2 samples were low or below the LOD, limiting the effectiveness of QC2 measurements for normalisation of study samples derived from fasting subjects, in whom acylcarnitine concentrations are typically elevated. In such cases, QC3, another QC sample provided with the kit with higher metabolite concentrations, may provide a more appropriate normalisation reference. Also, only one normalisation method was used throughout our study, without testing for variation of reproducibility outcomes depending on the use of different normalisation algorithms.

## Conclusion

With this study, we have demonstrated that targeted metabolomics kits provide a high level of reproducibility and standardisation allowing consistent results across multiple laboratories using different instrumentation. Results were reproducible for human plasma, the matrix for which the MxP® Quant 500 kit was validated by the manufacturer, but we also showed that this method can be applied to closely related matrices such as rat and mouse plasma. As expected, metabolites analysed with 7-point calibration had lower CVs than those with 1-point calibration, although 70 % of the targeted metabolites were reliably detected and reproducibly quantified across the laboratories. These results were mostly comparable with other metabolomics and lipidomics kits using the NIST SRM 1950 sample as a reference. Measured metabolite concentrations showed good concordance across studies for most metabolite classes with the notable exception of fatty acids, for which reproducibility was low. Altogether, these results support the use of this technology for applications where standardisation, reproducibility, and scalability of metabolomics are necessary while enabling measurement in multiple laboratories using different LC-MS/MS instrumentation, making the MxP® Quant 500 kit a relevant tool to apply metabolomics in epidemiology and multi-center studies.

## DECLARATION OF INTERESTS

G.K. is co-inventor (through Helmholtz Zentrum München) on patents regarding applications of metabolomics in diseases of the central nervous system and holds equity in Chymia LLC. A.L., T.H.P and T.K. were employed at biocrates life sciences ag at the time the experiment was conducted and/or at the time this manuscript was being written. K.C. is currently an AstraZeneca employee.

## AUTHOR CONTRIBUTIONS

Conceptualization, J.A. and G.K.; Investigation, J.L., J.K., G.P., H.M.G., S.S., X.L.G., D.S., D.W., J.Z., R.M., L.J.-W., K.A., J.W.T., M.P.S., K.C., S.C., N.A., S.A., A.Y., S.F.G., T.M.C., K.K., T.B., T.H.P., and T.K.; Data curation, J.L., and A.C.; Formal analysis, G.E.Z and G.K.; Visualization, G.E.Z, J.A., and G.K.; Resources, T.H.P. and T.K.; Supervision, D.W., J.W.T, M.S., S.F.G., T.B., and G.K.; Project management, J.A.; Writing – Original Draft, J.L., G.E.Z, J.A., and G.K.; Writing – Review & Editing, all authors.

## FUNDING SOURCES

G.K. received funding (through her institution) from the National Institutes of Health/National Institute on Aging through grants RF1AG057452, RF1AG058942, RF1AG059093, U01AG061359, and U19AG063744. G.P. received funding by the German Federal Ministry of Education and Research (BMBF) within the SMART-CARE consortium (fund no. 161L0212). S.F.G receives funding (through his Institution) from the National Institutes of Health through grants R01NS110838-01A1, R01ES032450, R01NS114409, R01AG079388-01, and R33AG067083. S.F.G also receives funding from Innovate UK, SBRI, and numerous internal funding programs. R.M. and D.W. received funding from the Canada Foundation for Innovation (CFI), Genome Alberta, a division of Genome Canada and Alberta Innovates.

---

**Supplementary Figure S1.**
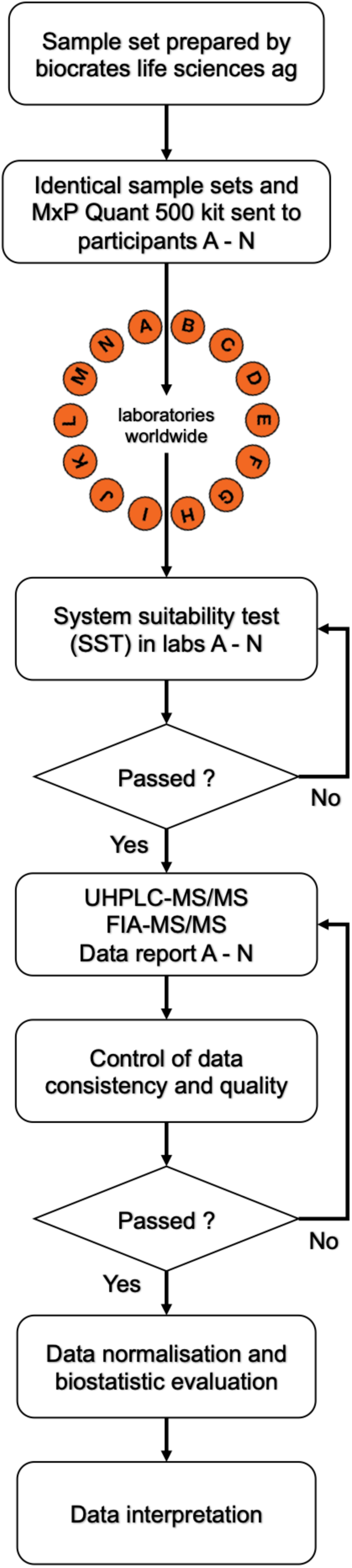
Description of the project workflow. The identical sample set was aliquoted by biocrates and sent directly to the participating laboratories. The laboratories, which were pseudonymised using capital letters A to N, performed a system suitability test (SST), which was checked by biocrates to proceed with the project sample analyses. The data from sample measurement were sent to the Helmholtz Zentrum München, where the consistency and quality of the data was checked before performing the statistical analyses. The resulting findings were interpreted by all partners from all participating laboratories.

## Notes

### Competing Interest Statement

G.K. is co-inventor (through Helmholtz Munich) on patents regarding applications of metabolomics in diseases of the central nervous system and holds equity in Chymia LLC. A.L., T.H.P and T.K. were employed at biocrates life sciences ag at the time the experiment was conducted and/or at the time this manuscript was being written. K.C. is currently an AstraZeneca employee.

