## Supplementary Data S1 for "Interlaboratory comparison of standardised metabolomics and lipidomics analyses in human and rodent blood using the MxP® Quant 500 kit": Cer(d16.1-22.0).pdf

**Presence of systematic differences between laboratories**

| | A | B | C | D | E | F | G | H | I | J | K | L | M | N |
|---|---|---|---|---|---|---|---|---|---|---|---|---|---|---|
| A | - | N | N | N | P | N | N | N | N | N | N | P | N | N |
| B | - | - | N | P | N | N | N | N | N | N | N | N | N | N |
| C | - | - | - | N | C | C | N | N | N | N | N | P | N | N |
| D | - | - | - | - | N | N | N | N | P | N | N | P | N | N |
| E | - | - | - | - | - | CP | N | N | C | N | N | C | C | N |
| F | - | - | - | - | - | - | N | N | CP | N | N | C | C | C |
| G | - | - | - | - | - | - | - | N | P | N | N | N | N | N |
| H | - | - | - | - | - | - | - | - | N | N | N | N | N | N |
| I | - | - | - | - | - | - | - | - | - | N | N | P | N | N |
| J | - | - | - | - | - | - | - | - | - | - | N | N | N | N |
| K | - | - | - | - | - | - | - | - | - | - | - | N | N | N |
| L | - | - | - | - | - | - | - | - | - | - | - | - | N | N |
| M | - | - | - | - | - | - | - | - | - | - | - | - | - | N |
| N | - | - | - | - | - | - | - | - | - | - | - | - | - | - |

| |
|---|
| N |
| C |
| P |
| CP |
| NA |

| |
|---|
| **No difference** |
| **Constanat difference** |
| **Proportional difference** |
| **Both constant and proportional difference** |
| **Calculation failed** |
