## Supplementary Data S1 for "Interlaboratory comparison of standardised metabolomics and lipidomics analyses in human and rodent blood using the MxP® Quant 500 kit": PC aa C28.1.pdf

**Presence of systematic differences between laboratories**

| | A | B | C | D | E | F | G | H | I | J | K | L | M | N |
|---|---|---|---|---|---|---|---|---|---|---|---|---|---|---|
| A | - | N | N | P | P | N | CP | N | CP | N | P | N | P | P |
| B | - | - | N | P | P | N | P | N | CP | N | P | N | P | P |
| C | - | - | - | P | P | N | P | N | CP | N | P | N | N | N |
| D | - | - | - | - | P | P | N | P | CP | P | CP | P | P | P |
| E | - | - | - | - | - | P | P | P | CP | P | P | P | P | P |
| F | - | - | - | - | - | - | P | P | CP | N | N | N | P | N |
| G | - | - | - | - | - | - | - | P | CP | P | N | CP | P | CP |
| H | - | - | - | - | - | - | - | - | CP | N | N | C | P | P |
| I | - | - | - | - | - | - | - | - | - | CP | CP | CP | CP | CP |
| J | - | - | - | - | - | - | - | - | - | - | N | N | N | N |
| K | - | - | - | - | - | - | - | - | - | - | - | N | N | N |
| L | - | - | - | - | - | - | - | - | - | - | - | - | N | N |
| M | - | - | - | - | - | - | - | - | - | - | - | - | - | N |
| N | - | - | - | - | - | - | - | - | - | - | - | - | - | - |

| | |
|---|---|
| N | No difference |
| C | Constanat difference |
| P | Proportional difference |
| CP | Both constant and proportional difference |
| NA | Calculation failed |
