## Supplementary Data S2 for "Interlaboratory comparison of standardised metabolomics and lipidomics analyses in human and rodent blood using the MxP® Quant 500 kit": 1-Met-His.pdf

**Amount of relative differences between laboratories**

| Min | A | B | C | D | E | F | G | H | I | J | K | L | M | N |
|---|---|---|---|---|---|---|---|---|---|---|---|---|---|---|
| A | - | 2.262 | -0.28 | 1.692 | 2.655 | 2.341 | 2.022 | 2.764 | 4.541 | 1.768 | 0.656 | 3.04 | 1.66 | 1.066 |
| B | - | - | 1.934 | 0.537 | 0.916 | -0.935 | 3.196 | 1.429 | 4.106 | 0.146 | 1.835 | 1.78 | 0.441 | 1.367 |
| C | - | - | - | 1.124 | 2.372 | 2.067 | 2.382 | 2.579 | 4.34 | 1.367 | -0.712 | 2.829 | 1.142 | 0.264 |
| D | - | - | - | - | 1.667 | 1.069 | 2.901 | 1.982 | 4.319 | -0.963 | 0.681 | 2.349 | -6.66 | 0.067 |
| E | - | - | - | - | - | 0.338 | 3.479 | 0.022 | 4.26 | 1.625 | 2.299 | 1.106 | 1.703 | 2.127 |
| F | - | - | - | - | - | - | 3.261 | 0.661 | 4.355 | 0.744 | 1.918 | 1.714 | 1.071 | 1.676 |
| G | - | - | - | - | - | - | - | 3.457 | 4.846 | 2.831 | 2.391 | 3.586 | 2.841 | 2.613 |
| H | - | - | - | - | - | - | - | - | 3.61 | 1.975 | 2.561 | 0.328 | 1.913 | 2.284 |
| I | - | - | - | - | - | - | - | - | - | 3.51 | 3.743 | 3.212 | 3.841 | 3.916 |
| J | - | - | - | - | - | - | - | - | - | - | 1.091 | 2.458 | -2.066 | -0.111 |
| K | - | - | - | - | - | - | - | - | - | - | - | 2.767 | 0.338 | -0.593 |
| L | - | - | - | - | - | - | - | - | - | - | - | - | 2.368 | 2.643 |
| M | - | - | - | - | - | - | - | - | - | - | - | - | - | 0.252 |
| N | - | - | - | - | - | - | - | - | - | - | - | - | - | - |

| | |
|---|---|
| | Larger than 10 |
| | Between 8 and 9,99 |
| | Between 6 and 7,99 |
| | Between 4 and 5,99 |
| | Between 2 and 3,99 |
| | Between 0 and 1,99 |
| | NA (Calculation failed) |

| Median | A | B | C | D | E | F | G | H | I | J | K | L | M | N |
|---|---|---|---|---|---|---|---|---|---|---|---|---|---|---|
| A | - | 1.745 | -0.301 | 0.528 | 2.548 | 2.075 | 1.765 | 2.829 | -0.074 | 2.23 | -1.352 | 2.013 | 2.047 | 1.143 |
| B | - | - | 2.103 | 0.986 | 1.408 | -0.256 | 2.802 | 1.958 | 1.947 | 0.487 | 1.612 | -0.473 | -0.233 | 0.224 |
| C | - | - | - | 1.141 | 2.736 | 2.294 | 1.359 | 2.959 | -0.909 | 2.448 | 0.305 | 2.274 | 2.301 | 1.586 |
| D | - | - | - | - | 2.19 | 1.468 | 2.284 | 2.513 | 0.918 | 1.739 | -0.01 | 1.43 | 1.476 | -0.305 |
| E | - | - | - | - | - | 0.826 | 3.3 | 0.452 | 2.858 | 0.323 | 2.534 | 0.943 | 0.869 | 1.953 |
| F | - | - | - | - | - | - | 2.979 | 1.575 | 2.395 | -0.862 | 1.972 | -3.149 | -5.049 | 1.035 |
| G | - | - | - | - | - | - | - | 3.362 | 0.96 | 2.968 | 1.864 | 2.847 | 2.868 | 2.446 |
| H | - | - | - | - | - | - | - | - | 3.025 | 1.405 | 2.865 | 1.748 | 1.684 | 2.398 |
| I | - | - | - | - | - | - | - | - | - | 0.688 | 1.278 | 0.066 | 0.075 | 0.13 |
| J | - | - | - | - | - | - | - | - | - | - | 2.195 | -0.533 | -0.804 | 1.362 |
| K | - | - | - | - | - | - | - | - | - | - | - | 1.875 | 1.892 | 0.857 |
| L | - | - | - | - | - | - | - | - | - | - | - | - | -3.672 | 0.987 |
| M | - | - | - | - | - | - | - | - | - | - | - | - | - | 1 |
| N | - | - | - | - | - | - | - | - | - | - | - | - | - | - |

| Max | A | B | C | D | E | F | G | H | I | J | K | L | M | N |
|---|---|---|---|---|---|---|---|---|---|---|---|---|---|---|
| A | - | 1.32 | 1.004 | -1.917 | 2.481 | 1.837 | 1.561 | 2.875 | 2.187 | 2.49 | -1.194 | 0.513 | 2.237 | 1.19 |
| B | - | - | 2.214 | 1.22 | 1.635 | 0.102 | 2.457 | 2.254 | 2.806 | 1.682 | 1.433 | -0.045 | 1.119 | -1.911 |
| C | - | - | - | 1.153 | 2.913 | 2.443 | -0.364 | 3.188 | 1.263 | 2.913 | 0.742 | 1.775 | 2.713 | 2.055 |
| D | - | - | - | - | 2.427 | 1.708 | 1.639 | 2.81 | 2.362 | 2.426 | -0.821 | 0.261 | 2.151 | 0.998 |
| E | - | - | - | - | - | 1.107 | 3.165 | 0.705 | 3.483 | -3.871 | 2.679 | 2.157 | -0.041 | 1.83 |
| F | - | - | - | - | - | - | 2.751 | 2.005 | 3.184 | 1.105 | 2.009 | 1.155 | 0.274 | 0.396 |
| G | - | - | - | - | - | - | - | 3.287 | 1.032 | 3.06 | 1.331 | 2.07 | 2.884 | 2.328 |
| H | - | - | - | - | - | - | - | - | 3.481 | 0.787 | 3.044 | 2.64 | 1.527 | 2.467 |
| I | - | - | - | - | - | - | - | - | - | 3.456 | 2.391 | 2.901 | 3.359 | 2.975 |
| J | - | - | - | - | - | - | - | - | - | - | 2.654 | 2.184 | 0.082 | 1.856 |
| K | - | - | - | - | - | - | - | - | - | - | - | 0.776 | 2.372 | 1.349 |
| L | - | - | - | - | - | - | - | - | - | - | - | - | 1.685 | -0.423 |
| M | - | - | - | - | - | - | - | - | - | - | - | - | - | 1.333 |
| N | - | - | - | - | - | - | - | - | - | - | - | - | - | - |
