## Supplementary Data S2 for "Interlaboratory comparison of standardised metabolomics and lipidomics analyses in human and rodent blood using the MxP® Quant 500 kit": 3-IAA.pdf

**Amount of relative differences between laboratories**

| Min | A | B | C | D | E | F | G | H | I | J | K | L | M | N |
|---|---|---|---|---|---|---|---|---|---|---|---|---|---|---|
| A | - | -4.941 | 3.129 | 3.892 | 5.201 | 2.436 | 1.307 | 1.931 | 4.223 | 1.757 | 5.384 | 5.692 | 3.518 | 2.703 |
| B | - | - | 3.3 | 3.774 | 5.217 | 2.16 | 0.399 | 0.909 | 4.082 | 1.888 | 5.345 | 5.675 | 3.346 | 3.042 |
| C | - | - | - | 4.621 | 5.55 | 3.897 | 3.568 | 2.989 | 4.791 | 2.684 | 5.042 | 5.481 | 4.377 | 0.544 |
| D | - | - | - | - | 4.844 | 3.082 | 3.563 | 4.152 | 2.652 | 4.077 | 5.743 | 5.938 | 1.47 | 4.258 |
| E | - | - | - | - | - | 4.769 | 4.98 | 5.139 | 4.27 | 5.123 | 5.937 | 6.124 | 4.489 | 5.276 |
| F | - | - | - | - | - | - | 1.935 | 3.342 | 3.977 | 3.254 | 5.652 | 5.848 | 2.669 | 3.576 |
| G | - | - | - | - | - | - | - | 1.956 | 4.036 | 2.314 | 5.397 | 5.7 | 3.103 | 3.163 |
| H | - | - | - | - | - | - | - | - | 4.075 | 1.119 | 5.228 | 5.607 | 3.502 | 2.893 |
| I | - | - | - | - | - | - | - | - | - | 4.731 | 6.027 | 6.126 | 3.529 | 4.836 |
| J | - | - | - | - | - | - | - | - | - | - | 5.273 | 5.637 | 3.854 | 2.163 |
| K | - | - | - | - | - | - | - | - | - | - | - | 4.247 | 5.835 | 5.048 |
| L | - | - | - | - | - | - | - | - | - | - | - | - | 6.359 | 5.783 |
| M | - | - | - | - | - | - | - | - | - | - | - | - | - | 4.072 |
| N | - | - | - | - | - | - | - | - | - | - | - | - | - | - |

| Median | A | B | C | D | E | F | G | H | I | J | K | L | M | N |
|---|---|---|---|---|---|---|---|---|---|---|---|---|---|---|
| A | - | 1.496 | 0.435 | 0.922 | 5.062 | 3.047 | 1.312 | 3.083 | 1.997 | 0.604 | 0.839 | 2.346 | -0.327 | 1.225 |
| B | - | - | 2.039 | -0.12 | 4.91 | 3.433 | -1.466 | 2.473 | 2.734 | 2.084 | 2.157 | 2.945 | 1.827 | -0.894 |
| C | - | - | - | 1.742 | 5.158 | 2.802 | 1.98 | 3.322 | 1.385 | -2.643 | -1.413 | 1.922 | -0.737 | 1.923 |
| D | - | - | - | - | 4.959 | 3.311 | -0.812 | 2.732 | 2.519 | 1.69 | 1.827 | 2.744 | 1.392 | -1.312 |
| E | - | - | - | - | - | 4.969 | 4.548 | 4.229 | 4.821 | 4.715 | 4.724 | 4.856 | 4.685 | 4.561 |
| F | - | - | - | - | - | - | 3.541 | 4.204 | 2.205 | 2.895 | 2.82 | 1.859 | 3.019 | 3.516 |
| G | - | - | - | - | - | - | - | 2.569 | 2.652 | 1.954 | 2.041 | 2.876 | 1.682 | -2.701 |
| H | - | - | - | - | - | - | - | - | 3.522 | 3.211 | 3.236 | 3.65 | 3.094 | 2.509 |
| I | - | - | - | - | - | - | - | - | - | 1.46 | 1.191 | -0.156 | 1.716 | 2.67 |
| J | - | - | - | - | - | - | - | - | - | - | -2.07 | 1.861 | -0.477 | 1.953 |
| K | - | - | - | - | - | - | - | - | - | - | - | 1.905 | 0.089 | 2.126 |
| L | - | - | - | - | - | - | - | - | - | - | - | - | 2.346 | 3.102 |
| M | - | - | - | - | - | - | - | - | - | - | - | - | - | 1.637 |
| N | - | - | - | - | - | - | - | - | - | - | - | - | - | - |

| Max | A | B | C | D | E | F | G | H | I | J | K | L | M | N |
|---|---|---|---|---|---|---|---|---|---|---|---|---|---|---|
| A | - | 1.687 | -1.773 | -4.213 | 5.014 | 3.331 | 1.313 | 3.341 | 2.91 | 0.34 | 2.249 | 1.898 | 1.354 | 1.963 |
| B | - | - | 1.717 | 1.583 | 4.794 | 3.679 | -0.705 | 2.713 | 3.368 | 2.109 | 0.549 | -0.996 | 2.464 | -0.286 |
| C | - | - | - | -1.641 | 5.003 | 3.3 | 1.466 | 3.361 | 2.873 | 0.084 | 2.309 | 1.976 | 1.199 | 2.072 |
| D | - | - | - | - | 4.995 | 3.339 | 1.277 | 3.314 | 2.926 | 0.444 | 2.213 | 1.856 | 1.38 | 1.945 |
| E | - | - | - | - | - | 4.993 | 4.471 | 4.046 | 4.891 | 4.65 | 4.373 | 4.409 | 4.711 | 4.388 |
| F | - | - | - | - | - | - | 3.803 | 4.478 | 1.524 | 3.287 | 4.04 | 3.926 | 3.063 | 3.96 |
| G | - | - | - | - | - | - | - | 2.86 | 3.295 | 1.897 | 1.077 | 0.141 | 2.294 | 0.485 |
| H | - | - | - | - | - | - | - | - | 3.991 | 3.354 | 2.271 | 2.514 | 3.51 | 2.429 |
| I | - | - | - | - | - | - | - | - | - | 2.756 | 3.737 | 3.601 | 2.437 | 3.647 |
| J | - | - | - | - | - | - | - | - | - | - | 2.597 | 2.346 | 0.327 | 2.418 |
| K | - | - | - | - | - | - | - | - | - | - | - | -0.209 | 2.803 | -0.504 |
| L | - | - | - | - | - | - | - | - | - | - | - | - | 2.578 | -1.379 |
| M | - | - | - | - | - | - | - | - | - | - | - | - | - | 2.728 |
| N | - | - | - | - | - | - | - | - | - | - | - | - | - | - |

| Colour | Range |
|---|---|
| Red | Larger than 10 |
| Orange | Between 8 and 9,99 |
| Light orange | Between 6 and 7,99 |
| Light yellow-green | Between 4 and 5,99 |
| Light green | Between 2 and 3,99 |
| Dark green | Between 0 and 1,99 |
| Light grey | NA (Calculation failed) |
