## Supplementary Data S2 for "Interlaboratory comparison of standardised metabolomics and lipidomics analyses in human and rodent blood using the MxP® Quant 500 kit": AA.pdf

**Amount of relative differences between laboratories**

| Min | A | B | C | D | E | F | G | H | I | J | K | L | M | N |
|---|---|---|---|---|---|---|---|---|---|---|---|---|---|---|
| A | - | 5.953 | 6.3 | 5.577 | 6.608 | 5.154 | 6.612 | 4.35 | 3.453 | 3.395 | 3.61 | 5.459 | 5.003 | 4.767 |
| B | - | - | 4.36 | 0.63 | 5.492 | 4.27 | 5.935 | 3.973 | 6.077 | 5.379 | 6.268 | 3.457 | 0.645 | 1.351 |
| C | - | - | - | 4.085 | 4.774 | 5.331 | 5.555 | 5.079 | 6.554 | 5.964 | 6.679 | 4.983 | 3.608 | 2.307 |
| D | - | - | - | - | 5.579 | 4.405 | 6.004 | 4.439 | 6.164 | 5.514 | 6.364 | 3.621 | 0.935 | 0.044 |
| E | - | - | - | - | - | 6.322 | 4.866 | 6.014 | 7.141 | 6.714 | 7.284 | 6.127 | 5.309 | 5.003 |
| F | - | - | - | - | - | - | 6.237 | 3.51 | 5.714 | 4.672 | 5.847 | 3.167 | 3.224 | 3.23 |
| G | - | - | - | - | - | - | - | 6.543 | 7.552 | 7.166 | 7.652 | 6.822 | 5.977 | 4.881 |
| H | - | - | - | - | - | - | - | - | 5.318 | 3.307 | 5.062 | 5.511 | 3.94 | 2.719 |
| I | - | - | - | - | - | - | - | - | - | 4.479 | 1.479 | 5.918 | 5.223 | 4.858 |
| J | - | - | - | - | - | - | - | - | - | - | 4.674 | 4.919 | 4.756 | 4.686 |
| K | - | - | - | - | - | - | - | - | - | - | - | 5.616 | 5.328 | 5.144 |
| L | - | - | - | - | - | - | - | - | - | - | - | - | 2.516 | 2.326 |
| M | - | - | - | - | - | - | - | - | - | - | - | - | - | 2.48 |
| N | - | - | - | - | - | - | - | - | - | - | - | - | - | - |

| | |
|---|---|
| | Larger than 10 |
| | Between 8 and 9,99 |
| | Between 6 and 7,99 |
| | Between 4 and 5,99 |
| | Between 2 and 3,99 |
| | Between 0 and 1,99 |
| | NA (Calculation failed) |

| Median | A | B | C | D | E | F | G | H | I | J | K | L | M | N |
|---|---|---|---|---|---|---|---|---|---|---|---|---|---|---|
| A | - | 2.541 | 3.729 | 3.605 | 4.566 | 3.517 | 5.24 | 4.533 | 2.53 | 2.292 | 3.983 | 3.288 | 4.165 | 4.009 |
| B | - | - | 2.556 | 2.662 | 4.318 | 2.13 | 4.768 | 4.216 | -2.865 | 3.41 | 4.727 | 1.369 | 3.585 | 3.828 |
| C | - | - | - | -0.551 | 3.977 | 0.53 | 4.358 | 3.717 | 2.609 | 4.052 | 5.102 | 1.736 | 2.7 | 3.547 |
| D | - | - | - | - | 3.852 | 0.934 | 4.564 | 3.622 | 2.683 | 4.191 | 5.118 | 1.942 | 2.619 | 3.181 |
| E | - | - | - | - | - | 4.4 | 2.137 | 1.285 | 4.681 | 5.265 | 5.925 | 4.546 | 3.492 | 2.315 |
| F | - | - | - | - | - | - | 4.687 | 3.62 | 2.484 | 4.084 | 4.983 | 0.932 | 2.927 | 3.104 |
| G | - | - | - | - | - | - | - | 1.359 | 4.695 | 5.208 | 5.954 | 4.586 | 3.576 | 2.625 |
| H | - | - | - | - | - | - | - | - | 4.518 | 5.176 | 5.588 | 3.7 | 2.65 | 3.322 |
| I | - | - | - | - | - | - | - | - | - | 3.505 | 4.409 | 2.396 | 3.734 | 3.235 |
| J | - | - | - | - | - | - | - | - | - | - | 3.565 | 3.725 | 4.417 | 4.426 |
| K | - | - | - | - | - | - | - | - | - | - | - | 4.473 | 4.874 | 4.782 |
| L | - | - | - | - | - | - | - | - | - | - | - | - | 3.241 | 3.389 |
| M | - | - | - | - | - | - | - | - | - | - | - | - | - | 1.859 |
| N | - | - | - | - | - | - | - | - | - | - | - | - | - | - |

| Max | A | B | C | D | E | F | G | H | I | J | K | L | M | N |
|---|---|---|---|---|---|---|---|---|---|---|---|---|---|---|
| A | - | 4.29 | 4.462 | 3.104 | 3.156 | -1.283 | 5.299 | 4.623 | 3.628 | 3.695 | 4.154 | 3.005 | 2.497 | 2.947 |
| B | - | - | 1.005 | 3.189 | 3.152 | 3.999 | 4.034 | 5.21 | 4.743 | 2.471 | 0.6 | 3.268 | 4.399 | 4.522 |
| C | - | - | - | 3.459 | 3.41 | 4.138 | 3.764 | 5.263 | 4.833 | 2.877 | 1.744 | 3.507 | 4.51 | 4.599 |
| D | - | - | - | - | -1.765 | 2.916 | 4.829 | 4.936 | 4.27 | 1.978 | 3.064 | -0.848 | 3.719 | 3.919 |
| E | - | - | - | - | - | 2.963 | 4.778 | 4.955 | 4.308 | 1.89 | 3.025 | -0.289 | 3.756 | 3.947 |
| F | - | - | - | - | - | - | 5.252 | 4.646 | 3.686 | 3.643 | 4.109 | 2.92 | 2.606 | 3.002 |
| G | - | - | - | - | - | - | - | 5.53 | 5.24 | 4.23 | 3.91 | 4.48 | 5.026 | 5.045 |
| H | - | - | - | - | - | - | - | - | 4.02 | 5.64 | 5.814 | 5.441 | 4.651 | 4.492 |
| I | - | - | - | - | - | - | - | - | - | 4.855 | 5.11 | 4.549 | 2.936 | 2.419 |
| J | - | - | - | - | - | - | - | - | - | - | 2.09 | 2.13 | 4.039 | 4.194 |
| K | - | - | - | - | - | - | - | - | - | - | - | 3.047 | 4.312 | 4.434 |
| L | - | - | - | - | - | - | - | - | - | - | - | - | 3.667 | 3.868 |
| M | - | - | - | - | - | - | - | - | - | - | - | - | - | 1.109 |
| N | - | - | - | - | - | - | - | - | - | - | - | - | - | - |
