## Supplementary Data S2 for "Interlaboratory comparison of standardised metabolomics and lipidomics analyses in human and rodent blood using the MxP® Quant 500 kit": ADMA.pdf

**Amount of relative differences between laboratories**

| Min | A | B | C | D | E | F | G | H | I | J | K | L | M | N |
|---|---|---|---|---|---|---|---|---|---|---|---|---|---|---|
| A | - | 3.545 | 3.717 | 3.042 | 4.349 | 3.853 | 3.125 | 3.369 | 2.974 | 3.714 | 2.251 | 3.892 | 4.23 | -0.486 |
| B | - | - | -1.303 | 1.893 | 3.317 | 1.304 | 2.218 | 0.755 | 2.331 | 0.118 | 4.263 | 1.508 | 2.589 | 3.567 |
| C | - | - | - | 1.583 | 3.528 | 1.854 | 2.189 | 0.661 | 1.826 | 0.968 | 4.032 | 2.23 | 2.028 | 3.206 |
| D | - | - | - | - | 3.815 | 2.706 | 0.387 | 0.282 | -0.104 | 2.291 | 3.725 | 2.857 | 2.85 | 2.642 |
| E | - | - | - | - | - | 3.148 | 4.182 | 3.885 | 4.096 | 3.422 | 4.98 | 2.953 | 3.048 | 4.556 |
| F | - | - | - | - | - | - | 3.043 | 2.477 | 2.806 | 0.75 | 4.393 | -0.002 | 0.287 | 3.79 |
| G | - | - | - | - | - | - | - | 1.376 | -1.45 | 2.532 | 3.657 | 3.005 | 3.21 | 2.578 |
| H | - | - | - | - | - | - | - | - | 1.651 | 1.829 | 4.031 | 2.41 | 2.974 | 3.172 |
| I | - | - | - | - | - | - | - | - | - | 2.478 | 3.802 | 2.84 | 3.473 | 2.817 |
| J | - | - | - | - | - | - | - | - | - | - | 4.293 | 1.242 | 1.984 | 3.67 |
| K | - | - | - | - | - | - | - | - | - | - | - | 4.238 | 4.616 | 2.46 |
| L | - | - | - | - | - | - | - | - | - | - | - | - | -1.783 | 3.885 |
| M | - | - | - | - | - | - | - | - | - | - | - | - | - | 3.948 |
| N | - | - | - | - | - | - | - | - | - | - | - | - | - | - |

| | |
|---|---|
| | Larger than 10 |
| | Between 8 and 9,99 |
| | Between 6 and 7,99 |
| | Between 4 and 5,99 |
| | Between 2 and 3,99 |
| | Between 0 and 1,99 |
| | NA (Calculation failed) |

| Median | A | B | C | D | E | F | G | H | I | J | K | L | M | N |
|---|---|---|---|---|---|---|---|---|---|---|---|---|---|---|
| A | - | 3.172 | 1.443 | -0.789 | 3.813 | 2.947 | 1.103 | 2.701 | 2.568 | 3.009 | 1.436 | 3.129 | 3.155 | -2.882 |
| B | - | - | 2.822 | 3.21 | 2.452 | 0.688 | 2.931 | 1.487 | 1.8 | -0.186 | 3.778 | -0.503 | -3.601 | 3.31 |
| C | - | - | - | 1.174 | 3.555 | 2.39 | -2.057 | 1.886 | 1.894 | 2.673 | 2.725 | 2.591 | 2.766 | 1.475 |
| D | - | - | - | - | 3.776 | 2.877 | 0.965 | 2.524 | 2.504 | 3.066 | 2.087 | 3.006 | 3.139 | -0.958 |
| E | - | - | - | - | - | 2.881 | 3.768 | 3.19 | 3.194 | 2.611 | 4.378 | 2.7 | 2.495 | 4.042 |
| F | - | - | - | - | - | - | 2.549 | 0.738 | 0.605 | 0.057 | 3.617 | -0.183 | 0.644 | 3.075 |
| G | - | - | - | - | - | - | - | 2.063 | 1.98 | 2.677 | 2.545 | 2.684 | 2.803 | 1.214 |
| H | - | - | - | - | - | - | - | - | -0.797 | 1.147 | 3.386 | 1.214 | 1.534 | 2.736 |
| I | - | - | - | - | - | - | - | - | - | 1.232 | 3.286 | 1.417 | 1.697 | 2.634 |
| J | - | - | - | - | - | - | - | - | - | - | 3.69 | -3.061 | -0.619 | 3.212 |
| K | - | - | - | - | - | - | - | - | - | - | - | 3.452 | 3.471 | 1.467 |
| L | - | - | - | - | - | - | - | - | - | - | - | - | -0.451 | 3.219 |
| M | - | - | - | - | - | - | - | - | - | - | - | - | - | 3.341 |
| N | - | - | - | - | - | - | - | - | - | - | - | - | - | - |

| Max | A | B | C | D | E | F | G | H | I | J | K | L | M | N |
|---|---|---|---|---|---|---|---|---|---|---|---|---|---|---|
| A | - | 1.517 | 4.234 | 4.27 | 2.092 | 3.025 | 3.301 | 0.516 | 1.04 | 0.614 | 1.322 | 2.918 | 2.557 | 0.523 |
| B | - | - | 4.493 | 4.563 | 2.918 | 3.488 | 3.706 | 2.355 | -1.435 | 2.199 | -2.521 | 3.35 | 3.207 | 2.223 |
| C | - | - | - | -1.563 | 3.631 | 3.182 | 2.934 | 3.87 | 4.195 | 3.88 | 4.177 | 3.261 | 3.445 | 3.884 |
| D | - | - | - | - | 3.659 | 3.215 | 2.982 | 3.921 | 4.245 | 3.902 | 4.21 | 3.341 | 3.487 | 3.913 |
| E | - | - | - | - | - | 1.94 | 2.421 | 1.355 | 2.773 | 1.376 | 2.707 | 1.662 | 0.746 | 1.443 |
| F | - | - | - | - | - | - | 0.586 | 2.679 | 3.246 | 2.629 | 3.305 | -0.825 | 1.053 | 2.648 |
| G | - | - | - | - | - | - | - | 2.899 | 3.478 | 2.917 | 3.503 | 1.104 | 1.832 | 2.928 |
| H | - | - | - | - | - | - | - | - | 2.196 | -1.671 | 2.166 | 2.509 | 2.186 | -1.962 |
| I | - | - | - | - | - | - | - | - | - | 1.905 | -0.199 | 3.222 | 3.096 | 1.975 |
| J | - | - | - | - | - | - | - | - | - | - | 2.017 | 2.557 | 2.107 | -2.739 |
| K | - | - | - | - | - | - | - | - | - | - | - | 3.357 | 3.181 | 2.227 |
| L | - | - | - | - | - | - | - | - | - | - | - | - | 0.478 | 2.466 |
| M | - | - | - | - | - | - | - | - | - | - | - | - | - | 2.099 |
| N | - | - | - | - | - | - | - | - | - | - | - | - | - | - |
