## Supplementary Data S2 for "Interlaboratory comparison of standardised metabolomics and lipidomics analyses in human and rodent blood using the MxP® Quant 500 kit": Asn.pdf

**Amount of relative differences between laboratories**

| Min | A | B | C | D | E | F | G | H | I | J | K | L | M | N |
|---|---|---|---|---|---|---|---|---|---|---|---|---|---|---|
| A | - | 11.345 | 9.333 | 8.501 | 5.543 | 8.329 | 8.436 | 8.896 | 8.605 | 8.586 | 9.606 | 11.383 | 6.624 | 7.67 |
| B | - | - | 2.114 | 1.991 | -0.225 | 2.017 | 2.736 | 1.522 | -1.719 | 2.235 | 2.725 | 1.001 | 0.57 | 3.007 |
| C | - | - | - | -1.001 | 2.31 | -1.605 | 1.174 | 2.811 | 2.072 | -1.112 | 1.286 | 1.13 | 1.511 | 1.842 |
| D | - | - | - | - | 1.962 | -1.851 | 1.688 | 2.832 | 1.717 | 0.511 | 1.918 | 0.526 | 1.088 | 2.149 |
| E | - | - | - | - | - | 2.159 | 2.9 | 1.581 | -0.528 | 2.492 | 2.985 | 1.36 | 0.969 | 3.124 |
| F | - | - | - | - | - | - | 1.684 | 2.916 | 1.736 | 0.529 | 2.052 | 0.71 | 1.137 | 2.121 |
| G | - | - | - | - | - | - | - | 3.293 | 2.639 | 0.789 | -0.997 | 2.175 | 2.368 | 0.245 |
| H | - | - | - | - | - | - | - | - | 1.615 | 3.056 | 3.399 | 2.421 | 2.312 | 3.522 |
| I | - | - | - | - | - | - | - | - | - | 2.282 | 2.676 | 1.261 | 0.746 | 3.042 |
| J | - | - | - | - | - | - | - | - | - | - | 1.363 | 1.54 | 1.856 | 1.534 |
| K | - | - | - | - | - | - | - | - | - | - | - | 2.487 | 2.594 | -2.492 |
| L | - | - | - | - | - | - | - | - | - | - | - | - | -0.774 | 2.579 |
| M | - | - | - | - | - | - | - | - | - | - | - | - | - | 2.727 |
| N | - | - | - | - | - | - | - | - | - | - | - | - | - | - |

| | |
|---|---|
| | Larger than 10 |
| | Between 8 and 9,99 |
| | Between 6 and 7,99 |
| | Between 4 and 5,99 |
| | Between 2 and 3,99 |
| | Between 0 and 1,99 |
| | NA (Calculation failed) |

| Median | A | B | C | D | E | F | G | H | I | J | K | L | M | N |
|---|---|---|---|---|---|---|---|---|---|---|---|---|---|---|
| A | - | 10.074 | 7.631 | 5.709 | 4.64 | 5.977 | 5.882 | 5.325 | 6.322 | 5.987 | 8.401 | 9.846 | 3.05 | 4.152 |
| B | - | - | 1.828 | -0.739 | 1.478 | -0.804 | 2.007 | 2.184 | 0.923 | -0.53 | 1.881 | -2.003 | 0.163 | 2.229 |
| C | - | - | - | 1.484 | 2.617 | 1.992 | -0.908 | 2.936 | 0.539 | 1.405 | -3.243 | 1.852 | 2.186 | 0.223 |
| D | - | - | - | - | 1.817 | 0.469 | 1.801 | 2.279 | 0.522 | -2.046 | 1.435 | -0.152 | 0.984 | 2.024 |
| E | - | - | - | - | - | 1.16 | 2.854 | 0.506 | 2.354 | 1.953 | 2.698 | 1.426 | 0.705 | 2.97 |
| F | - | - | - | - | - | - | 2.358 | 1.675 | 1.597 | 0.897 | 2.032 | -0.832 | -0.797 | 2.503 |
| G | - | - | - | - | - | - | - | 3.009 | 0.966 | 1.647 | -0.498 | 2.08 | 2.393 | -0.812 |
| H | - | - | - | - | - | - | - | - | 2.728 | 2.436 | 3.056 | 2.11 | 1.761 | 3.243 |
| I | - | - | - | - | - | - | - | - | - | 0.343 | 1.051 | 1.012 | 1.499 | 1.446 |
| J | - | - | - | - | - | - | - | - | - | - | 1.173 | 0.284 | 1.242 | 1.91 |
| K | - | - | - | - | - | - | - | - | - | - | - | 1.761 | 2.153 | 0.614 |
| L | - | - | - | - | - | - | - | - | - | - | - | - | 0.029 | 2.324 |
| M | - | - | - | - | - | - | - | - | - | - | - | - | - | 2.616 |
| N | - | - | - | - | - | - | - | - | - | - | - | - | - | - |

| Max | A | B | C | D | E | F | G | H | I | J | K | L | M | N |
|---|---|---|---|---|---|---|---|---|---|---|---|---|---|---|
| A | - | 10.998 | 9.404 | 8.608 | 7.47 | 8.53 | 8.746 | 8.888 | 9.064 | 8.835 | 9.852 | 11.09 | 7.685 | 8.236 |
| B | - | - | 1.022 | 2.077 | 2.786 | 3.053 | -1.092 | 2.704 | 2.648 | 2.539 | 2.232 | 2.083 | 2.568 | 0.101 |
| C | - | - | - | 2.603 | 3.11 | 3.334 | 1.261 | 3.072 | 2.007 | 2.943 | 2.715 | 2.616 | 2.944 | 1.578 |
| D | - | - | - | - | 1.449 | 2.058 | 1.974 | 1.125 | 3.437 | 0.743 | -1.422 | -4.116 | 0.77 | 1.779 |
| E | - | - | - | - | - | 0.585 | 2.772 | -1.208 | 3.796 | 0.043 | 1.267 | 1.536 | 0.036 | 2.661 |
| F | - | - | - | - | - | - | 3.066 | 0.955 | 3.969 | 1.346 | 1.973 | 2.155 | 1.278 | 2.988 |
| G | - | - | - | - | - | - | - | 2.581 | 2.71 | 2.435 | 2.058 | 1.896 | 2.438 | -1.036 |
| H | - | - | - | - | - | - | - | - | 3.777 | -0.499 | 0.95 | 1.307 | -1.081 | 2.585 |
| I | - | - | - | - | - | - | - | - | - | 3.486 | 3.361 | 3.297 | 3.479 | 2.754 |
| J | - | - | - | - | - | - | - | - | - | - | 0.429 | 0.83 | -4.32 | 2.39 |
| K | - | - | - | - | - | - | - | - | - | - | - | -0.91 | 0.597 | 1.947 |
| L | - | - | - | - | - | - | - | - | - | - | - | - | 0.867 | 1.741 |
| M | - | - | - | - | - | - | - | - | - | - | - | - | - | 2.402 |
| N | - | - | - | - | - | - | - | - | - | - | - | - | - | - |
