## Supplementary Data S2 for "Interlaboratory comparison of standardised metabolomics and lipidomics analyses in human and rodent blood using the MxP® Quant 500 kit": BABA.pdf

**Amount of relative differences between laboratories**

| Min | A | B | C | D | E | F | G | H | I | J | K | L | M | N |
|---|---|---|---|---|---|---|---|---|---|---|---|---|---|---|
| A | - | 4.819 | 5.253 | 6.272 | 4.154 | 4.915 | 3.504 | 4.989 | 11.177 | 5.263 | 4.915 | 6.535 | 4.954 | 2.469 |
| B | - | - | 4.473 | 8.706 | 7.486 | 7.513 | 7.37 | 5.227 | 6.023 | 4.7 | 6.125 | 8.668 | 5.279 | 6.693 |
| C | - | - | - | 7.93 | 6.975 | 6.543 | 7.103 | 4.297 | 7.537 | 2.314 | 4.287 | 8.222 | 4.539 | 5.62 |
| D | - | - | - | - | 5.538 | 6.085 | 5.562 | 5.536 | 6.234 | 5.663 | 5.05 | 2.667 | 5.503 | 4.77 |
| E | - | - | - | - | - | 4.232 | 3.325 | 4.49 | 9.254 | 5.267 | 3.726 | 6.588 | 4.385 | 1.3 |
| F | - | - | - | - | - | - | 4.911 | 3.873 | 13.7 | 5.12 | 3.479 | 6.953 | 3.66 | 3.43 |
| G | - | - | - | - | - | - | - | 5.069 | 8.712 | 5.631 | 4.729 | 6.173 | 5.007 | 3.742 |
| H | - | - | - | - | - | - | - | - | 11.492 | 4.24 | 2.726 | 7.763 | 1.81 | 4.724 |
| I | - | - | - | - | - | - | - | - | - | 13.217 | 12.546 | 15.966 | 13.756 | 13.921 |
| J | - | - | - | - | - | - | - | - | - | - | 3.477 | 8.385 | 4.789 | 5.83 |
| K | - | - | - | - | - | - | - | - | - | - | - | 7.717 | 2.308 | 4.679 |
| L | - | - | - | - | - | - | - | - | - | - | - | - | 5.54 | 4.7 |
| M | - | - | - | - | - | - | - | - | - | - | - | - | - | 4.569 |
| N | - | - | - | - | - | - | - | - | - | - | - | - | - | - |

| | |
|---|---|
| | Larger than 10 |
| | Between 8 and 9,99 |
| | Between 6 and 7,99 |
| | Between 4 and 5,99 |
| | Between 2 and 3,99 |
| | Between 0 and 1,99 |
| | NA (Calculation failed) |

| Median | A | B | C | D | E | F | G | H | I | J | K | L | M | N |
|---|---|---|---|---|---|---|---|---|---|---|---|---|---|---|
| A | - | 3.752 | 3.931 | 4.516 | 2.343 | 1.895 | 4.193 | 3.26 | 6.425 | 4.698 | 2.604 | 3.774 | 2.767 | -1.224 |
| B | - | - | 4.059 | 6.53 | 5.042 | 5.369 | 5.915 | 4.675 | 6.604 | 3.157 | 4.817 | 6.026 | 4.401 | 5.475 |
| C | - | - | - | 5.632 | 3.928 | 3.942 | 5.448 | 3.47 | 6.613 | 3.5 | 2.289 | 5.329 | 2.98 | 4.084 |
| D | - | - | - | - | 4.151 | 4.211 | 0.828 | 4.454 | 6.627 | 5.187 | 3.846 | 2.761 | 4.198 | 3.858 |
| E | - | - | - | - | - | -1.227 | 4.656 | 1.946 | 6.655 | 4.632 | -0.022 | 4.044 | 0.761 | 2.073 |
| F | - | - | - | - | - | - | 4.616 | 1.668 | 5.865 | 4.641 | 1.216 | 4.057 | 1.25 | 2.05 |
| G | - | - | - | - | - | - | - | 4.501 | 6.651 | 5.328 | 4.312 | 2.724 | 4.438 | 4.054 |
| H | - | - | - | - | - | - | - | - | 6.405 | 4.311 | 0.785 | 4.652 | 0.385 | 2.628 |
| I | - | - | - | - | - | - | - | - | - | 12.659 | 12.152 | 13.581 | 12.815 | 13 |
| J | - | - | - | - | - | - | - | - | - | - | 4.119 | 5.952 | 4.556 | 5.039 |
| K | - | - | - | - | - | - | - | - | - | - | - | 4.806 | 0.236 | 3.151 |
| L | - | - | - | - | - | - | - | - | - | - | - | - | 3.604 | 3.027 |
| M | - | - | - | - | - | - | - | - | - | - | - | - | - | 2.879 |
| N | - | - | - | - | - | - | - | - | - | - | - | - | - | - |

| Max | A | B | C | D | E | F | G | H | I | J | K | L | M | N |
|---|---|---|---|---|---|---|---|---|---|---|---|---|---|---|
| A | - | -0.459 | 3.185 | 3.189 | 1.11 | 1.606 | 4.731 | 3.53 | 5.261 | 3.484 | 3.042 | 0.497 | 2.899 | 1.609 |
| B | - | - | 3.432 | 3.349 | -0.897 | 2.385 | 4.954 | 3.827 | 6.614 | 3.498 | 1.782 | -7.077 | 3.417 | 2.1 |
| C | - | - | - | -1.161 | 3.336 | 2.182 | 4.128 | 1.542 | 6.585 | 4.235 | 1.931 | 3.189 | 0.226 | 2.317 |
| D | - | - | - | - | 3.486 | 2.563 | 3.949 | 0.494 | 6.634 | 4.29 | 1.689 | 3.215 | 1.282 | 2.535 |
| E | - | - | - | - | - | 2.417 | 4.88 | 3.796 | 6.858 | 3.099 | 3.238 | 0.364 | 3.267 | 2.432 |
| F | - | - | - | - | - | - | 4.522 | 2.956 | 7.81 | 3.737 | 1.549 | 1.934 | 1.877 | -1.407 |
| G | - | - | - | - | - | - | - | 3.612 | 6.802 | 4.862 | 3.992 | 4.435 | 3.955 | 4.233 |
| H | - | - | - | - | - | - | - | - | 4.648 | 4.392 | 2.74 | 3.524 | 2.051 | 2.927 |
| I | - | - | - | - | - | - | - | - | - | 11.478 | 11.853 | 11.215 | 11.687 | 11.646 |
| J | - | - | - | - | - | - | - | - | - | - | 4.398 | 3.309 | 4.398 | 4.046 |
| K | - | - | - | - | - | - | - | - | - | - | - | 2.775 | 0.069 | 1.333 |
| L | - | - | - | - | - | - | - | - | - | - | - | - | 3.02 | 2 |
| M | - | - | - | - | - | - | - | - | - | - | - | - | - | 1.926 |
| N | - | - | - | - | - | - | - | - | - | - | - | - | - | - |
