## Supplementary Data S2 for "Interlaboratory comparison of standardised metabolomics and lipidomics analyses in human and rodent blood using the MxP® Quant 500 kit": C8.pdf

**Amount of relative differences between laboratories**

| Min | A | B | C | D | E | F | G | H | I | J | K | L | M | N |
|---|---|---|---|---|---|---|---|---|---|---|---|---|---|---|
| A | - | NA | NA | NA | NA | NA | NA | NA | NA | NA | NA | NA | NA | NA |
| B | - | - | NA | NA | 3.888 | NA | 6.39 | NA | NA | NA | 6.28 | 6.01 | NA | NA |
| C | - | - | - | NA | NA | NA | NA | NA | NA | NA | NA | NA | NA | NA |
| D | - | - | - | - | NA | NA | NA | NA | NA | NA | NA | NA | NA | NA |
| E | - | - | - | - | - | NA | 6.701 | NA | NA | NA | 6.576 | 6.363 | NA | NA |
| F | - | - | - | - | - | - | NA | NA | NA | NA | NA | NA | NA | NA |
| G | - | - | - | - | - | - | - | NA | NA | NA | 2.332 | 2.856 | NA | NA |
| H | - | - | - | - | - | - | - | - | NA | NA | NA | NA | NA | NA |
| I | - | - | - | - | - | - | - | - | - | NA | NA | NA | NA | NA |
| J | - | - | - | - | - | - | - | - | - | - | NA | NA | NA | NA |
| K | - | - | - | - | - | - | - | - | - | - | - | 1.857 | NA | NA |
| L | - | - | - | - | - | - | - | - | - | - | - | - | NA | NA |
| M | - | - | - | - | - | - | - | - | - | - | - | - | - | NA |
| N | - | - | - | - | - | - | - | - | - | - | - | - | - | - |

| Median | A | B | C | D | E | F | G | H | I | J | K | L | M | N |
|---|---|---|---|---|---|---|---|---|---|---|---|---|---|---|
| A | - | NA | NA | NA | NA | NA | NA | NA | NA | NA | NA | NA | NA | NA |
| B | - | - | NA | NA | 1.98 | NA | 5.801 | NA | NA | NA | 5.798 | 5.211 | NA | NA |
| C | - | - | - | NA | NA | NA | NA | NA | NA | NA | NA | NA | NA | NA |
| D | - | - | - | - | NA | NA | NA | NA | NA | NA | NA | NA | NA | NA |
| E | - | - | - | - | - | NA | 5.987 | NA | NA | NA | 5.952 | 5.269 | NA | NA |
| F | - | - | - | - | - | - | NA | NA | NA | NA | NA | NA | NA | NA |
| G | - | - | - | - | - | - | - | NA | NA | NA | 2.066 | 0.018 | NA | NA |
| H | - | - | - | - | - | - | - | - | NA | NA | NA | NA | NA | NA |
| I | - | - | - | - | - | - | - | - | - | NA | NA | NA | NA | NA |
| J | - | - | - | - | - | - | - | - | - | - | NA | NA | NA | NA |
| K | - | - | - | - | - | - | - | - | - | - | - | 2.237 | NA | NA |
| L | - | - | - | - | - | - | - | - | - | - | - | - | NA | NA |
| M | - | - | - | - | - | - | - | - | - | - | - | - | - | NA |
| N | - | - | - | - | - | - | - | - | - | - | - | - | - | - |

| Max | A | B | C | D | E | F | G | H | I | J | K | L | M | N |
|---|---|---|---|---|---|---|---|---|---|---|---|---|---|---|
| A | - | NA | NA | NA | NA | NA | NA | NA | NA | NA | NA | NA | NA | NA |
| B | - | - | NA | NA | 4.279 | NA | 3.588 | NA | NA | NA | 3.906 | 3.21 | NA | NA |
| C | - | - | - | NA | NA | NA | NA | NA | NA | NA | NA | NA | NA | NA |
| D | - | - | - | - | NA | NA | NA | NA | NA | NA | NA | NA | NA | NA |
| E | - | - | - | - | - | NA | -2.295 | NA | NA | NA | -1.502 | 2.519 | NA | NA |
| F | - | - | - | - | - | - | NA | NA | NA | NA | NA | NA | NA | NA |
| G | - | - | - | - | - | - | - | NA | NA | NA | 1.394 | 2.405 | NA | NA |
| H | - | - | - | - | - | - | - | - | NA | NA | NA | NA | NA | NA |
| I | - | - | - | - | - | - | - | - | - | NA | NA | NA | NA | NA |
| J | - | - | - | - | - | - | - | - | - | - | NA | NA | NA | NA |
| K | - | - | - | - | - | - | - | - | - | - | - | 2.541 | NA | NA |
| L | - | - | - | - | - | - | - | - | - | - | - | - | NA | NA |
| M | - | - | - | - | - | - | - | - | - | - | - | - | - | NA |
| N | - | - | - | - | - | - | - | - | - | - | - | - | - | - |

| |
|---|
| Larger than 10 |
| Between 8 and 9,99 |
| Between 6 and 7,99 |
| Between 4 and 5,99 |
| Between 2 and 3,99 |
| Between 0 and 1,99 |
| NA (Calculation failed) |
