## Supplementary Data S2 for "Interlaboratory comparison of standardised metabolomics and lipidomics analyses in human and rodent blood using the MxP® Quant 500 kit": C14.1.pdf

**Amount of relative differences between laboratories**

| Min | A | B | C | D | E | F | G | H | I | J | K | L | M | N |
|---|---|---|---|---|---|---|---|---|---|---|---|---|---|---|
| A | - | NA | -0.39 | 3.85 | 4.892 | 3.962 | 3.66 | NA | -2.159 | NA | 4.391 | 3.867 | 5.976 | 2.363 |
| B | - | - | NA | NA | NA | NA | NA | NA | NA | NA | NA | NA | NA | NA |
| C | - | - | - | 3.771 | 5.508 | 3.845 | 5.186 | NA | 2.087 | NA | 5.786 | 5.669 | 6.977 | 2.389 |
| D | - | - | - | - | 3.809 | 1.082 | 2.976 | NA | 3.815 | NA | 0.82 | 3.301 | 6.391 | 1.55 |
| E | - | - | - | - | - | 4.539 | 5.404 | NA | 3.537 | NA | 5.133 | 4.886 | 4.357 | 3.572 |
| F | - | - | - | - | - | - | 4.649 | NA | 2.901 | NA | 5.123 | 3.214 | 6.674 | -4.276 |
| G | - | - | - | - | - | - | - | NA | 2.266 | NA | 2.74 | 3.489 | 5.511 | 3.171 |
| H | - | - | - | - | - | - | - | - | NA | NA | NA | NA | NA | NA |
| I | - | - | - | - | - | - | - | - | - | NA | 7.161 | 5.335 | 6.769 | 2.342 |
| J | - | - | - | - | - | - | - | - | - | - | NA | NA | NA | NA |
| K | - | - | - | - | - | - | - | - | - | - | - | 2.212 | 6.938 | 2.052 |
| L | - | - | - | - | - | - | - | - | - | - | - | - | 4.804 | 2.784 |
| M | - | - | - | - | - | - | - | - | - | - | - | - | - | 5.61 |
| N | - | - | - | - | - | - | - | - | - | - | - | - | - | - |

| | |
|---|---|
| | Larger than 10 |
| | Between 8 and 9,99 |
| | Between 6 and 7,99 |
| | Between 4 and 5,99 |
| | Between 2 and 3,99 |
| | Between 0 and 1,99 |
| | NA (Calculation failed) |

| Median | A | B | C | D | E | F | G | H | I | J | K | L | M | N |
|---|---|---|---|---|---|---|---|---|---|---|---|---|---|---|
| A | - | NA | 1.55 | 4.888 | 3.378 | 5.231 | 5.058 | NA | 3.321 | NA | 4.813 | 3.628 | 6.766 | 4.393 |
| B | - | - | NA | NA | NA | NA | NA | NA | NA | NA | NA | NA | NA | NA |
| C | - | - | - | 4.706 | 2.135 | 4.991 | 4.678 | NA | 3.037 | NA | 4.333 | 2.36 | 7.129 | 4.103 |
| D | - | - | - | - | 4.242 | 0.92 | 1.801 | NA | 4.175 | NA | 2.867 | 4.08 | 6.384 | 2.776 |
| E | - | - | - | - | - | 4.466 | 4.35 | NA | 1.969 | NA | 3.994 | 1.308 | 6.275 | 3.639 |
| F | - | - | - | - | - | - | 0.209 | NA | 3.783 | NA | 2.952 | 3.917 | 6.58 | 2.188 |
| G | - | - | - | - | - | - | - | NA | 3.601 | NA | 1.913 | 3.926 | 4.99 | 2.696 |
| H | - | - | - | - | - | - | - | - | NA | NA | NA | NA | NA | NA |
| I | - | - | - | - | - | - | - | - | - | NA | 1.845 | 1.759 | 6.881 | 3.309 |
| J | - | - | - | - | - | - | - | - | - | - | NA | NA | NA | NA |
| K | - | - | - | - | - | - | - | - | - | - | - | 3.565 | 1.992 | 1.353 |
| L | - | - | - | - | - | - | - | - | - | - | - | - | 5.81 | 3.403 |
| M | - | - | - | - | - | - | - | - | - | - | - | - | - | 5.64 |
| N | - | - | - | - | - | - | - | - | - | - | - | - | - | - |

| Max | A | B | C | D | E | F | G | H | I | J | K | L | M | N |
|---|---|---|---|---|---|---|---|---|---|---|---|---|---|---|
| A | - | NA | 2.66 | 5.854 | 4.324 | 5.835 | 5.344 | NA | 4.706 | NA | 5.157 | 4.088 | 7.937 | 5.424 |
| B | - | - | NA | NA | NA | NA | NA | NA | NA | NA | NA | NA | NA | NA |
| C | - | - | - | 5.621 | 4.034 | 5.564 | 5.149 | NA | 3.853 | NA | 4.989 | 3.885 | 7.505 | 5.073 |
| D | - | - | - | - | 4.329 | 0.758 | 2.405 | NA | 4.605 | NA | 3.026 | 4.427 | 6.363 | 3.617 |
| E | - | - | - | - | - | 4.399 | 4.078 | NA | 3.035 | NA | 3.698 | 1.132 | 7.946 | 3.727 |
| F | - | - | - | - | - | - | 1.412 | NA | 4.566 | NA | 1.834 | 4.271 | 6.259 | 3.43 |
| G | - | - | - | - | - | - | - | NA | 4.565 | NA | 1.73 | 3.989 | 8.03 | 1.581 |
| H | - | - | - | - | - | - | - | - | NA | NA | NA | NA | NA | NA |
| I | - | - | - | - | - | - | - | - | - | NA | 4.711 | 1.579 | 7.171 | 4.052 |
| J | - | - | - | - | - | - | - | - | - | - | NA | NA | NA | NA |
| K | - | - | - | - | - | - | - | - | - | - | - | 3.709 | 8.569 | -1.493 |
| L | - | - | - | - | - | - | - | - | - | - | - | - | 8.255 | 3.971 |
| M | - | - | - | - | - | - | - | - | - | - | - | - | - | 5.68 |
| N | - | - | - | - | - | - | - | - | - | - | - | - | - | - |
