## Supplementary Data S2 for "Interlaboratory comparison of standardised metabolomics and lipidomics analyses in human and rodent blood using the MxP® Quant 500 kit": C18.pdf

**Amount of relative differences between laboratories**

| Min | A | B | C | D | E | F | G | H | I | J | K | L | M | N |
|---|---|---|---|---|---|---|---|---|---|---|---|---|---|---|
| A | - | 5.481 | -0.83 | 3.577 | 5.761 | 5.074 | 4.921 | NA | 4.752 | 4.654 | 6.196 | 3.321 | 1.371 | 4.783 |
| B | - | - | 7.096 | 6.469 | 3.019 | 5.274 | 7.558 | NA | 2.819 | 5.396 | 5.572 | 7.263 | 5.667 | 5.595 |
| C | - | - | - | 3.642 | 5.55 | 5.104 | 4.912 | NA | 4.96 | 4.703 | 6.215 | 3.409 | 2.258 | 5.003 |
| D | - | - | - | - | 5.482 | 4.728 | 5.569 | NA | 4.696 | 4.148 | 5.858 | 4.727 | 2.748 | 4.527 |
| E | - | - | - | - | - | 5.069 | 7.453 | NA | 2.075 | 4.742 | 6.85 | 7.128 | 5.371 | 5.13 |
| F | - | - | - | - | - | - | 6.668 | NA | 3.538 | 2.65 | 5.91 | 6.247 | 4.903 | 0.554 |
| G | - | - | - | - | - | - | - | NA | 5.287 | 5.326 | 6.386 | 4.036 | 3.411 | 5.378 |
| H | - | - | - | - | - | - | - | - | NA | NA | NA | NA | NA | NA |
| I | - | - | - | - | - | - | - | - | - | 3.952 | 6.223 | 6.635 | 5.382 | 5.023 |
| J | - | - | - | - | - | - | - | - | - | - | 6.08 | 6.119 | 4.89 | -0.983 |
| K | - | - | - | - | - | - | - | - | - | - | - | 7.996 | 5.971 | 6.368 |
| L | - | - | - | - | - | - | - | - | - | - | - | - | 1.707 | 5.098 |
| M | - | - | - | - | - | - | - | - | - | - | - | - | - | 1.972 |
| N | - | - | - | - | - | - | - | - | - | - | - | - | - | - |

| | |
|---|---|
| | Larger than 10 |
| | Between 8 and 9,99 |
| | Between 6 and 7,99 |
| | Between 4 and 5,99 |
| | Between 2 and 3,99 |
| | Between 0 and 1,99 |
| | NA (Calculation failed) |

| Median | A | B | C | D | E | F | G | H | I | J | K | L | M | N |
|---|---|---|---|---|---|---|---|---|---|---|---|---|---|---|
| A | - | 5.266 | 2.256 | 2.283 | 5.203 | 4.477 | 4.378 | NA | 4.716 | 3.883 | 2.173 | 3.127 | 2.064 | 4.022 |
| B | - | - | 6.345 | 6.058 | 3.207 | 5.053 | 6.845 | NA | 2.857 | 5.18 | 4.725 | 6.497 | 5.679 | 5.399 |
| C | - | - | - | 3.122 | 5.197 | 4.683 | 3.997 | NA | 4.938 | 4.258 | 3.035 | 2.016 | 2.337 | 4.446 |
| D | - | - | - | - | 5.05 | 4.287 | 4.738 | NA | 4.676 | 3.669 | 3.462 | 3.736 | 2.87 | 3.898 |
| E | - | - | - | - | - | 4.632 | 6.648 | NA | 1.959 | 4.499 | 5.559 | 6.243 | 5.366 | 4.858 |
| F | - | - | - | - | - | - | 5.867 | NA | 3.556 | 2.251 | 2.342 | 5.348 | 4.859 | 1.842 |
| G | - | - | - | - | - | - | - | NA | 5.251 | 4.778 | 3.573 | 3.231 | 3.129 | 4.852 |
| H | - | - | - | - | - | - | - | - | NA | NA | NA | NA | NA | NA |
| I | - | - | - | - | - | - | - | - | - | 4.093 | 4.702 | 5.932 | 5.382 | 4.812 |
| J | - | - | - | - | - | - | - | - | - | - | 5.53 | 5.144 | 4.818 | 0.404 |
| K | - | - | - | - | - | - | - | - | - | - | - | 6.954 | 5.88 | 5.798 |
| L | - | - | - | - | - | - | - | - | - | - | - | - | 0.954 | 4.515 |
| M | - | - | - | - | - | - | - | - | - | - | - | - | - | 2.983 |
| N | - | - | - | - | - | - | - | - | - | - | - | - | - | - |

| Max | A | B | C | D | E | F | G | H | I | J | K | L | M | N |
|---|---|---|---|---|---|---|---|---|---|---|---|---|---|---|
| A | - | 4.176 | 3.164 | 2.512 | 3.121 | 1.508 | 3.435 | NA | 3.966 | -1.93 | 1.21 | 2.93 | 4.331 | 2.736 |
| B | - | - | 5.237 | 5.077 | 3.466 | 4.574 | 5.239 | NA | 3.355 | 4.799 | 5.079 | 5.102 | 5.794 | 4.894 |
| C | - | - | - | 1.61 | 4.341 | 3.444 | 0.151 | NA | 4.528 | 3.197 | 0.714 | 0.819 | 2.952 | -0.023 |
| D | - | - | - | - | 3.853 | 2.938 | 2.271 | NA | 4.326 | 2.463 | 2.668 | 0.764 | 3.727 | 0.689 |
| E | - | - | - | - | - | 3.304 | 4.437 | NA | 0.385 | 4.056 | 5.983 | 4.249 | 5.313 | 4.07 |
| F | - | - | - | - | - | - | 3.682 | NA | 3.81 | 1.366 | 3.528 | 3.266 | 4.325 | 2.978 |
| G | - | - | - | - | - | - | - | NA | 4.493 | 3.203 | 2.258 | 1.671 | 3.333 | 1.634 |
| H | - | - | - | - | - | - | - | - | NA | NA | NA | NA | NA | NA |
| I | - | - | - | - | - | - | - | - | - | 4.275 | 5.171 | 4.762 | 5.379 | 4.259 |
| J | - | - | - | - | - | - | - | - | - | - | 5.845 | 2.367 | 3.798 | 1.576 |
| K | - | - | - | - | - | - | - | - | - | - | - | 2.992 | 4.362 | -0.171 |
| L | - | - | - | - | - | - | - | - | - | - | - | - | 3.504 | -2.004 |
| M | - | - | - | - | - | - | - | - | - | - | - | - | - | 3.999 |
| N | - | - | - | - | - | - | - | - | - | - | - | - | - | - |
