## Supplementary Data S2 for "Interlaboratory comparison of standardised metabolomics and lipidomics analyses in human and rodent blood using the MxP® Quant 500 kit": CE(14.0).pdf

**Amount of relative differences between laboratories**

| Min | A | B | C | D | E | F | G | H | I | J | K | L | M | N |
|---|---|---|---|---|---|---|---|---|---|---|---|---|---|---|
| A | - | 4.445 | 2.771 | 4.757 | 7.233 | 5.822 | 6.916 | 3.11 | 5.496 | 6.229 | 6.364 | 6.837 | 1.698 | 3.498 |
| B | - | - | 2.993 | 2.888 | 6.915 | 6.515 | 6.731 | 3.898 | 6.257 | 5.845 | 6.106 | 6.716 | 4.871 | 5.345 |
| C | - | - | - | 6.878 | 8.271 | 3.947 | 8.599 | 6.855 | 6.172 | 8.674 | 8.457 | 8.251 | 7.053 | 5.556 |
| D | - | - | - | - | 7.106 | 6.592 | 6.483 | 4.497 | 6.054 | 4.945 | 5.532 | 6.533 | 5.373 | 5.322 |
| E | - | - | - | - | - | 7.811 | 4.926 | 6.981 | 6.972 | 7.152 | 6.596 | -0.325 | 7.392 | 6.989 |
| F | - | - | - | - | - | - | 7.298 | 5.611 | 3.082 | 7.056 | 7.047 | 7.192 | 5.493 | 4.862 |
| G | - | - | - | - | - | - | - | 6.714 | 7.005 | 6.519 | 5.624 | 4.435 | 7.124 | 6.866 |
| H | - | - | - | - | - | - | - | - | 5.842 | 6.097 | 6.313 | 6.766 | 3.6 | 4.52 |
| I | - | - | - | - | - | - | - | - | - | 7.436 | 7.303 | 7.373 | 6.171 | 5.635 |
| J | - | - | - | - | - | - | - | - | - | - | 4.474 | 6.104 | 6.278 | 6.088 |
| K | - | - | - | - | - | - | - | - | - | - | - | 5.901 | 6.79 | 6.54 |
| L | - | - | - | - | - | - | - | - | - | - | - | - | 7.759 | 7.493 |
| M | - | - | - | - | - | - | - | - | - | - | - | - | - | 3.994 |
| N | - | - | - | - | - | - | - | - | - | - | - | - | - | - |

| Median | A | B | C | D | E | F | G | H | I | J | K | L | M | N |
|---|---|---|---|---|---|---|---|---|---|---|---|---|---|---|
| A | - | 3.575 | 4.146 | 3.502 | 7.349 | 1.012 | 5.566 | 2.412 | 3.788 | -1.008 | 1.889 | 3.24 | 4.024 | 4.209 |
| B | - | - | 2.703 | 4.725 | 7.636 | 3.982 | 6.066 | 2.901 | 1.109 | 3.686 | 3.228 | 4.597 | 2.295 | 2.906 |
| C | - | - | - | 5.076 | 7.581 | 4.618 | 6.157 | 3.866 | 2.634 | 4.263 | 3.551 | 4.677 | -0.176 | -0.448 |
| D | - | - | - | - | 7.103 | 3.126 | 5.004 | 3.926 | 4.503 | 3.468 | 3.753 | 0.716 | 4.638 | 4.75 |
| E | - | - | - | - | - | 6.021 | 5.588 | 6.066 | 6.116 | 6.053 | 6.028 | 5.917 | 6.136 | 6.153 |
| F | - | - | - | - | - | - | 5.477 | 2.866 | 3.976 | 1.092 | 2.538 | 2.838 | 4.178 | 4.34 |
| G | - | - | - | - | - | - | - | 5.159 | 5.36 | 5.057 | 5.076 | 4.639 | 5.419 | 5.472 |
| H | - | - | - | - | - | - | - | - | 3.165 | 2.338 | 0.804 | 3.957 | 3.528 | 3.8 |
| I | - | - | - | - | - | - | - | - | - | 4.031 | 3.532 | 4.75 | 1.483 | 2.408 |
| J | - | - | - | - | - | - | - | - | - | - | 1.323 | 3.399 | 3.944 | 4.138 |
| K | - | - | - | - | - | - | - | - | - | - | - | 3.75 | 3.735 | 3.96 |
| L | - | - | - | - | - | - | - | - | - | - | - | - | 4.575 | 4.695 |
| M | - | - | - | - | - | - | - | - | - | - | - | - | - | 1.422 |
| N | - | - | - | - | - | - | - | - | - | - | - | - | - | - |

| Max | A | B | C | D | E | F | G | H | I | J | K | L | M | N |
|---|---|---|---|---|---|---|---|---|---|---|---|---|---|---|
| A | - | 3.504 | 4.473 | 3.901 | 8.675 | -1.079 | 6.068 | 2.358 | 4.141 | 1.243 | 0.393 | 4.53 | 4.1 | 4.322 |
| B | - | - | 3.644 | 4.889 | 8.812 | 3.626 | 6.448 | 2.815 | 2.859 | 3.961 | 3.844 | 5.266 | 2.69 | 3.315 |
| C | - | - | - | 5.584 | 9.189 | 4.721 | 6.968 | 4.472 | 0.949 | 5.155 | 5.124 | 5.955 | 2.793 | 0.7 |
| D | - | - | - | - | 8.464 | 3.758 | 5.534 | 4.121 | 4.831 | 3.365 | 3.51 | 3.069 | 4.802 | 4.922 |
| E | - | - | - | - | - | 6.288 | 5.841 | 6.29 | 6.357 | 6.217 | 6.24 | 6.066 | 6.341 | 6.344 |
| F | - | - | - | - | - | - | 6.091 | 2.281 | 4.113 | 1.537 | 1.173 | 4.59 | 4.077 | 4.308 |
| G | - | - | - | - | - | - | - | 5.457 | 5.702 | 5.252 | 5.286 | 4.66 | 5.673 | 5.717 |
| H | - | - | - | - | - | - | - | - | 3.731 | 2.979 | 2.793 | 4.87 | 3.656 | 3.982 |
| I | - | - | - | - | - | - | - | - | - | 4.562 | 4.484 | 5.59 | -1.568 | 1.608 |
| J | - | - | - | - | - | - | - | - | - | - | 0.315 | 4.271 | 4.292 | 4.457 |
| K | - | - | - | - | - | - | - | - | - | - | - | 4.429 | 4.242 | 4.399 |
| L | - | - | - | - | - | - | - | - | - | - | - | - | 5.117 | 5.185 |
| M | - | - | - | - | - | - | - | - | - | - | - | - | - | 1.859 |
| N | - | - | - | - | - | - | - | - | - | - | - | - | - | - |

| | |
|---|---|
| | Larger than 10 |
| | Between 8 and 9,99 |
| | Between 6 and 7,99 |
| | Between 4 and 5,99 |
| | Between 2 and 3,99 |
| | Between 0 and 1,99 |
| | NA (Calculation failed) |
