## Supplementary Data S2 for "Interlaboratory comparison of standardised metabolomics and lipidomics analyses in human and rodent blood using the MxP® Quant 500 kit": CE(14.1).pdf

**Amount of relative differences between laboratories**

| Min | A | B | C | D | E | F | G | H | I | J | K | L | M | N |
|---|---|---|---|---|---|---|---|---|---|---|---|---|---|---|
| A | - | 6.582 | 4.122 | 7.169 | 9.285 | 5.855 | 9.402 | 6.876 | 7.45 | 7.344 | 5.546 | 7.595 | 6.293 | 6.95 |
| B | - | - | 4.014 | 5.335 | 9.154 | 1.756 | 9.019 | 4.702 | 3.472 | 3.021 | 7.164 | 7.653 | 4.546 | 5.013 |
| C | - | - | - | 6.72 | 6.14 | 4.605 | 8.926 | -0.852 | 5.524 | 4.882 | 7.714 | 4.361 | 5.595 | 6.295 |
| D | - | - | - | - | 9.574 | 4.356 | 8.596 | 5.469 | 5.176 | 4.338 | 6.181 | 8.084 | 5.122 | 0.486 |
| E | - | - | - | - | - | 5.256 | 6.562 | 4.369 | 6.838 | 5.873 | 7.356 | 4.587 | 5.457 | 6.696 |
| F | - | - | - | - | - | - | 8.7 | 3.384 | 1.137 | 4.139 | 7.632 | 5.031 | 4.017 | 5.852 |
| G | - | - | - | - | - | - | - | 6.07 | 7.569 | 7.012 | 8.13 | 4.678 | 0.958 | 7.514 |
| H | - | - | - | - | - | - | - | - | 5.825 | 5.451 | 7.351 | 6.418 | 4.528 | 6.196 |
| I | - | - | - | - | - | - | - | - | - | 0.947 | 6.587 | 7.57 | 5.104 | 4.379 |
| J | - | - | - | - | - | - | - | - | - | - | 7.022 | 6.368 | 4.717 | 4.926 |
| K | - | - | - | - | - | - | - | - | - | - | - | 7.643 | 6.109 | 6.048 |
| L | - | - | - | - | - | - | - | - | - | - | - | - | 4.382 | 7.577 |
| M | - | - | - | - | - | - | - | - | - | - | - | - | - | 6.676 |
| N | - | - | - | - | - | - | - | - | - | - | - | - | - | - |

| Median | A | B | C | D | E | F | G | H | I | J | K | L | M | N |
|---|---|---|---|---|---|---|---|---|---|---|---|---|---|---|
| A | - | 2.782 | 4.773 | 4.637 | 5.901 | -1.33 | 6.213 | 5.23 | 5.782 | 4.057 | 5.243 | 4.387 | 2.545 | 2.808 |
| B | - | - | 5.108 | 5.18 | 6.349 | 3.48 | 6.437 | 5.692 | 5.99 | 4.681 | 5.681 | 4.818 | 1.41 | -0.132 |
| C | - | - | - | 2.279 | 6.741 | 4.387 | 3.092 | 4.046 | 4.414 | 3.654 | 4.403 | 5.279 | 4.883 | 4.508 |
| D | - | - | - | - | 3.293 | 4.028 | 5.219 | 3.522 | 4.331 | 2.761 | 3.454 | 4.333 | 4.517 | 4.691 |
| E | - | - | - | - | - | 5.985 | 5.043 | 5.353 | 5.447 | 5.843 | 5.396 | 5.263 | 6.07 | 5.962 |
| F | - | - | - | - | - | - | 6.016 | 5.163 | 5.4 | 3.516 | 5.288 | 5.911 | 2.995 | 3.319 |
| G | - | - | - | - | - | - | - | 3.714 | 4.03 | 5.156 | 3.861 | 2.775 | 5.519 | 5.476 |
| H | - | - | - | - | - | - | - | - | 1.892 | 4.271 | -0.647 | 2.706 | 5.075 | 5.105 |
| I | - | - | - | - | - | - | - | - | - | 4.462 | 2.583 | 4.672 | 5.142 | 5.232 |
| J | - | - | - | - | - | - | - | - | - | - | 4.422 | 4.999 | 4.121 | 4.3 |
| K | - | - | - | - | - | - | - | - | - | - | - | 4.215 | 5.013 | 5.042 |
| L | - | - | - | - | - | - | - | - | - | - | - | - | 5.461 | 5.211 |
| M | - | - | - | - | - | - | - | - | - | - | - | - | - | 0.855 |
| N | - | - | - | - | - | - | - | - | - | - | - | - | - | - |

| Max | A | B | C | D | E | F | G | H | I | J | K | L | M | N |
|---|---|---|---|---|---|---|---|---|---|---|---|---|---|---|
| A | - | 3.738 | 6.24 | 5.709 | 9.669 | 5.218 | 8.418 | 6.831 | 6.626 | 5.631 | 5.734 | 7.805 | 5.019 | 3.178 |
| B | - | - | 5.762 | 5.153 | 9.644 | 4.347 | 8.242 | 6.492 | 6.281 | 4.994 | 5.113 | 7.938 | 4.033 | 1.768 |
| C | - | - | - | 3.522 | 8.243 | 4.229 | 7.532 | 4.607 | 4.055 | 3.246 | 3.714 | 5.81 | 4.293 | 5.198 |
| D | - | - | - | - | 9.681 | 3.773 | 7.559 | 5.284 | 4.946 | 2.107 | -2.216 | 7.765 | 4.054 | 4.887 |
| E | - | - | - | - | - | 6.308 | 3.464 | 6.153 | 6.203 | 6.299 | 6.312 | 5.7 | 6.311 | 6.413 |
| F | - | - | - | - | - | - | 7.884 | 5.873 | 5.663 | 3.366 | 3.348 | 7.209 | 1.864 | 4.271 |
| G | - | - | - | - | - | - | - | 5.719 | 5.796 | 6.043 | 6.088 | 5.23 | 6.151 | 6.244 |
| H | - | - | - | - | - | - | - | - | 2.67 | 4.949 | 4.925 | 6.091 | 5.297 | 5.621 |
| I | - | - | - | - | - | - | - | - | - | 4.725 | 4.616 | 7.024 | 5.162 | 5.511 |
| J | - | - | - | - | - | - | - | - | - | - | 0.795 | 7.133 | 3.667 | 4.733 |
| K | - | - | - | - | - | - | - | - | - | - | - | 7.256 | 3.695 | 4.827 |
| L | - | - | - | - | - | - | - | - | - | - | - | - | 5.814 | 6.096 |
| M | - | - | - | - | - | - | - | - | - | - | - | - | - | 4.012 |
| N | - | - | - | - | - | - | - | - | - | - | - | - | - | - |

| | |
|---|---|
| | Larger than 10 |
| | Between 8 and 9,99 |
| | Between 6 and 7,99 |
| | Between 4 and 5,99 |
| | Between 2 and 3,99 |
| | Between 0 and 1,99 |
| | NA (Calculation failed) |
