## Supplementary Data S2 for "Interlaboratory comparison of standardised metabolomics and lipidomics analyses in human and rodent blood using the MxP® Quant 500 kit": CE(16.0).pdf

**Amount of relative differences between laboratories**

| Min | A | B | C | D | E | F | G | H | I | J | K | L | M | N |
|---|---|---|---|---|---|---|---|---|---|---|---|---|---|---|
| A | - | 4.634 | 3.758 | 8.439 | 9.546 | 4.727 | 5.814 | 0.751 | 1 | 3.411 | 4.799 | 4.703 | 3.894 | 3.253 |
| B | - | - | 5.539 | 8.51 | 9.604 | 5.966 | 3.688 | 4.932 | 4.422 | 4.197 | 1.304 | 2.572 | 3.923 | 3.954 |
| C | - | - | - | 8.322 | 9.486 | 3.649 | 5.973 | 3.192 | 2.046 | 4.358 | 5.16 | 4.852 | 4.509 | 4.164 |
| D | - | - | - | - | 8.533 | 9.052 | 7.782 | 8.751 | 8.176 | 8.906 | 8.567 | 8.496 | 8.725 | 8.298 |
| E | - | - | - | - | - | 6.886 | 6 | 6.239 | 5.276 | 6.412 | 5.818 | 5.176 | 6.416 | 5.779 |
| F | - | - | - | - | - | - | 6.184 | 4.36 | 4.412 | 5.219 | 5.677 | 5.857 | 5.255 | 4.855 |
| G | - | - | - | - | - | - | - | 5.56 | 1.301 | 6.024 | 2.864 | 2.477 | 4.906 | 2.643 |
| H | - | - | - | - | - | - | - | - | 1.438 | 3.497 | 4.857 | 4.652 | 3.923 | 3.811 |
| I | - | - | - | - | - | - | - | - | - | 3.817 | 5.198 | 4.996 | 4.219 | 3.322 |
| J | - | - | - | - | - | - | - | - | - | - | 4.27 | 4.264 | 1.757 | 3.801 |
| K | - | - | - | - | - | - | - | - | - | - | - | 2.018 | 4.351 | 3.524 |
| L | - | - | - | - | - | - | - | - | - | - | - | - | 4.917 | 2.357 |
| M | - | - | - | - | - | - | - | - | - | - | - | - | - | 1.471 |
| N | - | - | - | - | - | - | - | - | - | - | - | - | - | - |

| Median | A | B | C | D | E | F | G | H | I | J | K | L | M | N |
|---|---|---|---|---|---|---|---|---|---|---|---|---|---|---|
| A | - | 2.669 | 2.49 | 4.441 | 8.157 | 1.797 | 5.85 | 0.064 | 4.001 | 3.585 | 1.334 | 1.008 | -0.809 | 3.174 |
| B | - | - | -0.454 | 4.884 | 8.273 | 3.395 | 6.141 | 2.506 | 3.358 | 4.283 | 2.011 | 2.226 | 2.604 | 1.353 |
| C | - | - | - | 4.605 | 8.206 | 3.256 | 6.044 | 2.254 | 3.491 | 4.218 | 1.654 | 1.832 | 2.375 | 1.967 |
| D | - | - | - | - | 7.949 | 0.61 | 6.239 | 2.766 | 4.245 | 0.088 | 3.292 | 3.414 | 2.881 | 2.998 |
| E | - | - | - | - | - | 6.137 | 5.466 | 6.166 | 6.243 | 6.115 | 6.183 | 6.179 | 6.177 | 6.174 |
| F | - | - | - | - | - | - | 5.678 | 2.114 | 4.232 | 3.097 | 2.482 | 2.219 | 1.921 | 3.619 |
| G | - | - | - | - | - | - | - | 5.357 | 5.617 | 5.111 | 5.378 | 5.375 | 5.344 | 5.463 |
| H | - | - | - | - | - | - | - | - | 3.92 | 3.718 | 0.59 | 0.078 | -1.189 | 3.01 |
| I | - | - | - | - | - | - | - | - | - | 5.065 | 4.023 | 4.077 | 4.203 | 3.011 |
| J | - | - | - | - | - | - | - | - | - | - | 3.685 | 3.635 | 3.483 | 4.19 |
| K | - | - | - | - | - | - | - | - | - | - | - | -1.699 | 0.918 | 2.62 |
| L | - | - | - | - | - | - | - | - | - | - | - | - | 0.18 | 2.697 |
| M | - | - | - | - | - | - | - | - | - | - | - | - | - | 3.037 |
| N | - | - | - | - | - | - | - | - | - | - | - | - | - | - |

| Max | A | B | C | D | E | F | G | H | I | J | K | L | M | N |
|---|---|---|---|---|---|---|---|---|---|---|---|---|---|---|
| A | - | 2.391 | 2.637 | 8.713 | 8.509 | 1.242 | 6.659 | 0.242 | 4.104 | 3.69 | 0.326 | -3.365 | -2.446 | 3.166 |
| B | - | - | -0.509 | 8.732 | 8.452 | 3.003 | 6.677 | 2.147 | 3.695 | 4.286 | 2.123 | 2.488 | 2.485 | 2.099 |
| C | - | - | - | 8.558 | 8.22 | 3.234 | 6.845 | 2.451 | 3.567 | 4.334 | 2.349 | 2.679 | 2.736 | 1.36 |
| D | - | - | - | - | 6.199 | 4.765 | 4.36 | 4.85 | 5.499 | 4.176 | 4.872 | 5.434 | 4.91 | 5.282 |
| E | - | - | - | - | - | 6.313 | 6.416 | 6.308 | 6.417 | 6.238 | 6.306 | 6.357 | 6.334 | 6.399 |
| F | - | - | - | - | - | - | 6.62 | 1.812 | 4.215 | 3.377 | 1.723 | -5.79 | 0.981 | 3.419 |
| G | - | - | - | - | - | - | - | 5.505 | 5.741 | 5.268 | 5.461 | 5.512 | 5.471 | 5.577 |
| H | - | - | - | - | - | - | - | - | 4.017 | 3.821 | -2.949 | 0.08 | 0.336 | 2.904 |
| I | - | - | - | - | - | - | - | - | - | 5.144 | 4.241 | 4.403 | 4.348 | 3.291 |
| J | - | - | - | - | - | - | - | - | - | - | 3.648 | 3.565 | 3.535 | 4.223 |
| K | - | - | - | - | - | - | - | - | - | - | - | -5.653 | -0.131 | 2.971 |
| L | - | - | - | - | - | - | - | - | - | - | - | - | -1.321 | 2.925 |
| M | - | - | - | - | - | - | - | - | - | - | - | - | - | 3.125 |
| N | - | - | - | - | - | - | - | - | - | - | - | - | - | - |

| Colour | Legend |
|---|---|
| (red) | Larger than 10 |
| (orange) | Between 8 and 9,99 |
| (light orange) | Between 6 and 7,99 |
| (light green) | Between 4 and 5,99 |
| (green) | Between 2 and 3,99 |
| (dark green) | Between 0 and 1,99 |
| (grey) | NA (Calculation failed) |
