## Supplementary Data S2 for "Interlaboratory comparison of standardised metabolomics and lipidomics analyses in human and rodent blood using the MxP® Quant 500 kit": CE(16.1).pdf

**Amount of relative differences between laboratories**

| Min | A | B | C | D | E | F | G | H | I | J | K | L | M | N |
|---|---|---|---|---|---|---|---|---|---|---|---|---|---|---|
| A | - | 7.355 | 7.359 | 6.806 | 7.037 | 7.331 | 7.313 | 6.962 | 5.92 | 7.931 | 6.982 | 7.454 | 7.133 | 6.605 |
| B | - | - | 6.389 | 4.371 | 6.289 | 3.352 | 5.744 | 5.819 | 6.474 | 5.768 | 5.085 | 5.844 | 4.807 | 5.54 |
| C | - | - | - | 4.676 | 6.623 | 5.545 | 6.459 | 2.649 | 4.982 | 6.784 | 4.414 | 6.581 | 4.9 | 2.548 |
| D | - | - | - | - | 6.42 | 2.161 | 5.932 | 4.89 | 5.705 | 5.928 | 3.483 | 6.032 | 2.642 | 4.091 |
| E | - | - | - | - | - | 7.224 | 4.865 | 7.414 | 7.313 | 6.414 | 7.066 | 5.625 | 7.099 | 6.806 |
| F | - | - | - | - | - | - | 6.015 | 5.485 | 6.215 | 6.002 | 4.524 | 6.209 | 4.074 | 5.082 |
| G | - | - | - | - | - | - | - | 6.991 | 7.077 | 5.291 | 6.568 | 3.63 | 6.569 | 6.545 |
| H | - | - | - | - | - | - | - | - | 5.254 | 6.773 | 3.973 | 6.605 | 4.598 | 0.416 |
| I | - | - | - | - | - | - | - | - | - | 7.762 | 6.218 | 7.333 | 6.501 | 5.427 |
| J | - | - | - | - | - | - | - | - | - | - | 6.034 | 3.833 | 5.959 | 6.081 |
| K | - | - | - | - | - | - | - | - | - | - | - | 6.361 | 2.95 | 3.943 |
| L | - | - | - | - | - | - | - | - | - | - | - | - | 6.41 | 6.373 |
| M | - | - | - | - | - | - | - | - | - | - | - | - | - | 4.53 |
| N | - | - | - | - | - | - | - | - | - | - | - | - | - | - |

| Median | A | B | C | D | E | F | G | H | I | J | K | L | M | N |
|---|---|---|---|---|---|---|---|---|---|---|---|---|---|---|
| A | - | 3.417 | 1.915 | 3.276 | 2.729 | 3.188 | 0.252 | 0.192 | 4.189 | -0.907 | 1.463 | 1.552 | 3.247 | 4.09 |
| B | - | - | 3.912 | 4.756 | 4.35 | 1.458 | 3.628 | 3.541 | 2.942 | 3.631 | 3.058 | 3.359 | 1.207 | 2.399 |
| C | - | - | - | 3.195 | 2.031 | 3.481 | 1.596 | 1.783 | 4.333 | 1.338 | 2.577 | 2.267 | 3.528 | 4.194 |
| D | - | - | - | - | 2.427 | 4.262 | 3.54 | 3.61 | 4.777 | 3.542 | 3.894 | 3.733 | 4.289 | 4.673 |
| E | - | - | - | - | - | 3.887 | 2.667 | 2.923 | 4.562 | 2.965 | 3.518 | 3.015 | 3.93 | 4.392 |
| F | - | - | - | - | - | - | 3.212 | 3.115 | 3.345 | 3.238 | 2.46 | 2.848 | -1.225 | 2.977 |
| G | - | - | - | - | - | - | - | 0.087 | 4.186 | -0.082 | 2.232 | 0.889 | 3.216 | 3.965 |
| H | - | - | - | - | - | - | - | - | 4.113 | -0.099 | 1.402 | 0.531 | 3.068 | 3.941 |
| I | - | - | - | - | - | - | - | - | - | 4.498 | 4.203 | 4.234 | 3.423 | 0.909 |
| J | - | - | - | - | - | - | - | - | - | - | 1.895 | 0.361 | 3.113 | 3.925 |
| K | - | - | - | - | - | - | - | - | - | - | - | 0.533 | 2.53 | 3.687 |
| L | - | - | - | - | - | - | - | - | - | - | - | - | 2.908 | 3.813 |
| M | - | - | - | - | - | - | - | - | - | - | - | - | - | 2.915 |
| N | - | - | - | - | - | - | - | - | - | - | - | - | - | - |

| Max | A | B | C | D | E | F | G | H | I | J | K | L | M | N |
|---|---|---|---|---|---|---|---|---|---|---|---|---|---|---|
| A | - | 1.737 | 3.873 | 4.998 | 5.085 | 2.325 | 4.694 | 3.502 | 3.806 | 4.379 | 3.307 | 4.336 | 1.675 | 2.943 |
| B | - | - | 3.406 | 4.816 | 5.319 | 1.037 | 4.565 | 2.831 | 4.12 | 4.165 | 2.513 | 4.274 | -2.476 | 3.497 |
| C | - | - | - | 3.955 | 4.61 | 2.975 | 3.53 | 1.657 | 4.664 | 2.678 | 2.136 | 2.885 | 3.287 | 4.304 |
| D | - | - | - | - | 3.394 | 4.362 | 1.658 | 4.039 | 5.146 | 2.919 | 4.132 | 2.657 | 4.462 | 4.928 |
| E | - | - | - | - | - | 4.91 | 3.637 | 4.676 | 5.391 | 3.957 | 4.719 | 3.981 | 4.907 | 5.188 |
| F | - | - | - | - | - | - | 4.456 | 2.326 | 4.221 | 3.957 | 1.892 | 4.177 | 1.072 | 3.669 |
| G | - | - | - | - | - | - | - | 3.809 | 5.07 | 2.094 | 3.896 | 1.911 | 4.278 | 4.822 |
| H | - | - | - | - | - | - | - | - | 4.516 | 3.333 | 0.384 | 3.513 | 2.776 | 4.113 |
| I | - | - | - | - | - | - | - | - | - | 5.408 | 4.803 | 5.425 | 4.361 | 2.827 |
| J | - | - | - | - | - | - | - | - | - | - | 3.434 | -1.134 | 3.956 | 4.634 |
| K | - | - | - | - | - | - | - | - | - | - | - | 3.663 | 2.482 | 4.015 |
| L | - | - | - | - | - | - | - | - | - | - | - | - | 4.042 | 4.662 |
| M | - | - | - | - | - | - | - | - | - | - | - | - | - | 3.473 |
| N | - | - | - | - | - | - | - | - | - | - | - | - | - | - |

| | |
|---|---|
| | Larger than 10 |
| | Between 8 and 9,99 |
| | Between 6 and 7,99 |
| | Between 4 and 5,99 |
| | Between 2 and 3,99 |
| | Between 0 and 1,99 |
| | NA (Calculation failed) |
