## Supplementary Data S2 for "Interlaboratory comparison of standardised metabolomics and lipidomics analyses in human and rodent blood using the MxP® Quant 500 kit": CE(18.0).pdf

**Amount of relative differences between laboratories**

| Min | A | B | C | D | E | F | G | H | I | J | K | L | M | N |
|---|---|---|---|---|---|---|---|---|---|---|---|---|---|---|
| A | - | 3.721 | 3.485 | 5.777 | 5.734 | 4.54 | 5.283 | 4.206 | 2.597 | 1.591 | 5.138 | 5.625 | 1.393 | 1.957 |
| B | - | - | 4.784 | 5.609 | 5.803 | 5.345 | 4.836 | 5.2 | 4.364 | 3.733 | 4.73 | 5.428 | 4.173 | 3.682 |
| C | - | - | - | 5.942 | 5.609 | 3.251 | 5.428 | 2.496 | 0.582 | 3.413 | 5.389 | 5.671 | 3.05 | 3.903 |
| D | - | - | - | - | 4.925 | 6.503 | 1.964 | 6.421 | 5.563 | 6.268 | -1.682 | 3.771 | 5.971 | 5.4 |
| E | - | - | - | - | - | 7.636 | 5.226 | 7.373 | 6.663 | 7.187 | 5.099 | 4.191 | 7.164 | 6.292 |
| F | - | - | - | - | - | - | 6.044 | 1.854 | 3.961 | 5.059 | 5.765 | 6.392 | 4.034 | 4.652 |
| G | - | - | - | - | - | - | - | 6.314 | 5.484 | 6.117 | 1.482 | 3.317 | 5.837 | 5.249 |
| H | - | - | - | - | - | - | - | - | 3.306 | 4.077 | 5.536 | 5.918 | 3.789 | 4.289 |
| I | - | - | - | - | - | - | - | - | - | 4.317 | 5.652 | 5.811 | 2.928 | 4.595 |
| J | - | - | - | - | - | - | - | - | - | - | 5.264 | 5.587 | 1.871 | 4.06 |
| K | - | - | - | - | - | - | - | - | - | - | - | 2.962 | 5.82 | 5.461 |
| L | - | - | - | - | - | - | - | - | - | - | - | - | 6.588 | 5.516 |
| M | - | - | - | - | - | - | - | - | - | - | - | - | - | 2.719 |
| N | - | - | - | - | - | - | - | - | - | - | - | - | - | - |

| | |
|---|---|
| | Larger than 10 |
| | Between 8 and 9,99 |
| | Between 6 and 7,99 |
| | Between 4 and 5,99 |
| | Between 2 and 3,99 |
| | Between 0 and 1,99 |
| | NA (Calculation failed) |

| Median | A | B | C | D | E | F | G | H | I | J | K | L | M | N |
|---|---|---|---|---|---|---|---|---|---|---|---|---|---|---|
| A | - | 3.409 | 2.633 | 4.404 | 3.633 | 0.957 | 3.279 | 2.742 | 4.323 | 3.155 | 1.544 | 4.095 | 3.344 | 4.018 |
| B | - | - | 2.298 | 5.153 | 4.714 | 3.81 | 4.501 | 4.274 | 3.396 | 4.454 | 3.107 | 2.853 | -0.968 | 2.648 |
| C | - | - | - | 4.867 | 4.279 | 3.134 | 4.085 | 3.768 | 3.889 | 4.017 | 1.813 | 3.562 | 2.109 | 3.433 |
| D | - | - | - | - | 2.761 | 3.957 | 3.256 | 3.426 | 5.055 | 3.303 | 4.27 | 4.926 | 4.636 | 4.891 |
| E | - | - | - | - | - | 3.302 | 1.417 | 1.838 | 4.806 | 1.62 | 3.695 | 4.618 | 4.245 | 4.566 |
| F | - | - | - | - | - | - | 3.051 | 2.154 | 4.432 | 2.832 | 2.211 | 4.222 | 3.59 | 4.169 |
| G | - | - | - | - | - | - | - | 0.739 | 4.723 | -1.216 | 3.452 | 4.553 | 4.106 | 4.497 |
| H | - | - | - | - | - | - | - | - | 4.643 | 1.164 | 3.154 | 4.479 | 3.98 | 4.425 |
| I | - | - | - | - | - | - | - | - | - | 5.192 | 4.441 | 1.762 | 3.608 | 2.206 |
| J | - | - | - | - | - | - | - | - | - | - | 3.435 | 4.575 | 4.13 | 4.52 |
| K | - | - | - | - | - | - | - | - | - | - | - | 3.827 | 2.826 | 3.723 |
| L | - | - | - | - | - | - | - | - | - | - | - | - | 3.012 | 0.307 |
| M | - | - | - | - | - | - | - | - | - | - | - | - | - | 2.759 |
| N | - | - | - | - | - | - | - | - | - | - | - | - | - | - |

| Max | A | B | C | D | E | F | G | H | I | J | K | L | M | N |
|---|---|---|---|---|---|---|---|---|---|---|---|---|---|---|
| A | - | 3.379 | 2.863 | 4.886 | 5.224 | -1.026 | 3.951 | 2.543 | 4.491 | 3.212 | -1.015 | 3.566 | 3.477 | 4.094 |
| B | - | - | 1.816 | 5.489 | 5.845 | 3.611 | 4.843 | 4.176 | 3.752 | 4.488 | 3.624 | 0.479 | -0.258 | 2.928 |
| C | - | - | - | 5.293 | 5.504 | 3.125 | 4.589 | 3.829 | 4.032 | 4.156 | 3.055 | 2.444 | 2.003 | 3.393 |
| D | - | - | - | - | 1.14 | 4.508 | 3.478 | 4.156 | 5.353 | 3.922 | 4.417 | 5.05 | 4.982 | 5.118 |
| E | - | - | - | - | - | 4.589 | 2.89 | 4.001 | 5.33 | 3.599 | 4.165 | 4.978 | 4.974 | 4.948 |
| F | - | - | - | - | - | - | 4.093 | 2.334 | 4.469 | 3.26 | -0.685 | 3.006 | 3.549 | 4.128 |
| G | - | - | - | - | - | - | - | 2.932 | 5.051 | 2.225 | 3.561 | 4.682 | 4.52 | 4.746 |
| H | - | - | - | - | - | - | - | - | 4.724 | 1.767 | 2.306 | 4.008 | 3.994 | 4.434 |
| I | - | - | - | - | - | - | - | - | - | 5.321 | 4.874 | 3.709 | 3.777 | 2.731 |
| J | - | - | - | - | - | - | - | - | - | - | 2.982 | 4.315 | 4.258 | 4.547 |
| K | - | - | - | - | - | - | - | - | - | - | - | 3.93 | 3.583 | 4.107 |
| L | - | - | - | - | - | - | - | - | - | - | - | - | 0.367 | 0.922 |
| M | - | - | - | - | - | - | - | - | - | - | - | - | - | 2.762 |
| N | - | - | - | - | - | - | - | - | - | - | - | - | - | - |
