## Supplementary Data S2 for "Interlaboratory comparison of standardised metabolomics and lipidomics analyses in human and rodent blood using the MxP® Quant 500 kit": CE(18.1).pdf

**Amount of relative differences between laboratories**

| Min | A | B | C | D | E | F | G | H | I | J | K | L | M | N |
|---|---|---|---|---|---|---|---|---|---|---|---|---|---|---|
| A | - | 5.632 | 3.042 | 6.342 | 3.279 | 5.413 | 5.957 | 4.475 | 4.324 | 2.467 | 5.566 | 6.259 | 2.939 | 5.593 |
| B | - | - | 5.973 | 5.517 | 2.556 | 3.62 | 4.93 | 5.114 | 5.073 | 5.862 | 2.908 | 5.121 | 5.661 | 2.709 |
| C | - | - | - | 6.384 | 4.929 | 5.368 | 6.038 | 4.239 | 3.773 | 2.773 | 5.444 | 6.25 | 1.1 | 5.369 |
| D | - | - | - | - | 4.521 | 5.833 | -2.797 | 6.016 | 5.778 | 6.632 | 5.654 | 2.159 | 6.264 | 4.345 |
| E | - | - | - | - | - | 4.954 | 3.889 | 5.23 | 3.169 | 6.069 | 4.421 | 3.004 | 5.89 | 4.084 |
| F | - | - | - | - | - | - | 5.491 | 4.314 | 4.023 | 5.309 | 2.433 | 5.899 | 5.129 | 3.348 |
| G | - | - | - | - | - | - | - | 6.11 | 5.934 | 6.692 | 5.681 | 3.084 | 6.358 | 5.021 |
| H | - | - | - | - | - | - | - | - | -1.764 | 4 | 4.517 | 5.975 | 3.951 | 4.66 |
| I | - | - | - | - | - | - | - | - | - | 4.417 | 5.024 | 5.99 | 4.082 | 4.675 |
| J | - | - | - | - | - | - | - | - | - | - | 5.115 | 6.192 | 1.28 | 5.542 |
| K | - | - | - | - | - | - | - | - | - | - | - | 5.526 | 5.258 | 2.121 |
| L | - | - | - | - | - | - | - | - | - | - | - | - | 6.471 | 4.631 |
| M | - | - | - | - | - | - | - | - | - | - | - | - | - | 5.366 |
| N | - | - | - | - | - | - | - | - | - | - | - | - | - | - |

| Median | A | B | C | D | E | F | G | H | I | J | K | L | M | N |
|---|---|---|---|---|---|---|---|---|---|---|---|---|---|---|
| A | - | 3.908 | 2.488 | 4.091 | 3.938 | 2.911 | -0.086 | 2.417 | 4.178 | -2.577 | 2.268 | 2.327 | 3.931 | 4.647 |
| B | - | - | 3.466 | 5.23 | -1.06 | 3.142 | 4.077 | 3.53 | 1.817 | 4.138 | 3.559 | 3.587 | -2.633 | 3.533 |
| C | - | - | - | 4.596 | 3.313 | 0.964 | 2.351 | -1.546 | 3.725 | 2.563 | -0.212 | -0.557 | 3.349 | 4.365 |
| D | - | - | - | - | 4.701 | 4.351 | 3.823 | 4.175 | 4.865 | 3.791 | 4.21 | 4.125 | 4.741 | 5.115 |
| E | - | - | - | - | - | 3.012 | 4.008 | 3.467 | 1.833 | 4.087 | 3.463 | 3.484 | -0.536 | 3.554 |
| F | - | - | - | - | - | - | 2.91 | 1.325 | 3.498 | 3.017 | 1.464 | 1.656 | 3.046 | 4.227 |
| G | - | - | - | - | - | - | - | 2.185 | 4.126 | -1.163 | 2.285 | 2.044 | 3.87 | 4.59 |
| H | - | - | - | - | - | - | - | - | 3.754 | 2.458 | -1.571 | -1.248 | 3.392 | 4.378 |
| I | - | - | - | - | - | - | - | - | - | 4.45 | 4.017 | 3.991 | 1.787 | 3.097 |
| J | - | - | - | - | - | - | - | - | - | - | 2.281 | 2.272 | 3.925 | 4.637 |
| K | - | - | - | - | - | - | - | - | - | - | - | -1.517 | 3.415 | 4.381 |
| L | - | - | - | - | - | - | - | - | - | - | - | - | 3.472 | 4.382 |
| M | - | - | - | - | - | - | - | - | - | - | - | - | - | 3.543 |
| N | - | - | - | - | - | - | - | - | - | - | - | - | - | - |

| Max | A | B | C | D | E | F | G | H | I | J | K | L | M | N |
|---|---|---|---|---|---|---|---|---|---|---|---|---|---|---|
| A | - | 3.792 | 2.466 | 4.333 | 3.986 | 2.681 | 1.926 | 2.211 | 4.172 | -6.541 | 1.776 | -0.904 | 3.96 | 4.59 |
| B | - | - | 3.23 | 5.328 | 0.088 | 3.124 | 4.361 | 3.403 | 2.33 | 4.023 | 3.576 | 3.968 | 1.151 | 3.625 |
| C | - | - | - | 4.786 | 3.01 | -0.499 | 3.404 | 0.203 | 3.723 | 2.663 | 1.211 | 2.73 | 3.394 | 4.303 |
| D | - | - | - | - | 4.931 | 4.514 | 3.932 | 4.429 | 5.024 | 4.137 | 4.376 | 4.206 | 4.952 | 5.208 |
| E | - | - | - | - | - | 2.876 | 4.208 | 3.314 | 2.025 | 3.938 | 3.418 | 3.665 | 1.644 | 3.527 |
| F | - | - | - | - | - | - | 3.534 | 0.827 | 3.634 | 2.817 | 1.597 | 3.004 | 3.324 | 4.254 |
| G | - | - | - | - | - | - | - | 2.935 | 4.367 | 1.575 | 2.747 | 2.351 | 4.212 | 4.725 |
| H | - | - | - | - | - | - | - | - | 3.812 | 2.363 | 0.133 | 2.414 | 3.53 | 4.365 |
| I | - | - | - | - | - | - | - | - | - | 4.451 | 4.152 | 4.464 | 1.538 | 2.972 |
| J | - | - | - | - | - | - | - | - | - | - | 1.958 | -1.108 | 3.991 | 4.584 |
| K | - | - | - | - | - | - | - | - | - | - | - | 1.982 | 3.661 | 4.428 |
| L | - | - | - | - | - | - | - | - | - | - | - | - | 3.921 | 4.508 |
| M | - | - | - | - | - | - | - | - | - | - | - | - | - | 3.381 |
| N | - | - | - | - | - | - | - | - | - | - | - | - | - | - |

| | |
|---|---|
| | Larger than 10 |
| | Between 8 and 9,99 |
| | Between 6 and 7,99 |
| | Between 4 and 5,99 |
| | Between 2 and 3,99 |
| | Between 0 and 1,99 |
| | NA (Calculation failed) |
