## Supplementary Data S2 for "Interlaboratory comparison of standardised metabolomics and lipidomics analyses in human and rodent blood using the MxP® Quant 500 kit": CE(20.0).pdf

**Amount of relative differences between laboratories**

| Min | A | B | C | D | E | F | G | H | I | J | K | L | M | N |
|---|---|---|---|---|---|---|---|---|---|---|---|---|---|---|
| A | - | 5.422 | 5.172 | 1.849 | 5.084 | 2.74 | 3.307 | 3.733 | 4.837 | 5.343 | 5.863 | 7.356 | 4.067 | 1.342 |
| B | - | - | 5.308 | 3.183 | 5.596 | 3.598 | 6.096 | 3.726 | 4.858 | 6.015 | 3.377 | 5.416 | 4.842 | 4.697 |
| C | - | - | - | 1.998 | 6.432 | 2.835 | 6.561 | 4.203 | 5.772 | 6.157 | 0.897 | -0.27 | 5.98 | 5.643 |
| D | - | - | - | - | 4.33 | 1.694 | 4.076 | 3.024 | 3.257 | 5.701 | 5.426 | 7.325 | 2.322 | 4.402 |
| E | - | - | - | - | - | 4.118 | 5.268 | 4.099 | 5.942 | 5.45 | 5.558 | 6.796 | 4.362 | 4.335 |
| F | - | - | - | - | - | - | 4.146 | 0.893 | 4.428 | 5.798 | 5.315 | 7.359 | 3.436 | 4.754 |
| G | - | - | - | - | - | - | - | 4.495 | 7.326 | 2.928 | 7.324 | 7.927 | 6.586 | 6.529 |
| H | - | - | - | - | - | - | - | - | 7.52 | 2.086 | 7.034 | 8.28 | 6.817 | 6.725 |
| I | - | - | - | - | - | - | - | - | - | 5.644 | 5.651 | 5.384 | 4.564 | 4.269 |
| J | - | - | - | - | - | - | - | - | - | - | 6.961 | 7.639 | 6.489 | 6.144 |
| K | - | - | - | - | - | - | - | - | - | - | - | 5.189 | 4.677 | 4.846 |
| L | - | - | - | - | - | - | - | - | - | - | - | - | 6.167 | 5.881 |
| M | - | - | - | - | - | - | - | - | - | - | - | - | - | 1.498 |
| N | - | - | - | - | - | - | - | - | - | - | - | - | - | - |

| Median | A | B | C | D | E | F | G | H | I | J | K | L | M | N |
|---|---|---|---|---|---|---|---|---|---|---|---|---|---|---|
| A | - | 5.077 | 4.577 | 3.702 | 4.289 | 4.232 | 4.623 | 3.655 | 4.441 | 4.667 | 6.255 | 6.614 | 2.7 | 2.201 |
| B | - | - | 4.877 | 2.071 | 4.602 | 3.396 | 5.762 | 3.51 | 3.706 | 5.296 | 5.968 | 4.162 | 4.161 | 4.566 |
| C | - | - | - | 2.955 | 5.586 | 3.793 | 6.114 | 4.024 | 4.976 | 5.508 | 5.7 | 3.612 | 5.347 | 5.492 |
| D | - | - | - | - | 4.164 | 2.727 | 4.609 | 2.664 | 0.982 | 4.843 | 6.386 | 6.624 | 0.671 | 4.25 |
| E | - | - | - | - | - | 4.228 | 5.322 | 4.001 | 5.455 | 4.771 | 6.169 | 5.968 | 3.247 | 3.931 |
| F | - | - | - | - | - | - | 4.504 | -4.213 | 1.571 | 4.84 | 6.493 | 6.685 | 1.213 | 4.535 |
| G | - | - | - | - | - | - | - | 4.768 | 7.094 | 4.495 | 7.652 | 7.379 | 6.353 | 6.425 |
| H | - | - | - | - | - | - | - | - | 6.985 | 1.534 | 6.904 | 7.631 | 6.082 | 6.488 |
| I | - | - | - | - | - | - | - | - | - | 5.097 | 6.038 | 3.798 | 4.378 | 4.313 |
| J | - | - | - | - | - | - | - | - | - | - | 6.623 | 6.902 | 5.57 | 5.848 |
| K | - | - | - | - | - | - | - | - | - | - | - | 1.177 | 5.186 | 5.029 |
| L | - | - | - | - | - | - | - | - | - | - | - | - | 4.639 | 5.526 |
| M | - | - | - | - | - | - | - | - | - | - | - | - | - | 0.36 |
| N | - | - | - | - | - | - | - | - | - | - | - | - | - | - |

| Max | A | B | C | D | E | F | G | H | I | J | K | L | M | N |
|---|---|---|---|---|---|---|---|---|---|---|---|---|---|---|
| A | - | 3.608 | 2.811 | 3.887 | 1.108 | 4.387 | 5.452 | 1.518 | 2.906 | 4.279 | 6.624 | 5.461 | 3.509 | 4.223 |
| B | - | - | 1.797 | 1.714 | 3.598 | 3.357 | 5.168 | 3.834 | 4.322 | 5.157 | 6.984 | 5.153 | -0.109 | 3.078 |
| C | - | - | - | 3.081 | 3.311 | 3.912 | 5.193 | 3.79 | 4.049 | 4.948 | 6.882 | 5.203 | 0.705 | 3.489 |
| D | - | - | - | - | 3.87 | 2.851 | 5.095 | 4.225 | 4.694 | 5.33 | 7.06 | 5.107 | 2.121 | 2.208 |
| E | - | - | - | - | - | 4.247 | 5.39 | -1.346 | 2.883 | 4.403 | 6.677 | 5.396 | 3.293 | 4.194 |
| F | - | - | - | - | - | - | 4.865 | 4.738 | 5.149 | 5.677 | 7.245 | 4.878 | 3.644 | 1.475 |
| G | - | - | - | - | - | - | - | 6.535 | 6.446 | 6.803 | 7.973 | -1.937 | 5.827 | 5.394 |
| H | - | - | - | - | - | - | - | - | 3.139 | 4.584 | 6.727 | 5.471 | 3.424 | 4.284 |
| I | - | - | - | - | - | - | - | - | - | 3.622 | 6.403 | 5.528 | 3.981 | 4.579 |
| J | - | - | - | - | - | - | - | - | - | - | 6.06 | 5.712 | 4.711 | 5.001 |
| K | - | - | - | - | - | - | - | - | - | - | - | 6.161 | 5.788 | 5.888 |
| L | - | - | - | - | - | - | - | - | - | - | - | - | 5.831 | 5.377 |
| M | - | - | - | - | - | - | - | - | - | - | - | - | - | 3.206 |
| N | - | - | - | - | - | - | - | - | - | - | - | - | - | - |

| | |
|---|---|
| (red) | Larger than 10 |
| (orange) | Between 8 and 9,99 |
| (light orange) | Between 6 and 7,99 |
| (light green) | Between 4 and 5,99 |
| (green) | Between 2 and 3,99 |
| (dark green) | Between 0 and 1,99 |
| (grey) | NA (Calculation failed) |
