## Supplementary Data S2 for "Interlaboratory comparison of standardised metabolomics and lipidomics analyses in human and rodent blood using the MxP® Quant 500 kit": CE(20.1).pdf

**Amount of relative differences between laboratories**

| Min | A | B | C | D | E | F | G | H | I | J | K | L | M | N |
|---|---|---|---|---|---|---|---|---|---|---|---|---|---|---|
| A | - | 5.373 | 5.336 | 5.849 | 3.952 | 5.591 | 5.828 | 5.561 | 7.846 | 6.151 | 6.43 | 5.864 | 6.347 | 8.159 |
| B | - | - | 3.937 | 4.975 | 5.116 | 5.041 | 4.776 | 2.535 | 7.451 | 5.508 | 5.79 | 7.103 | 5.815 | 7.9 |
| C | - | - | - | 1.089 | 6.596 | 2.62 | 2.548 | 5.671 | 6.33 | 1.661 | 6.154 | 7.954 | 5.398 | 11.221 |
| D | - | - | - | - | 6.556 | 2.652 | 2.525 | 5.396 | 7.136 | 4.291 | 4.587 | 7.788 | 4.859 | 8.065 |
| E | - | - | - | - | - | 5.979 | 5.555 | 4.915 | 7.359 | 5.975 | 6.094 | 6.215 | 6.078 | 7.839 |
| F | - | - | - | - | - | - | 5.477 | 6.291 | 5.784 | 4.533 | 2.988 | 8.211 | 4.445 | 7.297 |
| G | - | - | - | - | - | - | - | 4.946 | 7.6 | 4.896 | 5.203 | 7.789 | 5.282 | 8.284 |
| H | - | - | - | - | - | - | - | - | 6.817 | 5.491 | 5.478 | 6.875 | 5.614 | 7.603 |
| I | - | - | - | - | - | - | - | - | - | 2.445 | 3.638 | 8.286 | 1.039 | 7.039 |
| J | - | - | - | - | - | - | - | - | - | - | 3.33 | 7.95 | 1.883 | 7.209 |
| K | - | - | - | - | - | - | - | - | - | - | - | 7.645 | 3.843 | 6.585 |
| L | - | - | - | - | - | - | - | - | - | - | - | - | 6.944 | 9.241 |
| M | - | - | - | - | - | - | - | - | - | - | - | - | - | 8.043 |
| N | - | - | - | - | - | - | - | - | - | - | - | - | - | - |

| Median | A | B | C | D | E | F | G | H | I | J | K | L | M | N |
|---|---|---|---|---|---|---|---|---|---|---|---|---|---|---|
| A | - | 4.878 | 3.533 | 5.653 | 2.526 | 5.321 | 5.553 | 4.677 | 5.69 | 5.814 | 4.979 | 5.336 | 5.944 | 6.032 |
| B | - | - | 0.318 | 4.815 | 5.094 | 4.767 | 4.693 | 2.473 | 5.029 | 5.073 | 2.967 | 6.683 | 5.529 | 5.8 |
| C | - | - | - | 0.333 | 6.338 | 2.875 | 1.859 | 5.47 | 2.869 | 0.823 | 4.228 | 7.553 | 5.293 | 9.125 |
| D | - | - | - | - | 6.404 | 1.79 | -0.366 | 5.509 | 3.113 | 3.068 | 4.553 | 7.413 | 4.715 | 5.443 |
| E | - | - | - | - | - | 5.794 | 5.495 | 4.409 | 5.653 | 5.731 | 4.975 | 5.618 | 5.9 | 6.076 |
| F | - | - | - | - | - | - | 4.682 | 6.102 | 3.357 | 3.634 | 4.972 | 7.829 | 4.571 | 4.21 |
| G | - | - | - | - | - | - | - | 5.419 | 3.373 | 3.797 | 4.541 | 7.432 | 4.79 | 5.4 |
| H | - | - | - | - | - | - | - | - | 5.217 | 5.253 | 3.973 | 6.367 | 5.626 | 5.945 |
| I | - | - | - | - | - | - | - | - | - | 2.69 | 4.863 | 7.874 | 4.23 | 5.45 |
| J | - | - | - | - | - | - | - | - | - | - | 4.392 | 7.515 | 4.602 | 5.641 |
| K | - | - | - | - | - | - | - | - | - | - | - | 7.151 | 5.207 | 5.865 |
| L | - | - | - | - | - | - | - | - | - | - | - | - | 6.279 | 6.803 |
| M | - | - | - | - | - | - | - | - | - | - | - | - | - | 4.535 |
| N | - | - | - | - | - | - | - | - | - | - | - | - | - | - |

| Max | A | B | C | D | E | F | G | H | I | J | K | L | M | N |
|---|---|---|---|---|---|---|---|---|---|---|---|---|---|---|
| A | - | 2.223 | -0.759 | 4.142 | 4.765 | 3.622 | 4.661 | 4.738 | 3.983 | 4.95 | 6.404 | 4.454 | 5.173 | 4.737 |
| B | - | - | 2.041 | 3.754 | 5.016 | 3.028 | 4.491 | 5.044 | 4.402 | 5.164 | 6.542 | 4.232 | 5.049 | 4.377 |
| C | - | - | - | 4.031 | 4.754 | 3.503 | 4.614 | 4.823 | 4.345 | 4.942 | 6.536 | 4.526 | 5.146 | 7.679 |
| D | - | - | - | - | 5.72 | 2.589 | 3.349 | 5.75 | 5.365 | 5.841 | 6.96 | 2.676 | 4.507 | 5.649 |
| E | - | - | - | - | - | 4.908 | 5.354 | -1.552 | 2.868 | 2.156 | 5.552 | 5.28 | 5.634 | -4.836 |
| F | - | - | - | - | - | - | 3.991 | 5.502 | 5.126 | 5.662 | 6.846 | 3.727 | 4.719 | 5.303 |
| G | - | - | - | - | - | - | - | 6.165 | 5.883 | 6.31 | 7.24 | 1.773 | 3.717 | 6.137 |
| H | - | - | - | - | - | - | - | - | 3.112 | 1.001 | 5.514 | 5.295 | 5.64 | 2.619 |
| I | - | - | - | - | - | - | - | - | - | 3.628 | 6.012 | 5.21 | 5.511 | 2.811 |
| J | - | - | - | - | - | - | - | - | - | - | 5.467 | 5.407 | 5.668 | 3.158 |
| K | - | - | - | - | - | - | - | - | - | - | - | 5.928 | 6.046 | 5.41 |
| L | - | - | - | - | - | - | - | - | - | - | - | - | 4.253 | 6.562 |
| M | - | - | - | - | - | - | - | - | - | - | - | - | - | 6.273 |
| N | - | - | - | - | - | - | - | - | - | - | - | - | - | - |

| | |
|---|---|
| | Larger than 10 |
| | Between 8 and 9,99 |
| | Between 6 and 7,99 |
| | Between 4 and 5,99 |
| | Between 2 and 3,99 |
| | Between 0 and 1,99 |
| | NA (Calculation failed) |
