## Supplementary Data S2 for "Interlaboratory comparison of standardised metabolomics and lipidomics analyses in human and rodent blood using the MxP® Quant 500 kit": CE(22.0).pdf

**Amount of relative differences between laboratories**

| Min | A | B | C | D | E | F | G | H | I | J | K | L | M | N |
|---|---|---|---|---|---|---|---|---|---|---|---|---|---|---|
| A | - | 6.909 | 5.471 | NA | 6.487 | 5.716 | NA | 7.87 | 7.663 | NA | 6.517 | NA | NA | 5.561 |
| B | - | - | 3.728 | NA | 7.004 | 4.671 | NA | 7.35 | 8.398 | NA | 7.643 | NA | NA | 6.621 |
| C | - | - | - | NA | 6.177 | 1.359 | NA | 10.374 | 7.57 | NA | 7.039 | NA | NA | 5.355 |
| D | - | - | - | - | NA | NA | NA | NA | NA | NA | NA | NA | NA | NA |
| E | - | - | - | - | - | 5.764 | NA | 6.719 | 5.449 | NA | 7.856 | NA | NA | 2.537 |
| F | - | - | - | - | - | - | NA | 9.286 | 7.617 | NA | 6.974 | NA | NA | 5.28 |
| G | - | - | - | - | - | - | - | NA | NA | NA | NA | NA | NA | NA |
| H | - | - | - | - | - | - | - | - | 7.056 | NA | 7.132 | NA | NA | 4.597 |
| I | - | - | - | - | - | - | - | - | - | NA | 10.339 | NA | NA | 5.223 |
| J | - | - | - | - | - | - | - | - | - | - | NA | NA | NA | NA |
| K | - | - | - | - | - | - | - | - | - | - | - | NA | NA | 4.967 |
| L | - | - | - | - | - | - | - | - | - | - | - | - | NA | NA |
| M | - | - | - | - | - | - | - | - | - | - | - | - | - | NA |
| N | - | - | - | - | - | - | - | - | - | - | - | - | - | - |

| Median | A | B | C | D | E | F | G | H | I | J | K | L | M | N |
|---|---|---|---|---|---|---|---|---|---|---|---|---|---|---|
| A | - | 5.948 | 4.7 | NA | 5.537 | 5.216 | NA | 4.379 | 4.444 | NA | 3.79 | NA | NA | 5.036 |
| B | - | - | 3.518 | NA | 5.984 | 3.475 | NA | 5.087 | 7.473 | NA | 7.399 | NA | NA | 6.405 |
| C | - | - | - | NA | 4.98 | 3.066 | NA | 6.765 | 6.813 | NA | 6.921 | NA | NA | 5.211 |
| D | - | - | - | - | NA | NA | NA | NA | NA | NA | NA | NA | NA | NA |
| E | - | - | - | - | - | 4.371 | NA | 1.59 | 5.927 | NA | 2.23 | NA | NA | 3.751 |
| F | - | - | - | - | - | - | NA | 6.327 | 6.976 | NA | 7.06 | NA | NA | 5.327 |
| G | - | - | - | - | - | - | - | NA | NA | NA | NA | NA | NA | NA |
| H | - | - | - | - | - | - | - | - | 6.347 | NA | 4.347 | NA | NA | 4.321 |
| I | - | - | - | - | - | - | - | - | - | NA | 8.428 | NA | NA | 5.066 |
| J | - | - | - | - | - | - | - | - | - | - | NA | NA | NA | NA |
| K | - | - | - | - | - | - | - | - | - | - | - | NA | NA | 5.053 |
| L | - | - | - | - | - | - | - | - | - | - | - | - | NA | NA |
| M | - | - | - | - | - | - | - | - | - | - | - | - | - | NA |
| N | - | - | - | - | - | - | - | - | - | - | - | - | - | - |

| Max | A | B | C | D | E | F | G | H | I | J | K | L | M | N |
|---|---|---|---|---|---|---|---|---|---|---|---|---|---|---|
| A | - | 4.819 | 3.285 | NA | 2.818 | 3.529 | NA | 4.981 | 7.025 | NA | 7.055 | NA | NA | 5.002 |
| B | - | - | 2.995 | NA | 4.092 | 3.679 | NA | 1.151 | 5.442 | NA | 6.782 | NA | NA | 4.529 |
| C | - | - | - | NA | 4.239 | 4.175 | NA | 7.575 | 5.563 | NA | 6.668 | NA | NA | 4.287 |
| D | - | - | - | - | NA | NA | NA | NA | NA | NA | NA | NA | NA | NA |
| E | - | - | - | - | - | 5.146 | NA | 4.89 | 6.234 | NA | 8.727 | NA | NA | 5.583 |
| F | - | - | - | - | - | - | NA | 5.741 | 6.062 | NA | 7.21 | NA | NA | 5.52 |
| G | - | - | - | - | - | - | - | NA | NA | NA | NA | NA | NA | NA |
| H | - | - | - | - | - | - | - | - | 5.249 | NA | 8.338 | NA | NA | -0.062 |
| I | - | - | - | - | - | - | - | - | - | NA | 10.503 | NA | NA | 4.02 |
| J | - | - | - | - | - | - | - | - | - | - | NA | NA | NA | NA |
| K | - | - | - | - | - | - | - | - | - | - | - | NA | NA | 5.387 |
| L | - | - | - | - | - | - | - | - | - | - | - | - | NA | NA |
| M | - | - | - | - | - | - | - | - | - | - | - | - | - | NA |
| N | - | - | - | - | - | - | - | - | - | - | - | - | - | - |

| | |
|---|---|
| | Larger than 10 |
| | Between 8 and 9,99 |
| | Between 6 and 7,99 |
| | Between 4 and 5,99 |
| | Between 2 and 3,99 |
| | Between 0 and 1,99 |
| | NA (Calculation failed) |
