## Supplementary Data S2 for "Interlaboratory comparison of standardised metabolomics and lipidomics analyses in human and rodent blood using the MxP® Quant 500 kit": CE(22.1).pdf

**Amount of relative differences between laboratories**

| Min | A | B | C | D | E | F | G | H | I | J | K | L | M | N |
|---|---|---|---|---|---|---|---|---|---|---|---|---|---|---|
| A | - | 2.613 | NA | 3.194 | 7.021 | -0.112 | 5.976 | NA | 6.451 | 3.262 | 10.694 | 8.557 | NA | 6.355 |
| B | - | - | NA | 8.072 | 8.509 | 7.171 | 7.965 | NA | 8.843 | 7.095 | 8.636 | 9.694 | NA | 8.802 |
| C | - | - | - | NA | NA | NA | NA | NA | NA | NA | NA | NA | NA | NA |
| D | - | - | - | - | 6.984 | 3.483 | 5.897 | NA | 5.602 | 2.982 | 6.709 | 8.56 | NA | 5.581 |
| E | - | - | - | - | - | 9.185 | 4.948 | NA | 7.914 | 9.968 | 7.857 | 7.277 | NA | 7.993 |
| F | - | - | - | - | - | - | 7.087 | NA | 8.334 | 4.431 | 8.848 | 9.186 | NA | 8.321 |
| G | - | - | - | - | - | - | - | NA | 7.65 | 7.515 | 7.66 | 7.981 | NA | 7.642 |
| H | - | - | - | - | - | - | - | - | NA | NA | NA | NA | NA | NA |
| I | - | - | - | - | - | - | - | - | - | 5.022 | 4.739 | 7.807 | NA | 2.887 |
| J | - | - | - | - | - | - | - | - | - | - | 7.277 | 8.953 | NA | 6.402 |
| K | - | - | - | - | - | - | - | - | - | - | - | 7.864 | NA | 4.417 |
| L | - | - | - | - | - | - | - | - | - | - | - | - | NA | 8.83 |
| M | - | - | - | - | - | - | - | - | - | - | - | - | - | NA |
| N | - | - | - | - | - | - | - | - | - | - | - | - | - | - |

| | |
|---|---|
| | Larger than 10 |
| | Between 8 and 9,99 |
| | Between 6 and 7,99 |
| | Between 4 and 5,99 |
| | Between 2 and 3,99 |
| | Between 0 and 1,99 |
| | NA (Calculation failed) |

| Median | A | B | C | D | E | F | G | H | I | J | K | L | M | N |
|---|---|---|---|---|---|---|---|---|---|---|---|---|---|---|
| A | - | 5.417 | NA | 3.253 | 6.269 | 4.642 | 5.01 | NA | 5.363 | 3.8 | 9.317 | 7.66 | NA | 4.084 |
| B | - | - | NA | 6.09 | 7.926 | 5.387 | 7.417 | NA | 7.804 | 6.623 | 7.706 | 8.886 | NA | 7.347 |
| C | - | - | - | NA | NA | NA | NA | NA | NA | NA | NA | NA | NA | NA |
| D | - | - | - | - | 6.263 | 4.086 | 4.965 | NA | 5.038 | 2.921 | 2.42 | 7.681 | NA | 3.849 |
| E | - | - | - | - | - | 5.941 | 4.774 | NA | 5.287 | 9.012 | 4.595 | 6.471 | NA | 4.18 |
| F | - | - | - | - | - | - | 6.484 | NA | 6.054 | 4.215 | 7.607 | 8.365 | NA | 5.251 |
| G | - | - | - | - | - | - | - | NA | 1.373 | 6.238 | 3.396 | 7.27 | NA | 3.907 |
| H | - | - | - | - | - | - | - | - | NA | NA | NA | NA | NA | NA |
| I | - | - | - | - | - | - | - | - | - | 4.896 | 0.113 | 6.787 | NA | 3.858 |
| J | - | - | - | - | - | - | - | - | - | - | 5.161 | 8.117 | NA | 5.075 |
| K | - | - | - | - | - | - | - | - | - | - | - | 6.837 | NA | 3.808 |
| L | - | - | - | - | - | - | - | - | - | - | - | - | NA | 5.734 |
| M | - | - | - | - | - | - | - | - | - | - | - | - | - | NA |
| N | - | - | - | - | - | - | - | - | - | - | - | - | - | - |

| Max | A | B | C | D | E | F | G | H | I | J | K | L | M | N |
|---|---|---|---|---|---|---|---|---|---|---|---|---|---|---|
| A | - | 5.989 | NA | 4.495 | 6.201 | 5.614 | 6.273 | NA | 4.115 | 4.864 | 9.514 | 6.309 | NA | 5.303 |
| B | - | - | NA | 6.169 | 6.67 | 5.729 | 6.79 | NA | 6.214 | 5.131 | 5.504 | 6.655 | NA | 6.384 |
| C | - | - | - | NA | NA | NA | NA | NA | NA | NA | NA | NA | NA | NA |
| D | - | - | - | - | 6.029 | 5.43 | 6.125 | NA | 3.702 | 2.683 | 7 | 6.19 | NA | 3.876 |
| E | - | - | - | - | - | 8.72 | 3.893 | NA | 7.756 | 10.208 | 7.994 | 4.211 | NA | 7.551 |
| F | - | - | - | - | - | - | 6.247 | NA | 8.006 | 3.015 | 7.341 | 6.293 | NA | 7.692 |
| G | - | - | - | - | - | - | - | NA | 7.952 | 8.232 | 8.212 | 2.149 | NA | 7.656 |
| H | - | - | - | - | - | - | - | - | NA | NA | NA | NA | NA | NA |
| I | - | - | - | - | - | - | - | - | - | 4.336 | 5.059 | 6.19 | NA | 4.37 |
| J | - | - | - | - | - | - | - | - | - | - | 6.959 | 6.226 | NA | 3.393 |
| K | - | - | - | - | - | - | - | - | - | - | - | 6.273 | NA | 5.484 |
| L | - | - | - | - | - | - | - | - | - | - | - | - | NA | 8.21 |
| M | - | - | - | - | - | - | - | - | - | - | - | - | - | NA |
| N | - | - | - | - | - | - | - | - | - | - | - | - | - | - |
