## Supplementary Data S2 for "Interlaboratory comparison of standardised metabolomics and lipidomics analyses in human and rodent blood using the MxP® Quant 500 kit": CE(22.6).pdf

**Amount of relative differences between laboratories**

| Min | A | B | C | D | E | F | G | H | I | J | K | L | M | N |
|---|---|---|---|---|---|---|---|---|---|---|---|---|---|---|
| A | - | 5.063 | 4.057 | 1.158 | 6.973 | 4.833 | 4.704 | 3.572 | 0.929 | 3.808 | -0.845 | 5.067 | 4.995 | 3.417 |
| B | - | - | 4.474 | 5.198 | 7.703 | 1.847 | 6.066 | 4.794 | 5.118 | 4.51 | 5.434 | 6.234 | 1.56 | 4.326 |
| C | - | - | - | 3.84 | 7.293 | 3.909 | 5.328 | 2.255 | 3.595 | 0.028 | 4.15 | 5.569 | 4.027 | -1.747 |
| D | - | - | - | - | 6.871 | 5.086 | 4.559 | 4.211 | 2.555 | 4.286 | 2.953 | 5.013 | 5.276 | 3.738 |
| E | - | - | - | - | - | 6.163 | 5.353 | 6.035 | 5.839 | 6.024 | 5.924 | 5.18 | 6.25 | 5.894 |
| F | - | - | - | - | - | - | 6.012 | 4.548 | 4.981 | 4.249 | 5.272 | 6.183 | 0.928 | 4.192 |
| G | - | - | - | - | - | - | - | 5.773 | 5.077 | 5.719 | 5.55 | 3.195 | 6.225 | 5.083 |
| H | - | - | - | - | - | - | - | - | 3.118 | 1.68 | 3.652 | 5.484 | 4.394 | 0.978 |
| I | - | - | - | - | - | - | - | - | - | 3.614 | -0.756 | 5.208 | 4.968 | 3.103 |
| J | - | - | - | - | - | - | - | - | - | - | 4.015 | 5.61 | 4.209 | -0.851 |
| K | - | - | - | - | - | - | - | - | - | - | - | 4.918 | 4.917 | 3.616 |
| L | - | - | - | - | - | - | - | - | - | - | - | - | 6.505 | 5.432 |
| M | - | - | - | - | - | - | - | - | - | - | - | - | - | 4.201 |
| N | - | - | - | - | - | - | - | - | - | - | - | - | - | - |

| | |
|---|---|
| | Larger than 10 |
| | Between 8 and 9,99 |
| | Between 6 and 7,99 |
| | Between 4 and 5,99 |
| | Between 2 and 3,99 |
| | Between 0 and 1,99 |
| | NA (Calculation failed) |

| Median | A | B | C | D | E | F | G | H | I | J | K | L | M | N |
|---|---|---|---|---|---|---|---|---|---|---|---|---|---|---|
| A | - | 3.883 | 0.928 | 2.087 | 6.356 | 4.131 | 2.218 | 1.628 | 1.909 | 2.598 | 2.565 | 2.943 | 3.948 | 4.218 |
| B | - | - | 3.996 | 3.936 | 6.883 | 1.402 | 3.864 | 3.775 | 3.879 | 3.385 | 4.719 | 3.421 | -1.528 | 1.889 |
| C | - | - | - | 0.045 | 6.449 | 4.001 | 0.653 | 0.815 | 0.324 | 2.208 | 3.122 | 2.27 | 3.806 | 4.084 |
| D | - | - | - | - | 6.499 | 3.687 | -1.265 | 1.45 | 0.129 | -0.965 | 3.491 | 1.75 | 3.397 | 3.83 |
| E | - | - | - | - | - | 5.746 | 5.578 | 5.497 | 5.547 | 5.554 | 5.372 | 5.624 | 5.713 | 5.769 |
| F | - | - | - | - | - | - | 4.068 | 4.089 | 4.118 | 3.765 | 4.875 | 3.677 | 1.263 | 0.253 |
| G | - | - | - | - | - | - | - | 2.834 | 1.321 | 1.701 | 3.662 | 1.494 | 2.921 | 3.557 |
| H | - | - | - | - | - | - | - | - | -6.057 | 1.577 | 3.333 | 1.941 | 3.651 | 3.971 |
| I | - | - | - | - | - | - | - | - | - | 1.103 | 3.326 | 2.061 | 3.583 | 3.94 |
| J | - | - | - | - | - | - | - | - | - | - | 3.75 | -0.023 | 3.305 | 3.722 |
| K | - | - | - | - | - | - | - | - | - | - | - | 3.661 | 4.354 | 4.545 |
| L | - | - | - | - | - | - | - | - | - | - | - | - | 1.758 | 3.046 |
| M | - | - | - | - | - | - | - | - | - | - | - | - | - | 1.77 |
| N | - | - | - | - | - | - | - | - | - | - | - | - | - | - |

| Max | A | B | C | D | E | F | G | H | I | J | K | L | M | N |
|---|---|---|---|---|---|---|---|---|---|---|---|---|---|---|
| A | - | 1.82 | 2.981 | 3.505 | 5.732 | 3.343 | 4.836 | 0.976 | 2.401 | 0.017 | 3.388 | 5.186 | 2.501 | 4.623 |
| B | - | - | 3.56 | 3.028 | 5.872 | 2.773 | 4.715 | 2.489 | 0.829 | 1.394 | 3.852 | 5.098 | 1.204 | 4.444 |
| C | - | - | - | 4.159 | 5.383 | 4.06 | 5.09 | 2.463 | 3.614 | 3.035 | 1.223 | 5.366 | 3.657 | 4.911 |
| D | - | - | - | - | 6.188 | 0.361 | 4.254 | 3.902 | 2.773 | 3.526 | 4.62 | 4.817 | 2.638 | 3.896 |
| E | - | - | - | - | - | 5.378 | 5.732 | 5.064 | 5.252 | 5.15 | 4.799 | 5.868 | 5.262 | 5.666 |
| F | - | - | - | - | - | - | 4.351 | 3.74 | 2.495 | 3.344 | 4.512 | 4.853 | 2.309 | 4 |
| G | - | - | - | - | - | - | - | 5.415 | 5.017 | 5.274 | 5.769 | 3.48 | 5.017 | 2.336 |
| H | - | - | - | - | - | - | - | - | 2.834 | 1.507 | 3.057 | 5.236 | 2.913 | 4.706 |
| I | - | - | - | - | - | - | - | - | - | 2.088 | 4.059 | 5.036 | -0.981 | 4.356 |
| J | - | - | - | - | - | - | - | - | - | - | 3.53 | 5.157 | 2.255 | 4.564 |
| K | - | - | - | - | - | - | - | - | - | - | - | 5.41 | 3.873 | 4.99 |
| L | - | - | - | - | - | - | - | - | - | - | - | - | 5.602 | 4.193 |
| M | - | - | - | - | - | - | - | - | - | - | - | - | - | 4.321 |
| N | - | - | - | - | - | - | - | - | - | - | - | - | - | - |
