## Supplementary Data S2 for "Interlaboratory comparison of standardised metabolomics and lipidomics analyses in human and rodent blood using the MxP® Quant 500 kit": Cer(d16.1-18.0).pdf

**Amount of relative differences between laboratories**

| Min | A | B | C | D | E | F | G | H | I | J | K | L | M | N |
|---|---|---|---|---|---|---|---|---|---|---|---|---|---|---|
| A | - | NA | 3.685 | 4.885 | 5.685 | 5.328 | 7.024 | 0.783 | 4.893 | 8.017 | NA | 5.04 | 4.342 | 3.721 |
| B | - | - | NA | NA | NA | NA | NA | NA | NA | NA | NA | NA | NA | NA |
| C | - | - | - | 4.781 | 5.585 | 4.047 | 3.433 | 4.385 | 0.631 | 6.75 | NA | 4.293 | 5.68 | 2.665 |
| D | - | - | - | - | 4.901 | 4.23 | 7.594 | 5.619 | 4.549 | 5.04 | NA | -0.571 | 2.039 | 4.832 |
| E | - | - | - | - | - | 6.182 | 8.755 | 6.694 | 6.177 | 1.577 | NA | 5.171 | 5.836 | 6.181 |
| F | - | - | - | - | - | - | 8.435 | 2.574 | 2.466 | 0.538 | NA | 4.884 | 4.454 | 3.703 |
| G | - | - | - | - | - | - | - | 3.536 | 3.148 | 6.66 | NA | 4.739 | 5.66 | 1.714 |
| H | - | - | - | - | - | - | - | - | 3.95 | 5.879 | NA | 5.001 | 3.428 | 4.044 |
| I | - | - | - | - | - | - | - | - | - | 2.732 | NA | 4.46 | 1.537 | 1.54 |
| J | - | - | - | - | - | - | - | - | - | - | NA | 3.147 | 5.386 | 5.138 |
| K | - | - | - | - | - | - | - | - | - | - | - | NA | NA | NA |
| L | - | - | - | - | - | - | - | - | - | - | - | - | 4.645 | 4.852 |
| M | - | - | - | - | - | - | - | - | - | - | - | - | - | 3.869 |
| N | - | - | - | - | - | - | - | - | - | - | - | - | - | - |

| Median | A | B | C | D | E | F | G | H | I | J | K | L | M | N |
|---|---|---|---|---|---|---|---|---|---|---|---|---|---|---|
| A | - | NA | 4.801 | 5.07 | 5.583 | 4.461 | 4.243 | 3.835 | -1.417 | 4.985 | NA | 5.262 | 4.346 | 3.621 |
| B | - | - | NA | NA | NA | NA | NA | NA | NA | NA | NA | NA | NA | NA |
| C | - | - | - | 3.097 | 4.858 | 6.083 | 3.43 | 5.817 | 5.256 | 1.509 | NA | 3.915 | 2.576 | 4.429 |
| D | - | - | - | - | 4.44 | 6.408 | 4.452 | 6.179 | 5.678 | 3.896 | NA | 2.834 | 4.16 | 5.011 |
| E | - | - | - | - | - | 7.088 | 5.793 | 6.91 | 6.53 | 5.524 | NA | 4.171 | 5.685 | 6.077 |
| F | - | - | - | - | - | - | 5.095 | 2.609 | 4.253 | 5.221 | NA | 5.662 | 5.074 | 4.943 |
| G | - | - | - | - | - | - | - | 5.368 | 4.619 | 2.819 | NA | 4.55 | 2.052 | 3.259 |
| H | - | - | - | - | - | - | - | - | 3.721 | 5.109 | NA | 5.547 | 4.949 | 4.633 |
| I | - | - | - | - | - | - | - | - | - | 4.518 | NA | 5.277 | 4.358 | 3.758 |
| J | - | - | - | - | - | - | - | - | - | - | NA | 4.296 | 2.146 | 3.709 |
| K | - | - | - | - | - | - | - | - | - | - | - | NA | NA | NA |
| L | - | - | - | - | - | - | - | - | - | - | - | - | 4.677 | 5.421 |
| M | - | - | - | - | - | - | - | - | - | - | - | - | - | 3.441 |
| N | - | - | - | - | - | - | - | - | - | - | - | - | - | - |

| Max | A | B | C | D | E | F | G | H | I | J | K | L | M | N |
|---|---|---|---|---|---|---|---|---|---|---|---|---|---|---|
| A | - | NA | 5.072 | 5.129 | 5.526 | 6.159 | 4.704 | 5.064 | 4.328 | 4.961 | NA | 5.375 | 4.348 | 3.535 |
| B | - | - | NA | NA | NA | NA | NA | NA | NA | NA | NA | NA | NA | NA |
| C | - | - | - | 0.974 | 4.251 | 7.075 | 3.562 | 6.627 | 5.995 | 5.061 | NA | 3.644 | 4.431 | 5.072 |
| D | - | - | - | - | 4.123 | 7.103 | 3.776 | 6.649 | 6.141 | 4.812 | NA | 3.436 | 4.578 | 5.14 |
| E | - | - | - | - | - | 7.543 | 5.228 | 7.132 | 6.734 | 5.896 | NA | 2.949 | 5.641 | 5.988 |
| F | - | - | - | - | - | - | 5.71 | 4.361 | 4.833 | 5.561 | NA | 5.971 | 5.61 | 5.487 |
| G | - | - | - | - | - | - | - | 6.264 | 5.582 | 4.243 | NA | 4.429 | 3.089 | 4.307 |
| H | - | - | - | - | - | - | - | - | 3.537 | 4.797 | NA | 5.786 | 5.185 | 4.974 |
| I | - | - | - | - | - | - | - | - | - | 4.776 | NA | 5.595 | 4.749 | 4.458 |
| J | - | - | - | - | - | - | - | - | - | - | NA | 5.168 | 2.528 | 2.179 |
| K | - | - | - | - | - | - | - | - | - | - | - | NA | NA | NA |
| L | - | - | - | - | - | - | - | - | - | - | - | - | 5.299 | 5.753 |
| M | - | - | - | - | - | - | - | - | - | - | - | - | - | 2.978 |
| N | - | - | - | - | - | - | - | - | - | - | - | - | - | - |

| | |
|---|---|
| (red) | Larger than 10 |
| (orange) | Between 8 and 9,99 |
| (light orange) | Between 6 and 7,99 |
| (light green) | Between 4 and 5,99 |
| (green) | Between 2 and 3,99 |
| (dark green) | Between 0 and 1,99 |
| (grey) | NA (Calculation failed) |
