## Supplementary Data S2 for "Interlaboratory comparison of standardised metabolomics and lipidomics analyses in human and rodent blood using the MxP® Quant 500 kit": Cer(d16.1-20.0).pdf

**Amount of relative differences between laboratories**

| Min | A | B | C | D | E | F | G | H | I | J | K | L | M | N |
|---|---|---|---|---|---|---|---|---|---|---|---|---|---|---|
| A | - | 4.769 | 2.258 | 4.668 | 5.673 | NA | 5.393 | NA | 6.21 | 4.921 | 5.87 | 8.136 | 6.113 | 5.346 |
| B | - | - | 6.551 | 2.473 | 9.104 | NA | 7.767 | NA | 4.686 | 6.69 | 8.255 | 9.422 | 6.866 | 4.324 |
| C | - | - | - | 3.156 | 5.439 | NA | 5.407 | NA | 5.619 | 5.922 | 6.255 | 8.073 | 3.033 | 4.813 |
| D | - | - | - | - | 8.386 | NA | 6.608 | NA | 5.106 | 6.292 | 7.5 | 8.21 | 5.436 | 4.309 |
| E | - | - | - | - | - | NA | 5.703 | NA | 6.734 | 4.32 | 6.779 | 7.689 | 5.08 | 5.93 |
| F | - | - | - | - | - | - | NA | NA | NA | NA | NA | NA | NA | NA |
| G | - | - | - | - | - | - | - | NA | 5.974 | 5.723 | 7.477 | 6.457 | 3.338 | 5.17 |
| H | - | - | - | - | - | - | - | - | NA | NA | NA | NA | NA | NA |
| I | - | - | - | - | - | - | - | - | - | 6.932 | 8.49 | 9.693 | 7.445 | 0.821 |
| J | - | - | - | - | - | - | - | - | - | - | 5.829 | 9.651 | 8.001 | 6.187 |
| K | - | - | - | - | - | - | - | - | - | - | - | 8.994 | 5.656 | 4.887 |
| L | - | - | - | - | - | - | - | - | - | - | - | - | 4.972 | 5.476 |
| M | - | - | - | - | - | - | - | - | - | - | - | - | - | 3.564 |
| N | - | - | - | - | - | - | - | - | - | - | - | - | - | - |

| | |
|---|---|
| | Larger than 10 |
| | Between 8 and 9,99 |
| | Between 6 and 7,99 |
| | Between 4 and 5,99 |
| | Between 2 and 3,99 |
| | Between 0 and 1,99 |
| | NA (Calculation failed) |

| Median | A | B | C | D | E | F | G | H | I | J | K | L | M | N |
|---|---|---|---|---|---|---|---|---|---|---|---|---|---|---|
| A | - | 1.58 | -1.449 | 2.245 | 3.887 | NA | 2.128 | NA | 3.662 | 3.777 | 3.579 | 3.24 | 3.983 | 5.051 |
| B | - | - | 1.391 | 2.999 | 4.046 | NA | 0.493 | NA | 3.863 | 3.58 | 3.588 | 3.603 | 4.046 | 4.989 |
| C | - | - | - | 1.411 | 3.915 | NA | 2.176 | NA | 3.29 | 3.979 | 3.55 | 3.291 | 3.744 | 4.825 |
| D | - | - | - | - | 4.43 | NA | 3.085 | NA | 2.721 | 4.185 | 2.546 | 4.066 | 3.221 | 4.579 |
| E | - | - | - | - | - | NA | 3.697 | NA | 5.076 | 1.249 | 4.967 | 2.56 | 5.155 | 5.821 |
| F | - | - | - | - | - | - | NA | NA | NA | NA | NA | NA | NA | NA |
| G | - | - | - | - | - | - | - | NA | 4.064 | 3.391 | 3.789 | 2.714 | 4.33 | 5.179 |
| H | - | - | - | - | - | - | - | - | NA | NA | NA | NA | NA | NA |
| I | - | - | - | - | - | - | - | - | - | 4.536 | 1.813 | 4.602 | -0.431 | 3.784 |
| J | - | - | - | - | - | - | - | - | - | - | 5.044 | 2.86 | 5.267 | 5.815 |
| K | - | - | - | - | - | - | - | - | - | - | - | 4.25 | -0.368 | 4.152 |
| L | - | - | - | - | - | - | - | - | - | - | - | - | 4.801 | 5.544 |
| M | - | - | - | - | - | - | - | - | - | - | - | - | - | 3.688 |
| N | - | - | - | - | - | - | - | - | - | - | - | - | - | - |

| Max | A | B | C | D | E | F | G | H | I | J | K | L | M | N |
|---|---|---|---|---|---|---|---|---|---|---|---|---|---|---|
| A | - | 4.274 | -1.923 | 2.847 | 4.1 | NA | 2.837 | NA | 3.742 | 3.143 | 4.331 | 4.007 | 3.068 | 4.261 |
| B | - | - | 4.033 | 3.984 | 1.341 | NA | 3.527 | NA | 3.294 | 3.892 | 5.61 | 1.68 | 5.131 | 5.743 |
| C | - | - | - | -0.068 | 4.102 | NA | 2.874 | NA | 2.793 | 1.375 | 4.455 | 4.013 | 3.82 | 4.848 |
| D | - | - | - | - | 3.378 | NA | 1.659 | NA | 2.386 | -0.934 | 4.782 | 3.375 | 4.057 | 4.978 |
| E | - | - | - | - | - | NA | 3.41 | NA | 1.523 | 3.16 | 5.574 | 0.407 | 5.165 | 5.594 |
| F | - | - | - | - | - | - | NA | NA | NA | NA | NA | NA | NA | NA |
| G | - | - | - | - | - | - | - | NA | 1.517 | 0.957 | 5.173 | 3.123 | 4.427 | 5.197 |
| H | - | - | - | - | - | - | - | - | NA | NA | NA | NA | NA | NA |
| I | - | - | - | - | - | - | - | - | - | 2.548 | 5.239 | 2.58 | 4.649 | 5.415 |
| J | - | - | - | - | - | - | - | - | - | - | 4.917 | 3.447 | 3.039 | 4.673 |
| K | - | - | - | - | - | - | - | - | - | - | - | 4.958 | 2.616 | 2.061 |
| L | - | - | - | - | - | - | - | - | - | - | - | - | 5.175 | 5.663 |
| M | - | - | - | - | - | - | - | - | - | - | - | - | - | 3.895 |
| N | - | - | - | - | - | - | - | - | - | - | - | - | - | - |
