## Supplementary Data S2 for "Interlaboratory comparison of standardised metabolomics and lipidomics analyses in human and rodent blood using the MxP® Quant 500 kit": Cer(d16.1-22.0).pdf

**Amount of relative differences between laboratories**

| Min | A | B | C | D | E | F | G | H | I | J | K | L | M | N |
|---|---|---|---|---|---|---|---|---|---|---|---|---|---|---|
| A | - | -2.236 | 3.173 | 7.567 | 2.714 | 5.92 | 1.301 | 4.428 | 3.545 | 4.973 | 3.619 | 8.619 | -2.222 | 5.66 |
| B | - | - | 0.087 | 5.532 | 6.645 | 7.375 | 2.199 | 2.571 | 2.547 | 3.877 | 2.092 | 7.173 | 3.223 | 6.24 |
| C | - | - | - | 5.425 | 8.288 | 7.568 | 1.874 | 3.386 | 2.764 | 10.301 | 2.914 | 8.183 | 3.145 | 6.825 |
| D | - | - | - | - | 7.123 | 6.832 | 3.94 | 5.587 | 4.579 | 8.986 | 2.456 | 5.974 | 3.424 | 6.154 |
| E | - | - | - | - | - | 5.864 | 5.412 | 2.347 | 4.97 | 5.882 | 5.181 | 8.825 | 5.584 | 6.972 |
| F | - | - | - | - | - | - | 5.637 | 6.45 | 5.286 | 8.828 | 7.612 | 10.209 | 5.736 | 8.374 |
| G | - | - | - | - | - | - | - | 4.31 | 3.16 | 10.267 | 4.927 | 3.79 | 2.317 | 6.968 |
| H | - | - | - | - | - | - | - | - | 3.461 | 5.801 | 0.837 | 9.148 | 2.084 | 6.263 |
| I | - | - | - | - | - | - | - | - | - | 10.096 | 4.718 | 7.926 | 4.02 | 7.101 |
| J | - | - | - | - | - | - | - | - | - | - | 4.044 | 9.013 | 4.553 | 6.089 |
| K | - | - | - | - | - | - | - | - | - | - | - | 7.57 | 3.283 | 6.197 |
| L | - | - | - | - | - | - | - | - | - | - | - | - | 3.223 | 5.09 |
| M | - | - | - | - | - | - | - | - | - | - | - | - | - | 6.583 |
| N | - | - | - | - | - | - | - | - | - | - | - | - | - | - |

| | |
|---|---|
| | Larger than 10 |
| | Between 8 and 9,99 |
| | Between 6 and 7,99 |
| | Between 4 and 5,99 |
| | Between 2 and 3,99 |
| | Between 0 and 1,99 |
| | NA (Calculation failed) |

| Median | A | B | C | D | E | F | G | H | I | J | K | L | M | N |
|---|---|---|---|---|---|---|---|---|---|---|---|---|---|---|
| A | - | 4.212 | 4.084 | -2.652 | 4.835 | 4.732 | 3.465 | 4.162 | 4.488 | 3.171 | 4.454 | 3.7 | 3.49 | 3.302 |
| B | - | - | -0.97 | 4.468 | 3.674 | 3.37 | 3.222 | 1.143 | 2.262 | 3.553 | 1.822 | 2.75 | 3.152 | 3.49 |
| C | - | - | - | 4.434 | 4.047 | 3.417 | 3.15 | 1.637 | 2.378 | 3.25 | 1.735 | 1.994 | 3.076 | 3.747 |
| D | - | - | - | - | 4.863 | 4.688 | 3.38 | 4.15 | 4.46 | 2.923 | 4.335 | 3.743 | 3.431 | 2.776 |
| E | - | - | - | - | - | 1.504 | 4.682 | 3.881 | 3.265 | 4.821 | 3.426 | 4.443 | 4.672 | 4.794 |
| F | - | - | - | - | - | - | 4.473 | 4.014 | 2.739 | 4.57 | 4.062 | 4.395 | 4.471 | 5.13 |
| G | - | - | - | - | - | - | - | 2.183 | 3.688 | -0.838 | 3.282 | 1.863 | -1.588 | 2.419 |
| H | - | - | - | - | - | - | - | - | 2.962 | 3.441 | 2.529 | 2.558 | 2.45 | 3.136 |
| I | - | - | - | - | - | - | - | - | - | 3.966 | 1.358 | 3.302 | 3.847 | 4.331 |
| J | - | - | - | - | - | - | - | - | - | - | 3.796 | 2.264 | 0.276 | -3.525 |
| K | - | - | - | - | - | - | - | - | - | - | - | 3.454 | 3.671 | 3.943 |
| L | - | - | - | - | - | - | - | - | - | - | - | - | 1.765 | 1.843 |
| M | - | - | - | - | - | - | - | - | - | - | - | - | - | 2.193 |
| N | - | - | - | - | - | - | - | - | - | - | - | - | - | - |

| Max | A | B | C | D | E | F | G | H | I | J | K | L | M | N |
|---|---|---|---|---|---|---|---|---|---|---|---|---|---|---|
| A | - | 4.369 | 4.583 | 2.846 | 4.876 | 4.586 | 3.947 | 4.115 | 4.764 | 3.258 | 4.545 | 4.174 | 4.106 | 4.161 |
| B | - | - | -5.363 | 4.409 | 3.245 | 0.49 | 3.555 | 1.804 | 2.118 | 3.597 | 2.228 | 2.326 | 3.113 | 1.431 |
| C | - | - | - | 4.38 | 2.434 | 1.5 | 3.529 | 0.966 | 2.17 | 4.756 | 2.301 | 2.936 | 3.039 | -2.608 |
| D | - | - | - | - | 4.642 | 4.257 | 2.981 | 3.699 | 4.404 | -0.504 | 4.482 | 3.675 | 3.434 | 4.105 |
| E | - | - | - | - | - | 2.896 | 4.091 | 4.158 | -3.334 | 4.881 | 2.805 | 4.033 | 3.783 | 3.883 |
| F | - | - | - | - | - | - | 3.022 | 2.305 | 2.905 | 4.156 | 3.268 | 2.717 | 2.564 | 2.121 |
| G | - | - | - | - | - | - | - | 3.067 | 3.87 | 4.132 | 3.967 | 1.811 | 1.637 | 3.399 |
| H | - | - | - | - | - | - | - | - | 2.673 | 3.357 | 2.784 | 0.13 | 2.609 | -2.138 |
| I | - | - | - | - | - | - | - | - | - | 4.95 | -0.098 | 3.702 | 3.747 | 2.193 |
| J | - | - | - | - | - | - | - | - | - | - | 3.756 | 3.497 | 3.818 | 3.19 |
| K | - | - | - | - | - | - | - | - | - | - | - | 3.126 | 3.839 | 2.941 |
| L | - | - | - | - | - | - | - | - | - | - | - | - | -1.752 | -1.128 |
| M | - | - | - | - | - | - | - | - | - | - | - | - | - | 2.915 |
| N | - | - | - | - | - | - | - | - | - | - | - | - | - | - |
