## Supplementary Data S2 for "Interlaboratory comparison of standardised metabolomics and lipidomics analyses in human and rodent blood using the MxP® Quant 500 kit": Cer(d16.1-24.0).pdf

**Amount of relative differences between laboratories**

| Min | A | B | C | D | E | F | G | H | I | J | K | L | M | N |
|---|---|---|---|---|---|---|---|---|---|---|---|---|---|---|
| A | - | 4.551 | 4.117 | 4.299 | 5.965 | 2.149 | 6.263 | 3.161 | 5.259 | 6.478 | 4.417 | 6.856 | 3.305 | 7.153 |
| B | - | - | 4.79 | 4.141 | 5.967 | 3.753 | 6.916 | 4.266 | 5.994 | 5.447 | 3.449 | 7.393 | 4.378 | 7.607 |
| C | - | - | - | 5.448 | 6.685 | 3.348 | 1.91 | 2.553 | 2.005 | 8.309 | 5.864 | 5.151 | 2.132 | 7.663 |
| D | - | - | - | - | 6.112 | 2.509 | 6.036 | -0.4 | 5.512 | 6.918 | 3.715 | 6.61 | -1.131 | 7.338 |
| E | - | - | - | - | - | 4.279 | 8.304 | 4.574 | 7.672 | 5.372 | 7.639 | 8.63 | 4.587 | 8.649 |
| F | - | - | - | - | - | - | 7.636 | 2.051 | 6.774 | 6.852 | 6.665 | 8.112 | 2.193 | 8.61 |
| G | - | - | - | - | - | - | - | 1.805 | 3.744 | 6.876 | 6.263 | 4.523 | 0.894 | 6.131 |
| H | - | - | - | - | - | - | - | - | 6.136 | 9.087 | 7.609 | 7.212 | 1.333 | 6.528 |
| I | - | - | - | - | - | - | - | - | - | 6.839 | 6.439 | 5.815 | 0.061 | 7.034 |
| J | - | - | - | - | - | - | - | - | - | - | 5.251 | 7.894 | -0.136 | 7.749 |
| K | - | - | - | - | - | - | - | - | - | - | - | 7.26 | 4.308 | 7.535 |
| L | - | - | - | - | - | - | - | - | - | - | - | - | 3.137 | 5.904 |
| M | - | - | - | - | - | - | - | - | - | - | - | - | - | 7.154 |
| N | - | - | - | - | - | - | - | - | - | - | - | - | - | - |

| | |
|---|---|
| | Larger than 10 |
| | Between 8 and 9,99 |
| | Between 6 and 7,99 |
| | Between 4 and 5,99 |
| | Between 2 and 3,99 |
| | Between 0 and 1,99 |
| | NA (Calculation failed) |

| Median | A | B | C | D | E | F | G | H | I | J | K | L | M | N |
|---|---|---|---|---|---|---|---|---|---|---|---|---|---|---|
| A | - | 3.571 | 3.516 | 2.743 | 3.227 | 2.42 | 2.945 | 0.433 | 1.979 | 4.203 | 3.51 | 1.641 | 2.302 | 2.566 |
| B | - | - | 4.639 | 4.347 | 1.183 | 4.151 | 4.47 | 3.741 | 4.143 | 5.096 | -1.004 | 3.308 | 4.127 | 4.333 |
| C | - | - | - | 1.896 | 4.213 | 2.51 | 1.456 | 3.217 | 2.545 | 1.53 | 4.301 | 3.643 | 2.58 | 1.051 |
| D | - | - | - | - | 3.91 | 1.111 | 0.242 | 2.53 | 1.046 | 3.348 | 4.017 | 3.174 | 1.256 | -3.19 |
| E | - | - | - | - | - | 4.019 | 4.345 | 3.557 | 4.026 | 4.937 | -0.774 | 3.062 | 3.96 | 4.251 |
| F | - | - | - | - | - | - | 1.953 | 1.929 | -4.992 | 3.425 | 3.842 | 2.657 | -0.978 | 2.196 |
| G | - | - | - | - | - | - | - | 2.755 | 1.852 | 3.333 | 4.145 | 3.335 | 1.728 | 0.738 |
| H | - | - | - | - | - | - | - | - | 1.887 | 3.559 | 3.58 | 2.083 | 1.937 | 2.891 |
| I | - | - | - | - | - | - | - | - | - | 3.764 | 3.908 | 2.724 | -2.728 | 1.45 |
| J | - | - | - | - | - | - | - | - | - | - | 4.643 | 4.156 | 3.35 | 3.353 |
| K | - | - | - | - | - | - | - | - | - | - | - | 3.224 | 4.074 | 4.289 |
| L | - | - | - | - | - | - | - | - | - | - | - | - | 2.951 | 3.229 |
| M | - | - | - | - | - | - | - | - | - | - | - | - | - | 1.904 |
| N | - | - | - | - | - | - | - | - | - | - | - | - | - | - |

| Max | A | B | C | D | E | F | G | H | I | J | K | L | M | N |
|---|---|---|---|---|---|---|---|---|---|---|---|---|---|---|
| A | - | 3.49 | 2.792 | 2.609 | 2.374 | 2.676 | 2.429 | 2.863 | 0.974 | 4.574 | 3.465 | 2.639 | 1.126 | -0.422 |
| B | - | - | 4.511 | 4.356 | 2.883 | 4.5 | 4.237 | 2.747 | 3.908 | 5.252 | -2.861 | 2.529 | 3.666 | 3.757 |
| C | - | - | - | 0.93 | 3.573 | -0.716 | 1.567 | 3.722 | 2.679 | 4.244 | 4.2 | 3.781 | 3.048 | 3.299 |
| D | - | - | - | - | 3.429 | 0.886 | -0.038 | 3.52 | 2.267 | 4.067 | 4.105 | 3.562 | 2.694 | 2.591 |
| E | - | - | - | - | - | 3.654 | 3.381 | 0.932 | 2.452 | 4.911 | 2.91 | 0.838 | 1.769 | 2.18 |
| F | - | - | - | - | - | - | 1.411 | 3.682 | 2.649 | 4.094 | 4.198 | 3.776 | 2.964 | 3.332 |
| G | - | - | - | - | - | - | - | 3.385 | 1.608 | 4.042 | 3.968 | 3.453 | 2.433 | 2.134 |
| H | - | - | - | - | - | - | - | - | 2.994 | 5.31 | 2.488 | -1.079 | 2.514 | 2.07 |
| I | - | - | - | - | - | - | - | - | - | 4.323 | 3.646 | 3.044 | 0.9 | 1.098 |
| J | - | - | - | - | - | - | - | - | - | - | 4.764 | 4.615 | 4.557 | 4.23 |
| K | - | - | - | - | - | - | - | - | - | - | - | 2.482 | 3.653 | 3.729 |
| L | - | - | - | - | - | - | - | - | - | - | - | - | 2.633 | 2.899 |
| M | - | - | - | - | - | - | - | - | - | - | - | - | - | 0.605 |
| N | - | - | - | - | - | - | - | - | - | - | - | - | - | - |
