## Supplementary Data S2 for "Interlaboratory comparison of standardised metabolomics and lipidomics analyses in human and rodent blood using the MxP® Quant 500 kit": Cer(d18.1-25.0).pdf

**Amount of relative differences between laboratories**

| Min | A | B | C | D | E | F | G | H | I | J | K | L | M | N |
|---|---|---|---|---|---|---|---|---|---|---|---|---|---|---|
| A | - | NA | NA | 5.076 | 5.596 | NA | 5.236 | 6.068 | 6.423 | 3.689 | 5.958 | 5.629 | 5.935 | 5.248 |
| B | - | - | NA | NA | NA | NA | NA | NA | NA | NA | NA | NA | NA | NA |
| C | - | - | - | NA | NA | NA | NA | NA | NA | NA | NA | NA | NA | NA |
| D | - | - | - | - | 4.595 | NA | 2.108 | 5.557 | 5.886 | 5.438 | 5.306 | 4.24 | 5.356 | 4.184 |
| E | - | - | - | - | - | NA | 4.693 | 5.056 | 5.591 | 6.396 | 4.285 | 2.796 | 4.555 | -1.945 |
| F | - | - | - | - | - | - | NA | NA | NA | NA | NA | NA | NA | NA |
| G | - | - | - | - | - | - | - | 5.383 | 5.883 | 5.535 | 5.315 | 3.926 | 5.277 | 3.971 |
| H | - | - | - | - | - | - | - | - | 3.444 | 6.848 | 2.439 | 5.991 | 2.684 | 5.002 |
| I | - | - | - | - | - | - | - | - | - | 7.259 | 4.353 | 6.468 | 4.661 | 5.639 |
| J | - | - | - | - | - | - | - | - | - | - | 6.203 | 6.141 | 6.277 | 5.792 |
| K | - | - | - | - | - | - | - | - | - | - | - | 5.696 | -1.058 | 4.898 |
| L | - | - | - | - | - | - | - | - | - | - | - | - | 4.893 | 3.105 |
| M | - | - | - | - | - | - | - | - | - | - | - | - | - | 4.367 |
| N | - | - | - | - | - | - | - | - | - | - | - | - | - | - |

| | |
|---|---|
| | Larger than 10 |
| | Between 8 and 9,99 |
| | Between 6 and 7,99 |
| | Between 4 and 5,99 |
| | Between 2 and 3,99 |
| | Between 0 and 1,99 |
| | NA (Calculation failed) |

| Median | A | B | C | D | E | F | G | H | I | J | K | L | M | N |
|---|---|---|---|---|---|---|---|---|---|---|---|---|---|---|
| A | - | NA | NA | 1.306 | 4.613 | NA | 1.456 | 2.038 | 3.497 | 2.731 | 0.237 | 3.42 | 3.167 | 2.115 |
| B | - | - | NA | NA | NA | NA | NA | NA | NA | NA | NA | NA | NA | NA |
| C | - | - | - | NA | NA | NA | NA | NA | NA | NA | NA | NA | NA | NA |
| D | - | - | - | - | 4.506 | NA | -1.287 | 2.616 | 3.319 | 2.184 | -0.918 | 3.103 | 2.592 | 2.521 |
| E | - | - | - | - | - | NA | 4.853 | 5.226 | 4.049 | 4.559 | 4.835 | 4.193 | 4.435 | 5.221 |
| F | - | - | - | - | - | - | NA | NA | NA | NA | NA | NA | NA | NA |
| G | - | - | - | - | - | - | - | 2.682 | 3.261 | 2.055 | -4.22 | 3.039 | 2.476 | 2.596 |
| H | - | - | - | - | - | - | - | - | 3.964 | 3.235 | 3.147 | 3.7 | 3.068 | 1.296 |
| I | - | - | - | - | - | - | - | - | - | 2.834 | 3.026 | 1.544 | 3.057 | 3.981 |
| J | - | - | - | - | - | - | - | - | - | - | 2.773 | 2.141 | 1.743 | 3.701 |
| K | - | - | - | - | - | - | - | - | - | - | - | 2.475 | 0.265 | 2.728 |
| L | - | - | - | - | - | - | - | - | - | - | - | - | 1.614 | 3.967 |
| M | - | - | - | - | - | - | - | - | - | - | - | - | - | 3.128 |
| N | - | - | - | - | - | - | - | - | - | - | - | - | - | - |

| Max | A | B | C | D | E | F | G | H | I | J | K | L | M | N |
|---|---|---|---|---|---|---|---|---|---|---|---|---|---|---|
| A | - | NA | NA | 0.245 | 4.453 | NA | 0.516 | 4.736 | 0.788 | 3.195 | 3.269 | 2.536 | 2.685 | 3.966 |
| B | - | - | NA | NA | NA | NA | NA | NA | NA | NA | NA | NA | NA | NA |
| C | - | - | - | NA | NA | NA | NA | NA | NA | NA | NA | NA | NA | NA |
| D | - | - | - | - | 4.496 | NA | -4.156 | 4.522 | 0.473 | 3.422 | 2.677 | 2.867 | 2.099 | 3.599 |
| E | - | - | - | - | - | NA | 4.87 | 5.902 | 4.8 | 3.977 | 5.212 | 4.302 | 5.128 | 5.564 |
| F | - | - | - | - | - | - | NA | NA | NA | NA | NA | NA | NA | NA |
| G | - | - | - | - | - | - | - | 4.436 | -0.067 | 3.415 | 2.807 | 2.876 | 2.086 | 3.566 |
| H | - | - | - | - | - | - | - | - | 4.038 | 4.671 | 3.51 | 4.503 | 3.673 | 3.543 |
| I | - | - | - | - | - | - | - | - | - | 3.547 | 2.577 | 2.982 | 1.725 | 2.746 |
| J | - | - | - | - | - | - | - | - | - | - | 4.4 | 1.982 | 4.215 | 4.966 |
| K | - | - | - | - | - | - | - | - | - | - | - | 3.679 | 0.764 | -1.188 |
| L | - | - | - | - | - | - | - | - | - | - | - | - | 3.672 | 4.466 |
| M | - | - | - | - | - | - | - | - | - | - | - | - | - | 2.477 |
| N | - | - | - | - | - | - | - | - | - | - | - | - | - | - |
