## Supplementary Data S2 for "Interlaboratory comparison of standardised metabolomics and lipidomics analyses in human and rodent blood using the MxP® Quant 500 kit": Cer(d18.2-14.0).pdf

**Amount of relative differences between laboratories**

| Min | A | B | C | D | E | F | G | H | I | J | K | L | M | N |
|---|---|---|---|---|---|---|---|---|---|---|---|---|---|---|
| A | - | NA | NA | NA | NA | NA | NA | NA | NA | NA | NA | NA | NA | NA |
| B | - | - | 3.796 | NA | 4.789 | NA | NA | NA | NA | 4.335 | NA | 4.745 | NA | 5.401 |
| C | - | - | - | NA | 4.672 | NA | NA | NA | NA | 4.718 | NA | 4.31 | NA | 5.943 |
| D | - | - | - | - | NA | NA | NA | NA | NA | NA | NA | NA | NA | NA |
| E | - | - | - | - | - | NA | NA | NA | NA | 3.594 | NA | 2.191 | NA | 5.158 |
| F | - | - | - | - | - | - | NA | NA | NA | NA | NA | NA | NA | NA |
| G | - | - | - | - | - | - | - | NA | NA | NA | NA | NA | NA | NA |
| H | - | - | - | - | - | - | - | - | NA | NA | NA | NA | NA | NA |
| I | - | - | - | - | - | - | - | - | - | NA | NA | NA | NA | NA |
| J | - | - | - | - | - | - | - | - | - | - | NA | 2.73 | NA | 6.319 |
| K | - | - | - | - | - | - | - | - | - | - | - | NA | NA | NA |
| L | - | - | - | - | - | - | - | - | - | - | - | - | NA | 4.288 |
| M | - | - | - | - | - | - | - | - | - | - | - | - | - | NA |
| N | - | - | - | - | - | - | - | - | - | - | - | - | - | - |

| Median | A | B | C | D | E | F | G | H | I | J | K | L | M | N |
|---|---|---|---|---|---|---|---|---|---|---|---|---|---|---|
| A | - | NA | NA | NA | NA | NA | NA | NA | NA | NA | NA | NA | NA | NA |
| B | - | - | 3.053 | NA | 5.437 | NA | NA | NA | NA | 4.335 | NA | 5.435 | NA | 2.779 |
| C | - | - | - | NA | 5.212 | NA | NA | NA | NA | 4.718 | NA | 5.167 | NA | 4.198 |
| D | - | - | - | - | NA | NA | NA | NA | NA | NA | NA | NA | NA | NA |
| E | - | - | - | - | - | NA | NA | NA | NA | 3.594 | NA | 1.079 | NA | 5.736 |
| F | - | - | - | - | - | - | NA | NA | NA | NA | NA | NA | NA | NA |
| G | - | - | - | - | - | - | - | NA | NA | NA | NA | NA | NA | NA |
| H | - | - | - | - | - | - | - | - | NA | NA | NA | NA | NA | NA |
| I | - | - | - | - | - | - | - | - | - | NA | NA | NA | NA | NA |
| J | - | - | - | - | - | - | - | - | - | - | NA | 4.526 | NA | 5.278 |
| K | - | - | - | - | - | - | - | - | - | - | - | NA | NA | NA |
| L | - | - | - | - | - | - | - | - | - | - | - | - | NA | 5.524 |
| M | - | - | - | - | - | - | - | - | - | - | - | - | - | NA |
| N | - | - | - | - | - | - | - | - | - | - | - | - | - | - |

| Max | A | B | C | D | E | F | G | H | I | J | K | L | M | N |
|---|---|---|---|---|---|---|---|---|---|---|---|---|---|---|
| A | - | NA | NA | NA | NA | NA | NA | NA | NA | NA | NA | NA | NA | NA |
| B | - | - | 4.533 | NA | 5.909 | NA | NA | NA | NA | 5.122 | NA | 5.888 | NA | 5.333 |
| C | - | - | - | NA | 5.627 | NA | NA | NA | NA | 2.329 | NA | 5.688 | NA | 5.292 |
| D | - | - | - | - | NA | NA | NA | NA | NA | NA | NA | NA | NA | NA |
| E | - | - | - | - | - | NA | NA | NA | NA | 6.072 | NA | -1.878 | NA | 6.262 |
| F | - | - | - | - | - | - | NA | NA | NA | NA | NA | NA | NA | NA |
| G | - | - | - | - | - | - | - | NA | NA | NA | NA | NA | NA | NA |
| H | - | - | - | - | - | - | - | - | NA | NA | NA | NA | NA | NA |
| I | - | - | - | - | - | - | - | - | - | NA | NA | NA | NA | NA |
| J | - | - | - | - | - | - | - | - | - | - | NA | 5.285 | NA | 3.991 |
| K | - | - | - | - | - | - | - | - | - | - | - | NA | NA | NA |
| L | - | - | - | - | - | - | - | - | - | - | - | - | NA | 6.345 |
| M | - | - | - | - | - | - | - | - | - | - | - | - | - | NA |
| N | - | - | - | - | - | - | - | - | - | - | - | - | - | - |

| | |
|---|---|
| | Larger than 10 |
| | Between 8 and 9,99 |
| | Between 6 and 7,99 |
| | Between 4 and 5,99 |
| | Between 2 and 3,99 |
| | Between 0 and 1,99 |
| | NA (Calculation failed) |
