## Supplementary Data S2 for "Interlaboratory comparison of standardised metabolomics and lipidomics analyses in human and rodent blood using the MxP® Quant 500 kit": Cer(d18.2-16.0).pdf

**Amount of relative differences between laboratories**

| Min | A | B | C | D | E | F | G | H | I | J | K | L | M | N |
|---|---|---|---|---|---|---|---|---|---|---|---|---|---|---|
| A | - | 4.593 | 2.603 | 4.876 | 5.978 | 5.085 | 5.269 | 4.869 | 3.056 | 5.385 | 4.593 | 6.053 | 4.955 | 3.139 |
| B | - | - | 4.172 | 6.612 | 7.277 | 4.705 | 6.824 | 3.76 | 3.445 | 6.745 | 3.146 | 7.131 | 2.916 | 5.478 |
| C | - | - | - | 5.59 | 6.507 | 5.457 | 5.85 | 4.709 | 3.732 | 5.536 | 5.423 | 6.325 | 4.76 | 2.182 |
| D | - | - | - | - | 4.775 | 5.435 | 2.769 | 5.511 | 3.874 | 4.105 | 5.716 | 4.524 | 5.556 | 4.283 |
| E | - | - | - | - | - | 5.269 | 4.077 | 5.747 | 2.533 | 5.551 | 5.711 | 2.732 | 5.839 | 5.301 |
| F | - | - | - | - | - | - | 7.793 | 5.462 | 1.865 | 7.934 | 6.08 | 8.122 | 5.448 | 7.203 |
| G | - | - | - | - | - | - | - | 5.649 | 4.278 | 4.583 | 5.541 | 3.948 | 5.668 | 4.388 |
| H | - | - | - | - | - | - | - | - | 2.727 | 7.315 | 1.454 | 7.42 | 1.172 | 6.136 |
| I | - | - | - | - | - | - | - | - | - | 3.364 | 7.034 | 6.534 | 2.478 | 6.6 |
| J | - | - | - | - | - | - | - | - | - | - | 5.676 | 5.235 | 5.433 | 4.481 |
| K | - | - | - | - | - | - | - | - | - | - | - | 7.361 | -0.082 | 5.478 |
| L | - | - | - | - | - | - | - | - | - | - | - | - | 5.738 | 5.248 |
| M | - | - | - | - | - | - | - | - | - | - | - | - | - | 5.915 |
| N | - | - | - | - | - | - | - | - | - | - | - | - | - | - |

| | |
|---|---|
| | Larger than 10 |
| | Between 8 and 9,99 |
| | Between 6 and 7,99 |
| | Between 4 and 5,99 |
| | Between 2 and 3,99 |
| | Between 0 and 1,99 |
| | NA (Calculation failed) |

| Median | A | B | C | D | E | F | G | H | I | J | K | L | M | N |
|---|---|---|---|---|---|---|---|---|---|---|---|---|---|---|
| A | - | 3.153 | 3.323 | -0.718 | 3.803 | 3.362 | 3.017 | 2.734 | 3.463 | 2.893 | 2.106 | 1.764 | 3.729 | 3.589 |
| B | - | - | -0.801 | 3.32 | 2.529 | 1.171 | 4.222 | 1.416 | 2.695 | 4.133 | 3.819 | 2.607 | 2.231 | 4.508 |
| C | - | - | - | 3.585 | 2.073 | 0.858 | 4.349 | 1.929 | 2.321 | 4.3 | 4.112 | 2.92 | 1.784 | 4.635 |
| D | - | - | - | - | 3.865 | 3.447 | 2.847 | 2.973 | 3.635 | 2.792 | 2.209 | 2.086 | 3.816 | 3.498 |
| E | - | - | - | - | - | 2.187 | 4.661 | 2.859 | 1.092 | 4.661 | 4.502 | 3.57 | -1.772 | 4.923 |
| F | - | - | - | - | - | - | 4.493 | 3.05 | 1.585 | 4.329 | 3.924 | 3.282 | -0.097 | 4.699 |
| G | - | - | - | - | - | - | - | 3.813 | 4.167 | -2.061 | 1.476 | 3.415 | 4.311 | 1.929 |
| H | - | - | - | - | - | - | - | - | 3.015 | 3.766 | 3.327 | 1.549 | 2.905 | 4.235 |
| I | - | - | - | - | - | - | - | - | - | 4.365 | 4.101 | 3.325 | -3.193 | 4.508 |
| J | - | - | - | - | - | - | - | - | - | - | 1.141 | 3.362 | 4.284 | 2.077 |
| K | - | - | - | - | - | - | - | - | - | - | - | 2.629 | 3.969 | 3.061 |
| L | - | - | - | - | - | - | - | - | - | - | - | - | 3.366 | 4.038 |
| M | - | - | - | - | - | - | - | - | - | - | - | - | - | 4.844 |
| N | - | - | - | - | - | - | - | - | - | - | - | - | - | - |

| Max | A | B | C | D | E | F | G | H | I | J | K | L | M | N |
|---|---|---|---|---|---|---|---|---|---|---|---|---|---|---|
| A | - | 2.696 | 3.446 | 0.211 | 4.122 | 1.334 | 2.558 | -1.117 | 3.77 | 2.512 | 2.973 | 2.858 | 3.306 | 3.618 |
| B | - | - | 2.252 | 2.2 | 3.698 | 2.476 | 3.563 | 2.831 | 1.145 | 3.705 | 4.085 | 0.396 | 2.054 | 4.396 |
| C | - | - | - | 3.29 | 3.082 | 3.585 | 4.143 | 3.588 | 1.589 | 4.199 | 4.631 | 1.829 | 0.066 | 4.725 |
| D | - | - | - | - | 4.04 | -1.714 | 2.852 | 1.323 | 3.358 | 3.025 | 3.555 | 2.524 | 2.969 | 3.776 |
| E | - | - | - | - | - | 4.224 | 4.825 | 4.583 | 3.29 | 4.854 | 5 | 3.689 | 3.488 | 5.161 |
| F | - | - | - | - | - | - | 3.018 | 0.617 | 3.235 | 3.143 | 3.111 | 2.359 | 3.233 | 4.045 |
| G | - | - | - | - | - | - | - | 2.056 | 4.051 | -0.071 | 1.419 | 3.598 | 3.805 | 2.52 |
| H | - | - | - | - | - | - | - | - | 3.247 | 2.654 | 3.452 | 2.673 | 3.224 | 3.889 |
| I | - | - | - | - | - | - | - | - | - | 4.47 | 5.155 | 2.447 | -0.108 | 5.004 |
| J | - | - | - | - | - | - | - | - | - | - | 1.961 | 3.693 | 3.905 | 2.653 |
| K | - | - | - | - | - | - | - | - | - | - | - | 3.964 | 4.206 | 2.465 |
| L | - | - | - | - | - | - | - | - | - | - | - | - | 1.091 | 4.37 |
| M | - | - | - | - | - | - | - | - | - | - | - | - | - | 4.714 |
| N | - | - | - | - | - | - | - | - | - | - | - | - | - | - |
