## Supplementary Data S2 for "Interlaboratory comparison of standardised metabolomics and lipidomics analyses in human and rodent blood using the MxP® Quant 500 kit": Cer(d18.2-18.0).pdf

**Amount of relative differences between laboratories**

| Min | A | B | C | D | E | F | G | H | I | J | K | L | M | N |
|---|---|---|---|---|---|---|---|---|---|---|---|---|---|---|
| A | - | 3.899 | 6.629 | 6.375 | 6.084 | NA | 6.576 | 3.205 | 4.949 | 4.846 | 6.585 | 7.108 | 4.812 | 5.209 |
| B | - | - | 7.062 | 6.279 | 7.438 | NA | 7.829 | 4.417 | 4.632 | 3.526 | 7.042 | 7.97 | 5.898 | 5.871 |
| C | - | - | - | 4.296 | 2.719 | NA | 3.612 | 6.033 | 3.962 | 5.504 | 3.566 | 5.192 | 5.228 | 4.959 |
| D | - | - | - | - | 7.885 | NA | 8.344 | 6.271 | 4.871 | 5.889 | 4.492 | 7.274 | 6.565 | 5.363 |
| E | - | - | - | - | - | NA | 3.905 | 5.764 | 2.77 | -0.074 | 3.806 | 5.393 | 5.117 | 3.411 |
| F | - | - | - | - | - | - | NA | NA | NA | NA | NA | NA | NA | NA |
| G | - | - | - | - | - | - | - | 6.056 | 4.229 | 4.441 | 4.697 | 4.225 | 5.469 | 5.071 |
| H | - | - | - | - | - | - | - | - | 2.709 | 4.43 | 7.535 | 8.459 | 6.566 | 6.608 |
| I | - | - | - | - | - | - | - | - | - | 4.86 | 5.515 | 6.13 | 5.151 | 2.924 |
| J | - | - | - | - | - | - | - | - | - | - | 6.922 | 7.774 | 6.091 | 6.19 |
| K | - | - | - | - | - | - | - | - | - | - | - | 5.765 | 4.399 | 3.793 |
| L | - | - | - | - | - | - | - | - | - | - | - | - | 5.632 | 4.899 |
| M | - | - | - | - | - | - | - | - | - | - | - | - | - | -1.533 |
| N | - | - | - | - | - | - | - | - | - | - | - | - | - | - |

| | |
|---|---|
| | Larger than 10 |
| | Between 8 and 9,99 |
| | Between 6 and 7,99 |
| | Between 4 and 5,99 |
| | Between 2 and 3,99 |
| | Between 0 and 1,99 |
| | NA (Calculation failed) |

| Median | A | B | C | D | E | F | G | H | I | J | K | L | M | N |
|---|---|---|---|---|---|---|---|---|---|---|---|---|---|---|
| A | - | 2.42 | 5.359 | 6.095 | 4.609 | NA | 6.021 | 5.005 | 4.342 | 5.35 | 6.155 | 5.064 | 5 | 4.955 |
| B | - | - | 5.147 | 5.904 | 4.251 | NA | 5.838 | 4.678 | 3.788 | 5.082 | 5.995 | 4.807 | 4.737 | 4.6 |
| C | - | - | - | 4.233 | 3.412 | NA | 4.25 | 2.279 | 3.803 | 1.5 | 4.429 | 2.017 | 2.489 | 2.44 |
| D | - | - | - | - | 4.763 | NA | 0.6 | 4.322 | 4.781 | 3.705 | 1.22 | 4.39 | 4.483 | 4.33 |
| E | - | - | - | - | - | NA | 5.044 | 2.621 | 1.5 | 3.802 | 5.157 | 2.837 | 2.418 | 2.568 |
| F | - | - | - | - | - | - | NA | NA | NA | NA | NA | NA | NA | NA |
| G | - | - | - | - | - | - | - | 4.322 | 4.749 | 3.847 | 1.178 | 4.273 | 4.369 | 4.327 |
| H | - | - | - | - | - | - | - | - | 3.106 | 2.501 | 5.113 | 1.905 | 1.894 | -4.373 |
| I | - | - | - | - | - | - | - | - | - | 4.254 | 5.404 | 3.283 | 3.134 | 3.325 |
| J | - | - | - | - | - | - | - | - | - | - | 4.655 | 1.898 | 1.679 | 2.592 |
| K | - | - | - | - | - | - | - | - | - | - | - | 4.419 | 4.503 | 4.565 |
| L | - | - | - | - | - | - | - | - | - | - | - | - | 0.946 | -0.231 |
| M | - | - | - | - | - | - | - | - | - | - | - | - | - | -0.768 |
| N | - | - | - | - | - | - | - | - | - | - | - | - | - | - |

| Max | A | B | C | D | E | F | G | H | I | J | K | L | M | N |
|---|---|---|---|---|---|---|---|---|---|---|---|---|---|---|
| A | - | 2.82 | 5.117 | 5.776 | 4.507 | NA | 5.995 | 5.358 | 3.75 | 5.438 | 6.098 | 4.894 | 5.022 | 4.918 |
| B | - | - | 4.627 | 5.441 | 3.711 | NA | 5.669 | 5.159 | 2.753 | 5.27 | 5.813 | 4.328 | 4.49 | 4.302 |
| C | - | - | - | 4.173 | 3.499 | NA | 4.269 | 3.039 | 3.689 | 3.7 | 4.659 | 2.456 | 0.634 | 1.025 |
| D | - | - | - | - | 4.255 | NA | 3.395 | 2.937 | 4.72 | 1.717 | 2.349 | 4.02 | 3.717 | 4.115 |
| E | - | - | - | - | - | NA | 5.073 | 4.262 | 3.176 | 4.061 | 5.365 | 2.563 | 3.385 | 3.021 |
| F | - | - | - | - | - | - | NA | NA | NA | NA | NA | NA | NA | NA |
| G | - | - | - | - | - | - | - | 3.318 | 5.006 | 3.683 | 2.434 | 4.37 | 4.141 | 4.193 |
| H | - | - | - | - | - | - | - | - | 4.199 | 1.207 | 4.155 | 3.106 | 2.98 | 3.681 |
| I | - | - | - | - | - | - | - | - | - | 4.87 | 5.391 | 3.651 | 3.851 | 3.371 |
| J | - | - | - | - | - | - | - | - | - | - | 3.858 | 3.539 | 3.62 | 4.041 |
| K | - | - | - | - | - | - | - | - | - | - | - | 4.59 | 4.516 | 4.642 |
| L | - | - | - | - | - | - | - | - | - | - | - | - | 2.053 | 1.573 |
| M | - | - | - | - | - | - | - | - | - | - | - | - | - | -0.691 |
| N | - | - | - | - | - | - | - | - | - | - | - | - | - | - |
