## Supplementary Data S2 for "Interlaboratory comparison of standardised metabolomics and lipidomics analyses in human and rodent blood using the MxP® Quant 500 kit": Cer(d18.2-20.0).pdf

**Amount of relative differences between laboratories**

| Min | A | B | C | D | E | F | G | H | I | J | K | L | M | N |
|---|---|---|---|---|---|---|---|---|---|---|---|---|---|---|
| A | - | NA | 3.835 | 5.233 | 3.168 | NA | 4.977 | 3.21 | 5.392 | 3.314 | 9.074 | 6.219 | 3.237 | 4.538 |
| B | - | - | NA | NA | NA | NA | NA | NA | NA | NA | NA | NA | NA | NA |
| C | - | - | - | 6.093 | 2.869 | NA | 5.16 | 2.872 | 5.135 | 5.74 | 13.762 | 5.307 | 3.826 | 3.474 |
| D | - | - | - | - | 5.401 | NA | 5.215 | 6.618 | 5.995 | 6.914 | 10.384 | 3.685 | 6.234 | 6.587 |
| E | - | - | - | - | - | NA | 5.383 | 3.74 | 4.731 | 5.183 | 8.7 | 5.25 | 4.453 | 3.482 |
| F | - | - | - | - | - | - | NA | NA | NA | NA | NA | NA | NA | NA |
| G | - | - | - | - | - | - | - | 3.632 | 5.522 | 6.461 | 11.486 | 4.688 | 5.029 | 5.201 |
| H | - | - | - | - | - | - | - | - | 5.273 | 7.69 | 9.705 | 6.536 | 5.33 | 5.32 |
| I | - | - | - | - | - | - | - | - | - | 6.55 | 12.079 | 5.061 | 4.185 | 4.304 |
| J | - | - | - | - | - | - | - | - | - | - | 7.088 | 5.979 | 1.638 | 4.126 |
| K | - | - | - | - | - | - | - | - | - | - | - | 7.757 | 6.237 | 6.945 |
| L | - | - | - | - | - | - | - | - | - | - | - | - | 5.05 | 4.662 |
| M | - | - | - | - | - | - | - | - | - | - | - | - | - | 0.922 |
| N | - | - | - | - | - | - | - | - | - | - | - | - | - | - |

| | |
|---|---|
| | Larger than 10 |
| | Between 8 and 9,99 |
| | Between 6 and 7,99 |
| | Between 4 and 5,99 |
| | Between 2 and 3,99 |
| | Between 0 and 1,99 |
| | NA (Calculation failed) |

| Median | A | B | C | D | E | F | G | H | I | J | K | L | M | N |
|---|---|---|---|---|---|---|---|---|---|---|---|---|---|---|
| A | - | NA | 2.983 | 3.228 | 4.223 | NA | 2.101 | 1.822 | 5.592 | 2.388 | 2.66 | 4.349 | 4.932 | 4.835 |
| B | - | - | NA | NA | NA | NA | NA | NA | NA | NA | NA | NA | NA | NA |
| C | - | - | - | 4.423 | 3.693 | NA | 1.976 | 1.161 | 5.45 | -0.3 | 8.027 | 3.918 | 4.677 | 4.492 |
| D | - | - | - | - | 4.512 | NA | 3.235 | 5.269 | 5.68 | 2.811 | -0.001 | 4.627 | 5.132 | 4.915 |
| E | - | - | - | - | - | NA | 4.298 | 2.678 | 5.135 | 4.141 | 3.481 | 1.138 | 3.928 | 3.366 |
| F | - | - | - | - | - | - | NA | NA | NA | NA | NA | NA | NA | NA |
| G | - | - | - | - | - | - | - | 2.406 | 5.522 | 1.397 | 5.999 | 4.245 | 4.845 | 4.624 |
| H | - | - | - | - | - | - | - | - | 5.443 | 1.703 | 4.516 | 3.735 | 4.792 | 4.527 |
| I | - | - | - | - | - | - | - | - | - | 6.325 | 7.128 | 5.639 | 4.98 | 5.296 |
| J | - | - | - | - | - | - | - | - | - | - | 2.937 | 3.976 | 4.73 | 4.569 |
| K | - | - | - | - | - | - | - | - | - | - | - | 2.896 | 4.187 | 4.183 |
| L | - | - | - | - | - | - | - | - | - | - | - | - | 3.764 | 2.816 |
| M | - | - | - | - | - | - | - | - | - | - | - | - | - | 2.377 |
| N | - | - | - | - | - | - | - | - | - | - | - | - | - | - |

| Max | A | B | C | D | E | F | G | H | I | J | K | L | M | N |
|---|---|---|---|---|---|---|---|---|---|---|---|---|---|---|
| A | - | NA | 2.658 | 3.209 | 4.593 | NA | 3.349 | 4.827 | 5.663 | 2.205 | 6.025 | 4.885 | 5.099 | 4.882 |
| B | - | - | NA | NA | NA | NA | NA | NA | NA | NA | NA | NA | NA | NA |
| C | - | - | - | 3.231 | 4.081 | NA | 0.388 | 2.933 | 5.555 | 2.995 | 10.616 | 4.35 | 4.787 | 4.617 |
| D | - | - | - | - | 4.247 | NA | 2.278 | 6.274 | 5.537 | 2.994 | 7.495 | 4.692 | 4.801 | 4.197 |
| E | - | - | - | - | - | NA | 4.004 | 5.485 | 5.265 | 4.63 | 6.098 | 2.606 | 3.81 | 3.344 |
| F | - | - | - | - | - | - | NA | NA | NA | NA | NA | NA | NA | NA |
| G | - | - | - | - | - | - | - | 3.086 | 5.521 | 3.111 | 8.288 | 4.194 | 4.81 | 4.49 |
| H | - | - | - | - | - | - | - | - | 5.504 | 4.558 | 6.428 | 4.558 | 4.671 | 4.325 |
| I | - | - | - | - | - | - | - | - | - | 6.688 | 8.721 | 5.684 | 5.085 | 5.42 |
| J | - | - | - | - | - | - | - | - | - | - | 3.258 | 4.545 | 4.942 | 4.959 |
| K | - | - | - | - | - | - | - | - | - | - | - | 4.86 | 5.128 | 5.443 |
| L | - | - | - | - | - | - | - | - | - | - | - | - | 3.335 | 1.913 |
| M | - | - | - | - | - | - | - | - | - | - | - | - | - | 2.533 |
| N | - | - | - | - | - | - | - | - | - | - | - | - | - | - |
