## Supplementary Data S2 for "Interlaboratory comparison of standardised metabolomics and lipidomics analyses in human and rodent blood using the MxP® Quant 500 kit": Cer(d18.2-22.0).pdf

**Amount of relative differences between laboratories**

| Min | A | B | C | D | E | F | G | H | I | J | K | L | M | N |
|---|---|---|---|---|---|---|---|---|---|---|---|---|---|---|
| A | - | 4.342 | 0.42 | 5.049 | 2.415 | 5 | 5.773 | 4.273 | 3.5 | 4.69 | 5.067 | 4.878 | 2.824 | 4.549 |
| B | - | - | 4.784 | 6.067 | 5.298 | 3.59 | 6.675 | 0.853 | 5.525 | 6.555 | 3.894 | 5.993 | 5.055 | 5.869 |
| C | - | - | - | 5.1 | 3.172 | 5.165 | 5.79 | 4.085 | 4.093 | 5.255 | 5.285 | 4.956 | 3.344 | 4.529 |
| D | - | - | - | - | 4.474 | 5.773 | 3.584 | 5.243 | 3.666 | 2.357 | 6.003 | 0.882 | 4.037 | 3.562 |
| E | - | - | - | - | - | 5.407 | 5.499 | 4.462 | 3.133 | 3.622 | 5.522 | 4.717 | 2.282 | 3.594 |
| F | - | - | - | - | - | - | 6.996 | 4.184 | 5.996 | 6.831 | 2.827 | 6.469 | 5.805 | 6.327 |
| G | - | - | - | - | - | - | - | 5.419 | 4.42 | 4.507 | 6.137 | 3.35 | 4.521 | 4.386 |
| H | - | - | - | - | - | - | - | - | 5.343 | 6.129 | 3.866 | 5.875 | 4.954 | 5.743 |
| I | - | - | - | - | - | - | - | - | - | 3.38 | 6.142 | 3.84 | 2.368 | 0.015 |
| J | - | - | - | - | - | - | - | - | - | - | 6.073 | 2.767 | 4.017 | 3.111 |
| K | - | - | - | - | - | - | - | - | - | - | - | 5.894 | 4.845 | 5.971 |
| L | - | - | - | - | - | - | - | - | - | - | - | - | 3.867 | 3.49 |
| M | - | - | - | - | - | - | - | - | - | - | - | - | - | 3.047 |
| N | - | - | - | - | - | - | - | - | - | - | - | - | - | - |

| | |
|---|---|
| | Larger than 10 |
| | Between 8 and 9,99 |
| | Between 6 and 7,99 |
| | Between 4 and 5,99 |
| | Between 2 and 3,99 |
| | Between 0 and 1,99 |
| | NA (Calculation failed) |

| Median | A | B | C | D | E | F | G | H | I | J | K | L | M | N |
|---|---|---|---|---|---|---|---|---|---|---|---|---|---|---|
| A | - | 3.848 | 3.551 | 2.088 | 3.801 | 4.837 | 2.44 | 3.891 | 3.035 | 2.696 | -0.104 | -0.185 | 3.113 | 3.689 |
| B | - | - | 1.346 | 4.403 | -2.482 | 4.083 | 4.478 | -0.006 | 2.74 | 4.559 | 4.113 | 4.107 | 2.636 | 0.484 |
| C | - | - | - | 4.175 | 1.3 | 4.262 | 4.291 | 1.843 | 1.972 | 4.358 | 3.841 | 3.833 | 1.759 | 0.633 |
| D | - | - | - | - | 4.107 | 4.983 | 0.549 | 4.199 | 3.585 | 1.071 | 1.635 | 1.694 | 3.643 | 4.07 |
| E | - | - | - | - | - | 4.113 | 4.496 | 0.374 | 2.681 | 4.554 | 4.095 | 4.085 | 2.541 | -0.603 |
| F | - | - | - | - | - | - | 5.569 | 4.265 | 4.833 | 5.611 | 5.369 | 5.365 | 4.802 | 4.46 |
| G | - | - | - | - | - | - | - | 4.296 | 3.739 | -0.814 | 2.198 | 2.24 | 3.794 | 4.183 |
| H | - | - | - | - | - | - | - | - | 2.958 | 4.64 | 4.207 | 4.204 | 2.863 | 1.239 |
| I | - | - | - | - | - | - | - | - | - | 4.001 | 3.351 | 3.314 | -0.834 | 2.445 |
| J | - | - | - | - | - | - | - | - | - | - | 2.334 | 2.394 | 3.835 | 4.205 |
| K | - | - | - | - | - | - | - | - | - | - | - | -4.317 | 3.211 | 3.718 |
| L | - | - | - | - | - | - | - | - | - | - | - | - | 3.259 | 3.812 |
| M | - | - | - | - | - | - | - | - | - | - | - | - | - | 2.281 |
| N | - | - | - | - | - | - | - | - | - | - | - | - | - | - |

| Max | A | B | C | D | E | F | G | H | I | J | K | L | M | N |
|---|---|---|---|---|---|---|---|---|---|---|---|---|---|---|
| A | - | 3.672 | 3.713 | -0.248 | 3.993 | 4.818 | 0.338 | 3.796 | 3.431 | 2.053 | 2.6 | 1.496 | 3.507 | 4.07 |
| B | - | - | -0.288 | 3.976 | 2.086 | 4.128 | 3.957 | 0.66 | 0.541 | 3.916 | 4.466 | 3.486 | 0.634 | 2.245 |
| C | - | - | - | 4.013 | 1.865 | 4.115 | 4.049 | -0.323 | 1.124 | 4.2 | 4.478 | 3.594 | 1.148 | 2.071 |
| D | - | - | - | - | 4.064 | 4.86 | -0.802 | 3.839 | 3.574 | 1.57 | 2.581 | 1.772 | 3.563 | 4.116 |
| E | - | - | - | - | - | 3.857 | 4.368 | 1.597 | 2.608 | 4.634 | 4.727 | 3.979 | 2.581 | -0.813 |
| F | - | - | - | - | - | - | 5.346 | 4.281 | 4.574 | 5.36 | 5.601 | 5.132 | 4.527 | 3.977 |
| G | - | - | - | - | - | - | - | 3.89 | 3.619 | 1.755 | 2.381 | 2.005 | 3.621 | 4.16 |
| H | - | - | - | - | - | - | - | - | 1.772 | 4.287 | 4.548 | 3.724 | 1.669 | 1.654 |
| I | - | - | - | - | - | - | - | - | - | 4.06 | 4.475 | 3.229 | -2.098 | 2.618 |
| J | - | - | - | - | - | - | - | - | - | - | 2.082 | 2.77 | 3.8 | 4.285 |
| K | - | - | - | - | - | - | - | - | - | - | - | 2.936 | 3.984 | 4.414 |
| L | - | - | - | - | - | - | - | - | - | - | - | - | 3.123 | 3.842 |
| M | - | - | - | - | - | - | - | - | - | - | - | - | - | 2.649 |
| N | - | - | - | - | - | - | - | - | - | - | - | - | - | - |
