## Supplementary Data S2 for "Interlaboratory comparison of standardised metabolomics and lipidomics analyses in human and rodent blood using the MxP® Quant 500 kit": Cer(d18.2-24.1).pdf

**Amount of relative differences between laboratories**

| Min | A | B | C | D | E | F | G | H | I | J | K | L | M | N |
|---|---|---|---|---|---|---|---|---|---|---|---|---|---|---|
| A | - | 5.412 | 6.056 | 0.589 | 3.024 | 5.894 | 0.564 | 5.525 | 2.804 | 5.401 | 5.061 | 3.994 | 4.196 | 3.567 |
| B | - | - | 4.868 | 5.768 | 6.115 | 5.06 | 5.789 | 4.347 | 4.964 | 7.093 | 4.912 | 5.013 | 3.429 | 5.276 |
| C | - | - | - | 6.456 | 6.645 | 3.974 | 6.543 | 1.423 | 5.683 | 7.474 | 6.065 | 6.168 | 4.758 | 6.304 |
| D | - | - | - | - | 1.297 | 5.784 | -1.911 | 5.559 | 3.378 | 5.32 | 3.984 | 4.209 | 4.555 | 2.997 |
| E | - | - | - | - | - | 5.986 | 2.769 | 6.043 | 5.287 | 5.601 | 5.593 | 4.607 | 4.985 | 4.563 |
| F | - | - | - | - | - | - | 6.943 | 4.036 | 6.355 | 7.788 | 6.645 | 6.554 | 5.643 | 6.787 |
| G | - | - | - | - | - | - | - | 5.584 | 3.719 | 4.869 | 3.575 | 4.15 | 4.737 | 2.97 |
| H | - | - | - | - | - | - | - | - | 5.899 | 7.186 | 6.117 | 6.194 | 5.038 | 6.315 |
| I | - | - | - | - | - | - | - | - | - | 5.399 | 2.902 | 3.575 | 3.862 | 2.914 |
| J | - | - | - | - | - | - | - | - | - | - | 6.462 | 5.277 | 4.821 | 5.582 |
| K | - | - | - | - | - | - | - | - | - | - | - | 1.408 | 4.066 | 2.141 |
| L | - | - | - | - | - | - | - | - | - | - | - | - | 3.927 | 3.14 |
| M | - | - | - | - | - | - | - | - | - | - | - | - | - | 4.577 |
| N | - | - | - | - | - | - | - | - | - | - | - | - | - | - |

| Median | A | B | C | D | E | F | G | H | I | J | K | L | M | N |
|---|---|---|---|---|---|---|---|---|---|---|---|---|---|---|
| A | - | 4.296 | 2.916 | -0.682 | 4.183 | 4.628 | 2.926 | 4.301 | 4.496 | 3.756 | 1.532 | 1.61 | 4.252 | 0.079 |
| B | - | - | 3.95 | 4.554 | 0.825 | 2.651 | 5.091 | -1.639 | 1.648 | 5.36 | 4.353 | 4.83 | 0.159 | 4.698 |
| C | - | - | - | 2.519 | 3.601 | 4.278 | 3.926 | 3.737 | 4.01 | 4.42 | 1.776 | 3.375 | 3.722 | 3.045 |
| D | - | - | - | - | 4.146 | 4.609 | 3.063 | 4.259 | 4.46 | 3.84 | 1.447 | 1.931 | 4.192 | 0.711 |
| E | - | - | - | - | - | 2.943 | 5.004 | 0.332 | 2.281 | 5.284 | 4.268 | 4.723 | -4.532 | 4.588 |
| F | - | - | - | - | - | - | 5.43 | 2.897 | 1.99 | 5.668 | 4.822 | 5.217 | 2.967 | 5.103 |
| G | - | - | - | - | - | - | - | 4.662 | 4.804 | 2.517 | 3.369 | 2.065 | 4.617 | 2.63 |
| H | - | - | - | - | - | - | - | - | 1.729 | 5.364 | 4.292 | 4.787 | -1.307 | 4.654 |
| I | - | - | - | - | - | - | - | - | - | 5.556 | 4.638 | 5.056 | 2.348 | 4.94 |
| J | - | - | - | - | - | - | - | - | - | - | 3.766 | 3.224 | 4.861 | 3.432 |
| K | - | - | - | - | - | - | - | - | - | - | - | 2.756 | 3.988 | 2.21 |
| L | - | - | - | - | - | - | - | - | - | - | - | - | 4.412 | 1.029 |
| M | - | - | - | - | - | - | - | - | - | - | - | - | - | 4.61 |
| N | - | - | - | - | - | - | - | - | - | - | - | - | - | - |

| Max | A | B | C | D | E | F | G | H | I | J | K | L | M | N |
|---|---|---|---|---|---|---|---|---|---|---|---|---|---|---|
| A | - | 4.183 | 1.079 | -0.758 | 4.227 | 4.478 | 2.973 | 4.117 | 4.567 | 3.595 | 0.594 | 1.938 | 4.255 | 0.567 |
| B | - | - | 4.29 | 4.483 | 1.001 | 2.13 | 5.053 | 1.126 | 2.493 | 5.183 | 4.321 | 4.822 | -0.837 | 4.674 |
| C | - | - | - | 1.454 | 4.202 | 4.297 | 3.579 | 3.837 | 4.393 | 3.769 | -0.877 | 2.972 | 3.955 | 2.566 |
| D | - | - | - | - | 4.214 | 4.475 | 3.12 | 4.056 | 4.514 | 3.706 | 1.092 | 2.242 | 4.168 | 0.967 |
| E | - | - | - | - | - | 1.914 | 5.075 | 2.192 | 1.249 | 5.267 | 4.481 | 4.833 | 0.903 | 4.679 |
| F | - | - | - | - | - | - | 5.315 | 2.731 | 0.751 | 5.405 | 4.645 | 5.124 | 2.431 | 4.993 |
| G | - | - | - | - | - | - | - | 4.541 | 4.858 | 2.187 | 3.359 | 1.859 | 4.61 | 2.617 |
| H | - | - | - | - | - | - | - | - | 2.98 | 5.169 | 4.116 | 4.686 | 0.551 | 4.548 |
| I | - | - | - | - | - | - | - | - | - | 5.564 | 4.72 | 5.135 | 2.638 | 4.998 |
| J | - | - | - | - | - | - | - | - | - | - | 3.353 | 3.024 | 4.863 | 3.258 |
| K | - | - | - | - | - | - | - | - | - | - | - | 2.836 | 3.983 | 2.212 |
| L | - | - | - | - | - | - | - | - | - | - | - | - | 4.435 | 1.262 |
| M | - | - | - | - | - | - | - | - | - | - | - | - | - | 4.611 |
| N | - | - | - | - | - | - | - | - | - | - | - | - | - | - |

| | |
|---|---|
| | Larger than 10 |
| | Between 8 and 9,99 |
| | Between 6 and 7,99 |
| | Between 4 and 5,99 |
| | Between 2 and 3,99 |
| | Between 0 and 1,99 |
| | NA (Calculation failed) |
