## Supplementary Data S2 for "Interlaboratory comparison of standardised metabolomics and lipidomics analyses in human and rodent blood using the MxP® Quant 500 kit": Cit.pdf

**Amount of relative differences between laboratories**

| Min | A | B | C | D | E | F | G | H | I | J | K | L | M | N |
|---|---|---|---|---|---|---|---|---|---|---|---|---|---|---|
| A | - | 3.887 | 0.77 | 1.732 | 3.675 | 3.809 | 0.674 | 3.85 | 3.142 | 2.865 | 2.145 | 2.542 | 2.956 | 3.495 |
| B | - | - | 4.231 | 3.728 | 1.207 | 0.789 | 4.181 | -0.24 | 2.583 | 3.193 | 3.611 | 3.175 | 2.996 | 2.246 |
| C | - | - | - | 2.413 | 3.87 | 4.002 | -5.034 | 4.033 | 3.539 | 3.187 | 2.648 | 2.852 | 3.249 | 3.745 |
| D | - | - | - | - | 3.283 | 3.407 | 2.233 | 3.481 | 2.613 | 1.922 | 0.03 | 1.665 | 2.205 | 2.973 |
| E | - | - | - | - | - | -0.14 | 3.972 | 0.639 | 1.892 | 2.687 | 3.258 | 2.785 | 2.472 | 1.155 |
| F | - | - | - | - | - | - | 4.012 | -0.919 | 1.399 | 2.872 | 3.37 | 2.807 | 2.648 | 1.713 |
| G | - | - | - | - | - | - | - | 3.887 | 3.388 | 2.999 | 2.401 | 2.938 | 3.126 | 3.522 |
| H | - | - | - | - | - | - | - | - | 2.222 | 2.97 | 3.445 | 2.915 | 2.733 | 1.947 |
| I | - | - | - | - | - | - | - | - | - | 2.234 | 2.868 | 1.403 | 1.552 | -4.919 |
| J | - | - | - | - | - | - | - | - | - | - | 1.54 | -2.143 | -0.172 | 1.943 |
| K | - | - | - | - | - | - | - | - | - | - | - | 0.998 | 1.843 | 2.796 |
| L | - | - | - | - | - | - | - | - | - | - | - | - | 0.84 | 2.416 |
| M | - | - | - | - | - | - | - | - | - | - | - | - | - | 1.59 |
| N | - | - | - | - | - | - | - | - | - | - | - | - | - | - |

| Median | A | B | C | D | E | F | G | H | I | J | K | L | M | N |
|---|---|---|---|---|---|---|---|---|---|---|---|---|---|---|
| A | - | 3.217 | -0.559 | -1.533 | 3.1 | 2.754 | 2.697 | 2.707 | 0.769 | 0.722 | 0.978 | 2.243 | 1.313 | -0.709 |
| B | - | - | 3.255 | 3.295 | -0.418 | 1.432 | 4.111 | 1.612 | 3.128 | 3.08 | 2.967 | 2.337 | 2.863 | 3.229 |
| C | - | - | - | -1.593 | 2.991 | 2.601 | 2.858 | 2.562 | 0.579 | -0.052 | 0.256 | 2.037 | 0.792 | -2.987 |
| D | - | - | - | - | 3.046 | 2.69 | 2.771 | 2.626 | 0.252 | 0.338 | 0.784 | 2.129 | 1.155 | -1.592 |
| E | - | - | - | - | - | 0.951 | 4.038 | 1.186 | 2.964 | 2.928 | 2.82 | 2.088 | 2.705 | 3.104 |
| F | - | - | - | - | - | - | 3.833 | -1.237 | 2.496 | 2.497 | 2.318 | 1.222 | 2.163 | 2.708 |
| G | - | - | - | - | - | - | - | 3.602 | 2.884 | 2.922 | 3.013 | 3.385 | 3.092 | 2.751 |
| H | - | - | - | - | - | - | - | - | 2.496 | 2.377 | 2.172 | 0.96 | 1.997 | 2.598 |
| I | - | - | - | - | - | - | - | - | - | -1.892 | 0.078 | 1.834 | 0.638 | -1.498 |
| J | - | - | - | - | - | - | - | - | - | - | -0.851 | 1.636 | 0.068 | -0.163 |
| K | - | - | - | - | - | - | - | - | - | - | - | 1.444 | -0.959 | 0.496 |
| L | - | - | - | - | - | - | - | - | - | - | - | - | 1.204 | 2.097 |
| M | - | - | - | - | - | - | - | - | - | - | - | - | - | 0.971 |
| N | - | - | - | - | - | - | - | - | - | - | - | - | - | - |

| Max | A | B | C | D | E | F | G | H | I | J | K | L | M | N |
|---|---|---|---|---|---|---|---|---|---|---|---|---|---|---|
| A | - | 2.298 | 1.324 | 0.941 | 2.418 | 0.773 | 3.438 | -1.192 | 2.74 | 1.422 | -1.7 | 1.884 | 0.575 | 2.847 |
| B | - | - | 1.306 | 2.85 | -1.15 | 1.753 | 4.047 | 2.252 | 3.583 | 2.987 | 2.275 | 0.334 | 2.755 | 3.674 |
| C | - | - | - | 2.213 | 1.497 | -0.256 | 3.78 | 1.121 | 3.383 | 2.403 | 1.166 | 0.183 | 2.044 | 3.327 |
| D | - | - | - | - | 2.834 | 1.837 | 3.124 | 1.214 | 2.205 | -0.563 | 1.136 | 2.464 | -1.211 | 2.371 |
| E | - | - | - | - | - | 1.965 | 4.094 | 2.388 | 3.658 | 3.089 | 2.423 | 0.795 | 2.859 | 3.73 |
| F | - | - | - | - | - | - | 3.652 | 0.214 | 3.198 | 2.123 | 0.44 | 1.079 | 1.646 | 3.157 |
| G | - | - | - | - | - | - | - | 3.336 | 1.908 | 2.859 | 3.316 | 3.711 | 3.065 | 1.698 |
| H | - | - | - | - | - | - | - | - | 2.763 | 1.662 | -2.492 | 1.759 | 0.993 | 2.938 |
| I | - | - | - | - | - | - | - | - | - | 2.032 | 2.748 | 3.245 | 2.303 | -0.763 |
| J | - | - | - | - | - | - | - | - | - | - | 1.536 | 2.652 | 0.226 | 2.125 |
| K | - | - | - | - | - | - | - | - | - | - | - | 1.769 | 0.848 | 2.912 |
| L | - | - | - | - | - | - | - | - | - | - | - | - | 2.43 | 3.5 |
| M | - | - | - | - | - | - | - | - | - | - | - | - | - | 2.486 |
| N | - | - | - | - | - | - | - | - | - | - | - | - | - | - |

| | |
|---|---|
| | Larger than 10 |
| | Between 8 and 9,99 |
| | Between 6 and 7,99 |
| | Between 4 and 5,99 |
| | Between 2 and 3,99 |
| | Between 0 and 1,99 |
| | NA (Calculation failed) |
