## Supplementary Data S2 for "Interlaboratory comparison of standardised metabolomics and lipidomics analyses in human and rodent blood using the MxP® Quant 500 kit": Cystine.pdf

**Amount of relative differences between laboratories**

| Min | A | B | C | D | E | F | G | H | I | J | K | L | M | N |
|---|---|---|---|---|---|---|---|---|---|---|---|---|---|---|
| A | - | 9.233 | 9.537 | 7.278 | -0.6 | 8.857 | 7.074 | 6.527 | 8.703 | 10.146 | 7.918 | 5.886 | 8.734 | 8.893 |
| B | - | - | 6.899 | 9.382 | 9.389 | 6.626 | 9.685 | 5.065 | 6.728 | 9.137 | 8.701 | 9.09 | 8.685 | 4.287 |
| C | - | - | - | 9.715 | 9.536 | 7.728 | 10.021 | 4.993 | 8.194 | 7.77 | 9.302 | 9.431 | 9.29 | 6.644 |
| D | - | - | - | - | 7.585 | 9.173 | 9.004 | 7.02 | 9.403 | 10.493 | 8.891 | 8.244 | 9.336 | 9.359 |
| E | - | - | - | - | - | 8.958 | 7.473 | 6.544 | 8.894 | 10.111 | 8.382 | 6.448 | 8.955 | 9.166 |
| F | - | - | - | - | - | - | 9.132 | 3.138 | 5.354 | 8.711 | 7.913 | 8.48 | 7.781 | 6.471 |
| G | - | - | - | - | - | - | - | 6.079 | 8.243 | 9.838 | 7.166 | 0.752 | 8.174 | 8.515 |
| H | - | - | - | - | - | - | - | - | 2.902 | 10.332 | 8.736 | 9.465 | 7.895 | 7.833 |
| I | - | - | - | - | - | - | - | - | - | 9.067 | 7.925 | 8.449 | 6.448 | 6.994 |
| J | - | - | - | - | - | - | - | - | - | - | 10.23 | 10.268 | 10.099 | 8.702 |
| K | - | - | - | - | - | - | - | - | - | - | - | 7.236 | 6.998 | 8.18 |
| L | - | - | - | - | - | - | - | - | - | - | - | - | 8.591 | 8.813 |
| M | - | - | - | - | - | - | - | - | - | - | - | - | - | 7.703 |
| N | - | - | - | - | - | - | - | - | - | - | - | - | - | - |

| | |
|---|---|
| | Larger than 10 |
| | Between 8 and 9,99 |
| | Between 6 and 7,99 |
| | Between 4 and 5,99 |
| | Between 2 and 3,99 |
| | Between 0 and 1,99 |
| | NA (Calculation failed) |

| Median | A | B | C | D | E | F | G | H | I | J | K | L | M | N |
|---|---|---|---|---|---|---|---|---|---|---|---|---|---|---|
| A | - | 0.582 | 5.13 | 2.087 | 3.825 | 4.209 | 3.144 | 1.076 | 1.637 | 3.908 | 2.293 | 3.371 | 3.217 | 2.428 |
| B | - | - | 5.025 | 2.342 | 3.915 | 4.092 | 2.987 | -0.691 | 1.2 | 3.998 | 2.028 | 3.465 | 3.044 | 2.068 |
| C | - | - | - | 4.838 | 5.17 | 3.549 | 4.218 | 4.598 | 4.513 | 5.198 | 4.428 | 5.04 | 4.2 | 4.419 |
| D | - | - | - | - | 3.369 | 4.566 | 3.778 | 2.776 | 2.958 | 3.464 | 3.255 | 2.666 | 3.819 | 3.349 |
| E | - | - | - | - | - | 5.247 | 4.745 | 4.251 | 4.331 | 0.391 | 4.477 | 2.16 | 4.773 | 4.514 |
| F | - | - | - | - | - | - | 2.971 | 3.813 | 3.667 | 4.831 | 3.475 | 4.587 | 2.915 | 3.459 |
| G | - | - | - | - | - | - | - | 2.653 | 2.4 | 4.462 | 1.851 | 4.138 | -1.288 | 1.705 |
| H | - | - | - | - | - | - | - | - | 0.506 | 4.135 | 1.866 | 3.473 | 2.939 | 1.69 |
| I | - | - | - | - | - | - | - | - | - | 4.171 | 0.829 | 3.698 | 2.591 | 0.976 |
| J | - | - | - | - | - | - | - | - | - | - | 4.609 | 2.681 | 4.883 | 4.602 |
| K | - | - | - | - | - | - | - | - | - | - | - | 3.861 | 2.051 | -1.664 |
| L | - | - | - | - | - | - | - | - | - | - | - | - | 4.453 | 4.125 |
| M | - | - | - | - | - | - | - | - | - | - | - | - | - | 1.893 |
| N | - | - | - | - | - | - | - | - | - | - | - | - | - | - |

| Max | A | B | C | D | E | F | G | H | I | J | K | L | M | N |
|---|---|---|---|---|---|---|---|---|---|---|---|---|---|---|
| A | - | 1.767 | 5.298 | 2.303 | 3.829 | 4.354 | 3.199 | 2.001 | 2.246 | 3.58 | 2.513 | 3.35 | 3.396 | 2.936 |
| B | - | - | 5.062 | 2.995 | 4.111 | 4.063 | 2.579 | -4.62 | 0.922 | 3.857 | 1.449 | 3.676 | 2.829 | 2.107 |
| C | - | - | - | 5.016 | 5.266 | 3.657 | 4.402 | 4.664 | 4.588 | 5.179 | 4.565 | 5.137 | 4.336 | 4.457 |
| D | - | - | - | - | 3.286 | 4.708 | 3.905 | 3.262 | 3.384 | 2.833 | 3.476 | 2.437 | 3.998 | 3.738 |
| E | - | - | - | - | - | 5.327 | 4.771 | 4.413 | 4.463 | 0.769 | 4.551 | 2.25 | 4.849 | 4.682 |
| F | - | - | - | - | - | - | 3.202 | 3.824 | 3.69 | 4.776 | 3.578 | 4.657 | 3.033 | 3.41 |
| G | - | - | - | - | - | - | - | 2.338 | 2.043 | 4.293 | 1.655 | 4.143 | 0.295 | 0.63 |
| H | - | - | - | - | - | - | - | - | 0.541 | 3.818 | 1.179 | 3.739 | 2.811 | 2.111 |
| I | - | - | - | - | - | - | - | - | - | 4.053 | -0.017 | 3.819 | 2.535 | 1.366 |
| J | - | - | - | - | - | - | - | - | - | - | 4.361 | 1.359 | 4.728 | 4.488 |
| K | - | - | - | - | - | - | - | - | - | - | - | 3.911 | 2.175 | 0.755 |
| L | - | - | - | - | - | - | - | - | - | - | - | - | 4.526 | 4.296 |
| M | - | - | - | - | - | - | - | - | - | - | - | - | - | 1.458 |
| N | - | - | - | - | - | - | - | - | - | - | - | - | - | - |
