## Supplementary Data S2 for "Interlaboratory comparison of standardised metabolomics and lipidomics analyses in human and rodent blood using the MxP® Quant 500 kit": DCA.pdf

**Amount of relative differences between laboratories**

| Min | A | B | C | D | E | F | G | H | I | J | K | L | M | N |
|---|---|---|---|---|---|---|---|---|---|---|---|---|---|---|
| A | - | 3.731 | -1.428 | 4.172 | 1.329 | 3.027 | 4.768 | 3.507 | 3.154 | 5.206 | 3.308 | 3.14 | 2.156 | 1.977 |
| B | - | - | 3.988 | 5.178 | 4.139 | 2.478 | 4.089 | 1.389 | 2.503 | 4.74 | 2.282 | 4.65 | 3.303 | 4.274 |
| C | - | - | - | 4.139 | 1.035 | 3.092 | 4.736 | 3.534 | 3.202 | 5.211 | 3.339 | 3.064 | 2.28 | 1.755 |
| D | - | - | - | - | 3.588 | 4.424 | 5.115 | 4.621 | 4.554 | 5.587 | 4.658 | 2.773 | 4.214 | 3.446 |
| E | - | - | - | - | - | 3.524 | 4.518 | 4.058 | 3.982 | 5.464 | 4.239 | 2.287 | 3.057 | -0.512 |
| F | - | - | - | - | - | - | 4.418 | 1.762 | -0.475 | 4.961 | 0.657 | 4.206 | 1.97 | 3.703 |
| G | - | - | - | - | - | - | - | 6.065 | 6.243 | 5.148 | 6.338 | 6.578 | 6.163 | 6.416 |
| H | - | - | - | - | - | - | - | - | 0.932 | 4.827 | 1.089 | 4.474 | 2.857 | 4.026 |
| I | - | - | - | - | - | - | - | - | - | 4.952 | 0.129 | 4.203 | 1.991 | 3.669 |
| J | - | - | - | - | - | - | - | - | - | - | 5.3 | 5.775 | 5.21 | 5.559 |
| K | - | - | - | - | - | - | - | - | - | - | - | 4.11 | 1.681 | 3.569 |
| L | - | - | - | - | - | - | - | - | - | - | - | - | 3.675 | 1.979 |
| M | - | - | - | - | - | - | - | - | - | - | - | - | - | 3.188 |
| N | - | - | - | - | - | - | - | - | - | - | - | - | - | - |

| | |
|---|---|
| | Larger than 10 |
| | Between 8 and 9,99 |
| | Between 6 and 7,99 |
| | Between 4 and 5,99 |
| | Between 2 and 3,99 |
| | Between 0 and 1,99 |
| | NA (Calculation failed) |

| Median | A | B | C | D | E | F | G | H | I | J | K | L | M | N |
|---|---|---|---|---|---|---|---|---|---|---|---|---|---|---|
| A | - | 3.306 | -0.406 | 3.142 | 3.565 | 2.927 | 1.236 | 3.209 | 2.204 | 3.134 | 0.094 | 2.596 | 2.868 | 2.482 |
| B | - | - | 3.569 | 4.381 | 1.057 | 1.362 | 3.771 | -0.394 | 2.561 | 0.439 | 3.609 | 2.107 | 1.542 | 2.259 |
| C | - | - | - | 3.002 | 3.643 | 3.053 | 0.819 | 3.312 | 2.413 | 3.241 | -1.697 | 2.755 | 2.999 | 2.655 |
| D | - | - | - | - | 4.249 | 3.914 | 2.468 | 4.054 | 3.623 | 4.027 | 2.827 | 3.771 | 3.887 | 3.72 |
| E | - | - | - | - | - | 2.315 | 4.088 | 1.602 | 3.065 | 1.69 | 3.891 | 2.746 | 2.407 | 2.881 |
| F | - | - | - | - | - | - | 3.424 | 0.829 | 1.698 | 0.325 | 3.231 | 0.756 | -1.595 | 1.125 |
| G | - | - | - | - | - | - | - | 3.09 | 2.082 | 2.656 | 0.409 | 2.348 | 2.744 | 2.368 |
| H | - | - | - | - | - | - | - | - | 2.353 | -0.818 | 3.506 | 1.818 | 1.098 | 2.006 |
| I | - | - | - | - | - | - | - | - | - | 2.102 | 2.572 | 0.586 | 1.496 | 0.071 |
| J | - | - | - | - | - | - | - | - | - | - | 3.303 | 1.148 | -0.3 | 1.288 |
| K | - | - | - | - | - | - | - | - | - | - | - | 2.811 | 3.048 | 2.731 |
| L | - | - | - | - | - | - | - | - | - | - | - | - | 0.398 | -0.724 |
| M | - | - | - | - | - | - | - | - | - | - | - | - | - | 0.901 |
| N | - | - | - | - | - | - | - | - | - | - | - | - | - | - |

| Max | A | B | C | D | E | F | G | H | I | J | K | L | M | N |
|---|---|---|---|---|---|---|---|---|---|---|---|---|---|---|
| A | - | 3.234 | -0.302 | 2.92 | 3.816 | 2.91 | 3.78 | 3.16 | 2.006 | 1.648 | 1.348 | 3.023 | 2.952 | 2.802 |
| B | - | - | 3.496 | 4.228 | 2.373 | 1.059 | 4.668 | -1.043 | 2.568 | 2.805 | 3.721 | 0.484 | 0.884 | 1.416 |
| C | - | - | - | 2.744 | 3.885 | 3.047 | 3.665 | 3.276 | 2.26 | 1.97 | 0.777 | 3.149 | 3.084 | 2.946 |
| D | - | - | - | - | 4.331 | 3.811 | 2.501 | 3.947 | 3.43 | 3.312 | 2.218 | 3.868 | 3.83 | 3.756 |
| E | - | - | - | - | - | 2.934 | 5.016 | 2.575 | 3.548 | 3.67 | 4.276 | 2.8 | 2.887 | 3.06 |
| F | - | - | - | - | - | - | 4.528 | 0.62 | 1.922 | 2.246 | 3.441 | -0.604 | -2.188 | -0.73 |
| G | - | - | - | - | - | - | - | 4.302 | 3.93 | 3.876 | 3.232 | 4.262 | 4.227 | 4.197 |
| H | - | - | - | - | - | - | - | - | 2.472 | 2.673 | 3.656 | -0.142 | 0.44 | 1.156 |
| I | - | - | - | - | - | - | - | - | - | 0.074 | 2.786 | 2.101 | 1.95 | 1.611 |
| J | - | - | - | - | - | - | - | - | - | - | 2.559 | 2.362 | 2.246 | 2.024 |
| K | - | - | - | - | - | - | - | - | - | - | - | 3.381 | 3.327 | 3.222 |
| L | - | - | - | - | - | - | - | - | - | - | - | - | -1.18 | 0.342 |
| M | - | - | - | - | - | - | - | - | - | - | - | - | - | -0.299 |
| N | - | - | - | - | - | - | - | - | - | - | - | - | - | - |
