## Supplementary Data S2 for "Interlaboratory comparison of standardised metabolomics and lipidomics analyses in human and rodent blood using the MxP® Quant 500 kit": DG(16.0_16.1).pdf

**Amount of relative differences between laboratories**

| Min | A | B | C | D | E | F | G | H | I | J | K | L | M | N |
|---|---|---|---|---|---|---|---|---|---|---|---|---|---|---|
| A | - | NA | NA | NA | NA | NA | NA | NA | NA | NA | NA | NA | NA | NA |
| B | - | - | NA | 2.475 | 8.267 | NA | NA | NA | 2.149 | -0.3 | 6.761 | 2.937 | NA | NA |
| C | - | - | - | NA | NA | NA | NA | NA | NA | NA | NA | NA | NA | NA |
| D | - | - | - | - | 5.837 | NA | NA | NA | 4.168 | 3.717 | 5.475 | 5.162 | NA | NA |
| E | - | - | - | - | - | NA | NA | NA | 2.206 | 4.312 | 7.949 | 6.334 | NA | NA |
| F | - | - | - | - | - | - | NA | NA | NA | NA | NA | NA | NA | NA |
| G | - | - | - | - | - | - | - | NA | NA | NA | NA | NA | NA | NA |
| H | - | - | - | - | - | - | - | - | NA | NA | NA | NA | NA | NA |
| I | - | - | - | - | - | - | - | - | - | 0.559 | 6.593 | 1.283 | NA | NA |
| J | - | - | - | - | - | - | - | - | - | - | 7.227 | 4.321 | NA | NA |
| K | - | - | - | - | - | - | - | - | - | - | - | 5.495 | NA | NA |
| L | - | - | - | - | - | - | - | - | - | - | - | - | NA | NA |
| M | - | - | - | - | - | - | - | - | - | - | - | - | - | NA |
| N | - | - | - | - | - | - | - | - | - | - | - | - | - | - |

| Median | A | B | C | D | E | F | G | H | I | J | K | L | M | N |
|---|---|---|---|---|---|---|---|---|---|---|---|---|---|---|
| A | - | NA | NA | NA | NA | NA | NA | NA | NA | NA | NA | NA | NA | NA |
| B | - | - | NA | 1.989 | 6.163 | NA | NA | NA | -0.537 | 4.435 | 5.376 | 1.762 | NA | NA |
| C | - | - | - | NA | NA | NA | NA | NA | NA | NA | NA | NA | NA | NA |
| D | - | - | - | - | 6.909 | NA | NA | NA | 3.574 | 3.917 | 4.143 | 4.766 | NA | NA |
| E | - | - | - | - | - | NA | NA | NA | 4.513 | 4.906 | 5.909 | 4.019 | NA | NA |
| F | - | - | - | - | - | - | NA | NA | NA | NA | NA | NA | NA | NA |
| G | - | - | - | - | - | - | - | NA | NA | NA | NA | NA | NA | NA |
| H | - | - | - | - | - | - | - | - | NA | NA | NA | NA | NA | NA |
| I | - | - | - | - | - | - | - | - | - | 4.666 | 5.32 | 2.174 | NA | NA |
| J | - | - | - | - | - | - | - | - | - | - | 5.255 | 3.043 | NA | NA |
| K | - | - | - | - | - | - | - | - | - | - | - | 5.023 | NA | NA |
| L | - | - | - | - | - | - | - | - | - | - | - | - | NA | NA |
| M | - | - | - | - | - | - | - | - | - | - | - | - | - | NA |
| N | - | - | - | - | - | - | - | - | - | - | - | - | - | - |

| Max | A | B | C | D | E | F | G | H | I | J | K | L | M | N |
|---|---|---|---|---|---|---|---|---|---|---|---|---|---|---|
| A | - | NA | NA | NA | NA | NA | NA | NA | NA | NA | NA | NA | NA | NA |
| B | - | - | NA | 1.63 | 7.638 | NA | NA | NA | 2.421 | 5.763 | 2.012 | 4.214 | NA | NA |
| C | - | - | - | NA | NA | NA | NA | NA | NA | NA | NA | NA | NA | NA |
| D | - | - | - | - | 7.587 | NA | NA | NA | 1.085 | 5.894 | -0.802 | 3.948 | NA | NA |
| E | - | - | - | - | - | NA | NA | NA | 6.069 | 5.519 | 6.069 | 6.174 | NA | NA |
| F | - | - | - | - | - | - | NA | NA | NA | NA | NA | NA | NA | NA |
| G | - | - | - | - | - | - | - | NA | NA | NA | NA | NA | NA | NA |
| H | - | - | - | - | - | - | - | - | NA | NA | NA | NA | NA | NA |
| I | - | - | - | - | - | - | - | - | - | 5.975 | 0.515 | 3.812 | NA | NA |
| J | - | - | - | - | - | - | - | - | - | - | 5.241 | 5.554 | NA | NA |
| K | - | - | - | - | - | - | - | - | - | - | - | 3.931 | NA | NA |
| L | - | - | - | - | - | - | - | - | - | - | - | - | NA | NA |
| M | - | - | - | - | - | - | - | - | - | - | - | - | - | NA |
| N | - | - | - | - | - | - | - | - | - | - | - | - | - | - |

| | |
|---|---|
| (red) | Larger than 10 |
| (orange) | Between 8 and 9,99 |
| (light orange) | Between 6 and 7,99 |
| (light green) | Between 4 and 5,99 |
| (green) | Between 2 and 3,99 |
| (dark green) | Between 0 and 1,99 |
| (grey) | NA (Calculation failed) |
