## Supplementary Data S2 for "Interlaboratory comparison of standardised metabolomics and lipidomics analyses in human and rodent blood using the MxP® Quant 500 kit": DG(16.0_18.1).pdf

**Amount of relative differences between laboratories**

| Min | A | B | C | D | E | F | G | H | I | J | K | L | M | N |
|---|---|---|---|---|---|---|---|---|---|---|---|---|---|---|
| A | - | 5.202 | 4.403 | 5.048 | 5.866 | 5.17 | 4.722 | 5.152 | 3.548 | 7.541 | 2.943 | 5.03 | 5.203 | 5.548 |
| B | - | - | 0.342 | -0.28 | 5.332 | 2.561 | 1.523 | 3.205 | 4.711 | 7.001 | 4.594 | -2.541 | 4.105 | 3.734 |
| C | - | - | - | 0.857 | 5.436 | 2.463 | 1.882 | 3.226 | 4.716 | 6.866 | 4.545 | -1.83 | 4.175 | 3.585 |
| D | - | - | - | - | 5.427 | 2.901 | 2.852 | 3.553 | 4.524 | 6.675 | 4.243 | 1.121 | 4.405 | 3.644 |
| E | - | - | - | - | - | 6.139 | 5.948 | 5.886 | 6.952 | 4.047 | 6.774 | 6.33 | 5.197 | 6.097 |
| F | - | - | - | - | - | - | -2.258 | 2.214 | 5.046 | 6.651 | 4.837 | 2.559 | 3.895 | 2.412 |
| G | - | - | - | - | - | - | - | 2.962 | 4.811 | 7.214 | 4.774 | -1.324 | 3.866 | 3.778 |
| H | - | - | - | - | - | - | - | - | 5.296 | 6.869 | 5.178 | 3.395 | 3.263 | 1.459 |
| I | - | - | - | - | - | - | - | - | - | 7.461 | 1.257 | 4.477 | 4.937 | 5.195 |
| J | - | - | - | - | - | - | - | - | - | - | 5.654 | 5.052 | 1.593 | 4.701 |
| K | - | - | - | - | - | - | - | - | - | - | - | 4.522 | 5.023 | 5.165 |
| L | - | - | - | - | - | - | - | - | - | - | - | - | 3.856 | 3.881 |
| M | - | - | - | - | - | - | - | - | - | - | - | - | - | 4.037 |
| N | - | - | - | - | - | - | - | - | - | - | - | - | - | - |

| | |
|---|---|
| | Larger than 10 |
| | Between 8 and 9,99 |
| | Between 6 and 7,99 |
| | Between 4 and 5,99 |
| | Between 2 and 3,99 |
| | Between 0 and 1,99 |
| | NA (Calculation failed) |

| Median | A | B | C | D | E | F | G | H | I | J | K | L | M | N |
|---|---|---|---|---|---|---|---|---|---|---|---|---|---|---|
| A | - | 2.665 | 1.076 | 1.931 | 4.66 | 1.468 | 3.711 | 3.645 | 1.337 | 3.809 | 1.759 | 3.88 | 3.564 | 3.371 |
| B | - | - | 2.058 | 3.565 | 4.254 | 2.107 | 2.722 | 2.545 | 2.07 | 4.352 | 3.535 | 3.142 | 2.438 | 2.198 |
| C | - | - | - | 2.933 | 4.563 | -1.893 | 3.346 | 3.239 | -3.219 | 3.952 | 2.885 | 3.634 | 3.184 | 3.071 |
| D | - | - | - | - | 4.818 | 2.792 | 4.053 | 3.989 | 2.843 | 2.79 | -1.938 | 4.235 | 3.967 | 3.916 |
| E | - | - | - | - | - | 4.948 | 4.209 | 4.252 | 4.916 | 5.416 | 5.4 | 3.758 | 4.291 | 4.146 |
| F | - | - | - | - | - | - | 3.373 | 3.264 | -2.001 | 3.893 | 2.854 | 3.675 | 3.223 | 3.137 |
| G | - | - | - | - | - | - | - | -0.445 | 3.544 | 4.861 | 4.275 | 1.23 | 0.401 | 1.237 |
| H | - | - | - | - | - | - | - | - | 3.399 | 4.791 | 4.218 | 1.719 | -1.372 | 0.237 |
| I | - | - | - | - | - | - | - | - | - | 4.001 | 2.682 | 3.638 | 3.237 | 3.028 |
| J | - | - | - | - | - | - | - | - | - | - | 1.894 | 4.524 | 4.286 | 4.331 |
| K | - | - | - | - | - | - | - | - | - | - | - | 4.161 | 3.904 | 3.797 |
| L | - | - | - | - | - | - | - | - | - | - | - | - | 1.853 | 2.278 |
| M | - | - | - | - | - | - | - | - | - | - | - | - | - | -1.818 |
| N | - | - | - | - | - | - | - | - | - | - | - | - | - | - |

| Max | A | B | C | D | E | F | G | H | I | J | K | L | M | N |
|---|---|---|---|---|---|---|---|---|---|---|---|---|---|---|
| A | - | 2.857 | 3.851 | 4.773 | 3.787 | 4.261 | 1.518 | 1.845 | 0.581 | 6.348 | 3.435 | 2.659 | 3.93 | 2.746 |
| B | - | - | 2.739 | 4.237 | 2.579 | 3.465 | 3.241 | 1.796 | 2.445 | 6.138 | 1.665 | 3.688 | 2.874 | -0.976 |
| C | - | - | - | 3.505 | -1.339 | 2.03 | 3.921 | 3.248 | 3.507 | 5.905 | 1.727 | 4.199 | -0.701 | 2.749 |
| D | - | - | - | - | 3.42 | 2.716 | 4.573 | 4.231 | 4.351 | 5.447 | 3.715 | 4.736 | 3.27 | 4.028 |
| E | - | - | - | - | - | 2.247 | 3.882 | 3.162 | 3.441 | 5.968 | 1.511 | 4.157 | 0.07 | 2.598 |
| F | - | - | - | - | - | - | 4.208 | 3.709 | 3.892 | 5.746 | 2.826 | 4.428 | 1.74 | 3.382 |
| G | - | - | - | - | - | - | - | 2.743 | 2.161 | 6.473 | 3.807 | 1.861 | 4.212 | 3.304 |
| H | - | - | - | - | - | - | - | - | 1.043 | 6.254 | 2.768 | 3.288 | 3.484 | 1.593 |
| I | - | - | - | - | - | - | - | - | - | 6.334 | 3.188 | 2.972 | 3.756 | 2.37 |
| J | - | - | - | - | - | - | - | - | - | - | 5.321 | 5.625 | 5.223 | 5.383 |
| K | - | - | - | - | - | - | - | - | - | - | - | 3.956 | 2.005 | 1.85 |
| L | - | - | - | - | - | - | - | - | - | - | - | - | 4.535 | 3.809 |
| M | - | - | - | - | - | - | - | - | - | - | - | - | - | 2.861 |
| N | - | - | - | - | - | - | - | - | - | - | - | - | - | - |
