## Supplementary Data S2 for "Interlaboratory comparison of standardised metabolomics and lipidomics analyses in human and rodent blood using the MxP® Quant 500 kit": DG(18.1_18.3).pdf

**Amount of relative differences between laboratories**

| Min | A | B | C | D | E | F | G | H | I | J | K | L | M | N |
|---|---|---|---|---|---|---|---|---|---|---|---|---|---|---|
| A | - | 5.406 | 4.137 | NA | 5.305 | NA | 2.015 | NA | 4.673 | NA | 10.912 | NA | 3.479 | 5.551 |
| B | - | - | 4.49 | NA | 3.52 | NA | 5.176 | NA | 2.457 | NA | 11.582 | NA | 1.365 | 3.03 |
| C | - | - | - | NA | 4.895 | NA | 4.029 | NA | 4.623 | NA | 8.193 | NA | 4.865 | 5.32 |
| D | - | - | - | - | NA | NA | NA | NA | NA | NA | NA | NA | NA | NA |
| E | - | - | - | - | - | NA | 6.085 | NA | 3.595 | NA | 8.296 | NA | 5.026 | 4.636 |
| F | - | - | - | - | - | - | NA | NA | NA | NA | NA | NA | NA | NA |
| G | - | - | - | - | - | - | - | NA | 0.002 | NA | 9.731 | NA | 5.214 | 1.959 |
| H | - | - | - | - | - | - | - | - | NA | NA | NA | NA | NA | NA |
| I | - | - | - | - | - | - | - | - | - | NA | 8.095 | NA | 1.07 | 1.009 |
| J | - | - | - | - | - | - | - | - | - | - | NA | NA | NA | NA |
| K | - | - | - | - | - | - | - | - | - | - | - | NA | 3.201 | 4.678 |
| L | - | - | - | - | - | - | - | - | - | - | - | - | NA | NA |
| M | - | - | - | - | - | - | - | - | - | - | - | - | - | 2.24 |
| N | - | - | - | - | - | - | - | - | - | - | - | - | - | - |

| Median | A | B | C | D | E | F | G | H | I | J | K | L | M | N |
|---|---|---|---|---|---|---|---|---|---|---|---|---|---|---|
| A | - | 3.734 | 5.071 | NA | 2.923 | NA | 3.796 | NA | 2.087 | NA | 7.506 | NA | 5.027 | 5.483 |
| B | - | - | 3.953 | NA | 4.418 | NA | 4.706 | NA | 2.848 | NA | 7.534 | NA | 2.699 | 4.235 |
| C | - | - | - | NA | 4.961 | NA | 5.24 | NA | 4.358 | NA | 5.538 | NA | 3.372 | 1.84 |
| D | - | - | - | - | NA | NA | NA | NA | NA | NA | NA | NA | NA | NA |
| E | - | - | - | - | - | NA | 2.434 | NA | 4.145 | NA | 6.843 | NA | 5.397 | 5.674 |
| F | - | - | - | - | - | - | NA | NA | NA | NA | NA | NA | NA | NA |
| G | - | - | - | - | - | - | - | NA | 4.721 | NA | 7.027 | NA | 6.083 | 6.379 |
| H | - | - | - | - | - | - | - | - | NA | NA | NA | NA | NA | NA |
| I | - | - | - | - | - | - | - | - | - | NA | 6.309 | NA | 3.735 | 4.719 |
| J | - | - | - | - | - | - | - | - | - | - | NA | NA | NA | NA |
| K | - | - | - | - | - | - | - | - | - | - | - | NA | 3.862 | 3.76 |
| L | - | - | - | - | - | - | - | - | - | - | - | - | NA | NA |
| M | - | - | - | - | - | - | - | - | - | - | - | - | - | 2.86 |
| N | - | - | - | - | - | - | - | - | - | - | - | - | - | - |

| Max | A | B | C | D | E | F | G | H | I | J | K | L | M | N |
|---|---|---|---|---|---|---|---|---|---|---|---|---|---|---|
| A | - | 5.879 | 5.991 | NA | 3.189 | NA | 4.066 | NA | 5.474 | NA | 10.516 | NA | 6.545 | 6.554 |
| B | - | - | 1.278 | NA | 4.731 | NA | 5.465 | NA | 3.25 | NA | 11.306 | NA | 4.255 | 4.619 |
| C | - | - | - | NA | 4.995 | NA | 5.46 | NA | 3.9 | NA | 7.537 | NA | 3.976 | 4.753 |
| D | - | - | - | - | NA | NA | NA | NA | NA | NA | NA | NA | NA | NA |
| E | - | - | - | - | - | NA | 4.737 | NA | 4.66 | NA | 6.199 | NA | 5.792 | 6.025 |
| F | - | - | - | - | - | - | NA | NA | NA | NA | NA | NA | NA | NA |
| G | - | - | - | - | - | - | - | NA | 5.986 | NA | 10.044 | NA | 6.787 | 6.976 |
| H | - | - | - | - | - | - | - | - | NA | NA | NA | NA | NA | NA |
| I | - | - | - | - | - | - | - | - | - | NA | 6.699 | NA | 5.118 | 5.377 |
| J | - | - | - | - | - | - | - | - | - | - | NA | NA | NA | NA |
| K | - | - | - | - | - | - | - | - | - | - | - | NA | 5.57 | 5.113 |
| L | - | - | - | - | - | - | - | - | - | - | - | - | NA | NA |
| M | - | - | - | - | - | - | - | - | - | - | - | - | - | 3.108 |
| N | - | - | - | - | - | - | - | - | - | - | - | - | - | - |

| | |
|---|---|
| | Larger than 10 |
| | Between 8 and 9,99 |
| | Between 6 and 7,99 |
| | Between 4 and 5,99 |
| | Between 2 and 3,99 |
| | Between 0 and 1,99 |
| | NA (Calculation failed) |
