## Supplementary Data S2 for "Interlaboratory comparison of standardised metabolomics and lipidomics analyses in human and rodent blood using the MxP® Quant 500 kit": DHA.pdf

**Amount of relative differences between laboratories**

| Min | A | B | C | D | E | F | G | H | I | J | K | L | M | N |
|---|---|---|---|---|---|---|---|---|---|---|---|---|---|---|
| A | - | 3.147 | -0.251 | 4.247 | 1.892 | 3.098 | 2.658 | 3.806 | 4.141 | 5.144 | 4.063 | 1.007 | -0.238 | -0.357 |
| B | - | - | 3.364 | 4.633 | 3.601 | 3.936 | 3.953 | 3.44 | 4.483 | 5.447 | 4.68 | 3.369 | 1.886 | 1.88 |
| C | - | - | - | 3.628 | 0.542 | 2.027 | 2.213 | 4.314 | 3.324 | 4.888 | 3.768 | -2.79 | 2.795 | 2.749 |
| D | - | - | - | - | 3.649 | 3.109 | 3.361 | 4.798 | 0.315 | 3.796 | 0.712 | 3.81 | 3.949 | 4.057 |
| E | - | - | - | - | - | 1.792 | 1.52 | 4.276 | 3.395 | 4.852 | 3.611 | 0.713 | 2.577 | 2.696 |
| F | - | - | - | - | - | - | 0.688 | 4.403 | 2.988 | 4.628 | 2.957 | 2.523 | 2.94 | 3.155 |
| G | - | - | - | - | - | - | - | 4.459 | 2.809 | 4.648 | 3.162 | 2.134 | 3.178 | 3.234 |
| H | - | - | - | - | - | - | - | - | 5.206 | 5.864 | 5.204 | 4.303 | 4.092 | 3.959 |
| I | - | - | - | - | - | - | - | - | - | 3.538 | 2.569 | 4.21 | 3.682 | 4.023 |
| J | - | - | - | - | - | - | - | - | - | - | 3.842 | 4.661 | 4.615 | 4.713 |
| K | - | - | - | - | - | - | - | - | - | - | - | 3.661 | 3.755 | 3.902 |
| L | - | - | - | - | - | - | - | - | - | - | - | - | 1.719 | 2.06 |
| M | - | - | - | - | - | - | - | - | - | - | - | - | - | 0.868 |
| N | - | - | - | - | - | - | - | - | - | - | - | - | - | - |

| | |
|---|---|
| | Larger than 10 |
| | Between 8 and 9,99 |
| | Between 6 and 7,99 |
| | Between 4 and 5,99 |
| | Between 2 and 3,99 |
| | Between 0 and 1,99 |
| | NA (Calculation failed) |

| Median | A | B | C | D | E | F | G | H | I | J | K | L | M | N |
|---|---|---|---|---|---|---|---|---|---|---|---|---|---|---|
| A | - | 2.53 | 4.274 | 4.341 | 3.675 | 3.425 | 4.167 | 3.615 | 2.273 | 4.89 | 4.035 | 3.002 | -2.269 | -2.676 |
| B | - | - | 3.711 | 3.81 | 2.887 | 2.518 | 3.541 | 4.107 | -8.569 | 4.546 | 3.426 | 1.113 | 2.509 | 2.17 |
| C | - | - | - | -0.293 | 2.311 | 2.671 | 0.374 | 4.749 | 3.52 | 3.198 | 1.002 | 3.278 | 4.058 | 3.952 |
| D | - | - | - | - | 2.631 | 2.973 | 0.942 | 4.759 | 3.677 | 2.968 | 1.675 | 3.355 | 4.093 | 4.06 |
| E | - | - | - | - | - | 0.702 | 2.07 | 4.503 | 2.795 | 3.897 | 1.659 | 2.217 | 3.578 | 3.474 |
| F | - | - | - | - | - | - | 2.671 | 4.388 | 2.419 | 4.089 | 2.331 | 1.402 | 3.329 | 3.261 |
| G | - | - | - | - | - | - | - | 4.691 | 3.41 | 3.372 | -0.048 | 3.085 | 3.955 | 3.871 |
| H | - | - | - | - | - | - | - | - | 4.304 | 5.586 | 5.039 | 4.534 | 3.784 | 3.847 |
| I | - | - | - | - | - | - | - | - | - | 4.587 | 3.506 | 2.015 | 1.903 | 2.142 |
| J | - | - | - | - | - | - | - | - | - | - | 3.354 | 4.046 | 4.513 | 4.511 |
| K | - | - | - | - | - | - | - | - | - | - | - | 2.849 | 3.833 | 3.804 |
| L | - | - | - | - | - | - | - | - | - | - | - | - | 2.934 | 2.798 |
| M | - | - | - | - | - | - | - | - | - | - | - | - | - | -1.957 |
| N | - | - | - | - | - | - | - | - | - | - | - | - | - | - |

| Max | A | B | C | D | E | F | G | H | I | J | K | L | M | N |
|---|---|---|---|---|---|---|---|---|---|---|---|---|---|---|
| A | - | 4.507 | 5.435 | 4.46 | 4.608 | 3.778 | 4.979 | 3.284 | 3.44 | 4.533 | 4.004 | 3.86 | 0.862 | 0.689 |
| B | - | - | 4.064 | -0.731 | 0.41 | 2.876 | 2.841 | 4.726 | 4.775 | -1.62 | 2.447 | 2.741 | 4.323 | 4.105 |
| C | - | - | - | 3.891 | 3.721 | 4.366 | 3.035 | 5.21 | 5.239 | 3.813 | 4.248 | 4.326 | 4.976 | 4.861 |
| D | - | - | - | - | 0.959 | 2.765 | 2.962 | 4.701 | 4.751 | -0.1 | 2.288 | 2.614 | 4.287 | 4.063 |
| E | - | - | - | - | - | 3.1 | 2.523 | 4.778 | 4.824 | -0.016 | 2.744 | 2.984 | 4.397 | 4.194 |
| F | - | - | - | - | - | - | 3.969 | 4.366 | 4.434 | 3.048 | 1.039 | -0.474 | 3.772 | 3.412 |
| G | - | - | - | - | - | - | - | 4.971 | 5.01 | 2.675 | 3.555 | 3.692 | 4.664 | 4.507 |
| H | - | - | - | - | - | - | - | - | 0.31 | 5.187 | 4.836 | 4.748 | 3.133 | 3.653 |
| I | - | - | - | - | - | - | - | - | - | 5.253 | 4.915 | 4.832 | 3.341 | 3.805 |
| J | - | - | - | - | - | - | - | - | - | - | 2.525 | 2.806 | 4.341 | 4.128 |
| K | - | - | - | - | - | - | - | - | - | - | - | 0.383 | 3.945 | 3.636 |
| L | - | - | - | - | - | - | - | - | - | - | - | - | 3.835 | 3.496 |
| M | - | - | - | - | - | - | - | - | - | - | - | - | - | 1.805 |
| N | - | - | - | - | - | - | - | - | - | - | - | - | - | - |
