## Supplementary Data S2 for "Interlaboratory comparison of standardised metabolomics and lipidomics analyses in human and rodent blood using the MxP® Quant 500 kit": DiCA(12.0).pdf

**Amount of relative differences between laboratories**

| Min | A | B | C | D | E | F | G | H | I | J | K | L | M | N |
|---|---|---|---|---|---|---|---|---|---|---|---|---|---|---|
| A | - | NA | NA | 4.358 | 3.277 | 0.347 | NA | NA | 5.163 | 6.362 | 4.587 | NA | NA | NA |
| B | - | - | NA | NA | NA | NA | NA | NA | NA | NA | NA | NA | NA | NA |
| C | - | - | - | NA | NA | NA | NA | NA | NA | NA | NA | NA | NA | NA |
| D | - | - | - | - | 6.031 | 2.397 | NA | NA | 4.912 | 6.495 | 4.404 | NA | NA | NA |
| E | - | - | - | - | - | 3.264 | NA | NA | 4.995 | 6.344 | 5.184 | NA | NA | NA |
| F | - | - | - | - | - | - | NA | NA | 5.19 | 6.521 | 4.257 | NA | NA | NA |
| G | - | - | - | - | - | - | - | NA | NA | NA | NA | NA | NA | NA |
| H | - | - | - | - | - | - | - | - | NA | NA | NA | NA | NA | NA |
| I | - | - | - | - | - | - | - | - | - | 6.733 | 7.59 | NA | NA | NA |
| J | - | - | - | - | - | - | - | - | - | - | 8.839 | NA | NA | NA |
| K | - | - | - | - | - | - | - | - | - | - | - | NA | NA | NA |
| L | - | - | - | - | - | - | - | - | - | - | - | - | NA | NA |
| M | - | - | - | - | - | - | - | - | - | - | - | - | - | NA |
| N | - | - | - | - | - | - | - | - | - | - | - | - | - | - |

| Median | A | B | C | D | E | F | G | H | I | J | K | L | M | N |
|---|---|---|---|---|---|---|---|---|---|---|---|---|---|---|
| A | - | NA | NA | 2.429 | 2.145 | 0.732 | NA | NA | 5.102 | 5.752 | -1.121 | NA | NA | NA |
| B | - | - | NA | NA | NA | NA | NA | NA | NA | NA | NA | NA | NA | NA |
| C | - | - | - | NA | NA | NA | NA | NA | NA | NA | NA | NA | NA | NA |
| D | - | - | - | - | 2.801 | 2.989 | NA | NA | 4.807 | 5.681 | 2.206 | NA | NA | NA |
| E | - | - | - | - | - | 3.085 | NA | NA | 4.941 | 5.702 | 2.214 | NA | NA | NA |
| F | - | - | - | - | - | - | NA | NA | 5.137 | 5.82 | 1.132 | NA | NA | NA |
| G | - | - | - | - | - | - | - | NA | NA | NA | NA | NA | NA | NA |
| H | - | - | - | - | - | - | - | - | NA | NA | NA | NA | NA | NA |
| I | - | - | - | - | - | - | - | - | - | 5.708 | 5.704 | NA | NA | NA |
| J | - | - | - | - | - | - | - | - | - | - | 7.117 | NA | NA | NA |
| K | - | - | - | - | - | - | - | - | - | - | - | NA | NA | NA |
| L | - | - | - | - | - | - | - | - | - | - | - | - | NA | NA |
| M | - | - | - | - | - | - | - | - | - | - | - | - | - | NA |
| N | - | - | - | - | - | - | - | - | - | - | - | - | - | - |

| Max | A | B | C | D | E | F | G | H | I | J | K | L | M | N |
|---|---|---|---|---|---|---|---|---|---|---|---|---|---|---|
| A | - | NA | NA | 3.815 | 1.241 | 1.798 | NA | NA | 4.14 | 3.745 | 2.478 | NA | NA | NA |
| B | - | - | NA | NA | NA | NA | NA | NA | NA | NA | NA | NA | NA | NA |
| C | - | - | - | NA | NA | NA | NA | NA | NA | NA | NA | NA | NA | NA |
| D | - | - | - | - | 3.985 | 4.351 | NA | NA | 2.122 | 2.082 | 3.487 | NA | NA | NA |
| E | - | - | - | - | - | 1.704 | NA | NA | 4.125 | 3.386 | 1.799 | NA | NA | NA |
| F | - | - | - | - | - | - | NA | NA | 4.366 | 2.594 | 2.934 | NA | NA | NA |
| G | - | - | - | - | - | - | - | NA | NA | NA | NA | NA | NA | NA |
| H | - | - | - | - | - | - | - | - | NA | NA | NA | NA | NA | NA |
| I | - | - | - | - | - | - | - | - | - | 4.639 | 4.017 | NA | NA | NA |
| J | - | - | - | - | - | - | - | - | - | - | 5.837 | NA | NA | NA |
| K | - | - | - | - | - | - | - | - | - | - | - | NA | NA | NA |
| L | - | - | - | - | - | - | - | - | - | - | - | - | NA | NA |
| M | - | - | - | - | - | - | - | - | - | - | - | - | - | NA |
| N | - | - | - | - | - | - | - | - | - | - | - | - | - | - |

| | |
|---|---|
| | Larger than 10 |
| | Between 8 and 9,99 |
| | Between 6 and 7,99 |
| | Between 4 and 5,99 |
| | Between 2 and 3,99 |
| | Between 0 and 1,99 |
| | NA (Calculation failed) |
