## Supplementary Data S2 for "Interlaboratory comparison of standardised metabolomics and lipidomics analyses in human and rodent blood using the MxP® Quant 500 kit": EPA.pdf

**Amount of relative differences between laboratories**

| Min | A | B | C | D | E | F | G | H | I | J | K | L | M | N |
|---|---|---|---|---|---|---|---|---|---|---|---|---|---|---|
| A | - | 4.861 | 4.518 | 5.163 | 2.789 | 4.033 | 5.925 | 6.641 | 3.698 | 5.661 | 5.157 | 2.288 | 5.145 | 5.436 |
| B | - | - | 2.735 | 0.56 | 4.637 | 4.693 | 5.467 | 6.475 | 3.307 | 4.658 | -6.124 | 5.022 | 0.1 | 4.884 |
| C | - | - | - | 4.458 | 5.527 | 5.655 | 4.986 | 6.343 | 4.718 | 3.507 | 4.356 | 5.837 | 4.175 | 4.265 |
| D | - | - | - | - | 4.359 | 4.611 | 5.617 | 6.441 | 1.987 | 4.701 | 0.01 | 4.885 | 0.791 | 5.13 |
| E | - | - | - | - | - | 2.576 | 5.784 | 6.571 | 3.088 | 5.465 | 4.749 | 1.786 | 4.753 | 5.341 |
| F | - | - | - | - | - | - | 5.746 | 6.548 | 3.155 | 5.351 | 4.406 | 2.832 | 4.451 | 5.38 |
| G | - | - | - | - | - | - | - | 5.823 | 6.427 | 5.9 | 6.82 | 7.299 | 6.78 | 3.808 |
| H | - | - | - | - | - | - | - | - | 8.039 | 7.802 | 8.457 | 8.717 | 8.4 | 6.948 |
| I | - | - | - | - | - | - | - | - | - | 5.354 | 4.524 | 3.468 | 4.518 | 5.121 |
| J | - | - | - | - | - | - | - | - | - | - | 5.189 | 6.143 | 5.067 | 4.051 |
| K | - | - | - | - | - | - | - | - | - | - | - | 5.024 | 0.222 | 4.904 |
| L | - | - | - | - | - | - | - | - | - | - | - | - | 4.859 | 5.427 |
| M | - | - | - | - | - | - | - | - | - | - | - | - | - | 4.895 |
| N | - | - | - | - | - | - | - | - | - | - | - | - | - | - |

| | |
|---|---|
| | Larger than 10 |
| | Between 8 and 9,99 |
| | Between 6 and 7,99 |
| | Between 4 and 5,99 |
| | Between 2 and 3,99 |
| | Between 0 and 1,99 |
| | NA (Calculation failed) |

| Median | A | B | C | D | E | F | G | H | I | J | K | L | M | N |
|---|---|---|---|---|---|---|---|---|---|---|---|---|---|---|
| A | - | 2 | 2.899 | 2.962 | 0.353 | 3.124 | 3.467 | 5.67 | 3.353 | 2.499 | 2.106 | 2.685 | 1.119 | 4.351 |
| B | - | - | -1.064 | 3.869 | 2.48 | 3.9 | 3.039 | 5.601 | 2.697 | 0.823 | -0.812 | 3.424 | 0.815 | 4.082 |
| C | - | - | - | 3.883 | 2.465 | 3.894 | 3.026 | 5.59 | 2.587 | 1.57 | -5.079 | 3.354 | -1.185 | 3.989 |
| D | - | - | - | - | 3.041 | -1.606 | 4.475 | 5.792 | 4.226 | 3.891 | 3.615 | 1.942 | 3.561 | 4.81 |
| E | - | - | - | - | - | 2.971 | 3.571 | 5.675 | 3.508 | 2.772 | 2.398 | 2.324 | 1.792 | 4.415 |
| F | - | - | - | - | - | - | 4.244 | 5.787 | 4.192 | 3.825 | 3.627 | 1.679 | 3.467 | 4.783 |
| G | - | - | - | - | - | - | - | 5.335 | 2.975 | 3.842 | 4.155 | 4.788 | 4.072 | 2.172 |
| H | - | - | - | - | - | - | - | - | 6.314 | 6.453 | 6.643 | 6.806 | 6.556 | 5.915 |
| I | - | - | - | - | - | - | - | - | - | 2.36 | 2.753 | 4.209 | 3.161 | 3.494 |
| J | - | - | - | - | - | - | - | - | - | - | 1.787 | 3.68 | 1.823 | 3.929 |
| K | - | - | - | - | - | - | - | - | - | - | - | 3.338 | 0.21 | 4.126 |
| L | - | - | - | - | - | - | - | - | - | - | - | - | 3.012 | 4.651 |
| M | - | - | - | - | - | - | - | - | - | - | - | - | - | 4.197 |
| N | - | - | - | - | - | - | - | - | - | - | - | - | - | - |

| Max | A | B | C | D | E | F | G | H | I | J | K | L | M | N |
|---|---|---|---|---|---|---|---|---|---|---|---|---|---|---|
| A | - | 3.124 | 4.403 | 5.105 | 2.337 | 4.301 | 5.512 | 4.621 | 3.109 | 4.549 | 3.256 | 3.445 | 3.574 | 2.664 |
| B | - | - | 3.454 | 4.5 | 1.734 | 3.34 | 5.039 | 3.8 | 3.992 | 3.745 | -0.295 | 1.103 | 1.565 | 1.135 |
| C | - | - | - | 3.371 | 3.722 | -0.52 | 4.226 | 1.525 | 4.622 | 1.3 | 3.184 | 3.048 | 2.814 | 3.579 |
| D | - | - | - | - | 4.426 | 3.355 | 3.072 | 2.755 | 4.99 | 2.927 | 4.178 | 4.088 | 4.007 | 4.317 |
| E | - | - | - | - | - | 3.796 | 5.227 | 4.079 | 3.703 | 4.114 | 2.096 | 2.491 | 2.701 | 0.153 |
| F | - | - | - | - | - | - | 4.369 | 2.016 | 4.564 | 1.617 | 3.078 | 2.862 | 2.694 | 3.461 |
| G | - | - | - | - | - | - | - | 3.932 | 5.215 | 3.905 | 4.599 | 4.557 | 4.501 | 4.709 |
| H | - | - | - | - | - | - | - | - | 4.704 | -7.615 | 3.466 | 3.295 | 3.21 | 3.75 |
| I | - | - | - | - | - | - | - | - | - | 5.125 | 4.313 | 4.421 | 4.489 | 4.022 |
| J | - | - | - | - | - | - | - | - | - | - | 3.477 | 3.319 | 3.198 | 3.773 |
| K | - | - | - | - | - | - | - | - | - | - | - | 0.347 | 1.099 | 1.521 |
| L | - | - | - | - | - | - | - | - | - | - | - | - | -0.279 | 2.068 |
| M | - | - | - | - | - | - | - | - | - | - | - | - | - | 2.323 |
| N | - | - | - | - | - | - | - | - | - | - | - | - | - | - |
