## Supplementary Data S2 for "Interlaboratory comparison of standardised metabolomics and lipidomics analyses in human and rodent blood using the MxP® Quant 500 kit": FA(18.2).pdf

**Amount of relative differences between laboratories**

| Min | A | B | C | D | E | F | G | H | I | J | K | L | M | N |
|---|---|---|---|---|---|---|---|---|---|---|---|---|---|---|
| A | - | 5.915 | 5.924 | 5.851 | 6.166 | 5.941 | 5.376 | 5.084 | 7.2 | 2.893 | 5.255 | 7.176 | 6.196 | 5.044 |
| B | - | - | 5.236 | 1.713 | 3.631 | 3.976 | 5.423 | 7.709 | 7.038 | 7.35 | 6.174 | 7.027 | 0.195 | 5.545 |
| C | - | - | - | 5.258 | 5.35 | 4.547 | 3.859 | 6.28 | 6.292 | 5.777 | 3.366 | 6.236 | 4.679 | 4.267 |
| D | - | - | - | - | 4.038 | 3.347 | 5.288 | 7.659 | 7.148 | 7.272 | 6.017 | 7.08 | 2.353 | 5.368 |
| E | - | - | - | - | - | 4.918 | 5.554 | 7.698 | 6.142 | 7.403 | 6.465 | 6.074 | 4.682 | 5.291 |
| F | - | - | - | - | - | - | 3.603 | 7.072 | 6.165 | 6.712 | 5.478 | 6.044 | 0.831 | 1.994 |
| G | - | - | - | - | - | - | - | 6.868 | 6.781 | 6.384 | 4.389 | 6.683 | 4.689 | 0.477 |
| H | - | - | - | - | - | - | - | - | 6.965 | 4.301 | 5.705 | 6.88 | 6.264 | 5.718 |
| I | - | - | - | - | - | - | - | - | - | 7.559 | 6.93 | 0.865 | 6.022 | 5.195 |
| J | - | - | - | - | - | - | - | - | - | - | 5.664 | 7.223 | 6.428 | 4.851 |
| K | - | - | - | - | - | - | - | - | - | - | - | 6.444 | 5.104 | 4.491 |
| L | - | - | - | - | - | - | - | - | - | - | - | - | 5.841 | 4.895 |
| M | - | - | - | - | - | - | - | - | - | - | - | - | - | 1.46 |
| N | - | - | - | - | - | - | - | - | - | - | - | - | - | - |

| Median | A | B | C | D | E | F | G | H | I | J | K | L | M | N |
|---|---|---|---|---|---|---|---|---|---|---|---|---|---|---|
| A | - | 5.665 | 0.844 | 5.636 | 5.521 | 4.86 | 4.853 | 5.17 | 4.931 | 3.39 | 1.642 | 4.648 | 4.191 | 4.505 |
| B | - | - | 6.575 | 0.952 | 3.335 | 5.504 | 5.461 | 7.5 | 5.093 | 6.379 | 6.566 | 5.446 | 5.897 | 5.785 |
| C | - | - | - | 5.557 | 5.416 | 4.585 | 4.656 | 5.593 | 5.048 | 2.229 | -2.363 | 4.851 | 4.3 | 4.374 |
| D | - | - | - | - | 2.991 | 5.405 | 5.377 | 7.459 | 4.991 | 6.314 | 6.508 | 5.373 | 5.837 | 5.659 |
| E | - | - | - | - | - | 5.031 | 4.941 | 7.312 | 4.132 | 6.069 | 6.235 | 4.645 | 5.356 | 5.251 |
| F | - | - | - | - | - | - | 0.582 | 6.58 | 3.55 | 4.694 | 5.001 | 2.603 | 2.578 | 1.52 |
| G | - | - | - | - | - | - | - | 6.58 | 2.74 | 4.759 | 5.133 | -3.879 | 3.459 | 2.768 |
| H | - | - | - | - | - | - | - | - | 5.616 | 5.105 | 4.857 | 5.493 | 5.329 | 5.491 |
| I | - | - | - | - | - | - | - | - | - | 5.723 | 5.855 | 2.368 | 4.464 | 4.472 |
| J | - | - | - | - | - | - | - | - | - | - | 3.343 | 3.436 | 2.224 | 3.55 |
| K | - | - | - | - | - | - | - | - | - | - | - | 4.781 | 4.231 | 4.399 |
| L | - | - | - | - | - | - | - | - | - | - | - | - | 4.002 | 4.021 |
| M | - | - | - | - | - | - | - | - | - | - | - | - | - | -0.312 |
| N | - | - | - | - | - | - | - | - | - | - | - | - | - | - |

| Max | A | B | C | D | E | F | G | H | I | J | K | L | M | N |
|---|---|---|---|---|---|---|---|---|---|---|---|---|---|---|
| A | - | 5.316 | 5.448 | 5.303 | 4.349 | 3.01 | 3.65 | 5.308 | 5.097 | 4.511 | 4.909 | 5.388 | 4.407 | 2.916 |
| B | - | - | 7.128 | -2.226 | 5.012 | 6.31 | 5.51 | 7.052 | 6.978 | 4.82 | 6.858 | 7.139 | 6.689 | 6.32 |
| C | - | - | - | 5.866 | 5.479 | 4.627 | 5.288 | 1.123 | 2.985 | 5.512 | 3.229 | 1.491 | 3.95 | 4.65 |
| D | - | - | - | - | 4.999 | 6.308 | 5.484 | 7.036 | 7.003 | 4.807 | 6.856 | 7.134 | 6.67 | 6.278 |
| E | - | - | - | - | - | 5.15 | 3.273 | 6.238 | 6.055 | 1.638 | 5.977 | 6.262 | 5.699 | 5.126 |
| F | - | - | - | - | - | - | 4.262 | 4.861 | 4.437 | 4.828 | 4.331 | 4.779 | 3.527 | -1.302 |
| G | - | - | - | - | - | - | - | 5.891 | 5.664 | 3.551 | 5.598 | 5.844 | 5.224 | 4.507 |
| H | - | - | - | - | - | - | - | - | 2.79 | 5.477 | 2.817 | 2.104 | 3.798 | 4.488 |
| I | - | - | - | - | - | - | - | - | - | 5.339 | 1.41 | 2.814 | 3.123 | 4.339 |
| J | - | - | - | - | - | - | - | - | - | - | 6.081 | 6.061 | 5.748 | 5.274 |
| K | - | - | - | - | - | - | - | - | - | - | - | 2.552 | 2.921 | 4.088 |
| L | - | - | - | - | - | - | - | - | - | - | - | - | 3.767 | 4.554 |
| M | - | - | - | - | - | - | - | - | - | - | - | - | - | 3.493 |
| N | - | - | - | - | - | - | - | - | - | - | - | - | - | - |

| | |
|---|---|
| (red) | Larger than 10 |
| (orange) | Between 8 and 9,99 |
| (light orange) | Between 6 and 7,99 |
| (light green) | Between 4 and 5,99 |
| (green) | Between 2 and 3,99 |
| (dark green) | Between 0 and 1,99 |
| (light grey) | NA (Calculation failed) |
