## Supplementary Data S2 for "Interlaboratory comparison of standardised metabolomics and lipidomics analyses in human and rodent blood using the MxP® Quant 500 kit": Hex3Cer(d18.1-22.0).pdf

**Amount of relative differences between laboratories**

| Min | A | B | C | D | E | F | G | H | I | J | K | L | M | N |
|---|---|---|---|---|---|---|---|---|---|---|---|---|---|---|
| A | - | 5.787 | 6.59 | 5.709 | 4.568 | 5.635 | 5.714 | 4.446 | 7.126 | 4.408 | 6.869 | 3.264 | 6.259 | 7.017 |
| B | - | - | 5.232 | 8.068 | 0.602 | 2.68 | 7.509 | 5.037 | 4.364 | 7.465 | 4.735 | 8.403 | 4.933 | 5.55 |
| C | - | - | - | 7.417 | 1.133 | -0.023 | 6.181 | 2.036 | 3.221 | 7.623 | 5.542 | 7.618 | 4.501 | 5.426 |
| D | - | - | - | - | 3.828 | 5.53 | 4.36 | 3.591 | 6.695 | 6.335 | 6.651 | 5.037 | 5.909 | 6.709 |
| E | - | - | - | - | - | 4.21 | 2.894 | 2.499 | 8.569 | 7.27 | 7.636 | 6.97 | 5.184 | 8.489 |
| F | - | - | - | - | - | - | 5.146 | 2.888 | 6.533 | 7.603 | 6.323 | 7.783 | 0.374 | 6.349 |
| G | - | - | - | - | - | - | - | 4.368 | 6.022 | 6.525 | 6.47 | 6.143 | 5.677 | 6.548 |
| H | - | - | - | - | - | - | - | - | 8.102 | 8.551 | 6.504 | 8.667 | 4.74 | 8.205 |
| I | - | - | - | - | - | - | - | - | - | 7.943 | 5.692 | 7.5 | 4.016 | 5.694 |
| J | - | - | - | - | - | - | - | - | - | - | 7.432 | 5.503 | 6.256 | 7.466 |
| K | - | - | - | - | - | - | - | - | - | - | - | 8.387 | 5.208 | 3.12 |
| L | - | - | - | - | - | - | - | - | - | - | - | - | 6.26 | 7.168 |
| M | - | - | - | - | - | - | - | - | - | - | - | - | - | 5.286 |
| N | - | - | - | - | - | - | - | - | - | - | - | - | - | - |

| | |
|---|---|
| | Larger than 10 |
| | Between 8 and 9,99 |
| | Between 6 and 7,99 |
| | Between 4 and 5,99 |
| | Between 2 and 3,99 |
| | Between 0 and 1,99 |
| | NA (Calculation failed) |

| Median | A | B | C | D | E | F | G | H | I | J | K | L | M | N |
|---|---|---|---|---|---|---|---|---|---|---|---|---|---|---|
| A | - | 4.185 | 2.879 | 0.6 | 4.118 | 4.058 | 2.767 | 1.831 | 2.368 | 1.866 | -0.663 | 3.604 | -1.253 | 1.574 |
| B | - | - | 4.847 | 4.385 | 0.522 | 0.295 | 5.09 | 4.329 | 3.934 | 3.991 | 4.54 | 3.185 | 4.673 | 4.183 |
| C | - | - | - | 2.709 | 4.357 | 4.523 | 1.562 | 2.439 | 3.336 | 3.005 | 2.162 | 4.013 | 1.731 | 2.938 |
| D | - | - | - | - | 3.939 | 3.919 | 3.14 | 0.8 | 1.909 | 0.758 | 0.235 | 3.416 | 1.191 | 0.58 |
| E | - | - | - | - | - | -1.049 | 4.863 | 3.966 | 3.099 | 4.115 | 3.733 | 2.813 | 4.504 | 2.382 |
| F | - | - | - | - | - | - | 4.962 | 3.712 | 3.846 | 4.151 | 4.376 | 2.979 | 4.63 | 4.112 |
| G | - | - | - | - | - | - | - | 3.439 | 3.577 | 3.352 | 2.822 | 4.178 | 2.592 | 3.269 |
| H | - | - | - | - | - | - | - | - | 2.917 | 1.57 | 0.457 | 2.45 | 2.961 | 3.142 |
| I | - | - | - | - | - | - | - | - | - | 1.532 | 2.599 | 2.757 | 2.872 | 1.425 |
| J | - | - | - | - | - | - | - | - | - | - | -1.19 | 3.182 | 2.188 | 0.421 |
| K | - | - | - | - | - | - | - | - | - | - | - | 3.651 | 0.398 | 1.951 |
| L | - | - | - | - | - | - | - | - | - | - | - | - | 3.839 | 3.402 |
| M | - | - | - | - | - | - | - | - | - | - | - | - | - | 2.18 |
| N | - | - | - | - | - | - | - | - | - | - | - | - | - | - |

| Max | A | B | C | D | E | F | G | H | I | J | K | L | M | N |
|---|---|---|---|---|---|---|---|---|---|---|---|---|---|---|
| A | - | 3.274 | 3.83 | -1.132 | 3.835 | 3.762 | 3.622 | 4.195 | 1.267 | 2.295 | 4.394 | 3.617 | 3.896 | 3.473 |
| B | - | - | 4.818 | 3.497 | 0.482 | -0.43 | 4.345 | 2.268 | 3.905 | 2.95 | 5.025 | 2.418 | 4.618 | 4.667 |
| C | - | - | - | 3.747 | 4.876 | 4.658 | 1.813 | 4.074 | 3.484 | 4.206 | 1.753 | 4.671 | -1.67 | 1.9 |
| D | - | - | - | - | 3.99 | 3.611 | 3.529 | 3.409 | 0.898 | 2.557 | 4.331 | 3.653 | 3.808 | 3.32 |
| E | - | - | - | - | - | 1.176 | 5.022 | 5.363 | 4.927 | 3.373 | 5.724 | 0.882 | 5.078 | 5.516 |
| F | - | - | - | - | - | - | 4.944 | 4.317 | 4.35 | 3.135 | 5.306 | 0.509 | 4.866 | 4.822 |
| G | - | - | - | - | - | - | - | -0.13 | 3.162 | 3.946 | 3.023 | 4.459 | 1.297 | -1.774 |
| H | - | - | - | - | - | - | - | - | 2.903 | 3.995 | 4.213 | 4.555 | 2.081 | 4.527 |
| I | - | - | - | - | - | - | - | - | - | 3.244 | 4.072 | 3.906 | 3.553 | 3.129 |
| J | - | - | - | - | - | - | - | - | - | - | 5.031 | 2.904 | 4.305 | 4.185 |
| K | - | - | - | - | - | - | - | - | - | - | - | 4.796 | 2.428 | 2.396 |
| L | - | - | - | - | - | - | - | - | - | - | - | - | 4.973 | 4.795 |
| M | - | - | - | - | - | - | - | - | - | - | - | - | - | -0.529 |
| N | - | - | - | - | - | - | - | - | - | - | - | - | - | - |
