## Supplementary Data S2 for "Interlaboratory comparison of standardised metabolomics and lipidomics analyses in human and rodent blood using the MxP® Quant 500 kit": Hex3Cer(d18.1-24.1).pdf

**Amount of relative differences between laboratories**

| Min | A | B | C | D | E | F | G | H | I | J | K | L | M | N |
|---|---|---|---|---|---|---|---|---|---|---|---|---|---|---|
| A | - | 4.387 | 1.993 | 5.062 | 4.703 | 5.887 | 4.829 | 6.008 | 3.735 | 6.213 | 6.735 | 4.865 | 5.76 | 6.187 |
| B | - | - | 3.132 | 5.567 | 4.999 | 6.022 | 5.601 | 6.137 | 4.335 | 5.84 | 6.737 | 5.339 | 6.026 | 6.185 |
| C | - | - | - | 5.674 | 5.002 | 6.06 | 5.311 | 6.162 | 4.556 | 6.307 | 6.803 | 5.292 | 6.011 | 6.328 |
| D | - | - | - | - | 3.084 | 5.444 | 2.818 | 5.779 | 2.633 | 6.908 | 6.45 | 3.134 | 5.334 | 5.876 |
| E | - | - | - | - | - | 5.321 | -0.864 | 5.807 | 0.781 | 7.083 | 6.63 | -1.457 | 5.124 | 5.743 |
| F | - | - | - | - | - | - | 5.77 | 4.74 | 3.283 | 7.883 | 5.742 | 5.666 | 0.9 | 4.682 |
| G | - | - | - | - | - | - | - | 5.665 | 4.176 | 6.981 | 6.44 | 1.058 | 5.123 | 5.805 |
| H | - | - | - | - | - | - | - | - | 3.634 | 8.392 | 2.112 | 6.751 | 5.483 | 3.653 |
| I | - | - | - | - | - | - | - | - | - | 8.233 | 7.11 | 6.397 | 3.124 | 5.891 |
| J | - | - | - | - | - | - | - | - | - | - | 7.142 | 6.15 | 6.476 | 6.614 |
| K | - | - | - | - | - | - | - | - | - | - | - | 6.871 | 5.73 | 4.284 |
| L | - | - | - | - | - | - | - | - | - | - | - | - | 5.096 | 5.752 |
| M | - | - | - | - | - | - | - | - | - | - | - | - | - | 4.569 |
| N | - | - | - | - | - | - | - | - | - | - | - | - | - | - |

| | |
|---|---|
| | Larger than 10 |
| | Between 8 and 9,99 |
| | Between 6 and 7,99 |
| | Between 4 and 5,99 |
| | Between 2 and 3,99 |
| | Between 0 and 1,99 |
| | NA (Calculation failed) |

| Median | A | B | C | D | E | F | G | H | I | J | K | L | M | N |
|---|---|---|---|---|---|---|---|---|---|---|---|---|---|---|
| A | - | -3.444 | -0.641 | 0.597 | 4.081 | 4.134 | 2.013 | 3.211 | 3.273 | 2.738 | 2.719 | 2.442 | 3.816 | 0.332 |
| B | - | - | -1.998 | 0.752 | 4.094 | 4.147 | 2.036 | 3.195 | 3.318 | 2.76 | 2.639 | 2.466 | 3.809 | -0.134 |
| C | - | - | - | 0.979 | 4.117 | 4.167 | 1.834 | 3.145 | 3.351 | 2.883 | 2.591 | 2.553 | 3.767 | -0.32 |
| D | - | - | - | - | 3.973 | 4.025 | 2.488 | 3.453 | 3.049 | 2.445 | 3.011 | 2.004 | 3.99 | 1.48 |
| E | - | - | - | - | - | -0.43 | 4.659 | 4.99 | 3.103 | 3.618 | 4.825 | 3.788 | 5.224 | 4.444 |
| F | - | - | - | - | - | - | 4.705 | 5.023 | 3.215 | 3.672 | 4.858 | 3.864 | 5.269 | 4.514 |
| G | - | - | - | - | - | - | - | 2.327 | 3.73 | 3.389 | 1.341 | 3.189 | 3.286 | 1.366 |
| H | - | - | - | - | - | - | - | - | 4.155 | 3.959 | 1.31 | 3.777 | 2.21 | 2.83 |
| I | - | - | - | - | - | - | - | - | - | 1.586 | 4.146 | 2.319 | 4.744 | 3.577 |
| J | - | - | - | - | - | - | - | - | - | - | 3.802 | 0.199 | 4.458 | 3.017 |
| K | - | - | - | - | - | - | - | - | - | - | - | 3.53 | 2.85 | 2.269 |
| L | - | - | - | - | - | - | - | - | - | - | - | - | 4.375 | 2.828 |
| M | - | - | - | - | - | - | - | - | - | - | - | - | - | 3.494 |
| N | - | - | - | - | - | - | - | - | - | - | - | - | - | - |

| Max | A | B | C | D | E | F | G | H | I | J | K | L | M | N |
|---|---|---|---|---|---|---|---|---|---|---|---|---|---|---|
| A | - | -1.461 | -0.904 | 0.207 | 4.062 | 4.05 | 2.274 | 3.694 | 3.105 | 2.872 | 3.562 | 2.299 | 4.097 | 2.305 |
| B | - | - | -5.49 | 0.228 | 4.063 | 4.053 | 2.432 | 3.716 | 2.805 | 2.865 | 3.517 | 2.254 | 4.131 | 2.193 |
| C | - | - | - | 0.506 | 4.088 | 4.07 | 2.212 | 3.686 | 2.656 | 3.013 | 3.519 | 2.365 | 4.094 | 2.273 |
| D | - | - | - | - | 3.988 | 3.967 | 2.566 | 3.832 | 3.151 | 2.69 | 3.64 | 1.967 | 4.2 | 2.548 |
| E | - | - | - | - | - | -2.371 | 4.695 | 5.177 | 3.406 | 3.497 | 5.097 | 3.82 | 5.342 | 4.673 |
| F | - | - | - | - | - | - | 4.666 | 5.149 | 3.195 | 3.453 | 5.036 | 3.785 | 5.335 | 4.658 |
| G | - | - | - | - | - | - | - | 2.965 | 3.569 | 3.533 | 2.736 | 3.213 | 3.552 | -3.788 |
| H | - | - | - | - | - | - | - | - | 4.278 | 4.199 | 1.26 | 4.029 | 1.512 | 3.016 |
| I | - | - | - | - | - | - | - | - | - | -0.698 | 4.638 | 1.716 | 4.787 | 3.952 |
| J | - | - | - | - | - | - | - | - | - | - | 4.4 | 1.433 | 4.749 | 3.741 |
| K | - | - | - | - | - | - | - | - | - | - | - | 3.84 | 2.373 | 2.591 |
| L | - | - | - | - | - | - | - | - | - | - | - | - | 4.535 | 3.341 |
| M | - | - | - | - | - | - | - | - | - | - | - | - | - | 3.413 |
| N | - | - | - | - | - | - | - | - | - | - | - | - | - | - |
