## Supplementary Data S2 for "Interlaboratory comparison of standardised metabolomics and lipidomics analyses in human and rodent blood using the MxP® Quant 500 kit": Hex3Cer(d18.1-26.1).pdf

**Amount of relative differences between laboratories**

| Min | A | B | C | D | E | F | G | H | I | J | K | L | M | N |
|---|---|---|---|---|---|---|---|---|---|---|---|---|---|---|
| A | - | 6.962 | 6.081 | 6.173 | 5.995 | NA | 7.487 | 4.519 | 5.266 | 3.894 | 3.155 | 8.778 | NA | 5.748 |
| B | - | - | 5.456 | 5.885 | 3.611 | NA | 5.264 | 1.698 | 3.657 | 4.178 | 3.152 | 7.896 | NA | 6.012 |
| C | - | - | - | 2.746 | 2.375 | NA | 6.291 | 5.68 | 6.995 | 6.591 | 5.291 | 7.34 | NA | 2.01 |
| D | - | - | - | - | 2.922 | NA | 4.842 | 7.058 | 4.856 | 5.373 | 5.493 | 6.936 | NA | 4.794 |
| E | - | - | - | - | - | NA | 6.783 | 8.554 | 5.336 | 5.416 | 11.675 | 6.58 | NA | 6.305 |
| F | - | - | - | - | - | - | NA | NA | NA | NA | NA | NA | NA | NA |
| G | - | - | - | - | - | - | - | 5.337 | 2.782 | 4.734 | 2.326 | 6.738 | NA | 3.519 |
| H | - | - | - | - | - | - | - | - | 5.246 | 8.074 | 5.501 | 10.204 | NA | 6.14 |
| I | - | - | - | - | - | - | - | - | - | 5.87 | 3.788 | 6.117 | NA | 3.209 |
| J | - | - | - | - | - | - | - | - | - | - | 4.214 | 7.808 | NA | 5.709 |
| K | - | - | - | - | - | - | - | - | - | - | - | 12.314 | NA | 1.021 |
| L | - | - | - | - | - | - | - | - | - | - | - | - | NA | 5.246 |
| M | - | - | - | - | - | - | - | - | - | - | - | - | - | NA |
| N | - | - | - | - | - | - | - | - | - | - | - | - | - | - |

| Median | A | B | C | D | E | F | G | H | I | J | K | L | M | N |
|---|---|---|---|---|---|---|---|---|---|---|---|---|---|---|
| A | - | 5.325 | 5.594 | 4.888 | 4.707 | NA | 4.322 | 3.385 | 2.382 | -0.127 | 4.023 | 4.158 | NA | 5.061 |
| B | - | - | 4.401 | -1.27 | 5.469 | NA | 4.027 | 3.647 | 4.439 | 4.671 | 5.177 | 5.305 | NA | 5.612 |
| C | - | - | - | 1.526 | 5.416 | NA | 4.018 | 3.49 | -1.505 | 4.463 | 4.83 | 5.117 | NA | 5.614 |
| D | - | - | - | - | 5.354 | NA | 3.556 | 6.196 | 4.433 | 4.251 | 5.189 | 5.017 | NA | 5.598 |
| E | - | - | - | - | - | NA | 5.397 | 7.586 | 4.671 | 5.297 | 8.551 | 4.176 | NA | 3.993 |
| F | - | - | - | - | - | - | NA | NA | NA | NA | NA | NA | NA | NA |
| G | - | - | - | - | - | - | - | 3.478 | 2.408 | 2.607 | 4.503 | 4.555 | NA | 5.389 |
| H | - | - | - | - | - | - | - | - | 3.345 | 4.594 | 3.182 | 3.871 | NA | 4.89 |
| I | - | - | - | - | - | - | - | - | - | 0.39 | 4.195 | 4.116 | NA | 5.183 |
| J | - | - | - | - | - | - | - | - | - | - | 3.973 | 4.243 | NA | 5.083 |
| K | - | - | - | - | - | - | - | - | - | - | - | 2.294 | NA | 4.46 |
| L | - | - | - | - | - | - | - | - | - | - | - | - | NA | 4.748 |
| M | - | - | - | - | - | - | - | - | - | - | - | - | - | NA |
| N | - | - | - | - | - | - | - | - | - | - | - | - | - | - |

| Max | A | B | C | D | E | F | G | H | I | J | K | L | M | N |
|---|---|---|---|---|---|---|---|---|---|---|---|---|---|---|
| A | - | 3.803 | 3.759 | 0.973 | 5.804 | NA | 4.456 | 4.626 | 4.855 | 3.264 | 5.45 | 5.23 | NA | 5.783 |
| B | - | - | 5.685 | 5.478 | 5.977 | NA | 4.941 | 5.831 | 5.831 | 4.956 | 6.043 | 5.686 | NA | 6.269 |
| C | - | - | - | -4.571 | 5.871 | NA | -2.833 | 6.664 | 7.026 | 5.088 | 6.675 | 5.432 | NA | 5.958 |
| D | - | - | - | - | 5.699 | NA | 2.871 | 6.576 | 3.811 | 1.112 | 4.764 | 5.287 | NA | 5.744 |
| E | - | - | - | - | - | NA | 6.361 | 8.346 | 6.541 | 6.66 | 11.442 | 4.522 | NA | -2.087 |
| F | - | - | - | - | - | - | NA | NA | NA | NA | NA | NA | NA | NA |
| G | - | - | - | - | - | - | - | 6.155 | 4.155 | 3.229 | 5.39 | 4.864 | NA | 5.629 |
| H | - | - | - | - | - | - | - | - | 4.198 | 7.329 | 6.095 | 6.046 | NA | 5.769 |
| I | - | - | - | - | - | - | - | - | - | 5.536 | 4.539 | 4.397 | NA | 5.428 |
| J | - | - | - | - | - | - | - | - | - | - | 5.729 | 4.878 | NA | 5.79 |
| K | - | - | - | - | - | - | - | - | - | - | - | 7.81 | NA | 4.807 |
| L | - | - | - | - | - | - | - | - | - | - | - | - | NA | 4.592 |
| M | - | - | - | - | - | - | - | - | - | - | - | - | - | NA |
| N | - | - | - | - | - | - | - | - | - | - | - | - | - | - |

| Colour | Legend |
|---|---|
| Red | Larger than 10 |
| Orange | Between 8 and 9,99 |
| Light orange | Between 6 and 7,99 |
| Light green | Between 4 and 5,99 |
| Green | Between 2 and 3,99 |
| Dark green | Between 0 and 1,99 |
| Grey | NA (Calculation failed) |
