## Supplementary Data S2 for "Interlaboratory comparison of standardised metabolomics and lipidomics analyses in human and rodent blood using the MxP® Quant 500 kit": HexCer(d16.1-22.0).pdf

**Amount of relative differences between laboratories**

| Min | A | B | C | D | E | F | G | H | I | J | K | L | M | N |
|---|---|---|---|---|---|---|---|---|---|---|---|---|---|---|
| A | - | 6.023 | 6.343 | 4.732 | 7.9 | 5.357 | 7.203 | 4.598 | 1.393 | 4.993 | 5.998 | 6.373 | 5.016 | 6.37 |
| B | - | - | 4.575 | 7.438 | 4.688 | 4.757 | 6.28 | 3.11 | 4.467 | 4.207 | 5.185 | 7.839 | 4.017 | 3.785 |
| C | - | - | - | 6.93 | 5.059 | 3.795 | 6.074 | 4.16 | 4.83 | 3.879 | 5.067 | 7.25 | 3.624 | 4.839 |
| D | - | - | - | - | 6.127 | 3.317 | 6.893 | 5.404 | 4.894 | 3.639 | 5.427 | 3.174 | 3.331 | 6.106 |
| E | - | - | - | - | - | 3.825 | 6.306 | 3.466 | 5.408 | 2.457 | 4.135 | 7.74 | 1.823 | 4.353 |
| F | - | - | - | - | - | - | 7.851 | 6.454 | 5.356 | 3.09 | 5.067 | 9.553 | 2.488 | 5.72 |
| G | - | - | - | - | - | - | - | 5.182 | 6.506 | 5.027 | 5.611 | 8.733 | 4.954 | 5.321 |
| H | - | - | - | - | - | - | - | - | 3.969 | 5.444 | 4.237 | 8.928 | 5.412 | 5.472 |
| I | - | - | - | - | - | - | - | - | - | 5.065 | 6.019 | 8.823 | 5.074 | 7.465 |
| J | - | - | - | - | - | - | - | - | - | - | 4.819 | 8.056 | 3.504 | 5.08 |
| K | - | - | - | - | - | - | - | - | - | - | - | 8.412 | 3.784 | 2.144 |
| L | - | - | - | - | - | - | - | - | - | - | - | - | 3.574 | 6.052 |
| M | - | - | - | - | - | - | - | - | - | - | - | - | - | 4.911 |
| N | - | - | - | - | - | - | - | - | - | - | - | - | - | - |

| | |
|---|---|
| | Larger than 10 |
| | Between 8 and 9,99 |
| | Between 6 and 7,99 |
| | Between 4 and 5,99 |
| | Between 2 and 3,99 |
| | Between 0 and 1,99 |
| | NA (Calculation failed) |

| Median | A | B | C | D | E | F | G | H | I | J | K | L | M | N |
|---|---|---|---|---|---|---|---|---|---|---|---|---|---|---|
| A | - | 3.074 | 1.875 | 3.472 | 3.504 | 1.373 | 4.16 | 4.823 | -2.558 | 4.224 | 2.264 | 2.533 | 3.428 | 3.47 |
| B | - | - | 1.846 | 1.557 | 2.009 | 2.51 | 3.31 | 5.261 | 3.135 | 3.491 | 3.742 | 1.33 | 2.032 | 1.477 |
| C | - | - | - | 2.721 | 2.84 | 1.545 | 3.776 | 5.056 | 2.546 | 3.735 | 3.24 | 0.271 | 2.881 | 2.697 |
| D | - | - | - | - | -5.012 | 3.425 | 2.868 | 5.458 | 3.777 | 2.819 | 4.144 | 2.588 | -4.605 | -3.998 |
| E | - | - | - | - | - | 3.474 | 2.9 | 5.499 | 3.861 | 2.603 | 4.169 | 2.711 | -1.133 | -0.265 |
| F | - | - | - | - | - | - | 4.213 | 5.005 | 1.539 | 3.935 | 2.607 | 1.92 | 3.558 | 3.099 |
| G | - | - | - | - | - | - | - | 5.73 | 4.63 | 2.073 | 4.715 | 3.854 | 2.64 | 3.129 |
| H | - | - | - | - | - | - | - | - | 4.515 | 5.215 | 4.224 | 4.699 | 4.99 | 4.865 |
| I | - | - | - | - | - | - | - | - | - | 4.24 | 2.233 | 2.819 | 3.428 | 3.71 |
| J | - | - | - | - | - | - | - | - | - | - | 4.723 | 3.771 | 2.579 | 2.874 |
| K | - | - | - | - | - | - | - | - | - | - | - | 3.214 | 4.003 | 3.789 |
| L | - | - | - | - | - | - | - | - | - | - | - | - | 2.531 | 2.486 |
| M | - | - | - | - | - | - | - | - | - | - | - | - | - | 0.06 |
| N | - | - | - | - | - | - | - | - | - | - | - | - | - | - |

| Max | A | B | C | D | E | F | G | H | I | J | K | L | M | N |
|---|---|---|---|---|---|---|---|---|---|---|---|---|---|---|
| A | - | 2.461 | 1.362 | 3.429 | 2.766 | 3.779 | 3.743 | 5.134 | -0.985 | 3.624 | 3.289 | 2.362 | 2.888 | 2.869 |
| B | - | - | 1.726 | 2.731 | 1.845 | 4.294 | 2.921 | 5.473 | 2.356 | 2.956 | 4.086 | -0.693 | 1.143 | 1.14 |
| C | - | - | - | 3.215 | 2.731 | 3.331 | 3.566 | 5.318 | -1.428 | 3.66 | 3.651 | 1.316 | 2.721 | 2.411 |
| D | - | - | - | - | 0.448 | 4.289 | 1.781 | 5.785 | 3.219 | 2.155 | 4.464 | 2.618 | 0.688 | 1.886 |
| E | - | - | - | - | - | 4.45 | 2.316 | 5.634 | 2.778 | 2.668 | 4.362 | 2.268 | 0.107 | 0.966 |
| F | - | - | - | - | - | - | 3.484 | 4.64 | 2.915 | 4.215 | 2.038 | 3.336 | 3.856 | 3.561 |
| G | - | - | - | - | - | - | - | 5.787 | 2.723 | 3.249 | 4.622 | 3.462 | 1.254 | 2.828 |
| H | - | - | - | - | - | - | - | - | 4.635 | 5.089 | 4.223 | 4.92 | 4.912 | 5.045 |
| I | - | - | - | - | - | - | - | - | - | 3.569 | 3.283 | 1.733 | 2.847 | 2.156 |
| J | - | - | - | - | - | - | - | - | - | - | 4.715 | 3.529 | 2.362 | 3.248 |
| K | - | - | - | - | - | - | - | - | - | - | - | 3.609 | 4.035 | 3.886 |
| L | - | - | - | - | - | - | - | - | - | - | - | - | 2.27 | 1.261 |
| M | - | - | - | - | - | - | - | - | - | - | - | - | - | 1.424 |
| N | - | - | - | - | - | - | - | - | - | - | - | - | - | - |
