## Supplementary Data S2 for "Interlaboratory comparison of standardised metabolomics and lipidomics analyses in human and rodent blood using the MxP® Quant 500 kit": HexCer(d16.1-24.0).pdf

**Amount of relative differences between laboratories**

| Min | A | B | C | D | E | F | G | H | I | J | K | L | M | N |
|---|---|---|---|---|---|---|---|---|---|---|---|---|---|---|
| A | - | 4.764 | 3.302 | 3.497 | 6.559 | 5.944 | 5.558 | NA | 4.727 | 5.616 | 6.092 | 4.577 | 5.288 | 5.949 |
| B | - | - | 4.613 | 5.146 | 7.281 | 5.59 | 5.295 | NA | 5.599 | 2.144 | 5.688 | 5.489 | 0.96 | 5.504 |
| C | - | - | - | 6.371 | 7.896 | 5.395 | 4.483 | NA | 7.661 | 5.051 | 6.248 | 6.635 | 6.854 | 5.94 |
| D | - | - | - | - | 6.083 | 5.927 | 5.555 | NA | 4.191 | 4.558 | 5.999 | 1.366 | 2.914 | 5.928 |
| E | - | - | - | - | - | 6.386 | 6.168 | NA | 4.553 | 2.805 | 6.374 | 5.64 | 4.899 | 6.381 |
| F | - | - | - | - | - | - | 5.511 | NA | 7.314 | 5.602 | 3.052 | 7.844 | 7.023 | 2.161 |
| G | - | - | - | - | - | - | - | NA | 6.177 | 2.851 | 4.259 | 6.69 | 5.838 | 4.72 |
| H | - | - | - | - | - | - | - | - | NA | NA | NA | NA | NA | NA |
| I | - | - | - | - | - | - | - | - | - | 2.335 | 6.501 | 3.125 | 4.622 | 6.16 |
| J | - | - | - | - | - | - | - | - | - | - | 7.406 | 6.676 | 5.831 | 4.229 |
| K | - | - | - | - | - | - | - | - | - | - | - | 7.144 | 6.276 | 2.895 |
| L | - | - | - | - | - | - | - | - | - | - | - | - | 0.833 | 6.227 |
| M | - | - | - | - | - | - | - | - | - | - | - | - | - | 6.132 |
| N | - | - | - | - | - | - | - | - | - | - | - | - | - | - |

| | |
|---|---|
| | Larger than 10 |
| | Between 8 and 9,99 |
| | Between 6 and 7,99 |
| | Between 4 and 5,99 |
| | Between 2 and 3,99 |
| | Between 0 and 1,99 |
| | NA (Calculation failed) |

| Median | A | B | C | D | E | F | G | H | I | J | K | L | M | N |
|---|---|---|---|---|---|---|---|---|---|---|---|---|---|---|
| A | - | 3.8 | 0.382 | 0.206 | 3.952 | 2.333 | 5.145 | NA | 2.391 | 4.81 | 6.18 | 2.782 | 1.625 | 2.606 |
| B | - | - | 3.509 | 3.431 | 0.662 | 4.004 | 4.326 | NA | 4.028 | 3.724 | 5.707 | 2.47 | 3.85 | 4.093 |
| C | - | - | - | -3.893 | 3.872 | 2.492 | 5.131 | NA | 2.165 | 4.749 | 6.121 | 2.572 | 2.145 | 1.96 |
| D | - | - | - | - | 3.833 | 2.673 | 5.066 | NA | 2.682 | 4.753 | 6.126 | 2.539 | 1.955 | 2.891 |
| E | - | - | - | - | - | 4.172 | 4.058 | NA | 4.182 | 3.558 | 5.603 | 2.855 | 3.963 | 4.244 |
| F | - | - | - | - | - | - | 5.404 | NA | -0.748 | 5.045 | 6.327 | 3.434 | -3.997 | 0.308 |
| G | - | - | - | - | - | - | - | NA | 4.909 | 2.715 | 4.714 | 4.436 | 4.881 | 4.939 |
| H | - | - | - | - | - | - | - | - | NA | NA | NA | NA | NA | NA |
| I | - | - | - | - | - | - | - | - | - | 5.179 | 6.367 | 3.686 | 1.621 | 0.168 |
| J | - | - | - | - | - | - | - | - | - | - | 5.028 | 4.065 | 4.627 | 4.616 |
| K | - | - | - | - | - | - | - | - | - | - | - | 5.25 | 5.492 | 5.489 |
| L | - | - | - | - | - | - | - | - | - | - | - | - | 3.203 | 3.592 |
| M | - | - | - | - | - | - | - | - | - | - | - | - | - | 1.95 |
| N | - | - | - | - | - | - | - | - | - | - | - | - | - | - |

| Max | A | B | C | D | E | F | G | H | I | J | K | L | M | N |
|---|---|---|---|---|---|---|---|---|---|---|---|---|---|---|
| A | - | 3.996 | -1.381 | -0.058 | 3.861 | -0.226 | 5.37 | NA | 2.677 | 4.634 | 6.419 | 2.729 | 1.881 | -2.038 |
| B | - | - | 3.138 | 3.544 | -1.895 | 3.666 | 4.606 | NA | 4.222 | 3.993 | 5.952 | 2.65 | 3.873 | 3.817 |
| C | - | - | - | 0.592 | 3.579 | 0.867 | 5.168 | NA | 3.583 | 4.696 | 6.377 | 2.198 | 2.615 | 3.425 |
| D | - | - | - | - | 3.766 | -0.519 | 5.298 | NA | 2.87 | 4.781 | 6.362 | 2.55 | 2.014 | 0.853 |
| E | - | - | - | - | - | 3.483 | 4.528 | NA | 4.166 | 3.874 | 5.926 | 2.726 | 3.944 | 3.607 |
| F | - | - | - | - | - | - | 5.396 | NA | 2.674 | 4.937 | 6.443 | 2.932 | 0.933 | 0.878 |
| G | - | - | - | - | - | - | - | NA | 5.077 | 3.118 | 4.928 | 4.552 | 4.94 | 4.96 |
| H | - | - | - | - | - | - | - | - | NA | NA | NA | NA | NA | NA |
| I | - | - | - | - | - | - | - | - | - | 5.412 | 6.623 | 3.721 | 1.794 | 2.648 |
| J | - | - | - | - | - | - | - | - | - | - | 5.683 | 4.207 | 4.693 | 4.861 |
| K | - | - | - | - | - | - | - | - | - | - | - | 5.345 | 5.546 | 5.61 |
| L | - | - | - | - | - | - | - | - | - | - | - | - | 3.219 | 2.45 |
| M | - | - | - | - | - | - | - | - | - | - | - | - | - | 1.628 |
| N | - | - | - | - | - | - | - | - | - | - | - | - | - | - |
