## Supplementary Data S2 for "Interlaboratory comparison of standardised metabolomics and lipidomics analyses in human and rodent blood using the MxP® Quant 500 kit": HexCer(d18.1-23.0).pdf

**Amount of relative differences between laboratories**

| Min | A | B | C | D | E | F | G | H | I | J | K | L | M | N |
|---|---|---|---|---|---|---|---|---|---|---|---|---|---|---|
| A | - | 3.818 | 3.661 | 1.702 | 4.375 | 3.924 | 5.466 | 4.066 | 2.658 | -0.566 | 4.896 | 5.205 | 3.212 | 3.563 |
| B | - | - | 1.053 | 4.376 | 2.95 | -0.129 | 4.625 | 1.318 | 3.143 | 4.11 | 3.497 | 3.953 | 2.567 | 1.859 |
| C | - | - | - | 4.183 | 3.268 | 1.55 | 4.767 | 2.164 | 2.661 | 3.915 | 3.71 | 4.106 | 1.933 | 0.613 |
| D | - | - | - | - | 4.576 | 4.125 | 5.363 | 4.277 | 3.24 | 1.306 | 4.835 | 5.214 | 3.63 | 3.777 |
| E | - | - | - | - | - | 3.021 | 4.288 | 2.355 | 4.152 | 4.771 | 3.006 | 3.823 | 3.929 | 3.707 |
| F | - | - | - | - | - | - | 4.805 | 0.983 | 3.313 | 4.186 | 3.702 | 4.156 | 2.693 | 2.068 |
| G | - | - | - | - | - | - | - | 4.037 | 4.881 | 5.258 | 3.648 | 3.781 | 4.809 | 4.555 |
| H | - | - | - | - | - | - | - | - | 3.56 | 4.352 | 2.873 | 3.811 | 3.161 | 2.659 |
| I | - | - | - | - | - | - | - | - | - | 2.985 | 4.466 | 4.842 | 1.585 | 2.35 |
| J | - | - | - | - | - | - | - | - | - | - | 5.106 | 5.376 | 3.528 | 3.756 |
| K | - | - | - | - | - | - | - | - | - | - | - | 0.817 | 3.99 | 3.511 |
| L | - | - | - | - | - | - | - | - | - | - | - | - | 4.208 | 3.88 |
| M | - | - | - | - | - | - | - | - | - | - | - | - | - | 1.16 |
| N | - | - | - | - | - | - | - | - | - | - | - | - | - | - |

| | |
|---|---|
| | Larger than 10 |
| | Between 8 and 9,99 |
| | Between 6 and 7,99 |
| | Between 4 and 5,99 |
| | Between 2 and 3,99 |
| | Between 0 and 1,99 |
| | NA (Calculation failed) |

| Median | A | B | C | D | E | F | G | H | I | J | K | L | M | N |
|---|---|---|---|---|---|---|---|---|---|---|---|---|---|---|
| A | - | 2.238 | 1.677 | 2.381 | 3.769 | 3.308 | 2.412 | 2.716 | 1.991 | 2.924 | 2.03 | 1.327 | 2.192 | 3.265 |
| B | - | - | 1.145 | 3.432 | 3.101 | 2.336 | 3.249 | 0.975 | 0.653 | 1.327 | 3.115 | 1.137 | 3.393 | 2.28 |
| C | - | - | - | 3.071 | 3.4 | 2.829 | 2.86 | 2.033 | -0.695 | 2.21 | 2.656 | -6.351 | 3.02 | 2.79 |
| D | - | - | - | - | 4.156 | 3.846 | -0.93 | 3.491 | 3.114 | 3.602 | 0.572 | 2.891 | -0.88 | 3.822 |
| E | - | - | - | - | - | 1.951 | 4.323 | 2.884 | 3.475 | 2.738 | 4.279 | 3.607 | 4.384 | 2.029 |
| F | - | - | - | - | - | - | 3.998 | 1.795 | 2.768 | 1.402 | 3.896 | 2.964 | 4.023 | -1.923 |
| G | - | - | - | - | - | - | - | 3.268 | 2.093 | 3.13 | -1.611 | 2.411 | 1.942 | 3.517 |
| H | - | - | - | - | - | - | - | - | 1.867 | -0.904 | 3.414 | 2.11 | 3.673 | 1.553 |
| I | - | - | - | - | - | - | - | - | - | 1.942 | 2.982 | 0.293 | 3.149 | 2.589 |
| J | - | - | - | - | - | - | - | - | - | - | 3.744 | 2.52 | 3.705 | 1 |
| K | - | - | - | - | - | - | - | - | - | - | - | 2.417 | 1.655 | 3.541 |
| L | - | - | - | - | - | - | - | - | - | - | - | - | 3.143 | 2.744 |
| M | - | - | - | - | - | - | - | - | - | - | - | - | - | 3.811 |
| N | - | - | - | - | - | - | - | - | - | - | - | - | - | - |

| Max | A | B | C | D | E | F | G | H | I | J | K | L | M | N |
|---|---|---|---|---|---|---|---|---|---|---|---|---|---|---|
| A | - | 1.746 | -0.29 | 2.463 | 3.553 | 3.149 | 3.245 | 2.302 | 1.781 | 3.222 | 2.977 | -0.037 | 3.024 | 3.2 |
| B | - | - | 1.167 | 3.209 | 3.139 | 2.552 | 3.63 | 0.911 | -3.315 | 2.741 | 3.446 | 1.997 | 3.546 | 2.697 |
| C | - | - | - | 2.784 | 3.433 | 2.987 | 3.345 | 2.011 | 1.148 | 3.136 | 3.083 | 0.793 | 3.205 | 3.094 |
| D | - | - | - | - | 4.021 | 3.784 | 1.532 | 3.313 | 3.084 | 3.826 | 0.627 | 2.139 | 1.325 | 3.83 |
| E | - | - | - | - | - | 1.601 | 4.545 | 2.954 | 3.26 | 1.209 | 4.491 | 3.893 | 4.481 | 1.241 |
| F | - | - | - | - | - | - | 4.298 | 2.12 | 2.606 | -0.634 | 4.171 | 3.379 | 4.231 | -1.032 |
| G | - | - | - | - | - | - | - | 3.789 | 3.589 | 4.224 | 0.822 | 2.858 | 1.256 | 4.181 |
| H | - | - | - | - | - | - | - | - | 0.825 | 2.311 | 3.663 | 2.627 | 3.78 | 2.235 |
| I | - | - | - | - | - | - | - | - | - | 2.661 | 3.494 | 2.09 | 3.576 | 2.63 |
| J | - | - | - | - | - | - | - | - | - | - | 4.229 | 3.396 | 4.276 | -2.28 |
| K | - | - | - | - | - | - | - | - | - | - | - | 2.512 | -2.947 | 4.005 |
| L | - | - | - | - | - | - | - | - | - | - | - | - | 2.667 | 3.434 |
| M | - | - | - | - | - | - | - | - | - | - | - | - | - | 4.027 |
| N | - | - | - | - | - | - | - | - | - | - | - | - | - | - |
