## Supplementary Data S2 for "Interlaboratory comparison of standardised metabolomics and lipidomics analyses in human and rodent blood using the MxP® Quant 500 kit": HexCer(d18.2-20.0).pdf

**Amount of relative differences between laboratories**

| Min | A | B | C | D | E | F | G | H | I | J | K | L | M | N |
|---|---|---|---|---|---|---|---|---|---|---|---|---|---|---|
| A | - | 3.889 | 5.233 | 4.345 | 0.199 | 5.51 | NA | NA | 4.199 | 4.317 | NA | NA | 5.564 | 4.352 |
| B | - | - | 4.666 | 0.626 | 5.129 | 4.675 | NA | NA | 3.435 | -0.37 | NA | NA | 3.794 | 2.89 |
| C | - | - | - | 3.974 | 6.232 | 0.929 | NA | NA | 5.093 | 4.241 | NA | NA | 5.984 | 4.732 |
| D | - | - | - | - | 4.601 | 4.184 | NA | NA | 3.533 | 2.988 | NA | NA | 5.604 | 2.655 |
| E | - | - | - | - | - | 5.238 | NA | NA | 4.49 | 2.51 | NA | NA | 4.451 | 4.354 |
| F | - | - | - | - | - | - | NA | NA | 2.74 | 2.121 | NA | NA | 6.657 | 2.162 |
| G | - | - | - | - | - | - | - | NA | NA | NA | NA | NA | NA | NA |
| H | - | - | - | - | - | - | - | - | NA | NA | NA | NA | NA | NA |
| I | - | - | - | - | - | - | - | - | - | 4.12 | NA | NA | 0.561 | 1.975 |
| J | - | - | - | - | - | - | - | - | - | - | NA | NA | 4.587 | 3.663 |
| K | - | - | - | - | - | - | - | - | - | - | - | NA | NA | NA |
| L | - | - | - | - | - | - | - | - | - | - | - | - | NA | NA |
| M | - | - | - | - | - | - | - | - | - | - | - | - | - | 1.725 |
| N | - | - | - | - | - | - | - | - | - | - | - | - | - | - |

| | |
|---|---|
| | Larger than 10 |
| | Between 8 and 9,99 |
| | Between 6 and 7,99 |
| | Between 4 and 5,99 |
| | Between 2 and 3,99 |
| | Between 0 and 1,99 |
| | NA (Calculation failed) |

| Median | A | B | C | D | E | F | G | H | I | J | K | L | M | N |
|---|---|---|---|---|---|---|---|---|---|---|---|---|---|---|
| A | - | 3.318 | 4.535 | 4.004 | 4.066 | 0.832 | NA | NA | -1.343 | 3.149 | NA | NA | 1.698 | 3.284 |
| B | - | - | 3.749 | 2.05 | 2.641 | 3.7 | NA | NA | 4.053 | 2.181 | NA | NA | 3.424 | 4.54 |
| C | - | - | - | 3.104 | 2.563 | 4.844 | NA | NA | 5.061 | 4.263 | NA | NA | 4.668 | 5.266 |
| D | - | - | - | - | 2.133 | 4.369 | NA | NA | 4.405 | 2.773 | NA | NA | 3.582 | 4.874 |
| E | - | - | - | - | - | 4.231 | NA | NA | 4.789 | 3.753 | NA | NA | 4.11 | 5.111 |
| F | - | - | - | - | - | - | NA | NA | -1.496 | 3.344 | NA | NA | 1.98 | 1.167 |
| G | - | - | - | - | - | - | - | NA | NA | NA | NA | NA | NA | NA |
| H | - | - | - | - | - | - | - | - | NA | NA | NA | NA | NA | NA |
| I | - | - | - | - | - | - | - | - | - | 3.592 | NA | NA | 2.919 | 2.009 |
| J | - | - | - | - | - | - | - | - | - | - | NA | NA | 1.909 | 4.01 |
| K | - | - | - | - | - | - | - | - | - | - | - | NA | NA | NA |
| L | - | - | - | - | - | - | - | - | - | - | - | - | NA | NA |
| M | - | - | - | - | - | - | - | - | - | - | - | - | - | 3.684 |
| N | - | - | - | - | - | - | - | - | - | - | - | - | - | - |

| Max | A | B | C | D | E | F | G | H | I | J | K | L | M | N |
|---|---|---|---|---|---|---|---|---|---|---|---|---|---|---|
| A | - | 3.111 | 3.875 | 3.455 | 4.196 | 5.063 | NA | NA | 4.629 | 3.499 | NA | NA | 1.472 | 4.995 |
| B | - | - | 2.68 | 2.855 | 3.377 | 5.276 | NA | NA | 4.625 | 3.829 | NA | NA | 3.365 | 5.125 |
| C | - | - | - | -2.902 | 0.093 | 5.62 | NA | NA | 5.015 | 4.302 | NA | NA | 4.34 | 5.562 |
| D | - | - | - | - | 1.254 | 5.542 | NA | NA | 5.113 | 5.077 | NA | NA | 4.34 | 5.531 |
| E | - | - | - | - | - | 5.821 | NA | NA | 5.125 | 4.791 | NA | NA | 4.499 | 5.489 |
| F | - | - | - | - | - | - | NA | NA | 3.086 | 4.374 | NA | NA | 4.191 | -0.718 |
| G | - | - | - | - | - | - | - | NA | NA | NA | NA | NA | NA | NA |
| H | - | - | - | - | - | - | - | - | NA | NA | NA | NA | NA | NA |
| I | - | - | - | - | - | - | - | - | - | 1.219 | NA | NA | 3.069 | 3.303 |
| J | - | - | - | - | - | - | - | - | - | - | NA | NA | -0.018 | 4.221 |
| K | - | - | - | - | - | - | - | - | - | - | - | NA | NA | NA |
| L | - | - | - | - | - | - | - | - | - | - | - | - | NA | NA |
| M | - | - | - | - | - | - | - | - | - | - | - | - | - | 4.313 |
| N | - | - | - | - | - | - | - | - | - | - | - | - | - | - |
