## Supplementary Data S2 for "Interlaboratory comparison of standardised metabolomics and lipidomics analyses in human and rodent blood using the MxP® Quant 500 kit": HexCer(d18.2-24.0).pdf

**Amount of relative differences between laboratories**

| Min | A | B | C | D | E | F | G | H | I | J | K | L | M | N |
|---|---|---|---|---|---|---|---|---|---|---|---|---|---|---|
| A | - | 4.385 | 3.642 | 0.409 | 4.472 | 5.044 | 2.979 | 5.532 | 3.595 | 0.369 | 5.706 | 5.236 | 4.657 | 4.765 |
| B | - | - | 2.731 | 4.65 | -0.515 | 4.356 | 2.561 | 5.062 | 3.286 | 3.936 | 5.35 | 4.602 | 2.29 | 2.756 |
| C | - | - | - | 3.679 | 3.609 | 4.953 | 0.412 | 5.24 | 0.978 | 2.107 | 5.625 | 4.934 | 3.58 | 3.942 |
| D | - | - | - | - | 4.589 | 5.207 | 3.39 | 5.539 | 3.61 | 2.646 | 5.734 | 5.252 | 4.542 | 4.648 |
| E | - | - | - | - | - | 4.6 | 3.248 | 4.904 | 3.239 | 3.996 | 5.327 | 4.536 | 0.462 | 0.557 |
| F | - | - | - | - | - | - | 5.148 | 3.082 | 5.684 | 5.713 | 4.95 | 2.003 | 5.077 | 4.864 |
| G | - | - | - | - | - | - | - | 4.867 | 3.389 | 3.702 | 5.312 | 4.6 | 1.88 | 0.26 |
| H | - | - | - | - | - | - | - | - | 5.676 | 5.638 | 3.978 | 2.314 | 5.512 | 5.396 |
| I | - | - | - | - | - | - | - | - | - | 2.961 | 5.42 | 4.858 | 3.462 | 4.077 |
| J | - | - | - | - | - | - | - | - | - | - | 5.706 | 5.155 | 4.726 | 4.854 |
| K | - | - | - | - | - | - | - | - | - | - | - | 4.818 | 6.286 | 6.244 |
| L | - | - | - | - | - | - | - | - | - | - | - | - | 5.416 | 5.249 |
| M | - | - | - | - | - | - | - | - | - | - | - | - | - | 1.441 |
| N | - | - | - | - | - | - | - | - | - | - | - | - | - | - |

| | |
|---|---|
| | Larger than 10 |
| | Between 8 and 9,99 |
| | Between 6 and 7,99 |
| | Between 4 and 5,99 |
| | Between 2 and 3,99 |
| | Between 0 and 1,99 |
| | NA (Calculation failed) |

| Median | A | B | C | D | E | F | G | H | I | J | K | L | M | N |
|---|---|---|---|---|---|---|---|---|---|---|---|---|---|---|
| A | - | 0.933 | 1.34 | 3.553 | 2.05 | 3.922 | 4.944 | 1.932 | 2.526 | 3.129 | 1.84 | 4.141 | 1.103 | 2.667 |
| B | - | - | -0.008 | 3.343 | 2.448 | 4.005 | 4.881 | 1.321 | 2.866 | 3.366 | 2.275 | 3.995 | -0.421 | 2.234 |
| C | - | - | - | 3.182 | 2.678 | 4.092 | 4.806 | 0.511 | 3.019 | 3.462 | 2.537 | 3.889 | -1.843 | 1.894 |
| D | - | - | - | - | 3.826 | 4.583 | 4.111 | 2.814 | 3.976 | 4.202 | 3.772 | 2.396 | 3.1 | 2.292 |
| E | - | - | - | - | - | 3.502 | 5.204 | 3.068 | 0.898 | 2.327 | -0.645 | 4.502 | 2.707 | 3.422 |
| F | - | - | - | - | - | - | 5.764 | 4.46 | 3.329 | 2.683 | 3.733 | 5.241 | 4.309 | 4.608 |
| G | - | - | - | - | - | - | - | 4.404 | 4.862 | 4.98 | 4.747 | 3.47 | 4.503 | 4.314 |
| H | - | - | - | - | - | - | - | - | 3.347 | 3.735 | 2.853 | 3.664 | 0.975 | 0.529 |
| I | - | - | - | - | - | - | - | - | - | 1.652 | 1.255 | 4.637 | 3.078 | 3.687 |
| J | - | - | - | - | - | - | - | - | - | - | 2.457 | 4.845 | 3.559 | 4.041 |
| K | - | - | - | - | - | - | - | - | - | - | - | 4.41 | 2.483 | 3.178 |
| L | - | - | - | - | - | - | - | - | - | - | - | - | 3.698 | 3.329 |
| M | - | - | - | - | - | - | - | - | - | - | - | - | - | 1.96 |
| N | - | - | - | - | - | - | - | - | - | - | - | - | - | - |

| Max | A | B | C | D | E | F | G | H | I | J | K | L | M | N |
|---|---|---|---|---|---|---|---|---|---|---|---|---|---|---|
| A | - | 1.807 | 1.913 | 3.665 | 1.378 | 3.426 | 5.319 | 3.566 | 2.405 | 3.229 | 3.028 | 4.852 | 2.17 | 3.196 |
| B | - | - | -0.915 | 3.135 | 2.552 | 3.904 | 5.131 | 3.044 | 3.096 | 3.639 | 1.941 | 4.592 | 0.3 | 2.502 |
| C | - | - | - | 3.126 | 2.561 | 3.767 | 5.125 | 2.947 | 3.097 | 3.622 | 2.235 | 4.587 | -0.466 | 2.41 |
| D | - | - | - | - | 3.737 | 4.377 | 4.576 | -0.498 | 4 | 4.28 | 2.041 | 3.814 | 2.87 | 1.681 |
| E | - | - | - | - | - | 3.026 | 5.437 | 3.861 | 1.431 | 2.761 | 3.391 | 4.999 | 2.801 | 3.553 |
| F | - | - | - | - | - | - | 5.878 | 4.749 | 2.808 | 1.354 | 4.583 | 5.481 | 4.219 | 4.584 |
| G | - | - | - | - | - | - | - | 4.314 | 5 | 5.142 | 4.57 | 3.072 | 4.64 | 4.437 |
| H | - | - | - | - | - | - | - | - | 3.877 | 4.219 | 2.37 | 3.832 | 2.598 | 0.976 |
| I | - | - | - | - | - | - | - | - | - | 2.022 | 3.733 | 5.163 | 3.339 | 3.943 |
| J | - | - | - | - | - | - | - | - | - | - | 4.279 | 5.386 | 3.914 | 4.341 |
| K | - | - | - | - | - | - | - | - | - | - | - | 4.311 | -0.66 | 1.788 |
| L | - | - | - | - | - | - | - | - | - | - | - | - | 4.151 | 3.816 |
| M | - | - | - | - | - | - | - | - | - | - | - | - | - | 2.152 |
| N | - | - | - | - | - | - | - | - | - | - | - | - | - | - |
