## Supplementary Data S2 for "Interlaboratory comparison of standardised metabolomics and lipidomics analyses in human and rodent blood using the MxP® Quant 500 kit": Ind-SO4.pdf

**Amount of relative differences between laboratories**

| Min | A | B | C | D | E | F | G | H | I | J | K | L | M | N |
|---|---|---|---|---|---|---|---|---|---|---|---|---|---|---|
| A | - | 4.122 | 2.869 | 4.754 | 2.896 | 3.936 | 3.831 | 4.926 | 3.651 | 0.998 | 4.507 | 2.507 | 4.276 | 0.455 |
| B | - | - | 3.399 | 3.579 | 4.529 | 0.247 | 0.018 | 3.719 | 2.249 | 4.014 | 2.434 | 3.063 | 0.933 | 4.046 |
| C | - | - | - | 4.357 | 3.786 | 3.069 | 2.898 | 4.528 | 2.418 | 2.654 | 3.935 | -0.906 | 3.571 | 2.542 |
| D | - | - | - | - | 4.951 | 3.512 | 3.57 | 0.611 | 3.911 | 4.871 | 2.637 | 4.178 | 3.19 | 4.695 |
| E | - | - | - | - | - | 4.655 | 4.551 | 5.382 | 4.457 | 3.066 | 5.045 | 3.845 | 4.865 | 3.232 |
| F | - | - | - | - | - | - | -1.788 | 3.871 | 1.972 | 3.845 | 2.746 | 2.827 | 1.641 | 3.981 |
| G | - | - | - | - | - | - | - | 3.828 | 1.419 | 3.624 | 2.788 | 2.895 | 1.868 | 3.774 |
| H | - | - | - | - | - | - | - | - | 4.221 | 4.925 | 3.019 | 4.262 | 3.579 | 4.957 |
| I | - | - | - | - | - | - | - | - | - | 3.544 | 3.307 | 2.29 | 2.696 | 3.498 |
| J | - | - | - | - | - | - | - | - | - | - | 4.186 | -0.209 | 3.873 | 0.455 |
| K | - | - | - | - | - | - | - | - | - | - | - | 3.633 | 1.953 | 4.564 |
| L | - | - | - | - | - | - | - | - | - | - | - | - | 3.535 | 2.984 |
| M | - | - | - | - | - | - | - | - | - | - | - | - | - | 4.191 |
| N | - | - | - | - | - | - | - | - | - | - | - | - | - | - |

| Median | A | B | C | D | E | F | G | H | I | J | K | L | M | N |
|---|---|---|---|---|---|---|---|---|---|---|---|---|---|---|
| A | - | 0.865 | 1.437 | 3.723 | 2.789 | 0.158 | 2.583 | 0.965 | 1.805 | 3.816 | 1.625 | 2.063 | 1.016 | 2.249 |
| B | - | - | 0.789 | 3.529 | 3.009 | 1.315 | 2.247 | -2.435 | 1.292 | 3.806 | 2.188 | 2.55 | -0.2 | 2.517 |
| C | - | - | - | 3.348 | 3.237 | 1.926 | 1.648 | -0.183 | -0.396 | 3.489 | 2.529 | 2.787 | -0.484 | 2.865 |
| D | - | - | - | - | 4.127 | 3.645 | 2.616 | 3.336 | 3.025 | 0.852 | 3.856 | 3.963 | 3.258 | 3.955 |
| E | - | - | - | - | - | 2.668 | 3.796 | 3.282 | 3.489 | 4.479 | 2.074 | 1.599 | 3.268 | 1.192 |
| F | - | - | - | - | - | - | 2.895 | 1.434 | 2.341 | 4.101 | 1.092 | 1.797 | 1.796 | 1.714 |
| G | - | - | - | - | - | - | - | 2.049 | 1.17 | 3.076 | 3.129 | 3.308 | 1.894 | 3.33 |
| H | - | - | - | - | - | - | - | - | 1.507 | 3.947 | 2.195 | 2.601 | 0.221 | 2.425 |
| I | - | - | - | - | - | - | - | - | - | 3.42 | 2.727 | 2.962 | 0.581 | 2.998 |
| J | - | - | - | - | - | - | - | - | - | - | 3.822 | 3.912 | 3.318 | 4.007 |
| K | - | - | - | - | - | - | - | - | - | - | - | 0.605 | 2.571 | -0.127 |
| L | - | - | - | - | - | - | - | - | - | - | - | - | 2.809 | -5.229 |
| M | - | - | - | - | - | - | - | - | - | - | - | - | - | 2.706 |
| N | - | - | - | - | - | - | - | - | - | - | - | - | - | - |

| Max | A | B | C | D | E | F | G | H | I | J | K | L | M | N |
|---|---|---|---|---|---|---|---|---|---|---|---|---|---|---|
| A | - | 3.042 | -6.623 | 1.112 | 2.714 | 3.5 | -5.172 | 3.954 | 0.972 | 4.425 | 4.137 | 3.565 | 3 | 3.018 |
| B | - | - | 3.159 | 3.488 | 0.852 | 1.72 | 3.162 | 2.96 | 2.762 | 5.02 | 3.344 | 1.969 | -1.886 | -2.641 |
| C | - | - | - | 0.936 | 2.715 | 3.507 | -5.596 | 3.961 | 0.998 | 4.457 | 4.138 | 3.566 | 2.998 | 3.015 |
| D | - | - | - | - | 3.115 | 3.721 | 0.952 | 4.107 | 1.996 | 4.38 | 4.258 | 3.77 | 3.296 | 3.299 |
| E | - | - | - | - | - | 2.416 | 2.789 | 3.307 | 2.255 | 4.946 | 3.582 | 2.531 | 0.706 | 0.673 |
| F | - | - | - | - | - | - | 3.662 | 2.207 | 3.39 | 5.188 | 2.828 | -0.584 | 1.878 | 1.827 |
| G | - | - | - | - | - | - | - | 3.944 | 0.997 | 4.402 | 4.133 | 3.562 | 2.995 | 3.014 |
| H | - | - | - | - | - | - | - | - | 3.98 | 5.442 | 1.377 | 2.042 | 3.127 | 3.113 |
| I | - | - | - | - | - | - | - | - | - | 4.609 | 3.991 | 3.329 | 2.609 | 2.629 |
| J | - | - | - | - | - | - | - | - | - | - | 5.015 | 4.777 | 4.596 | 4.591 |
| K | - | - | - | - | - | - | - | - | - | - | - | 2.793 | 3.537 | 3.522 |
| L | - | - | - | - | - | - | - | - | - | - | - | - | 2.124 | 2.096 |
| M | - | - | - | - | - | - | - | - | - | - | - | - | - | -3.017 |
| N | - | - | - | - | - | - | - | - | - | - | - | - | - | - |

| | |
|---|---|
| | Larger than 10 |
| | Between 8 and 9,99 |
| | Between 6 and 7,99 |
| | Between 4 and 5,99 |
| | Between 2 and 3,99 |
| | Between 0 and 1,99 |
| | NA (Calculation failed) |
