## Supplementary Data S2 for "Interlaboratory comparison of standardised metabolomics and lipidomics analyses in human and rodent blood using the MxP® Quant 500 kit": Lac.pdf

**Amount of relative differences between laboratories**

| Min | A | B | C | D | E | F | G | H | I | J | K | L | M | N |
|---|---|---|---|---|---|---|---|---|---|---|---|---|---|---|
| A | - | 2.761 | -4.861 | 1.337 | 3.536 | 3.153 | 2.371 | 3.7 | 0.021 | 2.6 | 3.053 | 2.244 | -0.774 | -1.097 |
| B | - | - | 2.898 | 2.229 | 2.433 | 1.011 | 3.654 | 2.71 | 2.975 | -1.318 | 0.308 | 1.011 | 2.663 | 2.918 |
| C | - | - | - | 1.451 | 3.561 | 3.185 | 2.306 | 3.719 | -0.763 | 2.673 | 3.04 | 2.308 | -0.432 | -3.322 |
| D | - | - | - | - | 3.274 | 2.689 | 2.939 | 3.419 | 1.646 | 1.976 | 2.494 | 1.322 | 0.706 | 1.451 |
| E | - | - | - | - | - | 1.922 | 4.252 | 0.079 | 3.795 | 2.642 | 2.088 | 2.948 | 3.614 | 3.78 |
| F | - | - | - | - | - | - | 3.899 | 2.206 | 3.316 | 1.38 | -0.449 | 2.009 | 3.098 | 3.303 |
| G | - | - | - | - | - | - | - | 4.137 | 1.976 | 3.447 | 3.68 | 3.249 | 2.479 | 2.215 |
| H | - | - | - | - | - | - | - | - | 3.88 | 2.882 | 2.46 | 3.128 | 3.725 | 3.876 |
| I | - | - | - | - | - | - | - | - | - | 2.863 | 3.243 | 2.595 | 0.618 | -0.19 |
| J | - | - | - | - | - | - | - | - | - | - | 0.973 | 0.763 | 2.533 | 2.809 |
| K | - | - | - | - | - | - | - | - | - | - | - | 1.605 | 2.95 | 3.104 |
| L | - | - | - | - | - | - | - | - | - | - | - | - | 2.211 | 2.43 |
| M | - | - | - | - | - | - | - | - | - | - | - | - | - | 0.083 |
| N | - | - | - | - | - | - | - | - | - | - | - | - | - | - |

| | |
|---|---|
| | Larger than 10 |
| | Between 8 and 9,99 |
| | Between 6 and 7,99 |
| | Between 4 and 5,99 |
| | Between 2 and 3,99 |
| | Between 0 and 1,99 |
| | NA (Calculation failed) |

| Median | A | B | C | D | E | F | G | H | I | J | K | L | M | N |
|---|---|---|---|---|---|---|---|---|---|---|---|---|---|---|
| A | - | 1.832 | 0.401 | -0.329 | 2.867 | 2.567 | 1.687 | 2.831 | 1.417 | 2.577 | 0.73 | 2.106 | 1.586 | 0.807 |
| B | - | - | 2.373 | 2.181 | 1.947 | 1.273 | 2.812 | 1.863 | -0.071 | 1.319 | 1.017 | -0.438 | -0.68 | 0.928 |
| C | - | - | - | -0.845 | 3.094 | 2.845 | 0.871 | 3.062 | 1.991 | 2.855 | 1.566 | 2.481 | 2.101 | 1.617 |
| D | - | - | - | - | 3.005 | 2.734 | 1.252 | 2.967 | 1.776 | 2.749 | 1.265 | 2.335 | 1.903 | 1.333 |
| E | - | - | - | - | - | 0.591 | 3.499 | -2.037 | 2.325 | 0.499 | 2.614 | 1.7 | 2.224 | 2.584 |
| F | - | - | - | - | - | - | 3.275 | 0.326 | 1.785 | -3.752 | 2.185 | 0.779 | 1.639 | 2.147 |
| G | - | - | - | - | - | - | - | 3.329 | 2.519 | 3.156 | 2.247 | 2.871 | 2.6 | 2.274 |
| H | - | - | - | - | - | - | - | - | 2.245 | 0.205 | 2.552 | 1.581 | 2.143 | 2.517 |
| I | - | - | - | - | - | - | - | - | - | 1.748 | 0.057 | 0.754 | -1.503 | -0.069 |
| J | - | - | - | - | - | - | - | - | - | - | 2.201 | 0.83 | 1.657 | 2.166 |
| K | - | - | - | - | - | - | - | - | - | - | - | 1.44 | 0.434 | -2.895 |
| L | - | - | - | - | - | - | - | - | - | - | - | - | 0.457 | 1.416 |
| M | - | - | - | - | - | - | - | - | - | - | - | - | - | 0.392 |
| N | - | - | - | - | - | - | - | - | - | - | - | - | - | - |

| Max | A | B | C | D | E | F | G | H | I | J | K | L | M | N |
|---|---|---|---|---|---|---|---|---|---|---|---|---|---|---|
| A | - | 1.51 | 0.74 | 0.666 | 2.625 | 2.399 | 1.449 | 2.502 | 1.872 | 2.572 | -4.642 | 2.068 | 1.863 | 1.19 |
| B | - | - | 2.189 | 2.169 | 1.788 | 1.325 | 2.492 | 1.547 | -0.147 | 1.677 | 1.495 | 0.544 | -0.192 | -0.877 |
| C | - | - | - | -4.697 | 2.942 | 2.758 | 0.066 | 2.841 | 2.378 | 2.896 | 0.748 | 2.523 | 2.384 | 1.937 |
| D | - | - | - | - | 2.926 | 2.744 | 0.078 | 2.83 | 2.375 | 2.887 | 0.692 | 2.513 | 2.365 | 1.919 |
| E | - | - | - | - | - | 0.009 | 3.226 | -0.825 | 1.394 | -2.038 | 2.717 | 1.057 | 1.425 | 2.048 |
| F | - | - | - | - | - | - | 3.064 | -1.236 | 0.765 | -0.468 | 2.453 | 0.154 | 0.752 | 1.643 |
| G | - | - | - | - | - | - | - | 3.033 | 2.635 | 3.076 | 1.476 | 2.754 | 2.631 | 2.288 |
| H | - | - | - | - | - | - | - | - | 1.045 | -1.497 | 2.573 | 0.575 | 1.073 | 1.837 |
| I | - | - | - | - | - | - | - | - | - | 1.269 | 1.849 | -0.869 | -4.554 | 0.479 |
| J | - | - | - | - | - | - | - | - | - | - | 2.632 | 0.847 | 1.299 | 1.957 |
| K | - | - | - | - | - | - | - | - | - | - | - | 2.076 | 1.901 | 1.159 |
| L | - | - | - | - | - | - | - | - | - | - | - | - | -0.95 | 1.011 |
| M | - | - | - | - | - | - | - | - | - | - | - | - | - | 0.458 |
| N | - | - | - | - | - | - | - | - | - | - | - | - | - | - |
