## Supplementary Data S2 for "Interlaboratory comparison of standardised metabolomics and lipidomics analyses in human and rodent blood using the MxP® Quant 500 kit": Lys.pdf

**Amount of relative differences between laboratories**

| Min | A | B | C | D | E | F | G | H | I | J | K | L | M | N |
|---|---|---|---|---|---|---|---|---|---|---|---|---|---|---|
| A | - | 3.312 | 2.389 | -1.747 | 3.466 | 2.064 | 2.69 | 3.579 | 2.97 | 1.829 | 1.868 | -2.147 | 1.791 | 2.553 |
| B | - | - | 4.155 | 3.442 | 0.732 | 2.493 | 1.705 | 1.465 | 1.456 | 2.721 | 2.939 | 3.546 | 2.839 | 2.2 |
| C | - | - | - | 2.86 | 4.152 | 3.518 | 3.792 | 4.205 | 3.716 | 3.381 | 3.056 | 2.419 | 3.255 | 3.538 |
| D | - | - | - | - | 3.516 | 2.199 | 2.79 | 3.604 | 2.954 | 1.911 | 1.742 | -1.342 | 1.854 | 2.551 |
| E | - | - | - | - | - | 2.959 | 2.372 | -0.087 | 1.843 | 3.095 | 3.105 | 3.689 | 3.116 | 2.57 |
| F | - | - | - | - | - | - | 1.255 | 2.988 | 1.878 | -0.29 | -0.162 | 2.289 | -0.048 | 0.536 |
| G | - | - | - | - | - | - | - | 2.65 | 1.633 | 1.355 | -0.049 | 2.523 | 1.157 | -0.507 |
| H | - | - | - | - | - | - | - | - | 2.192 | 3.256 | 3.246 | 3.803 | 3.274 | 2.794 |
| I | - | - | - | - | - | - | - | - | - | 2.319 | 2.236 | 3.106 | 2.345 | 1.031 |
| J | - | - | - | - | - | - | - | - | - | - | -0.108 | 2.215 | -1.384 | 0.927 |
| K | - | - | - | - | - | - | - | - | - | - | - | 1.988 | -2.525 | 1.302 |
| L | - | - | - | - | - | - | - | - | - | - | - | - | 2.017 | 2.624 |
| M | - | - | - | - | - | - | - | - | - | - | - | - | - | 1.276 |
| N | - | - | - | - | - | - | - | - | - | - | - | - | - | - |

| | |
|---|---|
| | Larger than 10 |
| | Between 8 and 9,99 |
| | Between 6 and 7,99 |
| | Between 4 and 5,99 |
| | Between 2 and 3,99 |
| | Between 0 and 1,99 |
| | NA (Calculation failed) |

| Median | A | B | C | D | E | F | G | H | I | J | K | L | M | N |
|---|---|---|---|---|---|---|---|---|---|---|---|---|---|---|
| A | - | 3.65 | -0.635 | 0.779 | 3.548 | 2.454 | 0.501 | 3.617 | 3.046 | 2.878 | 2.028 | 1.58 | 2.217 | 2.59 |
| B | - | - | 3.759 | 3.654 | -0.286 | 3.004 | 3.679 | -2.231 | 2.275 | 2.617 | 3.279 | 3.449 | 3.122 | 2.899 |
| C | - | - | - | -3.772 | 3.461 | 2.075 | -4.325 | 3.497 | 2.902 | 2.49 | 1.54 | 0.907 | 2.052 | 2.278 |
| D | - | - | - | - | 3.376 | 1.981 | -2.347 | 3.438 | 2.771 | 2.511 | 1.318 | 0.48 | 1.721 | 2.162 |
| E | - | - | - | - | - | 2.831 | 3.551 | -0.744 | 1.926 | 2.336 | 3.118 | 3.308 | 2.993 | 2.691 |
| F | - | - | - | - | - | - | 2.112 | 2.838 | 1.622 | 0.9 | 0.505 | 1.372 | -0.393 | -0.878 |
| G | - | - | - | - | - | - | - | 3.449 | 2.785 | 2.66 | 1.466 | 0.55 | 1.501 | 2.248 |
| H | - | - | - | - | - | - | - | - | 2.16 | 2.496 | 3.219 | 3.406 | 3.112 | 2.831 |
| I | - | - | - | - | - | - | - | - | - | 0.3 | 2.213 | 2.509 | 2.036 | 1.285 |
| J | - | - | - | - | - | - | - | - | - | - | 1.693 | 2.128 | 1.181 | 0.241 |
| K | - | - | - | - | - | - | - | - | - | - | - | 0.178 | -0.732 | 1.04 |
| L | - | - | - | - | - | - | - | - | - | - | - | - | 1.009 | 1.585 |
| M | - | - | - | - | - | - | - | - | - | - | - | - | - | 0.369 |
| N | - | - | - | - | - | - | - | - | - | - | - | - | - | - |

| Max | A | B | C | D | E | F | G | H | I | J | K | L | M | N |
|---|---|---|---|---|---|---|---|---|---|---|---|---|---|---|
| A | - | 4.088 | 3.815 | 2.195 | 3.68 | 2.902 | 2.837 | 3.679 | 3.201 | 3.656 | 2.353 | 3.215 | 2.831 | 2.658 |
| B | - | - | 1.914 | 3.909 | 2.357 | 3.548 | 4.86 | 2.396 | 3.244 | 2.458 | 3.867 | 3.237 | 3.576 | 3.694 |
| C | - | - | - | 3.343 | -0.438 | 2.786 | 4.549 | -0.246 | 2.337 | 0.016 | 3.33 | 2.3 | 2.813 | 3.017 |
| D | - | - | - | - | 3.097 | 1.566 | 3.619 | 3.103 | 2.258 | 3.066 | -1.149 | 2.292 | 1.398 | 0.864 |
| E | - | - | - | - | - | 2.608 | 4.513 | -6.422 | 2.095 | -2.235 | 3.148 | 2.019 | 2.701 | 2.899 |
| F | - | - | - | - | - | - | 3.975 | 2.541 | 0.818 | 2.479 | 1.421 | 0.938 | -1.573 | 0.25 |
| G | - | - | - | - | - | - | - | 4.234 | 3.932 | 4.217 | 3.549 | 3.962 | 3.734 | 3.643 |
| H | - | - | - | - | - | - | - | - | 2.087 | -2.093 | 3.151 | 2.01 | 2.706 | 2.899 |
| I | - | - | - | - | - | - | - | - | - | 1.987 | 2.156 | -5.6 | 1.053 | 1.673 |
| J | - | - | - | - | - | - | - | - | - | - | 3.131 | 1.941 | 2.625 | 2.825 |
| K | - | - | - | - | - | - | - | - | - | - | - | 2.144 | 1.143 | 0.345 |
| L | - | - | - | - | - | - | - | - | - | - | - | - | 1.137 | 1.613 |
| M | - | - | - | - | - | - | - | - | - | - | - | - | - | -0.196 |
| N | - | - | - | - | - | - | - | - | - | - | - | - | - | - |
