## Supplementary Data S2 for "Interlaboratory comparison of standardised metabolomics and lipidomics analyses in human and rodent blood using the MxP® Quant 500 kit": lysoPC a C14.0.pdf

**Amount of relative differences between laboratories**

| Min | A | B | C | D | E | F | G | H | I | J | K | L | M | N |
|---|---|---|---|---|---|---|---|---|---|---|---|---|---|---|
| A | - | 3.545 | 2.959 | 3.712 | 4.162 | 1.063 | -0.006 | 4.675 | 3.688 | 3.584 | 3.39 | 2.196 | 1.23 | 3.431 |
| B | - | - | 2.486 | 0.714 | 2.625 | 3.36 | 3.731 | 3.583 | -2.775 | 0.078 | 1.266 | 2.811 | 3.458 | 1.033 |
| C | - | - | - | 2.772 | 3.487 | 2.266 | 2.908 | 4.107 | 2.429 | 2.618 | 1.492 | 0.636 | 2.367 | 1.797 |
| D | - | - | - | - | 2.28 | 3.659 | 3.931 | 3.452 | 0.542 | -0.582 | 2.162 | 3.176 | 3.729 | 1.898 |
| E | - | - | - | - | - | 4.147 | 4.363 | 2.26 | 2.862 | 2.385 | 3.301 | 3.81 | 4.243 | 3.204 |
| F | - | - | - | - | - | - | 1.626 | 4.572 | 3.464 | 3.348 | 3.074 | 1.375 | -2.402 | 3.152 |
| G | - | - | - | - | - | - | - | 4.654 | 3.673 | 3.681 | 3.497 | 2.466 | 1.79 | 3.491 |
| H | - | - | - | - | - | - | - | - | 3.855 | 3.748 | 4.112 | 4.564 | 4.819 | 4.083 |
| I | - | - | - | - | - | - | - | - | - | -0.575 | 1.333 | 3.054 | 3.585 | 1.087 |
| J | - | - | - | - | - | - | - | - | - | - | 1.654 | 3.034 | 3.565 | 1.444 |
| K | - | - | - | - | - | - | - | - | - | - | - | 2.366 | 3.121 | -0.96 |
| L | - | - | - | - | - | - | - | - | - | - | - | - | 0.805 | 2.777 |
| M | - | - | - | - | - | - | - | - | - | - | - | - | - | 3.087 |
| N | - | - | - | - | - | - | - | - | - | - | - | - | - | - |

| | |
|---|---|
| | Larger than 10 |
| | Between 8 and 9,99 |
| | Between 6 and 7,99 |
| | Between 4 and 5,99 |
| | Between 2 and 3,99 |
| | Between 0 and 1,99 |
| | NA (Calculation failed) |

| Median | A | B | C | D | E | F | G | H | I | J | K | L | M | N |
|---|---|---|---|---|---|---|---|---|---|---|---|---|---|---|
| A | - | 2.951 | 1.42 | 3.101 | 3.594 | 0.773 | -0.721 | 4.36 | 3.215 | 3.225 | 2.711 | 2.592 | -2.596 | 2.811 |
| B | - | - | 2.279 | 0.376 | 2.149 | 2.568 | 3.157 | 3.706 | 0.931 | 1.058 | -0.74 | 0.764 | 3.044 | -0.89 |
| C | - | - | - | 2.569 | 3.169 | 0.147 | 1.929 | 4.116 | 2.713 | 2.735 | 1.99 | 1.621 | 1.673 | 2.072 |
| D | - | - | - | - | 1.716 | 2.917 | 3.386 | 3.596 | -0.344 | -0.452 | 1.001 | 1.682 | 3.3 | 0.874 |
| E | - | - | - | - | - | 3.439 | 3.822 | 3.114 | 1.364 | 1.271 | 2.359 | 2.727 | 3.745 | 2.372 |
| F | - | - | - | - | - | - | 1.252 | 4.256 | 2.954 | 2.96 | 2.3 | 2.126 | 0.946 | 2.441 |
| G | - | - | - | - | - | - | - | 4.356 | 3.224 | 3.299 | 2.798 | 2.721 | -1.245 | 2.871 |
| H | - | - | - | - | - | - | - | - | 3.712 | 3.691 | 4.011 | 4.146 | 4.663 | 4.002 |
| I | - | - | - | - | - | - | - | - | - | -2.047 | 1.459 | 1.974 | 3.419 | 1.333 |
| J | - | - | - | - | - | - | - | - | - | - | 1.521 | 1.922 | 3.396 | 1.414 |
| K | - | - | - | - | - | - | - | - | - | - | - | -0.06 | 2.912 | -1.94 |
| L | - | - | - | - | - | - | - | - | - | - | - | - | 2.734 | -0.271 |
| M | - | - | - | - | - | - | - | - | - | - | - | - | - | 2.85 |
| N | - | - | - | - | - | - | - | - | - | - | - | - | - | - |

| Max | A | B | C | D | E | F | G | H | I | J | K | L | M | N |
|---|---|---|---|---|---|---|---|---|---|---|---|---|---|---|
| A | - | 1.733 | 2.805 | 1.762 | 2.077 | -0.265 | -0.924 | 3.451 | 0.813 | 0.852 | -0.659 | 3.324 | 2.51 | 0.223 |
| B | - | - | 1.779 | -1.352 | -1.291 | 2.014 | 1.915 | 3.914 | 2.633 | 2.324 | 1.431 | 3.697 | 1.15 | 1.452 |
| C | - | - | - | 1.83 | 1.483 | 2.888 | 2.798 | 4.132 | 3.171 | 3.025 | 2.607 | 3.986 | 0.202 | 2.533 |
| D | - | - | - | - | 0.15 | 2.043 | 1.808 | 3.838 | 2.455 | 2.239 | 1.329 | 3.685 | 1.223 | 1.191 |
| E | - | - | - | - | - | 2.309 | 2.194 | 4.004 | 2.858 | 2.486 | 1.802 | 3.788 | 0.305 | 1.847 |
| F | - | - | - | - | - | - | -1.6 | 3.347 | -0.179 | -0.102 | 0.543 | 3.21 | 2.72 | 1.061 |
| G | - | - | - | - | - | - | - | 3.522 | 1.146 | 0.427 | -0.073 | 3.253 | 2.647 | 0.295 |
| H | - | - | - | - | - | - | - | - | 3.358 | 3.514 | 3.808 | 0.375 | 4.242 | 3.807 |
| I | - | - | - | - | - | - | - | - | - | -0.026 | 1.665 | 3.025 | 2.97 | 1.756 |
| J | - | - | - | - | - | - | - | - | - | - | 1.24 | 3.061 | 2.933 | 1.346 |
| K | - | - | - | - | - | - | - | - | - | - | - | 3.408 | 2.308 | -1.782 |
| L | - | - | - | - | - | - | - | - | - | - | - | - | 4.126 | 3.617 |
| M | - | - | - | - | - | - | - | - | - | - | - | - | - | 2.141 |
| N | - | - | - | - | - | - | - | - | - | - | - | - | - | - |
