## Supplementary Data S2 for "Interlaboratory comparison of standardised metabolomics and lipidomics analyses in human and rodent blood using the MxP® Quant 500 kit": lysoPC a C16.0.pdf

**Amount of relative differences between laboratories**

| Min | A | B | C | D | E | F | G | H | I | J | K | L | M | N |
|---|---|---|---|---|---|---|---|---|---|---|---|---|---|---|
| A | - | 4.319 | 2.59 | 2.123 | 3.127 | 4.31 | -0.289 | 4.985 | 1.324 | 3.831 | 3.099 | 2.094 | 2.884 | 2.952 |
| B | - | - | 3.942 | 3.87 | 3.19 | -1.115 | 4.347 | 3.676 | 3.959 | 1.989 | 3.496 | 3.851 | 3.71 | 3.616 |
| C | - | - | - | -1.338 | 2.035 | 3.825 | 2.274 | 4.672 | 0.803 | 3.232 | 1.589 | -1.894 | 0.812 | 1.185 |
| D | - | - | - | - | 1.406 | 4.026 | 1.923 | 4.841 | 1.225 | 3.369 | 1.869 | -2.478 | 1.768 | 1.836 |
| E | - | - | - | - | - | 3.574 | 3.153 | 4.708 | 2.469 | 2.994 | 0.542 | 1.903 | 0.351 | -2.65 |
| F | - | - | - | - | - | - | 4.427 | 3.638 | 4.074 | 2.297 | 3.55 | 3.973 | 3.815 | 3.745 |
| G | - | - | - | - | - | - | - | 4.976 | 0.55 | 3.767 | 2.954 | 1.739 | 2.787 | 2.815 |
| H | - | - | - | - | - | - | - | - | 4.875 | 4.111 | 4.668 | 4.832 | 4.798 | 4.748 |
| I | - | - | - | - | - | - | - | - | - | 3.649 | 2.655 | 1.096 | 2.563 | 2.595 |
| J | - | - | - | - | - | - | - | - | - | - | 2.813 | 3.408 | 3.081 | 2.974 |
| K | - | - | - | - | - | - | - | - | - | - | - | 1.985 | -0.163 | -0.856 |
| L | - | - | - | - | - | - | - | - | - | - | - | - | 2.203 | 2.24 |
| M | - | - | - | - | - | - | - | - | - | - | - | - | - | -1.207 |
| N | - | - | - | - | - | - | - | - | - | - | - | - | - | - |

| | |
|---|---|
| | Larger than 10 |
| | Between 8 and 9,99 |
| | Between 6 and 7,99 |
| | Between 4 and 5,99 |
| | Between 2 and 3,99 |
| | Between 0 and 1,99 |
| | NA (Calculation failed) |

| Median | A | B | C | D | E | F | G | H | I | J | K | L | M | N |
|---|---|---|---|---|---|---|---|---|---|---|---|---|---|---|
| A | - | 3.004 | -1.879 | 2.474 | 3.481 | 3.423 | 1.073 | 3.969 | 2.006 | 3.096 | 2.602 | 3.14 | 1.028 | 1.549 |
| B | - | - | 3.31 | 1.795 | 1.249 | 1.307 | 2.835 | 2.983 | 2.477 | -2.941 | 1.547 | -6.003 | 2.814 | 2.623 |
| C | - | - | - | 2.544 | 3.487 | 3.478 | 1.296 | 4.005 | 2.024 | 3.168 | 2.657 | 3.176 | 1.312 | 1.724 |
| D | - | - | - | - | 2.456 | 2.426 | 1.898 | 3.402 | 0.718 | 1.632 | -1.566 | 1.79 | 1.913 | 1.509 |
| E | - | - | - | - | - | -0.1 | 3.392 | 2.253 | 2.916 | 1.502 | 2.487 | 1.275 | 3.332 | 3.218 |
| F | - | - | - | - | - | - | 3.301 | 2.464 | 3.017 | 1.278 | 2.43 | 1.322 | 3.289 | 3.153 |
| G | - | - | - | - | - | - | - | 3.788 | 1.023 | 2.718 | 2.015 | 2.786 | -4.463 | -0.259 |
| H | - | - | - | - | - | - | - | - | 3.903 | 3.08 | 3.568 | 3.138 | 4.022 | 3.942 |
| I | - | - | - | - | - | - | - | - | - | 2.203 | 0.974 | 2.318 | 1.097 | 0.293 |
| J | - | - | - | - | - | - | - | - | - | - | 1.562 | -4.819 | 2.837 | 2.643 |
| K | - | - | - | - | - | - | - | - | - | - | - | 1.565 | 2.043 | 1.69 |
| L | - | - | - | - | - | - | - | - | - | - | - | - | 2.933 | 2.727 |
| M | - | - | - | - | - | - | - | - | - | - | - | - | - | -0.263 |
| N | - | - | - | - | - | - | - | - | - | - | - | - | - | - |

| Max | A | B | C | D | E | F | G | H | I | J | K | L | M | N |
|---|---|---|---|---|---|---|---|---|---|---|---|---|---|---|
| A | - | -0.868 | 2.511 | 2.741 | 3.792 | 1.845 | 1.652 | 2.006 | 2.54 | 1.754 | 1.942 | 3.711 | 1.278 | 0.323 |
| B | - | - | 2.383 | 2.822 | 3.916 | 2.018 | 1.824 | 2.123 | 2.661 | 1.951 | 2.181 | 3.758 | 0.981 | -0.205 |
| C | - | - | - | 3.551 | 4.254 | 3.134 | 3.061 | 3.199 | 3.458 | 3.107 | 3.209 | 4.153 | 1.628 | 2.069 |
| D | - | - | - | - | 3.108 | 1.665 | 1.877 | 1.38 | -0.284 | 1.803 | 1.389 | 2.782 | 3.286 | 3.073 |
| E | - | - | - | - | - | 3.457 | 3.559 | 3.432 | 3.314 | 3.664 | 3.574 | 0.24 | 4.289 | 4.13 |
| F | - | - | - | - | - | - | -1.175 | -1.101 | 1.211 | -2.673 | -1.96 | 3.309 | 2.63 | 2.313 |
| G | - | - | - | - | - | - | - | -0.074 | 1.437 | -2.536 | -0.438 | 3.356 | 2.548 | 2.184 |
| H | - | - | - | - | - | - | - | - | 1.115 | -1.264 | -2.022 | 3.276 | 2.717 | 2.428 |
| I | - | - | - | - | - | - | - | - | - | 1.396 | 0.926 | 2.938 | 3.148 | 2.913 |
| J | - | - | - | - | - | - | - | - | - | - | -0.553 | 3.328 | 2.597 | 2.278 |
| K | - | - | - | - | - | - | - | - | - | - | - | 3.242 | 2.769 | 2.471 |
| L | - | - | - | - | - | - | - | - | - | - | - | - | 4.159 | 4.041 |
| M | - | - | - | - | - | - | - | - | - | - | - | - | - | 0.239 |
| N | - | - | - | - | - | - | - | - | - | - | - | - | - | - |
