## Supplementary Data S2 for "Interlaboratory comparison of standardised metabolomics and lipidomics analyses in human and rodent blood using the MxP® Quant 500 kit": lysoPC a C16.1.pdf

**Amount of relative differences between laboratories**

| Min | A | B | C | D | E | F | G | H | I | J | K | L | M | N |
|---|---|---|---|---|---|---|---|---|---|---|---|---|---|---|
| A | - | 4.585 | 2.874 | 3.466 | 4.996 | 3.975 | 0.548 | 4.138 | 3.241 | 4.038 | 4.12 | 3.536 | 3.264 | 2.777 |
| B | - | - | 4.161 | 3.637 | 3.202 | 3.026 | 4.596 | 2.272 | 3.53 | 2.857 | 2.773 | 3.501 | 3.991 | 4.112 |
| C | - | - | - | 2.121 | 4.594 | 3.111 | 2.523 | 3.472 | 1.995 | 3.225 | 3.285 | 2.374 | 1.023 | -7.092 |
| D | - | - | - | - | 4.417 | 2.238 | 3.41 | 2.843 | -0.076 | 2.472 | 2.616 | -0.476 | 1.078 | 2.213 |
| E | - | - | - | - | - | 4.24 | 5.122 | 3.821 | 4.285 | 4.086 | 4.103 | 4.364 | 4.741 | 4.76 |
| F | - | - | - | - | - | - | 3.995 | 1.374 | 2.392 | -0.304 | 0.894 | 1.974 | 2.863 | 3.265 |
| G | - | - | - | - | - | - | - | 4.04 | 2.964 | 3.936 | 4.002 | 3.352 | 3.074 | 2.431 |
| H | - | - | - | - | - | - | - | - | 3.216 | 0.711 | -0.033 | 2.828 | 3.301 | 3.689 |
| I | - | - | - | - | - | - | - | - | - | 2.936 | 3.043 | 1.205 | -3.749 | 1.54 |
| J | - | - | - | - | - | - | - | - | - | - | -1.133 | 2.132 | 3.013 | 3.326 |
| K | - | - | - | - | - | - | - | - | - | - | - | 2.207 | 3.11 | 3.322 |
| L | - | - | - | - | - | - | - | - | - | - | - | - | 1.235 | 2.394 |
| M | - | - | - | - | - | - | - | - | - | - | - | - | - | 1.011 |
| N | - | - | - | - | - | - | - | - | - | - | - | - | - | - |

| | |
|---|---|
| | Larger than 10 |
| | Between 8 and 9,99 |
| | Between 6 and 7,99 |
| | Between 4 and 5,99 |
| | Between 2 and 3,99 |
| | Between 0 and 1,99 |
| | NA (Calculation failed) |

| Median | A | B | C | D | E | F | G | H | I | J | K | L | M | N |
|---|---|---|---|---|---|---|---|---|---|---|---|---|---|---|
| A | - | 3.569 | 0.438 | 2.968 | 3.876 | 3.431 | 1.27 | 3.988 | 3.192 | 3.196 | 2.928 | 3.442 | 1.632 | 2.255 |
| B | - | - | 3.629 | 2.334 | 1.552 | 0.232 | 3.469 | 1.917 | 1.901 | 1.72 | 2.27 | 0.452 | 3.331 | 3.038 |
| C | - | - | - | 2.708 | 3.756 | 3.291 | 0.164 | 3.86 | 2.954 | 3.001 | 2.702 | 3.282 | 0.948 | 1.84 |
| D | - | - | - | - | 2.883 | 1.749 | 2.542 | 3.107 | 0.424 | 0.609 | -2.953 | 1.747 | 2.325 | 1.691 |
| E | - | - | - | - | - | 2.117 | 3.875 | -1.168 | 2.843 | 2.703 | 3.001 | 2.188 | 3.754 | 3.534 |
| F | - | - | - | - | - | - | 3.258 | 2.414 | 1.077 | 0.96 | 1.822 | -5.086 | 3.121 | 2.776 |
| G | - | - | - | - | - | - | - | 3.789 | 2.784 | 2.788 | 2.389 | 3.111 | -0.545 | 1.258 |
| H | - | - | - | - | - | - | - | - | 2.998 | 2.973 | 3.28 | 2.546 | 3.909 | 3.708 |
| I | - | - | - | - | - | - | - | - | - | -3.761 | 0.772 | 0.945 | 2.723 | 2.253 |
| J | - | - | - | - | - | - | - | - | - | - | 0.662 | 0.901 | 2.72 | 2.261 |
| K | - | - | - | - | - | - | - | - | - | - | - | 1.732 | 2.318 | 1.664 |
| L | - | - | - | - | - | - | - | - | - | - | - | - | 3.12 | 2.766 |
| M | - | - | - | - | - | - | - | - | - | - | - | - | - | 0.757 |
| N | - | - | - | - | - | - | - | - | - | - | - | - | - | - |

| Max | A | B | C | D | E | F | G | H | I | J | K | L | M | N |
|---|---|---|---|---|---|---|---|---|---|---|---|---|---|---|
| A | - | -1.323 | 2.476 | 2.009 | 1.891 | 2.57 | 1.781 | 3.796 | 3.113 | 0.428 | 1.293 | 3.329 | 1.866 | 1.267 |
| B | - | - | 2.58 | 1.882 | 1.911 | 2.505 | 1.635 | 3.762 | 3.068 | -0.056 | 1.46 | 3.278 | 2.006 | 1.083 |
| C | - | - | - | 3.188 | 1.111 | 3.449 | 3.095 | 4.204 | 3.764 | 2.713 | 1.665 | 3.886 | 0.867 | 2.935 |
| D | - | - | - | - | 2.916 | 1.021 | -0.668 | 3.359 | 2.291 | 1.49 | 2.716 | 2.653 | 3.002 | 0.701 |
| E | - | - | - | - | - | 3.315 | 2.832 | 4.09 | 3.579 | 2.254 | 0.52 | 3.731 | -2.046 | 2.603 |
| F | - | - | - | - | - | - | 1.417 | 3.077 | 1.466 | 2.296 | 3.184 | 2.107 | 3.348 | 1.891 |
| G | - | - | - | - | - | - | - | 3.437 | 2.452 | 1.094 | 2.551 | 2.785 | 2.874 | 0.031 |
| H | - | - | - | - | - | - | - | - | 2.58 | 3.862 | 4.225 | 2.149 | 4.348 | 3.728 |
| I | - | - | - | - | - | - | - | - | - | 3.015 | 3.578 | 0.587 | 3.757 | 2.783 |
| J | - | - | - | - | - | - | - | - | - | - | 1.879 | 3.141 | 2.337 | 0.213 |
| K | - | - | - | - | - | - | - | - | - | - | - | 3.604 | 0.272 | 2.202 |
| L | - | - | - | - | - | - | - | - | - | - | - | - | 3.931 | 3.098 |
| M | - | - | - | - | - | - | - | - | - | - | - | - | - | 2.563 |
| N | - | - | - | - | - | - | - | - | - | - | - | - | - | - |
