## Supplementary Data S2 for "Interlaboratory comparison of standardised metabolomics and lipidomics analyses in human and rodent blood using the MxP® Quant 500 kit": lysoPC a C17.0.pdf

**Amount of relative differences between laboratories**

| Min | A | B | C | D | E | F | G | H | I | J | K | L | M | N |
|---|---|---|---|---|---|---|---|---|---|---|---|---|---|---|
| A | - | 3.429 | 2.305 | 2.637 | 2.175 | 4.369 | 2.245 | 4.218 | -0.448 | 3.764 | 1.386 | 2.126 | 2.042 | 3.188 |
| B | - | - | 2.735 | 2.274 | 2.853 | 3.419 | 2.853 | 3.11 | 3.66 | 1.67 | 3.217 | 2.829 | 2.896 | 1.074 |
| C | - | - | - | 1.141 | 0.388 | 4.046 | -2.181 | 3.844 | 2.096 | 3.236 | 0.805 | -1.807 | -0.548 | 2.211 |
| D | - | - | - | - | 0.834 | 3.878 | 1.176 | 3.66 | 2.758 | 2.892 | 2.163 | 0.894 | 1.473 | 1.42 |
| E | - | - | - | - | - | 4.155 | 0.741 | 4.079 | 2.319 | 3.494 | 1.059 | -0.272 | 1.131 | 2.749 |
| F | - | - | - | - | - | - | 4.321 | 1.375 | 4.499 | 3.008 | 4.366 | 4.195 | 4.341 | 3.792 |
| G | - | - | - | - | - | - | - | 3.861 | 2.133 | 3.258 | 0.559 | -1.987 | -1.18 | 2.259 |
| H | - | - | - | - | - | - | - | - | 4.458 | 2.537 | 4.24 | 4.045 | 4.133 | 3.507 |
| I | - | - | - | - | - | - | - | - | - | 3.918 | 1.973 | 2.497 | 2.621 | 3.433 |
| J | - | - | - | - | - | - | - | - | - | - | 3.607 | 3.312 | 3.486 | 2.414 |
| K | - | - | - | - | - | - | - | - | - | - | - | 0.858 | 0.894 | 2.715 |
| L | - | - | - | - | - | - | - | - | - | - | - | - | -1.587 | 2.446 |
| M | - | - | - | - | - | - | - | - | - | - | - | - | - | 2.365 |
| N | - | - | - | - | - | - | - | - | - | - | - | - | - | - |

| | |
|---|---|
| | Larger than 10 |
| | Between 8 and 9,99 |
| | Between 6 and 7,99 |
| | Between 4 and 5,99 |
| | Between 2 and 3,99 |
| | Between 0 and 1,99 |
| | NA (Calculation failed) |

| Median | A | B | C | D | E | F | G | H | I | J | K | L | M | N |
|---|---|---|---|---|---|---|---|---|---|---|---|---|---|---|
| A | - | 3.266 | -0.686 | 1.829 | 3.093 | 3.676 | 0.725 | 3.935 | 0.266 | 3.1 | 1.427 | 2.781 | -0.243 | 2.018 |
| B | - | - | 3.322 | 2.763 | 0.563 | 1.772 | 3.167 | 2.631 | 3.257 | 0.135 | 2.981 | 1.669 | 3.274 | 2.647 |
| C | - | - | - | 1.458 | 2.989 | 3.623 | -0.095 | 3.873 | -2.327 | 3.049 | 0.844 | 2.604 | -1.29 | 1.78 |
| D | - | - | - | - | 2.341 | 3.29 | 0.895 | 3.615 | 1.375 | 2.485 | 0.164 | 1.792 | 1.297 | -0.509 |
| E | - | - | - | - | - | 2.023 | 2.964 | 2.896 | 2.867 | -0.913 | 2.719 | 0.715 | 3.132 | 2.336 |
| F | - | - | - | - | - | - | 3.725 | 1.262 | 3.844 | 2.127 | 3.629 | 2.908 | 3.75 | 3.358 |
| G | - | - | - | - | - | - | - | 3.791 | -0.684 | 2.871 | -0.006 | 2.383 | -0.75 | 1.324 |
| H | - | - | - | - | - | - | - | - | 4.082 | 2.908 | 3.912 | 3.326 | 4.062 | 3.716 |
| I | - | - | - | - | - | - | - | - | - | 2.803 | 0.476 | 2.494 | -0.407 | 1.385 |
| J | - | - | - | - | - | - | - | - | - | - | 2.855 | 1.355 | 3.106 | 2.427 |
| K | - | - | - | - | - | - | - | - | - | - | - | 2.106 | 0.877 | 0.445 |
| L | - | - | - | - | - | - | - | - | - | - | - | - | 2.768 | 1.681 |
| M | - | - | - | - | - | - | - | - | - | - | - | - | - | 1.612 |
| N | - | - | - | - | - | - | - | - | - | - | - | - | - | - |

| Max | A | B | C | D | E | F | G | H | I | J | K | L | M | N |
|---|---|---|---|---|---|---|---|---|---|---|---|---|---|---|
| A | - | 3.076 | 1.958 | -2.26 | 3.843 | 2.061 | 0.709 | 3.543 | 2.24 | 1.804 | 1.479 | 3.331 | 1.241 | 0.835 |
| B | - | - | 3.753 | 3.253 | 2.776 | 2.179 | 3.44 | 1.813 | 1.981 | 2.443 | 2.595 | 0.854 | 3.568 | 3.467 |
| C | - | - | - | 1.812 | 4.134 | 2.951 | 1.116 | 3.904 | 3.061 | 2.831 | 2.689 | 3.748 | 0.566 | 1.043 |
| D | - | - | - | - | 3.906 | 2.142 | 0.52 | 3.564 | 2.322 | 1.907 | 1.719 | 3.354 | 1.101 | 0.651 |
| E | - | - | - | - | - | 3.451 | 4.161 | 1.558 | 3.523 | 3.643 | 3.643 | 2.257 | 4.286 | 4.218 |
| F | - | - | - | - | - | - | 2.59 | 2.978 | -0.635 | -0.651 | 0.614 | 2.639 | 2.759 | 2.611 |
| G | - | - | - | - | - | - | - | 3.71 | 2.639 | 2.332 | 2.084 | 3.527 | -0.425 | -2.962 |
| H | - | - | - | - | - | - | - | - | 2.955 | 3.206 | 3.322 | 0.85 | 3.989 | 3.925 |
| I | - | - | - | - | - | - | - | - | - | 0.346 | 0.957 | 2.491 | 2.911 | 2.771 |
| J | - | - | - | - | - | - | - | - | - | - | -0.438 | 2.768 | 2.602 | 2.442 |
| K | - | - | - | - | - | - | - | - | - | - | - | 2.902 | 2.448 | 2.212 |
| L | - | - | - | - | - | - | - | - | - | - | - | - | 3.789 | 3.72 |
| M | - | - | - | - | - | - | - | - | - | - | - | - | - | -0.633 |
| N | - | - | - | - | - | - | - | - | - | - | - | - | - | - |
