## Supplementary Data S2 for "Interlaboratory comparison of standardised metabolomics and lipidomics analyses in human and rodent blood using the MxP® Quant 500 kit": lysoPC a C28.0.pdf

**Amount of relative differences between laboratories**

| Min | A | B | C | D | E | F | G | H | I | J | K | L | M | N |
|---|---|---|---|---|---|---|---|---|---|---|---|---|---|---|
| A | - | NA | NA | 7.656 | 5.292 | 3.921 | 1.96 | 5.948 | 3.842 | NA | NA | 1.834 | NA | 1.651 |
| B | - | - | NA | NA | NA | NA | NA | NA | NA | NA | NA | NA | NA | NA |
| C | - | - | - | NA | NA | NA | NA | NA | NA | NA | NA | NA | NA | NA |
| D | - | - | - | - | 6.712 | 5.812 | 5.988 | 3.809 | 6.259 | NA | NA | 5.18 | NA | 6.061 |
| E | - | - | - | - | - | 4.109 | 4.572 | 5.864 | 1.421 | NA | NA | 4.942 | NA | 4.312 |
| F | - | - | - | - | - | - | 0.445 | 5.105 | 4.416 | NA | NA | 2.71 | NA | 1.9 |
| G | - | - | - | - | - | - | - | 5.271 | 4.131 | NA | NA | 1.81 | NA | 2.3 |
| H | - | - | - | - | - | - | - | - | 6.387 | NA | NA | 5.726 | NA | 5.748 |
| I | - | - | - | - | - | - | - | - | - | NA | NA | 1.761 | NA | 3.325 |
| J | - | - | - | - | - | - | - | - | - | - | NA | NA | NA | NA |
| K | - | - | - | - | - | - | - | - | - | - | - | NA | NA | NA |
| L | - | - | - | - | - | - | - | - | - | - | - | - | NA | 1.905 |
| M | - | - | - | - | - | - | - | - | - | - | - | - | - | NA |
| N | - | - | - | - | - | - | - | - | - | - | - | - | - | - |

| | |
|---|---|
| | Larger than 10 |
| | Between 8 and 9,99 |
| | Between 6 and 7,99 |
| | Between 4 and 5,99 |
| | Between 2 and 3,99 |
| | Between 0 and 1,99 |
| | NA (Calculation failed) |

| Median | A | B | C | D | E | F | G | H | I | J | K | L | M | N |
|---|---|---|---|---|---|---|---|---|---|---|---|---|---|---|
| A | - | NA | NA | 5.011 | 5.14 | 4.333 | 2.883 | 5.115 | 0.347 | NA | NA | 3.012 | NA | 3.665 |
| B | - | - | NA | NA | NA | NA | NA | NA | NA | NA | NA | NA | NA | NA |
| C | - | - | - | NA | NA | NA | NA | NA | NA | NA | NA | NA | NA | NA |
| D | - | - | - | - | 6.408 | 5.826 | 5.448 | 3.025 | 5.133 | NA | NA | 4.279 | NA | 6.012 |
| E | - | - | - | - | - | 3.583 | 4.305 | 5.595 | 4.388 | NA | NA | 4.963 | NA | 3.678 |
| F | - | - | - | - | - | - | 3.37 | 5.326 | 3.634 | NA | NA | 4.521 | NA | 1.356 |
| G | - | - | - | - | - | - | - | 5.099 | 1.466 | NA | NA | 3.749 | NA | 3.34 |
| H | - | - | - | - | - | - | - | - | 5.508 | NA | NA | 4.869 | NA | 5.908 |
| I | - | - | - | - | - | - | - | - | - | NA | NA | 3.282 | NA | 4.372 |
| J | - | - | - | - | - | - | - | - | - | - | NA | NA | NA | NA |
| K | - | - | - | - | - | - | - | - | - | - | - | NA | NA | NA |
| L | - | - | - | - | - | - | - | - | - | - | - | - | NA | 4.215 |
| M | - | - | - | - | - | - | - | - | - | - | - | - | - | NA |
| N | - | - | - | - | - | - | - | - | - | - | - | - | - | - |

| Max | A | B | C | D | E | F | G | H | I | J | K | L | M | N |
|---|---|---|---|---|---|---|---|---|---|---|---|---|---|---|
| A | - | NA | NA | 1.37 | 4.75 | 5.592 | 3.449 | 3.767 | 3.213 | NA | NA | 3.673 | NA | 5.625 |
| B | - | - | NA | NA | NA | NA | NA | NA | NA | NA | NA | NA | NA | NA |
| C | - | - | - | NA | NA | NA | NA | NA | NA | NA | NA | NA | NA | NA |
| D | - | - | - | - | 5.416 | 5.902 | 4.556 | 1.833 | 0.663 | NA | NA | 0.894 | NA | 5.806 |
| E | - | - | - | - | - | 4.136 | 3.971 | 5.345 | 5.322 | NA | NA | 4.985 | NA | 3.842 |
| F | - | - | - | - | - | - | 4.473 | 5.478 | 5.31 | NA | NA | 5.325 | NA | 0.96 |
| G | - | - | - | - | - | - | - | 4.949 | 4.258 | NA | NA | 4.573 | NA | 4.912 |
| H | - | - | - | - | - | - | - | - | 3.872 | NA | NA | 2.177 | NA | 6.397 |
| I | - | - | - | - | - | - | - | - | - | NA | NA | 4.548 | NA | 5.949 |
| J | - | - | - | - | - | - | - | - | - | - | NA | NA | NA | NA |
| K | - | - | - | - | - | - | - | - | - | - | - | NA | NA | NA |
| L | - | - | - | - | - | - | - | - | - | - | - | - | NA | 6.238 |
| M | - | - | - | - | - | - | - | - | - | - | - | - | - | NA |
| N | - | - | - | - | - | - | - | - | - | - | - | - | - | - |
