## Supplementary Data S2 for "Interlaboratory comparison of standardised metabolomics and lipidomics analyses in human and rodent blood using the MxP® Quant 500 kit": Met-SO.pdf

**Amount of relative differences between laboratories**

| Min | A | B | C | D | E | F | G | H | I | J | K | L | M | N |
|---|---|---|---|---|---|---|---|---|---|---|---|---|---|---|
| A | - | 5.766 | 8.147 | 5.627 | 5.837 | 4.401 | 5.812 | 0.571 | 6.179 | 6.085 | 3.092 | 5.584 | 5.525 | 5.739 |
| B | - | - | 8.749 | 2.387 | 4.244 | 6.647 | 5.172 | 6.794 | 5.281 | 4.371 | 6.509 | 3.055 | 2.045 | 1.867 |
| C | - | - | - | 7.542 | 7.138 | 8.435 | 6.548 | 7.962 | 7.309 | 7.596 | 7.483 | 7.2 | 7.474 | 7.708 |
| D | - | - | - | - | 4.528 | 6.494 | 5.4 | 6.643 | 5.39 | 4.629 | 6.352 | 3.695 | 0.029 | 0.655 |
| E | - | - | - | - | - | 7.191 | 4.922 | 7.29 | 4.67 | 1.232 | 7.087 | 3.707 | 4.842 | 4.784 |
| F | - | - | - | - | - | - | 5.553 | 4.565 | 6.094 | 5.902 | 4.514 | 5.386 | 5.243 | 5.447 |
| G | - | - | - | - | - | - | - | 8.579 | 5.548 | 6.548 | 8.338 | 6.624 | 7.05 | 7.136 |
| H | - | - | - | - | - | - | - | - | 6.264 | 6.144 | 2.45 | 5.619 | 5.62 | 5.838 |
| I | - | - | - | - | - | - | - | - | - | 5.056 | 7.604 | 5.31 | 5.998 | 6.111 |
| J | - | - | - | - | - | - | - | - | - | - | 7.012 | 3.31 | 4.762 | 4.856 |
| K | - | - | - | - | - | - | - | - | - | - | - | 5.722 | 5.718 | 5.935 |
| L | - | - | - | - | - | - | - | - | - | - | - | - | 3.727 | 3.592 |
| M | - | - | - | - | - | - | - | - | - | - | - | - | - | 0.751 |
| N | - | - | - | - | - | - | - | - | - | - | - | - | - | - |

| | |
|---|---|
| | Larger than 10 |
| | Between 8 and 9,99 |
| | Between 6 and 7,99 |
| | Between 4 and 5,99 |
| | Between 2 and 3,99 |
| | Between 0 and 1,99 |
| | NA (Calculation failed) |

| Median | A | B | C | D | E | F | G | H | I | J | K | L | M | N |
|---|---|---|---|---|---|---|---|---|---|---|---|---|---|---|
| A | - | 4.057 | 4.917 | 3.874 | 5.073 | 1.791 | 3.695 | 1.74 | 5.281 | 4.695 | -0.577 | 4.712 | 4.133 | 4.014 |
| B | - | - | 5.138 | 2.23 | 4.317 | 4.217 | 3.473 | 4.502 | 4.624 | 3.353 | 4.574 | 3.413 | 0.912 | 0.025 |
| C | - | - | - | 6.304 | 6.218 | 6.379 | 4.952 | 6.319 | 6.445 | 6.383 | 6.145 | 6.272 | 6.218 | 6.334 |
| D | - | - | - | - | 4.557 | 3.732 | 4 | 4.088 | 4.809 | 3.831 | 4.187 | 3.879 | 2.644 | 1.805 |
| E | - | - | - | - | - | 5.588 | 3.036 | 5.706 | 2.637 | 3.609 | 5.766 | 3.521 | 4.503 | 4.711 |
| F | - | - | - | - | - | - | 3.453 | 0.636 | 5.222 | 4.554 | 1.708 | 4.586 | 3.934 | 3.758 |
| G | - | - | - | - | - | - | - | 6.628 | 4.27 | 5.213 | 6.705 | 5.318 | 5.592 | 5.712 |
| H | - | - | - | - | - | - | - | - | 5.253 | 4.558 | 0.924 | 4.58 | 3.917 | 3.774 |
| I | - | - | - | - | - | - | - | - | - | 4.185 | 6.036 | 4.089 | 4.789 | 5.013 |
| J | - | - | - | - | - | - | - | - | - | - | 5.273 | -0.552 | 3.13 | 3.622 |
| K | - | - | - | - | - | - | - | - | - | - | - | 4.674 | 4.04 | 3.923 |
| L | - | - | - | - | - | - | - | - | - | - | - | - | 3.329 | 3.724 |
| M | - | - | - | - | - | - | - | - | - | - | - | - | - | 1.476 |
| N | - | - | - | - | - | - | - | - | - | - | - | - | - | - |

| Max | A | B | C | D | E | F | G | H | I | J | K | L | M | N |
|---|---|---|---|---|---|---|---|---|---|---|---|---|---|---|
| A | - | 3.283 | 1.896 | 3.845 | 3.702 | 1.028 | 4.152 | 2.421 | 0.784 | 2.931 | 2.552 | 2.122 | 1.554 | 3.154 |
| B | - | - | 2.604 | 2.085 | 4.377 | 2.77 | 2.731 | 3.824 | 2.897 | 0.89 | 3.822 | 3.702 | 2.632 | -0.409 |
| C | - | - | - | 3.416 | 4.012 | 0.517 | 3.764 | 3.186 | -0.526 | 2.142 | 3.272 | 2.847 | -2.421 | 2.519 |
| D | - | - | - | - | 4.583 | 3.409 | 1.411 | 4.138 | 3.487 | 2.546 | 4.139 | 4.044 | 3.323 | 2.259 |
| E | - | - | - | - | - | 4.135 | 5.153 | 3.121 | 4.092 | 4.574 | 3.069 | 3.305 | 4.198 | 4.659 |
| F | - | - | - | - | - | - | 3.872 | 2.94 | -2.319 | 2.454 | 2.961 | 2.622 | -0.074 | 2.746 |
| G | - | - | - | - | - | - | - | 4.276 | 3.807 | 2.998 | 4.287 | 4.255 | 3.621 | 2.794 |
| H | - | - | - | - | - | - | - | - | 3.127 | 3.815 | -3.095 | 0.513 | 3.229 | 3.963 |
| I | - | - | - | - | - | - | - | - | - | 2.623 | 2.845 | 2.516 | 0.663 | 2.909 |
| J | - | - | - | - | - | - | - | - | - | - | 3.633 | 3.502 | 2.152 | 0.167 |
| K | - | - | - | - | - | - | - | - | - | - | - | 0.877 | 3.255 | 3.983 |
| L | - | - | - | - | - | - | - | - | - | - | - | - | 2.955 | 3.806 |
| M | - | - | - | - | - | - | - | - | - | - | - | - | - | 2.533 |
| N | - | - | - | - | - | - | - | - | - | - | - | - | - | - |
