## Supplementary Data S2 for "Interlaboratory comparison of standardised metabolomics and lipidomics analyses in human and rodent blood using the MxP® Quant 500 kit": OH-GlutAcid.pdf

Amount of relative differences between laboratories

| Min | A | B | C | D | E | F | G | H | I | J | K | L | M | N |
|---|---|---|---|---|---|---|---|---|---|---|---|---|---|---|
| A | - | NA | NA | NA | NA | NA | NA | NA | NA | NA | NA | NA | NA | NA |
| B | - | - | NA | 4.586 | NA | 5.343 | NA | NA | NA | NA | NA | NA | 3.752 | NA |
| C | - | - | - | NA | NA | NA | NA | NA | NA | NA | NA | NA | NA | NA |
| D | - | - | - | - | NA | 5.605 | NA | NA | NA | NA | NA | NA | 4.186 | NA |
| E | - | - | - | - | - | NA | NA | NA | NA | NA | NA | NA | NA | NA |
| F | - | - | - | - | - | - | NA | NA | NA | NA | NA | NA | 4.252 | NA |
| G | - | - | - | - | - | - | - | NA | NA | NA | NA | NA | NA | NA |
| H | - | - | - | - | - | - | - | - | NA | NA | NA | NA | NA | NA |
| I | - | - | - | - | - | - | - | - | - | NA | NA | NA | NA | NA |
| J | - | - | - | - | - | - | - | - | - | - | NA | NA | NA | NA |
| K | - | - | - | - | - | - | - | - | - | - | - | NA | NA | NA |
| L | - | - | - | - | - | - | - | - | - | - | - | - | NA | NA |
| M | - | - | - | - | - | - | - | - | - | - | - | - | - | NA |
| N | - | - | - | - | - | - | - | - | - | - | - | - | - | - |

| | |
|---|---|
| | Larger than 10 |
| | Between 8 and 9,99 |
| | Between 6 and 7,99 |
| | Between 4 and 5,99 |
| | Between 2 and 3,99 |
| | Between 0 and 1,99 |
| | NA (Calculation failed) |

| Median | A | B | C | D | E | F | G | H | I | J | K | L | M | N |
|---|---|---|---|---|---|---|---|---|---|---|---|---|---|---|
| A | - | NA | NA | NA | NA | NA | NA | NA | NA | NA | NA | NA | NA | NA |
| B | - | - | NA | 5.542 | NA | 5.503 | NA | NA | NA | NA | NA | NA | 4.422 | NA |
| C | - | - | - | NA | NA | NA | NA | NA | NA | NA | NA | NA | NA | NA |
| D | - | - | - | - | NA | 5.394 | NA | NA | NA | NA | NA | NA | 3.967 | NA |
| E | - | - | - | - | - | NA | NA | NA | NA | NA | NA | NA | NA | NA |
| F | - | - | - | - | - | - | NA | NA | NA | NA | NA | NA | 4.058 | NA |
| G | - | - | - | - | - | - | - | NA | NA | NA | NA | NA | NA | NA |
| H | - | - | - | - | - | - | - | - | NA | NA | NA | NA | NA | NA |
| I | - | - | - | - | - | - | - | - | - | NA | NA | NA | NA | NA |
| J | - | - | - | - | - | - | - | - | - | - | NA | NA | NA | NA |
| K | - | - | - | - | - | - | - | - | - | - | - | NA | NA | NA |
| L | - | - | - | - | - | - | - | - | - | - | - | - | NA | NA |
| M | - | - | - | - | - | - | - | - | - | - | - | - | - | NA |
| N | - | - | - | - | - | - | - | - | - | - | - | - | - | - |

| Max | A | B | C | D | E | F | G | H | I | J | K | L | M | N |
|---|---|---|---|---|---|---|---|---|---|---|---|---|---|---|
| A | - | NA | NA | NA | NA | NA | NA | NA | NA | NA | NA | NA | NA | NA |
| B | - | - | NA | 6.917 | NA | 6.436 | NA | NA | NA | NA | NA | NA | 6.639 | NA |
| C | - | - | - | NA | NA | NA | NA | NA | NA | NA | NA | NA | NA | NA |
| D | - | - | - | - | NA | 3.935 | NA | NA | NA | NA | NA | NA | 3.249 | NA |
| E | - | - | - | - | - | NA | NA | NA | NA | NA | NA | NA | NA | NA |
| F | - | - | - | - | - | - | NA | NA | NA | NA | NA | NA | 2.791 | NA |
| G | - | - | - | - | - | - | - | NA | NA | NA | NA | NA | NA | NA |
| H | - | - | - | - | - | - | - | - | NA | NA | NA | NA | NA | NA |
| I | - | - | - | - | - | - | - | - | - | NA | NA | NA | NA | NA |
| J | - | - | - | - | - | - | - | - | - | - | NA | NA | NA | NA |
| K | - | - | - | - | - | - | - | - | - | - | - | NA | NA | NA |
| L | - | - | - | - | - | - | - | - | - | - | - | - | NA | NA |
| M | - | - | - | - | - | - | - | - | - | - | - | - | - | NA |
| N | - | - | - | - | - | - | - | - | - | - | - | - | - | - |
