## Supplementary Data S2 for "Interlaboratory comparison of standardised metabolomics and lipidomics analyses in human and rodent blood using the MxP® Quant 500 kit": p-Cresol-SO4.pdf

**Amount of relative differences between laboratories**

| Min | A | B | C | D | E | F | G | H | I | J | K | L | M | N |
|---|---|---|---|---|---|---|---|---|---|---|---|---|---|---|
| A | - | 6.696 | 7.827 | 6.5 | 4.876 | 7.921 | 5.792 | 8.029 | 7.681 | 5.842 | 6.169 | 4.314 | 7.71 | 7.475 |
| B | - | - | 6.712 | 7.52 | 6.88 | 7.091 | 5.575 | 7.252 | 6.658 | 4.514 | 4.645 | 6.049 | 6.657 | 6.61 |
| C | - | - | - | 7.962 | 7.533 | 5.535 | 7.045 | 5.9 | 3.197 | 6.509 | 6.672 | 7.096 | 3.389 | 4.726 |
| D | - | - | - | - | 5.871 | 8.414 | 7.226 | 8.503 | 8.229 | 7.043 | 7.362 | 6.667 | 8.291 | 7.966 |
| E | - | - | - | - | - | 8.222 | 6.574 | 8.317 | 8.018 | 6.552 | 6.794 | 5.703 | 8.041 | 7.803 |
| F | - | - | - | - | - | - | 7.652 | 3.593 | 5.446 | 7.136 | 7.357 | 7.606 | 5.395 | 3.452 |
| G | - | - | - | - | - | - | - | 7.715 | 7.257 | 4.001 | 4.567 | 4.682 | 7.276 | 7.125 |
| H | - | - | - | - | - | - | - | - | 5.763 | 7.196 | 7.402 | 7.638 | 5.71 | 4.354 |
| I | - | - | - | - | - | - | - | - | - | 6.661 | 6.825 | 7.224 | 0.468 | 4.474 |
| J | - | - | - | - | - | - | - | - | - | - | 0.203 | 5.742 | 7.287 | 7.247 |
| K | - | - | - | - | - | - | - | - | - | - | - | 5.529 | 7.113 | 6.961 |
| L | - | - | - | - | - | - | - | - | - | - | - | - | 7.693 | 7.527 |
| M | - | - | - | - | - | - | - | - | - | - | - | - | - | 4.085 |
| N | - | - | - | - | - | - | - | - | - | - | - | - | - | - |

| | |
|---|---|
| | Larger than 10 |
| | Between 8 and 9,99 |
| | Between 6 and 7,99 |
| | Between 4 and 5,99 |
| | Between 2 and 3,99 |
| | Between 0 and 1,99 |
| | NA (Calculation failed) |

| Median | A | B | C | D | E | F | G | H | I | J | K | L | M | N |
|---|---|---|---|---|---|---|---|---|---|---|---|---|---|---|
| A | - | 0.816 | 2.466 | 2.592 | 3.056 | 1.527 | -1.18 | -0.744 | -2.032 | 4.172 | 2.288 | 3.469 | -1.354 | 3.235 |
| B | - | - | 1.801 | 2.485 | 3.232 | 2.381 | 0.864 | 1.576 | 0.756 | 4.305 | 2.722 | 3.644 | 0.787 | 3.573 |
| C | - | - | - | 1.773 | 3.563 | 3.128 | 2.215 | 2.723 | 2.329 | 4.5 | 3.258 | 3.924 | 2.36 | 3.957 |
| D | - | - | - | - | 3.677 | 2.635 | 2.357 | 1.995 | 1.922 | 4.406 | 3.15 | 3.91 | 1.754 | 3.61 |
| E | - | - | - | - | - | 2.633 | 3.11 | 3.119 | 3.253 | 3.372 | 1.955 | 1.546 | 3.281 | -0.279 |
| F | - | - | - | - | - | - | 2.341 | 1.215 | 2.109 | 3.92 | -0.689 | 2.737 | 2.043 | 2.952 |
| G | - | - | - | - | - | - | - | -0.687 | -1.264 | 4.168 | 2.211 | 3.43 | -1.093 | 3.266 |
| H | - | - | - | - | - | - | - | - | 0.991 | 4.09 | 1.479 | 3.12 | 0.847 | 3.304 |
| I | - | - | - | - | - | - | - | - | - | 4.212 | 2.24 | 3.41 | -2.921 | 3.48 |
| J | - | - | - | - | - | - | - | - | - | - | 4.04 | 3.176 | 4.492 | 3.306 |
| K | - | - | - | - | - | - | - | - | - | - | - | 2.668 | 2.409 | 2.48 |
| L | - | - | - | - | - | - | - | - | - | - | - | - | 3.669 | 0.612 |
| M | - | - | - | - | - | - | - | - | - | - | - | - | - | 3.481 |
| N | - | - | - | - | - | - | - | - | - | - | - | - | - | - |

| Max | A | B | C | D | E | F | G | H | I | J | K | L | M | N |
|---|---|---|---|---|---|---|---|---|---|---|---|---|---|---|
| A | - | 2.52 | 3.073 | 3.718 | 2.754 | 4.34 | 2.125 | 4.241 | 3.797 | 4.588 | 3.393 | 3.754 | 3.861 | 4.681 |
| B | - | - | 1.533 | 4.29 | 0.379 | 3.929 | 0.483 | 3.792 | 3.133 | 4.287 | 2.393 | 3.054 | 3.178 | 4.419 |
| C | - | - | - | 4.465 | 0.185 | 3.661 | 2.104 | 3.491 | 2.586 | 4.1 | 1.492 | 2.505 | 2.643 | 4.26 |
| D | - | - | - | - | 4.179 | 4.945 | 3.999 | 4.89 | 4.629 | 5.063 | 4.43 | 4.594 | 4.699 | 5.126 |
| E | - | - | - | - | - | 3.858 | 1.353 | 3.702 | 2.944 | 4.223 | 1.985 | 2.852 | 3.07 | 4.34 |
| F | - | - | - | - | - | - | 4.288 | 0.713 | 2.929 | 2.35 | 3.501 | 3.009 | 2.891 | 2.9 |
| G | - | - | - | - | - | - | - | 3.941 | 3.337 | 4.371 | 2.745 | 3.265 | 3.401 | 4.495 |
| H | - | - | - | - | - | - | - | - | 2.554 | 2.738 | 3.229 | 2.632 | 2.502 | 3.163 |
| I | - | - | - | - | - | - | - | - | - | 3.576 | 1.687 | -2.13 | -2.058 | 3.812 |
| J | - | - | - | - | - | - | - | - | - | - | 4.135 | 3.8 | 3.696 | 1.278 |
| K | - | - | - | - | - | - | - | - | - | - | - | 1.698 | 2.068 | 4.075 |
| L | - | - | - | - | - | - | - | - | - | - | - | - | 0.202 | 3.84 |
| M | - | - | - | - | - | - | - | - | - | - | - | - | - | 3.766 |
| N | - | - | - | - | - | - | - | - | - | - | - | - | - | - |
