## Supplementary Data S2 for "Interlaboratory comparison of standardised metabolomics and lipidomics analyses in human and rodent blood using the MxP® Quant 500 kit": PC aa C28.1.pdf

**Amount of relative differences between laboratories**

| Min | A | B | C | D | E | F | G | H | I | J | K | L | M | N |
|---|---|---|---|---|---|---|---|---|---|---|---|---|---|---|
| A | - | 6.305 | 2.181 | 6.893 | 5.638 | 5.12 | 5.68 | 5.702 | 6.445 | 5.144 | 3.723 | 3.716 | 5.579 | 5.341 |
| B | - | - | 6.131 | 6.314 | 3.683 | 5.537 | 3.399 | 3.526 | 6.122 | 7.079 | 5.959 | 6.514 | 4.68 | 5.023 |
| C | - | - | - | 6.846 | 5.518 | 4.844 | 5.525 | 5.54 | 6.418 | 5.701 | 3.696 | 1.877 | 5.402 | 5.109 |
| D | - | - | - | - | 6.614 | 7.434 | 6.733 | 6.882 | 5.134 | 8.018 | 7.218 | 7.941 | 7.203 | 7.279 |
| E | - | - | - | - | - | 5.179 | -1.645 | -0.261 | 6.169 | 6.762 | 5.255 | 6.138 | 4.134 | 4.759 |
| F | - | - | - | - | - | - | 4.677 | 4.59 | 6.332 | 6.15 | 4.136 | 4.849 | 3.891 | 2.722 |
| G | - | - | - | - | - | - | - | 1.567 | 6.111 | 6.721 | 4.929 | 6.109 | 3.925 | 4.499 |
| H | - | - | - | - | - | - | - | - | 6.209 | 6.756 | 5.226 | 6.067 | 3.674 | 4.405 |
| I | - | - | - | - | - | - | - | - | - | 8.684 | 7.584 | 8.691 | 8.156 | 8.206 |
| J | - | - | - | - | - | - | - | - | - | - | 5.999 | 5.533 | 6.408 | 6.462 |
| K | - | - | - | - | - | - | - | - | - | - | - | 2.537 | 5.514 | 5.116 |
| L | - | - | - | - | - | - | - | - | - | - | - | - | 5.243 | 5.045 |
| M | - | - | - | - | - | - | - | - | - | - | - | - | - | 3.117 |
| N | - | - | - | - | - | - | - | - | - | - | - | - | - | - |

| | |
|---|---|
| | Larger than 10 |
| | Between 8 and 9,99 |
| | Between 6 and 7,99 |
| | Between 4 and 5,99 |
| | Between 2 and 3,99 |
| | Between 0 and 1,99 |
| | NA (Calculation failed) |

| Median | A | B | C | D | E | F | G | H | I | J | K | L | M | N |
|---|---|---|---|---|---|---|---|---|---|---|---|---|---|---|
| A | - | 2.551 | -1.716 | 2.453 | 3.743 | -1.197 | 2.999 | 1.876 | 3.606 | -0.295 | 2.836 | -1.866 | 1.1 | 1.166 |
| B | - | - | 2.727 | 3.601 | 3.004 | 2.749 | 3.881 | 1.226 | 4.272 | 2.791 | 3.795 | 2.57 | 3.094 | 3.112 |
| C | - | - | - | 2.364 | 3.772 | -3.008 | 2.936 | 1.991 | 3.565 | -1.249 | 2.763 | -0.752 | 0.869 | 0.943 |
| D | - | - | - | - | 4.171 | 2.278 | 1.176 | 3.145 | 2.692 | 2.301 | 0.599 | 2.478 | 1.706 | 1.668 |
| E | - | - | - | - | - | 3.998 | 4.627 | 3.49 | 4.889 | 4.034 | 4.57 | 3.923 | 4.165 | 4.174 |
| F | - | - | - | - | - | - | 2.911 | 2.036 | 3.556 | -1.65 | 2.738 | -0.495 | 0.759 | 0.837 |
| G | - | - | - | - | - | - | - | 3.435 | 1.978 | 2.763 | -0.361 | 2.935 | 2.435 | 2.411 |
| H | - | - | - | - | - | - | - | - | 4.053 | 2.188 | 3.492 | 1.824 | 2.598 | 2.621 |
| I | - | - | - | - | - | - | - | - | - | 3.481 | 2.284 | 3.515 | 3.22 | 3.211 |
| J | - | - | - | - | - | - | - | - | - | - | 2.647 | 0.222 | 0.377 | 0.545 |
| K | - | - | - | - | - | - | - | - | - | - | - | 2.791 | 2.222 | 2.191 |
| L | - | - | - | - | - | - | - | - | - | - | - | - | 1.278 | 1.344 |
| M | - | - | - | - | - | - | - | - | - | - | - | - | - | -3.513 |
| N | - | - | - | - | - | - | - | - | - | - | - | - | - | - |

| Max | A | B | C | D | E | F | G | H | I | J | K | L | M | N |
|---|---|---|---|---|---|---|---|---|---|---|---|---|---|---|
| A | - | 2.463 | -1.591 | 2.723 | 3.726 | -0.799 | 3.063 | 1.779 | 3.913 | -0.416 | 2.87 | -2.156 | 1.239 | 1.28 |
| B | - | - | 2.647 | 3.696 | 3.021 | 2.715 | 3.884 | 1.198 | 4.453 | 2.739 | 3.751 | 2.487 | 3.081 | 3.095 |
| C | - | - | - | 2.641 | 3.757 | -2.072 | 2.996 | 1.914 | 3.874 | -1.632 | 2.798 | -0.781 | 1.013 | 1.057 |
| D | - | - | - | - | 4.211 | 2.472 | 0.702 | 3.243 | 2.939 | 2.441 | -1.885 | 2.711 | 1.97 | 1.951 |
| E | - | - | - | - | - | 3.991 | 4.636 | 3.496 | 5.026 | 4.019 | 4.562 | 3.902 | 4.165 | 4.17 |
| F | - | - | - | - | - | - | 2.949 | 2.002 | 3.853 | -2.439 | 2.717 | -0.279 | 0.819 | 0.866 |
| G | - | - | - | - | - | - | - | 3.444 | 2.599 | 2.808 | 0.06 | 2.991 | 2.458 | 2.444 |
| H | - | - | - | - | - | - | - | - | 4.265 | 2.125 | 3.464 | 1.72 | 2.591 | 2.605 |
| I | - | - | - | - | - | - | - | - | - | 3.581 | 2.708 | 3.709 | 3.405 | 3.403 |
| J | - | - | - | - | - | - | - | - | - | - | 2.777 | -0.013 | 0.743 | 0.886 |
| K | - | - | - | - | - | - | - | - | - | - | - | 2.802 | 2.166 | 2.15 |
| L | - | - | - | - | - | - | - | - | - | - | - | - | 1.378 | 1.429 |
| M | - | - | - | - | - | - | - | - | - | - | - | - | - | -4.311 |
| N | - | - | - | - | - | - | - | - | - | - | - | - | - | - |
