## Supplementary Data S2 for "Interlaboratory comparison of standardised metabolomics and lipidomics analyses in human and rodent blood using the MxP® Quant 500 kit": PC aa C34.1.pdf

**Amount of relative differences between laboratories**

| Min | A | B | C | D | E | F | G | H | I | J | K | L | M | N |
|---|---|---|---|---|---|---|---|---|---|---|---|---|---|---|
| A | - | 5.518 | 3.387 | 1.003 | 5.092 | 4.47 | 4.945 | 6.489 | 4.886 | 6.81 | 4.324 | 2.424 | 4.468 | 4.255 |
| B | - | - | 5.097 | 5.595 | 1.787 | 4.566 | 6.27 | 5.359 | 3.962 | 5.51 | 4.548 | 5.579 | 4.568 | 4.305 |
| C | - | - | - | 3.281 | 4.6 | 3.344 | 5.345 | 6.201 | 4.206 | 6.385 | 3.219 | 3.889 | 3.498 | 3.366 |
| D | - | - | - | - | 5.017 | 4.309 | 5.073 | 6.429 | 4.797 | 6.659 | 4.129 | 2.901 | 4.352 | 4.134 |
| E | - | - | - | - | - | 4.226 | 6.348 | 5.931 | 3.246 | 6.268 | 4.311 | 5.617 | 4.211 | 4.142 |
| F | - | - | - | - | - | - | 5.752 | 6.083 | 2.95 | 6.33 | 0.312 | 4.701 | -3.243 | -0.618 |
| G | - | - | - | - | - | - | - | 6.905 | 5.94 | 7.144 | 5.598 | 4.529 | 5.743 | 5.543 |
| H | - | - | - | - | - | - | - | - | 5.623 | 1.095 | 5.751 | 6.215 | 5.838 | 5.541 |
| I | - | - | - | - | - | - | - | - | - | 6.016 | 3.121 | 5.012 | 2.876 | 2.615 |
| J | - | - | - | - | - | - | - | - | - | - | 5.915 | 6.461 | 6.035 | 5.744 |
| K | - | - | - | - | - | - | - | - | - | - | - | 4.563 | 0.246 | 1.038 |
| L | - | - | - | - | - | - | - | - | - | - | - | - | 4.913 | 4.7 |
| M | - | - | - | - | - | - | - | - | - | - | - | - | - | -0.712 |
| N | - | - | - | - | - | - | - | - | - | - | - | - | - | - |

| Legend |
|---|
| Larger than 10 |
| Between 8 and 9,99 |
| Between 6 and 7,99 |
| Between 4 and 5,99 |
| Between 2 and 3,99 |
| Between 0 and 1,99 |
| NA (Calculation failed) |

| Median | A | B | C | D | E | F | G | H | I | J | K | L | M | N |
|---|---|---|---|---|---|---|---|---|---|---|---|---|---|---|
| A | - | 1.4 | 1.177 | 0.613 | 4.185 | 2.065 | -9.345 | 2.935 | 0.313 | -0.422 | 0.003 | 2.411 | -2.089 | 3.735 |
| B | - | - | 2.31 | 0.444 | 3.993 | 0.591 | 1.589 | 3.404 | 0.659 | 1.456 | 1.8 | 1.374 | 1.364 | 3.435 |
| C | - | - | - | 1.857 | 4.32 | 2.645 | 1.072 | 2.406 | 1.75 | 0.824 | 0.365 | 2.877 | 1.257 | 3.923 |
| D | - | - | - | - | 4.081 | 1.43 | 0.637 | 3.221 | -1.775 | 1.128 | 1.357 | 1.944 | 0.394 | 3.582 |
| E | - | - | - | - | - | 4.099 | 4.482 | 4.98 | 4.376 | 4.505 | 4.542 | 3.981 | 4.458 | 2.586 |
| F | - | - | - | - | - | - | 2.154 | 3.623 | 1.631 | 2.249 | 2.396 | 0.206 | 2.046 | 3.248 |
| G | - | - | - | - | - | - | - | 2.977 | 0.179 | 0.366 | 0.487 | 2.386 | -3.45 | 3.727 |
| H | - | - | - | - | - | - | - | - | 3.002 | 2.943 | 2.745 | 3.557 | 2.839 | 4.264 |
| I | - | - | - | - | - | - | - | - | - | 0.479 | 1 | 2.041 | -0.013 | 3.61 |
| J | - | - | - | - | - | - | - | - | - | - | -0.016 | 2.18 | -1.14 | 3.656 |
| K | - | - | - | - | - | - | - | - | - | - | - | 2.599 | 0.064 | 3.807 |
| L | - | - | - | - | - | - | - | - | - | - | - | - | 2.429 | 3.08 |
| M | - | - | - | - | - | - | - | - | - | - | - | - | - | 3.71 |
| N | - | - | - | - | - | - | - | - | - | - | - | - | - | - |

| Max | A | B | C | D | E | F | G | H | I | J | K | L | M | N |
|---|---|---|---|---|---|---|---|---|---|---|---|---|---|---|
| A | - | -2.71 | 1.661 | 0.582 | 4.094 | 1.548 | 1.361 | 3.846 | -0.226 | 2.359 | 1.263 | 2.63 | 0.459 | 3.686 |
| B | - | - | 1.528 | 0.886 | 4.111 | 1.71 | 1.664 | 3.822 | -0.884 | 2.207 | 1.11 | 2.748 | 0.027 | 3.719 |
| C | - | - | - | 2.18 | 4.298 | 2.581 | 2.502 | 3.425 | 1.167 | 0.491 | -0.406 | 3.195 | 0.795 | 3.958 |
| D | - | - | - | - | 3.986 | 0.58 | 0.163 | 3.992 | 1.246 | 2.631 | 1.961 | 2.255 | 1.542 | 3.529 |
| E | - | - | - | - | - | 4.09 | 4.124 | 5.246 | 4.433 | 4.732 | 4.556 | 3.708 | 4.474 | 2.354 |
| F | - | - | - | - | - | - | -1.594 | 4.158 | 1.963 | 3.011 | 2.468 | 1.755 | 2.151 | 3.348 |
| G | - | - | - | - | - | - | - | 4.061 | 1.735 | 2.771 | 2.237 | 1.913 | 1.941 | 3.417 |
| H | - | - | - | - | - | - | - | - | 3.615 | 3.013 | 3.424 | 4.233 | 3.571 | 4.601 |
| I | - | - | - | - | - | - | - | - | - | 1.986 | 0.637 | 2.826 | -0.932 | 3.768 |
| J | - | - | - | - | - | - | - | - | - | - | 1.565 | 3.598 | 2.115 | 4.144 |
| K | - | - | - | - | - | - | - | - | - | - | - | 3.086 | 0.05 | 3.898 |
| L | - | - | - | - | - | - | - | - | - | - | - | - | 3 | 2.83 |
| M | - | - | - | - | - | - | - | - | - | - | - | - | - | 3.812 |
| N | - | - | - | - | - | - | - | - | - | - | - | - | - | - |
