## Supplementary Data S2 for "Interlaboratory comparison of standardised metabolomics and lipidomics analyses in human and rodent blood using the MxP® Quant 500 kit": PC aa C34.3.pdf

**Amount of relative differences between laboratories**

| Min | A | B | C | D | E | F | G | H | I | J | K | L | M | N |
|---|---|---|---|---|---|---|---|---|---|---|---|---|---|---|
| A | - | 4.318 | -0.632 | 2.047 | 4.232 | 2.256 | 3.524 | 2.927 | 2.316 | 4.841 | 3.013 | 0.534 | 3.027 | 2.445 |
| B | - | - | 4.484 | 4.806 | -5.522 | 4.083 | 5.17 | 3.804 | 4.067 | 2.705 | 3.743 | 4.371 | 3.745 | 4.013 |
| C | - | - | - | 2.212 | 4.165 | 2.101 | 3.558 | 2.811 | 2.167 | 4.708 | 2.884 | 0.208 | 2.932 | 2.349 |
| D | - | - | - | - | 4.488 | 3.09 | 2.983 | 3.555 | 3.115 | 5.059 | 3.551 | 2.273 | 3.628 | 3.231 |
| E | - | - | - | - | - | 4.089 | 5.158 | 3.81 | 4.081 | 3.047 | 3.706 | 4.345 | 3.74 | 4.019 |
| F | - | - | - | - | - | - | 4.077 | 1.571 | -2.82 | 4.528 | 1.713 | 1.781 | 1.826 | -0.504 |
| G | - | - | - | - | - | - | - | 4.104 | 3.903 | 5.18 | 4.139 | 3.558 | 4.165 | 3.96 |
| H | - | - | - | - | - | - | - | - | 1.336 | 4.227 | -0.859 | 2.544 | -0.915 | 0.966 |
| I | - | - | - | - | - | - | - | - | - | 4.455 | 1.589 | 1.75 | 1.725 | -0.694 |
| J | - | - | - | - | - | - | - | - | - | - | 4.003 | 4.561 | 4.104 | 4.232 |
| K | - | - | - | - | - | - | - | - | - | - | - | 2.488 | -2.159 | 1.27 |
| L | - | - | - | - | - | - | - | - | - | - | - | - | 2.739 | 2.025 |
| M | - | - | - | - | - | - | - | - | - | - | - | - | - | 1.365 |
| N | - | - | - | - | - | - | - | - | - | - | - | - | - | - |

| | |
|---|---|
| | Larger than 10 |
| | Between 8 and 9,99 |
| | Between 6 and 7,99 |
| | Between 4 and 5,99 |
| | Between 2 and 3,99 |
| | Between 0 and 1,99 |
| | NA (Calculation failed) |

| Median | A | B | C | D | E | F | G | H | I | J | K | L | M | N |
|---|---|---|---|---|---|---|---|---|---|---|---|---|---|---|
| A | - | 3.73 | 1.008 | 1.044 | 3.78 | 1.996 | 3.341 | 0.964 | 1.049 | 2.371 | 1.784 | -0.564 | -3.946 | 2.058 |
| B | - | - | 4.127 | 3.691 | -0.863 | 3.413 | 4.759 | 3.687 | 3.676 | 3.191 | 4.265 | 3.882 | 3.914 | 3.385 |
| C | - | - | - | 1.982 | 3.943 | 2.549 | 3.008 | 1.961 | 1.999 | 2.82 | 0.526 | 1.335 | 1.031 | 2.593 |
| D | - | - | - | - | 3.576 | 0.952 | 3.63 | -2.047 | -3.374 | 1.569 | 2.505 | 0.318 | 1.108 | 1.085 |
| E | - | - | - | - | - | 3.494 | 4.794 | 3.757 | 3.747 | 3.282 | 4.312 | 3.938 | 3.974 | 3.468 |
| F | - | - | - | - | - | - | 3.88 | 1.077 | 0.996 | 0.294 | 2.954 | 1.799 | 2.026 | -2.453 |
| G | - | - | - | - | - | - | - | 3.45 | 3.466 | 3.791 | 2.596 | 3.3 | 3.206 | 3.698 |
| H | - | - | - | - | - | - | - | - | -3.476 | 1.731 | 2.485 | 0.666 | 0.952 | 1.119 |
| I | - | - | - | - | - | - | - | - | - | 1.674 | 2.503 | 0.633 | 1.022 | 1.08 |
| J | - | - | - | - | - | - | - | - | - | - | 3.262 | 2.593 | 2.495 | 0.633 |
| K | - | - | - | - | - | - | - | - | - | - | - | 1.848 | 1.827 | 2.878 |
| L | - | - | - | - | - | - | - | - | - | - | - | - | -0.804 | 1.826 |
| M | - | - | - | - | - | - | - | - | - | - | - | - | - | 1.995 |
| N | - | - | - | - | - | - | - | - | - | - | - | - | - | - |

| Max | A | B | C | D | E | F | G | H | I | J | K | L | M | N |
|---|---|---|---|---|---|---|---|---|---|---|---|---|---|---|
| A | - | 3.527 | 1.478 | 1.946 | 3.604 | 1.907 | 3.265 | -1.22 | 0.257 | -0.613 | 2.868 | -1.452 | 1.229 | 1.916 |
| B | - | - | 4.003 | 3.098 | -0.484 | 3.126 | 4.564 | 3.654 | 3.535 | 3.602 | 4.404 | 3.627 | 3.96 | 3.123 |
| C | - | - | - | 2.686 | 3.866 | 2.662 | 2.726 | 1.611 | 1.944 | 1.761 | 2.135 | 1.621 | -1.401 | 2.659 |
| D | - | - | - | - | 3.113 | -3.808 | 3.817 | 1.794 | 1.447 | 1.524 | 3.523 | 1.886 | 2.696 | -3.427 |
| E | - | - | - | - | - | 3.25 | 4.626 | 3.742 | 3.629 | 3.721 | 4.469 | 3.734 | 4.037 | 3.247 |
| F | - | - | - | - | - | - | 3.797 | 1.797 | 1.398 | 1.574 | 3.516 | 1.807 | 2.665 | -5.406 |
| G | - | - | - | - | - | - | - | 3.208 | 3.304 | 3.252 | 1.119 | 3.182 | 2.729 | 3.607 |
| H | - | - | - | - | - | - | - | - | -0.193 | -1.492 | 2.937 | -3.015 | 1.417 | 1.777 |
| I | - | - | - | - | - | - | - | - | - | -1.269 | 3.1 | -0.286 | 1.856 | 1.362 |
| J | - | - | - | - | - | - | - | - | - | - | 2.895 | 0.064 | 1.135 | 1.888 |
| K | - | - | - | - | - | - | - | - | - | - | - | 2.87 | 2.224 | 3.368 |
| L | - | - | - | - | - | - | - | - | - | - | - | - | 1.45 | 1.759 |
| M | - | - | - | - | - | - | - | - | - | - | - | - | - | 2.581 |
| N | - | - | - | - | - | - | - | - | - | - | - | - | - | - |
