## Supplementary Data S2 for "Interlaboratory comparison of standardised metabolomics and lipidomics analyses in human and rodent blood using the MxP® Quant 500 kit": PC aa C36.1.pdf

**Amount of relative differences between laboratories**

| Min | A | B | C | D | E | F | G | H | I | J | K | L | M | N |
|---|---|---|---|---|---|---|---|---|---|---|---|---|---|---|
| A | - | 4.409 | 3.063 | 2.884 | 5.66 | 4.771 | 2.994 | 4.907 | 3.544 | 6.108 | 4.651 | 4.015 | 4.467 | 5.473 |
| B | - | - | 3.857 | 4.01 | 5.044 | 2.866 | 3.614 | 3.007 | 3.299 | 5.28 | 1.906 | 1.847 | -0.063 | 4.724 |
| C | - | - | - | -0.615 | 5.392 | 4.329 | 0.747 | 4.349 | 2.056 | 5.525 | 4.064 | 3.341 | 3.872 | 5.179 |
| D | - | - | - | - | 5.494 | 4.411 | 0.615 | 4.542 | 2.17 | 5.854 | 4.165 | 3.335 | 3.979 | 5.288 |
| E | - | - | - | - | - | 4.875 | 5.726 | 5.033 | 5.697 | 2.942 | 5.103 | 5.342 | 5.26 | 3.074 |
| F | - | - | - | - | - | - | 4.412 | -0.665 | 4.242 | 4.866 | 1.959 | 3.46 | 2.818 | 4.23 |
| G | - | - | - | - | - | - | - | 4.23 | 1.217 | 5.57 | 3.888 | 3.111 | 3.689 | 5.123 |
| H | - | - | - | - | - | - | - | - | 4.155 | 4.712 | 1.722 | 3.308 | 2.592 | 4.229 |
| I | - | - | - | - | - | - | - | - | - | 5.661 | 3.736 | 2.616 | 3.481 | 5.127 |
| J | - | - | - | - | - | - | - | - | - | - | 4.585 | 5.015 | 4.877 | -3.827 |
| K | - | - | - | - | - | - | - | - | - | - | - | 2.502 | 1.503 | 4.39 |
| L | - | - | - | - | - | - | - | - | - | - | - | - | 2.133 | 4.812 |
| M | - | - | - | - | - | - | - | - | - | - | - | - | - | 4.581 |
| N | - | - | - | - | - | - | - | - | - | - | - | - | - | - |

| Median | A | B | C | D | E | F | G | H | I | J | K | L | M | N |
|---|---|---|---|---|---|---|---|---|---|---|---|---|---|---|
| A | - | 2.651 | 2.817 | 1.594 | 3.544 | 2.321 | 2.719 | 1.544 | 1.793 | 1.736 | 2.925 | -1.192 | -0.142 | 2.467 |
| B | - | - | 3.821 | 1.804 | 2.576 | 0.332 | 3.77 | 1.834 | 1.56 | 1.371 | 3.866 | 2.611 | 2.933 | 3.651 |
| C | - | - | - | 3.223 | 4.138 | 3.497 | -1.226 | 3.213 | 3.293 | 3.339 | -1.373 | 2.815 | 2.514 | 0.471 |
| D | - | - | - | - | 3.166 | 1.051 | 3.304 | -3.381 | -1.075 | -0.993 | 3.444 | 1.391 | 2.011 | 3.137 |
| E | - | - | - | - | - | 2.985 | 4.412 | 3.4 | 3.304 | 3.102 | 4.454 | 3.71 | 3.898 | 4.334 |
| F | - | - | - | - | - | - | 3.62 | 1.22 | 0.785 | 0.161 | 3.711 | 2.261 | 2.668 | 3.488 |
| G | - | - | - | - | - | - | - | 3.151 | 3.235 | 3.283 | -0.314 | 2.732 | 2.407 | -0.056 |
| H | - | - | - | - | - | - | - | - | -0.749 | 0.328 | 3.393 | 1.262 | 1.975 | 3.121 |
| I | - | - | - | - | - | - | - | - | - | -2.678 | 3.507 | 1.629 | 2.181 | 3.221 |
| J | - | - | - | - | - | - | - | - | - | - | 3.618 | 2.065 | 2.551 | 3.416 |
| K | - | - | - | - | - | - | - | - | - | - | - | 2.867 | 2.553 | 0.645 |
| L | - | - | - | - | - | - | - | - | - | - | - | - | 0.537 | 2.609 |
| M | - | - | - | - | - | - | - | - | - | - | - | - | - | 2.196 |
| N | - | - | - | - | - | - | - | - | - | - | - | - | - | - |

| Max | A | B | C | D | E | F | G | H | I | J | K | L | M | N |
|---|---|---|---|---|---|---|---|---|---|---|---|---|---|---|
| A | - | 1.863 | 3.274 | 1.224 | 2.41 | 0.127 | 3.241 | 0.462 | 1.081 | 1.988 | 3.746 | 1.363 | 2.342 | 4.041 |
| B | - | - | 3.815 | 0.605 | 0.634 | 1.462 | 3.799 | 2.434 | 0.858 | 2.847 | 4.157 | 2.72 | 3.201 | 4.399 |
| C | - | - | - | 3.446 | 3.769 | 3.286 | -2.471 | 2.909 | 3.424 | 2.615 | 1.78 | 2.689 | 2.034 | 2.641 |
| D | - | - | - | - | 1.626 | 0.364 | 3.6 | 1.928 | -2.405 | 2.555 | 4.001 | 2.338 | 2.929 | 4.269 |
| E | - | - | - | - | - | 2.06 | 3.91 | 2.816 | 1.566 | 3.119 | 4.3 | 2.97 | 3.415 | 4.507 |
| F | - | - | - | - | - | - | 3.395 | 1.378 | 0.153 | 2.166 | 3.881 | 1.85 | 2.639 | 4.153 |
| G | - | - | - | - | - | - | - | 2.871 | 3.404 | 2.467 | 1.842 | 2.647 | 1.974 | 2.69 |
| H | - | - | - | - | - | - | - | - | 1.859 | 0.604 | 3.559 | -0.054 | 1.828 | 3.878 |
| I | - | - | - | - | - | - | - | - | - | 2.447 | 3.979 | 2.28 | 2.883 | 4.249 |
| J | - | - | - | - | - | - | - | - | - | - | 3.35 | 0.262 | 0.679 | 3.703 |
| K | - | - | - | - | - | - | - | - | - | - | - | 3.289 | 2.896 | 1.424 |
| L | - | - | - | - | - | - | - | - | - | - | - | - | 1.326 | 3.764 |
| M | - | - | - | - | - | - | - | - | - | - | - | - | - | 3.436 |
| N | - | - | - | - | - | - | - | - | - | - | - | - | - | - |

| Colour | Legend |
|---|---|
| (red) | Larger than 10 |
| (orange) | Between 8 and 9,99 |
| (light orange) | Between 6 and 7,99 |
| (light green) | Between 4 and 5,99 |
| (green) | Between 2 and 3,99 |
| (dark green) | Between 0 and 1,99 |
| (grey) | NA (Calculation failed) |
