## Supplementary Data S2 for "Interlaboratory comparison of standardised metabolomics and lipidomics analyses in human and rodent blood using the MxP® Quant 500 kit": PC aa C36.2.pdf

**Amount of relative differences between laboratories**

| Min | A | B | C | D | E | F | G | H | I | J | K | L | M | N |
|---|---|---|---|---|---|---|---|---|---|---|---|---|---|---|
| A | - | 3.701 | 1.481 | 3.042 | 4.436 | 4.065 | -1.181 | 4.499 | 3.769 | 4.705 | 2.697 | 1.44 | 3.448 | 4.034 |
| B | - | - | 3.475 | 2.624 | 3.417 | 1.894 | 3.837 | 3.396 | -1.781 | 3.027 | 2.5 | 3.163 | 0.995 | 2.29 |
| C | - | - | - | 2.313 | 4.274 | 3.789 | 1.559 | 4.325 | 3.421 | 4.335 | 2.128 | 0.126 | 3.058 | 3.843 |
| D | - | - | - | - | 3.95 | 3.235 | 3.028 | 3.979 | 2.577 | 3.915 | -0.934 | 1.944 | 1.866 | 3.297 |
| E | - | - | - | - | - | 2.725 | 4.791 | 0.189 | 3.404 | 0.941 | 4.25 | 4.546 | 3.78 | 2.704 |
| F | - | - | - | - | - | - | 4.212 | 2.764 | 1.938 | 2.349 | 3.219 | 3.688 | 2.616 | 0.639 |
| G | - | - | - | - | - | - | - | 4.526 | 3.797 | 4.651 | 2.747 | 1.692 | 3.511 | 4.067 |
| H | - | - | - | - | - | - | - | - | 3.481 | 2.699 | 3.837 | 4.1 | 3.672 | 2.004 |
| I | - | - | - | - | - | - | - | - | - | 2.883 | 2.483 | 3.12 | 1.074 | 2.262 |
| J | - | - | - | - | - | - | - | - | - | - | 3.566 | 4.002 | 3.212 | 0.33 |
| K | - | - | - | - | - | - | - | - | - | - | - | 1.766 | 2.092 | 3.379 |
| L | - | - | - | - | - | - | - | - | - | - | - | - | 3.221 | 3.849 |
| M | - | - | - | - | - | - | - | - | - | - | - | - | - | 2.759 |
| N | - | - | - | - | - | - | - | - | - | - | - | - | - | - |

| Median | A | B | C | D | E | F | G | H | I | J | K | L | M | N |
|---|---|---|---|---|---|---|---|---|---|---|---|---|---|---|
| A | - | 1.813 | 1.571 | 0.733 | 4.267 | 2.427 | -0.21 | -0.372 | -0.085 | 2.129 | 1.359 | 2.447 | 0.826 | 3.567 |
| B | - | - | 2.719 | 0.815 | 4.051 | 0.958 | 2.167 | 2.105 | 1.077 | -0.756 | 0.337 | 0.911 | 0.728 | 3.128 |
| C | - | - | - | 2.175 | 4.437 | 3.009 | 1.005 | 1.105 | 2.01 | 2.79 | 2.364 | 3.008 | 2.208 | 3.847 |
| D | - | - | - | - | 4.167 | 1.888 | 1.394 | 1.313 | -1.011 | 1.293 | -0.635 | 1.883 | -3.17 | 3.369 |
| E | - | - | - | - | - | 4.111 | 4.643 | 4.635 | 4.498 | 4.226 | 4.384 | 4.107 | 4.44 | 3.205 |
| F | - | - | - | - | - | - | 2.714 | 2.669 | 2.014 | 0.418 | 1.731 | -3.094 | 1.874 | 2.797 |
| G | - | - | - | - | - | - | - | -3.277 | 0.857 | 2.356 | 1.751 | 2.646 | 1.393 | 3.657 |
| H | - | - | - | - | - | - | - | - | 1.451 | 2.118 | 1.218 | 2.462 | 1.347 | 3.64 |
| I | - | - | - | - | - | - | - | - | - | 1.328 | -0.65 | 1.908 | -1.49 | 3.396 |
| J | - | - | - | - | - | - | - | - | - | - | 0.993 | 0.201 | 1.096 | 3.052 |
| K | - | - | - | - | - | - | - | - | - | - | - | 1.608 | -0.738 | 3.279 |
| L | - | - | - | - | - | - | - | - | - | - | - | - | 1.997 | 2.74 |
| M | - | - | - | - | - | - | - | - | - | - | - | - | - | 3.355 |
| N | - | - | - | - | - | - | - | - | - | - | - | - | - | - |

| Max | A | B | C | D | E | F | G | H | I | J | K | L | M | N |
|---|---|---|---|---|---|---|---|---|---|---|---|---|---|---|
| A | - | 0.687 | 2.589 | 0.301 | 4.181 | -1.027 | 0.104 | 3.845 | 2.176 | 2.389 | -3.178 | 2.829 | 1.528 | 3.242 |
| B | - | - | 2.068 | -2.479 | 4.272 | 0.037 | -1.429 | 3.692 | 1.652 | 1.461 | 0.797 | 3.071 | 0.57 | 3.43 |
| C | - | - | - | 2.111 | 4.507 | 2.342 | 2.158 | 3.045 | 0.207 | 0.061 | 2.506 | 3.612 | 1.45 | 3.85 |
| D | - | - | - | - | 4.258 | -0.413 | -3.125 | 3.711 | 1.713 | 1.654 | 0.527 | 3.04 | 0.756 | 3.406 |
| E | - | - | - | - | - | 4.51 | 4.563 | 5.32 | 4.795 | 4.799 | 4.469 | 3.759 | 4.687 | 3.413 |
| F | - | - | - | - | - | - | -0.68 | 3.808 | 2.045 | 2.065 | -0.375 | 2.92 | 1.269 | 3.294 |
| G | - | - | - | - | - | - | - | 3.726 | 1.761 | 1.903 | 0.108 | 3.015 | 0.881 | 3.381 |
| H | - | - | - | - | - | - | - | - | 3.123 | 3.192 | 3.774 | 4.26 | 3.369 | 4.405 |
| I | - | - | - | - | - | - | - | - | - | -1.486 | 2.29 | 3.492 | 0.686 | 3.762 |
| J | - | - | - | - | - | - | - | - | - | - | 2.245 | 3.522 | 0.577 | 3.739 |
| K | - | - | - | - | - | - | - | - | - | - | - | 2.793 | 1.621 | 3.223 |
| L | - | - | - | - | - | - | - | - | - | - | - | - | 3.429 | 1.375 |
| M | - | - | - | - | - | - | - | - | - | - | - | - | - | 3.592 |
| N | - | - | - | - | - | - | - | - | - | - | - | - | - | - |

| | |
|---|---|
| | Larger than 10 |
| | Between 8 and 9,99 |
| | Between 6 and 7,99 |
| | Between 4 and 5,99 |
| | Between 2 and 3,99 |
| | Between 0 and 1,99 |
| | NA (Calculation failed) |
