## Supplementary Data S2 for "Interlaboratory comparison of standardised metabolomics and lipidomics analyses in human and rodent blood using the MxP® Quant 500 kit": PC aa C36.3.pdf

**Amount of relative differences between laboratories**

| Min | A | B | C | D | E | F | G | H | I | J | K | L | M | N |
|---|---|---|---|---|---|---|---|---|---|---|---|---|---|---|
| A | - | 4.155 | -1.223 | 0.828 | 4.712 | 3.514 | 2.983 | 4.871 | 2.852 | 5.606 | 4.047 | 0.925 | 3.132 | 3.399 |
| B | - | - | 4.205 | 4.364 | 3.568 | 2.539 | 4.683 | 3.682 | 3.389 | 4.611 | -0.584 | 3.807 | 3.067 | 2.751 |
| C | - | - | - | 0.576 | 4.706 | 3.507 | 2.775 | 4.829 | 2.861 | 5.385 | 3.961 | 1.605 | 3.207 | 3.432 |
| D | - | - | - | - | 4.787 | 3.702 | 2.6 | 4.945 | 3.14 | 5.54 | 4.099 | 1.925 | 3.423 | 3.65 |
| E | - | - | - | - | - | 4.294 | 5.396 | 0.772 | 4.675 | 4.14 | 3.755 | 4.879 | 4.523 | 4.386 |
| F | - | - | - | - | - | - | 4.311 | 4.214 | 2.207 | 5.055 | 2.508 | 3.116 | 1.513 | 0.352 |
| G | - | - | - | - | - | - | - | 5.137 | 3.824 | 5.673 | 4.464 | 3.206 | 3.989 | 4.148 |
| H | - | - | - | - | - | - | - | - | 4.555 | 2.895 | 3.647 | 4.669 | 4.41 | 4.313 |
| I | - | - | - | - | - | - | - | - | - | 5.161 | 3.168 | 2.078 | 1.013 | 1.894 |
| J | - | - | - | - | - | - | - | - | - | - | 4.241 | 5.087 | 4.877 | 4.787 |
| K | - | - | - | - | - | - | - | - | - | - | - | 3.699 | 2.998 | 2.665 |
| L | - | - | - | - | - | - | - | - | - | - | - | - | 2.783 | 3.14 |
| M | - | - | - | - | - | - | - | - | - | - | - | - | - | 0.591 |
| N | - | - | - | - | - | - | - | - | - | - | - | - | - | - |

| | |
|---|---|
| | Larger than 10 |
| | Between 8 and 9,99 |
| | Between 6 and 7,99 |
| | Between 4 and 5,99 |
| | Between 2 and 3,99 |
| | Between 0 and 1,99 |
| | NA (Calculation failed) |

| Median | A | B | C | D | E | F | G | H | I | J | K | L | M | N |
|---|---|---|---|---|---|---|---|---|---|---|---|---|---|---|
| A | - | 1.143 | 2.563 | -0.513 | 3.906 | 1.347 | 2.557 | -0.153 | -2.45 | -0.462 | 0.421 | 0.716 | 1.436 | -0.825 |
| B | - | - | 3.033 | 1.574 | 3.715 | -1.119 | 3.054 | 1.552 | 1.193 | 1.342 | 1.928 | -0.469 | 2.265 | 0.614 |
| C | - | - | - | 2.288 | 4.302 | 2.997 | -5.438 | 2.256 | 2.439 | 2.336 | 2.075 | 2.818 | 1.613 | 2.615 |
| D | - | - | - | - | 3.962 | 1.684 | 2.364 | -2.221 | -0.988 | -6.47 | -0.647 | 1.227 | 1.006 | 0.316 |
| E | - | - | - | - | - | 3.867 | 4.624 | 4.21 | 4.15 | 4.179 | 4.282 | 3.992 | 4.369 | 4.078 |
| F | - | - | - | - | - | - | 3.128 | 1.79 | 1.472 | 1.653 | 2.054 | 0.107 | 2.414 | 1.008 |
| G | - | - | - | - | - | - | - | 2.225 | 2.423 | 2.284 | 2.103 | 2.834 | 1.572 | 2.601 |
| H | - | - | - | - | - | - | - | - | -0.892 | -0.908 | 0.34 | 0.805 | 0.891 | 0.344 |
| I | - | - | - | - | - | - | - | - | - | -1.46 | 0.401 | 0.753 | 1.319 | -0.431 |
| J | - | - | - | - | - | - | - | - | - | - | 1.014 | -0.024 | 1.294 | -0.601 |
| K | - | - | - | - | - | - | - | - | - | - | - | 1.57 | -0.184 | 1.176 |
| L | - | - | - | - | - | - | - | - | - | - | - | - | 2.142 | 0.125 |
| M | - | - | - | - | - | - | - | - | - | - | - | - | - | 1.651 |
| N | - | - | - | - | - | - | - | - | - | - | - | - | - | - |

| Max | A | B | C | D | E | F | G | H | I | J | K | L | M | N |
|---|---|---|---|---|---|---|---|---|---|---|---|---|---|---|
| A | - | -3.202 | 2.747 | -0.838 | 3.74 | 0.45 | 2.475 | 2.347 | 0.212 | 2.588 | 2.25 | 0.677 | 2.091 | -0.475 |
| B | - | - | 2.742 | -0.602 | 3.735 | 0.369 | 2.459 | 2.356 | 0.272 | 2.438 | 2.14 | 0.637 | 2.112 | -0.214 |
| C | - | - | - | 2.516 | 4.233 | 2.914 | 0.109 | 0.579 | 2.373 | 0.164 | 1.027 | 2.952 | 1.151 | 2.466 |
| D | - | - | - | - | 3.79 | 0.937 | 2.322 | 2.176 | -0.759 | 2.311 | 2.014 | 1.065 | 1.9 | -2.255 |
| E | - | - | - | - | - | 3.799 | 4.449 | 4.414 | 4.064 | 4.455 | 4.364 | 3.765 | 4.347 | 4.033 |
| F | - | - | - | - | - | - | 2.797 | 2.704 | 1.34 | 2.815 | 2.594 | -2.149 | 2.502 | 1.074 |
| G | - | - | - | - | - | - | - | -1.194 | 2.072 | -3.953 | -0.156 | 2.759 | 0.292 | 2.192 |
| H | - | - | - | - | - | - | - | - | 1.891 | -2.283 | -0.169 | 2.681 | -0.369 | 1.995 |
| I | - | - | - | - | - | - | - | - | - | 2.08 | 1.719 | 1.432 | 1.635 | -1.476 |
| J | - | - | - | - | - | - | - | - | - | - | 0.279 | 2.804 | 0.259 | 2.102 |
| K | - | - | - | - | - | - | - | - | - | - | - | 2.513 | -3.606 | 1.751 |
| L | - | - | - | - | - | - | - | - | - | - | - | - | 2.571 | 1.241 |
| M | - | - | - | - | - | - | - | - | - | - | - | - | - | 1.744 |
| N | - | - | - | - | - | - | - | - | - | - | - | - | - | - |
