## Supplementary Data S2 for "Interlaboratory comparison of standardised metabolomics and lipidomics analyses in human and rodent blood using the MxP® Quant 500 kit": PC ae C38.0.pdf

**Amount of relative differences between laboratories**

| Min | A | B | C | D | E | F | G | H | I | J | K | L | M | N |
|---|---|---|---|---|---|---|---|---|---|---|---|---|---|---|
| A | - | 3.647 | 1.554 | 0.509 | 3.69 | 3.205 | 3.547 | 1.661 | 1.078 | 4.014 | 1.658 | 2.571 | -0.768 | -2.868 |
| B | - | - | 4.061 | 4.045 | -0.122 | 2.076 | 4.791 | 3.378 | 3.472 | 1.306 | 4.177 | 2.877 | 3.736 | 3.784 |
| C | - | - | - | -4.481 | 3.93 | 3.519 | 3.297 | 2.509 | 2.174 | 4.041 | 0.497 | 3.03 | 1.777 | 1.401 |
| D | - | - | - | - | 3.771 | 3.244 | 3.509 | 1.953 | 2.218 | 4.229 | 1.43 | 2.649 | -0.785 | 0.726 |
| E | - | - | - | - | - | 1.879 | 4.701 | 3.356 | 3.579 | -0.25 | 4.065 | 2.753 | 3.76 | 3.869 |
| F | - | - | - | - | - | - | 4.471 | 2.706 | 2.529 | 2.657 | 3.761 | 1.716 | 3.254 | 3.378 |
| G | - | - | - | - | - | - | - | 3.69 | 3.577 | 4.581 | 3.051 | 3.96 | 3.316 | 3.33 |
| H | - | - | - | - | - | - | - | - | -0.665 | 3.609 | 2.754 | 1.485 | 1.414 | 1.781 |
| I | - | - | - | - | - | - | - | - | - | 3.299 | 2.173 | 2.267 | 0.609 | 1.077 |
| J | - | - | - | - | - | - | - | - | - | - | 4.098 | 3.174 | 3.814 | 3.856 |
| K | - | - | - | - | - | - | - | - | - | - | - | 3.188 | 2.156 | 2.021 |
| L | - | - | - | - | - | - | - | - | - | - | - | - | 2.638 | 2.717 |
| M | - | - | - | - | - | - | - | - | - | - | - | - | - | 0.017 |
| N | - | - | - | - | - | - | - | - | - | - | - | - | - | - |

| | |
|---|---|
| | Larger than 10 |
| | Between 8 and 9,99 |
| | Between 6 and 7,99 |
| | Between 4 and 5,99 |
| | Between 2 and 3,99 |
| | Between 0 and 1,99 |
| | NA (Calculation failed) |

| Median | A | B | C | D | E | F | G | H | I | J | K | L | M | N |
|---|---|---|---|---|---|---|---|---|---|---|---|---|---|---|
| A | - | 3.602 | 0.159 | 2.367 | 2.289 | 3.156 | 4.127 | 2.028 | 2.849 | 2.517 | 0.913 | 0.358 | 2.231 | 1.649 |
| B | - | - | 3.914 | 2.992 | 3.044 | 1.876 | 5.076 | 3.197 | 4.46 | 2.855 | 3.996 | 3.628 | 4.26 | 4.119 |
| C | - | - | - | 2.63 | 2.573 | 3.309 | 4.013 | 2.367 | 2.591 | 2.758 | -0.371 | 1.258 | 1.812 | 0.969 |
| D | - | - | - | - | -1.959 | 1.96 | 4.554 | 0.113 | 3.706 | -1.881 | 2.905 | 2.006 | 3.373 | 3.104 |
| E | - | - | - | - | - | 2.133 | 4.58 | -0.203 | 3.655 | 0.516 | 2.799 | 1.939 | 3.334 | 3.095 |
| F | - | - | - | - | - | - | 4.854 | 2.409 | 4.143 | 1.787 | 3.567 | 3.067 | 3.902 | 3.727 |
| G | - | - | - | - | - | - | - | 4.212 | 3.176 | 4.338 | 3.764 | 4.013 | 3.47 | 3.629 |
| H | - | - | - | - | - | - | - | - | 3.558 | 0.765 | 2.641 | 1.543 | 3.193 | 2.91 |
| I | - | - | - | - | - | - | - | - | - | 3.647 | 2.323 | 2.959 | 1.149 | 1.845 |
| J | - | - | - | - | - | - | - | - | - | - | 3.084 | 2.419 | 3.54 | 3.327 |
| K | - | - | - | - | - | - | - | - | - | - | - | 1.652 | 1.441 | 0.21 |
| L | - | - | - | - | - | - | - | - | - | - | - | - | 2.593 | 2.169 |
| M | - | - | - | - | - | - | - | - | - | - | - | - | - | 0.514 |
| N | - | - | - | - | - | - | - | - | - | - | - | - | - | - |

| Max | A | B | C | D | E | F | G | H | I | J | K | L | M | N |
|---|---|---|---|---|---|---|---|---|---|---|---|---|---|---|
| A | - | 3.592 | -0.57 | 2.752 | 1.387 | 3.144 | 4.33 | 2.116 | 3.308 | 2.006 | 0.608 | -1.767 | 2.632 | 2.002 |
| B | - | - | 3.876 | 2.563 | 3.439 | 1.822 | 5.19 | 3.143 | 4.654 | 3.141 | 3.941 | 3.813 | 4.38 | 4.203 |
| C | - | - | - | 2.942 | 1.732 | 3.252 | 4.251 | 2.324 | 3.183 | 2.372 | -0.752 | -1.191 | 2.456 | 1.719 |
| D | - | - | - | - | 2.069 | 1.326 | 4.86 | 1.305 | 4.185 | 1.461 | 3.144 | 2.906 | 3.752 | 3.533 |
| E | - | - | - | - | - | 2.675 | 4.52 | 0.815 | 3.676 | 0.61 | 2.073 | 1.512 | 3.185 | 2.762 |
| F | - | - | - | - | - | - | 5.001 | 2.314 | 4.401 | 2.336 | 3.508 | 3.336 | 4.043 | 3.815 |
| G | - | - | - | - | - | - | - | 4.33 | 3.035 | 4.293 | 3.914 | 4.031 | 3.511 | 3.706 |
| H | - | - | - | - | - | - | - | - | 3.907 | -5.661 | 2.607 | 2.277 | 3.459 | 3.121 |
| I | - | - | - | - | - | - | - | - | - | 3.696 | 2.942 | 3.133 | 1.715 | 2.391 |
| J | - | - | - | - | - | - | - | - | - | - | 2.596 | 2.037 | 3.451 | 3.129 |
| K | - | - | - | - | - | - | - | - | - | - | - | 0.169 | 2.25 | 1.416 |
| L | - | - | - | - | - | - | - | - | - | - | - | - | 2.58 | 1.961 |
| M | - | - | - | - | - | - | - | - | - | - | - | - | - | 1.095 |
| N | - | - | - | - | - | - | - | - | - | - | - | - | - | - |
