## Supplementary Data S2 for "Interlaboratory comparison of standardised metabolomics and lipidomics analyses in human and rodent blood using the MxP® Quant 500 kit": PC ae C38.1.pdf

**Amount of relative differences between laboratories**

| Min | A | B | C | D | E | F | G | H | I | J | K | L | M | N |
|---|---|---|---|---|---|---|---|---|---|---|---|---|---|---|
| A | - | 6.411 | 6.71 | 7.065 | 6.326 | 6.189 | 7.011 | 6.355 | 6.572 | 4.556 | 6.988 | 5.003 | 6.549 | 7.118 |
| B | - | - | 4.486 | 6.499 | 5.399 | 5.726 | 6.532 | 4.49 | 5.856 | 6.165 | 6.03 | 8.174 | 5.591 | 4.715 |
| C | - | - | - | 6.472 | 5.773 | 4.317 | 6.528 | 5.214 | 6.039 | 5.136 | 6.152 | 7.664 | 5.877 | 5.32 |
| D | - | - | - | - | 6.139 | 8.605 | -5.436 | 7.605 | 5.933 | 8.599 | 6.451 | 10.116 | 6.242 | 7.974 |
| E | - | - | - | - | - | 7.499 | 5.71 | 6.017 | 3.352 | 7.583 | 0.378 | 9.212 | 0.102 | 6.41 |
| F | - | - | - | - | - | - | 6.702 | 5.734 | 6.236 | 4.219 | 6.406 | 7.268 | 6.075 | 6.01 |
| G | - | - | - | - | - | - | - | 7.533 | 5.807 | 8.519 | 6.367 | 10.029 | 6.122 | 7.894 |
| H | - | - | - | - | - | - | - | - | 5.519 | 6.585 | 5.564 | 8.453 | 5.18 | 1.699 |
| I | - | - | - | - | - | - | - | - | - | 7.79 | 3.295 | 9.378 | 3.541 | 6.734 |
| J | - | - | - | - | - | - | - | - | - | - | 6.604 | 6.697 | 6.246 | 6.421 |
| K | - | - | - | - | - | - | - | - | - | - | - | 9.134 | 1.231 | 6.426 |
| L | - | - | - | - | - | - | - | - | - | - | - | - | 6.53 | 6.981 |
| M | - | - | - | - | - | - | - | - | - | - | - | - | - | 6.381 |
| N | - | - | - | - | - | - | - | - | - | - | - | - | - | - |

| | |
|---|---|
| | Larger than 10 |
| | Between 8 and 9,99 |
| | Between 6 and 7,99 |
| | Between 4 and 5,99 |
| | Between 2 and 3,99 |
| | Between 0 and 1,99 |
| | NA (Calculation failed) |

| Median | A | B | C | D | E | F | G | H | I | J | K | L | M | N |
|---|---|---|---|---|---|---|---|---|---|---|---|---|---|---|
| A | - | 5.287 | 4.913 | 6.257 | 5.864 | 4.877 | 5.967 | 5.113 | 5.99 | 3.139 | 5.763 | 2.059 | 5.764 | 5.591 |
| B | - | - | 4.819 | 5.698 | 4.855 | 5.044 | 5.436 | 2.051 | 5.235 | 5.209 | 4.192 | 6.082 | 4.662 | -1.068 |
| C | - | - | - | 5.955 | 5.494 | 1.435 | 5.814 | 4.679 | 5.682 | 2.572 | 5.202 | 4.864 | 5.403 | 4.525 |
| D | - | - | - | - | 5.791 | 7.609 | 2.292 | 6.676 | 5.418 | 7.522 | 6.083 | 8.043 | 5.683 | 6.824 |
| E | - | - | - | - | - | 6.476 | 4.802 | 5.028 | 3.15 | 6.442 | 3.493 | 7.063 | -0.418 | 5.264 |
| F | - | - | - | - | - | - | 5.796 | 4.735 | 5.746 | 2.342 | 5.215 | 4.718 | 5.41 | 4.626 |
| G | - | - | - | - | - | - | - | 6.537 | 5.192 | 7.407 | 5.906 | 7.929 | 5.466 | 6.681 |
| H | - | - | - | - | - | - | - | - | 5.065 | 5.381 | 3.95 | 6.216 | 4.522 | 1.9 |
| I | - | - | - | - | - | - | - | - | - | 6.703 | 4.417 | 7.282 | 3.308 | 5.684 |
| J | - | - | - | - | - | - | - | - | - | - | 5.328 | 4.239 | 5.494 | 4.86 |
| K | - | - | - | - | - | - | - | - | - | - | - | 6.804 | 3.264 | 4.664 |
| L | - | - | - | - | - | - | - | - | - | - | - | - | 5.697 | 5.325 |
| M | - | - | - | - | - | - | - | - | - | - | - | - | - | 5.229 |
| N | - | - | - | - | - | - | - | - | - | - | - | - | - | - |

| Max | A | B | C | D | E | F | G | H | I | J | K | L | M | N |
|---|---|---|---|---|---|---|---|---|---|---|---|---|---|---|
| A | - | -0.338 | 5.032 | 3.613 | 3.816 | 3.714 | 4.162 | 3.727 | 2.835 | 2.193 | 5.596 | 2.303 | 3.894 | 4.244 |
| B | - | - | 5.05 | 2.917 | 3.876 | 3.759 | 4.036 | 3.85 | 2.952 | 2.423 | 5.592 | 2.128 | 3.908 | 4.219 |
| C | - | - | - | 4.378 | 3.744 | 3.819 | 3.659 | 3.805 | 4.253 | 4.367 | 3.543 | 4.782 | 3.734 | 3.392 |
| D | - | - | - | - | 2.507 | 2.467 | 3.433 | 3.09 | 0.738 | 0.827 | 5.322 | 3.586 | 3.099 | 3.458 |
| E | - | - | - | - | - | -0.662 | 0.753 | -3.506 | 2.562 | 3.06 | 4.858 | 4.009 | -2.503 | 1.97 |
| F | - | - | - | - | - | - | 1.712 | -1.792 | 2.353 | 2.86 | 4.918 | 3.94 | 0.416 | 2.241 |
| G | - | - | - | - | - | - | - | -0.629 | 2.811 | 3.265 | 4.809 | 4.146 | -0.615 | 1.32 |
| H | - | - | - | - | - | - | - | - | 2.522 | 2.959 | 4.885 | 3.975 | -1.232 | 2.033 |
| I | - | - | - | - | - | - | - | - | - | 1.255 | 5.236 | 3.411 | 2.774 | 3.411 |
| J | - | - | - | - | - | - | - | - | - | - | 5.342 | 3.138 | 3.212 | 3.686 |
| K | - | - | - | - | - | - | - | - | - | - | - | 5.13 | 4.485 | 4.288 |
| L | - | - | - | - | - | - | - | - | - | - | - | - | 4.296 | 4.571 |
| M | - | - | - | - | - | - | - | - | - | - | - | - | - | 1.833 |
| N | - | - | - | - | - | - | - | - | - | - | - | - | - | - |
