## Supplementary Data S2 for "Interlaboratory comparison of standardised metabolomics and lipidomics analyses in human and rodent blood using the MxP® Quant 500 kit": PC ae C38.2.pdf

**Amount of relative differences between laboratories**

| Min | A | B | C | D | E | F | G | H | I | J | K | L | M | N |
|---|---|---|---|---|---|---|---|---|---|---|---|---|---|---|
| A | - | 4.369 | 1.982 | 4.734 | 4.349 | 4.28 | 4.848 | 4.348 | 4.769 | 3.785 | 3.583 | 1.349 | 4.462 | 4.393 |
| B | - | - | 4.284 | 3.239 | 1.726 | -2.053 | 3.511 | 5.697 | 3.033 | 3.209 | 3.162 | 4.689 | 2.025 | 0.845 |
| C | - | - | - | 4.584 | 4.219 | 4.033 | 4.69 | 4.747 | 4.541 | 3.138 | 3.106 | 2.473 | 4.278 | 4.15 |
| D | - | - | - | - | 2.606 | 3.352 | 0.982 | 6.111 | 1.071 | 4.449 | 4.36 | 5.262 | 2.54 | 3.107 |
| E | - | - | - | - | - | 0.97 | 3.289 | 5.74 | 2.843 | 3.33 | 3.522 | 4.868 | 0.587 | -3.233 |
| F | - | - | - | - | - | - | 3.478 | 5.724 | 2.923 | 3.323 | 3.23 | 4.679 | 1.945 | 0.58 |
| G | - | - | - | - | - | - | - | 6.151 | 1.801 | 4.521 | 4.539 | 5.396 | 3.111 | 3.446 |
| H | - | - | - | - | - | - | - | - | 5.303 | 4.83 | 4.712 | 3.771 | 5.107 | 5.083 |
| I | - | - | - | - | - | - | - | - | - | 4.264 | 4.167 | 5.144 | 1.849 | 2.663 |
| J | - | - | - | - | - | - | - | - | - | - | -0.564 | 4.032 | 3.49 | 3.248 |
| K | - | - | - | - | - | - | - | - | - | - | - | 3.936 | 3.575 | 3.327 |
| L | - | - | - | - | - | - | - | - | - | - | - | - | 4.617 | 4.622 |
| M | - | - | - | - | - | - | - | - | - | - | - | - | - | 1.226 |
| N | - | - | - | - | - | - | - | - | - | - | - | - | - | - |

| Median | A | B | C | D | E | F | G | H | I | J | K | L | M | N |
|---|---|---|---|---|---|---|---|---|---|---|---|---|---|---|
| A | - | 3.921 | -1.942 | 4.494 | 4.266 | 3.773 | 4.659 | 4.072 | 4.263 | 3.138 | 2.822 | 0.777 | 4.171 | 4.046 |
| B | - | - | 4.187 | 3.009 | 2.46 | 0.063 | 3.604 | 5.241 | 2.438 | 2.586 | 3.034 | 4.078 | 1.826 | 1.161 |
| C | - | - | - | 4.476 | 4.314 | 3.844 | 4.679 | 4.055 | 4.309 | 3.326 | 3.032 | 0.383 | 4.192 | 4.101 |
| D | - | - | - | - | 1.111 | 3.204 | 2.297 | 5.63 | 1.141 | 3.838 | 4.069 | 4.759 | 2.175 | 2.503 |
| E | - | - | - | - | - | 2.788 | 2.876 | 5.495 | -3.337 | 3.657 | 3.898 | 4.485 | 0.837 | 1.762 |
| F | - | - | - | - | - | - | 3.701 | 5.196 | 2.677 | 2.236 | 2.815 | 3.996 | 2.121 | 1.699 |
| G | - | - | - | - | - | - | - | 5.828 | 2.959 | 4.384 | 4.551 | 5.032 | 3.31 | 3.509 |
| H | - | - | - | - | - | - | - | - | 4.934 | 4.42 | 4.322 | 4.011 | 4.893 | 4.823 |
| I | - | - | - | - | - | - | - | - | - | 3.557 | 3.831 | 4.602 | 1.182 | 1.754 |
| J | - | - | - | - | - | - | - | - | - | - | 1.061 | 3.221 | 3.225 | 3.001 |
| K | - | - | - | - | - | - | - | - | - | - | - | 2.922 | 3.456 | 3.289 |
| L | - | - | - | - | - | - | - | - | - | - | - | - | 3.956 | 3.749 |
| M | - | - | - | - | - | - | - | - | - | - | - | - | - | 0.279 |
| N | - | - | - | - | - | - | - | - | - | - | - | - | - | - |

| Max | A | B | C | D | E | F | G | H | I | J | K | L | M | N |
|---|---|---|---|---|---|---|---|---|---|---|---|---|---|---|
| A | - | 2.373 | 4.218 | 2.877 | 3.711 | 3.618 | 3.449 | 2.996 | 3.328 | 1.685 | 3.419 | 4.483 | 2.119 | -0.035 |
| B | - | - | 3.665 | 1.089 | 4.112 | 2.773 | 3.929 | 1.463 | 2.211 | 0.866 | 2.446 | 4.715 | -0.63 | 1.98 |
| C | - | - | - | 3.252 | 4.739 | 2.407 | 4.635 | 3.155 | 2.862 | 3.698 | 2.724 | 5.11 | 3.59 | 3.892 |
| D | - | - | - | - | 4.246 | 2.238 | 4.076 | -0.683 | 1.37 | 1.957 | 1.694 | 4.792 | 1.415 | 2.539 |
| E | - | - | - | - | - | 4.868 | 1.36 | 4.601 | 4.728 | 4.232 | 4.771 | 3.414 | 4.321 | 4.017 |
| F | - | - | - | - | - | - | 4.365 | 1.931 | 1.052 | 3.012 | 0.536 | 4.947 | 2.832 | 3.332 |
| G | - | - | - | - | - | - | - | 4.407 | 4.547 | 3.986 | 4.596 | 3.685 | 4.084 | 3.73 |
| H | - | - | - | - | - | - | - | - | 0.873 | 2.154 | 1.301 | 4.812 | 1.765 | 2.7 |
| I | - | - | - | - | - | - | - | - | - | 2.632 | -0.516 | 4.88 | 2.355 | 3.028 |
| J | - | - | - | - | - | - | - | - | - | - | 2.845 | 4.632 | 0.101 | 1.131 |
| K | - | - | - | - | - | - | - | - | - | - | - | 4.9 | 2.523 | 3.131 |
| L | - | - | - | - | - | - | - | - | - | - | - | - | 5.104 | 4.911 |
| M | - | - | - | - | - | - | - | - | - | - | - | - | - | 1.665 |
| N | - | - | - | - | - | - | - | - | - | - | - | - | - | - |

| | |
|---|---|
| | Larger than 10 |
| | Between 8 and 9,99 |
| | Between 6 and 7,99 |
| | Between 4 and 5,99 |
| | Between 2 and 3,99 |
| | Between 0 and 1,99 |
| | NA (Calculation failed) |
