## Supplementary Data S2 for "Interlaboratory comparison of standardised metabolomics and lipidomics analyses in human and rodent blood using the MxP® Quant 500 kit": PC ae C38.3.pdf

**Amount of relative differences between laboratories**

| Min | A | B | C | D | E | F | G | H | I | J | K | L | M | N |
|---|---|---|---|---|---|---|---|---|---|---|---|---|---|---|
| A | - | 3.1 | 0.765 | 3.093 | 4.745 | 2.819 | 3.873 | 2.509 | 3.345 | 4.128 | 3.663 | 3.737 | 3.41 | 3.218 |
| B | - | - | 2.804 | 1.306 | 4.078 | -0.898 | 0.464 | 0.996 | -0.09 | 2.201 | 0.291 | 2.263 | 0.455 | 0.413 |
| C | - | - | - | 2.567 | 4.612 | 2.6 | 3.411 | 2.209 | 3.042 | 3.631 | 3.176 | 3.573 | 3.139 | 2.754 |
| D | - | - | - | - | 4.398 | 1.736 | 2.694 | 1.954 | -0.298 | 3.144 | 1.293 | 2.315 | 0.721 | -3.419 |
| E | - | - | - | - | - | 4.499 | 4.227 | 4.569 | 4.373 | 4.049 | 4.301 | 4.008 | 4.326 | 4.527 |
| F | - | - | - | - | - | - | 2.424 | 0.579 | 1.189 | 2.861 | 1.807 | 2.626 | 1.551 | -0.646 |
| G | - | - | - | - | - | - | - | 3.545 | 2.695 | 0.391 | 2.286 | 1.192 | 2.514 | 2.908 |
| H | - | - | - | - | - | - | - | - | 1.82 | 2.96 | 2.073 | 2.887 | 2.022 | 0.857 |
| I | - | - | - | - | - | - | - | - | - | 2.218 | 0.079 | 2.057 | -0.423 | 0.525 |
| J | - | - | - | - | - | - | - | - | - | - | 1.185 | 0.241 | 1.212 | 2.341 |
| K | - | - | - | - | - | - | - | - | - | - | - | 1.657 | -3.28 | 1.43 |
| L | - | - | - | - | - | - | - | - | - | - | - | - | 1.78 | 2.532 |
| M | - | - | - | - | - | - | - | - | - | - | - | - | - | 1.21 |
| N | - | - | - | - | - | - | - | - | - | - | - | - | - | - |

| Median | A | B | C | D | E | F | G | H | I | J | K | L | M | N |
|---|---|---|---|---|---|---|---|---|---|---|---|---|---|---|
| A | - | 1.663 | -2.124 | 3.627 | 4.43 | 2.566 | 4.277 | 1.978 | 3.186 | 0.301 | 2.262 | 3.561 | 3.079 | 3.103 |
| B | - | - | 2.018 | 3.114 | 4.23 | 0.967 | 3.997 | -5.511 | 2.472 | 0.745 | -2.768 | 3.146 | 2.243 | 2.217 |
| C | - | - | - | 3.605 | 4.431 | 2.533 | 4.261 | 1.936 | 3.171 | 0.633 | 2.089 | 3.568 | 3.057 | 3.064 |
| D | - | - | - | - | 3.347 | 2.862 | 2.985 | 3.249 | 1.958 | 3.733 | 3.111 | 0.175 | 2.177 | 2.108 |
| E | - | - | - | - | - | 4.36 | 1.723 | 4.515 | 4.019 | 4.657 | 4.501 | 3.614 | 4.102 | 4.106 |
| F | - | - | - | - | - | - | 3.834 | 1.047 | 1.785 | 2.156 | 0.656 | 2.702 | 1.44 | 1.455 |
| G | - | - | - | - | - | - | - | 4.239 | 3.684 | 4.534 | 4.163 | 3.316 | 3.757 | 3.729 |
| H | - | - | - | - | - | - | - | - | 2.441 | 1.124 | -1.496 | 3.086 | 2.228 | 2.232 |
| I | - | - | - | - | - | - | - | - | - | 3.054 | 2.354 | 1.611 | -0.462 | -0.611 |
| J | - | - | - | - | - | - | - | - | - | - | -0.474 | 3.35 | 2.483 | 2.437 |
| K | - | - | - | - | - | - | - | - | - | - | - | 3.162 | 2.22 | 2.189 |
| L | - | - | - | - | - | - | - | - | - | - | - | - | 2.006 | 1.988 |
| M | - | - | - | - | - | - | - | - | - | - | - | - | - | -5.083 |
| N | - | - | - | - | - | - | - | - | - | - | - | - | - | - |

| Max | A | B | C | D | E | F | G | H | I | J | K | L | M | N |
|---|---|---|---|---|---|---|---|---|---|---|---|---|---|---|
| A | - | 0.903 | 0.912 | 3.927 | 4.22 | 2.027 | 4.469 | 0.524 | 3.019 | 3.076 | 2.072 | 3.408 | 2.632 | 2.986 |
| B | - | - | -2.585 | 4.086 | 4.308 | 2.585 | 4.613 | 1.537 | 3.304 | 2.435 | 0.118 | 3.583 | 3.035 | 3.358 |
| C | - | - | - | 4.074 | 4.319 | 2.418 | 4.594 | 1.427 | 3.283 | 2.291 | 0.281 | 3.565 | 2.967 | 3.304 |
| D | - | - | - | - | 2.042 | 3.773 | 3.132 | 4.134 | 3.052 | 4.726 | 4.414 | 2.548 | 3.444 | 3.047 |
| E | - | - | - | - | - | 4.1 | 2.506 | 4.434 | 3.582 | 4.925 | 4.7 | 3.214 | 3.828 | 3.556 |
| F | - | - | - | - | - | - | 4.281 | 1.528 | 2.183 | 3.515 | 2.884 | 2.757 | 1.315 | 2.236 |
| G | - | - | - | - | - | - | - | 4.868 | 4.235 | 5.238 | 5.037 | 3.99 | 4.431 | 4.222 |
| H | - | - | - | - | - | - | - | - | 2.85 | 3.019 | 2.019 | 3.223 | 2.413 | 2.887 |
| I | - | - | - | - | - | - | - | - | - | 4.001 | 3.605 | 1.137 | 1.165 | -3.64 |
| J | - | - | - | - | - | - | - | - | - | - | 2.051 | 4.102 | 3.78 | 3.93 |
| K | - | - | - | - | - | - | - | - | - | - | - | 3.747 | 3.258 | 3.506 |
| L | - | - | - | - | - | - | - | - | - | - | - | - | 2.206 | 1.181 |
| M | - | - | - | - | - | - | - | - | - | - | - | - | - | 1.154 |
| N | - | - | - | - | - | - | - | - | - | - | - | - | - | - |

| | |
|---|---|
| | Larger than 10 |
| | Between 8 and 9,99 |
| | Between 6 and 7,99 |
| | Between 4 and 5,99 |
| | Between 2 and 3,99 |
| | Between 0 and 1,99 |
| | NA (Calculation failed) |
