## Supplementary Data S2 for "Interlaboratory comparison of standardised metabolomics and lipidomics analyses in human and rodent blood using the MxP® Quant 500 kit": PC ae C38.4.pdf

**Amount of relative differences between laboratories**

| Min | A | B | C | D | E | F | G | H | I | J | K | L | M | N |
|---|---|---|---|---|---|---|---|---|---|---|---|---|---|---|
| A | - | 3.69 | -4.49 | 0.226 | 4.58 | 3.885 | 1.368 | 3.669 | 2.854 | 4.522 | 3.445 | 3.386 | 2.045 | 2.624 |
| B | - | - | 3.659 | 3.908 | 3.706 | 1.667 | 3.969 | 0.126 | 2.243 | 2.734 | 1.35 | 0.412 | 3.008 | 2.714 |
| C | - | - | - | 0.606 | 4.54 | 3.88 | 1.415 | 3.665 | 2.788 | 3.942 | 3.194 | 3.401 | 2.153 | 2.618 |
| D | - | - | - | - | 4.627 | 3.971 | 0.577 | 3.815 | 2.985 | 4.401 | 3.454 | 3.563 | 2.391 | 2.91 |
| E | - | - | - | - | - | 3.513 | 5.017 | 3.735 | 4.391 | 3.221 | 4.151 | 4.032 | 4.609 | 4.462 |
| F | - | - | - | - | - | - | 4.286 | 1.085 | 3.084 | 1.67 | 2.51 | 2.223 | 3.551 | 3.383 |
| G | - | - | - | - | - | - | - | 3.907 | 3.223 | 4.399 | 3.564 | 3.682 | 2.751 | 3.095 |
| H | - | - | - | - | - | - | - | - | 2.649 | 2.191 | 1.943 | 1.334 | 3.226 | 2.936 |
| I | - | - | - | - | - | - | - | - | - | 3.378 | 1.28 | 1.955 | 1.564 | -0.334 |
| J | - | - | - | - | - | - | - | - | - | - | 2.735 | 2.875 | 3.817 | 3.435 |
| K | - | - | - | - | - | - | - | - | - | - | - | 0.352 | 2.397 | 1.788 |
| L | - | - | - | - | - | - | - | - | - | - | - | - | 2.739 | 2.359 |
| M | - | - | - | - | - | - | - | - | - | - | - | - | - | 0.833 |
| N | - | - | - | - | - | - | - | - | - | - | - | - | - | - |

| Median | A | B | C | D | E | F | G | H | I | J | K | L | M | N |
|---|---|---|---|---|---|---|---|---|---|---|---|---|---|---|
| A | - | 1.797 | 1.13 | 0.599 | 3.926 | 2.519 | 1.895 | 2.845 | 2.507 | 2.26 | 1.389 | 1.069 | 1.47 | 1.865 |
| B | - | - | 2.548 | 0.993 | 3.61 | 1.299 | 2.894 | 1.953 | 1.228 | 0.66 | 2.625 | 0.471 | -0.357 | -1.863 |
| C | - | - | - | 1.86 | 4.091 | 2.958 | 0.583 | 3.197 | 2.949 | 2.819 | -1.511 | 2.068 | 2.272 | 2.516 |
| D | - | - | - | - | 3.797 | 2.092 | 2.409 | 2.524 | 2.083 | 1.769 | 2.064 | -0.787 | 0.374 | 1.095 |
| E | - | - | - | - | - | 3.473 | 4.486 | 3.25 | 3.489 | 3.591 | 4.388 | 3.954 | 3.889 | 3.767 |
| F | - | - | - | - | - | - | 3.344 | 0.525 | -3.11 | -0.159 | 3.146 | 1.982 | 1.738 | 1.158 |
| G | - | - | - | - | - | - | - | 3.394 | 3.182 | 3.056 | 0.152 | 2.487 | 2.643 | 2.829 |
| H | - | - | - | - | - | - | - | - | 0.67 | 1.239 | 3.387 | 2.452 | 2.267 | 1.913 |
| I | - | - | - | - | - | - | - | - | - | -0.403 | 3.129 | 1.933 | 1.64 | 1.119 |
| J | - | - | - | - | - | - | - | - | - | - | 2.992 | 1.61 | 1.314 | 0.49 |
| K | - | - | - | - | - | - | - | - | - | - | - | 2.164 | 2.328 | 2.598 |
| L | - | - | - | - | - | - | - | - | - | - | - | - | -0.744 | 0.711 |
| M | - | - | - | - | - | - | - | - | - | - | - | - | - | -0.107 |
| N | - | - | - | - | - | - | - | - | - | - | - | - | - | - |

| Max | A | B | C | D | E | F | G | H | I | J | K | L | M | N |
|---|---|---|---|---|---|---|---|---|---|---|---|---|---|---|
| A | - | 1.234 | 1.303 | 0.861 | 3.839 | 2.226 | 1.956 | 2.704 | 2.461 | 1.842 | 2.021 | -0.027 | 1.371 | 1.747 |
| B | - | - | 2.322 | -0.927 | 3.6 | 1.249 | 2.648 | 2.073 | 1.665 | 0.315 | 2.712 | 0.479 | -2.26 | 0.067 |
| C | - | - | - | 2.074 | 4.036 | 2.799 | 0.413 | 3.129 | 2.966 | 2.702 | 0.448 | 1.741 | 2.287 | 2.504 |
| D | - | - | - | - | 3.677 | 1.552 | 2.549 | 2.247 | 1.928 | 1.159 | 2.535 | -0.118 | -0.401 | 0.611 |
| E | - | - | - | - | - | 3.468 | 4.391 | 3.179 | 3.334 | 3.613 | 4.411 | 3.943 | 3.756 | 3.662 |
| F | - | - | - | - | - | - | 3.142 | 0.902 | -0.213 | 0.221 | 3.2 | 1.946 | 1.133 | 0.383 |
| G | - | - | - | - | - | - | - | 3.317 | 3.177 | 2.901 | -4.993 | 2.214 | 2.627 | 2.795 |
| H | - | - | - | - | - | - | - | - | 0.062 | 1.495 | 3.481 | 2.553 | 2.069 | 1.735 |
| I | - | - | - | - | - | - | - | - | - | 0.58 | 3.313 | 2.277 | 1.65 | 1.219 |
| J | - | - | - | - | - | - | - | - | - | - | 3.017 | 1.311 | -0.145 | -1.553 |
| K | - | - | - | - | - | - | - | - | - | - | - | 2.296 | 2.683 | 2.834 |
| L | - | - | - | - | - | - | - | - | - | - | - | - | 0.667 | 1.264 |
| M | - | - | - | - | - | - | - | - | - | - | - | - | - | -0.266 |
| N | - | - | - | - | - | - | - | - | - | - | - | - | - | - |

- Larger than 10
- Between 8 and 9,99
- Between 6 and 7,99
- Between 4 and 5,99
- Between 2 and 3,99
- Between 0 and 1,99
- NA (Calculation failed)
