## Supplementary Data S2 for "Interlaboratory comparison of standardised metabolomics and lipidomics analyses in human and rodent blood using the MxP® Quant 500 kit": PC ae C38.5.pdf

**Amount of relative differences between laboratories**

| Min | A | B | C | D | E | F | G | H | I | J | K | L | M | N |
|---|---|---|---|---|---|---|---|---|---|---|---|---|---|---|
| A | - | 3.943 | 2.226 | -2.587 | 4.737 | 3.505 | 2.674 | 2.746 | 4.063 | 4.413 | 2.43 | 3.511 | -1.686 | 2.529 |
| B | - | - | 4.364 | 4.033 | 3.94 | 1.436 | 4.522 | 2.817 | 1.752 | 0.268 | 3.474 | 1.258 | 3.968 | 3.052 |
| C | - | - | - | 2.136 | 4.887 | 3.966 | 1.139 | 3.552 | 4.354 | 4.218 | 2.899 | 4.012 | 2.234 | 3.468 |
| D | - | - | - | - | 4.748 | 3.556 | 2.719 | 3.05 | 4.098 | 4.277 | 1.997 | 3.619 | -3.545 | 2.814 |
| E | - | - | - | - | - | 4.34 | 5.396 | 4.586 | 3.772 | 3.841 | 4.838 | 4.223 | 5.154 | 4.695 |
| F | - | - | - | - | - | - | 4.319 | 2.203 | 2.582 | 2.819 | 2.903 | -0.815 | 3.692 | 2.683 |
| G | - | - | - | - | - | - | - | 3.799 | 4.529 | 4.627 | 3.337 | 4.146 | 2.668 | 3.664 |
| H | - | - | - | - | - | - | - | - | 3.316 | 3.034 | 2.166 | 2.272 | 2.981 | -0.369 |
| I | - | - | - | - | - | - | - | - | - | 1.813 | 3.742 | 2.029 | 4.212 | 3.407 |
| J | - | - | - | - | - | - | - | - | - | - | 3.475 | 2.536 | 4.227 | 3.294 |
| K | - | - | - | - | - | - | - | - | - | - | - | 2.998 | 2.219 | 1.442 |
| L | - | - | - | - | - | - | - | - | - | - | - | - | 3.741 | 2.671 |
| M | - | - | - | - | - | - | - | - | - | - | - | - | - | 2.674 |
| N | - | - | - | - | - | - | - | - | - | - | - | - | - | - |

| | |
|---|---|
| | Larger than 10 |
| | Between 8 and 9,99 |
| | Between 6 and 7,99 |
| | Between 4 and 5,99 |
| | Between 2 and 3,99 |
| | Between 0 and 1,99 |
| | NA (Calculation failed) |

| Median | A | B | C | D | E | F | G | H | I | J | K | L | M | N |
|---|---|---|---|---|---|---|---|---|---|---|---|---|---|---|
| A | - | 3.21 | -3.76 | 1.936 | 4.182 | 2.673 | 1.3 | 3.301 | 2.657 | 2.659 | 0.005 | 1.732 | 1.534 | 2.149 |
| B | - | - | 3.366 | 2.594 | 3.288 | 1.677 | 3.696 | -0.648 | 1.72 | 1.607 | 3.186 | 2.717 | 2.814 | 2.421 |
| C | - | - | - | 1.978 | 4.189 | 2.689 | 1.235 | 3.305 | 2.678 | 2.713 | 0.186 | 1.76 | 1.561 | 2.167 |
| D | - | - | - | - | 3.895 | 1.387 | 2.713 | 2.628 | 1.354 | 1.382 | 1.551 | -0.804 | 0.022 | -0.793 |
| E | - | - | - | - | - | 3.845 | 4.658 | 3.339 | 3.857 | 3.829 | 4.389 | 4.181 | 4.217 | 4.069 |
| F | - | - | - | - | - | - | 3.242 | 1.899 | -3.375 | -4.106 | 2.52 | 1.713 | 1.89 | 1.051 |
| G | - | - | - | - | - | - | - | 3.579 | 3.093 | 3.102 | 1.765 | 2.489 | 2.376 | 2.743 |
| H | - | - | - | - | - | - | - | - | 1.96 | 1.89 | 3.282 | 2.852 | 2.946 | 2.584 |
| I | - | - | - | - | - | - | - | - | - | -1.612 | 2.495 | 1.676 | 1.846 | 1.003 |
| J | - | - | - | - | - | - | - | - | - | - | 2.567 | 1.763 | 1.937 | 1.174 |
| K | - | - | - | - | - | - | - | - | - | - | - | 1.208 | 0.927 | 1.776 |
| L | - | - | - | - | - | - | - | - | - | - | - | - | -1.308 | 0.21 |
| M | - | - | - | - | - | - | - | - | - | - | - | - | - | 0.668 |
| N | - | - | - | - | - | - | - | - | - | - | - | - | - | - |

| Max | A | B | C | D | E | F | G | H | I | J | K | L | M | N |
|---|---|---|---|---|---|---|---|---|---|---|---|---|---|---|
| A | - | 3.177 | -3.356 | 1.997 | 4.158 | 2.627 | 1.177 | 3.315 | 2.564 | 2.583 | -0.283 | 1.564 | 1.589 | 2.132 |
| B | - | - | 3.312 | 2.478 | 3.259 | 1.686 | 3.638 | -0.209 | 1.826 | 1.65 | 3.173 | 2.758 | 2.751 | 2.39 |
| C | - | - | - | 1.97 | 4.157 | 2.604 | 1.24 | 3.297 | 2.553 | 2.655 | -0.191 | 1.49 | 1.53 | 2.08 |
| D | - | - | - | - | 3.854 | 1.167 | 2.712 | 2.613 | 1.005 | 1.165 | 1.685 | 0.207 | 0.067 | -1.647 |
| E | - | - | - | - | - | 3.821 | 4.608 | 3.279 | 3.86 | 3.829 | 4.367 | 4.179 | 4.17 | 4.037 |
| F | - | - | - | - | - | - | 3.157 | 1.991 | -1.619 | -3.413 | 2.502 | 1.788 | 1.759 | 0.924 |
| G | - | - | - | - | - | - | - | 3.572 | 2.997 | 3.042 | 1.642 | 2.341 | 2.365 | 2.691 |
| H | - | - | - | - | - | - | - | - | 2.141 | 1.988 | 3.314 | 2.957 | 2.944 | 2.634 |
| I | - | - | - | - | - | - | - | - | - | -1.284 | 2.411 | 1.658 | 1.62 | 0.731 |
| J | - | - | - | - | - | - | - | - | - | - | 2.514 | 1.716 | 1.726 | 0.966 |
| K | - | - | - | - | - | - | - | - | - | - | - | 1.038 | 1.091 | 1.788 |
| L | - | - | - | - | - | - | - | - | - | - | - | - | -4.318 | 0.543 |
| M | - | - | - | - | - | - | - | - | - | - | - | - | - | 0.483 |
| N | - | - | - | - | - | - | - | - | - | - | - | - | - | - |
