## Supplementary Data S2 for "Interlaboratory comparison of standardised metabolomics and lipidomics analyses in human and rodent blood using the MxP® Quant 500 kit": PC ae C38.6.pdf

**Amount of relative differences between laboratories**

| Min | A | B | C | D | E | F | G | H | I | J | K | L | M | N |
|---|---|---|---|---|---|---|---|---|---|---|---|---|---|---|
| A | - | 2.6 | 3.578 | 2.139 | 4.193 | 3.243 | 1.552 | 1.834 | 4.382 | 3.754 | 4.198 | 3.194 | 2.492 | -0.613 |
| B | - | - | 4.254 | 0.781 | 3.761 | 1.781 | 3.276 | 1.009 | 3.956 | 4.518 | 4.737 | 1.663 | 3.607 | 2.358 |
| C | - | - | - | 4.053 | 4.739 | 4.245 | 3.076 | 3.865 | 4.937 | 2.155 | 3.136 | 4.349 | 2.554 | 3.658 |
| D | - | - | - | - | 3.849 | 2.256 | 2.961 | -1.355 | 4.011 | 4.379 | 4.635 | 2.312 | 3.411 | 1.773 |
| E | - | - | - | - | - | 3.39 | 4.673 | 3.966 | 1.947 | 5.309 | 5.437 | 3.332 | 4.886 | 4.289 |
| F | - | - | - | - | - | - | 3.764 | 2.527 | 3.881 | 4.717 | 4.93 | -5.736 | 4.061 | 3.193 |
| G | - | - | - | - | - | - | - | 2.845 | 4.523 | 3.438 | 3.871 | 3.558 | 1.287 | 2.056 |
| H | - | - | - | - | - | - | - | - | 4.18 | 4.372 | 4.646 | 2.525 | 3.348 | 1.46 |
| I | - | - | - | - | - | - | - | - | - | 5.396 | 5.387 | 3.476 | 4.828 | 4.364 |
| J | - | - | - | - | - | - | - | - | - | - | 2.826 | 4.553 | 2.95 | 4.068 |
| K | - | - | - | - | - | - | - | - | - | - | - | 4.886 | 3.835 | 4.481 |
| L | - | - | - | - | - | - | - | - | - | - | - | - | 4.001 | 3.127 |
| M | - | - | - | - | - | - | - | - | - | - | - | - | - | 2.815 |
| N | - | - | - | - | - | - | - | - | - | - | - | - | - | - |

| | |
|---|---|
| (red) | Larger than 10 |
| (orange) | Between 8 and 9,99 |
| (light orange) | Between 6 and 7,99 |
| (light green) | Between 4 and 5,99 |
| (green) | Between 2 and 3,99 |
| (dark green) | Between 0 and 1,99 |
| (grey) | NA (Calculation failed) |

| Median | A | B | C | D | E | F | G | H | I | J | K | L | M | N |
|---|---|---|---|---|---|---|---|---|---|---|---|---|---|---|
| A | - | 2.678 | -2.54 | 2.542 | 4.151 | 3.174 | 2.085 | 2.954 | 2.847 | 2.849 | -1.109 | 1.968 | 2.082 | 1.428 |
| B | - | - | 2.812 | -0.71 | 3.602 | 1.487 | 3.507 | 0.516 | -0.228 | -0.204 | 2.664 | 1.409 | 1.212 | 1.988 |
| C | - | - | - | 2.599 | 4.165 | 3.196 | 1.997 | 2.97 | 2.885 | 2.892 | -0.556 | 2.035 | 2.13 | 1.507 |
| D | - | - | - | - | 3.664 | 1.76 | 3.418 | 1.012 | 0.542 | 0.554 | 2.51 | 1.026 | 0.767 | 1.741 |
| E | - | - | - | - | - | 3.409 | 4.742 | 3.609 | 3.684 | 3.682 | 4.395 | 4.074 | 4.039 | 4.197 |
| F | - | - | - | - | - | - | 3.866 | 0.494 | 1.014 | 0.999 | 3.234 | 2.491 | 2.396 | 2.8 |
| G | - | - | - | - | - | - | - | 3.524 | 3.456 | 3.458 | 2.175 | 2.969 | 3.025 | 2.734 |
| H | - | - | - | - | - | - | - | - | -0.852 | -0.913 | 2.971 | 2.041 | 1.92 | 2.448 |
| I | - | - | - | - | - | - | - | - | - | -6.231 | 2.862 | 1.821 | 1.647 | 2.267 |
| J | - | - | - | - | - | - | - | - | - | - | 2.867 | 1.854 | 1.742 | 2.325 |
| K | - | - | - | - | - | - | - | - | - | - | - | 1.776 | 1.879 | 1.089 |
| L | - | - | - | - | - | - | - | - | - | - | - | - | -1.483 | 0.331 |
| M | - | - | - | - | - | - | - | - | - | - | - | - | - | 0.684 |
| N | - | - | - | - | - | - | - | - | - | - | - | - | - | - |

| Max | A | B | C | D | E | F | G | H | I | J | K | L | M | N |
|---|---|---|---|---|---|---|---|---|---|---|---|---|---|---|
| A | - | 2.683 | -0.709 | 2.565 | 4.148 | 3.169 | 2.118 | 3.004 | 2.647 | 3.101 | 1.061 | 1.815 | 2.278 | 1.502 |
| B | - | - | 2.637 | -0.894 | 3.589 | 1.466 | 3.523 | 0.728 | -2.77 | 1.292 | 2.197 | 1.629 | 0.765 | 1.958 |
| C | - | - | - | 2.422 | 4.111 | 3.09 | 2.306 | 2.886 | 2.537 | 3.038 | 0.582 | 1.521 | 2.099 | 1.111 |
| D | - | - | - | - | 3.649 | 1.721 | 3.448 | 1.121 | -1.134 | 1.56 | 2.016 | 1.359 | 0.193 | 1.739 |
| E | - | - | - | - | - | 3.411 | 4.747 | 3.582 | 3.806 | 3.466 | 4.257 | 4.117 | 3.966 | 4.19 |
| F | - | - | - | - | - | - | 3.874 | 0.166 | 1.661 | -1.382 | 2.925 | 2.595 | 2.185 | 2.768 |
| G | - | - | - | - | - | - | - | 3.559 | 3.344 | 3.636 | 2.625 | 2.913 | 3.138 | 2.771 |
| H | - | - | - | - | - | - | - | - | 0.871 | -0.306 | 2.669 | 2.281 | 1.759 | 2.497 |
| I | - | - | - | - | - | - | - | - | - | 1.477 | 2.101 | 1.566 | 0.631 | 1.892 |
| J | - | - | - | - | - | - | - | - | - | - | 2.87 | 2.5 | 2.014 | 2.707 |
| K | - | - | - | - | - | - | - | - | - | - | - | 0.539 | 1.592 | -0.475 |
| L | - | - | - | - | - | - | - | - | - | - | - | - | 0.467 | -0.442 |
| M | - | - | - | - | - | - | - | - | - | - | - | - | - | 1.144 |
| N | - | - | - | - | - | - | - | - | - | - | - | - | - | - |
