## Supplementary Data S2 for "Interlaboratory comparison of standardised metabolomics and lipidomics analyses in human and rodent blood using the MxP® Quant 500 kit": PC ae C40.1.pdf

**Amount of relative differences between laboratories**

| Min | A | B | C | D | E | F | G | H | I | J | K | L | M | N |
|---|---|---|---|---|---|---|---|---|---|---|---|---|---|---|
| A | - | 2.659 | 3.553 | 2.916 | 2.667 | 3.053 | 1.734 | 1.25 | -1.634 | 0.027 | 3.117 | 3.056 | 1.886 | 1.384 |
| B | - | - | 3.974 | 1.923 | 1.455 | 3.721 | -5.32 | 2.54 | 3.26 | 2.383 | 3.608 | 3.51 | 0.228 | -3.384 |
| C | - | - | - | 4.084 | 4.011 | 1.494 | 3.772 | 3.1 | 2.852 | 3.215 | 1.551 | 1.777 | 3.71 | 3.725 |
| D | - | - | - | - | -1.369 | 4.094 | 1.696 | 3.203 | 3.786 | 3.148 | 3.984 | 3.883 | 2.422 | 1.686 |
| E | - | - | - | - | - | 3.956 | 1.677 | 3.216 | 1.767 | 2.993 | 4.006 | 3.974 | 1.447 | 1.983 |
| F | - | - | - | - | - | - | 3.436 | 2.471 | 3.017 | 2.778 | -2.134 | -3.387 | 3.467 | 3.356 |
| G | - | - | - | - | - | - | - | 2.533 | 2.714 | 2.306 | 3.621 | 3.539 | -0.114 | -1.566 |
| H | - | - | - | - | - | - | - | - | 2.312 | 0.803 | 2.629 | 2.604 | 2.833 | 2.211 |
| I | - | - | - | - | - | - | - | - | - | 2.141 | -1.635 | -1.436 | 2.885 | 3.475 |
| J | - | - | - | - | - | - | - | - | - | - | 2.801 | 2.688 | 1.994 | 2.124 |
| K | - | - | - | - | - | - | - | - | - | - | - | -1.389 | 3.485 | 3.384 |
| L | - | - | - | - | - | - | - | - | - | - | - | - | 3.802 | 3.333 |
| M | - | - | - | - | - | - | - | - | - | - | - | - | - | 0.277 |
| N | - | - | - | - | - | - | - | - | - | - | - | - | - | - |

| Median | A | B | C | D | E | F | G | H | I | J | K | L | M | N |
|---|---|---|---|---|---|---|---|---|---|---|---|---|---|---|
| A | - | 0.037 | 3.918 | -4.046 | 2.71 | 3.15 | -6.417 | -1.289 | 4.436 | 2.363 | 3.236 | 1.448 | 1.468 | -1.392 |
| B | - | - | 3.729 | 0.363 | 2.955 | 2.986 | 0.258 | 0.581 | 4.422 | 1.757 | 2.876 | 0.587 | 0.624 | -0.297 |
| C | - | - | - | 3.692 | 4.224 | 2.354 | 3.696 | 3.729 | 2.74 | 3.121 | 2.353 | 3.397 | 3.394 | 3.654 |
| D | - | - | - | - | 2.754 | 3.215 | -6.129 | -1.983 | 4.508 | 2.132 | 3.057 | 1.458 | 1.514 | -1.114 |
| E | - | - | - | - | - | 4.04 | 2.806 | 2.708 | 4.866 | 3.65 | 4.097 | 3.31 | 3.313 | 2.881 |
| F | - | - | - | - | - | - | 3.03 | 3.094 | 3.536 | 1.822 | -1.308 | 2.5 | 2.498 | 2.97 |
| G | - | - | - | - | - | - | - | -1.464 | 4.466 | 2.241 | 3.146 | 1.457 | 1.49 | -1.294 |
| H | - | - | - | - | - | - | - | - | 4.462 | 2.58 | 3.38 | 1.673 | 1.672 | -0.305 |
| I | - | - | - | - | - | - | - | - | - | 3.905 | 3.603 | 3.96 | 3.992 | 4.147 |
| J | - | - | - | - | - | - | - | - | - | - | 1.971 | 0.926 | 0.921 | 2.053 |
| K | - | - | - | - | - | - | - | - | - | - | - | 2.592 | 2.586 | 3.031 |
| L | - | - | - | - | - | - | - | - | - | - | - | - | -4.873 | 1.214 |
| M | - | - | - | - | - | - | - | - | - | - | - | - | - | 1.219 |
| N | - | - | - | - | - | - | - | - | - | - | - | - | - | - |

| Max | A | B | C | D | E | F | G | H | I | J | K | L | M | N |
|---|---|---|---|---|---|---|---|---|---|---|---|---|---|---|
| A | - | 3.44 | 4.374 | 3.857 | 2.78 | 3.298 | 2.761 | 2.438 | 5.884 | 3.534 | 3.403 | 2.901 | 3.698 | 2.9 |
| B | - | - | 3.199 | 1.793 | 4.008 | -0.204 | 1.835 | 3.871 | 5.38 | -0.974 | -3.255 | 4.049 | 1.087 | 1.569 |
| C | - | - | - | 2.393 | 4.521 | 3.161 | 3.528 | 4.438 | 4.924 | 2.959 | 3.08 | 4.554 | 2.687 | 3.468 |
| D | - | - | - | - | 4.217 | 1.897 | 2.706 | 4.109 | 5.248 | 1.388 | 1.675 | 4.26 | 0.135 | 2.601 |
| E | - | - | - | - | - | 4.169 | 3.876 | 0.666 | 6.191 | 4.303 | 4.227 | -0.622 | 4.415 | 3.95 |
| F | - | - | - | - | - | - | 1.48 | 3.798 | 5.496 | 0.725 | -0.57 | 3.982 | 1.531 | 1.103 |
| G | - | - | - | - | - | - | - | 3.513 | 5.602 | 2.135 | 1.83 | 3.751 | 2.533 | -0.801 |
| H | - | - | - | - | - | - | - | - | 6.088 | 4.173 | 4.078 | 1.058 | 4.285 | 3.775 |
| I | - | - | - | - | - | - | - | - | - | 4.976 | 4.967 | 5.436 | 4.877 | 5.06 |
| J | - | - | - | - | - | - | - | - | - | - | -0.415 | 4.098 | 0.498 | 1.868 |
| K | - | - | - | - | - | - | - | - | - | - | - | 4.03 | 1.139 | 1.497 |
| L | - | - | - | - | - | - | - | - | - | - | - | - | 4.456 | 4.005 |
| M | - | - | - | - | - | - | - | - | - | - | - | - | - | 2.314 |
| N | - | - | - | - | - | - | - | - | - | - | - | - | - | - |

| | |
|---|---|
| | Larger than 10 |
| | Between 8 and 9,99 |
| | Between 6 and 7,99 |
| | Between 4 and 5,99 |
| | Between 2 and 3,99 |
| | Between 0 and 1,99 |
| | NA (Calculation failed) |
