## Supplementary Data S2 for "Interlaboratory comparison of standardised metabolomics and lipidomics analyses in human and rodent blood using the MxP® Quant 500 kit": PC ae C40.2.pdf

**Amount of relative differences between laboratories**

| Min | A | B | C | D | E | F | G | H | I | J | K | L | M | N |
|---|---|---|---|---|---|---|---|---|---|---|---|---|---|---|
| A | - | 2.872 | 4.668 | 2.568 | 2.765 | 2.946 | 3.267 | -3.387 | 1.809 | 4.583 | 3.481 | 0.988 | 1.105 | 0.837 |
| B | - | - | 4.095 | 0.429 | -3.052 | -0.783 | 1.66 | 2.728 | 0.831 | 4.103 | 2.236 | 2.153 | 2.413 | 3.12 |
| C | - | - | - | 4.198 | 3.872 | 3.784 | 3.643 | 4.381 | 4.041 | -0.517 | 3.231 | 4.386 | 4.438 | 4.613 |
| D | - | - | - | - | 0.132 | 1.36 | 2.123 | 2.378 | -1.263 | 4.209 | 2.65 | 1.846 | 1.886 | 2.85 |
| E | - | - | - | - | - | -0.182 | 1.209 | 3.192 | 1.677 | 4.031 | 2.811 | 3.14 | 3.447 | 3.769 |
| F | - | - | - | - | - | - | 0.546 | 2.907 | 2.354 | 3.99 | 1.945 | 2.741 | 2.823 | 3.393 |
| G | - | - | - | - | - | - | - | 3.259 | 2.231 | 3.856 | 2.095 | 3.312 | 3.484 | 3.849 |
| H | - | - | - | - | - | - | - | - | 2.334 | 4.682 | 3.596 | 0.889 | 0.643 | 1.104 |
| I | - | - | - | - | - | - | - | - | - | 4.366 | 2.702 | 1.503 | -0.134 | 2.754 |
| J | - | - | - | - | - | - | - | - | - | - | 3.202 | 4.425 | 4.525 | 4.739 |
| K | - | - | - | - | - | - | - | - | - | - | - | 3.718 | 3.765 | 4.108 |
| L | - | - | - | - | - | - | - | - | - | - | - | - | -0.448 | 2.11 |
| M | - | - | - | - | - | - | - | - | - | - | - | - | - | 1.85 |
| N | - | - | - | - | - | - | - | - | - | - | - | - | - | - |

| | |
|---|---|
| | Larger than 10 |
| | Between 8 and 9,99 |
| | Between 6 and 7,99 |
| | Between 4 and 5,99 |
| | Between 2 and 3,99 |
| | Between 0 and 1,99 |
| | NA (Calculation failed) |

| Median | A | B | C | D | E | F | G | H | I | J | K | L | M | N |
|---|---|---|---|---|---|---|---|---|---|---|---|---|---|---|
| A | - | 2.052 | 2.269 | 0.208 | 4.227 | 0.051 | 3.796 | 1.553 | 2.05 | 1.554 | 2.756 | 3.669 | 1.148 | 2.99 |
| B | - | - | -0.533 | 1.47 | 4.456 | 2.314 | 4.107 | 2.753 | -3.18 | 2.755 | 3.382 | 4.011 | 2.602 | 3.528 |
| C | - | - | - | 1.656 | 4.5 | 2.538 | 4.134 | 2.867 | -3.436 | 2.903 | 3.471 | 4.044 | 2.749 | 3.586 |
| D | - | - | - | - | 4.305 | 1.199 | 3.895 | 2.035 | 1.576 | 2.049 | 2.983 | 3.779 | 1.772 | 3.175 |
| E | - | - | - | - | - | 4.442 | 2.775 | 4.322 | 4.89 | 4.294 | 3.907 | 3.037 | 4.371 | 3.804 |
| F | - | - | - | - | - | - | 3.67 | 0.803 | 2.507 | 0.884 | 2.511 | 3.532 | 0.111 | 2.76 |
| G | - | - | - | - | - | - | - | 3.63 | 4.411 | 3.593 | 2.949 | 0.362 | 3.705 | 2.743 |
| H | - | - | - | - | - | - | - | - | 2.878 | -6.391 | 1.984 | 3.33 | -0.478 | 2.363 |
| I | - | - | - | - | - | - | - | - | - | 2.756 | 3.387 | 4.016 | 2.604 | 3.536 |
| J | - | - | - | - | - | - | - | - | - | - | 1.956 | 3.26 | -0.184 | 2.253 |
| K | - | - | - | - | - | - | - | - | - | - | - | 2.594 | 2.301 | 0.002 |
| L | - | - | - | - | - | - | - | - | - | - | - | - | 3.558 | 2.432 |
| M | - | - | - | - | - | - | - | - | - | - | - | - | - | 2.54 |
| N | - | - | - | - | - | - | - | - | - | - | - | - | - | - |

| Max | A | B | C | D | E | F | G | H | I | J | K | L | M | N |
|---|---|---|---|---|---|---|---|---|---|---|---|---|---|---|
| A | - | 1.802 | -0.151 | -1.606 | 4.563 | 1.61 | 4.133 | 1.84 | 2.092 | 2.94 | 3.374 | 3.902 | 1.6 | 3.172 |
| B | - | - | 1.278 | 1.593 | 4.711 | 2.645 | 4.351 | 2.758 | -1.064 | 3.455 | 3.752 | 4.161 | 2.635 | 3.589 |
| C | - | - | - | -0.94 | 4.595 | 1.989 | 4.198 | 2.141 | 1.631 | 3.157 | 3.515 | 3.984 | 1.976 | 3.299 |
| D | - | - | - | - | 4.572 | 1.773 | 4.159 | 1.95 | 1.866 | 3.022 | 3.428 | 3.932 | 1.75 | 3.224 |
| E | - | - | - | - | - | 4.73 | 2.917 | 4.712 | 5.172 | 4.335 | 4.054 | 3.431 | 4.732 | 4.229 |
| F | - | - | - | - | - | - | 3.901 | -0.945 | 2.961 | 2.313 | 2.93 | 3.622 | -4.738 | 2.616 |
| G | - | - | - | - | - | - | - | 4.092 | 4.719 | 3.541 | 3.074 | 1.576 | 4.134 | 3.373 |
| H | - | - | - | - | - | - | - | - | 2.963 | 2.181 | 2.846 | 3.568 | -0.835 | 2.502 |
| I | - | - | - | - | - | - | - | - | - | 3.533 | 3.795 | 4.186 | 2.817 | 3.637 |
| J | - | - | - | - | - | - | - | - | - | - | 1.488 | 2.96 | 2.416 | -0.063 |
| K | - | - | - | - | - | - | - | - | - | - | - | 2.31 | 3.067 | 0.863 |
| L | - | - | - | - | - | - | - | - | - | - | - | - | 3.819 | 2.824 |
| M | - | - | - | - | - | - | - | - | - | - | - | - | - | 2.632 |
| N | - | - | - | - | - | - | - | - | - | - | - | - | - | - |
