## Supplementary Data S2 for "Interlaboratory comparison of standardised metabolomics and lipidomics analyses in human and rodent blood using the MxP® Quant 500 kit": PC ae C40.3.pdf

**Amount of relative differences between laboratories**

| Min | A | B | C | D | E | F | G | H | I | J | K | L | M | N |
|---|---|---|---|---|---|---|---|---|---|---|---|---|---|---|
| A | - | 1.221 | 3.809 | 4.68 | 4.188 | 3.237 | 5.348 | 4.714 | 4.216 | -0.716 | 4.592 | 2.376 | 4.534 | 5.662 |
| B | - | - | 3.531 | 4.806 | 4.336 | 2.833 | 5.405 | 4.539 | 4.367 | 1.657 | 4.683 | 2.931 | 4.655 | 5.702 |
| C | - | - | - | 5.223 | 4.83 | 1.883 | 5.66 | 3.375 | 4.887 | 3.794 | 5.148 | 4.1 | 5.074 | 5.873 |
| D | - | - | - | - | 1.274 | 5.108 | 4.078 | 5.853 | 2.153 | 4.823 | 1.651 | 4.245 | 0.431 | 4.996 |
| E | - | - | - | - | - | 5.042 | 4.616 | 5.763 | -1.315 | 4.508 | 2.171 | 4.008 | 2.388 | 5.246 |
| F | - | - | - | - | - | - | 5.628 | 3.862 | 4.759 | 3.48 | 5.073 | 3.836 | 4.975 | 5.852 |
| G | - | - | - | - | - | - | - | 6.376 | 4.487 | 5.639 | 4.469 | 5.262 | 3.966 | 4.21 |
| H | - | - | - | - | - | - | - | - | 5.243 | 4.6 | 5.527 | 4.669 | 5.38 | 6.054 |
| I | - | - | - | - | - | - | - | - | - | 4.491 | 1.725 | 3.888 | 2.489 | 5.187 |
| J | - | - | - | - | - | - | - | - | - | - | 4.474 | 2.432 | 4.518 | 5.632 |
| K | - | - | - | - | - | - | - | - | - | - | - | 4.113 | 1.707 | 5.093 |
| L | - | - | - | - | - | - | - | - | - | - | - | - | 4.241 | 5.577 |
| M | - | - | - | - | - | - | - | - | - | - | - | - | - | 4.972 |
| N | - | - | - | - | - | - | - | - | - | - | - | - | - | - |

| Median | A | B | C | D | E | F | G | H | I | J | K | L | M | N |
|---|---|---|---|---|---|---|---|---|---|---|---|---|---|---|
| A | - | 0.537 | 3.217 | 3.585 | 3.946 | 2.433 | 4.21 | 3.529 | 3.502 | 1.976 | 3.351 | 2.162 | 3.812 | 4.957 |
| B | - | - | 2.983 | 3.731 | 4.051 | 2.003 | 4.278 | 3.323 | 3.64 | 1.394 | 3.476 | 2.534 | 3.934 | 4.997 |
| C | - | - | - | 4.322 | 4.505 | 1.905 | 4.646 | 0.879 | 4.232 | 2.214 | 4.123 | 3.651 | 4.445 | 5.213 |
| D | - | - | - | - | 2.278 | 4.208 | 3.278 | 4.675 | -0.093 | 3.994 | -2.006 | 2.796 | 1.288 | 4.464 |
| E | - | - | - | - | - | 4.618 | 2.02 | 4.995 | 2.208 | 4.495 | 2.531 | 3.677 | 0.931 | 4.207 |
| F | - | - | - | - | - | - | 4.478 | 2.532 | 3.971 | 0.109 | 3.843 | 3.211 | 4.237 | 5.124 |
| G | - | - | - | - | - | - | - | 5.321 | 3.344 | 4.837 | 3.379 | 4.25 | 3.198 | 3.7 |
| H | - | - | - | - | - | - | - | - | 4.316 | 2.54 | 4.187 | 3.799 | 4.569 | 5.268 |
| I | - | - | - | - | - | - | - | - | - | 4.095 | -0.118 | 2.974 | 1.103 | 4.438 |
| J | - | - | - | - | - | - | - | - | - | - | 3.774 | 2.994 | 4.08 | 5.067 |
| K | - | - | - | - | - | - | - | - | - | - | - | 2.854 | 1.309 | 4.467 |
| L | - | - | - | - | - | - | - | - | - | - | - | - | 3.318 | 4.799 |
| M | - | - | - | - | - | - | - | - | - | - | - | - | - | 4.328 |
| N | - | - | - | - | - | - | - | - | - | - | - | - | - | - |

| Max | A | B | C | D | E | F | G | H | I | J | K | L | M | N |
|---|---|---|---|---|---|---|---|---|---|---|---|---|---|---|
| A | - | -1.159 | 0.888 | 3.363 | 3.322 | 1.185 | 3.358 | 2.962 | -2.746 | 3.337 | 2.735 | 1.729 | 2.405 | 2.995 |
| B | - | - | 1.099 | 3.42 | 3.265 | 0.953 | 3.364 | 2.888 | -1.058 | 3.39 | 2.702 | 1.498 | 2.513 | 3.025 |
| C | - | - | - | 3.059 | 3.542 | 1.946 | 2.99 | 3.23 | 0.766 | 2.983 | 2.144 | 2.367 | 1.775 | 2.545 |
| D | - | - | - | - | 4.234 | 3.59 | -1.303 | 4.073 | 3.243 | -0.198 | 2.123 | 3.643 | 2.251 | 1.283 |
| E | - | - | - | - | - | 3.098 | 4.486 | 1.285 | 3.498 | 4.478 | 4.189 | 2.899 | 4.091 | 4.345 |
| F | - | - | - | - | - | - | 3.684 | 2.518 | 1.244 | 3.658 | 3.178 | 0.194 | 2.965 | 3.427 |
| G | - | - | - | - | - | - | - | 4.056 | 3.213 | -1.871 | 1.931 | 3.615 | 2.167 | 1.012 |
| H | - | - | - | - | - | - | - | - | 3.093 | 4.267 | 3.963 | 2.236 | 3.83 | 4.098 |
| I | - | - | - | - | - | - | - | - | - | 3.285 | 2.595 | 1.754 | 2.379 | 2.948 |
| J | - | - | - | - | - | - | - | - | - | - | 1.733 | 3.6 | 2.068 | 0.601 |
| K | - | - | - | - | - | - | - | - | - | - | - | 3.179 | -0.304 | 0.872 |
| L | - | - | - | - | - | - | - | - | - | - | - | - | 3.165 | 3.561 |
| M | - | - | - | - | - | - | - | - | - | - | - | - | - | 1.29 |
| N | - | - | - | - | - | - | - | - | - | - | - | - | - | - |

| | |
|---|---|
| | Larger than 10 |
| | Between 8 and 9,99 |
| | Between 6 and 7,99 |
| | Between 4 and 5,99 |
| | Between 2 and 3,99 |
| | Between 0 and 1,99 |
| | NA (Calculation failed) |
