## Supplementary Data S2 for "Interlaboratory comparison of standardised metabolomics and lipidomics analyses in human and rodent blood using the MxP® Quant 500 kit": PC ae C40.4.pdf

**Amount of relative differences between laboratories**

| Min | A | B | C | D | E | F | G | H | I | J | K | L | M | N |
|---|---|---|---|---|---|---|---|---|---|---|---|---|---|---|
| A | - | 3.168 | 1.035 | 2.658 | 3.908 | 2.458 | 3.148 | 1.454 | 2.529 | 2.943 | 2.566 | 1.446 | 3.436 | 4.168 |
| B | - | - | 3.673 | 1.936 | 2.621 | 2.078 | -2.853 | 3.748 | 1.95 | 1.59 | 2.177 | 2.866 | 0.39 | 3.074 |
| C | - | - | - | 3.005 | 4.039 | 2.972 | 3.488 | -1.416 | 2.971 | 3.468 | 3.026 | 2.297 | 3.668 | 4.224 |
| D | - | - | - | - | 3.194 | -0.967 | 1.855 | 3.235 | -3.394 | 1.183 | -1.452 | 1.853 | 2.299 | 3.551 |
| E | - | - | - | - | - | 3.444 | 2.759 | 4.361 | 3.414 | 3.264 | 3.498 | 3.847 | 2.21 | 1.978 |
| F | - | - | - | - | - | - | 2.115 | 3.172 | -0.934 | 1.527 | -2.874 | 1.67 | 2.341 | 3.515 |
| G | - | - | - | - | - | - | - | 3.767 | 2.01 | 1.175 | 2.308 | 2.914 | -0.079 | 2.946 |
| H | - | - | - | - | - | - | - | - | 3.052 | 3.631 | 3.114 | 2.449 | 3.743 | 4.294 |
| I | - | - | - | - | - | - | - | - | - | 0.575 | -4.217 | 1.627 | 2.408 | 3.585 |
| J | - | - | - | - | - | - | - | - | - | - | 1.175 | 2.311 | 1.962 | 3.153 |
| K | - | - | - | - | - | - | - | - | - | - | - | 1.071 | 2.583 | 3.529 |
| L | - | - | - | - | - | - | - | - | - | - | - | - | 3.119 | 3.952 |
| M | - | - | - | - | - | - | - | - | - | - | - | - | - | 2.793 |
| N | - | - | - | - | - | - | - | - | - | - | - | - | - | - |

| Median | A | B | C | D | E | F | G | H | I | J | K | L | M | N |
|---|---|---|---|---|---|---|---|---|---|---|---|---|---|---|
| A | - | 2.992 | 1.617 | 2.098 | 3.775 | 0.465 | 2.885 | 1.401 | 2.47 | 1.121 | -0.63 | 1.728 | 2.974 | 3.829 |
| B | - | - | 3.629 | 2.041 | 2.657 | 2.809 | -0.138 | 3.542 | 1.528 | 3.358 | 2.986 | 2.314 | -2.356 | 2.746 |
| C | - | - | - | 2.868 | 4.045 | 2.229 | 3.358 | -0.638 | 3.075 | 0.378 | 1.981 | 2.7 | 3.435 | 4.078 |
| D | - | - | - | - | 3.32 | 1.463 | 1.6 | 2.863 | 0.162 | 2.587 | 1.848 | -0.258 | 1.911 | 3.379 |
| E | - | - | - | - | - | 3.828 | 2.939 | 4.253 | 3.288 | 4.15 | 3.924 | 3.59 | 2.773 | -0.683 |
| F | - | - | - | - | - | - | 2.318 | 2.248 | 1.711 | 1.392 | -0.154 | 0.83 | 2.605 | 3.659 |
| G | - | - | - | - | - | - | - | 3.382 | 0.947 | 3.102 | 2.722 | 1.883 | -0.123 | 2.96 |
| H | - | - | - | - | - | - | - | - | 2.997 | -3.69 | 1.685 | 2.538 | 3.358 | 4.03 |
| I | - | - | - | - | - | - | - | - | - | 2.986 | 2.359 | 1.186 | 1.315 | 3.189 |
| J | - | - | - | - | - | - | - | - | - | - | -0.822 | 1.675 | 2.874 | 3.781 |
| K | - | - | - | - | - | - | - | - | - | - | - | 1.458 | 2.808 | 3.743 |
| L | - | - | - | - | - | - | - | - | - | - | - | - | 2.262 | 3.492 |
| M | - | - | - | - | - | - | - | - | - | - | - | - | - | 2.795 |
| N | - | - | - | - | - | - | - | - | - | - | - | - | - | - |

| Max | A | B | C | D | E | F | G | H | I | J | K | L | M | N |
|---|---|---|---|---|---|---|---|---|---|---|---|---|---|---|
| A | - | 2.691 | 2.439 | 0.497 | 3.599 | 1.988 | 1.687 | 1.325 | 2.345 | 3.312 | 2.185 | 2.003 | 2.225 | 3.402 |
| B | - | - | 3.524 | 2.178 | 2.699 | 3.41 | 1.627 | 3.199 | -0.099 | 3.996 | 3.494 | 1.156 | 0.856 | 2.338 |
| C | - | - | - | 2.65 | 4.051 | -0.779 | 2.91 | 1.027 | 3.262 | 2.672 | -2.428 | 3.067 | 3.134 | 3.921 |
| D | - | - | - | - | 3.457 | 2.504 | 0.458 | 2.118 | 1.825 | 3.645 | 2.762 | 1.297 | 1.333 | 3.191 |
| E | - | - | - | - | - | 4.211 | 3.353 | 4.091 | 2.997 | 4.595 | 4.248 | 3.213 | 3.202 | 1.369 |
| F | - | - | - | - | - | - | 2.773 | 0.135 | 3.131 | 2.797 | 0.721 | 2.937 | 2.843 | 3.786 |
| G | - | - | - | - | - | - | - | 2.6 | 1.178 | 3.762 | 3.039 | -0.188 | -0.17 | 2.973 |
| H | - | - | - | - | - | - | - | - | 2.882 | 3.263 | 1.102 | 2.636 | 2.785 | 3.72 |
| I | - | - | - | - | - | - | - | - | - | 3.937 | 3.34 | 0.389 | -0.897 | 2.662 |
| J | - | - | - | - | - | - | - | - | - | - | 1.742 | 3.685 | 3.461 | 4.204 |
| K | - | - | - | - | - | - | - | - | - | - | - | 3.087 | 3.016 | 3.924 |
| L | - | - | - | - | - | - | - | - | - | - | - | - | -0.688 | 2.841 |
| M | - | - | - | - | - | - | - | - | - | - | - | - | - | 2.797 |
| N | - | - | - | - | - | - | - | - | - | - | - | - | - | - |

| | |
|---|---|
| | Larger than 10 |
| | Between 8 and 9,99 |
| | Between 6 and 7,99 |
| | Between 4 and 5,99 |
| | Between 2 and 3,99 |
| | Between 0 and 1,99 |
| | NA (Calculation failed) |
