## Supplementary Data S2 for "Interlaboratory comparison of standardised metabolomics and lipidomics analyses in human and rodent blood using the MxP® Quant 500 kit": PC ae C40.5.pdf

**Amount of relative differences between laboratories**

| Min | A | B | C | D | E | F | G | H | I | J | K | L | M | N |
|---|---|---|---|---|---|---|---|---|---|---|---|---|---|---|
| A | - | 2.351 | 1.383 | 0.408 | 3.919 | 3.537 | 1.532 | 2.888 | 0.03 | 3.106 | 1.489 | 2.963 | -2.346 | 1.235 |
| B | - | - | 1.713 | 2.075 | 3.503 | 2.627 | 3.274 | 1.181 | 2.413 | -1.107 | 2.165 | 1.537 | 2.504 | 3.185 |
| C | - | - | - | 0.101 | 3.766 | 3.322 | 2.492 | 2.426 | 0.946 | 1.439 | 0.138 | 2.53 | 1.115 | 2.378 |
| D | - | - | - | - | 3.921 | 3.503 | 2.245 | 2.377 | 0.42 | 2.645 | 0.14 | 2.874 | 0.936 | 2.045 |
| E | - | - | - | - | - | 2.242 | 4.51 | 3.061 | 4.245 | 3.321 | 4.097 | 3.232 | 4.3 | 4.446 |
| F | - | - | - | - | - | - | 4.111 | 1.692 | 3.683 | 2.365 | 3.457 | 2.378 | 3.789 | 4.094 |
| G | - | - | - | - | - | - | - | 3.395 | 1.738 | 3.823 | 1.87 | 3.552 | 1.304 | -0.715 |
| H | - | - | - | - | - | - | - | - | 3.002 | 0.92 | 2.741 | -2.834 | 3.124 | 3.591 |
| I | - | - | - | - | - | - | - | - | - | 2.773 | -0.676 | 3.002 | -0.633 | 1.563 |
| J | - | - | - | - | - | - | - | - | - | - | 1.993 | 0.329 | 2.883 | 3.369 |
| K | - | - | - | - | - | - | - | - | - | - | - | 2.751 | 1.057 | 2.032 |
| L | - | - | - | - | - | - | - | - | - | - | - | - | 3.136 | 3.497 |
| M | - | - | - | - | - | - | - | - | - | - | - | - | - | 1.065 |
| N | - | - | - | - | - | - | - | - | - | - | - | - | - | - |

| Median | A | B | C | D | E | F | G | H | I | J | K | L | M | N |
|---|---|---|---|---|---|---|---|---|---|---|---|---|---|---|
| A | - | 3.092 | 1.537 | 3.594 | 4.265 | 2.88 | 3.467 | 2.653 | 2.712 | 2.239 | 2.309 | 3.219 | 2.818 | 2.791 |
| B | - | - | 2.586 | 1.989 | 3.556 | 0.135 | 1.54 | 1.325 | 0.948 | 1.991 | 1.902 | 0.007 | 0.627 | 0.683 |
| C | - | - | - | 3.239 | 4.069 | 2.199 | 3.07 | 1.783 | 1.912 | 0.966 | 1.097 | 2.723 | 2.086 | 2.051 |
| D | - | - | - | - | 3.025 | 2.347 | 0.051 | 2.815 | 2.535 | 3.05 | 2.987 | 1.568 | 2.532 | 2.518 |
| E | - | - | - | - | - | 3.865 | 3.329 | 4.031 | 3.954 | 4.161 | 4.136 | 3.607 | 3.914 | 3.923 |
| F | - | - | - | - | - | - | 2.026 | -0.121 | -0.029 | 1.438 | 1.325 | 1.111 | -2.629 | -1.378 |
| G | - | - | - | - | - | - | - | 2.634 | 2.201 | 2.864 | 2.772 | 0.921 | 2.247 | 2.216 |
| H | - | - | - | - | - | - | - | - | -2.663 | 0.731 | 0.546 | 1.662 | -0.351 | -0.661 |
| I | - | - | - | - | - | - | - | - | - | 1.117 | 0.913 | 1.48 | -2.161 | -2.316 |
| J | - | - | - | - | - | - | - | - | - | - | -2.432 | 2.249 | 1.277 | 1.178 |
| K | - | - | - | - | - | - | - | - | - | - | - | 2.195 | 1.093 | 1.036 |
| L | - | - | - | - | - | - | - | - | - | - | - | - | 1.34 | 1.402 |
| M | - | - | - | - | - | - | - | - | - | - | - | - | - | -6.42 |
| N | - | - | - | - | - | - | - | - | - | - | - | - | - | - |

| Max | A | B | C | D | E | F | G | H | I | J | K | L | M | N |
|---|---|---|---|---|---|---|---|---|---|---|---|---|---|---|
| A | - | 3.189 | 1.576 | 3.812 | 4.33 | 2.664 | 3.766 | 2.605 | 3.016 | 2.033 | 2.436 | 3.263 | 3.088 | 3.119 |
| B | - | - | 2.753 | 2.451 | 3.568 | 1.511 | 2.38 | 1.768 | -0.023 | 2.23 | 1.84 | -0.623 | -0.475 | -2.46 |
| C | - | - | - | 3.506 | 4.128 | 1.718 | 3.454 | 1.621 | 2.419 | 0.873 | 1.497 | 2.756 | 2.467 | 2.566 |
| D | - | - | - | - | 2.724 | 3.177 | -1.478 | 3.224 | 2.815 | 3.409 | 3.229 | 2.292 | 2.713 | 2.593 |
| E | - | - | - | - | - | 4.082 | 2.949 | 4.158 | 3.862 | 4.26 | 4.144 | 3.668 | 3.823 | 3.798 |
| F | - | - | - | - | - | - | 2.995 | -1.048 | 1.215 | 1.211 | -0.486 | 1.824 | 1.361 | 1.481 |
| G | - | - | - | - | - | - | - | 3.227 | 2.767 | 3.424 | 3.177 | 2.153 | 2.63 | 2.497 |
| H | - | - | - | - | - | - | - | - | 1.282 | 0.698 | -1.49 | 1.92 | 1.424 | 1.606 |
| I | - | - | - | - | - | - | - | - | - | 1.856 | 1.271 | 0.857 | -1.332 | -0.32 |
| J | - | - | - | - | - | - | - | - | - | - | 0.092 | 2.435 | 2.132 | 2.219 |
| K | - | - | - | - | - | - | - | - | - | - | - | 2.062 | 1.573 | 1.687 |
| L | - | - | - | - | - | - | - | - | - | - | - | - | 0.356 | -0.004 |
| M | - | - | - | - | - | - | - | - | - | - | - | - | - | -1.38 |
| N | - | - | - | - | - | - | - | - | - | - | - | - | - | - |

| Colour | Legend |
|---|---|
| (red) | Larger than 10 |
| (orange) | Between 8 and 9,99 |
| (light orange) | Between 6 and 7,99 |
| (light yellow-green) | Between 4 and 5,99 |
| (light green) | Between 2 and 3,99 |
| (dark green) | Between 0 and 1,99 |
| (light grey) | NA (Calculation failed) |
