## Supplementary Data S2 for "Interlaboratory comparison of standardised metabolomics and lipidomics analyses in human and rodent blood using the MxP® Quant 500 kit": PC ae C40.6.pdf

**Amount of relative differences between laboratories**

| Min | A | B | C | D | E | F | G | H | I | J | K | L | M | N |
|---|---|---|---|---|---|---|---|---|---|---|---|---|---|---|
| A | - | -1.824 | 3.131 | 1.883 | 3.573 | 0.96 | 2.302 | 2.971 | 1.675 | 3.189 | 3.931 | 2.841 | 2.928 | 2.576 |
| B | - | - | 3.075 | -0.406 | 3.62 | 0.045 | 1.384 | 3.27 | 2.916 | 3.276 | 4.122 | 3.332 | 2.678 | 2.067 |
| C | - | - | - | 2.883 | 4.291 | 2.714 | 2.386 | 4.086 | 3.778 | 1.33 | 2.936 | 4.133 | 1.288 | 2.084 |
| D | - | - | - | - | 3.86 | -0.306 | 0.465 | 3.455 | 2.728 | 2.93 | 3.761 | 3.386 | 2.144 | 1.391 |
| E | - | - | - | - | - | 3.958 | 4.19 | 1.671 | 3.066 | 4.643 | 5.016 | 1.872 | 4.46 | 4.284 |
| F | - | - | - | - | - | - | 1.135 | 3.371 | 2.891 | 3.021 | 3.86 | 3.365 | 2.439 | 1.915 |
| G | - | - | - | - | - | - | - | 3.622 | 3.024 | 2.433 | 3.464 | 3.528 | 1.428 | 0.19 |
| H | - | - | - | - | - | - | - | - | 2.315 | 4.382 | 4.749 | -0.148 | 4.074 | 3.922 |
| I | - | - | - | - | - | - | - | - | - | 3.963 | 4.365 | 1.96 | 3.481 | 3.317 |
| J | - | - | - | - | - | - | - | - | - | - | 2.78 | 4.25 | 1.678 | 2.587 |
| K | - | - | - | - | - | - | - | - | - | - | - | 4.606 | 3.251 | 3.587 |
| L | - | - | - | - | - | - | - | - | - | - | - | - | 3.959 | 3.782 |
| M | - | - | - | - | - | - | - | - | - | - | - | - | - | 0.89 |
| N | - | - | - | - | - | - | - | - | - | - | - | - | - | - |

| | |
|---|---|
| (red) | Larger than 10 |
| (orange) | Between 8 and 9,99 |
| (light orange) | Between 6 and 7,99 |
| (light yellow-green) | Between 4 and 5,99 |
| (light green) | Between 2 and 3,99 |
| (dark green) | Between 0 and 1,99 |
| (grey) | NA (Calculation failed) |

| Median | A | B | C | D | E | F | G | H | I | J | K | L | M | N |
|---|---|---|---|---|---|---|---|---|---|---|---|---|---|---|
| A | - | 3.593 | 1.64 | 1.352 | 4.156 | 2.54 | 0.869 | 3.393 | 2.035 | 2.915 | -0.414 | 2.158 | 2.26 | 1.818 |
| B | - | - | 3.351 | 3.428 | 2.712 | 2.898 | 3.991 | 1.043 | 3.207 | 2.468 | 3.849 | 3.118 | 3.089 | 3.304 |
| C | - | - | - | -1.01 | 3.928 | 1.337 | 2.367 | 2.892 | -0.179 | 2.094 | 1.934 | 0.488 | 0.65 | -1.641 |
| D | - | - | - | - | 3.969 | 1.677 | 2.187 | 3.006 | 0.575 | 2.307 | 1.752 | 0.942 | 1.129 | -0.127 |
| E | - | - | - | - | - | 3.889 | 4.584 | 3.181 | 4.072 | 3.671 | 4.492 | 4.025 | 4 | 4.129 |
| F | - | - | - | - | - | - | 2.994 | 2.323 | 0.782 | 0.853 | 2.738 | 0.334 | 0.039 | 1.223 |
| G | - | - | - | - | - | - | - | 3.594 | 2.537 | 3.192 | 0.122 | 2.628 | 2.694 | 2.39 |
| H | - | - | - | - | - | - | - | - | 2.835 | 1.739 | 3.646 | 2.746 | 2.671 | 2.958 |
| I | - | - | - | - | - | - | - | - | - | 1.834 | 2.349 | -1.391 | -0.478 | -0.725 |
| J | - | - | - | - | - | - | - | - | - | - | 3.04 | 1.53 | 1.476 | 2.017 |
| K | - | - | - | - | - | - | - | - | - | - | - | 2.312 | 2.343 | 1.963 |
| L | - | - | - | - | - | - | - | - | - | - | - | - | -1.405 | -0.066 |
| M | - | - | - | - | - | - | - | - | - | - | - | - | - | 0.346 |
| N | - | - | - | - | - | - | - | - | - | - | - | - | - | - |

| Max | A | B | C | D | E | F | G | H | I | J | K | L | M | N |
|---|---|---|---|---|---|---|---|---|---|---|---|---|---|---|
| A | - | 3.836 | 2.417 | 1.836 | 4.239 | 2.81 | 0.374 | 3.451 | 2.085 | 3.302 | 1.347 | 1.987 | 2.726 | 2.342 |
| B | - | - | 3.397 | 3.631 | 2.465 | 3.078 | 4.183 | 1.985 | 3.572 | 2.308 | 3.779 | 3.6 | 3.141 | 3.433 |
| C | - | - | - | 0.833 | 3.851 | 0.929 | 2.793 | 2.554 | 0.2 | 2.382 | 1.561 | 0.642 | 0.526 | -1.656 |
| D | - | - | - | - | 3.988 | 1.834 | 2.349 | 2.918 | -0.565 | 2.744 | 0.073 | -1.496 | 1.66 | 0.577 |
| E | - | - | - | - | - | 3.878 | 4.641 | 3.325 | 4.182 | 3.464 | 4.341 | 4.217 | 3.918 | 4.102 |
| F | - | - | - | - | - | - | 3.162 | 2.049 | 1.663 | 1.673 | 2.289 | 1.758 | -1.397 | 1.071 |
| G | - | - | - | - | - | - | - | 3.589 | 2.442 | 3.481 | 1.942 | 2.378 | 2.968 | 2.651 |
| H | - | - | - | - | - | - | - | - | 2.903 | -0.154 | 3.21 | 2.958 | 2.271 | 2.715 |
| I | - | - | - | - | - | - | - | - | - | 2.64 | 0.649 | -1.926 | 1.305 | -0.122 |
| J | - | - | - | - | - | - | - | - | - | - | 3.093 | 2.789 | 1.873 | 2.507 |
| K | - | - | - | - | - | - | - | - | - | - | - | 0.493 | 2.143 | 1.365 |
| L | - | - | - | - | - | - | - | - | - | - | - | - | 1.447 | 0.173 |
| M | - | - | - | - | - | - | - | - | - | - | - | - | - | 0.832 |
| N | - | - | - | - | - | - | - | - | - | - | - | - | - | - |
