## Supplementary Data S2 for "Interlaboratory comparison of standardised metabolomics and lipidomics analyses in human and rodent blood using the MxP® Quant 500 kit": PC ae C42.1.pdf

**Amount of relative differences between laboratories**

| Min | A | B | C | D | E | F | G | H | I | J | K | L | M | N |
|---|---|---|---|---|---|---|---|---|---|---|---|---|---|---|
| A | - | 5.088 | 2.886 | 5.17 | 3.122 | 3.144 | 3.7 | -1.048 | 4.92 | 3.713 | 0.598 | 2.46 | 4.898 | 5.016 |
| B | - | - | 4.724 | 2.008 | 4.738 | 4.794 | 4.333 | 5.19 | 5.148 | 4.386 | 5.228 | 4.857 | 1.33 | 1.686 |
| C | - | - | - | 4.85 | 0.194 | -0.062 | 2.558 | 3.047 | 3.976 | 2.42 | 3.222 | 1.018 | 4.498 | 4.701 |
| D | - | - | - | - | 4.746 | 4.848 | 4.223 | 5.127 | 5.148 | 4.433 | 5.136 | 4.786 | 2.405 | 1.103 |
| E | - | - | - | - | - | -1.868 | 2.256 | 3.319 | 5.208 | 2.184 | 3.496 | 1.809 | 4.536 | 4.712 |
| F | - | - | - | - | - | - | 2.437 | 3.076 | 4.022 | 2.347 | 3.228 | 1.291 | 4.418 | 4.648 |
| G | - | - | - | - | - | - | - | 3.856 | 3.276 | -2.811 | 3.962 | 3.028 | 3.981 | 4.324 |
| H | - | - | - | - | - | - | - | - | 4.797 | 3.74 | 0.038 | 2.56 | 4.917 | 5.017 |
| I | - | - | - | - | - | - | - | - | - | 1.189 | 3.01 | 2.733 | 4.281 | 4.539 |
| J | - | - | - | - | - | - | - | - | - | - | 3.813 | 2.902 | 4.068 | 4.39 |
| K | - | - | - | - | - | - | - | - | - | - | - | 2.696 | 4.746 | 4.94 |
| L | - | - | - | - | - | - | - | - | - | - | - | - | 4.736 | 4.85 |
| M | - | - | - | - | - | - | - | - | - | - | - | - | - | 2.605 |
| N | - | - | - | - | - | - | - | - | - | - | - | - | - | - |

| Median | A | B | C | D | E | F | G | H | I | J | K | L | M | N |
|---|---|---|---|---|---|---|---|---|---|---|---|---|---|---|
| A | - | 4.27 | 2.239 | 3.396 | 3.12 | 1.609 | 2.787 | 0.053 | 3.189 | 2.151 | 3.121 | 2.669 | 3.988 | 4.133 |
| B | - | - | 4.08 | 3.22 | 3.662 | 4.374 | 3.979 | 4.721 | 5.35 | 4.154 | 5.081 | 3.979 | 1.89 | 1.169 |
| C | - | - | - | 2.627 | 2.023 | 1.078 | 0.993 | 2.71 | 4.006 | -1.06 | 3.826 | 0.614 | 3.533 | 3.725 |
| D | - | - | - | - | 1.972 | 3.607 | 2.827 | 4.158 | 4.878 | 3.115 | 4.535 | 2.79 | 2.169 | 2.465 |
| E | - | - | - | - | - | 2.62 | 0.955 | 3.41 | 4.077 | 2.236 | 4.253 | 1.353 | 2.978 | 3.287 |
| F | - | - | - | - | - | - | 2.025 | 2.156 | 3.692 | 0.96 | 3.522 | 1.891 | 3.775 | 3.922 |
| G | - | - | - | - | - | - | - | 3.171 | 4.152 | 1.023 | 4.011 | -1.734 | 3.33 | 3.524 |
| H | - | - | - | - | - | - | - | - | 3.064 | 2.461 | 2.903 | 2.884 | 4.077 | 4.203 |
| I | - | - | - | - | - | - | - | - | - | 3.763 | 1.651 | 3.888 | 4.68 | 4.705 |
| J | - | - | - | - | - | - | - | - | - | - | 3.763 | 0.766 | 3.595 | 3.749 |
| K | - | - | - | - | - | - | - | - | - | - | - | 3.769 | 4.494 | 4.572 |
| L | - | - | - | - | - | - | - | - | - | - | - | - | 3.351 | 3.583 |
| M | - | - | - | - | - | - | - | - | - | - | - | - | - | 0.533 |
| N | - | - | - | - | - | - | - | - | - | - | - | - | - | - |

| Max | A | B | C | D | E | F | G | H | I | J | K | L | M | N |
|---|---|---|---|---|---|---|---|---|---|---|---|---|---|---|
| A | - | 2.684 | 0.513 | 3.824 | 3.117 | 2.274 | 1.222 | 1.009 | 6.135 | 2.792 | 4.316 | 2.893 | 0.896 | 1.43 |
| B | - | - | 2.368 | 4.476 | 1.354 | 3.481 | 3.193 | 3.098 | 5.617 | 3.78 | 4.816 | -0.438 | 2.294 | 3.28 |
| C | - | - | - | 3.981 | 2.855 | 2.654 | 1.97 | 1.808 | 6.032 | 3.023 | 4.443 | 2.579 | -1.017 | 2.081 |
| D | - | - | - | - | 4.388 | 3.386 | 3.384 | 3.618 | 4.323 | 3.051 | 2.163 | 4.244 | 3.842 | 3.34 |
| E | - | - | - | - | - | 3.897 | 3.599 | 3.558 | 6.651 | 4.075 | 4.965 | 0.451 | 2.912 | 3.667 |
| F | - | - | - | - | - | - | 1.024 | 1.348 | 5.864 | 1.122 | 3.887 | 3.512 | 2.575 | 0.684 |
| G | - | - | - | - | - | - | - | -0.651 | 5.917 | 2.174 | 4.084 | 3.251 | 2.101 | -2.795 |
| H | - | - | - | - | - | - | - | - | 6.011 | 2.261 | 4.134 | 3.208 | 1.8 | -0.631 |
| I | - | - | - | - | - | - | - | - | - | 5.103 | 4.366 | 5.38 | 4.994 | 4.905 |
| J | - | - | - | - | - | - | - | - | - | - | 3.681 | 3.72 | 2.883 | 1.665 |
| K | - | - | - | - | - | - | - | - | - | - | - | 4.496 | 4.186 | 3.841 |
| L | - | - | - | - | - | - | - | - | - | - | - | - | 2.661 | 3.464 |
| M | - | - | - | - | - | - | - | - | - | - | - | - | - | 2.289 |
| N | - | - | - | - | - | - | - | - | - | - | - | - | - | - |

| | |
|---|---|
| (red) | Larger than 10 |
| (orange) | Between 8 and 9,99 |
| (light orange) | Between 6 and 7,99 |
| (light green) | Between 4 and 5,99 |
| (green) | Between 2 and 3,99 |
| (dark green) | Between 0 and 1,99 |
| (grey) | NA (Calculation failed) |
