## Supplementary Data S2 for "Interlaboratory comparison of standardised metabolomics and lipidomics analyses in human and rodent blood using the MxP® Quant 500 kit": PC ae C42.2.pdf

**Amount of relative differences between laboratories**

| Min | A | B | C | D | E | F | G | H | I | J | K | L | M | N |
|---|---|---|---|---|---|---|---|---|---|---|---|---|---|---|
| A | - | 3.119 | 2.571 | 1.562 | 3.166 | 2.636 | 2.371 | 2.941 | 3.395 | 1.31 | 2.957 | 2.022 | 1.718 | 2.169 |
| B | - | - | 4.008 | 2.718 | -1.987 | 1.556 | 3.93 | 0.126 | 4.501 | 3.745 | 4.185 | 2.352 | 3.709 | 2.215 |
| C | - | - | - | 2.993 | 3.805 | 3.498 | -0.607 | 3.632 | 2.715 | 1.017 | 0.732 | 3.241 | 1.25 | 3.295 |
| D | - | - | - | - | 2.785 | 1.979 | 2.978 | 2.451 | 3.859 | 2.613 | 3.341 | 0.598 | 2.572 | 1.062 |
| E | - | - | - | - | - | 1.591 | 3.966 | -1.406 | 4.131 | 3.694 | 4.21 | 2.432 | 3.765 | 2.282 |
| F | - | - | - | - | - | - | 3.605 | 1.043 | 4.022 | 3.324 | 3.878 | 1.231 | 3.346 | 0.935 |
| G | - | - | - | - | - | - | - | 3.66 | 2.591 | 1.097 | 1.365 | 3.126 | 0.824 | 3.188 |
| H | - | - | - | - | - | - | - | - | 4.456 | 3.487 | 3.752 | 1.143 | 3.366 | 0.862 |
| I | - | - | - | - | - | - | - | - | - | 2.194 | 2.168 | 3.79 | 2.761 | 3.942 |
| J | - | - | - | - | - | - | - | - | - | - | 2.461 | 2.747 | -1.989 | 2.955 |
| K | - | - | - | - | - | - | - | - | - | - | - | 3.538 | 2.251 | 3.582 |
| L | - | - | - | - | - | - | - | - | - | - | - | - | 2.936 | -0.768 |
| M | - | - | - | - | - | - | - | - | - | - | - | - | - | 2.906 |
| N | - | - | - | - | - | - | - | - | - | - | - | - | - | - |

| Median | A | B | C | D | E | F | G | H | I | J | K | L | M | N |
|---|---|---|---|---|---|---|---|---|---|---|---|---|---|---|
| A | - | 3.077 | 2.913 | -1.834 | 3.132 | 2.238 | 2.26 | -0.151 | 4.39 | 1.217 | 2.437 | 1.823 | 1.065 | 2.069 |
| B | - | - | 4.122 | 3.251 | -1.583 | 2.012 | 3.852 | 3.348 | 5.007 | 2.755 | 3.922 | 2.416 | 3.522 | 2.206 |
| C | - | - | - | 2.741 | 3.903 | 3.513 | 1.288 | 2.591 | 3.653 | 3.172 | 0.872 | 3.356 | 2.308 | 3.441 |
| D | - | - | - | - | 3.135 | 2.36 | 2.206 | -0.535 | 4.36 | 1.322 | 2.399 | 1.912 | 0.932 | 2.16 |
| E | - | - | - | - | - | 2.135 | 3.892 | 3.385 | 5.016 | 2.819 | 3.959 | 2.511 | 3.572 | 2.314 |
| F | - | - | - | - | - | - | 3.341 | 2.578 | 4.75 | 1.413 | 3.435 | 0.427 | 2.867 | -0.658 |
| G | - | - | - | - | - | - | - | 1.91 | 3.956 | 2.753 | -0.737 | 2.988 | 1.362 | 3.097 |
| H | - | - | - | - | - | - | - | - | 4.258 | 1.775 | 2.139 | 2.239 | 0.05 | 2.452 |
| I | - | - | - | - | - | - | - | - | - | 4.223 | 3.614 | 4.315 | 3.91 | 4.367 |
| J | - | - | - | - | - | - | - | - | - | - | 2.934 | 0.395 | 2.111 | 0.977 |
| K | - | - | - | - | - | - | - | - | - | - | - | 3.076 | 1.649 | 3.172 |
| L | - | - | - | - | - | - | - | - | - | - | - | - | 2.548 | -0.513 |
| M | - | - | - | - | - | - | - | - | - | - | - | - | - | 2.613 |
| N | - | - | - | - | - | - | - | - | - | - | - | - | - | - |

| Max | A | B | C | D | E | F | G | H | I | J | K | L | M | N |
|---|---|---|---|---|---|---|---|---|---|---|---|---|---|---|
| A | - | 3.04 | 3.124 | 1.332 | 3.095 | 1.845 | 2.163 | 3.089 | 4.92 | 2.573 | 1.531 | 1.631 | 0.182 | 1.987 |
| B | - | - | 4.202 | 3.53 | -1.253 | 2.279 | 3.786 | 4.19 | 5.344 | 0.789 | 3.572 | 2.468 | 3.352 | 2.199 |
| C | - | - | - | 2.536 | 4 | 3.524 | 1.958 | -1.422 | 4.165 | 3.837 | 2.48 | 3.446 | 2.81 | 3.545 |
| D | - | - | - | - | 3.429 | 2.591 | 0.98 | 2.621 | 4.695 | 3.171 | -4.013 | 2.502 | 0.406 | 2.659 |
| E | - | - | - | - | - | 2.44 | 3.829 | 4.276 | 5.51 | 1.387 | 3.629 | 2.575 | 3.396 | 2.338 |
| F | - | - | - | - | - | - | 3.086 | 3.711 | 5.186 | 1.53 | 2.73 | -1.006 | 2.32 | -1.486 |
| G | - | - | - | - | - | - | - | 2.042 | 4.583 | 3.353 | 0.605 | 2.86 | 1.682 | 3.022 |
| H | - | - | - | - | - | - | - | - | 4.055 | 3.856 | 2.415 | 3.404 | 2.675 | 3.475 |
| I | - | - | - | - | - | - | - | - | - | 4.874 | 4.362 | 4.647 | 4.436 | 4.628 |
| J | - | - | - | - | - | - | - | - | - | - | 3.312 | 1.701 | 2.921 | 1.331 |
| K | - | - | - | - | - | - | - | - | - | - | - | 2.522 | 0.877 | 2.749 |
| L | - | - | - | - | - | - | - | - | - | - | - | - | 2.139 | -0.342 |
| M | - | - | - | - | - | - | - | - | - | - | - | - | - | 2.336 |
| N | - | - | - | - | - | - | - | - | - | - | - | - | - | - |

| Colour | Legend |
|---|---|
| (red) | Larger than 10 |
| (orange) | Between 8 and 9,99 |
| (light orange) | Between 6 and 7,99 |
| (light yellow-green) | Between 4 and 5,99 |
| (light green) | Between 2 and 3,99 |
| (dark green) | Between 0 and 1,99 |
| (grey) | NA (Calculation failed) |
