## Supplementary Data S2 for "Interlaboratory comparison of standardised metabolomics and lipidomics analyses in human and rodent blood using the MxP® Quant 500 kit": PC ae C42.3.pdf

**Amount of relative differences between laboratories**

| Min | A | B | C | D | E | F | G | H | I | J | K | L | M | N |
|---|---|---|---|---|---|---|---|---|---|---|---|---|---|---|
| A | - | 2.414 | 3.754 | 1.714 | 2.626 | 3.671 | 0.519 | 4.042 | -1.11 | 4.036 | 2.745 | -1.458 | 1.507 | 3.151 |
| B | - | - | 3.048 | 1.132 | 3.401 | 2.794 | 1.748 | 3.598 | 2.745 | 3.553 | -0.052 | 2.466 | 3.04 | 3.762 |
| C | - | - | - | 3.151 | 3.922 | -0.614 | 3.165 | 1.707 | 3.789 | 2.469 | 2.587 | 3.57 | 3.862 | 4.291 |
| D | - | - | - | - | 3.235 | 3.367 | 1.212 | 4.155 | 1.254 | 3.748 | 1.638 | 1.9 | 2.69 | 3.567 |
| E | - | - | - | - | - | 4.308 | 3.043 | 4.374 | 1.167 | 4.631 | 3.767 | 2.64 | 1.848 | 2.13 |
| F | - | - | - | - | - | - | 3.329 | 2.111 | 3.799 | 1.673 | 2.566 | 3.549 | 3.819 | 4.308 |
| G | - | - | - | - | - | - | - | 3.87 | 0.18 | 3.956 | 2.353 | 0.89 | 2.07 | 3.358 |
| H | - | - | - | - | - | - | - | - | 4.648 | 0.562 | 3.367 | 3.844 | 4.16 | 4.579 |
| I | - | - | - | - | - | - | - | - | - | 4.12 | 2.438 | -0.111 | 2.418 | 3.401 |
| J | - | - | - | - | - | - | - | - | - | - | 3.13 | 3.866 | 4.09 | 4.495 |
| K | - | - | - | - | - | - | - | - | - | - | - | 2.844 | 3.196 | 3.867 |
| L | - | - | - | - | - | - | - | - | - | - | - | - | 1.186 | 3.132 |
| M | - | - | - | - | - | - | - | - | - | - | - | - | - | 2.516 |
| N | - | - | - | - | - | - | - | - | - | - | - | - | - | - |

| Median | A | B | C | D | E | F | G | H | I | J | K | L | M | N |
|---|---|---|---|---|---|---|---|---|---|---|---|---|---|---|
| A | - | 2.54 | 3.466 | 1.758 | 2.947 | 3.628 | -1.058 | 3.42 | 2.898 | 3.508 | 2.07 | 1.703 | 1.302 | 2.166 |
| B | - | - | 2.294 | 1.22 | 3.678 | 2.634 | 2.579 | 2.231 | 0.699 | 2.385 | 0.62 | 3.1 | 2.973 | 3.286 |
| C | - | - | - | 2.783 | 4.08 | -0.115 | 3.379 | -1.24 | 1.491 | -1.91 | 2.683 | 3.699 | 3.634 | 3.811 |
| D | - | - | - | - | 3.424 | 3.142 | 1.9 | 2.876 | 1.913 | 2.951 | -0.359 | 2.683 | 2.512 | 2.928 |
| E | - | - | - | - | - | 4.428 | 2.975 | 4.3 | 4.104 | 4.373 | 3.668 | 2.25 | 2.464 | 1.795 |
| F | - | - | - | - | - | - | 3.51 | 0.495 | 1.998 | -0.259 | 2.874 | 3.796 | 3.726 | 3.905 |
| G | - | - | - | - | - | - | - | 3.473 | 3.022 | 3.575 | 2.16 | 1.534 | 1.138 | 2.02 |
| H | - | - | - | - | - | - | - | - | -0.634 | -0.122 | 2.671 | 3.656 | 3.62 | 3.762 |
| I | - | - | - | - | - | - | - | - | - | 2.177 | 0.696 | 3.153 | 3.011 | 3.363 |
| J | - | - | - | - | - | - | - | - | - | - | 2.798 | 3.74 | 3.691 | 3.84 |
| K | - | - | - | - | - | - | - | - | - | - | - | 2.816 | 2.682 | 3.018 |
| L | - | - | - | - | - | - | - | - | - | - | - | - | -0.586 | 0.206 |
| M | - | - | - | - | - | - | - | - | - | - | - | - | - | 1.076 |
| N | - | - | - | - | - | - | - | - | - | - | - | - | - | - |

| Max | A | B | C | D | E | F | G | H | I | J | K | L | M | N |
|---|---|---|---|---|---|---|---|---|---|---|---|---|---|---|
| A | - | 2.627 | 3.275 | 1.784 | 3.105 | 3.594 | 0.636 | 2.733 | 3.358 | 3.123 | 0.892 | 2.377 | 1.105 | 0.706 |
| B | - | - | 1.592 | 1.27 | 3.816 | 2.499 | 2.896 | -0.978 | 2.19 | 0.82 | 1.955 | 3.409 | 2.914 | 2.864 |
| C | - | - | - | 2.506 | 4.164 | 1.089 | 3.487 | 1.591 | 0.106 | 1.341 | 2.767 | 3.78 | 3.411 | 3.384 |
| D | - | - | - | - | 3.523 | 2.943 | 2.179 | -6.934 | 2.57 | 2.234 | 0.488 | 3.041 | 2.343 | 2.28 |
| E | - | - | - | - | - | 4.513 | 2.935 | 4.245 | 4.491 | 4.214 | 3.567 | 1.913 | 2.833 | 3.099 |
| F | - | - | - | - | - | - | 3.603 | 2.441 | -1.021 | 1.572 | 3.112 | 3.94 | 3.642 | 3.565 |
| G | - | - | - | - | - | - | - | 3.111 | 3.51 | 3.322 | 1.952 | 1.846 | -0.936 | -5.286 |
| H | - | - | - | - | - | - | - | - | 2.996 | 1.132 | 1.42 | 3.514 | 2.929 | 2.775 |
| I | - | - | - | - | - | - | - | - | - | 1.209 | 3.036 | 3.831 | 3.371 | 3.338 |
| J | - | - | - | - | - | - | - | - | - | - | 2.398 | 3.649 | 3.242 | 3.166 |
| K | - | - | - | - | - | - | - | - | - | - | - | 2.797 | 2.039 | 1.955 |
| L | - | - | - | - | - | - | - | - | - | - | - | - | 1.648 | 1.924 |
| M | - | - | - | - | - | - | - | - | - | - | - | - | - | -2.07 |
| N | - | - | - | - | - | - | - | - | - | - | - | - | - | - |

| | |
|---|---|
| | Larger than 10 |
| | Between 8 and 9,99 |
| | Between 6 and 7,99 |
| | Between 4 and 5,99 |
| | Between 2 and 3,99 |
| | Between 0 and 1,99 |
| | NA (Calculation failed) |
