## Supplementary Data S2 for "Interlaboratory comparison of standardised metabolomics and lipidomics analyses in human and rodent blood using the MxP® Quant 500 kit": PC ae C42.4.pdf

**Amount of relative differences between laboratories**

| Min | A | B | C | D | E | F | G | H | I | J | K | L | M | N |
|---|---|---|---|---|---|---|---|---|---|---|---|---|---|---|
| A | - | 3.358 | -0.606 | 2.597 | -0.686 | 3.225 | 4.114 | 2.146 | 3.786 | 3.785 | 1.979 | 3.138 | 3.178 | 4.335 |
| B | - | - | 3.364 | 4.257 | 3.627 | 1.609 | 5.05 | 3.058 | 2.002 | 4.955 | 3.161 | 4.492 | 0.469 | 3.219 |
| C | - | - | - | 2.983 | 1.219 | 2.837 | 4.371 | 0.997 | 3.699 | 4.259 | 0.463 | 3.439 | 3.04 | 4.175 |
| D | - | - | - | - | 2.839 | 3.964 | 3.302 | 3.4 | 4.128 | 3.033 | 2.955 | 0.501 | 3.773 | 4.75 |
| E | - | - | - | - | - | 3.509 | 4.115 | 3.026 | 3.577 | 4.015 | -0.962 | 3.164 | 2.738 | 4.712 |
| F | - | - | - | - | - | - | 4.868 | 2.445 | 2.456 | 4.745 | 2.653 | 4.257 | -0.765 | 3.706 |
| G | - | - | - | - | - | - | - | 4.253 | 4.512 | 0.58 | 3.795 | 2.847 | 4.277 | 5.164 |
| H | - | - | - | - | - | - | - | - | 3.486 | 4.34 | -1.183 | 3.765 | 2.609 | 4.11 |
| I | - | - | - | - | - | - | - | - | - | 5.243 | 3.772 | 4.821 | 2.536 | 2.107 |
| J | - | - | - | - | - | - | - | - | - | - | 3.802 | 2.573 | 4.288 | 5.079 |
| K | - | - | - | - | - | - | - | - | - | - | - | 3.409 | 2.709 | 4.501 |
| L | - | - | - | - | - | - | - | - | - | - | - | - | 3.866 | 4.869 |
| M | - | - | - | - | - | - | - | - | - | - | - | - | - | 3.612 |
| N | - | - | - | - | - | - | - | - | - | - | - | - | - | - |

| Median | A | B | C | D | E | F | G | H | I | J | K | L | M | N |
|---|---|---|---|---|---|---|---|---|---|---|---|---|---|---|
| A | - | 2.189 | 2.02 | 1.333 | 4.034 | 2.832 | 2.79 | -0.466 | 2.522 | 0.814 | 3.328 | 2.42 | 3.164 | 4.17 |
| B | - | - | 3.216 | 1.217 | 3.541 | 1.415 | 1.219 | 2.497 | 0.012 | 2.87 | 2.342 | -1.692 | 2.005 | 3.772 |
| C | - | - | - | 2.673 | 4.267 | 3.444 | 3.413 | 1.735 | 3.237 | 0.934 | 3.724 | 3.142 | 3.612 | 4.395 |
| D | - | - | - | - | 3.891 | 2.195 | 2.182 | 1.752 | 1.769 | 1.882 | 3.065 | 1.694 | 2.822 | 4.011 |
| E | - | - | - | - | - | 3.724 | 3.695 | 4.513 | 3.864 | 4.454 | 2.95 | 3.812 | 3.24 | -0.198 |
| F | - | - | - | - | - | - | -1.884 | 3.081 | 0.628 | 3.284 | 1.565 | 1.063 | 0.84 | 3.527 |
| G | - | - | - | - | - | - | - | 3.075 | 0.245 | 2.949 | 2.284 | -0.233 | 1.722 | 3.638 |
| H | - | - | - | - | - | - | - | - | 2.653 | 0.484 | 3.367 | 2.532 | 3.216 | 4.196 |
| I | - | - | - | - | - | - | - | - | - | 3.118 | 1.936 | -0.203 | 1.504 | 3.665 |
| J | - | - | - | - | - | - | - | - | - | - | 3.542 | 2.719 | 3.369 | 4.243 |
| K | - | - | - | - | - | - | - | - | - | - | - | 2.48 | 0.522 | 3.057 |
| L | - | - | - | - | - | - | - | - | - | - | - | - | 2.001 | 3.728 |
| M | - | - | - | - | - | - | - | - | - | - | - | - | - | 3.312 |
| N | - | - | - | - | - | - | - | - | - | - | - | - | - | - |

| Max | A | B | C | D | E | F | G | H | I | J | K | L | M | N |
|---|---|---|---|---|---|---|---|---|---|---|---|---|---|---|
| A | - | 1.644 | 2.456 | 1.997 | 4.357 | 2.765 | 3.342 | 0.94 | 1.695 | 0.123 | 3.575 | 3.071 | 3.159 | 4.125 |
| B | - | - | 3.168 | -0.83 | 4.124 | 1.806 | 2.816 | 2.329 | -3.743 | 0.763 | 3.23 | 2.387 | 2.6 | 3.885 |
| C | - | - | - | 3.143 | 4.612 | 3.514 | 3.89 | 1.863 | 3.057 | 2.786 | 4.075 | 3.708 | 3.781 | 4.446 |
| D | - | - | - | - | 4.061 | 1.575 | 2.69 | 2.721 | -0.035 | 1.39 | 3.097 | 2.083 | 2.193 | 3.739 |
| E | - | - | - | - | - | 4.069 | 3.629 | 4.908 | 4.512 | 4.541 | 3.359 | 3.921 | 3.96 | 2.487 |
| F | - | - | - | - | - | - | 1.839 | 3.195 | 1.902 | 2.51 | 2.537 | 0.662 | 1.167 | 3.477 |
| G | - | - | - | - | - | - | - | 3.888 | 3.115 | 3.308 | 1.07 | 1.371 | 1.485 | 2.694 |
| H | - | - | - | - | - | - | - | - | 2.251 | 1.616 | 3.769 | 3.316 | 3.392 | 4.217 |
| I | - | - | - | - | - | - | - | - | - | 0.823 | 3.203 | 2.333 | 2.611 | 3.89 |
| J | - | - | - | - | - | - | - | - | - | - | 3.452 | 2.749 | 2.774 | 3.92 |
| K | - | - | - | - | - | - | - | - | - | - | - | 2.168 | 2.157 | 2.221 |
| L | - | - | - | - | - | - | - | - | - | - | - | - | -7.085 | 3.196 |
| M | - | - | - | - | - | - | - | - | - | - | - | - | - | 3.224 |
| N | - | - | - | - | - | - | - | - | - | - | - | - | - | - |

| | |
|---|---|
| (red) | Larger than 10 |
| (orange) | Between 8 and 9,99 |
| (light orange) | Between 6 and 7,99 |
| (light green) | Between 4 and 5,99 |
| (green) | Between 2 and 3,99 |
| (dark green) | Between 0 and 1,99 |
| (grey) | NA (Calculation failed) |
