## Supplementary Data S2 for "Interlaboratory comparison of standardised metabolomics and lipidomics analyses in human and rodent blood using the MxP® Quant 500 kit": PC ae C42.5.pdf

**Amount of relative differences between laboratories**

| Min | A | B | C | D | E | F | G | H | I | J | K | L | M | N |
|---|---|---|---|---|---|---|---|---|---|---|---|---|---|---|
| A | - | -3.512 | 1.919 | 0.399 | 3.482 | 1.435 | 0.162 | 1.497 | 0.457 | -1.564 | 1.329 | 2.101 | 2.483 | 2.629 |
| B | - | - | 1.847 | 0.329 | 3.472 | 1.404 | 0.637 | 1.709 | 0.47 | -0.086 | 1.522 | 2.102 | 2.373 | 2.51 |
| C | - | - | - | 1.159 | 3.832 | 2.616 | 1.035 | 2.676 | 1.186 | 1.527 | -0.027 | 2.936 | 3.157 | 3.221 |
| D | - | - | - | - | 3.576 | 1.86 | -6.898 | 2.485 | -1.513 | -1.267 | 0.772 | 2.428 | 2.54 | 2.724 |
| E | - | - | - | - | - | 3.224 | 3.738 | 2.73 | 3.806 | 3.716 | 3.96 | 2.925 | 2.777 | 2.516 |
| F | - | - | - | - | - | - | 1.946 | -3.316 | 2.023 | 1.677 | 2.463 | 0.725 | 1.678 | 1.844 |
| G | - | - | - | - | - | - | - | 2.836 | -4.284 | -1.181 | 0.719 | 2.41 | 2.483 | 2.698 |
| H | - | - | - | - | - | - | - | - | 1.999 | 1.814 | 2.444 | 0.508 | 1.633 | 1.775 |
| I | - | - | - | - | - | - | - | - | - | -1.109 | 0.559 | 2.507 | 2.672 | 2.805 |
| J | - | - | - | - | - | - | - | - | - | - | 1.048 | 2.288 | 2.456 | 2.716 |
| K | - | - | - | - | - | - | - | - | - | - | - | 2.822 | 2.863 | 3.05 |
| L | - | - | - | - | - | - | - | - | - | - | - | - | -0.056 | 0.66 |
| M | - | - | - | - | - | - | - | - | - | - | - | - | - | -0.8 |
| N | - | - | - | - | - | - | - | - | - | - | - | - | - | - |

| Median | A | B | C | D | E | F | G | H | I | J | K | L | M | N |
|---|---|---|---|---|---|---|---|---|---|---|---|---|---|---|
| A | - | 1.097 | 1.804 | 1.444 | 3.727 | 0.696 | 1.861 | 0.007 | 0.384 | 1.425 | 0.279 | 2.573 | 2.732 | 2.839 |
| B | - | - | 2.501 | -0.415 | 3.522 | -1.475 | 0.687 | 0.47 | -0.473 | -0.583 | -0.575 | 2.008 | 2.308 | 2.357 |
| C | - | - | - | 2.585 | 4.015 | 2.305 | 2.779 | 2.107 | 2.214 | 2.572 | 2.183 | 3.189 | 3.295 | 3.361 |
| D | - | - | - | - | 3.434 | 0.156 | -0.253 | 1.369 | 0.6 | -3.789 | 0.352 | 1.711 | 2.139 | 2.091 |
| E | - | - | - | - | - | 3.741 | 3.477 | 3.889 | 3.787 | 3.609 | 3.774 | 3.077 | 2.88 | 2.854 |
| F | - | - | - | - | - | - | 1.061 | -1.259 | -1.8 | 0.123 | -0.942 | 2.146 | 2.323 | 2.516 |
| G | - | - | - | - | - | - | - | 1.888 | 1.231 | -0.155 | 1.026 | 1.299 | 1.882 | 1.769 |
| H | - | - | - | - | - | - | - | - | -3.845 | 0.496 | -2.387 | 2.247 | 2.395 | 2.603 |
| I | - | - | - | - | - | - | - | - | - | 0.473 | -3.188 | 2.237 | 2.513 | 2.526 |
| J | - | - | - | - | - | - | - | - | - | - | 0.305 | 1.74 | 2.144 | 2.126 |
| K | - | - | - | - | - | - | - | - | - | - | - | 2.156 | 2.489 | 2.433 |
| L | - | - | - | - | - | - | - | - | - | - | - | - | -0.105 | 0.219 |
| M | - | - | - | - | - | - | - | - | - | - | - | - | - | -2.028 |
| N | - | - | - | - | - | - | - | - | - | - | - | - | - | - |

| Max | A | B | C | D | E | F | G | H | I | J | K | L | M | N |
|---|---|---|---|---|---|---|---|---|---|---|---|---|---|---|
| A | - | 1.871 | 1.699 | 2.629 | 3.948 | -0.863 | 2.886 | -3.088 | 1.95 | 2.4 | 2.198 | 2.873 | 3.056 | 2.996 |
| B | - | - | 2.89 | 1.305 | 3.573 | 1.734 | 2.041 | 1.871 | -3.546 | 1.034 | 0.403 | 1.925 | 2.197 | 2.212 |
| C | - | - | - | 3.207 | 4.186 | 1.913 | 3.415 | 1.718 | 2.786 | 3.1 | 2.958 | 3.37 | 3.49 | 3.47 |
| D | - | - | - | - | 3.268 | 2.535 | 0.582 | 2.717 | 1.395 | -1.027 | -0.174 | 0.613 | 1.113 | 1.184 |
| E | - | - | - | - | - | 4.116 | 3.23 | 4.234 | 3.768 | 3.51 | 3.581 | 3.192 | 3.03 | 3.09 |
| F | - | - | - | - | - | - | 2.763 | -0.576 | 1.732 | 2.252 | 2.001 | 2.748 | 2.977 | 2.912 |
| G | - | - | - | - | - | - | - | 3.169 | 2.182 | 1.033 | 1.256 | -3.304 | -0.619 | -0.324 |
| H | - | - | - | - | - | - | - | - | 1.983 | 2.494 | 2.191 | 2.906 | 3.121 | 3.057 |
| I | - | - | - | - | - | - | - | - | - | 1.125 | 0.627 | 1.969 | 2.217 | 2.238 |
| J | - | - | - | - | - | - | - | - | - | - | -1.056 | 1.048 | 1.442 | 1.321 |
| K | - | - | - | - | - | - | - | - | - | - | - | 1.199 | 1.569 | 1.561 |
| L | - | - | - | - | - | - | - | - | - | - | - | - | -0.185 | -0.301 |
| M | - | - | - | - | - | - | - | - | - | - | - | - | - | -4.879 |
| N | - | - | - | - | - | - | - | - | - | - | - | - | - | - |

| | |
|---|---|
| (red) | Larger than 10 |
| (orange) | Between 8 and 9,99 |
| (yellow) | Between 6 and 7,99 |
| (light yellow-green) | Between 4 and 5,99 |
| (light green) | Between 2 and 3,99 |
| (dark green) | Between 0 and 1,99 |
| (light grey) | NA (Calculation failed) |
