## Supplementary Data S2 for "Interlaboratory comparison of standardised metabolomics and lipidomics analyses in human and rodent blood using the MxP® Quant 500 kit": PC ae C44.3.pdf

**Amount of relative differences between laboratories**

| Min | A | B | C | D | E | F | G | H | I | J | K | L | M | N |
|---|---|---|---|---|---|---|---|---|---|---|---|---|---|---|
| A | - | 4.322 | 5.329 | NA | 1.716 | 4.962 | 3.543 | 6.005 | -0.835 | 2.934 | -2.695 | 2.909 | 3.07 | 1.826 |
| B | - | - | 3.057 | NA | 1.924 | 0.99 | 5.443 | 4.166 | 5.188 | 5.813 | 8.091 | -0.092 | 4.121 | 5.05 |
| C | - | - | - | NA | 4.809 | 2.576 | 6.793 | 3.171 | 5.433 | 6.422 | 8.041 | 4.901 | 4.59 | 5.289 |
| D | - | - | - | - | NA | NA | NA | NA | NA | NA | NA | NA | NA | NA |
| E | - | - | - | - | - | 4.772 | 3.679 | 5.444 | 5.545 | 3.185 | -1.696 | 2.023 | 0.944 | 2.703 |
| F | - | - | - | - | - | - | 6.998 | 4.134 | 5.185 | 6.334 | 7.981 | 4.992 | 4.194 | 4.961 |
| G | - | - | - | - | - | - | - | 6.592 | 4.521 | 4.429 | 3.837 | 4.174 | 4.424 | 4.002 |
| H | - | - | - | - | - | - | - | - | 5.66 | 7.01 | 10.176 | 4.989 | 4.938 | 5.509 |
| I | - | - | - | - | - | - | - | - | - | 3.938 | 5.198 | 4.595 | 1.237 | -0.131 |
| J | - | - | - | - | - | - | - | - | - | - | 2.378 | 2.106 | 2.216 | -0.048 |
| K | - | - | - | - | - | - | - | - | - | - | - | 2.017 | 4.517 | 3.678 |
| L | - | - | - | - | - | - | - | - | - | - | - | - | 0.299 | 3.068 |
| M | - | - | - | - | - | - | - | - | - | - | - | - | - | 3.315 |
| N | - | - | - | - | - | - | - | - | - | - | - | - | - | - |

| | |
|---|---|
| | Larger than 10 |
| | Between 8 and 9,99 |
| | Between 6 and 7,99 |
| | Between 4 and 5,99 |
| | Between 2 and 3,99 |
| | Between 0 and 1,99 |
| | NA (Calculation failed) |

| Median | A | B | C | D | E | F | G | H | I | J | K | L | M | N |
|---|---|---|---|---|---|---|---|---|---|---|---|---|---|---|
| A | - | 2.23 | 3.547 | NA | -2.214 | 3.505 | 1.906 | 4.842 | 2.378 | 3.038 | -3.014 | -4.19 | 1.742 | 0.339 |
| B | - | - | 3.038 | NA | -0.022 | 2.764 | 1.785 | 4.15 | 1.584 | 2.818 | 3.88 | 0.592 | 2.576 | -0.031 |
| C | - | - | - | NA | 2.97 | 0.423 | 3.64 | 3.142 | 2.318 | 1.046 | 2.637 | 2.389 | 3.716 | 2.745 |
| D | - | - | - | - | NA | NA | NA | NA | NA | NA | NA | NA | NA | NA |
| E | - | - | - | - | - | 3.453 | 1.768 | 4.564 | 0.29 | 3.118 | -1.663 | -1.001 | 1.975 | 0.312 |
| F | - | - | - | - | - | - | 3.669 | 3.49 | 2.046 | 0.713 | 2.567 | 2.302 | 3.603 | 2.625 |
| G | - | - | - | - | - | - | - | 5.329 | 2.72 | 3.65 | 1.589 | 1.736 | 0.555 | 2.412 |
| H | - | - | - | - | - | - | - | - | 3.69 | 3.532 | 5.415 | 3.703 | 4.337 | 3.836 |
| I | - | - | - | - | - | - | - | - | - | 1.487 | 2.942 | 2.689 | 3.114 | 2.083 |
| J | - | - | - | - | - | - | - | - | - | - | 2.97 | 2.909 | 3.332 | 2.656 |
| K | - | - | - | - | - | - | - | - | - | - | - | -2.311 | 0.284 | 0.806 |
| L | - | - | - | - | - | - | - | - | - | - | - | - | 2.291 | -0.871 |
| M | - | - | - | - | - | - | - | - | - | - | - | - | - | 2.612 |
| N | - | - | - | - | - | - | - | - | - | - | - | - | - | - |

| Max | A | B | C | D | E | F | G | H | I | J | K | L | M | N |
|---|---|---|---|---|---|---|---|---|---|---|---|---|---|---|
| A | - | 4.459 | 4.019 | NA | 1.327 | 3.85 | -0.527 | 3.796 | 3.219 | 3.089 | -2.043 | 1.347 | 4.011 | -1.618 |
| B | - | - | 3.019 | NA | -0.225 | 3.705 | 4.252 | 4.129 | 4.963 | 5.324 | 7.104 | 0.76 | 2.696 | 4.631 |
| C | - | - | - | NA | 2.431 | 2.223 | 5.42 | 3.106 | 4.48 | 5.378 | 6.924 | 1.519 | 0.307 | 3.991 |
| D | - | - | - | - | NA | NA | NA | NA | NA | NA | NA | NA | NA | NA |
| E | - | - | - | - | - | 3.369 | 0.904 | -2.608 | 5.148 | 3.081 | -0.779 | -0.573 | 3.365 | 2.734 |
| F | - | - | - | - | - | - | 5.699 | 1.873 | 4.242 | 5.307 | 6.864 | 1.968 | 2.448 | 3.494 |
| G | - | - | - | - | - | - | - | 4.767 | 4.792 | 2.976 | 0.859 | 0.593 | 4.343 | 1.099 |
| H | - | - | - | - | - | - | - | - | 3.809 | 5.633 | 9.117 | 2.779 | 3.143 | 2.912 |
| I | - | - | - | - | - | - | - | - | - | 1.9 | 4.737 | 4.046 | 4.368 | 2.713 |
| J | - | - | - | - | - | - | - | - | - | - | 3.164 | 3.099 | 4.705 | 3.326 |
| K | - | - | - | - | - | - | - | - | - | - | - | 0.088 | 4.779 | 2.576 |
| L | - | - | - | - | - | - | - | - | - | - | - | - | 3.151 | 2.762 |
| M | - | - | - | - | - | - | - | - | - | - | - | - | - | 4.115 |
| N | - | - | - | - | - | - | - | - | - | - | - | - | - | - |
