## Supplementary Data S2 for "Interlaboratory comparison of standardised metabolomics and lipidomics analyses in human and rodent blood using the MxP® Quant 500 kit": PC ae C44.4.pdf

**Amount of relative differences between laboratories**

| Min | A | B | C | D | E | F | G | H | I | J | K | L | M | N |
|---|---|---|---|---|---|---|---|---|---|---|---|---|---|---|
| A | - | 5.544 | 5.735 | 1.119 | 2.5 | 5.023 | 3.382 | 4.097 | 5.506 | 3.48 | 3.69 | 3.169 | 4.512 | 4.911 |
| B | - | - | 1.348 | 6.442 | 6.365 | 3.858 | 6.635 | 4.028 | 0.771 | 6.644 | 6.619 | 6.584 | 5.173 | 5.251 |
| C | - | - | - | 6.22 | 6.136 | 3.267 | 6.432 | 3.193 | 0.223 | 6.499 | 6.39 | 6.373 | 4.792 | 4.972 |
| D | - | - | - | - | 3.012 | 4.978 | 3.755 | 3.912 | 5.596 | 3.889 | 3.993 | 3.566 | 4.398 | 4.867 |
| E | - | - | - | - | - | 5.261 | 1.252 | 3.964 | 5.875 | 1.391 | 2.588 | 0.408 | 4.879 | 5.445 |
| F | - | - | - | - | - | - | 6.063 | 2.929 | 3.492 | 6.122 | 6.072 | 6.002 | 4.032 | 3.967 |
| G | - | - | - | - | - | - | - | 4.423 | 5.737 | 0.325 | 1.691 | 0.301 | 4.91 | 5.258 |
| H | - | - | - | - | - | - | - | - | 3.877 | 5.881 | 5.82 | 5.745 | 3.046 | 2.957 |
| I | - | - | - | - | - | - | - | - | - | 6.573 | 6.472 | 6.423 | 4.922 | 5.058 |
| J | - | - | - | - | - | - | - | - | - | - | 0.77 | 1.467 | 4.913 | 5.246 |
| K | - | - | - | - | - | - | - | - | - | - | - | 2.017 | 5.01 | 5.358 |
| L | - | - | - | - | - | - | - | - | - | - | - | - | 4.891 | 5.266 |
| M | - | - | - | - | - | - | - | - | - | - | - | - | - | 1.23 |
| N | - | - | - | - | - | - | - | - | - | - | - | - | - | - |

| | |
|---|---|
| | Larger than 10 |
| | Between 8 and 9,99 |
| | Between 6 and 7,99 |
| | Between 4 and 5,99 |
| | Between 2 and 3,99 |
| | Between 0 and 1,99 |
| | NA (Calculation failed) |

| Median | A | B | C | D | E | F | G | H | I | J | K | L | M | N |
|---|---|---|---|---|---|---|---|---|---|---|---|---|---|---|
| A | - | 0.612 | 3.863 | 2.362 | 3.896 | -0.384 | 0.402 | 2.566 | 3.673 | 2.992 | -0.911 | 1.806 | 2.043 | 1.292 |
| B | - | - | 3.725 | 3.006 | 4.073 | 1.577 | 2.323 | 2.931 | 3.406 | 1.772 | 1.585 | 2.463 | 0.284 | 2.753 |
| C | - | - | - | 4.236 | 4.732 | 3.887 | 4.026 | 2.149 | 1.337 | 3.192 | 3.852 | 4.049 | 3.425 | 4.171 |
| D | - | - | - | - | 3.361 | 2.666 | 2.014 | 3.518 | 4.255 | 3.787 | 2.585 | 0.768 | 3.29 | 1.545 |
| E | - | - | - | - | - | 4.256 | 4.031 | 4.531 | 5.038 | 4.78 | 4.208 | 3.735 | 4.498 | 3.96 |
| F | - | - | - | - | - | - | 0.885 | 3.156 | 3.717 | 2.796 | -2.96 | 1.602 | 2.023 | 1.834 |
| G | - | - | - | - | - | - | - | 2.721 | 3.823 | 3.238 | 0.876 | 1.266 | 2.436 | 0.079 |
| H | - | - | - | - | - | - | - | - | 1.852 | 1.026 | 3.044 | 3.472 | 2.195 | 3.537 |
| I | - | - | - | - | - | - | - | - | - | 2.778 | 3.622 | 3.849 | 3.081 | 3.987 |
| J | - | - | - | - | - | - | - | - | - | - | 2.786 | 3.412 | 1.856 | 3.263 |
| K | - | - | - | - | - | - | - | - | - | - | - | 2.069 | 1.806 | 1.479 |
| L | - | - | - | - | - | - | - | - | - | - | - | - | 3.009 | 0.557 |
| M | - | - | - | - | - | - | - | - | - | - | - | - | - | 2.814 |
| N | - | - | - | - | - | - | - | - | - | - | - | - | - | - |

| Max | A | B | C | D | E | F | G | H | I | J | K | L | M | N |
|---|---|---|---|---|---|---|---|---|---|---|---|---|---|---|
| A | - | 4.503 | 5.732 | 2.482 | 4.19 | 3.082 | 1.58 | 4.516 | 5.371 | 2.919 | 0.436 | 2.436 | 3.544 | 0.829 |
| B | - | - | 4.271 | 4.48 | 5.049 | 2.814 | 4.311 | 0.3 | 3.904 | 3.083 | 4.204 | 4.396 | 3.096 | 3.764 |
| C | - | - | - | 5.04 | 5.408 | 4.354 | 4.949 | 3.851 | 1.676 | 4.419 | 4.879 | 4.991 | 4.426 | 4.676 |
| D | - | - | - | - | 3.732 | 3.917 | 1.43 | 4.953 | 5.708 | 3.774 | 2.167 | -2.739 | 4.203 | 2.968 |
| E | - | - | - | - | - | 5.018 | 4.197 | 5.69 | 6.283 | 4.933 | 4.349 | 3.921 | 5.216 | 4.621 |
| F | - | - | - | - | - | - | 3.538 | 3.302 | 4.652 | -0.367 | 3.336 | 3.735 | -0.688 | 2.701 |
| G | - | - | - | - | - | - | - | 4.768 | 5.565 | 3.385 | 0.699 | 1.37 | 3.94 | 2.211 |
| H | - | - | - | - | - | - | - | - | 3.656 | 3.254 | 4.179 | 4.393 | 3.031 | 3.865 |
| I | - | - | - | - | - | - | - | - | - | 4.28 | 4.792 | 4.895 | 4.272 | 4.547 |
| J | - | - | - | - | - | - | - | - | - | - | 3.037 | 3.567 | 1.926 | 2.396 |
| K | - | - | - | - | - | - | - | - | - | - | - | 2.077 | 3.755 | 1.61 |
| L | - | - | - | - | - | - | - | - | - | - | - | - | 4.218 | 2.921 |
| M | - | - | - | - | - | - | - | - | - | - | - | - | - | 2.954 |
| N | - | - | - | - | - | - | - | - | - | - | - | - | - | - |
