## Supplementary Data S2 for "Interlaboratory comparison of standardised metabolomics and lipidomics analyses in human and rodent blood using the MxP® Quant 500 kit": PC ae C44.5.pdf

**Amount of relative differences between laboratories**

| Min | A | B | C | D | E | F | G | H | I | J | K | L | M | N |
|---|---|---|---|---|---|---|---|---|---|---|---|---|---|---|
| A | - | 4.671 | 4.676 | 3.578 | 5.297 | 4.525 | 5.497 | 5.34 | 4.462 | 5.432 | 5.24 | 5.444 | 4.433 | 5.085 |
| B | - | - | 0.663 | 5.004 | 6.281 | 1.435 | 6.589 | 4.068 | 0.636 | 6.558 | 6.322 | 6.618 | 2.784 | 1.981 |
| C | - | - | - | 5.011 | 6.352 | 2.422 | 6.658 | 3.856 | 1.875 | 6.633 | 6.371 | 6.671 | 3.245 | 0.125 |
| D | - | - | - | - | 5.323 | 4.849 | 5.001 | 5.624 | 4.516 | 5.307 | 5.139 | 4.693 | 4.892 | 5.66 |
| E | - | - | - | - | - | 5.713 | 3.938 | 6.17 | 5.596 | 2.919 | 2.858 | 4.498 | 5.807 | 6.252 |
| F | - | - | - | - | - | - | 6.52 | 4.203 | 0.405 | 6.443 | 6.238 | 6.51 | 2.042 | 2.537 |
| G | - | - | - | - | - | - | - | 5.931 | 5.421 | 2.184 | -0.403 | 1.894 | 5.541 | 5.917 |
| H | - | - | - | - | - | - | - | - | 4.216 | 7.01 | 6.778 | 7.084 | 4.756 | 4.124 |
| I | - | - | - | - | - | - | - | - | - | 6.528 | 6.278 | 6.586 | 2.505 | 2.24 |
| J | - | - | - | - | - | - | - | - | - | - | 1.687 | 3.372 | 5.62 | 6.088 |
| K | - | - | - | - | - | - | - | - | - | - | - | 2.676 | 5.557 | 5.972 |
| L | - | - | - | - | - | - | - | - | - | - | - | - | 5.475 | 5.906 |
| M | - | - | - | - | - | - | - | - | - | - | - | - | - | 3.455 |
| N | - | - | - | - | - | - | - | - | - | - | - | - | - | - |

| | |
|---|---|
| | Larger than 10 |
| | Between 8 and 9,99 |
| | Between 6 and 7,99 |
| | Between 4 and 5,99 |
| | Between 2 and 3,99 |
| | Between 0 and 1,99 |
| | NA (Calculation failed) |

| Median | A | B | C | D | E | F | G | H | I | J | K | L | M | N |
|---|---|---|---|---|---|---|---|---|---|---|---|---|---|---|
| A | - | -0.189 | 0.126 | 2.062 | 3.99 | 0.123 | 2.021 | 1.371 | 2.115 | 1.846 | 0.711 | 1.22 | -0.211 | 1.781 |
| B | - | - | 1.067 | 2.225 | 3.918 | -2.063 | 2.299 | 0.853 | 2.33 | 1.502 | 1.349 | 0.655 | -4.282 | 1.389 |
| C | - | - | - | 1.274 | 4.077 | 1.212 | 1.453 | 1.947 | 1.488 | 2.282 | -1.227 | 1.858 | 1.094 | 2.215 |
| D | - | - | - | - | 4.25 | 2.318 | -3.55 | 2.712 | -1.69 | 2.884 | 1.182 | 2.645 | 2.273 | 2.904 |
| E | - | - | - | - | - | 4.144 | 4.577 | 4.002 | 4.615 | 3.883 | 4.382 | 4.031 | 4.172 | 3.901 |
| F | - | - | - | - | - | - | 2.371 | 0.656 | 2.409 | 1.367 | 1.48 | 0.402 | -2.393 | 1.261 |
| G | - | - | - | - | - | - | - | 2.67 | -1.362 | 2.881 | 1.253 | 2.608 | 2.24 | 2.85 |
| H | - | - | - | - | - | - | - | - | 2.779 | 0.058 | 2.17 | -2.152 | 0.848 | -0.318 |
| I | - | - | - | - | - | - | - | - | - | 2.9 | 1.219 | 2.648 | 2.267 | 2.858 |
| J | - | - | - | - | - | - | - | - | - | - | 2.421 | 0.46 | 1.506 | -2.459 |
| K | - | - | - | - | - | - | - | - | - | - | - | 1.922 | 1.26 | 2.302 |
| L | - | - | - | - | - | - | - | - | - | - | - | - | 0.548 | 0.239 |
| M | - | - | - | - | - | - | - | - | - | - | - | - | - | 1.366 |
| N | - | - | - | - | - | - | - | - | - | - | - | - | - | - |

| Max | A | B | C | D | E | F | G | H | I | J | K | L | M | N |
|---|---|---|---|---|---|---|---|---|---|---|---|---|---|---|
| A | - | -0.216 | 1.696 | 1.903 | 4.166 | -1.456 | 1.436 | -5.612 | 2.636 | 2.334 | -2.306 | 1.839 | -1.418 | 0.855 |
| B | - | - | 1.258 | 1.655 | 4.216 | -1.203 | 0.939 | -0.334 | 2.399 | 2.544 | -0.928 | 2.102 | -1.102 | 1.35 |
| C | - | - | - | -0.405 | 4.362 | 1.498 | -1.021 | 1.649 | 1.456 | 3.015 | 1.506 | 2.7 | 1.5 | 2.273 |
| D | - | - | - | - | 4.409 | 1.768 | 0.19 | 1.871 | 0.977 | 3.143 | 1.91 | 2.834 | 1.743 | 2.374 |
| E | - | - | - | - | - | 4.481 | 4.652 | 4.458 | 4.893 | 3.949 | 4.457 | 4.122 | 4.488 | 4.308 |
| F | - | - | - | - | - | - | 1.217 | -1.199 | 2.532 | 2.42 | -3.404 | 1.941 | -4.496 | 1.147 |
| G | - | - | - | - | - | - | - | 1.416 | 1.799 | 2.951 | 1.327 | 2.624 | 1.154 | 2.128 |
| H | - | - | - | - | - | - | - | - | 2.592 | 2.339 | -1.632 | 1.793 | -1.764 | 0.828 |
| I | - | - | - | - | - | - | - | - | - | 3.399 | 2.459 | 3.179 | 2.443 | 2.885 |
| J | - | - | - | - | - | - | - | - | - | - | 2.51 | 0.761 | 2.572 | 1.934 |
| K | - | - | - | - | - | - | - | - | - | - | - | 1.895 | -1.867 | 0.884 |
| L | - | - | - | - | - | - | - | - | - | - | - | - | 2.073 | 0.968 |
| M | - | - | - | - | - | - | - | - | - | - | - | - | - | 1.091 |
| N | - | - | - | - | - | - | - | - | - | - | - | - | - | - |
