## Supplementary Data S2 for "Interlaboratory comparison of standardised metabolomics and lipidomics analyses in human and rodent blood using the MxP® Quant 500 kit": PC ae C44.6.pdf

**Amount of relative differences between laboratories**

| Min | A | B | C | D | E | F | G | H | I | J | K | L | M | N |
|---|---|---|---|---|---|---|---|---|---|---|---|---|---|---|
| A | - | 2.976 | 4.959 | 2.718 | 4.444 | 3.109 | 5.071 | 1.554 | 3.496 | 4.496 | 5.103 | 5.271 | 2.728 | 4.372 |
| B | - | - | 4.629 | 1.306 | 4.02 | -1.132 | 4.7 | 3.408 | 4.256 | 4.13 | 4.789 | 4.886 | -0.024 | 4.903 |
| C | - | - | - | 4.444 | 2.987 | 4.486 | 2.477 | 4.948 | 5.185 | 3.441 | 0.363 | -7 | 4.535 | 5.558 |
| D | - | - | - | - | 4.316 | 0.057 | 4.57 | 3.585 | 4.254 | 3.898 | 4.963 | 4.873 | 0.176 | 5.143 |
| E | - | - | - | - | - | 4.315 | 2.146 | 4.904 | 5.147 | 0.157 | 2.692 | 2.403 | 4.292 | 5.657 |
| F | - | - | - | - | - | - | 4.577 | 3.424 | 4.2 | 3.877 | 4.678 | 4.822 | 0.597 | 4.839 |
| G | - | - | - | - | - | - | - | 5.048 | 5.286 | 2.641 | 2.247 | -2.275 | 4.596 | 5.764 |
| H | - | - | - | - | - | - | - | - | 3.099 | 4.686 | 5.3 | 5.426 | 3.314 | 4.228 |
| I | - | - | - | - | - | - | - | - | - | 5.169 | 5.705 | 5.814 | 4.326 | 3.401 |
| J | - | - | - | - | - | - | - | - | - | - | 3.614 | 2.891 | 4.128 | 5.667 |
| K | - | - | - | - | - | - | - | - | - | - | - | 0.443 | 4.609 | 5.64 |
| L | - | - | - | - | - | - | - | - | - | - | - | - | 4.549 | 5.613 |
| M | - | - | - | - | - | - | - | - | - | - | - | - | - | 4.788 |
| N | - | - | - | - | - | - | - | - | - | - | - | - | - | - |

| | |
|---|---|
| | Larger than 10 |
| | Between 8 and 9,99 |
| | Between 6 and 7,99 |
| | Between 4 and 5,99 |
| | Between 2 and 3,99 |
| | Between 0 and 1,99 |
| | NA (Calculation failed) |

| Median | A | B | C | D | E | F | G | H | I | J | K | L | M | N |
|---|---|---|---|---|---|---|---|---|---|---|---|---|---|---|
| A | - | 2.875 | 2.412 | 1.599 | 4.035 | 0.076 | 2.159 | 2.023 | 2.759 | -4.3 | -3.925 | -0.945 | 1.632 | 3.161 |
| B | - | - | 3.769 | 2.239 | 3.265 | 2.804 | 3.661 | 1.826 | -0.863 | 2.981 | 3.005 | 2.905 | 2.2 | 0.694 |
| C | - | - | - | 3.032 | 4.349 | 2.52 | 0.428 | 3.149 | 3.524 | 2.444 | 2.299 | 2.421 | 2.992 | 3.738 |
| D | - | - | - | - | 3.73 | 1.107 | 2.929 | 0.147 | 1.969 | 1.607 | 1.861 | 1.471 | -6.872 | 2.686 |
| E | - | - | - | - | - | 4.202 | 4.582 | 3.833 | 3.446 | 4.235 | 4.262 | 4.233 | 3.949 | 3.131 |
| F | - | - | - | - | - | - | 2.42 | 1.652 | 2.561 | -0.215 | 0.049 | -1.011 | 1.116 | 3.002 |
| G | - | - | - | - | - | - | - | 3.04 | 3.448 | 2.063 | 1.924 | 2.178 | 2.848 | 3.708 |
| H | - | - | - | - | - | - | - | - | 1.497 | 2.02 | 2.112 | 1.889 | 0.009 | 2.346 |
| I | - | - | - | - | - | - | - | - | - | 2.81 | 2.878 | 2.734 | 1.97 | 1.265 |
| J | - | - | - | - | - | - | - | - | - | - | -1.212 | -1.518 | 1.626 | 3.217 |
| K | - | - | - | - | - | - | - | - | - | - | - | -1.289 | 1.667 | 3.14 |
| L | - | - | - | - | - | - | - | - | - | - | - | - | 1.473 | 3.078 |
| M | - | - | - | - | - | - | - | - | - | - | - | - | - | 2.577 |
| N | - | - | - | - | - | - | - | - | - | - | - | - | - | - |

| Max | A | B | C | D | E | F | G | H | I | J | K | L | M | N |
|---|---|---|---|---|---|---|---|---|---|---|---|---|---|---|
| A | - | 3.201 | 0.649 | 2.249 | 4.359 | 1.159 | -1.526 | 2.079 | 2.589 | 1.727 | 1.977 | 2.13 | 2.148 | 2.877 |
| B | - | - | 3.587 | 2.358 | 3.633 | 2.96 | 3.428 | 2.477 | 1.934 | 2.703 | 2.521 | 2.409 | 2.422 | 1.154 |
| C | - | - | - | 2.521 | 4.441 | 1.89 | -0.416 | 2.52 | 2.881 | 2.22 | 2.497 | 2.557 | 2.553 | 3.129 |
| D | - | - | - | - | 4.075 | 1.347 | 2.415 | -1.163 | 0.479 | 0.418 | -1.26 | -2.693 | -2.603 | 1.329 |
| E | - | - | - | - | - | 4.528 | 4.739 | 4.361 | 4.217 | 4.44 | 4.37 | 4.332 | 4.34 | 4.071 |
| F | - | - | - | - | - | - | 1.451 | 1.044 | 1.936 | 0.178 | 0.873 | 1.167 | 1.174 | 2.381 |
| G | - | - | - | - | - | - | - | 2.226 | 2.64 | 1.957 | 2.26 | 2.308 | 2.284 | 2.9 |
| H | - | - | - | - | - | - | - | - | 0.901 | -0.107 | -2.134 | -2.536 | -2.191 | 1.691 |
| I | - | - | - | - | - | - | - | - | - | 1.504 | 1.086 | 0.817 | 0.773 | 0.374 |
| J | - | - | - | - | - | - | - | - | - | - | -0.079 | 0.157 | 0.157 | 1.872 |
| K | - | - | - | - | - | - | - | - | - | - | - | -1.664 | -1.9 | 1.698 |
| L | - | - | - | - | - | - | - | - | - | - | - | - | -4.249 | 1.559 |
| M | - | - | - | - | - | - | - | - | - | - | - | - | - | 1.598 |
| N | - | - | - | - | - | - | - | - | - | - | - | - | - | - |
