## Supplementary Data S2 for "Interlaboratory comparison of standardised metabolomics and lipidomics analyses in human and rodent blood using the MxP® Quant 500 kit": Sarcosine.pdf

**Amount of relative differences between laboratories**

| Min | A | B | C | D | E | F | G | H | I | J | K | L | M | N |
|---|---|---|---|---|---|---|---|---|---|---|---|---|---|---|
| A | - | -1.799 | 0.717 | 3.247 | 3.897 | 2.033 | 1.273 | 5.047 | 3.856 | 3.737 | 2.099 | 4.261 | 2.277 | 3.658 |
| B | - | - | 0.593 | 3.256 | 4.239 | 1.98 | 1.162 | 5.1 | 3.739 | 3.751 | 2.149 | 4.09 | 2.301 | 3.692 |
| C | - | - | - | 3.423 | 4.073 | 2.482 | 0.314 | 5.104 | 3.605 | 3.884 | 1.569 | 4.454 | 2.671 | 3.81 |
| D | - | - | - | - | 4.874 | 2.506 | 3.751 | 4.762 | -5.426 | 2.294 | 3.922 | 5.138 | 2.276 | 1.816 |
| E | - | - | - | - | - | 4.035 | 3.454 | 5.581 | 4.383 | 4.763 | 3.561 | 1.87 | 4.111 | 4.641 |
| F | - | - | - | - | - | - | 2.773 | 4.913 | 3.685 | 3.291 | 3.071 | 4.724 | -0.424 | 3.132 |
| G | - | - | - | - | - | - | - | 5.177 | 3.582 | 3.988 | 0.697 | 4.013 | 2.87 | 3.887 |
| H | - | - | - | - | - | - | - | - | 5.291 | 4.695 | 5.792 | 6.502 | 5.315 | 4.893 |
| I | - | - | - | - | - | - | - | - | - | 1.423 | 3.879 | 5.194 | 2.806 | 1.512 |
| J | - | - | - | - | - | - | - | - | - | - | 4.408 | 5.45 | 3.328 | 0.561 |
| K | - | - | - | - | - | - | - | - | - | - | - | 3.032 | 3.095 | 4.051 |
| L | - | - | - | - | - | - | - | - | - | - | - | - | 4.457 | 4.83 |
| M | - | - | - | - | - | - | - | - | - | - | - | - | - | 3.013 |
| N | - | - | - | - | - | - | - | - | - | - | - | - | - | - |

| | |
|---|---|
| | Larger than 10 |
| | Between 8 and 9,99 |
| | Between 6 and 7,99 |
| | Between 4 and 5,99 |
| | Between 2 and 3,99 |
| | Between 0 and 1,99 |
| | NA (Calculation failed) |

| Median | A | B | C | D | E | F | G | H | I | J | K | L | M | N |
|---|---|---|---|---|---|---|---|---|---|---|---|---|---|---|
| A | - | 2.619 | 0.024 | 1.647 | 4.202 | 2.534 | 1.437 | 3.708 | 1.385 | 2.976 | 1.207 | 4.596 | 2.711 | 1.794 |
| B | - | - | 2.977 | 1.721 | 4.701 | -0.615 | 3.295 | 2.845 | 1.841 | 0.626 | 2.004 | 5.02 | -2.338 | 1.622 |
| C | - | - | - | 2.044 | 4.107 | 2.752 | 0.726 | 3.801 | 1.822 | 3.135 | 1.73 | 4.512 | 2.904 | 2.151 |
| D | - | - | - | - | 4.499 | 1.563 | 2.51 | 3.361 | 0 | 2.343 | 0.036 | 4.816 | 1.876 | -1.051 |
| E | - | - | - | - | - | 4.367 | 3.752 | 4.723 | 4.139 | 4.473 | 4.132 | 2.252 | 4.408 | 4.216 |
| F | - | - | - | - | - | - | 3.176 | 2.956 | 1.671 | 1.133 | 1.846 | 4.986 | -0.289 | 1.286 |
| G | - | - | - | - | - | - | - | 3.934 | 2.334 | 3.36 | 2.283 | 4.388 | 3.169 | 2.585 |
| H | - | - | - | - | - | - | - | - | 3.827 | 2.458 | 3.761 | 5.433 | 2.85 | 3.432 |
| I | - | - | - | - | - | - | - | - | - | 2.641 | -9.497 | 4.744 | 2.264 | 0.656 |
| J | - | - | - | - | - | - | - | - | - | - | 2.681 | 5.128 | 0.57 | 2.273 |
| K | - | - | - | - | - | - | - | - | - | - | - | 4.777 | 2.018 | -0.178 |
| L | - | - | - | - | - | - | - | - | - | - | - | - | 4.629 | 4.476 |
| M | - | - | - | - | - | - | - | - | - | - | - | - | - | 1.715 |
| N | - | - | - | - | - | - | - | - | - | - | - | - | - | - |

| Max | A | B | C | D | E | F | G | H | I | J | K | L | M | N |
|---|---|---|---|---|---|---|---|---|---|---|---|---|---|---|
| A | - | 3.029 | -0.261 | -0.23 | 4.299 | 2.645 | 1.476 | 2.562 | 0.522 | 2.624 | 2.123 | 4.676 | 2.813 | -0.859 |
| B | - | - | 3.28 | 2.982 | 4.839 | 0.928 | 3.55 | 1.341 | 3.226 | 1.12 | 1.957 | 5.196 | 0.191 | 3.031 |
| C | - | - | - | 0.815 | 4.12 | 2.818 | 0.813 | 2.726 | -0.751 | 2.791 | 2.398 | 4.528 | 2.964 | 0.475 |
| D | - | - | - | - | 4.334 | 2.406 | 1.907 | 2.082 | 0.432 | 2.358 | 1.75 | 4.713 | 2.605 | -1.045 |
| E | - | - | - | - | - | 4.445 | 3.818 | 4.227 | 4.045 | 4.37 | 4.272 | 2.341 | 4.482 | 4.049 |
| F | - | - | - | - | - | - | 3.261 | -1.995 | 3.114 | -2.502 | 1.072 | 5.05 | -0.253 | 2.586 |
| G | - | - | - | - | - | - | - | 2.964 | 1.42 | 3.09 | 2.783 | 4.475 | 3.244 | 1.68 |
| H | - | - | - | - | - | - | - | - | 2.507 | -0.206 | -0.841 | 4.917 | 1.014 | 2.254 |
| I | - | - | - | - | - | - | - | - | - | 2.877 | 2.192 | 4.592 | 3.026 | 0.226 |
| J | - | - | - | - | - | - | - | - | - | - | 0.857 | 5.026 | -0.09 | 2.561 |
| K | - | - | - | - | - | - | - | - | - | - | - | 5.031 | 1.474 | 2.04 |
| L | - | - | - | - | - | - | - | - | - | - | - | - | 4.675 | 4.343 |
| M | - | - | - | - | - | - | - | - | - | - | - | - | - | 2.786 |
| N | - | - | - | - | - | - | - | - | - | - | - | - | - | - |
