## Supplementary Data S2 for "Interlaboratory comparison of standardised metabolomics and lipidomics analyses in human and rodent blood using the MxP® Quant 500 kit": SM (OH) C14.1.pdf

**Amount of relative differences between laboratories**

| Min | A | B | C | D | E | F | G | H | I | J | K | L | M | N |
|---|---|---|---|---|---|---|---|---|---|---|---|---|---|---|
| A | - | 4.981 | 4.993 | 3.201 | 4.301 | 3.258 | 2.994 | 3.054 | -2.402 | 6.3 | 3.593 | 3.422 | 4.518 | 5.324 |
| B | - | - | 0.126 | 4.656 | 5.579 | 4.7 | 5.045 | 5.456 | 5.353 | 5.684 | 4.002 | 5.697 | 3.472 | 2.013 |
| C | - | - | - | 4.658 | 5.395 | 4.785 | 5.047 | 5.438 | 5.15 | 5.763 | 4.18 | 5.698 | 3.702 | 0.825 |
| D | - | - | - | - | 4.579 | 0.698 | 1.905 | 4.295 | 3.662 | 6.194 | 2.068 | 4.488 | 3.57 | 4.82 |
| E | - | - | - | - | - | 5.049 | 4.746 | 3.332 | 3.968 | 6.923 | 5.204 | 2.628 | 5.584 | 6.15 |
| F | - | - | - | - | - | - | 1.582 | 4.219 | 3.596 | 6.377 | 2.885 | 4.583 | 3.907 | 5.032 |
| G | - | - | - | - | - | - | - | 3.983 | 2.678 | 6.331 | 3.154 | 4.265 | 4.017 | 5.039 |
| H | - | - | - | - | - | - | - | - | 3.143 | 6.741 | 4.724 | 1.507 | 5.186 | 5.78 |
| I | - | - | - | - | - | - | - | - | - | 6.529 | 4.054 | 3.857 | 4.656 | 5.396 |
| J | - | - | - | - | - | - | - | - | - | - | 5.837 | 6.615 | 6.006 | 5.617 |
| K | - | - | - | - | - | - | - | - | - | - | - | 4.958 | 2.884 | 4.591 |
| L | - | - | - | - | - | - | - | - | - | - | - | - | 5.267 | 5.825 |
| M | - | - | - | - | - | - | - | - | - | - | - | - | - | 3.979 |
| N | - | - | - | - | - | - | - | - | - | - | - | - | - | - |

| Median | A | B | C | D | E | F | G | H | I | J | K | L | M | N |
|---|---|---|---|---|---|---|---|---|---|---|---|---|---|---|
| A | - | 3.545 | 3.096 | -0.263 | 4.372 | 3.597 | 2.366 | 3.756 | 2.907 | 3.538 | 3.079 | 3.458 | 2.71 | 3.211 |
| B | - | - | 1.827 | 3.627 | 3.347 | -1.244 | 2.902 | 1.028 | 2.269 | -2.568 | 1.882 | -0.324 | 2.573 | 1.459 |
| C | - | - | - | 3.087 | 3.736 | 1.934 | 1.924 | 2.433 | 0.23 | 1.736 | -2.742 | 1.404 | 1.221 | -0.4 |
| D | - | - | - | - | 4.328 | 3.504 | 2.118 | 3.674 | 2.743 | 3.447 | 2.943 | 3.356 | 2.517 | 3.087 |
| E | - | - | - | - | - | 3.447 | 4.301 | 3.19 | 4.067 | 3.524 | 3.954 | 3.622 | 4.165 | 3.856 |
| F | - | - | - | - | - | - | 2.982 | 0.698 | 2.389 | -0.956 | 2.044 | 0.31 | 2.669 | 1.68 |
| G | - | - | - | - | - | - | - | 3.14 | 1.318 | 2.779 | 1.81 | 2.624 | 0.528 | 2.117 |
| H | - | - | - | - | - | - | - | - | 2.798 | 1.088 | 2.542 | 1.541 | 3.011 | 2.296 |
| I | - | - | - | - | - | - | - | - | - | 2.176 | 0.08 | 1.906 | 0.063 | 0.928 |
| J | - | - | - | - | - | - | - | - | - | - | 1.774 | -1.049 | 2.547 | 1.309 |
| K | - | - | - | - | - | - | - | - | - | - | - | 1.47 | 1.101 | -0.165 |
| L | - | - | - | - | - | - | - | - | - | - | - | - | 2.317 | 0.957 |
| M | - | - | - | - | - | - | - | - | - | - | - | - | - | 1.577 |
| N | - | - | - | - | - | - | - | - | - | - | - | - | - | - |

| Max | A | B | C | D | E | F | G | H | I | J | K | L | M | N |
|---|---|---|---|---|---|---|---|---|---|---|---|---|---|---|
| A | - | 3.621 | 3.204 | -0.033 | 4.373 | 3.631 | 2.416 | 3.764 | 2.928 | 3.657 | 3.131 | 3.458 | 2.783 | 3.322 |
| B | - | - | 1.843 | 3.688 | 3.277 | -3.626 | 3.006 | 0.518 | 2.464 | -1.327 | 1.994 | 0.638 | 2.62 | 1.449 |
| C | - | - | - | 3.167 | 3.696 | 1.809 | 2.105 | 2.277 | 0.792 | 1.989 | -1.237 | 0.954 | 1.326 | -0.33 |
| D | - | - | - | - | 4.325 | 3.52 | 2.121 | 3.663 | 2.723 | 3.565 | 2.976 | 3.331 | 2.567 | 3.177 |
| E | - | - | - | - | - | 3.407 | 4.293 | 3.188 | 4.068 | 3.363 | 3.924 | 3.632 | 4.137 | 3.769 |
| F | - | - | - | - | - | - | 3.009 | 0.46 | 2.46 | -1.326 | 2.027 | 0.68 | 2.646 | 1.465 |
| G | - | - | - | - | - | - | - | 3.123 | 1.283 | 2.969 | 1.885 | 2.58 | 0.718 | 2.292 |
| H | - | - | - | - | - | - | - | - | 2.792 | -0.052 | 2.463 | 1.581 | 2.951 | 2.057 |
| I | - | - | - | - | - | - | - | - | - | 2.483 | 0.403 | 1.846 | -0.382 | 1.358 |
| J | - | - | - | - | - | - | - | - | - | - | 2.115 | 0.926 | 2.733 | 1.701 |
| K | - | - | - | - | - | - | - | - | - | - | - | 1.24 | 1.06 | 0.341 |
| L | - | - | - | - | - | - | - | - | - | - | - | - | 2.201 | 0.164 |
| M | - | - | - | - | - | - | - | - | - | - | - | - | - | 1.707 |
| N | - | - | - | - | - | - | - | - | - | - | - | - | - | - |

| | |
|---|---|
| | Larger than 10 |
| | Between 8 and 9,99 |
| | Between 6 and 7,99 |
| | Between 4 and 5,99 |
| | Between 2 and 3,99 |
| | Between 0 and 1,99 |
| | NA (Calculation failed) |
