## Supplementary Data S2 for "Interlaboratory comparison of standardised metabolomics and lipidomics analyses in human and rodent blood using the MxP® Quant 500 kit": SM (OH) C16.1.pdf

**Amount of relative differences between laboratories**

| Min | A | B | C | D | E | F | G | H | I | J | K | L | M | N |
|---|---|---|---|---|---|---|---|---|---|---|---|---|---|---|
| A | - | 2.677 | 3.679 | 2.796 | 3.764 | 0.77 | 3.502 | 3.623 | 3.632 | 5.377 | 1.491 | 3.399 | 2.29 | 1.152 |
| B | - | - | 4.448 | 3.859 | 2.425 | 2.909 | 4.147 | 2.885 | 4.181 | 5.806 | 3.327 | 2.869 | 3.43 | 2.885 |
| C | - | - | - | 2.895 | 4.535 | 3.65 | 2.476 | 4.649 | 1.661 | 4.869 | 3.465 | 4.819 | 3.218 | 3.765 |
| D | - | - | - | - | 4.144 | 2.44 | 1.872 | 4.233 | 2.033 | 5.128 | 1.675 | 4.172 | 1.173 | 2.376 |
| E | - | - | - | - | - | 4.045 | 4.658 | -1.121 | 4.633 | 6.017 | 4.13 | -3.794 | 4.328 | 4.017 |
| F | - | - | - | - | - | - | 3.255 | 3.818 | 3.362 | 5.497 | 1.179 | 3.858 | 1.74 | -2.927 |
| G | - | - | - | - | - | - | - | 4.425 | 0.592 | 5.065 | 2.616 | 4.466 | 2.432 | 3.136 |
| H | - | - | - | - | - | - | - | - | 4.735 | 5.986 | 4.138 | 1.252 | 4.284 | 4.066 |
| I | - | - | - | - | - | - | - | - | - | 5.069 | 2.847 | 4.614 | 2.415 | 3.278 |
| J | - | - | - | - | - | - | - | - | - | - | 5.135 | 5.791 | 5.184 | 5.384 |
| K | - | - | - | - | - | - | - | - | - | - | - | 3.99 | 0.598 | 0.912 |
| L | - | - | - | - | - | - | - | - | - | - | - | - | 4.075 | 3.822 |
| M | - | - | - | - | - | - | - | - | - | - | - | - | - | 1.603 |
| N | - | - | - | - | - | - | - | - | - | - | - | - | - | - |

| Median | A | B | C | D | E | F | G | H | I | J | K | L | M | N |
|---|---|---|---|---|---|---|---|---|---|---|---|---|---|---|
| A | - | 3.748 | 2.237 | -0.174 | 4.251 | 3.203 | 1.298 | 3.408 | 2.004 | 2.97 | 2.322 | 3.071 | 2.137 | 3.058 |
| B | - | - | 3.324 | 4.043 | 2.71 | 2.28 | 3.661 | 1.704 | 3.44 | 2.675 | 3.277 | 2.535 | 3.385 | 2.562 |
| C | - | - | - | 2.557 | 3.909 | 2.23 | 1.271 | 2.612 | -0.454 | 1.711 | -1.843 | 1.93 | -1.281 | 1.888 |
| D | - | - | - | - | 4.305 | 3.321 | 1.725 | 3.507 | 2.277 | 3.114 | 2.547 | 3.2 | 2.381 | 3.187 |
| E | - | - | - | - | - | 3.602 | 4.359 | 3.385 | 4.218 | 3.791 | 4.117 | 3.718 | 4.179 | 3.729 |
| F | - | - | - | - | - | - | 2.894 | 0.598 | 2.513 | 0.56 | 2.207 | -0.155 | 2.415 | 0 |
| G | - | - | - | - | - | - | - | 3.057 | 0.695 | 2.472 | 1.395 | 2.605 | 0.964 | 2.58 |
| H | - | - | - | - | - | - | - | - | 2.891 | 1.616 | 2.65 | 1.31 | 2.797 | 1.351 |
| I | - | - | - | - | - | - | - | - | - | 2.003 | 0.039 | 2.177 | -1.486 | 2.148 |
| J | - | - | - | - | - | - | - | - | - | - | 1.594 | -0.711 | 1.952 | -1.375 |
| K | - | - | - | - | - | - | - | - | - | - | - | 1.829 | -0.579 | 1.792 |
| L | - | - | - | - | - | - | - | - | - | - | - | - | 2.124 | -3.914 |
| M | - | - | - | - | - | - | - | - | - | - | - | - | - | 2.04 |
| N | - | - | - | - | - | - | - | - | - | - | - | - | - | - |

| Max | A | B | C | D | E | F | G | H | I | J | K | L | M | N |
|---|---|---|---|---|---|---|---|---|---|---|---|---|---|---|
| A | - | 3.764 | 2.395 | -0.51 | 4.26 | 3.252 | 1.504 | 3.401 | 2.189 | 3.171 | 2.391 | 3.061 | 2.213 | 3.104 |
| B | - | - | 3.27 | 4.048 | 2.716 | 2.257 | 3.645 | 1.838 | 3.408 | 2.382 | 3.276 | 2.617 | 3.384 | 2.553 |
| C | - | - | - | 2.653 | 3.891 | 2.158 | 1.396 | 2.473 | -0.217 | 2.016 | -4.064 | 1.668 | -0.598 | 1.786 |
| D | - | - | - | - | 4.309 | 3.34 | 1.806 | 3.478 | 2.363 | 3.277 | 2.568 | 3.164 | 2.402 | 3.203 |
| E | - | - | - | - | - | 3.587 | 4.35 | 3.429 | 4.202 | 3.659 | 4.116 | 3.754 | 4.175 | 3.721 |
| F | - | - | - | - | - | - | 2.882 | 0.188 | 2.474 | -1.187 | 2.23 | 0.374 | 2.429 | 0.031 |
| G | - | - | - | - | - | - | - | 2.988 | 0.785 | 2.695 | 1.333 | 2.503 | 0.898 | 2.562 |
| H | - | - | - | - | - | - | - | - | 2.761 | 0.71 | 2.566 | 1.312 | 2.729 | 1.13 |
| I | - | - | - | - | - | - | - | - | - | 2.293 | -0.263 | 2.002 | -2.123 | 2.103 |
| J | - | - | - | - | - | - | - | - | - | - | 2.077 | -0.313 | 2.293 | -0.579 |
| K | - | - | - | - | - | - | - | - | - | - | - | 1.694 | -0.626 | 1.809 |
| L | - | - | - | - | - | - | - | - | - | - | - | - | 2.014 | -1.773 |
| M | - | - | - | - | - | - | - | - | - | - | - | - | - | 2.05 |
| N | - | - | - | - | - | - | - | - | - | - | - | - | - | - |

| | |
|---|---|
| | Larger than 10 |
| | Between 8 and 9,99 |
| | Between 6 and 7,99 |
| | Between 4 and 5,99 |
| | Between 2 and 3,99 |
| | Between 0 and 1,99 |
| | NA (Calculation failed) |
