## Supplementary Data S2 for "Interlaboratory comparison of standardised metabolomics and lipidomics analyses in human and rodent blood using the MxP® Quant 500 kit": SM C18.0.pdf

**Amount of relative differences between laboratories**

| Min | A | B | C | D | E | F | G | H | I | J | K | L | M | N |
|---|---|---|---|---|---|---|---|---|---|---|---|---|---|---|
| A | - | 2.583 | 3.474 | 4.148 | 3.31 | 2.647 | 3.475 | 1.392 | 3.604 | 5.419 | 0.059 | 2.736 | 3.299 | 2.719 |
| B | - | - | 4.217 | 4.747 | 1.398 | -0.866 | 4.116 | 1.859 | 4.287 | 5.84 | 3.111 | -0.145 | 4.109 | 3.592 |
| C | - | - | - | 2.584 | 3.741 | 4.287 | 1.656 | 3.825 | -0.072 | 5.238 | 3.307 | 4.281 | 1.318 | 3.074 |
| D | - | - | - | - | 4.331 | 4.551 | 3.161 | 4.289 | 2.835 | 4.734 | 3.914 | 4.607 | 3.006 | 3.835 |
| E | - | - | - | - | - | 0.947 | 3.919 | 3.084 | 3.818 | 5.834 | 2.867 | -3.847 | 4.055 | 3.108 |
| F | - | - | - | - | - | - | 4.238 | 2.102 | 4.252 | 5.858 | 3.07 | -2.189 | 4.138 | 3.799 |
| G | - | - | - | - | - | - | - | 3.519 | 1.307 | 5.195 | 2.805 | 4.011 | -1.392 | 2.34 |
| H | - | - | - | - | - | - | - | - | 3.577 | 5.706 | 2.21 | 2.308 | 3.797 | 3.065 |
| I | - | - | - | - | - | - | - | - | - | 5.244 | 3.304 | 4.364 | 0.006 | 3.025 |
| J | - | - | - | - | - | - | - | - | - | - | 5.19 | 5.53 | 5.155 | 5.42 |
| K | - | - | - | - | - | - | - | - | - | - | - | 3.217 | 3.065 | 1.269 |
| L | - | - | - | - | - | - | - | - | - | - | - | - | 4.155 | 3.643 |
| M | - | - | - | - | - | - | - | - | - | - | - | - | - | 2.317 |
| N | - | - | - | - | - | - | - | - | - | - | - | - | - | - |

| | |
|---|---|
| | Larger than 10 |
| | Between 8 and 9,99 |
| | Between 6 and 7,99 |
| | Between 4 and 5,99 |
| | Between 2 and 3,99 |
| | Between 0 and 1,99 |
| | NA (Calculation failed) |

| Median | A | B | C | D | E | F | G | H | I | J | K | L | M | N |
|---|---|---|---|---|---|---|---|---|---|---|---|---|---|---|
| A | - | 2.898 | 2.086 | 2.146 | 4.013 | 2.517 | -0.713 | 3.285 | 1.822 | 2.84 | 1.763 | 2.344 | -1.672 | 1.908 |
| B | - | - | 1.793 | 3.68 | 3.244 | 0.907 | 3.125 | 1.308 | 2.081 | -1.956 | 2.129 | 1.36 | 2.948 | 2.006 |
| C | - | - | - | 3.183 | 3.647 | 0.585 | 2.347 | 2.518 | -0.498 | 1.633 | -0.141 | -0.183 | 2.054 | -0.831 |
| D | - | - | - | - | 4.306 | 3.279 | 1.868 | 3.762 | 2.934 | 3.477 | 2.905 | 3.189 | 2.176 | 2.963 |
| E | - | - | - | - | - | 3.641 | 4.329 | 2.933 | 3.922 | 3.432 | 3.936 | 3.726 | 4.248 | 3.899 |
| F | - | - | - | - | - | - | 2.747 | 2.096 | 1.219 | 0.679 | 1.3 | -0.546 | 2.519 | 1.061 |
| G | - | - | - | - | - | - | - | 3.364 | 2.052 | 2.959 | 1.998 | 2.504 | -0.135 | 2.11 |
| H | - | - | - | - | - | - | - | - | 2.787 | 1.503 | 2.814 | 2.367 | 3.386 | 2.734 |
| I | - | - | - | - | - | - | - | - | - | 1.926 | -2.436 | 0.654 | 1.759 | -2.646 |
| J | - | - | - | - | - | - | - | - | - | - | 2.002 | 1.053 | 2.9 | 1.89 |
| K | - | - | - | - | - | - | - | - | - | - | - | 0.8 | 1.678 | -1.57 |
| L | - | - | - | - | - | - | - | - | - | - | - | - | 2.324 | 0.497 |
| M | - | - | - | - | - | - | - | - | - | - | - | - | - | 1.785 |
| N | - | - | - | - | - | - | - | - | - | - | - | - | - | - |

| Max | A | B | C | D | E | F | G | H | I | J | K | L | M | N |
|---|---|---|---|---|---|---|---|---|---|---|---|---|---|---|
| A | - | 2.901 | 2.137 | 2.106 | 4.019 | 2.516 | -0.95 | 3.292 | 1.897 | 2.896 | 1.778 | 2.34 | -1.36 | 1.935 |
| B | - | - | 1.728 | 3.666 | 3.264 | 0.919 | 3.113 | 1.333 | 2.017 | -4 | 2.117 | 1.376 | 2.934 | 1.987 |
| C | - | - | - | 3.188 | 3.645 | 0.473 | 2.368 | 2.503 | -0.504 | 1.736 | -0.004 | -0.457 | 2.072 | -0.684 |
| D | - | - | - | - | 4.306 | 3.266 | 1.85 | 3.758 | 2.935 | 3.505 | 2.893 | 3.169 | 2.168 | 2.955 |
| E | - | - | - | - | - | 3.648 | 4.332 | 2.932 | 3.923 | 3.396 | 3.942 | 3.738 | 4.249 | 3.903 |
| F | - | - | - | - | - | - | 2.724 | 2.115 | 1.09 | 0.953 | 1.271 | -0.53 | 2.496 | 1.005 |
| G | - | - | - | - | - | - | - | 3.363 | 2.079 | 3.004 | 1.989 | 2.482 | -0.12 | 2.108 |
| H | - | - | - | - | - | - | - | - | 2.775 | 1.354 | 2.818 | 2.39 | 3.383 | 2.732 |
| I | - | - | - | - | - | - | - | - | - | 2.012 | -1.895 | 0.503 | 1.773 | -3.236 |
| J | - | - | - | - | - | - | - | - | - | - | 2.119 | 1.305 | 2.96 | 2.007 |
| K | - | - | - | - | - | - | - | - | - | - | - | 0.748 | 1.661 | -1.493 |
| L | - | - | - | - | - | - | - | - | - | - | - | - | 2.296 | 0.419 |
| M | - | - | - | - | - | - | - | - | - | - | - | - | - | 1.781 |
| N | - | - | - | - | - | - | - | - | - | - | - | - | - | - |
