## Supplementary Data S2 for "Interlaboratory comparison of standardised metabolomics and lipidomics analyses in human and rodent blood using the MxP® Quant 500 kit": SM C18.1.pdf

**Amount of relative differences between laboratories**

| Min | A | B | C | D | E | F | G | H | I | J | K | L | M | N |
|---|---|---|---|---|---|---|---|---|---|---|---|---|---|---|
| A | - | 2.621 | 3.571 | 3.346 | 1.344 | 4.215 | 2.982 | 3.733 | -2.854 | 2.879 | 2.838 | 3.821 | 3.084 | 4.285 |
| B | - | - | 2.791 | 4.185 | 3.705 | 3.959 | 2.081 | 3.248 | 1.88 | 4.298 | -1.294 | 3.388 | 1.691 | 4.055 |
| C | - | - | - | 4.401 | 4.326 | 3.368 | 0.537 | 2.052 | 3.617 | 4.731 | 1.39 | 2.08 | 0.533 | 3.432 |
| D | - | - | - | - | 1.57 | 4.907 | 4.306 | 4.641 | 3.621 | 0.916 | 4.032 | 4.705 | 4.243 | 5.021 |
| E | - | - | - | - | - | 5.153 | 4.854 | 4.795 | 4.272 | 3.206 | 4.326 | 4.916 | 4.43 | 5.344 |
| F | - | - | - | - | - | - | 3.749 | 2.216 | 4.459 | 5.199 | 3.666 | 2.619 | 3.445 | -4.605 |
| G | - | - | - | - | - | - | - | 2.831 | 3.28 | 4.468 | -0.744 | 2.549 | 0.867 | 3.631 |
| H | - | - | - | - | - | - | - | - | 4.016 | 4.965 | 3.045 | -0.667 | 2.719 | 2.464 |
| I | - | - | - | - | - | - | - | - | - | 3.677 | 2.796 | 4.147 | 2.818 | 4.447 |
| J | - | - | - | - | - | - | - | - | - | - | 3.946 | 4.453 | 4.379 | 5.048 |
| K | - | - | - | - | - | - | - | - | - | - | - | 3.18 | 1.102 | 3.89 |
| L | - | - | - | - | - | - | - | - | - | - | - | - | 2.582 | 2.747 |
| M | - | - | - | - | - | - | - | - | - | - | - | - | - | 3.394 |
| N | - | - | - | - | - | - | - | - | - | - | - | - | - | - |

| | |
|---|---|
| | Larger than 10 |
| | Between 8 and 9,99 |
| | Between 6 and 7,99 |
| | Between 4 and 5,99 |
| | Between 2 and 3,99 |
| | Between 0 and 1,99 |
| | NA (Calculation failed) |

| Median | A | B | C | D | E | F | G | H | I | J | K | L | M | N |
|---|---|---|---|---|---|---|---|---|---|---|---|---|---|---|
| A | - | 3.203 | 1.974 | 1.639 | 4.076 | 2.771 | 0.247 | 3.039 | 2.42 | 1.665 | 2.362 | 2.33 | 0.146 | 1.644 |
| B | - | - | 2.54 | 3.762 | 3.083 | 1.394 | 3.518 | 0.123 | 2.086 | 2.721 | 2.157 | 2.203 | 3.158 | 2.747 |
| C | - | - | - | 2.877 | 3.757 | 1.573 | 2.423 | 2.148 | 0.535 | -0.458 | 0.316 | 0.155 | 1.563 | -0.201 |
| D | - | - | - | - | 4.28 | 3.269 | 0.901 | 3.459 | 3.041 | 2.612 | 3.002 | 2.981 | 2.03 | 2.591 |
| E | - | - | - | - | - | 3.599 | 4.447 | 3.381 | 3.792 | 4.054 | 3.83 | 3.834 | 4.253 | 4.057 |
| F | - | - | - | - | - | - | 3.101 | 0.573 | 0.676 | 1.933 | 0.822 | 0.943 | 2.61 | 1.981 |
| G | - | - | - | - | - | - | - | 3.214 | 2.69 | 2.134 | 2.65 | 2.619 | 1.186 | 2.091 |
| H | - | - | - | - | - | - | - | - | 1.645 | 2.435 | 1.732 | 1.794 | 2.949 | 2.468 |
| I | - | - | - | - | - | - | - | - | - | 1.187 | -2.024 | -1.415 | 2.168 | 1.255 |
| J | - | - | - | - | - | - | - | - | - | - | 1.01 | 0.956 | 1.136 | -3.352 |
| K | - | - | - | - | - | - | - | - | - | - | - | -2.857 | 2.092 | 1.102 |
| L | - | - | - | - | - | - | - | - | - | - | - | - | 2.036 | 1.001 |
| M | - | - | - | - | - | - | - | - | - | - | - | - | - | 1.026 |
| N | - | - | - | - | - | - | - | - | - | - | - | - | - | - |

| Max | A | B | C | D | E | F | G | H | I | J | K | L | M | N |
|---|---|---|---|---|---|---|---|---|---|---|---|---|---|---|
| A | - | 3.209 | 1.933 | 1.579 | 4.109 | 2.73 | 0.405 | 3.022 | 2.443 | 1.714 | 2.353 | 2.288 | -0.004 | 1.536 |
| B | - | - | 2.583 | 3.753 | 3.154 | 1.547 | 3.548 | 0.37 | 2.089 | 2.691 | 2.178 | 2.276 | 3.187 | 2.814 |
| C | - | - | - | 2.827 | 3.827 | 1.514 | 2.439 | 2.15 | 0.745 | -1.125 | 0.386 | 0.089 | 1.574 | 0.043 |
| D | - | - | - | - | 4.304 | 3.219 | 0.675 | 3.423 | 3.029 | 2.622 | 2.977 | 2.927 | 1.95 | 2.501 |
| E | - | - | - | - | - | 3.688 | 4.497 | 3.48 | 3.848 | 4.077 | 3.884 | 3.904 | 4.298 | 4.124 |
| F | - | - | - | - | - | - | 3.089 | 0.683 | 0.333 | 1.797 | 0.672 | 0.891 | 2.593 | 2.001 |
| G | - | - | - | - | - | - | - | 3.221 | 2.749 | 2.222 | 2.675 | 2.62 | 1.19 | 2.053 |
| H | - | - | - | - | - | - | - | - | 1.541 | 2.361 | 1.698 | 1.812 | 2.952 | 2.507 |
| I | - | - | - | - | - | - | - | - | - | 1.116 | -1.47 | -0.592 | 2.222 | 1.443 |
| J | - | - | - | - | - | - | - | - | - | - | 0.848 | 0.698 | 1.343 | -0.852 |
| K | - | - | - | - | - | - | - | - | - | - | - | -1.812 | 2.123 | 1.251 |
| L | - | - | - | - | - | - | - | - | - | - | - | - | 2.026 | 1.085 |
| M | - | - | - | - | - | - | - | - | - | - | - | - | - | 0.941 |
| N | - | - | - | - | - | - | - | - | - | - | - | - | - | - |
