## Supplementary Data S2 for "Interlaboratory comparison of standardised metabolomics and lipidomics analyses in human and rodent blood using the MxP® Quant 500 kit": SM C24.0.pdf

**Amount of relative differences between laboratories**

| Min | A | B | C | D | E | F | G | H | I | J | K | L | M | N |
|---|---|---|---|---|---|---|---|---|---|---|---|---|---|---|
| A | - | 3.392 | 3.386 | 2.693 | 3.743 | 3.667 | 1.648 | 2.628 | 0.613 | 3.073 | -2.219 | 2.281 | 1.937 | -1.602 |
| B | - | - | -1.464 | 2.437 | 0.257 | 1.637 | 2.987 | 2.153 | 3.351 | 4.508 | 3.563 | 2.631 | 2.816 | 3.566 |
| C | - | - | - | 2.396 | 0.479 | 1.502 | 3.02 | 2.251 | 3.355 | 4.503 | 3.6 | 2.702 | 2.876 | 3.596 |
| D | - | - | - | - | 3.111 | 2.678 | 1.471 | 0.173 | 1.653 | 3.921 | 2.559 | -4.96 | 0.736 | 2.616 |
| E | - | - | - | - | - | 2.287 | 2.707 | 2.014 | 3.349 | 4.408 | 3.431 | 2.816 | 2.498 | 3.508 |
| F | - | - | - | - | - | - | 3.375 | 2.706 | 3.493 | 4.584 | 3.784 | 2.93 | 3.227 | 3.817 |
| G | - | - | - | - | - | - | - | 1.687 | 0.742 | 3.713 | 1.832 | 0.771 | -0.405 | 1.828 |
| H | - | - | - | - | - | - | - | - | 2.468 | 4.093 | 2.83 | 1.01 | 1.267 | 2.765 |
| I | - | - | - | - | - | - | - | - | - | 3.375 | 0.295 | 1.775 | 1.729 | -0.252 |
| J | - | - | - | - | - | - | - | - | - | - | 3.04 | 3.551 | 3.709 | 3.253 |
| K | - | - | - | - | - | - | - | - | - | - | - | 2.076 | 2.184 | -1.062 |
| L | - | - | - | - | - | - | - | - | - | - | - | - | -4.275 | 2.132 |
| M | - | - | - | - | - | - | - | - | - | - | - | - | - | 2.125 |
| N | - | - | - | - | - | - | - | - | - | - | - | - | - | - |

| Median | A | B | C | D | E | F | G | H | I | J | K | L | M | N |
|---|---|---|---|---|---|---|---|---|---|---|---|---|---|---|
| A | - | 3.619 | 3.568 | 0.276 | 4.303 | 3.023 | 1.626 | 3.363 | 2.951 | 1.846 | 3.072 | 3.327 | 2.333 | 2.785 |
| B | - | - | -1.027 | 3.946 | 3.075 | 2.256 | 3.397 | 1.204 | 2.401 | 3.315 | 2.133 | 1.381 | 3.036 | 2.623 |
| C | - | - | - | 3.891 | 3.152 | 2.089 | 3.319 | 0.845 | 2.245 | 3.234 | 1.964 | 1.067 | 2.943 | 2.495 |
| D | - | - | - | - | 4.378 | 3.237 | 2.165 | 3.531 | 3.204 | 2.264 | 3.234 | 3.502 | 2.619 | 3.025 |
| E | - | - | - | - | - | 3.862 | 4.382 | 3.562 | 3.939 | 4.317 | 3.771 | 3.607 | 4.158 | 3.994 |
| F | - | - | - | - | - | - | 2.452 | 1.233 | -1.67 | 2.338 | -3.317 | 1.062 | 1.819 | 0.48 |
| G | - | - | - | - | - | - | - | 2.894 | 2.262 | -1.047 | 2.449 | 2.841 | 1.011 | 1.973 |
| H | - | - | - | - | - | - | - | - | 1.57 | 2.891 | 0.967 | -1.714 | 2.507 | 1.904 |
| I | - | - | - | - | - | - | - | - | - | 2.067 | 0.209 | 1.404 | 1.307 | -0.607 |
| J | - | - | - | - | - | - | - | - | - | - | 2.389 | 2.687 | 0.815 | 1.748 |
| K | - | - | - | - | - | - | - | - | - | - | - | 0.489 | 1.882 | 0.931 |
| L | - | - | - | - | - | - | - | - | - | - | - | - | 2.402 | 1.766 |
| M | - | - | - | - | - | - | - | - | - | - | - | - | - | 0.938 |
| N | - | - | - | - | - | - | - | - | - | - | - | - | - | - |

| Max | A | B | C | D | E | F | G | H | I | J | K | L | M | N |
|---|---|---|---|---|---|---|---|---|---|---|---|---|---|---|
| A | - | 3.687 | 3.616 | 1.964 | 4.463 | 2.802 | 1.62 | 3.522 | 3.224 | 2.761 | 3.491 | 3.548 | 2.444 | 3.13 |
| B | - | - | -0.45 | 4.241 | 3.468 | 2.758 | 3.491 | 0.745 | 2.002 | 2.714 | 0.859 | 0.535 | 3.101 | 2.231 |
| C | - | - | - | 4.186 | 3.538 | 2.595 | 3.39 | -0.106 | 1.73 | 2.56 | 0.074 | -0.573 | 2.965 | 1.991 |
| D | - | - | - | - | 4.657 | 3.35 | 2.692 | 3.869 | 3.634 | 3.335 | 3.879 | 3.905 | 3.126 | 3.563 |
| E | - | - | - | - | - | 4.28 | 4.629 | 3.816 | 4.06 | 4.292 | 3.872 | 3.791 | 4.452 | 4.096 |
| F | - | - | - | - | - | - | 2.049 | 2.307 | 1.334 | -4.717 | 2.294 | 2.383 | 0.676 | 0.912 |
| G | - | - | - | - | - | - | - | 3.115 | 2.694 | 2.031 | 3.102 | 3.158 | 1.283 | 2.549 |
| H | - | - | - | - | - | - | - | - | 1.205 | 2.282 | -2.567 | -2.025 | 2.759 | 1.563 |
| I | - | - | - | - | - | - | - | - | - | 1.351 | 0.986 | 1.26 | 2.134 | -0.714 |
| J | - | - | - | - | - | - | - | - | - | - | 2.071 | 2.244 | 0.99 | 0.822 |
| K | - | - | - | - | - | - | - | - | - | - | - | -1.008 | 2.68 | 1.334 |
| L | - | - | - | - | - | - | - | - | - | - | - | - | 2.811 | 1.654 |
| M | - | - | - | - | - | - | - | - | - | - | - | - | - | 1.822 |
| N | - | - | - | - | - | - | - | - | - | - | - | - | - | - |

| | |
|---|---|
| | Larger than 10 |
| | Between 8 and 9,99 |
| | Between 6 and 7,99 |
| | Between 4 and 5,99 |
| | Between 2 and 3,99 |
| | Between 0 and 1,99 |
| | NA (Calculation failed) |
