## Supplementary Data S2 for "Interlaboratory comparison of standardised metabolomics and lipidomics analyses in human and rodent blood using the MxP® Quant 500 kit": SM C26.0.pdf

**Amount of relative differences between laboratories**

| Min | A | B | C | D | E | F | G | H | I | J | K | L | M | N |
|---|---|---|---|---|---|---|---|---|---|---|---|---|---|---|
| A | - | 3.368 | 1.25 | 5.763 | 2.611 | 5.284 | 3.757 | 4.419 | 5.008 | 6.035 | 4.603 | 1.938 | 4.335 | 1.893 |
| B | - | - | 3.928 | 6.227 | 4.647 | 4.887 | 4.793 | 2.707 | 3.946 | 6.291 | 4.629 | 3.296 | 3.506 | 3.729 |
| C | - | - | - | 5.675 | 2.38 | 5.305 | 3.373 | 4.414 | 5.025 | 5.924 | 3.967 | 1.925 | 4.456 | 2.248 |
| D | - | - | - | - | 4.777 | 6.043 | 4.711 | 5.9 | 5.936 | 4.622 | 3.304 | 4.98 | 5.568 | 5.471 |
| E | - | - | - | - | - | 5.598 | 2.769 | 5.313 | 5.799 | 5.554 | 4.136 | 2.753 | 4.632 | 4.127 |
| F | - | - | - | - | - | - | 6.296 | 5.006 | 3.897 | 7.066 | 5.467 | 5.588 | 4.43 | 5.955 |
| G | - | - | - | - | - | - | - | 5.234 | 5.405 | 5.486 | 4.284 | 3.871 | 4.782 | 4.402 |
| H | - | - | - | - | - | - | - | - | 3.599 | 6.427 | 5.068 | 4.43 | 2.023 | 4.506 |
| I | - | - | - | - | - | - | - | - | - | 6.793 | 4.797 | 4.908 | 2.808 | 5.478 |
| J | - | - | - | - | - | - | - | - | - | - | 3.321 | 5.362 | 5.912 | 5.972 |
| K | - | - | - | - | - | - | - | - | - | - | - | 1.685 | 4.533 | 5.227 |
| L | - | - | - | - | - | - | - | - | - | - | - | - | 4.021 | 2.504 |
| M | - | - | - | - | - | - | - | - | - | - | - | - | - | 4.878 |
| N | - | - | - | - | - | - | - | - | - | - | - | - | - | - |

| | |
|---|---|
| | Larger than 10 |
| | Between 8 and 9,99 |
| | Between 6 and 7,99 |
| | Between 4 and 5,99 |
| | Between 2 and 3,99 |
| | Between 0 and 1,99 |
| | NA (Calculation failed) |

| Median | A | B | C | D | E | F | G | H | I | J | K | L | M | N |
|---|---|---|---|---|---|---|---|---|---|---|---|---|---|---|
| A | - | 1.48 | 2.347 | 1.978 | 0.884 | 2.925 | -1.882 | -1.381 | 2.389 | 3.919 | 4.661 | 1.797 | 3.124 | 2.732 |
| B | - | - | 2.97 | 2.765 | 0.056 | 2.318 | 1.298 | 1.451 | 2.983 | 4.191 | 4.811 | -0.367 | 3.543 | 3.259 |
| C | - | - | - | 0.079 | 2.704 | 3.588 | 2.34 | 2.193 | -2.796 | 3.257 | 4.246 | 3.021 | 1.794 | 0.603 |
| D | - | - | - | - | 2.593 | 3.497 | 1.993 | 1.198 | 0.592 | 3.47 | 4.363 | 2.872 | 2.286 | 1.359 |
| E | - | - | - | - | - | 2.529 | 0.804 | 1.653 | 2.972 | 4.153 | 4.774 | 0.676 | 3.455 | 3.143 |
| F | - | - | - | - | - | - | 2.898 | 2.777 | 3.712 | 4.564 | 5.117 | 2.127 | 4.079 | 3.916 |
| G | - | - | - | - | - | - | - | 0.24 | 2.485 | 3.972 | 4.674 | 1.724 | 3.176 | 2.791 |
| H | - | - | - | - | - | - | - | - | 2.404 | 3.934 | 4.676 | 1.657 | 3.142 | 2.795 |
| I | - | - | - | - | - | - | - | - | - | 3.202 | 4.242 | 2.955 | 1.734 | 0.77 |
| J | - | - | - | - | - | - | - | - | - | - | 2.978 | 4.067 | 2.49 | 2.96 |
| K | - | - | - | - | - | - | - | - | - | - | - | 4.489 | 3.698 | 3.922 |
| L | - | - | - | - | - | - | - | - | - | - | - | - | 3.656 | 3.398 |
| M | - | - | - | - | - | - | - | - | - | - | - | - | - | 0.648 |
| N | - | - | - | - | - | - | - | - | - | - | - | - | - | - |

| Max | A | B | C | D | E | F | G | H | I | J | K | L | M | N |
|---|---|---|---|---|---|---|---|---|---|---|---|---|---|---|
| A | - | 0.686 | 2.419 | -1.188 | 1.493 | 1.915 | 0.478 | 0.251 | 3.223 | 3.408 | 4.693 | 1.783 | 3.627 | 2.902 |
| B | - | - | 2.832 | 1.048 | 0.916 | 0.987 | -1.307 | 1.621 | 3.387 | 3.688 | 4.905 | 0.877 | 3.864 | 3.214 |
| C | - | - | - | 2.227 | 2.99 | 3.115 | 2.659 | 1.983 | 1.899 | 2.318 | 4.383 | 3.096 | 2.738 | 1.016 |
| D | - | - | - | - | 1.777 | 2.21 | 1.12 | -2.563 | 3.068 | 3.303 | 4.735 | 2.292 | 3.52 | 2.661 |
| E | - | - | - | - | - | -0.829 | 0.415 | 2.243 | 3.889 | 3.924 | 5.039 | 0.115 | 3.952 | 3.451 |
| F | - | - | - | - | - | - | 0.738 | 2.566 | 3.69 | 3.785 | 4.873 | -6.734 | 4.028 | 3.502 |
| G | - | - | - | - | - | - | - | 1.388 | 3.441 | 3.71 | 4.855 | 1.117 | 3.77 | 3.197 |
| H | - | - | - | - | - | - | - | - | 2.848 | 3.163 | 4.395 | 2.492 | 3.385 | 2.504 |
| I | - | - | - | - | - | - | - | - | - | -3.354 | 3.797 | 3.528 | 1.517 | 0.791 |
| J | - | - | - | - | - | - | - | - | - | - | 4.17 | 3.84 | 0.337 | 1.755 |
| K | - | - | - | - | - | - | - | - | - | - | - | 4.608 | 3.547 | 3.741 |
| L | - | - | - | - | - | - | - | - | - | - | - | - | 4.021 | 3.565 |
| M | - | - | - | - | - | - | - | - | - | - | - | - | - | 2.018 |
| N | - | - | - | - | - | - | - | - | - | - | - | - | - | - |
