## Supplementary Data S2 for "Interlaboratory comparison of standardised metabolomics and lipidomics analyses in human and rodent blood using the MxP® Quant 500 kit": SM C26.1.pdf

**Amount of relative differences between laboratories**

| Min | A | B | C | D | E | F | G | H | I | J | K | L | M | N |
|---|---|---|---|---|---|---|---|---|---|---|---|---|---|---|
| A | - | 4.543 | 5.937 | 3.928 | 1.552 | 5.867 | 4.319 | 5.551 | 3.233 | 3.415 | 4.253 | 5.169 | 5.052 | 6.454 |
| B | - | - | 4.757 | 1.726 | 3.356 | 4.495 | 3.58 | 3.41 | 1.994 | 3.641 | 0.619 | 3.388 | 2.725 | 5.478 |
| C | - | - | - | 4.872 | 5.188 | 2.379 | 5.485 | 4.354 | 4.802 | 5.387 | 4.83 | 3.947 | 4.403 | 4.252 |
| D | - | - | - | - | 3.327 | 5.102 | 2.002 | 4.271 | 1.183 | 2.973 | 0.419 | 4.106 | 3.467 | 5.916 |
| E | - | - | - | - | - | 6.058 | 4.647 | 5.701 | 2.476 | 3.276 | 4.134 | 5.319 | 5.092 | 6.659 |
| F | - | - | - | - | - | - | 5.452 | 4.082 | 4.91 | 5.397 | 4.831 | 3.868 | 4.291 | 4.758 |
| G | - | - | - | - | - | - | - | 4.456 | -1.918 | 2.261 | 1.948 | 4.35 | 3.828 | 5.962 |
| H | - | - | - | - | - | - | - | - | 5.253 | 5.396 | 4.85 | 2.146 | 3.639 | 5.834 |
| I | - | - | - | - | - | - | - | - | - | 2.471 | 2.5 | 4.475 | 4.073 | 5.971 |
| J | - | - | - | - | - | - | - | - | - | - | 3.21 | 4.652 | 4.41 | 6.14 |
| K | - | - | - | - | - | - | - | - | - | - | - | 4.1 | 3.459 | 5.881 |
| L | - | - | - | - | - | - | - | - | - | - | - | - | 2.095 | 5.372 |
| M | - | - | - | - | - | - | - | - | - | - | - | - | - | 5.659 |
| N | - | - | - | - | - | - | - | - | - | - | - | - | - | - |

| Colour | Legend |
|---|---|
| Red | Larger than 10 |
| Orange | Between 8 and 9,99 |
| Light orange | Between 6 and 7,99 |
| Light green | Between 4 and 5,99 |
| Green | Between 2 and 3,99 |
| Dark green | Between 0 and 1,99 |
| Grey | NA (Calculation failed) |

| Median | A | B | C | D | E | F | G | H | I | J | K | L | M | N |
|---|---|---|---|---|---|---|---|---|---|---|---|---|---|---|
| A | - | 3.844 | 0.847 | 1.747 | 3.41 | 2.729 | 2.458 | 4.663 | 2.443 | 2.209 | 1.174 | 0.19 | 2.291 | 2.239 |
| B | - | - | 3.491 | 3.296 | 4.443 | 4.174 | 3.005 | 5.115 | 2.863 | 3.182 | 3.443 | 3.759 | 4.082 | 3.129 |
| C | - | - | - | 0.704 | 3.545 | 2.895 | 1.854 | 4.716 | 2.264 | 1.205 | -1.314 | 1.307 | 2.668 | 1.476 |
| D | - | - | - | - | 3.739 | 3.228 | 1.017 | 4.788 | 1.283 | 0.002 | 0.16 | 2.078 | 2.984 | 0.318 |
| E | - | - | - | - | - | 2.145 | 4.184 | 4.037 | 4.162 | 4.112 | 3.865 | 3.413 | 2.704 | 4.115 |
| F | - | - | - | - | - | - | 3.603 | 4.342 | 3.719 | 3.437 | 3.118 | 2.454 | 0.391 | 3.494 |
| G | - | - | - | - | - | - | - | 4.861 | -1.361 | -0.013 | 1.62 | 2.617 | 3.285 | -0.4 |
| H | - | - | - | - | - | - | - | - | 5.376 | 5.311 | 5.198 | 5.015 | 4.769 | 5.322 |
| I | - | - | - | - | - | - | - | - | - | 0.491 | 1.799 | 2.709 | 3.341 | 0.224 |
| J | - | - | - | - | - | - | - | - | - | - | 1.106 | 2.402 | 3.158 | -2.381 |
| K | - | - | - | - | - | - | - | - | - | - | - | 1.656 | 2.783 | 1.258 |
| L | - | - | - | - | - | - | - | - | - | - | - | - | 1.971 | 2.478 |
| M | - | - | - | - | - | - | - | - | - | - | - | - | - | 3.312 |
| N | - | - | - | - | - | - | - | - | - | - | - | - | - | - |

| Max | A | B | C | D | E | F | G | H | I | J | K | L | M | N |
|---|---|---|---|---|---|---|---|---|---|---|---|---|---|---|
| A | - | 4.284 | 2.927 | 2.477 | 3.535 | 0.699 | 2.922 | 4.579 | 3.043 | 2.6 | 2.223 | 1.337 | 1.016 | 3.651 |
| B | - | - | 3.291 | 3.508 | 4.687 | 4.139 | 3.273 | 5.181 | 3.233 | 3.473 | 3.618 | 3.793 | 4.169 | 2.513 |
| C | - | - | - | 0.732 | 4.145 | 3.115 | -2.072 | 4.877 | -1.176 | 0.501 | 1.47 | 2.124 | 3.194 | 2.213 |
| D | - | - | - | - | 4.025 | 2.797 | 0.896 | 4.816 | 1.301 | -1.486 | 0.128 | 1.514 | 2.925 | 2.695 |
| E | - | - | - | - | - | 3.498 | 4.441 | 3.812 | 4.481 | 4.31 | 4.186 | 3.998 | 3.412 | 4.775 |
| F | - | - | - | - | - | - | 3.257 | 4.508 | 3.319 | 2.979 | 2.687 | 2.176 | -0.878 | 3.86 |
| G | - | - | - | - | - | - | - | 4.884 | -0.964 | 0.618 | 1.578 | 2.212 | 3.216 | 2.159 |
| H | - | - | - | - | - | - | - | - | 5.398 | 5.303 | 5.233 | 5.14 | 4.847 | 5.592 |
| I | - | - | - | - | - | - | - | - | - | 1.042 | 1.693 | 2.289 | 3.241 | 2.019 |
| J | - | - | - | - | - | - | - | - | - | - | 0.432 | 1.676 | 2.94 | 2.631 |
| K | - | - | - | - | - | - | - | - | - | - | - | 0.747 | 2.692 | 2.925 |
| L | - | - | - | - | - | - | - | - | - | - | - | - | 2.249 | 3.257 |
| M | - | - | - | - | - | - | - | - | - | - | - | - | - | 3.913 |
| N | - | - | - | - | - | - | - | - | - | - | - | - | - | - |
