## Supplementary Data S2 for "Interlaboratory comparison of standardised metabolomics and lipidomics analyses in human and rodent blood using the MxP® Quant 500 kit": t4-OH-Pro.pdf

**Amount of relative differences between laboratories**

| Min | A | B | C | D | E | F | G | H | I | J | K | L | M | N |
|---|---|---|---|---|---|---|---|---|---|---|---|---|---|---|
| A | - | 2.678 | 0.38 | -0.479 | 2.549 | 3.171 | 2.643 | -1.472 | 1.432 | 2.386 | 0.208 | 1.69 | 2.632 | 1.383 |
| B | - | - | 3.101 | 2.688 | 0.063 | 0.757 | 3.802 | 2.646 | 3.232 | 0.958 | 2.959 | 1.593 | -0.8 | 2.169 |
| C | - | - | - | 0.867 | 2.819 | 3.342 | 2.325 | 0.594 | 0.111 | 2.682 | -2.003 | 2.208 | 2.896 | 1.961 |
| D | - | - | - | - | 2.388 | 3.094 | 2.812 | -0.899 | 1.681 | 2.214 | 0.783 | 1.413 | 2.508 | 1.064 |
| E | - | - | - | - | - | 1.604 | 3.686 | 2.514 | 3.082 | -0.37 | 2.818 | 1.26 | -1.326 | 1.782 |
| F | - | - | - | - | - | - | 4.037 | 3.169 | 3.522 | 2.007 | 3.363 | 2.44 | 1.453 | 2.75 |
| G | - | - | - | - | - | - | - | 2.548 | 1.89 | 3.474 | 2.232 | 3.156 | 3.584 | 3.113 |
| H | - | - | - | - | - | - | - | - | 1.555 | 2.329 | 0.648 | 1.533 | 2.631 | 1.351 |
| I | - | - | - | - | - | - | - | - | - | 2.839 | 0.301 | 2.528 | 3.058 | 2.263 |
| J | - | - | - | - | - | - | - | - | - | - | 2.716 | 0.928 | 0.144 | 1.425 |
| K | - | - | - | - | - | - | - | - | - | - | - | 2.14 | 2.878 | 1.923 |
| L | - | - | - | - | - | - | - | - | - | - | - | - | 1.993 | -1.178 |
| M | - | - | - | - | - | - | - | - | - | - | - | - | - | 1.978 |
| N | - | - | - | - | - | - | - | - | - | - | - | - | - | - |

| Median | A | B | C | D | E | F | G | H | I | J | K | L | M | N |
|---|---|---|---|---|---|---|---|---|---|---|---|---|---|---|
| A | - | 1.91 | 0.891 | -1.56 | 2.098 | 2.74 | 2.422 | -2.217 | 1.842 | 2.166 | 0.336 | 2.697 | 1.211 | -3.797 |
| B | - | - | 2.556 | 1.776 | -0.661 | 1.66 | 3.25 | 1.973 | 2.96 | 0.16 | 2.293 | 1.538 | 0.433 | 1.871 |
| C | - | - | - | 1.09 | 2.597 | 3.076 | 1.799 | 0.927 | 0.708 | 2.656 | -0.641 | 3.039 | 2.05 | 0.959 |
| D | - | - | - | - | 1.991 | 2.687 | 2.521 | -1.872 | 1.964 | 2.079 | 0.638 | 2.628 | 1.03 | -2.314 |
| E | - | - | - | - | - | 1.339 | 3.345 | 2.173 | 3.044 | -1.506 | 2.506 | 1.192 | 1.022 | 2.124 |
| F | - | - | - | - | - | - | 3.71 | 2.876 | 3.462 | 1.164 | 3.075 | -1.902 | 2.237 | 2.829 |
| G | - | - | - | - | - | - | - | 2.389 | 0.964 | 3.219 | 1.936 | 3.496 | 2.874 | 2.379 |
| H | - | - | - | - | - | - | - | - | 1.906 | 2.127 | 0.55 | 2.656 | 1.17 | -6.187 |
| I | - | - | - | - | - | - | - | - | - | 2.982 | 1.209 | 3.29 | 2.525 | 1.849 |
| J | - | - | - | - | - | - | - | - | - | - | 2.597 | 1.021 | 1.211 | 2.219 |
| K | - | - | - | - | - | - | - | - | - | - | - | 2.939 | 1.833 | 0.425 |
| L | - | - | - | - | - | - | - | - | - | - | - | - | 2.197 | 2.808 |
| M | - | - | - | - | - | - | - | - | - | - | - | - | - | 1.188 |
| N | - | - | - | - | - | - | - | - | - | - | - | - | - | - |

| Max | A | B | C | D | E | F | G | H | I | J | K | L | M | N |
|---|---|---|---|---|---|---|---|---|---|---|---|---|---|---|
| A | - | -0.291 | 1.442 | -2.36 | 1.051 | 1.947 | 2.009 | -3.507 | 2.436 | 1.87 | 0.517 | 3.559 | 1.78 | 1.772 |
| B | - | - | 0.734 | -0.258 | 1.578 | 2.306 | 1.512 | -0.074 | 2.156 | 2.194 | -1.51 | 3.648 | 1.322 | 1.324 |
| C | - | - | - | 1.356 | 2.213 | 2.669 | 0.239 | 1.442 | 1.472 | 2.626 | 0.334 | 3.815 | -0.419 | -0.443 |
| D | - | - | - | - | 1.142 | 1.961 | 1.929 | -1.857 | 2.414 | 1.909 | 0.373 | 3.573 | 1.748 | 1.745 |
| E | - | - | - | - | - | 0.933 | 2.606 | 1.003 | 2.963 | 0.7 | 1.811 | 3.307 | 2.492 | 2.492 |
| F | - | - | - | - | - | - | 3.014 | 1.961 | 3.33 | -2.119 | 2.453 | 3.023 | 2.933 | 2.931 |
| G | - | - | - | - | - | - | - | 1.987 | 0.733 | 2.864 | 1.293 | 3.92 | -1.199 | -1.129 |
| H | - | - | - | - | - | - | - | - | 2.436 | 1.858 | 0.382 | 3.566 | 1.845 | 1.823 |
| I | - | - | - | - | - | - | - | - | - | 3.129 | 2.007 | 4.028 | 0.962 | 0.967 |
| J | - | - | - | - | - | - | - | - | - | - | 2.385 | 3.075 | 2.883 | 2.881 |
| K | - | - | - | - | - | - | - | - | - | - | - | 3.698 | 1.035 | 1.015 |
| L | - | - | - | - | - | - | - | - | - | - | - | - | 4.11 | 4.107 |
| M | - | - | - | - | - | - | - | - | - | - | - | - | - | -8.158 |
| N | - | - | - | - | - | - | - | - | - | - | - | - | - | - |

| | |
|---|---|
| | Larger than 10 |
| | Between 8 and 9,99 |
| | Between 6 and 7,99 |
| | Between 4 and 5,99 |
| | Between 2 and 3,99 |
| | Between 0 and 1,99 |
| | NA (Calculation failed) |
