## Supplementary Data S2 for "Interlaboratory comparison of standardised metabolomics and lipidomics analyses in human and rodent blood using the MxP® Quant 500 kit": Taurine.pdf

**Amount of relative differences between laboratories**

| Min | A | B | C | D | E | F | G | H | I | J | K | L | M | N |
|---|---|---|---|---|---|---|---|---|---|---|---|---|---|---|
| A | - | 2.499 | 3.73 | 2.328 | -0.211 | 1.696 | 2.593 | 3.019 | 2.441 | 1.816 | 0.499 | -1.436 | -4.713 | 1.455 |
| B | - | - | 4.367 | 3.526 | 2.505 | 1.2 | 3.57 | 1.763 | 3.709 | 3.21 | 1.855 | 2.903 | 2.453 | 3.11 |
| C | - | - | - | 3.055 | 3.629 | 4.076 | 3.278 | 4.542 | 2.124 | 3.56 | 4.165 | 2.713 | 3.914 | 3.563 |
| D | - | - | - | - | 2.294 | 3.112 | -1.164 | 3.832 | 1.451 | 1.425 | 3.014 | -0.074 | 2.629 | 1.605 |
| E | - | - | - | - | - | 2.166 | 1.904 | 3.508 | 3.124 | 0.486 | 2.278 | 3.053 | 1.238 | 0.305 |
| F | - | - | - | - | - | - | 3.199 | 2.559 | 3.501 | 2.733 | -0.423 | 2.049 | 1.544 | 2.629 |
| G | - | - | - | - | - | - | - | 3.789 | 0.701 | 1.222 | 2.918 | 1.2 | 2.52 | 1.506 |
| H | - | - | - | - | - | - | - | - | 3.99 | 3.7 | 2.852 | 2.298 | 3.176 | 3.605 |
| I | - | - | - | - | - | - | - | - | - | 2.495 | 3.422 | -0.639 | 3.199 | 2.587 |
| J | - | - | - | - | - | - | - | - | - | - | 2.514 | -1.579 | 1.827 | -1.452 |
| K | - | - | - | - | - | - | - | - | - | - | - | 0.963 | 0.519 | 2.082 |
| L | - | - | - | - | - | - | - | - | - | - | - | - | 1.53 | 0.154 |
| M | - | - | - | - | - | - | - | - | - | - | - | - | - | 1.616 |
| N | - | - | - | - | - | - | - | - | - | - | - | - | - | - |

| | |
|---|---|
| | Larger than 10 |
| | Between 8 and 9,99 |
| | Between 6 and 7,99 |
| | Between 4 and 5,99 |
| | Between 2 and 3,99 |
| | Between 0 and 1,99 |
| | NA (Calculation failed) |

| Median | A | B | C | D | E | F | G | H | I | J | K | L | M | N |
|---|---|---|---|---|---|---|---|---|---|---|---|---|---|---|
| A | - | 3.263 | 0.49 | 1.828 | 3.574 | 3.302 | -3.379 | 3.414 | -0.305 | 2.699 | 1.017 | 2.649 | 2.575 | 0.698 |
| B | - | - | 3.178 | 2.72 | 1.299 | -1.383 | 3.383 | 0.228 | 3.167 | 1.914 | 3.073 | 1.615 | 2.081 | 3.119 |
| C | - | - | - | 1.285 | 3.376 | 3.134 | 0.185 | 3.244 | 0.779 | 2.018 | -0.59 | 2.693 | 1.969 | -0.729 |
| D | - | - | - | - | 3.06 | 2.702 | 1.857 | 2.835 | 0.741 | 1.135 | 0.922 | 1.82 | 0.926 | 0.963 |
| E | - | - | - | - | - | 0.949 | 3.694 | 0.196 | 3.341 | 2.798 | 3.469 | 2.477 | 2.84 | 3.467 |
| F | - | - | - | - | - | - | 3.44 | -0.437 | 3.157 | 2.139 | 3.145 | 1.289 | 2.263 | 3.183 |
| G | - | - | - | - | - | - | - | 3.395 | 0.519 | 2.569 | 0.868 | 2.694 | 2.471 | 0.67 |
| H | - | - | - | - | - | - | - | - | 3.398 | 2.287 | 3.273 | 1.583 | 2.426 | 3.328 |
| I | - | - | - | - | - | - | - | - | - | 1.727 | -0.445 | 2.053 | 1.63 | -0.757 |
| J | - | - | - | - | - | - | - | - | - | - | 2.049 | 0.917 | -1.608 | 2.101 |
| K | - | - | - | - | - | - | - | - | - | - | - | 2.443 | 2.008 | -1.437 |
| L | - | - | - | - | - | - | - | - | - | - | - | - | 1.564 | 2.801 |
| M | - | - | - | - | - | - | - | - | - | - | - | - | - | 2.071 |
| N | - | - | - | - | - | - | - | - | - | - | - | - | - | - |

| Max | A | B | C | D | E | F | G | H | I | J | K | L | M | N |
|---|---|---|---|---|---|---|---|---|---|---|---|---|---|---|
| A | - | 3.948 | 4.371 | 3.986 | 4.741 | 4.236 | 3.045 | 3.804 | 4.055 | 4.107 | 1.504 | 4.15 | 3.714 | 3.003 |
| B | - | - | 2.625 | -1.073 | 3.742 | 2.005 | 3.089 | 0.777 | 0.488 | 1.079 | 3.888 | 2.468 | 1.456 | 3.133 |
| C | - | - | - | 2.581 | 2.946 | 1.119 | 3.96 | 3.079 | 2.317 | 2.087 | 4.481 | 2.652 | 3.236 | 3.985 |
| D | - | - | - | - | 3.694 | 1.849 | 3.177 | 1.059 | 0.035 | 0.742 | 3.924 | 3.097 | 1.688 | 3.212 |
| E | - | - | - | - | - | 3.409 | 4.655 | 4.12 | 3.779 | 3.69 | 5.013 | -1.979 | 4.212 | 4.67 |
| F | - | - | - | - | - | - | 3.709 | 2.572 | 1.499 | 0.996 | 4.295 | -0.23 | 2.817 | 3.736 |
| G | - | - | - | - | - | - | - | 2.64 | 3.2 | 3.292 | 2.538 | 3.867 | 2.411 | -2.063 |
| H | - | - | - | - | - | - | - | - | 1.592 | 1.934 | 3.692 | -0.419 | -0.001 | 2.791 |
| I | - | - | - | - | - | - | - | - | - | -0.292 | 3.999 | 3.457 | 2.062 | 3.361 |
| J | - | - | - | - | - | - | - | - | - | - | 4.109 | 2.297 | 2.31 | 3.475 |
| K | - | - | - | - | - | - | - | - | - | - | - | 3.612 | 3.398 | 2.405 |
| L | - | - | - | - | - | - | - | - | - | - | - | - | 3.312 | 3.96 |
| M | - | - | - | - | - | - | - | - | - | - | - | - | - | 2.553 |
| N | - | - | - | - | - | - | - | - | - | - | - | - | - | - |
