## Supplementary Data S2 for "Interlaboratory comparison of standardised metabolomics and lipidomics analyses in human and rodent blood using the MxP® Quant 500 kit": TCDCA.pdf

**Amount of relative differences between laboratories**

| Min | A | B | C | D | E | F | G | H | I | J | K | L | M | N |
|---|---|---|---|---|---|---|---|---|---|---|---|---|---|---|
| A | - | 3.978 | 2.216 | 2.459 | 5.626 | 2.468 | 5.471 | 4.512 | 2.484 | 4.618 | 2.431 | 6.134 | 3.023 | 3.653 |
| B | - | - | 3.283 | 4.108 | 5.042 | 2.348 | 4.88 | 3.499 | 4.235 | 5.165 | 4.076 | 5.713 | 0.653 | 4.34 |
| C | - | - | - | 3.382 | 5.445 | -2.018 | 5.138 | 3.882 | 3.147 | 4.771 | 3.207 | 5.9 | 0.551 | 4.25 |
| D | - | - | - | - | 5.828 | 3.561 | 5.678 | 4.839 | 1.263 | 4.243 | 0.465 | 6.291 | 3.782 | 3.247 |
| E | - | - | - | - | - | 4.771 | -0.442 | 3.476 | 5.614 | 5.888 | 5.383 | 4.125 | 4.408 | 5.072 |
| F | - | - | - | - | - | - | 5.18 | 3.972 | 3.681 | 4.928 | 3.681 | 5.898 | 1.163 | 4.176 |
| G | - | - | - | - | - | - | - | 3.666 | 5.413 | 5.766 | 5.213 | 4.134 | 4.524 | 5.165 |
| H | - | - | - | - | - | - | - | - | 5.046 | 5.53 | 4.793 | 5.141 | 3.542 | 4.744 |
| I | - | - | - | - | - | - | - | - | - | 4.379 | -0.349 | 6.209 | 3.675 | 3.562 |
| J | - | - | - | - | - | - | - | - | - | - | 4.452 | 6.79 | 5.061 | 2.094 |
| K | - | - | - | - | - | - | - | - | - | - | - | 6.228 | 3.606 | 3.629 |
| L | - | - | - | - | - | - | - | - | - | - | - | - | 4.982 | 5.377 |
| M | - | - | - | - | - | - | - | - | - | - | - | - | - | 4.263 |
| N | - | - | - | - | - | - | - | - | - | - | - | - | - | - |

| | |
|---|---|
| | Larger than 10 |
| | Between 8 and 9,99 |
| | Between 6 and 7,99 |
| | Between 4 and 5,99 |
| | Between 2 and 3,99 |
| | Between 0 and 1,99 |
| | NA (Calculation failed) |

| Median | A | B | C | D | E | F | G | H | I | J | K | L | M | N |
|---|---|---|---|---|---|---|---|---|---|---|---|---|---|---|
| A | - | 0.505 | 3.582 | 0.078 | -0.365 | 2.654 | 3.88 | 2.037 | 0.596 | 2.871 | 2.25 | 3.063 | 2.342 | 2.235 |
| B | - | - | 3.375 | 1.316 | -0.45 | 2.268 | 3.702 | 1.38 | -2.923 | 3.114 | 1.663 | 2.765 | 1.839 | 2.629 |
| C | - | - | - | 3.541 | 3.302 | 2.331 | 1.304 | 2.812 | 3.226 | 4.106 | 2.665 | 1.694 | 2.616 | 3.895 |
| D | - | - | - | - | 0.896 | 2.899 | 3.996 | 2.386 | 1.389 | 2.657 | 2.581 | 3.247 | 2.64 | 1.889 |
| E | - | - | - | - | - | 2.536 | 3.798 | 1.804 | 0.165 | 2.974 | 1.925 | 2.954 | 2.16 | 2.461 |
| F | - | - | - | - | - | - | 2.984 | 1.06 | 2.198 | 3.689 | 0.665 | 0.916 | 0.208 | 3.38 |
| G | - | - | - | - | - | - | - | 3.196 | 3.496 | 4.257 | 3.121 | 2.442 | 3.057 | 4.086 |
| H | - | - | - | - | - | - | - | - | 1.261 | 3.456 | -1.045 | 2.034 | -0.09 | 3.099 |
| I | - | - | - | - | - | - | - | - | - | 3.121 | 1.721 | 2.744 | 1.859 | 2.625 |
| J | - | - | - | - | - | - | - | - | - | - | 3.726 | 4.075 | 3.746 | 1.506 |
| K | - | - | - | - | - | - | - | - | - | - | - | 1.714 | -1.875 | 3.197 |
| L | - | - | - | - | - | - | - | - | - | - | - | - | 1.599 | 3.622 |
| M | - | - | - | - | - | - | - | - | - | - | - | - | - | 3.226 |
| N | - | - | - | - | - | - | - | - | - | - | - | - | - | - |

| Max | A | B | C | D | E | F | G | H | I | J | K | L | M | N |
|---|---|---|---|---|---|---|---|---|---|---|---|---|---|---|
| A | - | -3.819 | 3.688 | -1.081 | 1.963 | 2.68 | 3.459 | 0.496 | 1.184 | 2.348 | 2.652 | 1.402 | 2.185 | 1.488 |
| B | - | - | 3.685 | -0.854 | 1.991 | 2.656 | 3.453 | 0.434 | 1.111 | 2.364 | 2.629 | 1.368 | 2.156 | 1.476 |
| C | - | - | - | 3.56 | 3.907 | 2.542 | 0.714 | 3.344 | 3.235 | 3.992 | 2.562 | 3.178 | 2.893 | 3.794 |
| D | - | - | - | - | 1.777 | 2.767 | 3.527 | 0.894 | 1.402 | 2.226 | 2.739 | 1.651 | 2.308 | 1.203 |
| E | - | - | - | - | - | 3.429 | 3.96 | 2.485 | 2.696 | 0.296 | 3.414 | 2.777 | 3.147 | 0.374 |
| F | - | - | - | - | - | - | 2.122 | 2.216 | 1.891 | 3.411 | -3.211 | 1.784 | 0.795 | 3.097 |
| G | - | - | - | - | - | - | - | 3.113 | 2.987 | 3.864 | 2.144 | 2.918 | 2.539 | 3.634 |
| H | - | - | - | - | - | - | - | - | -0.184 | 2.685 | 2.244 | 0.278 | 1.626 | 2.055 |
| I | - | - | - | - | - | - | - | - | - | 2.835 | 1.967 | -1.035 | 1.048 | 2.264 |
| J | - | - | - | - | - | - | - | - | - | - | 3.576 | 3.034 | 3.324 | 1.318 |
| K | - | - | - | - | - | - | - | - | - | - | - | 1.739 | 0.815 | 3.07 |
| L | - | - | - | - | - | - | - | - | - | - | - | - | 0.885 | 2.39 |
| M | - | - | - | - | - | - | - | - | - | - | - | - | - | 2.803 |
| N | - | - | - | - | - | - | - | - | - | - | - | - | - | - |
