## Supplementary Data S2 for "Interlaboratory comparison of standardised metabolomics and lipidomics analyses in human and rodent blood using the MxP® Quant 500 kit": TG(14.0_34.0).pdf

**Amount of relative differences between laboratories**

| Min | A | B | C | D | E | F | G | H | I | J | K | L | M | N |
|---|---|---|---|---|---|---|---|---|---|---|---|---|---|---|
| A | - | 7.003 | 5.817 | 9.13 | 7.622 | 6.982 | 7.479 | 6.835 | 5.696 | 8.111 | 6.668 | 6.777 | 6.394 | 7.226 |
| B | - | - | 4.792 | 8.73 | 7.312 | 4.5 | 4.459 | 4.798 | 7.394 | 7.469 | 3.002 | 6.521 | 5.338 | 1.307 |
| C | - | - | - | 8.82 | 7.27 | 5.489 | 4.8 | 2.88 | 6.805 | 7.466 | 4.312 | 5.325 | 5.707 | 5.205 |
| D | - | - | - | - | 6.249 | 6.856 | 8.2 | 7.622 | 8.687 | 3.355 | 7.152 | 8.44 | 4.46 | 7.646 |
| E | - | - | - | - | - | 9.03 | 10.02 | 9.465 | 10.267 | 7.927 | 9.1 | 10.097 | 6.715 | 9.549 |
| F | - | - | - | - | - | - | 5.99 | 5.626 | 7.641 | 7.314 | 4.828 | 7.02 | 5.508 | 4.942 |
| G | - | - | - | - | - | - | - | 1.103 | 6.907 | 7.403 | 3.335 | 5.89 | 5.544 | 4.133 |
| H | - | - | - | - | - | - | - | - | 6.869 | 7.479 | 3.554 | 5.812 | 5.613 | 4.329 |
| I | - | - | - | - | - | - | - | - | - | 8.125 | 6.292 | 6.074 | 6.194 | 6.943 |
| J | - | - | - | - | - | - | - | - | - | - | 7.083 | 8.374 | 1.127 | 7.611 |
| K | - | - | - | - | - | - | - | - | - | - | - | 6.282 | 5.545 | 1.879 |
| L | - | - | - | - | - | - | - | - | - | - | - | - | 5.752 | 6.07 |
| M | - | - | - | - | - | - | - | - | - | - | - | - | - | 7.616 |
| N | - | - | - | - | - | - | - | - | - | - | - | - | - | - |

| Median | A | B | C | D | E | F | G | H | I | J | K | L | M | N |
|---|---|---|---|---|---|---|---|---|---|---|---|---|---|---|
| A | - | 4.638 | 2.775 | 5.15 | 0.896 | 4.743 | 4 | 4.106 | 3.984 | 4.193 | 4.37 | 4.09 | 3.867 | 4.905 |
| B | - | - | 4.284 | 4.488 | 2.431 | 2.126 | 3.813 | 4 | 3.703 | 3.285 | 3.49 | 3.683 | 1.315 | 2.789 |
| C | - | - | - | 5.472 | 3.275 | 4.311 | 2.759 | 1.979 | 2.253 | 2.999 | 2.946 | 2.662 | 4.049 | 4.524 |
| D | - | - | - | - | 2.789 | 2.501 | 4.935 | 5.047 | 4.619 | 4.934 | 4.897 | 4.863 | 2.111 | 2.414 |
| E | - | - | - | - | - | 6.679 | 7.244 | 7.319 | 7.123 | 6.562 | 7.28 | 7.21 | 3.879 | 6.627 |
| F | - | - | - | - | - | - | 4.291 | 4.325 | 4.016 | 3.736 | 4.04 | 4.148 | 2.709 | 1.115 |
| G | - | - | - | - | - | - | - | -3.981 | 1.21 | 1.318 | 1.624 | 0.625 | 3.701 | 4.164 |
| H | - | - | - | - | - | - | - | - | 1.213 | 2.352 | 1.661 | 0.658 | 3.75 | 4.165 |
| I | - | - | - | - | - | - | - | - | - | 1.828 | 2.197 | -0.109 | 1.064 | 4.06 |
| J | - | - | - | - | - | - | - | - | - | - | 3.739 | 3.517 | 4.72 | 3.28 |
| K | - | - | - | - | - | - | - | - | - | - | - | 0.444 | 3.75 | 3.963 |
| L | - | - | - | - | - | - | - | - | - | - | - | - | 2.18 | 4.02 |
| M | - | - | - | - | - | - | - | - | - | - | - | - | - | 4.181 |
| N | - | - | - | - | - | - | - | - | - | - | - | - | - | - |

| Max | A | B | C | D | E | F | G | H | I | J | K | L | M | N |
|---|---|---|---|---|---|---|---|---|---|---|---|---|---|---|
| A | - | 3.184 | 4.627 | 6.294 | 7.572 | 3.595 | 4.484 | 4.421 | 3.163 | 7.334 | 4.539 | 1.908 | 5.223 | -1.649 |
| B | - | - | 3.831 | 6.02 | 7.386 | 1.465 | 3.606 | 3.498 | 4.05 | 7.127 | 3.698 | 2.286 | 4.694 | 3.006 |
| C | - | - | - | 5.463 | 7.053 | 3.332 | 0.836 | 1.368 | 4.754 | 6.774 | 0.159 | 4.065 | 3.337 | 4.285 |
| D | - | - | - | - | 5.935 | 5.222 | 4.984 | 5.011 | 5.624 | 5.482 | 4.96 | 5.401 | 4.585 | 5.458 |
| E | - | - | - | - | - | 5.943 | 5.858 | 5.867 | 6.106 | 3.196 | 5.849 | 6.01 | 5.722 | 6.033 |
| F | - | - | - | - | - | - | 3.2 | 3.06 | 4.231 | 7.059 | 3.317 | 2.891 | 4.496 | 3.39 |
| G | - | - | - | - | - | - | - | -0.382 | 4.679 | 6.827 | -0.487 | 3.926 | 3.602 | 4.172 |
| H | - | - | - | - | - | - | - | - | 4.646 | 6.851 | 0.578 | 3.862 | 3.709 | 4.12 |
| I | - | - | - | - | - | - | - | - | - | 7.556 | 5.146 | 3.806 | 5.674 | 3.353 |
| J | - | - | - | - | - | - | - | - | - | - | 5.724 | 5.917 | 5.575 | 5.945 |
| K | - | - | - | - | - | - | - | - | - | - | - | 3.982 | 3.509 | 4.217 |
| L | - | - | - | - | - | - | - | - | - | - | - | - | 5.016 | 1.737 |
| M | - | - | - | - | - | - | - | - | - | - | - | - | - | 4.749 |
| N | - | - | - | - | - | - | - | - | - | - | - | - | - | - |

| | |
|---|---|
| | Larger than 10 |
| | Between 8 and 9,99 |
| | Between 6 and 7,99 |
| | Between 4 and 5,99 |
| | Between 2 and 3,99 |
| | Between 0 and 1,99 |
| | NA (Calculation failed) |
