## Supplementary Data S2 for "Interlaboratory comparison of standardised metabolomics and lipidomics analyses in human and rodent blood using the MxP® Quant 500 kit": TG(14.0_35.2).pdf

**Amount of relative differences between laboratories**

| Min | A | B | C | D | E | F | G | H | I | J | K | L | M | N |
|---|---|---|---|---|---|---|---|---|---|---|---|---|---|---|
| A | - | 6.11 | 4.562 | 4.9 | 6.713 | 5.146 | 5.421 | 5.201 | 3.266 | 7.278 | 3.735 | 5.348 | 6.344 | 4.973 |
| B | - | - | 4.658 | 3.575 | 5.583 | 3.11 | 3.716 | 2.682 | 7.018 | 6.153 | 5.719 | 3.746 | 4.381 | 6.444 |
| C | - | - | - | 3.59 | 5.712 | 4.871 | 4.993 | 4.735 | 6.403 | 6.358 | 4.57 | 3.399 | 1.196 | 5.777 |
| D | - | - | - | - | 5.936 | 3.484 | 4.453 | 2.598 | 6.543 | 6.318 | 5.168 | 1.288 | 3.233 | 5.768 |
| E | - | - | - | - | - | 5.07 | 1.887 | 4.487 | 7.88 | 5.622 | 6.831 | 5.628 | 6.879 | 7.625 |
| F | - | - | - | - | - | - | 0.133 | 3.753 | 7.745 | 5.243 | 6.791 | 5.274 | 6.234 | 7.338 |
| G | - | - | - | - | - | - | - | 3.969 | 7.303 | 6.194 | 6.305 | 4.791 | 4.773 | 6.769 |
| H | - | - | - | - | - | - | - | - | 6.999 | 6.364 | 6.157 | 3.753 | 4.703 | 6.613 |
| I | - | - | - | - | - | - | - | - | - | 6.902 | 4.538 | 5.35 | 6.079 | 4.785 |
| J | - | - | - | - | - | - | - | - | - | - | 7.297 | 6.237 | 7.253 | 7.978 |
| K | - | - | - | - | - | - | - | - | - | - | - | 4.817 | 4.803 | 4.09 |
| L | - | - | - | - | - | - | - | - | - | - | - | - | 4.244 | 5.632 |
| M | - | - | - | - | - | - | - | - | - | - | - | - | - | 5.86 |
| N | - | - | - | - | - | - | - | - | - | - | - | - | - | - |

| Median | A | B | C | D | E | F | G | H | I | J | K | L | M | N |
|---|---|---|---|---|---|---|---|---|---|---|---|---|---|---|
| A | - | 4.635 | 1.303 | 4.003 | 0.784 | 3.881 | 5.002 | 2.599 | 1.287 | 5.571 | 2.955 | 4.892 | 4.479 | 1.975 |
| B | - | - | 4.602 | 2.636 | 4.661 | -2.16 | 3.918 | 3.143 | 4.634 | 2.85 | 5.085 | 3.078 | 2.072 | 4.682 |
| C | - | - | - | 3.964 | 2.351 | 4.4 | 5.03 | 4.195 | -0.027 | 4.799 | 3.127 | 4.749 | 3.794 | 1.628 |
| D | - | - | - | - | 4.505 | -2.403 | 4.325 | 2.236 | 4.186 | 3.184 | 4.797 | 3.863 | 1.797 | 4.162 |
| E | - | - | - | - | - | 5.26 | 4.567 | 5.276 | 3.752 | 5.311 | 5.003 | 3.695 | -3.046 | 4.502 |
| F | - | - | - | - | - | - | 3.356 | 0.978 | 5.1 | 0.946 | 5.727 | 1.704 | 3.43 | 5.224 |
| G | - | - | - | - | - | - | - | 4.733 | 5.582 | 3.315 | 5.929 | 2.873 | 4.431 | 5.6 |
| H | - | - | - | - | - | - | - | - | 4.148 | 4.435 | 5.243 | 3.6 | -1.317 | 4.498 |
| I | - | - | - | - | - | - | - | - | - | 4.714 | 1.365 | 4.895 | 4.143 | 1.46 |
| J | - | - | - | - | - | - | - | - | - | - | 5.935 | 2.108 | 4.309 | 5.601 |
| K | - | - | - | - | - | - | - | - | - | - | - | 5.053 | 4.349 | 2.371 |
| L | - | - | - | - | - | - | - | - | - | - | - | - | 4.014 | 5.322 |
| M | - | - | - | - | - | - | - | - | - | - | - | - | - | 4.272 |
| N | - | - | - | - | - | - | - | - | - | - | - | - | - | - |

| Max | A | B | C | D | E | F | G | H | I | J | K | L | M | N |
|---|---|---|---|---|---|---|---|---|---|---|---|---|---|---|
| A | - | 3.95 | 5.376 | 3.322 | 7.679 | 5.507 | 2.871 | 4.858 | -2.578 | 7.05 | 4.943 | 4.246 | 3.29 | 3.283 |
| B | - | - | 4.515 | 2.181 | 7.365 | 4.7 | 4.296 | 3.536 | 3.723 | 6.639 | 3.755 | 4.894 | 2.278 | 2.275 |
| C | - | - | - | 4.481 | 6.842 | 1.37 | 5.11 | 3.144 | 4.877 | 5.972 | 2.936 | 5.419 | 4.503 | 4.5 |
| D | - | - | - | - | 7.513 | 4.992 | 3.984 | 4.087 | 3.176 | 6.77 | 4.241 | 4.72 | -2.007 | -1.801 |
| E | - | - | - | - | - | 5.702 | 6.119 | 5.842 | 6.069 | 4.567 | 5.837 | 6.204 | 6.005 | 6.004 |
| F | - | - | - | - | - | - | 5.178 | 3.482 | 4.959 | 5.862 | 3.335 | 5.464 | 4.604 | 4.613 |
| G | - | - | - | - | - | - | - | 5.289 | 2.995 | 7.219 | 5.361 | 3.645 | 4.204 | 4.201 |
| H | - | - | - | - | - | - | - | - | 4.478 | 6.32 | 0.426 | 5.212 | 3.887 | 3.892 |
| I | - | - | - | - | - | - | - | - | - | 7.02 | 4.942 | 4.249 | 3.249 | 3.26 |
| J | - | - | - | - | - | - | - | - | - | - | 5.447 | 6.021 | 5.712 | 5.719 |
| K | - | - | - | - | - | - | - | - | - | - | - | 5.25 | 4.025 | 4.019 |
| L | - | - | - | - | - | - | - | - | - | - | - | - | 5.14 | 5.139 |
| M | - | - | - | - | - | - | - | - | - | - | - | - | - | -4.008 |
| N | - | - | - | - | - | - | - | - | - | - | - | - | - | - |

| | |
|---|---|
| | Larger than 10 |
| | Between 8 and 9,99 |
| | Between 6 and 7,99 |
| | Between 4 and 5,99 |
| | Between 2 and 3,99 |
| | Between 0 and 1,99 |
| | NA (Calculation failed) |
