## Supplementary Data S2 for "Interlaboratory comparison of standardised metabolomics and lipidomics analyses in human and rodent blood using the MxP® Quant 500 kit": TG(14.0_36.1).pdf

**Amount of relative differences between laboratories**

| Min | A | B | C | D | E | F | G | H | I | J | K | L | M | N |
|---|---|---|---|---|---|---|---|---|---|---|---|---|---|---|
| A | - | 6.562 | 7.386 | 8.129 | 8.241 | 7.262 | 6.736 | 7.148 | 3.08 | 10.273 | 7.046 | 6.657 | 6.597 | 6.565 |
| B | - | - | 6.589 | 7.302 | 7.793 | 6.745 | 1.994 | 6.229 | 6.418 | 9.995 | 5.583 | 1.459 | 1.499 | 2.214 |
| C | - | - | - | 4.063 | 6.571 | 5.453 | 6.995 | 3.782 | 7.641 | 9.328 | 5.971 | 6.96 | 6.88 | 6.791 |
| D | - | - | - | - | 6.643 | 5.671 | 6.656 | -3.431 | 7.372 | 9.207 | 5.444 | 6.639 | 6.547 | 6.433 |
| E | - | - | - | - | - | 3.702 | 8.008 | 6.602 | 8.358 | 8.754 | 7.401 | 7.997 | 7.923 | 7.833 |
| F | - | - | - | - | - | - | 7.96 | 6.403 | 8.376 | 9.318 | 7.294 | 7.923 | 7.853 | 7.781 |
| G | - | - | - | - | - | - | - | 6.126 | 6.371 | 9.853 | 5.418 | -0.281 | -2.186 | 0.868 |
| H | - | - | - | - | - | - | - | - | 7.415 | 9.305 | 5.462 | 6.646 | 6.561 | 6.463 |
| I | - | - | - | - | - | - | - | - | - | 10.473 | 7.134 | 6.721 | 6.648 | 6.614 |
| J | - | - | - | - | - | - | - | - | - | - | 7.371 | 7.87 | 7.786 | 7.702 |
| K | - | - | - | - | - | - | - | - | - | - | - | 5.672 | 5.59 | 5.459 |
| L | - | - | - | - | - | - | - | - | - | - | - | - | -2.042 | 1.088 |
| M | - | - | - | - | - | - | - | - | - | - | - | - | - | 0.913 |
| N | - | - | - | - | - | - | - | - | - | - | - | - | - | - |

| |
|---|
| Larger than 10 |
| Between 8 and 9,99 |
| Between 6 and 7,99 |
| Between 4 and 5,99 |
| Between 2 and 3,99 |
| Between 0 and 1,99 |
| NA (Calculation failed) |

| Median | A | B | C | D | E | F | G | H | I | J | K | L | M | N |
|---|---|---|---|---|---|---|---|---|---|---|---|---|---|---|
| A | - | 3.522 | 4.485 | 3.968 | 4.82 | 5.279 | 2.887 | 3.637 | 3.491 | -2.852 | 4.012 | 3.722 | 4.008 | 4.258 |
| B | - | - | 3.376 | 0.772 | 4.403 | 4.933 | 2.535 | -0.45 | -6.897 | 3.6 | 1.206 | -0.813 | 2.357 | 3.075 |
| C | - | - | - | 3.496 | 3.992 | 4.576 | 4.044 | 3.093 | 3.081 | 4.477 | 3.315 | 3.326 | 1.767 | -4.589 |
| D | - | - | - | - | 4.606 | 4.975 | 2.233 | 1.249 | 1.353 | 3.408 | 0.219 | -0.117 | 2.785 | 3.338 |
| E | - | - | - | - | - | 2.945 | 5.358 | 4.96 | 4.923 | 5.577 | 5.125 | 5.095 | 4.644 | 4.367 |
| F | - | - | - | - | - | - | 5.734 | 5.406 | 5.388 | 5.89 | 5.51 | 5.488 | 5.15 | 4.957 |
| G | - | - | - | - | - | - | - | 2.638 | 2.482 | 2.695 | 2.901 | 2.579 | 3.379 | 3.756 |
| H | - | - | - | - | - | - | - | - | -5.414 | 3.617 | 0.191 | 0.371 | 2.236 | 2.981 |
| I | - | - | - | - | - | - | - | - | - | 3.493 | 2.534 | 1.237 | 2.462 | 3.165 |
| J | - | - | - | - | - | - | - | - | - | - | 2.253 | 2.745 | 3.827 | 4.08 |
| K | - | - | - | - | - | - | - | - | - | - | - | -0.13 | 2.135 | 2.938 |
| L | - | - | - | - | - | - | - | - | - | - | - | - | 2.206 | 2.987 |
| M | - | - | - | - | - | - | - | - | - | - | - | - | - | 1.807 |
| N | - | - | - | - | - | - | - | - | - | - | - | - | - | - |

| Max | A | B | C | D | E | F | G | H | I | J | K | L | M | N |
|---|---|---|---|---|---|---|---|---|---|---|---|---|---|---|
| A | - | 2.786 | 4.764 | 5.314 | 4.732 | 2.797 | 3.958 | 5.153 | 3.562 | 7.457 | 4.272 | 2.763 | -0.129 | 1.572 |
| B | - | - | 4.248 | 4.956 | 4.243 | -3.311 | 3.01 | 4.747 | 4.132 | 7.31 | 3.542 | -3.779 | 2.487 | 3.208 |
| C | - | - | - | 3.343 | -3.417 | 3.989 | 3.2 | 2.72 | 4.941 | 6.866 | 2.628 | 4.004 | 4.371 | 4.569 |
| D | - | - | - | - | 3.21 | 4.565 | 4.131 | 1.689 | 5.211 | 6.585 | 3.89 | 4.575 | 4.809 | 4.942 |
| E | - | - | - | - | - | 3.997 | 3.175 | 2.724 | 4.938 | 6.855 | 2.61 | 4.018 | 4.382 | 4.568 |
| F | - | - | - | - | - | - | 2.989 | 4.739 | 4.137 | 7.307 | 3.526 | -2.371 | 2.516 | 3.222 |
| G | - | - | - | - | - | - | - | 4.12 | 4.566 | 7.115 | 1.73 | 2.911 | 3.66 | 4.001 |
| H | - | - | - | - | - | - | - | - | 5.129 | 6.697 | 3.583 | 4.408 | 4.678 | 4.83 |
| I | - | - | - | - | - | - | - | - | - | 7.745 | 5.145 | 4.401 | 3.853 | 3.33 |
| J | - | - | - | - | - | - | - | - | - | - | 5.829 | 5.943 | 5.993 | 6.023 |
| K | - | - | - | - | - | - | - | - | - | - | - | 3.391 | 3.95 | 4.225 |
| L | - | - | - | - | - | - | - | - | - | - | - | - | 2.468 | 3.197 |
| M | - | - | - | - | - | - | - | - | - | - | - | - | - | 1.943 |
| N | - | - | - | - | - | - | - | - | - | - | - | - | - | - |
