## Supplementary Data S2 for "Interlaboratory comparison of standardised metabolomics and lipidomics analyses in human and rodent blood using the MxP® Quant 500 kit": TG(16.0_30.2).pdf

**Amount of relative differences between laboratories**

| Min | A | B | C | D | E | F | G | H | I | J | K | L | M | N |
|---|---|---|---|---|---|---|---|---|---|---|---|---|---|---|
| A | - | 6.464 | 5.405 | 5.292 | 6.711 | 5.471 | 5.414 | 5.897 | 5.326 | 8.328 | 5.668 | 5.348 | 5.612 | 5.404 |
| B | - | - | 4.187 | 2.55 | 6.224 | 3.296 | 5.361 | 3.987 | 4.678 | 7.599 | 5.398 | 5.731 | 5.361 | 4.592 |
| C | - | - | - | 4.135 | 6.27 | 4.867 | 4.255 | 1.149 | 2.615 | 7.485 | 4.331 | 4.876 | 4.256 | 2.27 |
| D | - | - | - | - | 6.086 | 3.947 | 5.486 | 4.092 | 4.863 | 7.788 | 5.408 | 5.749 | 5.412 | 4.665 |
| E | - | - | - | - | - | 5.952 | 7.925 | 7.602 | 7.56 | 5.344 | 8.026 | 8.129 | 8.006 | 7.518 |
| F | - | - | - | - | - | - | 6.119 | 5.196 | 5.692 | 7.988 | 6.056 | 6.322 | 6.058 | 5.554 |
| G | - | - | - | - | - | - | - | 4.363 | 2.884 | 7.905 | 1.36 | 2.852 | 1.312 | 3.197 |
| H | - | - | - | - | - | - | - | - | 3.307 | 7.631 | 4.512 | 5.027 | 4.456 | 3.077 |
| I | - | - | - | - | - | - | - | - | - | 8.031 | 2.138 | 3.951 | 2.222 | 1.029 |
| J | - | - | - | - | - | - | - | - | - | - | 6.243 | 6.364 | 6.174 | 5.194 |
| K | - | - | - | - | - | - | - | - | - | - | - | 3.155 | -1.399 | 3.27 |
| L | - | - | - | - | - | - | - | - | - | - | - | - | 3.279 | 3.755 |
| M | - | - | - | - | - | - | - | - | - | - | - | - | - | 2.668 |
| N | - | - | - | - | - | - | - | - | - | - | - | - | - | - |

| | |
|---|---|
| | Larger than 10 |
| | Between 8 and 9,99 |
| | Between 6 and 7,99 |
| | Between 4 and 5,99 |
| | Between 2 and 3,99 |
| | Between 0 and 1,99 |
| | NA (Calculation failed) |

| Median | A | B | C | D | E | F | G | H | I | J | K | L | M | N |
|---|---|---|---|---|---|---|---|---|---|---|---|---|---|---|
| A | - | 4.663 | 3.676 | 4.294 | 5.933 | 4.952 | 3.53 | 4.095 | 4.205 | 3.494 | 2.471 | 3.689 | 3.84 | 4.266 |
| B | - | - | 4.176 | 3.132 | 5.482 | 3.044 | 4.199 | 3.49 | 2.891 | 4.493 | 4.744 | 4.176 | 3.624 | 3.449 |
| C | - | - | - | 3.667 | 5.711 | 4.709 | -2.52 | 2.548 | 3.179 | 1.982 | 2.88 | -5.029 | 2.294 | 2.617 |
| D | - | - | - | - | 5.514 | 3.987 | 3.561 | 1.873 | 1.119 | 2.596 | 4.144 | 3.365 | 2.15 | 2.362 |
| E | - | - | - | - | - | 5.377 | 6.608 | 6.41 | 6.279 | 6.614 | 6.79 | 6.64 | 6.383 | 6.473 |
| F | - | - | - | - | - | - | 5.028 | 4.52 | 4.415 | 3.38 | 5.309 | 4.947 | 4.584 | 4.62 |
| G | - | - | - | - | - | - | - | 2.612 | 3.122 | 1.246 | 2.815 | -0.455 | 2.136 | 2.837 |
| H | - | - | - | - | - | - | - | - | 1.713 | 3.304 | 3.81 | 2.595 | 0.216 | -1.148 |
| I | - | - | - | - | - | - | - | - | - | 3.517 | 4.104 | 3.127 | 2.256 | 0.637 |
| J | - | - | - | - | - | - | - | - | - | - | -2.119 | 2.597 | 3.782 | 3.42 |
| K | - | - | - | - | - | - | - | - | - | - | - | 2.828 | 3.46 | 3.696 |
| L | - | - | - | - | - | - | - | - | - | - | - | - | 1.715 | 2.722 |
| M | - | - | - | - | - | - | - | - | - | - | - | - | - | 1.601 |
| N | - | - | - | - | - | - | - | - | - | - | - | - | - | - |

| Max | A | B | C | D | E | F | G | H | I | J | K | L | M | N |
|---|---|---|---|---|---|---|---|---|---|---|---|---|---|---|
| A | - | 2.584 | 4.642 | 4.456 | 3.134 | 0.493 | 3.22 | 3.839 | 0.406 | 7.177 | 4.575 | 1.768 | 1.786 | 1.887 |
| B | - | - | 4.162 | 3.902 | 1.779 | 2.155 | 1.634 | 2.966 | 2.796 | 7.037 | 4.071 | 1.296 | 1.289 | 1.128 |
| C | - | - | - | 1.236 | 3.672 | 4.236 | 3.646 | 3.093 | 4.396 | 6.586 | -0.123 | 4.109 | 4.107 | 4.088 |
| D | - | - | - | - | 3.331 | 4.077 | 3.373 | 2.639 | 4.256 | 6.673 | 0.687 | 3.923 | 3.921 | 3.903 |
| E | - | - | - | - | - | 2.931 | -3.685 | 2.111 | 3.344 | 6.987 | 3.736 | 2.551 | 2.52 | 2.333 |
| F | - | - | - | - | - | - | 2.973 | 3.68 | 1.425 | 7.164 | 4.474 | 1.037 | 1.056 | 1.223 |
| G | - | - | - | - | - | - | - | 2.193 | 3.286 | 6.956 | 3.73 | 2.43 | 2.427 | 2.36 |
| H | - | - | - | - | - | - | - | - | 3.776 | 6.84 | 3.057 | 3.251 | 3.25 | 3.214 |
| I | - | - | - | - | - | - | - | - | - | 7.217 | 4.672 | 2.262 | 2.273 | 2.358 |
| J | - | - | - | - | - | - | - | - | - | - | 5.63 | 5.855 | 5.856 | 5.85 |
| K | - | - | - | - | - | - | - | - | - | - | - | 4.043 | 4.041 | 4.022 |
| L | - | - | - | - | - | - | - | - | - | - | - | - | -5.797 | -1.936 |
| M | - | - | - | - | - | - | - | - | - | - | - | - | - | -1.904 |
| N | - | - | - | - | - | - | - | - | - | - | - | - | - | - |
