## Supplementary Data S2 for "Interlaboratory comparison of standardised metabolomics and lipidomics analyses in human and rodent blood using the MxP® Quant 500 kit": TG(16.0_32.2).pdf

**Amount of relative differences between laboratories**

| Min | A | B | C | D | E | F | G | H | I | J | K | L | M | N |
|---|---|---|---|---|---|---|---|---|---|---|---|---|---|---|
| A | - | 5.762 | 5.15 | 5.927 | 6.93 | 6.003 | 5.464 | 4.933 | 4.519 | 9.04 | 5.472 | 5.53 | 5.145 | 5.478 |
| B | - | - | 4.882 | 2.941 | 6.363 | 4.262 | 3.414 | 3.957 | 4.586 | 8.665 | 1.177 | 2.166 | 4.134 | 1.894 |
| C | - | - | - | 4.954 | 6.581 | 5.286 | 3.837 | 2.782 | -0.229 | 8.721 | 4.284 | 4.231 | 2.846 | 4.224 |
| D | - | - | - | - | 6.156 | 3.707 | 4.088 | 4.236 | 4.752 | 8.4 | 2.815 | 3.366 | 4.513 | 3.135 |
| E | - | - | - | - | - | 6.197 | 6.942 | 6.811 | 6.959 | 7.488 | 6.638 | 6.793 | 7.009 | 6.695 |
| F | - | - | - | - | - | - | 5.114 | 5.206 | 5.532 | 8.555 | 4.537 | 4.748 | 5.37 | 4.655 |
| G | - | - | - | - | - | - | - | 2.439 | 3.753 | 8.624 | 2.842 | 2.46 | 2.752 | 2.583 |
| H | - | - | - | - | - | - | - | - | 3.021 | 8.77 | 3.724 | 3.61 | -4.313 | 3.621 |
| I | - | - | - | - | - | - | - | - | - | 9.1 | 4.601 | 4.63 | 3.583 | 4.577 |
| J | - | - | - | - | - | - | - | - | - | - | 5.853 | 6.084 | 6.324 | 5.939 |
| K | - | - | - | - | - | - | - | - | - | - | - | 1.163 | 3.904 | 0.478 |
| L | - | - | - | - | - | - | - | - | - | - | - | - | 3.714 | -1.296 |
| M | - | - | - | - | - | - | - | - | - | - | - | - | - | 3.64 |
| N | - | - | - | - | - | - | - | - | - | - | - | - | - | - |

| Median | A | B | C | D | E | F | G | H | I | J | K | L | M | N |
|---|---|---|---|---|---|---|---|---|---|---|---|---|---|---|
| A | - | 4.352 | 3.305 | 4.432 | 5.526 | 4.977 | 3.986 | 3.822 | 3.947 | 5.529 | 4.183 | 4.441 | 3.94 | 4.509 |
| B | - | - | 3.999 | 0.917 | 5.034 | 3.514 | 2.961 | 3.112 | 2.176 | 4.471 | 1.619 | -1.325 | 2.475 | 1.674 |
| C | - | - | - | 3.637 | 5.37 | 4.57 | 2.795 | 2.545 | 3.128 | 5.111 | 3.4 | 3.741 | 3.021 | 3.99 |
| D | - | - | - | - | 5.104 | 3.724 | 2.416 | 2.547 | 0.369 | 4.371 | -1.436 | 0.843 | 1.344 | 2.551 |
| E | - | - | - | - | - | 5.104 | 5.939 | 5.929 | 5.689 | 5.103 | 5.707 | 5.686 | 5.769 | 5.392 |
| F | - | - | - | - | - | - | 4.419 | 4.461 | 4.07 | 3.138 | 3.972 | 3.717 | 4.184 | 3.087 |
| G | - | - | - | - | - | - | - | -0.129 | 1.438 | 4.774 | 2.078 | 2.78 | 0.89 | 3.327 |
| H | - | - | - | - | - | - | - | - | 1.68 | 4.948 | 2.331 | 2.984 | 1.323 | 3.421 |
| I | - | - | - | - | - | - | - | - | - | 5.14 | 1.491 | 2.668 | -1.444 | 3.066 |
| J | - | - | - | - | - | - | - | - | - | - | 2.209 | 2.326 | 2.435 | 1.49 |
| K | - | - | - | - | - | - | - | - | - | - | - | 1.358 | 1.261 | 2.611 |
| L | - | - | - | - | - | - | - | - | - | - | - | - | 2.456 | 1.725 |
| M | - | - | - | - | - | - | - | - | - | - | - | - | - | 3.084 |
| N | - | - | - | - | - | - | - | - | - | - | - | - | - | - |

| Max | A | B | C | D | E | F | G | H | I | J | K | L | M | N |
|---|---|---|---|---|---|---|---|---|---|---|---|---|---|---|
| A | - | 2.394 | 3.419 | 4.096 | 2.697 | 0.563 | 3.41 | 2.495 | 3.104 | 6.943 | 3.228 | 0.65 | -0.168 | 2.093 |
| B | - | - | 2.366 | 3.5 | 0.64 | 1.84 | 2.348 | -1.549 | 3.72 | 6.821 | 1.972 | 1.803 | 2.06 | 3.178 |
| C | - | - | - | 2.548 | 1.861 | 3.055 | -4.034 | 2.196 | 4.125 | 6.672 | 0.246 | 3.04 | 3.149 | 3.756 |
| D | - | - | - | - | 3.137 | 3.742 | 2.482 | 3.299 | 4.46 | 6.509 | 2.73 | 3.734 | 3.798 | 4.193 |
| E | - | - | - | - | - | 2.318 | 1.894 | 0.057 | 3.863 | 6.778 | 1.301 | 2.322 | 2.501 | 3.361 |
| F | - | - | - | - | - | - | 3.169 | 2.024 | 3.318 | 6.914 | 2.96 | -3.397 | -0.699 | 2.504 |
| G | - | - | - | - | - | - | - | 2.178 | 4.121 | 6.673 | 0.169 | 3.029 | 3.14 | 3.751 |
| H | - | - | - | - | - | - | - | - | 3.753 | 6.809 | 1.833 | 1.935 | 2.167 | 3.226 |
| I | - | - | - | - | - | - | - | - | - | 7.187 | 4.302 | 3.478 | 3.376 | 2.256 |
| J | - | - | - | - | - | - | - | - | - | - | 5.679 | 5.775 | 5.785 | 5.84 |
| K | - | - | - | - | - | - | - | - | - | - | - | 2.834 | 2.962 | 3.642 |
| L | - | - | - | - | - | - | - | - | - | - | - | - | -0.513 | 2.528 |
| M | - | - | - | - | - | - | - | - | - | - | - | - | - | 2.348 |
| N | - | - | - | - | - | - | - | - | - | - | - | - | - | - |

| | |
|---|---|
| | Larger than 10 |
| | Between 8 and 9,99 |
| | Between 6 and 7,99 |
| | Between 4 and 5,99 |
| | Between 2 and 3,99 |
| | Between 0 and 1,99 |
| | NA (Calculation failed) |
