## Supplementary Data S2 for "Interlaboratory comparison of standardised metabolomics and lipidomics analyses in human and rodent blood using the MxP® Quant 500 kit": TG(16.0_32.3).pdf

**Amount of relative differences between laboratories**

| Min | A | B | C | D | E | F | G | H | I | J | K | L | M | N |
|---|---|---|---|---|---|---|---|---|---|---|---|---|---|---|
| A | - | 6.075 | 5.586 | 5.737 | 6.764 | 5.54 | 5.555 | 5.65 | 4.641 | 8.67 | 5.589 | 5.442 | 5.071 | 5.587 |
| B | - | - | 4.074 | 3.578 | 5.454 | 1.134 | 3.995 | 2.91 | 4.502 | 8.031 | 0.428 | 4.363 | 4.754 | 0.614 |
| C | - | - | - | 2.312 | 5.909 | 3.782 | -1.239 | 2.905 | 3.341 | 8.214 | 3.538 | 1.959 | 3.476 | 3.521 |
| D | - | - | - | - | 5.644 | 3.65 | 1.866 | 1.976 | 3.272 | 7.944 | 3.225 | 2.964 | 3.759 | 3.222 |
| E | - | - | - | - | - | 5.013 | 5.969 | 5.668 | 5.942 | 7.558 | 5.295 | 6.116 | 6.212 | 5.29 |
| F | - | - | - | - | - | - | 4.146 | 3.213 | 4.696 | 8.143 | 1.721 | 4.511 | 4.884 | 1.716 |
| G | - | - | - | - | - | - | - | 2.82 | 3.531 | 8.252 | 3.451 | 2.114 | 3.586 | 3.439 |
| H | - | - | - | - | - | - | - | - | 4.157 | 8.228 | 2.235 | 3.61 | 4.295 | 2.193 |
| I | - | - | - | - | - | - | - | - | - | 8.713 | 4.721 | 4.028 | 2.605 | 4.71 |
| J | - | - | - | - | - | - | - | - | - | - | 5.199 | 6.135 | 6.124 | 5.158 |
| K | - | - | - | - | - | - | - | - | - | - | - | 4.231 | 4.661 | -4.238 |
| L | - | - | - | - | - | - | - | - | - | - | - | - | 3.004 | 3.804 |
| M | - | - | - | - | - | - | - | - | - | - | - | - | - | 4.362 |
| N | - | - | - | - | - | - | - | - | - | - | - | - | - | - |

| Median | A | B | C | D | E | F | G | H | I | J | K | L | M | N |
|---|---|---|---|---|---|---|---|---|---|---|---|---|---|---|
| A | - | 4.661 | 4.249 | 4.187 | 5.094 | 4.753 | 4.463 | 4.524 | 4.231 | 5.717 | 4.625 | 4.351 | 3.802 | 4.607 |
| B | - | - | 2.866 | 3.547 | 4.135 | 1.767 | 2.581 | -0.045 | 2.272 | 4.308 | -0.073 | 2.813 | 3.577 | 0.368 |
| C | - | - | - | 1.799 | 4.476 | 3.307 | 0.786 | 2.511 | 0.782 | 4.841 | 2.617 | -0.864 | 2.237 | 2.953 |
| D | - | - | - | - | 4.709 | 3.755 | 2.346 | 3.261 | 2.62 | 4.827 | 3.259 | 2.051 | -2.006 | 3.541 |
| E | - | - | - | - | - | 4.254 | 4.993 | 4.598 | 4.817 | 2.913 | 4.65 | 5.049 | 5.227 | 4.392 |
| F | - | - | - | - | - | - | 3.247 | 2.177 | 3.113 | 4.23 | 2.135 | 3.427 | 3.992 | 1.073 |
| G | - | - | - | - | - | - | - | 1.975 | -2.106 | 4.763 | 2.141 | 0.268 | 2.748 | 2.584 |
| H | - | - | - | - | - | - | - | - | 2.044 | 4.585 | -1.607 | 2.475 | 3.459 | 1.114 |
| I | - | - | - | - | - | - | - | - | - | 5.267 | 2.668 | -0.482 | 2.634 | 2.745 |
| J | - | - | - | - | - | - | - | - | - | - | 1.603 | 3.289 | 3.44 | 1.397 |
| K | - | - | - | - | - | - | - | - | - | - | - | 2.577 | 3.448 | 1.17 |
| L | - | - | - | - | - | - | - | - | - | - | - | - | 2.51 | 2.775 |
| M | - | - | - | - | - | - | - | - | - | - | - | - | - | 3.577 |
| N | - | - | - | - | - | - | - | - | - | - | - | - | - | - |

| Max | A | B | C | D | E | F | G | H | I | J | K | L | M | N |
|---|---|---|---|---|---|---|---|---|---|---|---|---|---|---|
| A | - | 3.601 | 3.283 | 4.635 | 3.228 | 2.597 | 2.637 | 2.47 | 3.35 | 6.973 | 3.643 | 1.983 | 2.871 | 2.722 |
| B | - | - | 1.115 | 3.508 | 1.162 | 2.445 | 2.404 | 2.562 | 4.317 | 6.667 | -1.62 | 2.864 | 2.118 | 2.316 |
| C | - | - | - | 3.789 | -2.637 | 1.754 | 1.677 | 1.934 | 4.185 | 6.729 | 1.346 | 2.389 | 1.15 | 1.517 |
| D | - | - | - | - | 3.612 | 3.917 | 3.905 | 3.956 | 4.813 | 6.341 | 3.313 | 4.067 | 3.819 | 3.875 |
| E | - | - | - | - | - | 1.683 | 1.6 | 1.899 | 4.176 | 6.728 | 1.425 | 2.38 | 1.135 | 1.429 |
| F | - | - | - | - | - | - | -2.366 | -1.103 | 3.938 | 6.83 | 2.615 | 0.969 | 0.253 | -0.842 |
| G | - | - | - | - | - | - | - | -0.633 | 3.954 | 6.812 | 2.564 | 1.069 | 0.01 | -1.531 |
| H | - | - | - | - | - | - | - | - | 3.9 | 6.84 | 2.723 | 0.548 | 0.737 | -0.02 |
| I | - | - | - | - | - | - | - | - | - | 7.257 | 4.664 | 3.978 | 4.285 | 4.229 |
| J | - | - | - | - | - | - | - | - | - | - | 5.653 | 5.758 | 5.722 | 5.722 |
| K | - | - | - | - | - | - | - | - | - | - | - | 2.92 | 2.216 | 2.395 |
| L | - | - | - | - | - | - | - | - | - | - | - | - | 1.671 | 1.308 |
| M | - | - | - | - | - | - | - | - | - | - | - | - | - | -0.591 |
| N | - | - | - | - | - | - | - | - | - | - | - | - | - | - |

| | |
|---|---|
| | Larger than 10 |
| | Between 8 and 9,99 |
| | Between 6 and 7,99 |
| | Between 4 and 5,99 |
| | Between 2 and 3,99 |
| | Between 0 and 1,99 |
| | NA (Calculation failed) |
