## Supplementary Data S2 for "Interlaboratory comparison of standardised metabolomics and lipidomics analyses in human and rodent blood using the MxP® Quant 500 kit": TG(16.0_34.2).pdf

**Amount of relative differences between laboratories**

| Min | A | B | C | D | E | F | G | H | I | J | K | L | M | N |
|---|---|---|---|---|---|---|---|---|---|---|---|---|---|---|
| A | - | 4.86 | 4.229 | 5.094 | 5.778 | 4.685 | 4.32 | 4.245 | 3.993 | 7.995 | 4.326 | 4.832 | 4.597 | 5.082 |
| B | - | - | 3.713 | 1.961 | 4.925 | 1.553 | 3.578 | 3.598 | 4.591 | 7.516 | 3.434 | 1.384 | 2.518 | 1.958 |
| C | - | - | - | 3.924 | 5.274 | 3.168 | 0.096 | -0.201 | 3.237 | 7.468 | 0.997 | 3.233 | 2.711 | 3.133 |
| D | - | - | - | - | 4.687 | 2.085 | 3.793 | 3.753 | 4.874 | 6.87 | 3.616 | 2.516 | 2.914 | 3.463 |
| E | - | - | - | - | - | 5.284 | 5.696 | 5.679 | 6.195 | 6.719 | 5.633 | 5.374 | 5.441 | 5.633 |
| F | - | - | - | - | - | - | 3.077 | 3.165 | 4.198 | 7.787 | 2.94 | 0.419 | 1.328 | 1.506 |
| G | - | - | - | - | - | - | - | -1.669 | 3.321 | 7.577 | -0.25 | 3.124 | 2.463 | 3.144 |
| H | - | - | - | - | - | - | - | - | 3.11 | 7.714 | 0.257 | 3.245 | 2.569 | 3.426 |
| I | - | - | - | - | - | - | - | - | - | 8.135 | 2.454 | 3.925 | 3.365 | 4.396 |
| J | - | - | - | - | - | - | - | - | - | - | 4.292 | 3.963 | 3.893 | 4.834 |
| K | - | - | - | - | - | - | - | - | - | - | - | 3.033 | 2.236 | 3.165 |
| L | - | - | - | - | - | - | - | - | - | - | - | - | 1.727 | -1.117 |
| M | - | - | - | - | - | - | - | - | - | - | - | - | - | 1.708 |
| N | - | - | - | - | - | - | - | - | - | - | - | - | - | - |

| Median | A | B | C | D | E | F | G | H | I | J | K | L | M | N |
|---|---|---|---|---|---|---|---|---|---|---|---|---|---|---|
| A | - | 3.88 | 2.47 | 3.302 | 4.963 | 4.175 | 2.963 | 3.301 | 3.837 | 4.138 | 3.239 | 3.806 | 3.671 | 4.154 |
| B | - | - | 3.589 | 2.794 | 4.203 | 0.944 | 3.11 | 2.705 | -1.692 | 2.022 | 2.603 | -1.081 | 1.709 | 2.024 |
| C | - | - | - | 2.178 | 4.762 | 3.646 | 1.582 | 2.296 | 3.445 | 2.776 | 2.389 | 3.353 | 2.966 | 3.837 |
| D | - | - | - | - | 4.567 | 2.993 | 0.012 | -0.355 | 3.003 | 0.767 | 0.774 | 2.809 | 1.878 | 3.499 |
| E | - | - | - | - | - | 4.372 | 5.007 | 4.902 | 4.332 | 5.102 | 4.833 | 4.432 | 4.708 | 3.994 |
| F | - | - | - | - | - | - | 3.509 | 3.186 | 1.433 | 2.07 | 3.168 | 1.739 | 2.516 | 0.402 |
| G | - | - | - | - | - | - | - | 1.007 | 2.999 | 2.266 | 1.165 | 2.882 | 2.328 | 3.528 |
| H | - | - | - | - | - | - | - | - | 2.557 | 2.146 | -3.198 | 2.407 | 1.633 | 3.256 |
| I | - | - | - | - | - | - | - | - | - | 2.268 | 2.51 | -1.423 | 0.957 | 2.039 |
| J | - | - | - | - | - | - | - | - | - | - | 3.624 | 4.12 | 3.701 | 4.42 |
| K | - | - | - | - | - | - | - | - | - | - | - | 2.383 | 1.54 | 3.244 |
| L | - | - | - | - | - | - | - | - | - | - | - | - | 1.397 | 2.169 |
| M | - | - | - | - | - | - | - | - | - | - | - | - | - | 2.791 |
| N | - | - | - | - | - | - | - | - | - | - | - | - | - | - |

| Max | A | B | C | D | E | F | G | H | I | J | K | L | M | N |
|---|---|---|---|---|---|---|---|---|---|---|---|---|---|---|
| A | - | 2.143 | 3.926 | 4.909 | 1.601 | 2.523 | 3.13 | 1.583 | 3.721 | 6.953 | 2.12 | 1.252 | 2.028 | 2.573 |
| B | - | - | 3.371 | 4.627 | 0.169 | 3.284 | 2.052 | 0.451 | 4.077 | 6.852 | -3.737 | 0.957 | -1.577 | 3.311 |
| C | - | - | - | 3.698 | 3.385 | 4.186 | 2.489 | 3.407 | 4.625 | 6.562 | 3.24 | 3.478 | 3.274 | 4.199 |
| D | - | - | - | - | 4.372 | 4.786 | 4.042 | 4.385 | 5.059 | 6.178 | 4.312 | 4.419 | 4.33 | 4.795 |
| E | - | - | - | - | - | 3.116 | 2.43 | -2.24 | 3.991 | 6.877 | 0.135 | -0.196 | -0.397 | 3.15 |
| F | - | - | - | - | - | - | 3.943 | 3.214 | 2.986 | 7.111 | 3.419 | 3.108 | 3.381 | -2.307 |
| G | - | - | - | - | - | - | - | 2.403 | 4.336 | 6.728 | 2.022 | 2.547 | 2.104 | 3.757 |
| H | - | - | - | - | - | - | - | - | 3.976 | 6.883 | 0.381 | -0.773 | 0.068 | 3.118 |
| I | - | - | - | - | - | - | - | - | - | 7.319 | 4.337 | 4.166 | 4.314 | 3.066 |
| J | - | - | - | - | - | - | - | - | - | - | 5.749 | 5.772 | 5.751 | 5.863 |
| K | - | - | - | - | - | - | - | - | - | - | - | 0.899 | -1.902 | 3.301 |
| L | - | - | - | - | - | - | - | - | - | - | - | - | 0.714 | 3.027 |
| M | - | - | - | - | - | - | - | - | - | - | - | - | - | 3.265 |
| N | - | - | - | - | - | - | - | - | - | - | - | - | - | - |

| Colour | Legend |
|---|---|
| (red) | Larger than 10 |
| (orange) | Between 8 and 9,99 |
| (light orange) | Between 6 and 7,99 |
| (light yellow-green) | Between 4 and 5,99 |
| (light green) | Between 2 and 3,99 |
| (dark green) | Between 0 and 1,99 |
| (grey) | NA (Calculation failed) |
