## Supplementary Data S2 for "Interlaboratory comparison of standardised metabolomics and lipidomics analyses in human and rodent blood using the MxP® Quant 500 kit": TG(16.0_34.4).pdf

**Amount of relative differences between laboratories**

| Min | A | B | C | D | E | F | G | H | I | J | K | L | M | N |
|---|---|---|---|---|---|---|---|---|---|---|---|---|---|---|
| A | - | 4.658 | 4.287 | 4.947 | 5.92 | 5.649 | 4.212 | 4.571 | 4.217 | 7.569 | 5.319 | 4.612 | 4.697 | 4.996 |
| B | - | - | 2.981 | 0.132 | 5.239 | 4.967 | 3.087 | -0.422 | 3.688 | 7.154 | 4.143 | -0.994 | 1.531 | 1.88 |
| C | - | - | - | 2.939 | 5.368 | 5.174 | -1.017 | 2.761 | 2.324 | 7.014 | 4.556 | 2.792 | 3.356 | 3.35 |
| D | - | - | - | - | 5.148 | 4.962 | 2.927 | -1.67 | 3.706 | 6.884 | 4.151 | -1.523 | 2.051 | 1.558 |
| E | - | - | - | - | - | 1.027 | 5.831 | 5.497 | 6.07 | 6.345 | 4.418 | 5.515 | 5.293 | 5.484 |
| F | - | - | - | - | - | - | 5.78 | 5.438 | 6.008 | 6.617 | 4.264 | 5.445 | 5.231 | 5.381 |
| G | - | - | - | - | - | - | - | 2.829 | 1.889 | 7.155 | 4.6 | 2.889 | 3.378 | 3.542 |
| H | - | - | - | - | - | - | - | - | 3.648 | 7.066 | 4.171 | -2.316 | 1.884 | 1.927 |
| I | - | - | - | - | - | - | - | - | - | 7.556 | 4.729 | 3.097 | 3.417 | 3.993 |
| J | - | - | - | - | - | - | - | - | - | - | 0.666 | 4.215 | 3.537 | 4.581 |
| K | - | - | - | - | - | - | - | - | - | - | - | 4.382 | 4.01 | 4.189 |
| L | - | - | - | - | - | - | - | - | - | - | - | - | 1.781 | 1.967 |
| M | - | - | - | - | - | - | - | - | - | - | - | - | - | 0.997 |
| N | - | - | - | - | - | - | - | - | - | - | - | - | - | - |

| | |
|---|---|
| | Larger than 10 |
| | Between 8 and 9,99 |
| | Between 6 and 7,99 |
| | Between 4 and 5,99 |
| | Between 2 and 3,99 |
| | Between 0 and 1,99 |
| | NA (Calculation failed) |

| Median | A | B | C | D | E | F | G | H | I | J | K | L | M | N |
|---|---|---|---|---|---|---|---|---|---|---|---|---|---|---|
| A | - | 3.933 | 2.873 | 3.445 | 5.211 | 5.28 | 3.192 | 3.819 | 3.822 | 5.154 | 4.536 | 3.792 | 4.26 | 4.32 |
| B | - | - | 3.265 | 2.24 | 4.658 | 4.706 | 2.889 | 0.884 | -3.39 | 3.964 | 3.562 | 0.103 | 2.195 | 3.041 |
| C | - | - | - | 2.185 | 4.989 | 5.018 | 1.009 | 2.818 | 3.149 | 4.424 | 4.313 | 2.979 | 3.698 | 4.057 |
| D | - | - | - | - | 4.824 | 4.847 | 1.459 | 1.311 | 2.245 | 3.983 | 4.042 | 1.825 | 3.134 | 3.713 |
| E | - | - | - | - | - | -3.619 | 5.413 | 5.157 | 4.969 | 4.549 | 3.849 | 5.072 | 4.736 | 4.242 |
| F | - | - | - | - | - | - | 5.436 | 5.177 | 5.014 | 4.408 | 3.949 | 5.099 | 4.765 | 4.325 |
| G | - | - | - | - | - | - | - | 2.383 | 2.743 | 4.415 | 4.143 | 2.55 | 3.48 | 3.841 |
| H | - | - | - | - | - | - | - | - | 0.927 | 3.999 | 3.777 | -0.184 | 2.669 | 3.354 |
| I | - | - | - | - | - | - | - | - | - | 4.511 | 3.453 | -1.141 | 2.465 | 2.861 |
| J | - | - | - | - | - | - | - | - | - | - | 4.437 | 2.494 | 3.313 | 4.237 |
| K | - | - | - | - | - | - | - | - | - | - | - | 3.973 | 3.245 | 1.997 |
| L | - | - | - | - | - | - | - | - | - | - | - | - | 2.477 | 3.218 |
| M | - | - | - | - | - | - | - | - | - | - | - | - | - | 1.743 |
| N | - | - | - | - | - | - | - | - | - | - | - | - | - | - |

| Max | A | B | C | D | E | F | G | H | I | J | K | L | M | N |
|---|---|---|---|---|---|---|---|---|---|---|---|---|---|---|
| A | - | 1.875 | 4.103 | 4.076 | 2.352 | 3.256 | 3.193 | 3.037 | 3.268 | 6.814 | -1.931 | 2.55 | 2.255 | 3.194 |
| B | - | - | 3.707 | 3.68 | 3.043 | 3.668 | 2.401 | 2.133 | 3.684 | 6.73 | 1.917 | 1.096 | 3.027 | 3.627 |
| C | - | - | - | -2.284 | 4.234 | 4.511 | 2.784 | 2.941 | 4.519 | 6.358 | 3.896 | 3.273 | 4.23 | 4.49 |
| D | - | - | - | - | 4.221 | 4.501 | 2.745 | 2.905 | 4.507 | 6.364 | 3.878 | 3.245 | 4.217 | 4.48 |
| E | - | - | - | - | - | 2.314 | 3.883 | 3.779 | 2.311 | 6.954 | 2.28 | 3.486 | -4.425 | 2.168 |
| F | - | - | - | - | - | - | 4.366 | 4.291 | -2.891 | 7.086 | 3.363 | 4.088 | 2.384 | -1.165 |
| G | - | - | - | - | - | - | - | -0.239 | 4.106 | 6.57 | 3.107 | 1.572 | 3.673 | 4.066 |
| H | - | - | - | - | - | - | - | - | 4.045 | 6.605 | 2.966 | 1.1 | 3.586 | 4.003 |
| I | - | - | - | - | - | - | - | - | - | 7.087 | 3.378 | 4.104 | 2.431 | -0.811 |
| J | - | - | - | - | - | - | - | - | - | - | 5.743 | 5.662 | 5.796 | 5.848 |
| K | - | - | - | - | - | - | - | - | - | - | - | 2.618 | 2.187 | 3.156 |
| L | - | - | - | - | - | - | - | - | - | - | - | - | 3.333 | 3.827 |
| M | - | - | - | - | - | - | - | - | - | - | - | - | - | 2.193 |
| N | - | - | - | - | - | - | - | - | - | - | - | - | - | - |
