## Supplementary Data S2 for "Interlaboratory comparison of standardised metabolomics and lipidomics analyses in human and rodent blood using the MxP® Quant 500 kit": TG(16.0_35.1).pdf

**Amount of relative differences between laboratories**

| Min | A | B | C | D | E | F | G | H | I | J | K | L | M | N |
|---|---|---|---|---|---|---|---|---|---|---|---|---|---|---|
| A | - | 5.558 | 5.394 | 7.179 | 7.916 | 6.195 | 5.542 | 5.447 | 3.061 | 9.163 | 3.414 | 5.628 | 5.63 | 5.09 |
| B | - | - | -2.459 | 6.731 | 7.716 | 5.235 | 3.81 | 1.638 | 5.678 | 8.945 | 6.596 | 2.185 | 2.294 | 4.453 |
| C | - | - | - | 6.772 | 7.76 | 5.296 | 3.543 | 1.188 | 5.685 | 9.013 | 6.583 | 1.426 | 2.557 | 4.369 |
| D | - | - | - | - | 5.912 | 3.796 | 6.59 | 5.915 | 6.845 | 6.481 | 7.644 | 6.313 | 5.714 | 6.492 |
| E | - | - | - | - | - | 5.324 | 7.112 | 6.511 | 7.245 | 5.894 | 8.006 | 6.873 | 6.366 | 6.986 |
| F | - | - | - | - | - | - | 6.277 | 5.713 | 6.835 | 8.594 | 7.553 | 5.942 | 5.402 | 6.332 |
| G | - | - | - | - | - | - | - | 3.034 | 5.178 | 8.9 | 6.246 | 3.024 | 3.966 | 3.066 |
| H | - | - | - | - | - | - | - | - | 5.575 | 8.969 | 6.511 | -0.867 | 2.999 | 4.172 |
| I | - | - | - | - | - | - | - | - | - | 9.236 | 4.444 | 5.442 | 5.466 | 4.765 |
| J | - | - | - | - | - | - | - | - | - | - | 7.527 | 6.164 | 5.481 | 6.344 |
| K | - | - | - | - | - | - | - | - | - | - | - | 5.543 | 5.688 | 5.249 |
| L | - | - | - | - | - | - | - | - | - | - | - | - | 3.098 | 4.114 |
| M | - | - | - | - | - | - | - | - | - | - | - | - | - | 4.8 |
| N | - | - | - | - | - | - | - | - | - | - | - | - | - | - |

| | |
|---|---|
| | Larger than 10 |
| | Between 8 and 9,99 |
| | Between 6 and 7,99 |
| | Between 4 and 5,99 |
| | Between 2 and 3,99 |
| | Between 0 and 1,99 |
| | NA (Calculation failed) |

| Median | A | B | C | D | E | F | G | H | I | J | K | L | M | N |
|---|---|---|---|---|---|---|---|---|---|---|---|---|---|---|
| A | - | 4.368 | 4.668 | 5.007 | 4.618 | 5.289 | 3.996 | 4.431 | 2.571 | 3.141 | 5.88 | 4.37 | 4.475 | 3.8 |
| B | - | - | 1.436 | 3.586 | 2.948 | 4.361 | 2.896 | -2.226 | 4.283 | 3.738 | 6.628 | 0.418 | 1.41 | 3.136 |
| C | - | - | - | 3.076 | 1.694 | 4.198 | 3.413 | 1.683 | 4.533 | 4.222 | 6.72 | 2.133 | -0.946 | 3.603 |
| D | - | - | - | - | 1.869 | 3.809 | 3.58 | 2.155 | 4.531 | 4.238 | 6.65 | 2.654 | 0.668 | 3.466 |
| E | - | - | - | - | - | 3.059 | 4.135 | 3.32 | 4.877 | 4.488 | 6.761 | 3.554 | 1.09 | 3.995 |
| F | - | - | - | - | - | - | 5.057 | 4.603 | 5.571 | 5.375 | 7.164 | 4.685 | 4.262 | 5.082 |
| G | - | - | - | - | - | - | - | 2.75 | 3.5 | 2.924 | 6.422 | 2.511 | 3.192 | 0.488 |
| H | - | - | - | - | - | - | - | - | 4.268 | 3.857 | 6.627 | 0.146 | 1.426 | 3.11 |
| I | - | - | - | - | - | - | - | - | - | 2.255 | 6.103 | 3.97 | 4.127 | 3.098 |
| J | - | - | - | - | - | - | - | - | - | - | 6.143 | 3.159 | 4.044 | 2.307 |
| K | - | - | - | - | - | - | - | - | - | - | - | 5.615 | 5.674 | 5.501 |
| L | - | - | - | - | - | - | - | - | - | - | - | - | 1.916 | 2.897 |
| M | - | - | - | - | - | - | - | - | - | - | - | - | - | 3.557 |
| N | - | - | - | - | - | - | - | - | - | - | - | - | - | - |

| Max | A | B | C | D | E | F | G | H | I | J | K | L | M | N |
|---|---|---|---|---|---|---|---|---|---|---|---|---|---|---|
| A | - | 0.754 | 2.903 | 5.928 | 6.824 | 2.424 | 0.893 | 1.924 | 2.093 | 7.15 | 6.603 | 1.277 | 1.471 | 0.261 |
| B | - | - | 2.542 | 6.039 | 6.919 | 1.866 | 1.856 | 1.087 | 1.403 | 7.215 | 6.652 | -0.515 | 0.074 | -1.031 |
| C | - | - | - | 6.225 | 7.048 | 1.214 | 3.331 | 1.986 | 1.769 | 7.337 | 6.82 | 2.449 | 2.348 | 2.758 |
| D | - | - | - | - | 5.013 | 5.379 | 5.213 | 5.344 | 5.355 | 5.578 | 4.418 | 5.326 | 5.328 | 5.283 |
| E | - | - | - | - | - | 5.812 | 5.719 | 5.795 | 5.805 | 3.649 | 3.248 | 5.784 | 5.786 | 5.759 |
| F | - | - | - | - | - | - | 2.92 | 0.68 | 0.088 | 7.284 | 6.757 | 1.621 | 1.438 | 2.122 |
| G | - | - | - | - | - | - | - | 2.471 | 2.597 | 7.101 | 6.547 | 2.057 | 2.172 | 1.59 |
| H | - | - | - | - | - | - | - | - | -0.939 | 7.251 | 6.713 | 0.546 | 0.136 | 1.429 |
| I | - | - | - | - | - | - | - | - | - | 7.276 | 6.728 | 0.969 | 0.677 | 1.688 |
| J | - | - | - | - | - | - | - | - | - | - | 4.293 | 5.908 | 5.91 | 5.894 |
| K | - | - | - | - | - | - | - | - | - | - | - | 5.658 | 5.663 | 5.641 |
| L | - | - | - | - | - | - | - | - | - | - | - | - | -1.474 | 0.267 |
| M | - | - | - | - | - | - | - | - | - | - | - | - | - | 0.668 |
| N | - | - | - | - | - | - | - | - | - | - | - | - | - | - |
