## Supplementary Data S2 for "Interlaboratory comparison of standardised metabolomics and lipidomics analyses in human and rodent blood using the MxP® Quant 500 kit": TG(16.0_36.3).pdf

**Amount of relative differences between laboratories**

| Min | A | B | C | D | E | F | G | H | I | J | K | L | M | N |
|---|---|---|---|---|---|---|---|---|---|---|---|---|---|---|
| A | - | 4.64 | 4.107 | 4.27 | 5.499 | 4.813 | 4.036 | 4.227 | 3.647 | 7.226 | 4.157 | 4.636 | 4.348 | 4.849 |
| B | - | - | 3.318 | 2.885 | 4.559 | 2.536 | 3.507 | 3.06 | 4.479 | 6.804 | 3.064 | 1.372 | 2.482 | 2.426 |
| C | - | - | - | 1.191 | 4.925 | 3.843 | 0.346 | 0.588 | 3.495 | 6.741 | 0.538 | 2.725 | 2 | 2.088 |
| D | - | - | - | - | 4.811 | 3.655 | 1.86 | -0.083 | 3.905 | 6.682 | -0.643 | 1.977 | 0.792 | -1.833 |
| E | - | - | - | - | - | 4.302 | 5.401 | 5.277 | 5.885 | 6.438 | 5.247 | 5.036 | 5.124 | 5.342 |
| F | - | - | - | - | - | - | 4.184 | 3.91 | 4.896 | 6.793 | 3.907 | 3.174 | 3.598 | 3.549 |
| G | - | - | - | - | - | - | - | 1.431 | 3.283 | 6.76 | 1.398 | 2.984 | 2.363 | 2.508 |
| H | - | - | - | - | - | - | - | - | 3.437 | 7.045 | -1.353 | 2.618 | 1.2 | 2.694 |
| I | - | - | - | - | - | - | - | - | - | 7.498 | 2.409 | 3.789 | 3.03 | 4.288 |
| J | - | - | - | - | - | - | - | - | - | - | 3.771 | 3.828 | 3.476 | 4.888 |
| K | - | - | - | - | - | - | - | - | - | - | - | 2.52 | 1.35 | 2.055 |
| L | - | - | - | - | - | - | - | - | - | - | - | - | 1.551 | 1.461 |
| M | - | - | - | - | - | - | - | - | - | - | - | - | - | 1.122 |
| N | - | - | - | - | - | - | - | - | - | - | - | - | - | - |

| Median | A | B | C | D | E | F | G | H | I | J | K | L | M | N |
|---|---|---|---|---|---|---|---|---|---|---|---|---|---|---|
| A | - | 3.631 | 1.842 | 1.648 | 4.758 | 4.136 | 1.568 | 3.47 | 3.879 | 3.824 | 2.849 | 3.359 | 3.379 | 3.888 |
| B | - | - | 3.327 | 3.501 | 4.02 | 2.51 | 3.426 | 1.184 | 1.394 | 4.921 | 2.666 | 1.358 | 1.549 | 1.199 |
| C | - | - | - | 0.25 | 4.575 | 3.837 | -0.625 | 2.818 | 3.524 | 4.225 | 1.741 | 2.76 | 2.691 | 3.487 |
| D | - | - | - | - | 4.629 | 3.944 | -0.472 | 3.019 | 3.667 | 4.051 | 2.186 | 3.001 | 2.916 | 3.619 |
| E | - | - | - | - | - | 3.617 | 4.983 | 4.467 | 3.976 | 5.776 | 4.739 | 4.463 | 4.511 | 4.04 |
| F | - | - | - | - | - | - | 4.123 | 3.08 | 1.697 | 5.257 | 3.674 | 3.133 | 3.194 | 1.84 |
| G | - | - | - | - | - | - | - | 2.94 | 3.593 | 4.166 | 1.993 | 2.885 | 2.823 | 3.56 |
| H | - | - | - | - | - | - | - | - | 2.174 | 4.787 | 2.047 | -0.864 | -0.517 | 2.126 |
| I | - | - | - | - | - | - | - | - | - | 5.022 | 3.132 | 2.383 | 2.325 | -2.492 |
| J | - | - | - | - | - | - | - | - | - | - | 4.188 | 4.442 | 4.355 | 4.644 |
| K | - | - | - | - | - | - | - | - | - | - | - | 1.816 | 1.694 | 3.026 |
| L | - | - | - | - | - | - | - | - | - | - | - | - | -1.505 | 2.243 |
| M | - | - | - | - | - | - | - | - | - | - | - | - | - | 2.342 |
| N | - | - | - | - | - | - | - | - | - | - | - | - | - | - |

| Max | A | B | C | D | E | F | G | H | I | J | K | L | M | N |
|---|---|---|---|---|---|---|---|---|---|---|---|---|---|---|
| A | - | -2.469 | 3.307 | 3.953 | 3.277 | 2.537 | 3.324 | 1.91 | 3.988 | 6.692 | 2.03 | 1.201 | -6.625 | 2.776 |
| B | - | - | 3.334 | 3.987 | 3.199 | 2.481 | 3.343 | 1.825 | 3.98 | 6.75 | 2.074 | 1.346 | -2.831 | 2.733 |
| C | - | - | - | 2.376 | 4.136 | 3.831 | -4.394 | 3.629 | 4.555 | 6.445 | 2.42 | 2.789 | 3.174 | 3.923 |
| D | - | - | - | - | 4.431 | 4.211 | 2.299 | 4.069 | 4.771 | 6.349 | 3.331 | 3.524 | 3.769 | 4.28 |
| E | - | - | - | - | - | 1.976 | 4.416 | 2.616 | 2.837 | 7.043 | 3.88 | 3.679 | 3.345 | 1.472 |
| F | - | - | - | - | - | - | 4.061 | 1.109 | 3.427 | 6.901 | 3.378 | 3.097 | 2.598 | 0.046 |
| G | - | - | - | - | - | - | - | 3.637 | 4.559 | 6.439 | 2.435 | 2.794 | 3.186 | 3.931 |
| H | - | - | - | - | - | - | - | - | 3.656 | 6.84 | 3.019 | 2.655 | 1.938 | 1.664 |
| I | - | - | - | - | - | - | - | - | - | 7.216 | 4.567 | 4.432 | 4.232 | 3.433 |
| J | - | - | - | - | - | - | - | - | - | - | 5.702 | 5.718 | 5.742 | 5.807 |
| K | - | - | - | - | - | - | - | - | - | - | - | 0.692 | 1.983 | 3.38 |
| L | - | - | - | - | - | - | - | - | - | - | - | - | 1.229 | 3.167 |
| M | - | - | - | - | - | - | - | - | - | - | - | - | - | 2.746 |
| N | - | - | - | - | - | - | - | - | - | - | - | - | - | - |

| | |
|---|---|
| | Larger than 10 |
| | Between 8 and 9,99 |
| | Between 6 and 7,99 |
| | Between 4 and 5,99 |
| | Between 2 and 3,99 |
| | Between 0 and 1,99 |
| | NA (Calculation failed) |
