## Supplementary Data S2 for "Interlaboratory comparison of standardised metabolomics and lipidomics analyses in human and rodent blood using the MxP® Quant 500 kit": TG(16.0_36.4).pdf

**Amount of relative differences between laboratories**

| Min | A | B | C | D | E | F | G | H | I | J | K | L | M | N |
|---|---|---|---|---|---|---|---|---|---|---|---|---|---|---|
| A | - | 4.365 | 3.917 | 3.379 | 5.055 | 5.063 | 3.797 | 4.398 | 3.807 | 5.674 | 3.852 | 3.719 | 4.258 | 4.242 |
| B | - | - | 2.703 | 3.647 | 4.139 | 3.745 | 3.066 | -1.707 | 3.726 | 4.81 | 2.576 | 2.945 | 1.027 | 2.768 |
| C | - | - | - | 2.515 | 4.496 | 4.138 | 0.883 | 2.432 | 3.002 | 4.861 | -0.15 | 0.004 | 2.014 | 0.697 |
| D | - | - | - | - | 4.776 | 4.546 | 1.915 | 3.445 | -1.196 | 5.147 | 2.532 | 2.195 | 3.195 | 2.509 |
| E | - | - | - | - | - | 2.2 | 4.988 | 4.478 | 5.231 | 3.168 | 4.869 | 4.982 | 4.578 | 4.873 |
| F | - | - | - | - | - | - | 4.546 | 3.877 | 4.972 | 3.404 | 4.339 | 4.509 | 4.023 | 4.542 |
| G | - | - | - | - | - | - | - | 2.874 | 2.367 | 4.978 | 1.287 | -0.248 | 2.536 | -0.168 |
| H | - | - | - | - | - | - | - | - | 3.656 | 4.811 | 2.522 | 2.886 | 0.821 | 2.621 |
| I | - | - | - | - | - | - | - | - | - | 5.56 | 1.075 | -0.41 | 2.783 | 2.606 |
| J | - | - | - | - | - | - | - | - | - | - | 3.372 | 3.67 | 3.242 | 4.55 |
| K | - | - | - | - | - | - | - | - | - | - | - | 0.476 | 1.974 | -7.257 |
| L | - | - | - | - | - | - | - | - | - | - | - | - | 2.547 | 1.48 |
| M | - | - | - | - | - | - | - | - | - | - | - | - | - | 2.149 |
| N | - | - | - | - | - | - | - | - | - | - | - | - | - | - |

| Median | A | B | C | D | E | F | G | H | I | J | K | L | M | N |
|---|---|---|---|---|---|---|---|---|---|---|---|---|---|---|
| A | - | 3.623 | 1.469 | 1.123 | 4.744 | 4.256 | 1.634 | 3.637 | 3.516 | 4.248 | 3.104 | 3.225 | 3.365 | 3.85 |
| B | - | - | 3.413 | 3.457 | 4.079 | 3.007 | 3.319 | -1.177 | -0.938 | 5.047 | 2.234 | 1.64 | 1.322 | 1.292 |
| C | - | - | - | -2.492 | 4.63 | 4.08 | -0.471 | 3.331 | 3.214 | 4.273 | 2.344 | 2.749 | 2.852 | 3.57 |
| D | - | - | - | - | 4.667 | 4.1 | -0.114 | 3.363 | 3.224 | 4.357 | 2.545 | 2.862 | 2.937 | 3.605 |
| E | - | - | - | - | - | 3.376 | 5.014 | 4.312 | 4.379 | 5.914 | 4.719 | 4.608 | 4.545 | 4.101 |
| F | - | - | - | - | - | - | 4.283 | 3.045 | 3.205 | 5.459 | 3.851 | 3.638 | 3.549 | 2.598 |
| G | - | - | - | - | - | - | - | 3.241 | 3.102 | 4.383 | 2.222 | 2.637 | 2.744 | 3.492 |
| H | - | - | - | - | - | - | - | - | -0.037 | 5.086 | 2.346 | 1.816 | 1.544 | 1.04 |
| I | - | - | - | - | - | - | - | - | - | 5.043 | 1.696 | 1.162 | 0.467 | 1.755 |
| J | - | - | - | - | - | - | - | - | - | - | 3.737 | 4.06 | 4.045 | 4.443 |
| K | - | - | - | - | - | - | - | - | - | - | - | 0.568 | 1.037 | 2.752 |
| L | - | - | - | - | - | - | - | - | - | - | - | - | -0.4 | 2.403 |
| M | - | - | - | - | - | - | - | - | - | - | - | - | - | 2.305 |
| N | - | - | - | - | - | - | - | - | - | - | - | - | - | - |

| Max | A | B | C | D | E | F | G | H | I | J | K | L | M | N |
|---|---|---|---|---|---|---|---|---|---|---|---|---|---|---|
| A | - | 1.726 | 3.535 | 2.562 | 4.275 | 2.724 | 2.992 | 2.147 | 3.312 | 6.432 | -1.112 | 2.37 | -1.436 | 3.515 |
| B | - | - | 3.951 | 3.242 | 4.005 | 1.745 | 3.539 | 0.184 | 2.802 | 6.521 | 1.561 | 0.924 | 1.595 | 3.102 |
| C | - | - | - | 2.397 | 4.772 | 4.028 | 1.744 | 3.846 | 4.283 | 6.041 | 3.446 | 3.923 | 3.432 | 4.379 |
| D | - | - | - | - | 4.529 | 3.553 | 0.996 | 3.283 | 3.917 | 6.215 | 2.562 | 3.352 | 2.579 | 4.041 |
| E | - | - | - | - | - | 3.948 | 5.04 | 4.14 | 3.448 | 6.987 | 4.48 | 4.042 | 4.507 | 3.188 |
| F | - | - | - | - | - | - | 3.953 | 1.196 | 1.944 | 6.592 | 2.664 | 0.209 | 2.72 | 2.424 |
| G | - | - | - | - | - | - | - | 3.514 | 4.061 | 6.155 | 2.957 | 3.597 | 2.947 | 4.179 |
| H | - | - | - | - | - | - | - | - | 2.558 | 6.549 | 2.058 | -0.49 | 2.074 | 2.905 |
| I | - | - | - | - | - | - | - | - | - | 6.782 | 3.418 | 2.461 | 3.416 | 0.66 |
| J | - | - | - | - | - | - | - | - | - | - | 5.534 | 5.61 | 5.514 | 5.691 |
| K | - | - | - | - | - | - | - | - | - | - | - | 2.209 | -4.146 | 3.485 |
| L | - | - | - | - | - | - | - | - | - | - | - | - | 2.31 | 2.796 |
| M | - | - | - | - | - | - | - | - | - | - | - | - | - | 3.48 |
| N | - | - | - | - | - | - | - | - | - | - | - | - | - | - |

| Colour | Legend |
|---|---|
| (red) | Larger than 10 |
| (orange) | Between 8 and 9,99 |
| (light orange) | Between 6 and 7,99 |
| (light green) | Between 4 and 5,99 |
| (green) | Between 2 and 3,99 |
| (dark green) | Between 0 and 1,99 |
| (grey) | NA (Calculation failed) |
