## Supplementary Data S2 for "Interlaboratory comparison of standardised metabolomics and lipidomics analyses in human and rodent blood using the MxP® Quant 500 kit": TG(16.0_36.5).pdf

**Amount of relative differences between laboratories**

| Min | A | B | C | D | E | F | G | H | I | J | K | L | M | N |
|---|---|---|---|---|---|---|---|---|---|---|---|---|---|---|
| A | - | 4.309 | 4.601 | 4.524 | 5.758 | 5.258 | 4.515 | 4.758 | 4.098 | 6.647 | 4.919 | 4.671 | 4.626 | 4.901 |
| B | - | - | 1.87 | 1.745 | 5.351 | 4.459 | 1.37 | 3.002 | 2.677 | 6.276 | 3.773 | 2.94 | 2.419 | 3.042 |
| C | - | - | - | -0.437 | 5.103 | 4.266 | -0.051 | 2.311 | 3.448 | 5.954 | 3.481 | 1.932 | 1.304 | 1.856 |
| D | - | - | - | - | 5.173 | 4.302 | -1.894 | 2.456 | 3.377 | 5.998 | 3.548 | 2.429 | 1.377 | 1.958 |
| E | - | - | - | - | - | 4.227 | 5.651 | 5.368 | 6.038 | 4.993 | 4.977 | 5.431 | 5.517 | 5.477 |
| F | - | - | - | - | - | - | 4.642 | 4.149 | 5.228 | 5.702 | 3.282 | 4.291 | 4.389 | 4.25 |
| G | - | - | - | - | - | - | - | 2.551 | 3.312 | 5.939 | 3.581 | 2.43 | 1.713 | 2.043 |
| H | - | - | - | - | - | - | - | - | 4.05 | 5.937 | 2.701 | -0.645 | 1.417 | 0.633 |
| I | - | - | - | - | - | - | - | - | - | 6.616 | 4.124 | 3.623 | 3.3 | 3.912 |
| J | - | - | - | - | - | - | - | - | - | - | 2.803 | 4.263 | 4.31 | 4.682 |
| K | - | - | - | - | - | - | - | - | - | - | - | 2.927 | 3.35 | 3.239 |
| L | - | - | - | - | - | - | - | - | - | - | - | - | 1.273 | -0.689 |
| M | - | - | - | - | - | - | - | - | - | - | - | - | - | 1.48 |
| N | - | - | - | - | - | - | - | - | - | - | - | - | - | - |

| Median | A | B | C | D | E | F | G | H | I | J | K | L | M | N |
|---|---|---|---|---|---|---|---|---|---|---|---|---|---|---|
| A | - | 3.03 | 1.766 | 1.677 | 4.945 | 4.442 | 1.116 | 3.263 | 3.601 | 4.572 | 3.773 | 3.675 | 3.453 | 3.93 |
| B | - | - | 2.345 | 2.404 | 4.641 | 3.883 | 2.662 | 0.008 | 2.04 | 5.036 | 2.445 | 2.325 | 1.4 | 2.906 |
| C | - | - | - | -1.242 | 4.807 | 4.232 | 0.234 | 2.435 | 3.099 | 4.679 | 3.25 | 3.2 | 2.825 | 3.563 |
| D | - | - | - | - | 4.834 | 4.257 | -0.389 | 2.53 | 3.155 | 4.644 | 3.305 | 3.289 | 2.878 | 3.598 |
| E | - | - | - | - | - | 3.623 | 5.328 | 4.982 | 4.749 | 6.18 | 4.701 | 4.712 | 4.876 | 4.479 |
| F | - | - | - | - | - | - | 4.619 | 4.047 | 3.657 | 5.771 | 3.511 | 3.579 | 3.857 | 3.092 |
| G | - | - | - | - | - | - | - | 2.699 | 3.266 | 4.597 | 3.398 | 3.376 | 3.023 | 3.675 |
| H | - | - | - | - | - | - | - | - | 1.742 | 5.038 | 2.122 | 2.042 | 0.814 | 2.749 |
| I | - | - | - | - | - | - | - | - | - | 5.302 | 0.807 | 0.043 | 0.497 | 1.843 |
| J | - | - | - | - | - | - | - | - | - | - | 4.447 | 4.486 | 4.361 | 4.663 |
| K | - | - | - | - | - | - | - | - | - | - | - | -3.129 | 1.431 | 1.363 |
| L | - | - | - | - | - | - | - | - | - | - | - | - | 1.324 | 1.343 |
| M | - | - | - | - | - | - | - | - | - | - | - | - | - | 2.352 |
| N | - | - | - | - | - | - | - | - | - | - | - | - | - | - |

| Max | A | B | C | D | E | F | G | H | I | J | K | L | M | N |
|---|---|---|---|---|---|---|---|---|---|---|---|---|---|---|
| A | - | -2.111 | 3.428 | 3.361 | 4.133 | 3.268 | 3.574 | 2.093 | 3.263 | 6.52 | 0.583 | 2.071 | -0.168 | 2.902 |
| B | - | - | 3.461 | 3.482 | 3.995 | 3.241 | 3.632 | 2.19 | 3.238 | 6.544 | 1.048 | 1.686 | -0.606 | 2.829 |
| C | - | - | - | -2.265 | 4.608 | 4.207 | 0.4 | 2.527 | 4.205 | 6.203 | 3.002 | 3.68 | 3.387 | 4.015 |
| D | - | - | - | - | 4.599 | 4.223 | 0.138 | 2.587 | 4.224 | 6.205 | 3.041 | 3.67 | 3.437 | 4.042 |
| E | - | - | - | - | - | 2.933 | 5.058 | 4.579 | 2.85 | 7.02 | 4.392 | 3.888 | 4.116 | 3.394 |
| F | - | - | - | - | - | - | 4.603 | 3.961 | -5.943 | 6.821 | 3.693 | 2.767 | 3.287 | 1.561 |
| G | - | - | - | - | - | - | - | 2.807 | 4.287 | 6.152 | 3.207 | 3.781 | 3.545 | 4.118 |
| H | - | - | - | - | - | - | - | - | 3.748 | 6.4 | 1.254 | 2.873 | 2.322 | 3.474 |
| I | - | - | - | - | - | - | - | - | - | 6.844 | 3.654 | 2.718 | 3.271 | 1.319 |
| J | - | - | - | - | - | - | - | - | - | - | 5.589 | 5.657 | 5.647 | 5.711 |
| K | - | - | - | - | - | - | - | - | - | - | - | 2.303 | 1.481 | 3.162 |
| L | - | - | - | - | - | - | - | - | - | - | - | - | 1.361 | 2.075 |
| M | - | - | - | - | - | - | - | - | - | - | - | - | - | 2.663 |
| N | - | - | - | - | - | - | - | - | - | - | - | - | - | - |

| | |
|---|---|
| | Larger than 10 |
| | Between 8 and 9,99 |
| | Between 6 and 7,99 |
| | Between 4 and 5,99 |
| | Between 2 and 3,99 |
| | Between 0 and 1,99 |
| | NA (Calculation failed) |
