## Supplementary Data S2 for "Interlaboratory comparison of standardised metabolomics and lipidomics analyses in human and rodent blood using the MxP® Quant 500 kit": TG(16.0_38.2).pdf

**Amount of relative differences between laboratories**

| Min | A | B | C | D | E | F | G | H | I | J | K | L | M | N |
|---|---|---|---|---|---|---|---|---|---|---|---|---|---|---|
| A | - | 4.909 | 4.938 | 5.812 | 6.32 | 5.669 | 5.316 | 4.28 | 1.483 | 7.762 | 1.075 | 5.335 | 5.262 | 4.302 |
| B | - | - | 2.095 | 4.892 | 5.884 | 4.716 | 3.121 | 4.124 | 5.614 | 7.732 | 5.717 | 3.341 | 3.051 | 4.777 |
| C | - | - | - | 4.86 | 5.727 | 4.87 | 3.559 | 3.349 | 5.477 | 7.299 | 5.525 | 3.782 | 3.671 | 4.724 |
| D | - | - | - | - | 4.759 | 2.131 | 4.311 | 5.519 | 6.607 | 6.903 | 6.597 | 4.077 | 4.102 | 6.25 |
| E | - | - | - | - | - | 4.344 | 5.683 | 6.338 | 7.234 | 6.192 | 7.202 | 5.558 | 5.545 | 6.982 |
| F | - | - | - | - | - | - | 4.658 | 5.73 | 6.73 | 7.051 | 6.726 | 4.461 | 4.498 | 6.382 |
| G | - | - | - | - | - | - | - | 4.73 | 5.99 | 7.535 | 6.052 | 0.941 | -1.275 | 5.378 |
| H | - | - | - | - | - | - | - | - | 4.772 | 7.565 | 4.92 | 4.542 | 4.446 | 3.192 |
| I | - | - | - | - | - | - | - | - | - | 7.958 | 2.525 | 5.257 | 5.161 | 4.246 |
| J | - | - | - | - | - | - | - | - | - | - | 6.899 | 4.948 | 4.861 | 6.76 |
| K | - | - | - | - | - | - | - | - | - | - | - | 5.361 | 5.319 | 4.03 |
| L | - | - | - | - | - | - | - | - | - | - | - | - | 0.843 | 5.433 |
| M | - | - | - | - | - | - | - | - | - | - | - | - | - | 5.31 |
| N | - | - | - | - | - | - | - | - | - | - | - | - | - | - |

| | |
|---|---|
| | Larger than 10 |
| | Between 8 and 9,99 |
| | Between 6 and 7,99 |
| | Between 4 and 5,99 |
| | Between 2 and 3,99 |
| | Between 0 and 1,99 |
| | NA (Calculation failed) |

| Median | A | B | C | D | E | F | G | H | I | J | K | L | M | N |
|---|---|---|---|---|---|---|---|---|---|---|---|---|---|---|
| A | - | 4.389 | 2.815 | 4.455 | 4.81 | 4.651 | 4.5 | 2.607 | 3.221 | 1.873 | 4.439 | 4.731 | 4.707 | 1.316 |
| B | - | - | 4.183 | -0.45 | 3.255 | 2.464 | 0.936 | 4.217 | 3.94 | 4.408 | 5.75 | 2.801 | 2.777 | 4.528 |
| C | - | - | - | 3.975 | 4.58 | 4.358 | 4.1 | -2.189 | 1.221 | 1.86 | 4.887 | 4.428 | 4.442 | 1.822 |
| D | - | - | - | - | 3.425 | 2.723 | 1.147 | 4.108 | 3.971 | 4.534 | 5.714 | 2.894 | 3.031 | 4.435 |
| E | - | - | - | - | - | 2.267 | 3.351 | 4.983 | 4.945 | 5.305 | 6.159 | 2.074 | 1.546 | 5.19 |
| F | - | - | - | - | - | - | 2.208 | 4.674 | 4.565 | 4.963 | 5.983 | -0.008 | 0.742 | 4.919 |
| G | - | - | - | - | - | - | - | 4.368 | 4.149 | 4.664 | 5.823 | 2.384 | 2.407 | 4.655 |
| H | - | - | - | - | - | - | - | - | 1.424 | 1.465 | 4.894 | 4.429 | 4.424 | 1.917 |
| I | - | - | - | - | - | - | - | - | - | 2.34 | 5.105 | 4.239 | 4.192 | 2.957 |
| J | - | - | - | - | - | - | - | - | - | - | 4.403 | 4.706 | 4.8 | 1.994 |
| K | - | - | - | - | - | - | - | - | - | - | - | 5.304 | 5.302 | 4.322 |
| L | - | - | - | - | - | - | - | - | - | - | - | - | -5.612 | 5.01 |
| M | - | - | - | - | - | - | - | - | - | - | - | - | - | 5.006 |
| N | - | - | - | - | - | - | - | - | - | - | - | - | - | - |

| Max | A | B | C | D | E | F | G | H | I | J | K | L | M | N |
|---|---|---|---|---|---|---|---|---|---|---|---|---|---|---|
| A | - | 3.679 | 3.756 | 3.206 | 3.443 | -2.312 | 3.366 | 2.063 | 3.603 | 7.094 | 5.116 | 4.089 | 4.167 | 2.454 |
| B | - | - | 4.918 | 4.679 | 4.789 | 3.864 | 1.556 | 4.285 | -0.359 | 7.451 | 5.771 | 2.259 | 2.56 | 4.397 |
| C | - | - | - | 1.855 | 1.081 | 3.591 | 4.391 | 3.042 | 4.499 | 6.776 | 4.225 | 4.749 | 4.794 | 2.816 |
| D | - | - | - | - | 0.614 | 3.13 | 4.17 | 2.263 | 4.303 | 6.871 | 4.532 | 4.597 | 4.648 | 1.826 |
| E | - | - | - | - | - | 3.348 | 4.268 | 2.661 | 4.395 | 6.828 | 4.404 | 4.667 | 4.715 | 2.336 |
| F | - | - | - | - | - | - | 3.342 | 2.114 | 3.588 | 7.118 | 5.128 | 4.077 | 4.158 | 2.508 |
| G | - | - | - | - | - | - | - | 4.011 | 1.061 | 7.357 | 5.649 | 2.906 | 3.099 | 4.14 |
| H | - | - | - | - | - | - | - | - | 3.97 | 7.012 | 4.876 | 4.345 | 4.41 | 0.376 |
| I | - | - | - | - | - | - | - | - | - | 7.425 | 5.739 | 2.468 | 2.727 | 4.331 |
| J | - | - | - | - | - | - | - | - | - | - | 5.456 | 6.031 | 6.04 | 5.805 |
| K | - | - | - | - | - | - | - | - | - | - | - | 5.264 | 5.29 | 4.439 |
| L | - | - | - | - | - | - | - | - | - | - | - | - | 0.216 | 4.762 |
| M | - | - | - | - | - | - | - | - | - | - | - | - | - | 4.841 |
| N | - | - | - | - | - | - | - | - | - | - | - | - | - | - |
