## Supplementary Data S2 for "Interlaboratory comparison of standardised metabolomics and lipidomics analyses in human and rodent blood using the MxP® Quant 500 kit": TG(16.0_38.4).pdf

**Amount of relative differences between laboratories**

| Min | A | B | C | D | E | F | G | H | I | J | K | L | M | N |
|---|---|---|---|---|---|---|---|---|---|---|---|---|---|---|
| A | - | 5.4 | 5.66 | 5.891 | 6.513 | 5.972 | 5.587 | 5.292 | 4.635 | 7.644 | 5.518 | 5.683 | 5.1 | 5.887 |
| B | - | - | 2.939 | 4.221 | 5.926 | 4.724 | 3.03 | 2.212 | 4.625 | 7.297 | 1.42 | 3.386 | -0.155 | 3.38 |
| C | - | - | - | 3.298 | 5.661 | 4.273 | 0.812 | 0.109 | 4.965 | 6.997 | 2.157 | 1.541 | 2.013 | -1.564 |
| D | - | - | - | - | 5.404 | 3.446 | 2.777 | 2.596 | 5.384 | 6.814 | 3.827 | 2.684 | 3.515 | 3.594 |
| E | - | - | - | - | - | 5.404 | 6.068 | 5.958 | 6.898 | 6.29 | 6.313 | 6.107 | 6.124 | 6.353 |
| F | - | - | - | - | - | - | 4.295 | 4.179 | 5.864 | 6.759 | 4.796 | 4.26 | 4.57 | 4.707 |
| G | - | - | - | - | - | - | - | 0.538 | 5.094 | 7.06 | 2.677 | 0.498 | 2.785 | 0.333 |
| H | - | - | - | - | - | - | - | - | 4.942 | 7.314 | 0.945 | 2.558 | 2.53 | 2.607 |
| I | - | - | - | - | - | - | - | - | - | 7.587 | 4.658 | 5.002 | 3.983 | 5.225 |
| J | - | - | - | - | - | - | - | - | - | - | 6.131 | 5.941 | 5.641 | 6.357 |
| K | - | - | - | - | - | - | - | - | - | - | - | 2.931 | 0.752 | 2.533 |
| L | - | - | - | - | - | - | - | - | - | - | - | - | 3.255 | 1.047 |
| M | - | - | - | - | - | - | - | - | - | - | - | - | - | 3.616 |
| N | - | - | - | - | - | - | - | - | - | - | - | - | - | - |

| Median | A | B | C | D | E | F | G | H | I | J | K | L | M | N |
|---|---|---|---|---|---|---|---|---|---|---|---|---|---|---|
| A | - | 3.325 | 1.374 | 1.474 | 4.812 | 3.27 | 2.9 | 3.807 | 3.286 | 4.899 | 2.561 | 3.582 | 3.364 | 4.036 |
| B | - | - | 3.084 | 3.049 | 4.337 | -4.74 | 1.51 | 2.232 | -1.514 | 5.298 | 2.362 | 1.124 | -1.724 | 2.782 |
| C | - | - | - | -3.919 | 4.72 | 3.003 | 2.398 | 3.626 | 3.085 | 4.73 | 1.559 | 3.301 | 3.033 | 3.821 |
| D | - | - | - | - | 4.721 | 3.029 | 2.424 | 3.642 | 3.126 | 4.656 | 1.509 | 3.307 | 3.038 | 3.818 |
| E | - | - | - | - | - | 4.599 | 4.817 | 4.202 | 4.538 | 6.056 | 5.002 | 4.476 | 4.61 | 4.052 |
| F | - | - | - | - | - | - | 1.631 | 2.229 | -0.659 | 5.168 | 2.555 | 0.891 | -4.277 | 2.683 |
| G | - | - | - | - | - | - | - | 2.892 | 1.728 | 5.097 | 1.275 | 2.275 | 1.613 | 3.227 |
| H | - | - | - | - | - | - | - | - | 2.191 | 5.532 | 3.365 | 1.419 | 2.203 | 1.194 |
| I | - | - | - | - | - | - | - | - | - | 5.396 | 2.238 | 1.173 | -1.569 | 2.826 |
| J | - | - | - | - | - | - | - | - | - | - | 3.845 | 4.42 | 4.331 | 4.633 |
| K | - | - | - | - | - | - | - | - | - | - | - | 2.819 | 2.394 | 3.52 |
| L | - | - | - | - | - | - | - | - | - | - | - | - | 0.911 | 2.25 |
| M | - | - | - | - | - | - | - | - | - | - | - | - | - | 2.728 |
| N | - | - | - | - | - | - | - | - | - | - | - | - | - | - |

| Max | A | B | C | D | E | F | G | H | I | J | K | L | M | N |
|---|---|---|---|---|---|---|---|---|---|---|---|---|---|---|
| A | - | 1.617 | 4.283 | 4.625 | -1.352 | 4.041 | 3.492 | 0.839 | 2.362 | 7.039 | 3.575 | 2.055 | 1.112 | 1.567 |
| B | - | - | 3.991 | 4.416 | 0.946 | 3.705 | 2.986 | 2.242 | 3 | 6.98 | 3.109 | 0.273 | -0.111 | 2.528 |
| C | - | - | - | 2.226 | 3.95 | 1.294 | 2.785 | 4.154 | 4.367 | 6.566 | 2.644 | 3.666 | 3.861 | 4.223 |
| D | - | - | - | - | 4.261 | 2.769 | 3.459 | 4.423 | 4.595 | 6.428 | 3.385 | 4.051 | 4.202 | 4.481 |
| E | - | - | - | - | - | 3.938 | 3.358 | 1.479 | 2.613 | 7.029 | 3.441 | 1.652 | -0.158 | 1.954 |
| F | - | - | - | - | - | - | 2.177 | 3.976 | 4.22 | 6.64 | 1.962 | 3.387 | 3.632 | 4.056 |
| G | - | - | - | - | - | - | - | 3.552 | 3.884 | 6.76 | -0.629 | 2.642 | 3.039 | 3.666 |
| H | - | - | - | - | - | - | - | - | 1.782 | 7.099 | 3.812 | 2.628 | 2 | 0.128 |
| I | - | - | - | - | - | - | - | - | - | 7.187 | 4.177 | 3.314 | 2.943 | 1.278 |
| J | - | - | - | - | - | - | - | - | - | - | 5.701 | 5.79 | 5.82 | 5.866 |
| K | - | - | - | - | - | - | - | - | - | - | - | 2.77 | 3.147 | 3.729 |
| L | - | - | - | - | - | - | - | - | - | - | - | - | 1.07 | 2.787 |
| M | - | - | - | - | - | - | - | - | - | - | - | - | - | 2.281 |
| N | - | - | - | - | - | - | - | - | - | - | - | - | - | - |

| Colour | Legend |
|---|---|
| Red | Larger than 10 |
| Orange | Between 8 and 9,99 |
| Light orange | Between 6 and 7,99 |
| Light green | Between 4 and 5,99 |
| Green | Between 2 and 3,99 |
| Dark green | Between 0 and 1,99 |
| Grey | NA (Calculation failed) |
