## Supplementary Data S2 for "Interlaboratory comparison of standardised metabolomics and lipidomics analyses in human and rodent blood using the MxP® Quant 500 kit": TG(16.0_38.5).pdf

**Amount of relative differences between laboratories**

| Min | A | B | C | D | E | F | G | H | I | J | K | L | M | N |
|---|---|---|---|---|---|---|---|---|---|---|---|---|---|---|
| A | - | 3.714 | 4.565 | 4.483 | 5.852 | 4.944 | 4.311 | 4.214 | 1.502 | 6.592 | 4.786 | 4.421 | 4.164 | 4.763 |
| B | - | - | 3.515 | 3.201 | 5.543 | 4.252 | 2.545 | 2.665 | 3.747 | 6.339 | 4.039 | 2.958 | 2.519 | 3.375 |
| C | - | - | - | 1.834 | 5.129 | 3.039 | 2.827 | 2.351 | 4.837 | 6.042 | 2.54 | 2.37 | 2.501 | 2.202 |
| D | - | - | - | - | 5.171 | 3.49 | 1.852 | -0.047 | 4.586 | 5.962 | 3.221 | 0.694 | 0.678 | 1.388 |
| E | - | - | - | - | - | 4.928 | 5.706 | 5.531 | 6.348 | 4.744 | 4.989 | 5.643 | 5.562 | 5.782 |
| F | - | - | - | - | - | - | 4.084 | 3.781 | 5.321 | 5.783 | 0.977 | 3.877 | 3.838 | 4.021 |
| G | - | - | - | - | - | - | - | 0.734 | 4.331 | 6.124 | 3.579 | 0.974 | 0.117 | 1.42 |
| H | - | - | - | - | - | - | - | - | 4.418 | 6.26 | 3.431 | 0.19 | -0.666 | 1.855 |
| I | - | - | - | - | - | - | - | - | - | 6.717 | 4.662 | 4.276 | 3.878 | 4.773 |
| J | - | - | - | - | - | - | - | - | - | - | 3.749 | 5.013 | 4.64 | 5.494 |
| K | - | - | - | - | - | - | - | - | - | - | - | 3.512 | 3.629 | 3.503 |
| L | - | - | - | - | - | - | - | - | - | - | - | - | 0.351 | 0.338 |
| M | - | - | - | - | - | - | - | - | - | - | - | - | - | 1.914 |
| N | - | - | - | - | - | - | - | - | - | - | - | - | - | - |

| | |
|---|---|
| | Larger than 10 |
| | Between 8 and 9,99 |
| | Between 6 and 7,99 |
| | Between 4 and 5,99 |
| | Between 2 and 3,99 |
| | Between 0 and 1,99 |
| | NA (Calculation failed) |

| Median | A | B | C | D | E | F | G | H | I | J | K | L | M | N |
|---|---|---|---|---|---|---|---|---|---|---|---|---|---|---|
| A | - | 2.196 | 3.325 | 2.041 | 4.838 | 3.574 | 2.367 | 3.228 | 2.973 | 3.835 | 3.975 | 3.166 | 3.019 | 3.988 |
| B | - | - | 2.449 | 0.165 | 4.64 | 2.997 | -4.405 | 2.384 | 1.826 | 4.184 | 3.457 | 1.96 | 1.941 | 3.59 |
| C | - | - | - | 2.892 | 4.328 | 1.629 | 2.571 | -3.206 | 0.809 | 4.584 | 2.441 | 1.079 | 0.463 | 2.87 |
| D | - | - | - | - | 4.685 | 3.338 | 0.479 | 2.874 | 2.483 | 3.852 | 3.569 | 2.296 | 2.558 | 3.808 |
| E | - | - | - | - | - | 4.296 | 5.015 | 4.548 | 4.69 | 5.668 | 4.22 | 4.813 | 4.67 | 3.85 |
| F | - | - | - | - | - | - | 3.304 | 1.658 | 2.356 | 4.77 | 0.873 | 2.582 | 2.242 | 2.106 |
| G | - | - | - | - | - | - | - | 2.492 | 1.957 | 4.108 | 3.429 | 1.885 | 2.076 | 3.635 |
| H | - | - | - | - | - | - | - | - | 0.842 | 4.627 | 2.578 | 0.66 | 0.549 | 2.85 |
| I | - | - | - | - | - | - | - | - | - | 4.606 | 3.176 | 0.691 | -2.103 | 3.119 |
| J | - | - | - | - | - | - | - | - | - | - | 4.016 | 3.274 | 3.832 | 4.457 |
| K | - | - | - | - | - | - | - | - | - | - | - | 2.976 | 2.849 | 1.031 |
| L | - | - | - | - | - | - | - | - | - | - | - | - | -1.461 | 3.149 |
| M | - | - | - | - | - | - | - | - | - | - | - | - | - | 3.099 |
| N | - | - | - | - | - | - | - | - | - | - | - | - | - | - |

| Max | A | B | C | D | E | F | G | H | I | J | K | L | M | N |
|---|---|---|---|---|---|---|---|---|---|---|---|---|---|---|
| A | - | 1.864 | 2.201 | 4.083 | 1.532 | 2.737 | 3.262 | -0.591 | 3.454 | 6.843 | -0.682 | 1.837 | 0.694 | 3.292 |
| B | - | - | -0.082 | 3.703 | 2.538 | 1.544 | 2.527 | 2.056 | 3.815 | 6.757 | 2.011 | -5.321 | 0.976 | 3.694 |
| C | - | - | - | 3.57 | 2.798 | 0.966 | 2.259 | 2.335 | 3.896 | 6.721 | 2.304 | -0.044 | 1.517 | 3.778 |
| D | - | - | - | - | 4.058 | 3.17 | 2.682 | 3.926 | 4.582 | 6.394 | 3.917 | 3.524 | 3.734 | 4.521 |
| E | - | - | - | - | - | 3.172 | 3.629 | 0.802 | 3.123 | 6.922 | 0.946 | 2.6 | 1.963 | 2.894 |
| F | - | - | - | - | - | - | 1.474 | 2.783 | 4.045 | 6.672 | 2.738 | 1.482 | 2.247 | 3.941 |
| G | - | - | - | - | - | - | - | 3.23 | 4.231 | 6.588 | 3.213 | 2.442 | 2.871 | 4.143 |
| H | - | - | - | - | - | - | - | - | 3.373 | 6.866 | -2.575 | 2.112 | 1.194 | 3.193 |
| I | - | - | - | - | - | - | - | - | - | 7.148 | 3.55 | 4.026 | 3.817 | 0.434 |
| J | - | - | - | - | - | - | - | - | - | - | 5.753 | 5.701 | 5.729 | 5.859 |
| K | - | - | - | - | - | - | - | - | - | - | - | 2.103 | 1.064 | 3.208 |
| L | - | - | - | - | - | - | - | - | - | - | - | - | 0.99 | 3.697 |
| M | - | - | - | - | - | - | - | - | - | - | - | - | - | 3.479 |
| N | - | - | - | - | - | - | - | - | - | - | - | - | - | - |
