## Supplementary Data S2 for "Interlaboratory comparison of standardised metabolomics and lipidomics analyses in human and rodent blood using the MxP® Quant 500 kit": TG(16.0_40.6).pdf

**Amount of relative differences between laboratories**

| Min | A | B | C | D | E | F | G | H | I | J | K | L | M | N |
|---|---|---|---|---|---|---|---|---|---|---|---|---|---|---|
| A | - | 4.689 | 4.061 | 4.02 | 5.353 | 5.166 | 4.2 | 4.602 | 4.295 | 5.182 | 4.228 | 4.44 | 4.322 | 5.021 |
| B | - | - | 3.674 | 3.862 | 4.135 | 3.129 | 3.516 | 1.717 | 3.996 | 2.789 | 3.202 | 2.94 | 3.233 | -0.087 |
| C | - | - | - | 0.985 | 4.699 | 4.128 | -0.215 | 3.033 | 2.455 | 3.78 | 1.719 | 1.993 | 1.391 | 3.231 |
| D | - | - | - | - | 4.771 | 4.237 | 1.49 | 3.317 | 1.909 | 3.949 | 2.419 | 2.531 | 2.176 | 3.423 |
| E | - | - | - | - | - | 3.628 | 5.014 | 4.497 | 5.478 | 4.285 | 4.812 | 4.846 | 4.913 | 4.647 |
| F | - | - | - | - | - | - | 4.139 | 3.168 | 4.841 | 3.078 | 3.819 | 3.865 | 3.968 | 3.485 |
| G | - | - | - | - | - | - | - | 2.886 | 2.493 | 3.717 | 1.163 | 1.685 | 0.854 | 3.24 |
| H | - | - | - | - | - | - | - | - | 3.661 | 3.126 | 2.501 | 2.098 | 2.576 | 1.938 |
| I | - | - | - | - | - | - | - | - | - | 4.444 | -1.459 | 1.785 | 0.486 | 3.971 |
| J | - | - | - | - | - | - | - | - | - | - | 2.271 | 1.081 | 0.988 | 0.568 |
| K | - | - | - | - | - | - | - | - | - | - | - | -0.195 | -0.742 | 2.848 |
| L | - | - | - | - | - | - | - | - | - | - | - | - | 0.579 | 2.762 |
| M | - | - | - | - | - | - | - | - | - | - | - | - | - | 3.253 |
| N | - | - | - | - | - | - | - | - | - | - | - | - | - | - |

| Median | A | B | C | D | E | F | G | H | I | J | K | L | M | N |
|---|---|---|---|---|---|---|---|---|---|---|---|---|---|---|
| A | - | 4.424 | 3.392 | 2.961 | 4.888 | 4.324 | 3.553 | 4.254 | 4.138 | 2.431 | 3.751 | 3.921 | 3.796 | 4.328 |
| B | - | - | 3.915 | 4.18 | 3.369 | 0.383 | 3.721 | 1.811 | 2.292 | 4.481 | 3.429 | 3.182 | 3.206 | -0.861 |
| C | - | - | - | 1.318 | 4.48 | 3.643 | 0.811 | 3.36 | 3.194 | 2.86 | 1.942 | 2.391 | 2.491 | 3.741 |
| D | - | - | - | - | 4.6 | 3.878 | 2.051 | 3.638 | 3.514 | 2.176 | 2.637 | 2.909 | 2.999 | 3.964 |
| E | - | - | - | - | - | 3.608 | 4.746 | 3.976 | 4.075 | 5.306 | 4.588 | 4.497 | 4.4 | 3.411 |
| F | - | - | - | - | - | - | 3.693 | 1.611 | 2.052 | 4.668 | 3.401 | 3.201 | 2.973 | 0.48 |
| G | - | - | - | - | - | - | - | 3.111 | 2.893 | 3.223 | 1.07 | 1.838 | 1.917 | 3.531 |
| H | - | - | - | - | - | - | - | - | 0.463 | 4.226 | 2.809 | 2.432 | 2.462 | 1.538 |
| I | - | - | - | - | - | - | - | - | - | 3.862 | 2.358 | 1.686 | 2.171 | 1.684 |
| J | - | - | - | - | - | - | - | - | - | - | 3.793 | 3.864 | 4.121 | 4.666 |
| K | - | - | - | - | - | - | - | - | - | - | - | 0.549 | 0.763 | 3.281 |
| L | - | - | - | - | - | - | - | - | - | - | - | - | -2.958 | 3.031 |
| M | - | - | - | - | - | - | - | - | - | - | - | - | - | 2.985 |
| N | - | - | - | - | - | - | - | - | - | - | - | - | - | - |

| Max | A | B | C | D | E | F | G | H | I | J | K | L | M | N |
|---|---|---|---|---|---|---|---|---|---|---|---|---|---|---|
| A | - | 2.92 | 3.921 | 4.315 | -1.105 | 3.621 | 3.295 | 1.895 | 3.93 | 6.853 | 3.01 | 2.217 | -0.856 | 2.507 |
| B | - | - | 4.624 | 4.9 | 3.173 | 4.444 | 4.238 | 2.058 | 3.017 | 7.096 | 4.102 | 3.734 | 3.14 | 1.192 |
| C | - | - | - | 2.064 | 3.652 | 1.211 | 2.219 | 4.037 | 4.719 | 6.483 | 2.606 | 3.181 | 3.668 | 4.169 |
| D | - | - | - | - | 4.005 | 2.638 | 3.084 | 4.308 | 4.871 | 6.37 | 3.29 | 3.672 | 4.019 | 4.411 |
| E | - | - | - | - | - | 3.568 | 3.18 | 2.177 | 3.973 | 6.884 | 2.909 | 1.965 | -2.209 | 2.616 |
| F | - | - | - | - | - | - | 1.196 | 3.856 | 4.623 | 6.522 | 1.974 | 2.817 | 3.406 | 4.003 |
| G | - | - | - | - | - | - | - | 3.623 | 4.509 | 6.605 | 0.613 | 2.221 | 3.077 | 3.809 |
| H | - | - | - | - | - | - | - | - | 3.568 | 6.979 | 3.645 | 3.133 | 2.15 | 0.867 |
| I | - | - | - | - | - | - | - | - | - | 7.292 | 4.782 | 4.546 | 4.207 | 3.496 |
| J | - | - | - | - | - | - | - | - | - | - | 5.664 | 5.705 | 5.751 | 5.823 |
| K | - | - | - | - | - | - | - | - | - | - | - | 1.698 | 2.833 | 3.662 |
| L | - | - | - | - | - | - | - | - | - | - | - | - | 1.987 | 3.292 |
| M | - | - | - | - | - | - | - | - | - | - | - | - | - | 2.594 |
| N | - | - | - | - | - | - | - | - | - | - | - | - | - | - |

| Colour | Meaning |
|---|---|
| (red) | Larger than 10 |
| (orange) | Between 8 and 9,99 |
| (light orange) | Between 6 and 7,99 |
| (light green) | Between 4 and 5,99 |
| (green) | Between 2 and 3,99 |
| (dark green) | Between 0 and 1,99 |
| (light grey) | NA (Calculation failed) |
