## Supplementary Data S2 for "Interlaboratory comparison of standardised metabolomics and lipidomics analyses in human and rodent blood using the MxP® Quant 500 kit": TG(16.0_40.7).pdf

**Amount of relative differences between laboratories**

| Min | A | B | C | D | E | F | G | H | I | J | K | L | M | N |
|---|---|---|---|---|---|---|---|---|---|---|---|---|---|---|
| A | - | 2.926 | 4.267 | 4.445 | 5.752 | 5.052 | 4.688 | 4.235 | 3.97 | 7.023 | 3.308 | 4.717 | 4.483 | 4.974 |
| B | - | - | 3.394 | 3.754 | 5.554 | 4.753 | 4.077 | 3.346 | 3.042 | 6.946 | -1.946 | 4.211 | 3.92 | 4.676 |
| C | - | - | - | 1.85 | 5.256 | 4.271 | 2.618 | -3.071 | -0.54 | 6.62 | 3.521 | 3.227 | 2.752 | 4.208 |
| D | - | - | - | - | 5.155 | 4.054 | 1.174 | 1.946 | 2.034 | 6.578 | 3.975 | 2.591 | 1.788 | 3.955 |
| E | - | - | - | - | - | 4.505 | 5.547 | 5.769 | 5.718 | 5.797 | 6.172 | 5.317 | 5.392 | 4.54 |
| F | - | - | - | - | - | - | 4.107 | 4.62 | 4.627 | 6.61 | 5.267 | 3.697 | 3.952 | -0.828 |
| G | - | - | - | - | - | - | - | 2.632 | 2.689 | 6.396 | 4.21 | 1.956 | 0.376 | 3.759 |
| H | - | - | - | - | - | - | - | - | 0.124 | 6.652 | 3.469 | 3.237 | 2.675 | 4.161 |
| I | - | - | - | - | - | - | - | - | - | 7.096 | 2.845 | 3.616 | 2.934 | 4.217 |
| J | - | - | - | - | - | - | - | - | - | - | 5.569 | 4.267 | 4.161 | -0.809 |
| K | - | - | - | - | - | - | - | - | - | - | - | 4.233 | 3.995 | 4.728 |
| L | - | - | - | - | - | - | - | - | - | - | - | - | 1.668 | 3.234 |
| M | - | - | - | - | - | - | - | - | - | - | - | - | - | 3.658 |
| N | - | - | - | - | - | - | - | - | - | - | - | - | - | - |

| | |
|---|---|
| | Larger than 10 |
| | Between 8 and 9,99 |
| | Between 6 and 7,99 |
| | Between 4 and 5,99 |
| | Between 2 and 3,99 |
| | Between 0 and 1,99 |
| | NA (Calculation failed) |

| Median | A | B | C | D | E | F | G | H | I | J | K | L | M | N |
|---|---|---|---|---|---|---|---|---|---|---|---|---|---|---|
| A | - | 0.547 | 1.651 | 2.556 | 4.889 | 4.196 | 2.763 | 2.44 | 3.864 | 4.681 | 2.914 | 3.776 | 3.765 | 4.513 |
| B | - | - | 0.661 | 1.948 | 4.834 | 4.11 | 2.356 | 2.088 | 3.748 | 4.69 | 3.137 | 3.62 | 3.626 | 4.44 |
| C | - | - | - | 0.775 | 4.772 | 4.02 | 1.761 | 1.656 | 3.625 | 4.641 | 3.318 | 3.445 | 3.477 | 4.363 |
| D | - | - | - | - | 4.712 | 3.905 | 0.837 | 0.781 | 3.462 | 4.72 | 3.535 | 3.245 | 3.286 | 4.271 |
| E | - | - | - | - | - | 3.864 | 5.053 | 5.015 | 4.319 | 6.056 | 5.646 | 4.495 | 4.45 | 3.164 |
| F | - | - | - | - | - | - | 3.963 | 3.96 | 2.225 | 5.594 | 4.961 | 2.718 | 2.622 | 2.357 |
| G | - | - | - | - | - | - | - | -4.756 | 3.235 | 4.813 | 3.765 | 2.97 | 3.023 | 4.156 |
| H | - | - | - | - | - | - | - | - | 3.252 | 4.887 | 3.762 | 3.025 | 3.055 | 4.173 |
| I | - | - | - | - | - | - | - | - | - | 5.551 | 4.691 | 0.107 | 0.243 | 3.268 |
| J | - | - | - | - | - | - | - | - | - | - | 1.789 | 4.286 | 4.366 | 4.858 |
| K | - | - | - | - | - | - | - | - | - | - | - | 4.252 | 4.263 | 4.796 |
| L | - | - | - | - | - | - | - | - | - | - | - | - | -2.181 | 3.429 |
| M | - | - | - | - | - | - | - | - | - | - | - | - | - | 3.418 |
| N | - | - | - | - | - | - | - | - | - | - | - | - | - | - |

| Max | A | B | C | D | E | F | G | H | I | J | K | L | M | N |
|---|---|---|---|---|---|---|---|---|---|---|---|---|---|---|
| A | - | 2.04 | 3.304 | 3.313 | 3.584 | 2.306 | 3.086 | 1.97 | 3.785 | 6.376 | 4.288 | 1.796 | 2.66 | 3.917 |
| B | - | - | 2.494 | 2.517 | 3.914 | 3.094 | 2.118 | -2.947 | 4.094 | 6.335 | 3.909 | 2.856 | 3.325 | 4.189 |
| C | - | - | - | -2.542 | 4.278 | 3.75 | 0.237 | 2.441 | 4.424 | 6.137 | 3.16 | 3.601 | 3.886 | 4.489 |
| D | - | - | - | - | 4.274 | 3.757 | 0.494 | 2.485 | 4.433 | 6.155 | 3.128 | 3.619 | 3.906 | 4.505 |
| E | - | - | - | - | - | 2.854 | 4.478 | 4.075 | 1.543 | 6.801 | 5.127 | 3.085 | 2.36 | 1.96 |
| F | - | - | - | - | - | - | 3.836 | 3.211 | 3.226 | 6.674 | 4.705 | 0.478 | 0.658 | 3.355 |
| G | - | - | - | - | - | - | - | 2.09 | 4.359 | 6.119 | 3.352 | 3.466 | 3.786 | 4.438 |
| H | - | - | - | - | - | - | - | - | 4.084 | 6.277 | 3.933 | 2.826 | 3.309 | 4.184 |
| I | - | - | - | - | - | - | - | - | - | 6.84 | 5.273 | 3.573 | 3.089 | 0.658 |
| J | - | - | - | - | - | - | - | - | - | - | 5.265 | 5.671 | 5.717 | 5.796 |
| K | - | - | - | - | - | - | - | - | - | - | - | 4.267 | 4.453 | 4.855 |
| L | - | - | - | - | - | - | - | - | - | - | - | - | 1.605 | 3.585 |
| M | - | - | - | - | - | - | - | - | - | - | - | - | - | 3.162 |
| N | - | - | - | - | - | - | - | - | - | - | - | - | - | - |
