## Supplementary Data S2 for "Interlaboratory comparison of standardised metabolomics and lipidomics analyses in human and rodent blood using the MxP® Quant 500 kit": TG(16.1_32.2).pdf

**Amount of relative differences between laboratories**

| Min | A | B | C | D | E | F | G | H | I | J | K | L | M | N |
|---|---|---|---|---|---|---|---|---|---|---|---|---|---|---|
| A | - | 6.276 | 4.928 | 5.953 | 10.218 | 6.176 | 5.676 | 5.504 | 4.065 | 8.59 | 5.73 | 5.305 | 5.369 | 5.338 |
| B | - | - | 0.497 | 4.426 | 9.83 | 4.454 | 5.653 | 2.728 | 5.541 | 7.758 | 3.704 | 4.813 | 3.746 | 4.223 |
| C | - | - | - | 4.941 | 11.064 | 5.706 | 2.229 | 3.293 | 5.14 | 9.155 | 3.558 | 0.752 | 1.871 | -1.601 |
| D | - | - | - | - | 9.979 | 5.124 | 4.605 | 2.735 | 4.928 | 8.031 | 1.949 | 3.445 | -0.682 | 1.852 |
| E | - | - | - | - | - | 3.345 | 6.882 | 4.67 | 5.909 | 6.815 | 5.634 | 5.873 | 4.988 | 5.327 |
| F | - | - | - | - | - | - | 6.538 | 5.083 | 6.376 | 7.689 | 5.448 | 5.949 | 5.409 | 5.622 |
| G | - | - | - | - | - | - | - | 4.181 | 4.104 | 8.292 | 4.266 | 2.196 | 3.641 | 3.253 |
| H | - | - | - | - | - | - | - | - | 5.339 | 8.198 | 0.983 | 4.193 | 2.734 | 3.439 |
| I | - | - | - | - | - | - | - | - | - | 8.662 | 5.325 | 4.616 | 4.818 | 4.75 |
| J | - | - | - | - | - | - | - | - | - | - | 5.982 | 6.281 | 5.689 | 5.914 |
| K | - | - | - | - | - | - | - | - | - | - | - | 3.858 | 1.335 | 2.767 |
| L | - | - | - | - | - | - | - | - | - | - | - | - | 3.068 | 2.345 |
| M | - | - | - | - | - | - | - | - | - | - | - | - | - | 1.759 |
| N | - | - | - | - | - | - | - | - | - | - | - | - | - | - |

| | |
|---|---|
| | Larger than 10 |
| | Between 8 and 9,99 |
| | Between 6 and 7,99 |
| | Between 4 and 5,99 |
| | Between 2 and 3,99 |
| | Between 0 and 1,99 |
| | NA (Calculation failed) |

| Median | A | B | C | D | E | F | G | H | I | J | K | L | M | N |
|---|---|---|---|---|---|---|---|---|---|---|---|---|---|---|
| A | - | 4.232 | 4.274 | 4.237 | 8.771 | 5.287 | 3.785 | 4.591 | 3.515 | 5.659 | 4.182 | 4.295 | 4.491 | 4.465 |
| B | - | - | 4.162 | 2.657 | 8.311 | 4.173 | 3.079 | 1.841 | 3.027 | 4.322 | 2.469 | 1.758 | 2.593 | 0.168 |
| C | - | - | - | 2.021 | 9.933 | 4.396 | 3.585 | 1.029 | 4.275 | 6.501 | 3.499 | 2.876 | 2.478 | 2.27 |
| D | - | - | - | - | 8.489 | 4.544 | 1.563 | 3.119 | 1.735 | 4.805 | -1.524 | 1.214 | 3.337 | 2.66 |
| E | - | - | - | - | - | 4.978 | 4.151 | 4.681 | 4.444 | 3.891 | 4.054 | 4.225 | 5.051 | 4.615 |
| F | - | - | - | - | - | - | 4.972 | 4.085 | 4.943 | -1.705 | 4.819 | 4.679 | 3.755 | 4.319 |
| G | - | - | - | - | - | - | - | 3.524 | 0.294 | 5.092 | 1.93 | 2.514 | 3.556 | 3.2 |
| H | - | - | - | - | - | - | - | - | 3.749 | 4.498 | 3.168 | 2.778 | 0.3 | 1.365 |
| I | - | - | - | - | - | - | - | - | - | 5.486 | 2.819 | 3.101 | 3.653 | 3.544 |
| J | - | - | - | - | - | - | - | - | - | - | 3.439 | 3.117 | 2.625 | 1.375 |
| K | - | - | - | - | - | - | - | - | - | - | - | 1.039 | 3.335 | 2.606 |
| L | - | - | - | - | - | - | - | - | - | - | - | - | 2.678 | 1.874 |
| M | - | - | - | - | - | - | - | - | - | - | - | - | - | 1.579 |
| N | - | - | - | - | - | - | - | - | - | - | - | - | - | - |

| Max | A | B | C | D | E | F | G | H | I | J | K | L | M | N |
|---|---|---|---|---|---|---|---|---|---|---|---|---|---|---|
| A | - | 4.499 | 3.608 | 3.951 | 9.82 | 2.818 | 2.57 | 1.361 | 2.799 | 6.883 | 4.286 | 0.31 | 2.542 | 1.775 |
| B | - | - | 4.834 | 2.532 | 9.524 | 3.664 | 3.77 | 4.034 | 4.595 | 6.302 | 1.349 | 4.283 | 4.538 | 4.41 |
| C | - | - | - | 4.988 | 10.091 | 4.468 | 4.379 | 4.078 | 2.583 | 7.236 | 5.178 | 3.652 | 2.861 | 3.331 |
| D | - | - | - | - | 9.638 | 2.869 | 3.058 | 3.489 | 4.285 | 6.487 | 1.782 | 3.861 | 4.209 | 4.037 |
| E | - | - | - | - | - | 6.481 | 6.482 | 6.488 | 6.502 | 6.304 | 6.46 | 6.494 | 6.5 | 6.496 |
| F | - | - | - | - | - | - | 0.129 | 2.086 | 3.723 | 6.705 | 3.535 | 2.963 | 3.599 | 3.298 |
| G | - | - | - | - | - | - | - | 1.668 | 3.61 | 6.734 | 3.681 | 2.755 | 3.476 | 3.143 |
| H | - | - | - | - | - | - | - | - | 3.22 | 6.831 | 4.048 | 1.889 | 3.036 | 2.544 |
| I | - | - | - | - | - | - | - | - | - | 7.093 | 4.835 | 2.622 | 0.295 | 1.946 |
| J | - | - | - | - | - | - | - | - | - | - | 5.51 | 5.785 | 5.839 | 5.809 |
| K | - | - | - | - | - | - | - | - | - | - | - | 4.117 | 4.409 | 4.263 |
| L | - | - | - | - | - | - | - | - | - | - | - | - | 2.214 | 1.157 |
| M | - | - | - | - | - | - | - | - | - | - | - | - | - | 1.37 |
| N | - | - | - | - | - | - | - | - | - | - | - | - | - | - |
