## Supplementary Data S2 for "Interlaboratory comparison of standardised metabolomics and lipidomics analyses in human and rodent blood using the MxP® Quant 500 kit": TG(16.1_33.1).pdf

**Amount of relative differences between laboratories**

| Min | A | B | C | D | E | F | G | H | I | J | K | L | M | N |
|---|---|---|---|---|---|---|---|---|---|---|---|---|---|---|
| A | - | 6.002 | NA | 6.408 | 12.329 | 5.079 | 6.064 | 5.956 | 5.297 | 8.634 | 5.746 | 5.71 | 3.708 | 6.091 |
| B | - | - | NA | 4.929 | 12.185 | 1.898 | 5.937 | 4.475 | 6.159 | 7.937 | 2.672 | 5.7 | 6.363 | 4.646 |
| C | - | - | - | NA | NA | NA | NA | NA | NA | NA | NA | NA | NA | NA |
| D | - | - | - | - | 12.149 | 2.325 | 4.719 | 1.386 | 5.197 | 8.083 | 3.493 | 4.31 | 6.066 | 0.68 |
| E | - | - | - | - | - | 5.523 | 8.432 | 7.602 | 8.16 | 8.223 | 6.749 | 8.016 | 7.259 | 7.815 |
| F | - | - | - | - | - | - | 6.227 | 4.878 | 6.322 | 7.766 | 3.125 | 5.921 | 6.203 | 5.187 |
| G | - | - | - | - | - | - | - | 4.005 | 4.469 | 8.517 | 3.984 | 1.841 | 6.194 | 4.204 |
| H | - | - | - | - | - | - | - | - | 5.394 | 8.372 | 2.877 | 4.535 | 6.205 | -2.666 |
| I | - | - | - | - | - | - | - | - | - | 8.575 | 4.922 | 4.014 | 5.532 | 5.169 |
| J | - | - | - | - | - | - | - | - | - | - | 5.903 | 7.119 | 6.999 | 6.771 |
| K | - | - | - | - | - | - | - | - | - | - | - | 5.572 | 5.955 | 4.76 |
| L | - | - | - | - | - | - | - | - | - | - | - | - | 5.857 | 4.388 |
| M | - | - | - | - | - | - | - | - | - | - | - | - | - | 5.638 |
| N | - | - | - | - | - | - | - | - | - | - | - | - | - | - |

| | |
|---|---|
| | Larger than 10 |
| | Between 8 and 9,99 |
| | Between 6 and 7,99 |
| | Between 4 and 5,99 |
| | Between 2 and 3,99 |
| | Between 0 and 1,99 |
| | NA (Calculation failed) |

| Median | A | B | C | D | E | F | G | H | I | J | K | L | M | N |
|---|---|---|---|---|---|---|---|---|---|---|---|---|---|---|
| A | - | 4.65 | NA | 4.361 | 10.501 | 3.877 | 4.728 | 4.75 | 3.938 | 5.632 | 4.358 | 4.391 | 4.814 | 4.719 |
| B | - | - | NA | 3.401 | 10.352 | 3.069 | 1.591 | -0.717 | 3.451 | 4.217 | 3.625 | 3.502 | 6.079 | -5.489 |
| C | - | - | - | NA | NA | NA | NA | NA | NA | NA | NA | NA | NA | NA |
| D | - | - | - | - | 10.312 | 0.97 | 2.837 | 3.255 | -2.102 | 4.763 | -0.06 | -0.192 | 5.844 | 3.268 |
| E | - | - | - | - | - | 3.248 | 5.057 | 4.584 | 4.556 | 4.941 | 5.23 | 5.334 | 6.158 | 4.462 |
| F | - | - | - | - | - | - | 1.526 | 2.415 | 1.772 | 4.986 | 2.64 | 2.383 | 5.746 | 2.439 |
| G | - | - | - | - | - | - | - | 0.792 | 3.451 | 4.592 | 3.234 | 2.815 | 6.205 | 0.506 |
| H | - | - | - | - | - | - | - | - | 3.532 | 4.569 | 3.428 | 3.228 | 6.125 | -4.962 |
| I | - | - | - | - | - | - | - | - | - | 5.193 | 1.597 | 2.098 | 5.719 | 3.541 |
| J | - | - | - | - | - | - | - | - | - | - | 4.773 | 4.353 | 6.263 | 2.62 |
| K | - | - | - | - | - | - | - | - | - | - | - | -1.111 | 5.503 | 3.259 |
| L | - | - | - | - | - | - | - | - | - | - | - | - | 5.878 | 3.016 |
| M | - | - | - | - | - | - | - | - | - | - | - | - | - | 5.405 |
| N | - | - | - | - | - | - | - | - | - | - | - | - | - | - |

| Max | A | B | C | D | E | F | G | H | I | J | K | L | M | N |
|---|---|---|---|---|---|---|---|---|---|---|---|---|---|---|
| A | - | 3.795 | NA | 3.77 | 10.915 | 5.256 | 3.316 | 1.347 | 2.197 | 6.688 | 5.338 | 0.785 | 5.855 | 1.296 |
| B | - | - | NA | -5.749 | 10.778 | 4.414 | 4.379 | 3.846 | 4.017 | 6.337 | 4.543 | 3.77 | 5.267 | 3.834 |
| C | - | - | - | NA | NA | NA | NA | NA | NA | NA | NA | NA | NA | NA |
| D | - | - | - | - | 10.748 | 4.411 | 4.382 | 3.843 | 4.009 | 6.322 | 4.55 | 3.765 | 5.268 | 3.83 |
| E | - | - | - | - | - | 6.517 | 6.561 | 6.554 | 6.557 | 6.443 | 6.513 | 6.553 | 6.496 | 6.554 |
| F | - | - | - | - | - | - | 5.117 | 4.878 | 4.95 | 5.642 | 0.717 | 4.849 | 3.827 | 4.874 |
| G | - | - | - | - | - | - | - | 3.034 | 2.564 | 6.979 | 5.802 | 3.195 | 6.227 | 3.052 |
| H | - | - | - | - | - | - | - | - | 1.063 | 6.799 | 5.461 | -0.184 | 5.951 | -2.833 |
| I | - | - | - | - | - | - | - | - | - | 6.843 | 5.559 | 1.591 | 6.03 | 1.17 |
| J | - | - | - | - | - | - | - | - | - | - | 5.019 | 5.712 | 4.608 | 5.72 |
| K | - | - | - | - | - | - | - | - | - | - | - | 4.906 | 3.628 | 4.932 |
| L | - | - | - | - | - | - | - | - | - | - | - | - | 5.919 | -0.418 |
| M | - | - | - | - | - | - | - | - | - | - | - | - | - | 5.252 |
| N | - | - | - | - | - | - | - | - | - | - | - | - | - | - |
