## Supplementary Data S2 for "Interlaboratory comparison of standardised metabolomics and lipidomics analyses in human and rodent blood using the MxP® Quant 500 kit": TG(16.1_34.2).pdf

**Amount of relative differences between laboratories**

| Min | A | B | C | D | E | F | G | H | I | J | K | L | M | N |
|---|---|---|---|---|---|---|---|---|---|---|---|---|---|---|
| A | - | 5.286 | 4.691 | 4.999 | 6.096 | 5.457 | 4.78 | 4.665 | 3.996 | 8.332 | 4.704 | 5.017 | 4.72 | 5.068 |
| B | - | - | 3.721 | 2.546 | 5.417 | 3.091 | 3.165 | 3.129 | 4.068 | 7.907 | 3.069 | -3.276 | 2.794 | -0.21 |
| C | - | - | - | 2.749 | 5.65 | 4.31 | 1.821 | 1.742 | 2.175 | 7.927 | 1.878 | 3.481 | 2.285 | 3.436 |
| D | - | - | - | - | 5.501 | 3.736 | 1.419 | 1.18 | 3.238 | 7.774 | 0.99 | 2.523 | -0.872 | 2.291 |
| E | - | - | - | - | - | 5.636 | 6.116 | 6.058 | 6.207 | 7.049 | 6.069 | 5.795 | 5.996 | 5.853 |
| F | - | - | - | - | - | - | 4.173 | 4.13 | 4.672 | 7.838 | 4.105 | 3.072 | 3.955 | 3.235 |
| G | - | - | - | - | - | - | - | -3.876 | 2.999 | 7.904 | -1.831 | 3.004 | 0.615 | 2.902 |
| H | - | - | - | - | - | - | - | - | 3.068 | 8.021 | -1.244 | 3.038 | 0.662 | 2.982 |
| I | - | - | - | - | - | - | - | - | - | 8.377 | 3.417 | 4.176 | 3.487 | 4.24 |
| J | - | - | - | - | - | - | - | - | - | - | 5.193 | 4.781 | 5.045 | 4.947 |
| K | - | - | - | - | - | - | - | - | - | - | - | 2.946 | 0.356 | 2.827 |
| L | - | - | - | - | - | - | - | - | - | - | - | - | 2.798 | -0.63 |
| M | - | - | - | - | - | - | - | - | - | - | - | - | - | 2.765 |
| N | - | - | - | - | - | - | - | - | - | - | - | - | - | - |

| Median | A | B | C | D | E | F | G | H | I | J | K | L | M | N |
|---|---|---|---|---|---|---|---|---|---|---|---|---|---|---|
| A | - | 3.793 | 2.824 | 3.096 | 4.964 | 4.306 | 3.195 | 3.445 | 3.555 | 4.793 | 3.205 | 3.993 | 3.768 | 4.15 |
| B | - | - | 3.175 | 3.006 | 4.451 | 2.366 | 2.669 | 2.023 | 1.132 | 2.967 | 2.796 | 0.73 | -2.171 | 2.307 |
| C | - | - | - | 0.028 | 4.803 | 3.713 | 1.272 | 2.164 | 2.597 | 4 | 0.962 | 3.284 | 2.982 | 3.656 |
| D | - | - | - | - | 4.802 | 3.609 | 0.717 | 1.92 | 2.483 | 3.802 | 0.056 | 3.179 | 2.885 | 3.609 |
| E | - | - | - | - | - | 4.661 | 5.28 | 5.161 | 5.018 | 4.996 | 5.351 | 4.847 | 4.914 | 4.508 |
| F | - | - | - | - | - | - | 3.579 | 3.252 | 2.891 | 0.635 | 3.666 | 1.826 | 2.429 | -3.128 |
| G | - | - | - | - | - | - | - | 1.09 | 1.916 | 3.725 | -1.004 | 2.913 | 2.501 | 3.391 |
| H | - | - | - | - | - | - | - | - | 0.677 | 3.71 | 1.368 | 2.468 | 1.827 | 3.078 |
| I | - | - | - | - | - | - | - | - | - | 4.162 | 1.721 | 2.17 | 1.041 | 2.78 |
| J | - | - | - | - | - | - | - | - | - | - | 0.942 | 3.467 | 3.444 | 4.034 |
| K | - | - | - | - | - | - | - | - | - | - | - | 3.001 | 2.629 | 3.459 |
| L | - | - | - | - | - | - | - | - | - | - | - | - | 0.985 | 1.692 |
| M | - | - | - | - | - | - | - | - | - | - | - | - | - | 2.337 |
| N | - | - | - | - | - | - | - | - | - | - | - | - | - | - |

| Max | A | B | C | D | E | F | G | H | I | J | K | L | M | N |
|---|---|---|---|---|---|---|---|---|---|---|---|---|---|---|
| A | - | 2.736 | 3.565 | 4.155 | 1.959 | 2.259 | 3.393 | 2.433 | 2.873 | 6.955 | 3.601 | 1.467 | -1.159 | 1.998 |
| B | - | - | 2.274 | 3.393 | 1.125 | 0.792 | 1.831 | 0.269 | 3.717 | 6.806 | 2.354 | 1.872 | 2.748 | 3.324 |
| C | - | - | - | 2.431 | 2.774 | 2.65 | 0.28 | 2.526 | 4.102 | 6.661 | -2.028 | 3.021 | 3.463 | 3.824 |
| D | - | - | - | - | 3.522 | 3.469 | 2.651 | 3.405 | 4.419 | 6.519 | 2.286 | 3.68 | 3.962 | 4.214 |
| E | - | - | - | - | - | -1.102 | 2.616 | 0.144 | 3.483 | 6.865 | 2.894 | 0.692 | 2.228 | 2.964 |
| F | - | - | - | - | - | - | 2.435 | -0.89 | 3.537 | 6.853 | 2.8 | 0.983 | 2.344 | 3.073 |
| G | - | - | - | - | - | - | - | 2.202 | 4.014 | 6.692 | 0.573 | 2.802 | 3.313 | 3.714 |
| H | - | - | - | - | - | - | - | - | 3.597 | 6.833 | 2.676 | 1.315 | 2.48 | 3.158 |
| I | - | - | - | - | - | - | - | - | - | 7.172 | 4.394 | 3.448 | 2.889 | 1.853 |
| J | - | - | - | - | - | - | - | - | - | - | 5.665 | 5.777 | 5.815 | 5.847 |
| K | - | - | - | - | - | - | - | - | - | - | - | 3.061 | 3.493 | 3.848 |
| L | - | - | - | - | - | - | - | - | - | - | - | - | 1.664 | 2.725 |
| M | - | - | - | - | - | - | - | - | - | - | - | - | - | 1.822 |
| N | - | - | - | - | - | - | - | - | - | - | - | - | - | - |

| | |
|---|---|
| | Larger than 10 |
| | Between 8 and 9,99 |
| | Between 6 and 7,99 |
| | Between 4 and 5,99 |
| | Between 2 and 3,99 |
| | Between 0 and 1,99 |
| | NA (Calculation failed) |
