## Supplementary Data S2 for "Interlaboratory comparison of standardised metabolomics and lipidomics analyses in human and rodent blood using the MxP® Quant 500 kit": TG(16.1_34.3).pdf

**Amount of relative differences between laboratories**

| Min | A | B | C | D | E | F | G | H | I | J | K | L | M | N |
|---|---|---|---|---|---|---|---|---|---|---|---|---|---|---|
| A | - | 5.038 | 4.55 | 4.702 | 6.13 | 5.611 | 4.44 | 4.706 | 3.946 | 8.217 | 4.63 | 4.95 | 4.254 | 5.129 |
| B | - | - | 3.909 | 3.11 | 5.671 | 4.159 | 3.614 | 2.572 | 4.182 | 8.126 | 2.305 | -1.271 | 3.677 | 1.708 |
| C | - | - | - | 2.569 | 5.807 | 4.784 | 1.759 | 3.134 | 0.13 | 7.823 | 3.352 | 3.732 | 1.829 | 3.999 |
| D | - | - | - | - | 5.714 | 4.45 | 1.201 | 1.762 | 2.687 | 7.774 | 2.458 | 3.003 | 0.696 | 3.385 |
| E | - | - | - | - | - | 5.86 | 6.413 | 6.24 | 6.459 | 6.745 | 6.068 | 6.134 | 6.315 | 6.097 |
| F | - | - | - | - | - | - | 4.762 | 4.337 | 4.975 | 7.705 | 4.059 | 3.906 | 4.67 | 3.63 |
| G | - | - | - | - | - | - | - | 2.48 | 2.223 | 7.847 | 2.816 | 3.368 | -2.908 | 3.704 |
| H | - | - | - | - | - | - | - | - | 3.327 | 7.733 | 1.21 | 2.238 | 2.316 | 2.818 |
| I | - | - | - | - | - | - | - | - | - | 8.387 | 3.469 | 4.163 | 2.13 | 4.503 |
| J | - | - | - | - | - | - | - | - | - | - | 4.183 | 4.587 | 4.735 | 4.618 |
| K | - | - | - | - | - | - | - | - | - | - | - | 2.13 | 2.916 | 3.005 |
| L | - | - | - | - | - | - | - | - | - | - | - | - | 3.455 | 1.51 |
| M | - | - | - | - | - | - | - | - | - | - | - | - | - | 3.874 |
| N | - | - | - | - | - | - | - | - | - | - | - | - | - | - |

| | |
|---|---|
| (red) | Larger than 10 |
| (orange) | Between 8 and 9,99 |
| (light orange) | Between 6 and 7,99 |
| (light green) | Between 4 and 5,99 |
| (green) | Between 2 and 3,99 |
| (dark green) | Between 0 and 1,99 |
| (grey) | NA (Calculation failed) |

| Median | A | B | C | D | E | F | G | H | I | J | K | L | M | N |
|---|---|---|---|---|---|---|---|---|---|---|---|---|---|---|
| A | - | 4.273 | 2.134 | 2.649 | 5.101 | 4.516 | 2.82 | 3.137 | 3.8 | 4.916 | 3.957 | 3.946 | 3.339 | 4.154 |
| B | - | - | 4.318 | 4.057 | 4.333 | 1.611 | 3.993 | 3.788 | 2.647 | 2.981 | 2.206 | 2.37 | 3.606 | 0.483 |
| C | - | - | - | 1.55 | 5.08 | 4.256 | 1.805 | 2.388 | 3.559 | 4.351 | 3.708 | 3.646 | 2.726 | 3.983 |
| D | - | - | - | - | 5.009 | 4.042 | -0.906 | 1.209 | 3.22 | 4.107 | 3.404 | 3.309 | 1.902 | 3.747 |
| E | - | - | - | - | - | 4.869 | 5.682 | 5.614 | 5.164 | 5.476 | 5.122 | 5.204 | 5.554 | 4.886 |
| F | - | - | - | - | - | - | 4.22 | 4.052 | 2.962 | 1.39 | 2.679 | 2.892 | 3.895 | 1.527 |
| G | - | - | - | - | - | - | - | 0.85 | 3.1 | 4.108 | 3.31 | 3.228 | 1.689 | 3.672 |
| H | - | - | - | - | - | - | - | - | 2.857 | 3.816 | 3.089 | 2.967 | 0.542 | 3.52 |
| I | - | - | - | - | - | - | - | - | - | 4.403 | 0.864 | 0.73 | 2.194 | 2.198 |
| J | - | - | - | - | - | - | - | - | - | - | 4.117 | 3.965 | 3.147 | 4.408 |
| K | - | - | - | - | - | - | - | - | - | - | - | -0.911 | 2.855 | 1.623 |
| L | - | - | - | - | - | - | - | - | - | - | - | - | 2.759 | 1.915 |
| M | - | - | - | - | - | - | - | - | - | - | - | - | - | 3.263 |
| N | - | - | - | - | - | - | - | - | - | - | - | - | - | - |

| Max | A | B | C | D | E | F | G | H | I | J | K | L | M | N |
|---|---|---|---|---|---|---|---|---|---|---|---|---|---|---|
| A | - | 3.184 | 3.715 | 3.709 | 2.237 | -1.691 | 3.178 | 3.583 | 3.662 | 6.851 | 2.384 | -2.559 | 0.36 | 2.215 |
| B | - | - | 4.613 | 4.615 | 2.427 | 3.293 | 4.32 | 4.539 | 1.977 | 7.129 | 2.085 | 3.299 | 3.51 | 2.313 |
| C | - | - | - | -6.03 | 3.948 | 3.559 | 1.855 | -0.026 | 4.513 | 6.522 | 4.024 | 3.556 | 3.394 | 3.971 |
| D | - | - | - | - | 3.939 | 3.561 | 1.868 | 0.019 | 4.513 | 6.533 | 4.024 | 3.558 | 3.399 | 3.973 |
| E | - | - | - | - | - | 2.065 | 3.812 | 4.091 | 3.142 | 6.991 | 0.303 | 2.007 | 2.452 | -2.182 |
| F | - | - | - | - | - | - | 3.215 | 3.614 | 3.644 | 6.879 | 2.339 | -5.985 | 0.5 | 2.135 |
| G | - | - | - | - | - | - | - | 1.45 | 4.317 | 6.626 | 3.714 | 3.076 | 2.84 | 3.647 |
| H | - | - | - | - | - | - | - | - | 4.464 | 6.557 | 3.948 | 3.442 | 3.269 | 3.892 |
| I | - | - | - | - | - | - | - | - | - | 7.226 | 3.09 | 3.841 | 3.997 | 3.21 |
| J | - | - | - | - | - | - | - | - | - | - | 5.833 | 5.771 | 5.755 | 5.814 |
| K | - | - | - | - | - | - | - | - | - | - | - | 2.422 | 2.774 | -0.496 |
| L | - | - | - | - | - | - | - | - | - | - | - | - | 0.507 | 2.163 |
| M | - | - | - | - | - | - | - | - | - | - | - | - | - | 2.518 |
| N | - | - | - | - | - | - | - | - | - | - | - | - | - | - |
