## Supplementary Data S2 for "Interlaboratory comparison of standardised metabolomics and lipidomics analyses in human and rodent blood using the MxP® Quant 500 kit": TG(16.1_36.2).pdf

**Amount of relative differences between laboratories**

| Min | A | B | C | D | E | F | G | H | I | J | K | L | M | N |
|---|---|---|---|---|---|---|---|---|---|---|---|---|---|---|
| A | - | 6.357 | 6.256 | 6.544 | 6.748 | 6.628 | 6.294 | 5.904 | 4.584 | 9.763 | 6.088 | 6.264 | 5.844 | 6.105 |
| B | - | - | 3.84 | -1.661 | 5.359 | 4.233 | 3.769 | 4.305 | 5.698 | 9.543 | 4.419 | 2.826 | 4.518 | 3.522 |
| C | - | - | - | 3.808 | 5.643 | 4.919 | -0.641 | 2.607 | 5.204 | 9.45 | 2.779 | 2.628 | 3.188 | 0.85 |
| D | - | - | - | - | 5.298 | 4.172 | 3.604 | 3.966 | 5.441 | 9.309 | 4.187 | 2.398 | 4.219 | 3.051 |
| E | - | - | - | - | - | 4.934 | 6.154 | 6.141 | 6.682 | 9.369 | 6.281 | 5.913 | 6.221 | 5.967 |
| F | - | - | - | - | - | - | 5.2 | 5.366 | 6.242 | 9.5 | 5.483 | 4.831 | 5.488 | 5.026 |
| G | - | - | - | - | - | - | - | 2.636 | 5.184 | 9.363 | 2.889 | 2.505 | 3.212 | 0.333 |
| H | - | - | - | - | - | - | - | - | 4.999 | 9.54 | -2.047 | 3.719 | 1.546 | 3.009 |
| I | - | - | - | - | - | - | - | - | - | 10.073 | 5.629 | 5.927 | 5.254 | 5.696 |
| J | - | - | - | - | - | - | - | - | - | - | 7.358 | 7.19 | 7.192 | 7.092 |
| K | - | - | - | - | - | - | - | - | - | - | - | 3.61 | 0.99 | 3.007 |
| L | - | - | - | - | - | - | - | - | - | - | - | - | 3.961 | 2.181 |
| M | - | - | - | - | - | - | - | - | - | - | - | - | - | 3.559 |
| N | - | - | - | - | - | - | - | - | - | - | - | - | - | - |

| | |
|---|---|
| | Larger than 10 |
| | Between 8 and 9,99 |
| | Between 6 and 7,99 |
| | Between 4 and 5,99 |
| | Between 2 and 3,99 |
| | Between 0 and 1,99 |
| | NA (Calculation failed) |

| Median | A | B | C | D | E | F | G | H | I | J | K | L | M | N |
|---|---|---|---|---|---|---|---|---|---|---|---|---|---|---|
| A | - | 3.822 | 3.033 | 2.761 | 4.577 | 4.486 | 2.982 | 3.156 | 4.019 | 2.414 | 1.591 | 3.672 | 3.536 | 4.152 |
| B | - | - | 2.93 | 3.45 | 3.562 | 3.164 | 3.281 | 2.762 | 0.664 | 3.231 | 3.989 | 1.875 | 1.419 | 2.102 |
| C | - | - | - | 1.561 | 4.167 | 3.946 | 0.877 | -0.362 | 3.095 | 1.388 | 2.913 | 1.913 | 2.175 | 3.463 |
| D | - | - | - | - | 4.376 | 4.176 | 0.11 | 1.983 | 3.519 | -0.113 | 2.156 | 2.716 | 2.952 | 3.808 |
| E | - | - | - | - | - | 1.895 | 4.687 | 4.43 | 3.602 | 4.71 | 5.042 | 4.229 | 4.041 | 3.114 |
| F | - | - | - | - | - | - | 4.375 | 4.11 | 3.029 | 4.472 | 4.786 | 3.813 | 3.66 | 2.33 |
| G | - | - | - | - | - | - | - | 1.383 | 3.35 | 0.74 | 2.492 | 2.462 | 2.655 | 3.663 |
| H | - | - | - | - | - | - | - | - | 2.971 | 1.505 | 3.031 | 1.659 | 1.882 | 3.358 |
| I | - | - | - | - | - | - | - | - | - | 3.062 | 4.003 | 2.094 | 2.422 | 1.011 |
| J | - | - | - | - | - | - | - | - | - | - | 2.262 | 2.825 | 3.641 | 4.198 |
| K | - | - | - | - | - | - | - | - | - | - | - | 3.394 | 3.489 | 4.11 |
| L | - | - | - | - | - | - | - | - | - | - | - | - | -0.215 | 2.924 |
| M | - | - | - | - | - | - | - | - | - | - | - | - | - | 2.778 |
| N | - | - | - | - | - | - | - | - | - | - | - | - | - | - |

| Max | A | B | C | D | E | F | G | H | I | J | K | L | M | N |
|---|---|---|---|---|---|---|---|---|---|---|---|---|---|---|
| A | - | 1.727 | 3.222 | 4.145 | -0.51 | 1.799 | 3.604 | 2.664 | 3.807 | 6.775 | 4.164 | 2.593 | -0.783 | 2.365 |
| B | - | - | 2.561 | 3.819 | 1.888 | 2.707 | 3.105 | 1.534 | 4.069 | 6.779 | 3.818 | 1.415 | 1.435 | 3.035 |
| C | - | - | - | 2.953 | 3.185 | 3.554 | 1.345 | 1.502 | 4.422 | 6.584 | 2.953 | 1.613 | 3.024 | 3.734 |
| D | - | - | - | - | 3.964 | 4.178 | 2.276 | 3.295 | 4.758 | 6.38 | -5.223 | 3.328 | 3.882 | 4.286 |
| E | - | - | - | - | - | 1.549 | 3.681 | 2.783 | 3.763 | 6.878 | 4.211 | 2.721 | 0.016 | 2.222 |
| F | - | - | - | - | - | - | 4.017 | 3.335 | 3.463 | 6.924 | 4.463 | 3.298 | 2.027 | 0.841 |
| G | - | - | - | - | - | - | - | 2.392 | 4.55 | 6.483 | 2.336 | 2.449 | 3.383 | 3.953 |
| H | - | - | - | - | - | - | - | - | 4.259 | 6.676 | 3.444 | -2.151 | 2.443 | 3.432 |
| I | - | - | - | - | - | - | - | - | - | 7.255 | 5.215 | 4.545 | 4.08 | 3.366 |
| J | - | - | - | - | - | - | - | - | - | - | 5.539 | 5.699 | 5.771 | 5.823 |
| K | - | - | - | - | - | - | - | - | - | - | - | 3.325 | 3.881 | 4.289 |
| L | - | - | - | - | - | - | - | - | - | - | - | - | 2.386 | 3.405 |
| M | - | - | - | - | - | - | - | - | - | - | - | - | - | 2.496 |
| N | - | - | - | - | - | - | - | - | - | - | - | - | - | - |
