## Supplementary Data S2 for "Interlaboratory comparison of standardised metabolomics and lipidomics analyses in human and rodent blood using the MxP® Quant 500 kit": TG(16.1_36.3).pdf

**Amount of relative differences between laboratories**

| Min | A | B | C | D | E | F | G | H | I | J | K | L | M | N |
|---|---|---|---|---|---|---|---|---|---|---|---|---|---|---|
| A | - | 4.199 | 4.016 | 4.126 | 5.378 | 4.958 | 4.03 | 3.954 | 3.504 | 8.045 | 4.449 | 4.492 | 3.733 | 4.364 |
| B | - | - | 1.788 | 1.41 | 4.832 | 3.712 | 1.854 | 1.769 | 2.999 | 7.991 | 2.401 | 2.208 | 2.504 | 0.794 |
| C | - | - | - | -2.592 | 4.934 | 3.934 | -1.873 | -0.848 | 1.894 | 7.619 | 3.206 | 2.977 | 0.477 | 2.328 |
| D | - | - | - | - | 4.941 | 3.902 | -1.637 | -0.287 | 1.68 | 7.585 | 3.273 | 3.005 | 0.141 | 2.324 |
| E | - | - | - | - | - | 4.573 | 5.474 | 5.392 | 5.563 | 7.433 | 4.84 | 5.016 | 5.473 | 5.203 |
| F | - | - | - | - | - | - | 4.157 | 4.045 | 4.415 | 7.634 | 2.605 | 3.006 | 4.216 | 3.539 |
| G | - | - | - | - | - | - | - | -0.274 | 1.796 | 7.588 | 3.246 | 3.032 | 0.195 | 2.386 |
| H | - | - | - | - | - | - | - | - | 2.117 | 7.758 | 3.108 | 2.915 | 1.009 | 2.229 |
| I | - | - | - | - | - | - | - | - | - | 8.287 | 3.585 | 3.658 | 1.042 | 3.402 |
| J | - | - | - | - | - | - | - | - | - | - | 3.289 | 4.06 | 4.596 | 4.315 |
| K | - | - | - | - | - | - | - | - | - | - | - | 0.305 | 3.566 | 2.159 |
| L | - | - | - | - | - | - | - | - | - | - | - | - | 3.362 | 1.731 |
| M | - | - | - | - | - | - | - | - | - | - | - | - | - | 2.871 |
| N | - | - | - | - | - | - | - | - | - | - | - | - | - | - |

| Median | A | B | C | D | E | F | G | H | I | J | K | L | M | N |
|---|---|---|---|---|---|---|---|---|---|---|---|---|---|---|
| A | - | 3.231 | -0.179 | 0.85 | 4.584 | 3.672 | -2.556 | 1.868 | 3.556 | 2.301 | 3.549 | 3.258 | 2.501 | 3.544 |
| B | - | - | 3.255 | 3.656 | 3.993 | 1.845 | 3.356 | 2.72 | 1.353 | 3.996 | 0.897 | -2.591 | 2.14 | 1.271 |
| C | - | - | - | 1.529 | 4.529 | 3.588 | -0.78 | 1.415 | 3.471 | 3.1 | 3.34 | 3.156 | 2.215 | 3.454 |
| D | - | - | - | - | 4.658 | 3.861 | 1.196 | 2.435 | 3.771 | 2.33 | 3.648 | 3.528 | 2.868 | 3.754 |
| E | - | - | - | - | - | 3.811 | 4.921 | 4.719 | 3.921 | 5.403 | 4.11 | 4.17 | 4.576 | 3.946 |
| F | - | - | - | - | - | - | 3.839 | 3.418 | 0.066 | 4.56 | 1.24 | 1.807 | 3.077 | 0.283 |
| G | - | - | - | - | - | - | - | 1.687 | 3.535 | 3.036 | 3.409 | 3.24 | 2.372 | 3.519 |
| H | - | - | - | - | - | - | - | - | 3.108 | 3.401 | 2.961 | 2.675 | 1.031 | 3.085 |
| I | - | - | - | - | - | - | - | - | - | 4.06 | -7.547 | 1.314 | 2.792 | -3.406 |
| J | - | - | - | - | - | - | - | - | - | - | 4.387 | 4.483 | 4.159 | 4.594 |
| K | - | - | - | - | - | - | - | - | - | - | - | 0.507 | 2.632 | -0.255 |
| L | - | - | - | - | - | - | - | - | - | - | - | - | 2.227 | 1.143 |
| M | - | - | - | - | - | - | - | - | - | - | - | - | - | 2.707 |
| N | - | - | - | - | - | - | - | - | - | - | - | - | - | - |

| Max | A | B | C | D | E | F | G | H | I | J | K | L | M | N |
|---|---|---|---|---|---|---|---|---|---|---|---|---|---|---|
| A | - | 1.214 | 3.556 | 4.141 | 2.516 | 0.146 | 3.669 | 2.755 | 3.591 | 6.745 | -0.608 | -0.167 | -0.135 | 2.39 |
| B | - | - | 3.866 | 4.378 | 1.521 | 1.846 | 3.957 | 3.229 | 3.308 | 6.892 | 0.894 | 1.78 | 1.729 | 1.537 |
| C | - | - | - | 2.431 | 3.939 | 3.258 | -0.27 | 2.176 | 4.42 | 6.436 | 3.474 | 3.286 | 3.298 | 3.929 |
| D | - | - | - | - | 4.29 | 3.829 | 2.129 | 3.237 | 4.667 | 6.321 | 3.97 | 3.845 | 3.862 | 4.293 |
| E | - | - | - | - | - | 2.734 | 4.248 | 3.655 | 2.853 | 6.955 | 2.287 | 2.675 | 2.619 | -5.804 |
| F | - | - | - | - | - | - | 3.524 | 2.479 | 3.71 | 6.756 | 0.76 | -2.595 | -1.765 | 2.65 |
| G | - | - | - | - | - | - | - | 2.411 | 4.465 | 6.401 | 3.568 | 3.389 | 3.409 | 3.996 |
| H | - | - | - | - | - | - | - | - | 4.142 | 6.598 | 2.78 | 2.447 | 2.469 | 3.488 |
| I | - | - | - | - | - | - | - | - | - | 7.147 | 3.707 | 3.878 | 3.874 | 2.958 |
| J | - | - | - | - | - | - | - | - | - | - | 5.764 | 5.753 | 5.757 | 5.804 |
| K | - | - | - | - | - | - | - | - | - | - | - | 0.652 | 0.42 | 2.219 |
| L | - | - | - | - | - | - | - | - | - | - | - | - | -2.857 | 2.63 |
| M | - | - | - | - | - | - | - | - | - | - | - | - | - | 2.576 |
| N | - | - | - | - | - | - | - | - | - | - | - | - | - | - |

| | |
|---|---|
| | Larger than 10 |
| | Between 8 and 9,99 |
| | Between 6 and 7,99 |
| | Between 4 and 5,99 |
| | Between 2 and 3,99 |
| | Between 0 and 1,99 |
| | NA (Calculation failed) |
