## Supplementary Data S2 for "Interlaboratory comparison of standardised metabolomics and lipidomics analyses in human and rodent blood using the MxP® Quant 500 kit": TG(16.1_36.5).pdf

**Amount of relative differences between laboratories**

| Min | A | B | C | D | E | F | G | H | I | J | K | L | M | N |
|---|---|---|---|---|---|---|---|---|---|---|---|---|---|---|
| A | - | 4.396 | 0.58 | 4.544 | 5.924 | 4.95 | 4.256 | 4.711 | 3.468 | 6.05 | 5.249 | 4.506 | 4.525 | 4.605 |
| B | - | - | 4.478 | 1.052 | 5.441 | 3.206 | 1.771 | 2.108 | 3.188 | 5.612 | 3.958 | 2.024 | 1.03 | 2.062 |
| C | - | - | - | 4.336 | 5.752 | 4.692 | 3.974 | 4.462 | 3.434 | 5.803 | 4.968 | 4.428 | 4.334 | 4.451 |
| D | - | - | - | - | 5.313 | 2.701 | 2.495 | 0.98 | 3.377 | 5.394 | 3.551 | 1.399 | -3.39 | 1.215 |
| E | - | - | - | - | - | 5.458 | 5.903 | 5.643 | 5.979 | 2.467 | 5.35 | 5.516 | 5.68 | 5.572 |
| F | - | - | - | - | - | - | 3.584 | 2.123 | 4.007 | 4.931 | 2.103 | 1.635 | 2.534 | 1.774 |
| G | - | - | - | - | - | - | - | 2.832 | 2.197 | 5.446 | 4.021 | 2.96 | 2.438 | 2.906 |
| H | - | - | - | - | - | - | - | - | 3.605 | 5.272 | 3.265 | -0.247 | 0.815 | -0.972 |
| I | - | - | - | - | - | - | - | - | - | 6.001 | 4.89 | 3.696 | 3.654 | 3.844 |
| J | - | - | - | - | - | - | - | - | - | - | 4.2 | 3.753 | 4.257 | 3.971 |
| K | - | - | - | - | - | - | - | - | - | - | - | 2.057 | 3.024 | 2.457 |
| L | - | - | - | - | - | - | - | - | - | - | - | - | 1.231 | -2.936 |
| M | - | - | - | - | - | - | - | - | - | - | - | - | - | 1.134 |
| N | - | - | - | - | - | - | - | - | - | - | - | - | - | - |

| | |
|---|---|
| | Larger than 10 |
| | Between 8 and 9,99 |
| | Between 6 and 7,99 |
| | Between 4 and 5,99 |
| | Between 2 and 3,99 |
| | Between 0 and 1,99 |
| | NA (Calculation failed) |

| Median | A | B | C | D | E | F | G | H | I | J | K | L | M | N |
|---|---|---|---|---|---|---|---|---|---|---|---|---|---|---|
| A | - | 2.853 | 3.711 | 2.026 | 4.674 | 2.723 | -0.031 | 1.803 | 2.884 | 3.1 | -0.165 | 2.962 | 2.753 | 3.806 |
| B | - | - | 4.435 | 1.235 | 4.452 | -0.746 | 3.027 | 1.128 | -1.419 | -0.24 | 2.746 | 0.584 | -3.858 | 2.798 |
| C | - | - | - | 3.985 | 5.157 | 4.109 | 3.47 | 3.993 | 4.177 | 4 | 3.609 | 4.245 | 4.157 | 4.557 |
| D | - | - | - | - | 4.581 | 0.493 | 2.53 | -1.403 | 1.298 | 0.177 | 2.047 | 1.899 | 1.179 | 3.13 |
| E | - | - | - | - | - | 5.012 | 5.363 | 4.985 | 4.94 | 5.432 | 5.242 | 4.829 | 4.935 | 4.521 |
| F | - | - | - | - | - | - | 2.813 | -1.126 | 0.175 | 2.573 | 2.348 | 1.338 | 0.134 | 2.882 |
| G | - | - | - | - | - | - | - | 2.54 | 2.954 | 1.89 | 0.621 | 3.165 | 2.925 | 3.778 |
| H | - | - | - | - | - | - | - | - | 0.961 | 1.416 | 2.178 | 1.701 | 0.84 | 3.04 |
| I | - | - | - | - | - | - | - | - | - | 1.405 | 3.02 | -0.537 | -0.889 | 2.82 |
| J | - | - | - | - | - | - | - | - | - | - | 3.814 | 4.205 | 4.082 | 4.357 |
| K | - | - | - | - | - | - | - | - | - | - | - | 2.795 | 2.421 | 3.457 |
| L | - | - | - | - | - | - | - | - | - | - | - | - | 0.559 | 2.415 |
| M | - | - | - | - | - | - | - | - | - | - | - | - | - | 2.759 |
| N | - | - | - | - | - | - | - | - | - | - | - | - | - | - |

| Max | A | B | C | D | E | F | G | H | I | J | K | L | M | N |
|---|---|---|---|---|---|---|---|---|---|---|---|---|---|---|
| A | - | 2.553 | 4.796 | 3.583 | 2.526 | 3.972 | 4.231 | 3.701 | 1.978 | 6.674 | 4.835 | 2.703 | 2.907 | 2.144 |
| B | - | - | 4.379 | 2.596 | 3.291 | 3.263 | 3.626 | 2.788 | 3.21 | 6.59 | 4.493 | -0.056 | 0.729 | 3.252 |
| C | - | - | - | 3.608 | 4.67 | 3.228 | 2.801 | 3.522 | 4.638 | 5.918 | 0.461 | 4.044 | 3.983 | 4.649 |
| D | - | - | - | - | 3.911 | 1.705 | 2.56 | -0.358 | 3.856 | 6.393 | 3.944 | 2.255 | 2.057 | 3.878 |
| E | - | - | - | - | - | 4.43 | 4.62 | 4.201 | -1.41 | 6.851 | 5.16 | 3.562 | 3.644 | -2.445 |
| F | - | - | - | - | - | - | 1.392 | 1.247 | 4.108 | 6.247 | 3.556 | 2.986 | 2.832 | 4.117 |
| G | - | - | - | - | - | - | - | 2.283 | 4.27 | 6.186 | 3.154 | 3.338 | 3.249 | 4.283 |
| H | - | - | - | - | - | - | - | - | 3.922 | 6.356 | 3.865 | 2.474 | 2.291 | 3.941 |
| I | - | - | - | - | - | - | - | - | - | 6.848 | 5.127 | 3.469 | 3.587 | -1.671 |
| J | - | - | - | - | - | - | - | - | - | - | 5.214 | 5.616 | 5.599 | 5.754 |
| K | - | - | - | - | - | - | - | - | - | - | - | 4.136 | 4.1 | 4.712 |
| L | - | - | - | - | - | - | - | - | - | - | - | - | -0.496 | 3.377 |
| M | - | - | - | - | - | - | - | - | - | - | - | - | - | 3.46 |
| N | - | - | - | - | - | - | - | - | - | - | - | - | - | - |
