## Supplementary Data S2 for "Interlaboratory comparison of standardised metabolomics and lipidomics analyses in human and rodent blood using the MxP® Quant 500 kit": TG(16.1_38.3).pdf

**Amount of relative differences between laboratories**

| Min | A | B | C | D | E | F | G | H | I | J | K | L | M | N |
|---|---|---|---|---|---|---|---|---|---|---|---|---|---|---|
| A | - | 3.749 | 4.023 | 4.772 | 6.013 | 5.914 | 4.79 | 4.324 | 3.327 | 6.672 | 4.34 | 5.02 | 4.714 | 5.137 |
| B | - | - | -0.037 | 3.505 | 5.737 | 5.625 | 3.793 | 2.477 | 3.544 | 6.496 | 0.804 | 4.031 | 3.679 | 4.044 |
| C | - | - | - | 2.889 | 5.495 | 5.437 | 3.585 | 2.235 | 4.295 | 6.037 | 2.525 | 3.513 | 3.568 | 2.565 |
| D | - | - | - | - | 5.442 | 5.318 | 2.109 | 2.052 | 4.791 | 6.178 | 3.747 | 2.224 | 1.895 | -0.237 |
| E | - | - | - | - | - | 0.289 | 5.692 | 5.992 | 6.795 | 4.683 | 6.425 | 5.824 | 5.696 | 6.114 |
| F | - | - | - | - | - | - | 5.502 | 5.805 | 6.722 | 3.607 | 6.342 | 5.667 | 5.475 | 6.064 |
| G | - | - | - | - | - | - | - | 3.254 | 4.919 | 6.356 | 3.913 | 1.426 | 0.165 | 0.899 |
| H | - | - | - | - | - | - | - | - | 4.361 | 6.357 | 2.627 | 3.294 | 2.928 | 3.106 |
| I | - | - | - | - | - | - | - | - | - | 6.835 | 3.596 | 4.599 | 3.988 | 4.922 |
| J | - | - | - | - | - | - | - | - | - | - | 5.835 | 4.968 | 4.328 | 5.689 |
| K | - | - | - | - | - | - | - | - | - | - | - | 3.996 | 3.884 | 3.618 |
| L | - | - | - | - | - | - | - | - | - | - | - | - | 1.803 | 0.091 |
| M | - | - | - | - | - | - | - | - | - | - | - | - | - | 1.883 |
| N | - | - | - | - | - | - | - | - | - | - | - | - | - | - |

| | |
|---|---|
| | Larger than 10 |
| | Between 8 and 9,99 |
| | Between 6 and 7,99 |
| | Between 4 and 5,99 |
| | Between 2 and 3,99 |
| | Between 0 and 1,99 |
| | NA (Calculation failed) |

| Median | A | B | C | D | E | F | G | H | I | J | K | L | M | N |
|---|---|---|---|---|---|---|---|---|---|---|---|---|---|---|
| A | - | 3.1 | 0.696 | 3.555 | 5.127 | 4.938 | 4.307 | 3.323 | 4.036 | 3.714 | -2.274 | 4.434 | 4.246 | 4.294 |
| B | - | - | 3.467 | 1.673 | 4.891 | 4.601 | 3.672 | 1.042 | 3.187 | 4.37 | 3.334 | 3.864 | 3.597 | 3.56 |
| C | - | - | - | 3.641 | 5.229 | 4.987 | 4.456 | 3.63 | 4.237 | 2.484 | -1.472 | 4.562 | 4.443 | 4.317 |
| D | - | - | - | - | 4.815 | 4.444 | 3.39 | -0.991 | 2.779 | 4.48 | 3.851 | 3.62 | 3.33 | 3.104 |
| E | - | - | - | - | - | 3.345 | 4.672 | 5.358 | 4.925 | 6.064 | 5.975 | 4.532 | 4.68 | 4.898 |
| F | - | - | - | - | - | - | 3.75 | 4.779 | 4.149 | 5.626 | 5.602 | 3.464 | 3.722 | 4.177 |
| G | - | - | - | - | - | - | - | 3.616 | 2.053 | 5.27 | 4.729 | 1.087 | -0.515 | 0.342 |
| H | - | - | - | - | - | - | - | - | 2.858 | 4.484 | 3.701 | 3.67 | 3.376 | 3.267 |
| I | - | - | - | - | - | - | - | - | - | 5.164 | 4.239 | 2.637 | 1.603 | 2.148 |
| J | - | - | - | - | - | - | - | - | - | - | 2.866 | 4.439 | 4.382 | 3.891 |
| K | - | - | - | - | - | - | - | - | - | - | - | 4.55 | 4.415 | 4.331 |
| L | - | - | - | - | - | - | - | - | - | - | - | - | 1.486 | 1.812 |
| M | - | - | - | - | - | - | - | - | - | - | - | - | - | -1.261 |
| N | - | - | - | - | - | - | - | - | - | - | - | - | - | - |

| Max | A | B | C | D | E | F | G | H | I | J | K | L | M | N |
|---|---|---|---|---|---|---|---|---|---|---|---|---|---|---|
| A | - | -1.263 | 4.839 | 2.428 | 3.609 | 2.696 | 3.6 | 0.539 | 4.284 | 6.928 | 4.442 | 3.754 | 3.567 | 3.197 |
| B | - | - | 4.87 | 2.535 | 3.524 | 2.839 | 3.544 | 0.962 | 4.258 | 6.97 | 4.495 | 3.725 | 3.511 | 3.111 |
| C | - | - | - | 4.175 | 4.985 | 4.101 | 4.985 | 4.403 | 5.227 | 6.187 | 2.411 | 5.034 | 4.975 | 4.875 |
| D | - | - | - | - | 4.035 | -0.072 | 4.044 | 1.872 | 4.557 | 6.804 | 3.953 | 4.171 | 4.024 | 3.786 |
| E | - | - | - | - | - | 4.416 | -4.243 | 3.92 | 3.101 | 7.27 | 5.272 | 1.029 | -4.779 | 1.744 |
| F | - | - | - | - | - | - | 4.138 | 2.339 | 4.623 | 6.748 | 3.839 | 4.235 | 4.126 | 3.874 |
| G | - | - | - | - | - | - | - | 3.943 | 3.078 | 7.251 | 5.269 | 0.766 | -1.8 | 1.752 |
| H | - | - | - | - | - | - | - | - | 4.369 | 6.918 | 4.336 | 3.899 | 3.711 | 3.366 |
| I | - | - | - | - | - | - | - | - | - | 7.471 | 5.686 | 2.867 | 3.254 | 3.692 |
| J | - | - | - | - | - | - | - | - | - | - | 5.496 | 5.942 | 5.93 | 5.892 |
| K | - | - | - | - | - | - | - | - | - | - | - | 4.859 | 4.795 | 4.666 |
| L | - | - | - | - | - | - | - | - | - | - | - | - | 1.088 | 2.366 |
| M | - | - | - | - | - | - | - | - | - | - | - | - | - | 1.665 |
| N | - | - | - | - | - | - | - | - | - | - | - | - | - | - |
