## Supplementary Data S2 for "Interlaboratory comparison of standardised metabolomics and lipidomics analyses in human and rodent blood using the MxP® Quant 500 kit": TG(17.0_34.1).pdf

**Amount of relative differences between laboratories**

| Min | A | B | C | D | E | F | G | H | I | J | K | L | M | N |
|---|---|---|---|---|---|---|---|---|---|---|---|---|---|---|
| A | - | 5.795 | 5.384 | 7.578 | 8.238 | 6.168 | 5.84 | 5.51 | 2.589 | 9.053 | 5.295 | 5.946 | 5.646 | 5.619 |
| B | - | - | 4.038 | 7.288 | 8.274 | 4.987 | 3.218 | 4.857 | 5.997 | 9.03 | 3.984 | -0.476 | 3.06 | 4.121 |
| C | - | - | - | 7.298 | 8.186 | 5.419 | 3.145 | 3.076 | 5.445 | 8.958 | -3.983 | 4.124 | 2.994 | 0.048 |
| D | - | - | - | - | 6.648 | 3.718 | 6.402 | 6.763 | 6.729 | 6.667 | 5.88 | 6.162 | 6.075 | 6.428 |
| E | - | - | - | - | - | 6.302 | 7.585 | 7.838 | 7.685 | 6.483 | 7.142 | 7.431 | 7.321 | 7.58 |
| F | - | - | - | - | - | - | 5.982 | 6.442 | 6.701 | 8.382 | 5.696 | 5.611 | 5.691 | 6.095 |
| G | - | - | - | - | - | - | - | 4.108 | 5.49 | 8.708 | 1.714 | 3.194 | 1.081 | 2.695 |
| H | - | - | - | - | - | - | - | - | 4.803 | 8.737 | 3.3 | 4.532 | 3.989 | 3.163 |
| I | - | - | - | - | - | - | - | - | - | 9.347 | 5.218 | 6.025 | 5.667 | 5.678 |
| J | - | - | - | - | - | - | - | - | - | - | 6.135 | 6.473 | 6.338 | 6.701 |
| K | - | - | - | - | - | - | - | - | - | - | - | 3.208 | 1.893 | 2.394 |
| L | - | - | - | - | - | - | - | - | - | - | - | - | 2.837 | 4.078 |
| M | - | - | - | - | - | - | - | - | - | - | - | - | - | 3.133 |
| N | - | - | - | - | - | - | - | - | - | - | - | - | - | - |

| | |
|---|---|
| | Larger than 10 |
| | Between 8 and 9,99 |
| | Between 6 and 7,99 |
| | Between 4 and 5,99 |
| | Between 2 and 3,99 |
| | Between 0 and 1,99 |
| | NA (Calculation failed) |

| Median | A | B | C | D | E | F | G | H | I | J | K | L | M | N |
|---|---|---|---|---|---|---|---|---|---|---|---|---|---|---|
| A | - | 4.677 | 4.353 | 4.287 | 4.897 | 4.923 | 3.823 | 2.967 | 3.535 | 3.771 | 2.787 | 4.327 | 4.095 | 4.28 |
| B | - | - | 3.313 | 3.06 | 3.322 | 2.08 | 4.054 | 4.767 | 4.259 | 3.941 | 4.738 | 3.093 | 3.712 | 3.331 |
| C | - | - | - | -7.219 | 3.965 | 3.68 | 2.677 | 3.975 | 3.151 | 2.84 | 3.958 | 0.001 | 1.632 | -1.745 |
| D | - | - | - | - | 4.381 | 3.744 | 1.374 | 3.947 | 1.613 | 2.854 | 3.548 | 1.993 | -0.158 | 1.151 |
| E | - | - | - | - | - | 3.694 | 4.994 | 5.671 | 5 | 5.231 | 5.468 | 4.568 | 4.935 | 4.725 |
| F | - | - | - | - | - | - | 4.352 | 5.057 | 4.455 | 4.433 | 4.95 | 3.59 | 4.11 | 3.828 |
| G | - | - | - | - | - | - | - | 3.22 | 1.032 | 0.21 | 3.11 | 2.797 | 1.56 | 2.477 |
| H | - | - | - | - | - | - | - | - | 2.651 | 3.036 | -1.355 | 3.872 | 3.469 | 3.751 |
| I | - | - | - | - | - | - | - | - | - | 2.56 | 2.337 | 3.295 | 2.706 | 3.203 |
| J | - | - | - | - | - | - | - | - | - | - | 2.437 | 3.315 | 2.348 | 2.945 |
| K | - | - | - | - | - | - | - | - | - | - | - | 3.843 | 3.402 | 3.699 |
| L | - | - | - | - | - | - | - | - | - | - | - | - | 2.106 | 0.612 |
| M | - | - | - | - | - | - | - | - | - | - | - | - | - | 1.424 |
| N | - | - | - | - | - | - | - | - | - | - | - | - | - | - |

| Max | A | B | C | D | E | F | G | H | I | J | K | L | M | N |
|---|---|---|---|---|---|---|---|---|---|---|---|---|---|---|
| A | - | 3.531 | 2.714 | 6.004 | 6.383 | 2.52 | 2.609 | 3.545 | 3.846 | 7.011 | 4.325 | -2.428 | 1.291 | 2.51 |
| B | - | - | 2.517 | 6.47 | 6.841 | 4.295 | 4.325 | 4.718 | 1.68 | 7.355 | 5.159 | 3.695 | 3.993 | 2.738 |
| C | - | - | - | 6.264 | 6.63 | 3.719 | 3.761 | 4.285 | 3.078 | 7.192 | 4.829 | 2.778 | 3.271 | -0.13 |
| D | - | - | - | - | 3.611 | 5.188 | 5.166 | 5.03 | 5.597 | 5.219 | 4.785 | 5.326 | 5.267 | 5.427 |
| E | - | - | - | - | - | 5.469 | 5.438 | 5.352 | 5.777 | 4.538 | 5.188 | 5.569 | 5.525 | 5.636 |
| F | - | - | - | - | - | - | -1.554 | 2.48 | 4.253 | 6.901 | 3.753 | 2.479 | 1.632 | 3.437 |
| G | - | - | - | - | - | - | - | 2.386 | 4.276 | 6.868 | 3.71 | 2.56 | 1.783 | 3.477 |
| H | - | - | - | - | - | - | - | - | 4.546 | 6.743 | 2.903 | 3.403 | 3.04 | 3.959 |
| I | - | - | - | - | - | - | - | - | - | 7.426 | 5.331 | 4.056 | 4.3 | 3.35 |
| J | - | - | - | - | - | - | - | - | - | - | 5.602 | 5.838 | 5.807 | 5.891 |
| K | - | - | - | - | - | - | - | - | - | - | - | 4.07 | 3.871 | 4.421 |
| L | - | - | - | - | - | - | - | - | - | - | - | - | 1.392 | 2.478 |
| M | - | - | - | - | - | - | - | - | - | - | - | - | - | 2.996 |
| N | - | - | - | - | - | - | - | - | - | - | - | - | - | - |
