## Supplementary Data S2 for "Interlaboratory comparison of standardised metabolomics and lipidomics analyses in human and rodent blood using the MxP® Quant 500 kit": TG(17.1_32.1).pdf

**Amount of relative differences between laboratories**

| Min | A | B | C | D | E | F | G | H | I | J | K | L | M | N |
|---|---|---|---|---|---|---|---|---|---|---|---|---|---|---|
| A | - | 7.017 | 5.271 | 6.639 | NA | 5.409 | 5.543 | 6.307 | 4.667 | 9.062 | 5.794 | 5.863 | 4.393 | 5.637 |
| B | - | - | 3.23 | 3.912 | NA | 4.246 | 4.727 | 1.942 | 5.104 | 8.071 | 4.53 | 3.743 | 3.164 | 3.614 |
| C | - | - | - | 4.21 | NA | 4.271 | 5.307 | 3.72 | 5.54 | 8.505 | 4.636 | 4.541 | 4.373 | 4.596 |
| D | - | - | - | - | NA | 4.104 | 4.239 | 2.658 | 4.861 | 8.343 | 4.669 | 2.401 | 3.243 | 2.587 |
| E | - | - | - | - | - | NA | NA | NA | NA | NA | NA | NA | NA | NA |
| F | - | - | - | - | - | - | 6.553 | 6.003 | 6.645 | 8.064 | 2.547 | 6.233 | 5.801 | 6.152 |
| G | - | - | - | - | - | - | - | 4.726 | 3.685 | 8.627 | 5.137 | 3.741 | 0.468 | 3.339 |
| H | - | - | - | - | - | - | - | - | 5.296 | 8.484 | 4.351 | 3.677 | 4.138 | 3.797 |
| I | - | - | - | - | - | - | - | - | - | 8.988 | 5.417 | 4.938 | 2.423 | 4.597 |
| J | - | - | - | - | - | - | - | - | - | - | 4.635 | 6.737 | 5.771 | 6.568 |
| K | - | - | - | - | - | - | - | - | - | - | - | 5.933 | 5.499 | 5.828 |
| L | - | - | - | - | - | - | - | - | - | - | - | - | 3.497 | 1.528 |
| M | - | - | - | - | - | - | - | - | - | - | - | - | - | 2.93 |
| N | - | - | - | - | - | - | - | - | - | - | - | - | - | - |

| Legend |
|---|
| Larger than 10 |
| Between 8 and 9,99 |
| Between 6 and 7,99 |
| Between 4 and 5,99 |
| Between 2 and 3,99 |
| Between 0 and 1,99 |
| NA (Calculation failed) |

| Median | A | B | C | D | E | F | G | H | I | J | K | L | M | N |
|---|---|---|---|---|---|---|---|---|---|---|---|---|---|---|
| A | - | 3.977 | 4.127 | 4.575 | NA | 4.657 | 3.996 | 4.718 | 4.076 | 6.169 | 5.009 | 4.618 | 3.371 | 4.344 |
| B | - | - | 3.739 | 1.787 | NA | 3.881 | 0.686 | 3.586 | 3.083 | 4.923 | 4.239 | 3.315 | 0.662 | 2.58 |
| C | - | - | - | 3.387 | NA | 3.291 | 3.827 | 1.332 | 2.424 | 5.021 | 3.79 | 1.956 | 3.65 | 3.168 |
| D | - | - | - | - | NA | 3.311 | 1.227 | 3.052 | 2.052 | 4.885 | 4.028 | 2.728 | 1.72 | 1.274 |
| E | - | - | - | - | - | NA | NA | NA | NA | NA | NA | NA | NA | NA |
| F | - | - | - | - | - | - | 4.704 | 3.396 | 4.001 | 4.847 | 0.426 | 3.941 | 4.795 | 4.401 |
| G | - | - | - | - | - | - | - | 3.335 | 2.43 | 5.219 | 4.314 | 3.197 | -5.768 | 2.3 |
| H | - | - | - | - | - | - | - | - | 2.217 | 4.643 | 3.244 | 0.78 | 3.587 | 2.661 |
| I | - | - | - | - | - | - | - | - | - | 5.381 | 4.198 | 2.466 | 2.283 | 0.02 |
| J | - | - | - | - | - | - | - | - | - | - | 1.272 | 3.173 | 3.791 | 3.753 |
| K | - | - | - | - | - | - | - | - | - | - | - | 3.799 | 4.594 | 4.229 |
| L | - | - | - | - | - | - | - | - | - | - | - | - | 3.39 | 2.217 |
| M | - | - | - | - | - | - | - | - | - | - | - | - | - | 2.313 |
| N | - | - | - | - | - | - | - | - | - | - | - | - | - | - |

| Max | A | B | C | D | E | F | G | H | I | J | K | L | M | N |
|---|---|---|---|---|---|---|---|---|---|---|---|---|---|---|
| A | - | 5.029 | 3.052 | 3.939 | NA | 4.987 | 2.319 | 2.156 | 3.57 | 7.021 | 4.328 | 0.741 | 2.866 | 2.124 |
| B | - | - | 4.185 | 3.701 | NA | -0.393 | 4.379 | 4.409 | 5.066 | 6.204 | 3.233 | 4.69 | 4.25 | 4.41 |
| C | - | - | - | 2.658 | NA | 4.435 | 1.654 | 1.883 | 4.222 | 6.805 | 3.449 | 3.219 | -0.076 | 1.904 |
| D | - | - | - | - | NA | 3.816 | 3.163 | 3.24 | 4.557 | 6.632 | 2.05 | 3.88 | 2.791 | 3.245 |
| E | - | - | - | - | - | NA | NA | NA | NA | NA | NA | NA | NA | NA |
| F | - | - | - | - | - | - | 4.342 | 4.371 | 5.048 | 6.237 | 3.13 | 4.663 | 4.214 | 4.375 |
| G | - | - | - | - | - | - | - | -0.838 | 4.004 | 6.895 | 3.851 | 2.66 | 1.163 | -0.8 |
| H | - | - | - | - | - | - | - | - | 3.961 | 6.911 | 3.912 | 2.538 | 1.479 | -5.395 |
| I | - | - | - | - | - | - | - | - | - | 7.337 | 5.184 | 3.53 | 4.445 | 4.203 |
| J | - | - | - | - | - | - | - | - | - | - | 5.583 | 5.842 | 5.745 | 5.775 |
| K | - | - | - | - | - | - | - | - | - | - | - | 4.182 | 3.42 | 3.714 |
| L | - | - | - | - | - | - | - | - | - | - | - | - | 3.194 | 2.616 |
| M | - | - | - | - | - | - | - | - | - | - | - | - | - | 1.46 |
| N | - | - | - | - | - | - | - | - | - | - | - | - | - | - |
