## Supplementary Data S2 for "Interlaboratory comparison of standardised metabolomics and lipidomics analyses in human and rodent blood using the MxP® Quant 500 kit": TG(17.1_34.1).pdf

**Amount of relative differences between laboratories**

| Min | A | B | C | D | E | F | G | H | I | J | K | L | M | N |
|---|---|---|---|---|---|---|---|---|---|---|---|---|---|---|
| A | - | 5.736 | 5.734 | 6.062 | 6.612 | 6.665 | 5.507 | 5.765 | 4.35 | 9.332 | 5.418 | 5.587 | 5.49 | 5.379 |
| B | - | - | 4.13 | 1.512 | 5.862 | 5.429 | 4.535 | 3.096 | 5.215 | 9.053 | 4.39 | 4.035 | 4.159 | 4.336 |
| C | - | - | - | 3.701 | 6.005 | 5.725 | 2.349 | 2.896 | 4.205 | 8.872 | 1.87 | -0.984 | -0.297 | 1.556 |
| D | - | - | - | - | 5.834 | 5.366 | 4.09 | 1.8 | 4.753 | 8.775 | 3.827 | 3.423 | 3.532 | 3.709 |
| E | - | - | - | - | - | 5.272 | 6.788 | 6.492 | 6.75 | 7.979 | 6.669 | 6.653 | 6.626 | 6.613 |
| F | - | - | - | - | - | - | 6.154 | 5.725 | 6.261 | 8.62 | 6.033 | 5.953 | 5.953 | 5.979 |
| G | - | - | - | - | - | - | - | 3.619 | 3.924 | 8.983 | -0.133 | 2.45 | 1.912 | 0.427 |
| H | - | - | - | - | - | - | - | - | 4.876 | 9.084 | 3.627 | 2.948 | 3.22 | 3.552 |
| I | - | - | - | - | - | - | - | - | - | 9.593 | 4.681 | 4.999 | 4.821 | 4.617 |
| J | - | - | - | - | - | - | - | - | - | - | 6.371 | 6.387 | 6.33 | 6.273 |
| K | - | - | - | - | - | - | - | - | - | - | - | 2.087 | 1.464 | -0.915 |
| L | - | - | - | - | - | - | - | - | - | - | - | - | 0.906 | 2.17 |
| M | - | - | - | - | - | - | - | - | - | - | - | - | - | 1.508 |
| N | - | - | - | - | - | - | - | - | - | - | - | - | - | - |

| Median | A | B | C | D | E | F | G | H | I | J | K | L | M | N |
|---|---|---|---|---|---|---|---|---|---|---|---|---|---|---|
| A | - | 4.364 | 3.153 | 3.424 | 4.919 | 4.845 | 3.171 | 4.218 | 4.079 | 2.217 | 2.928 | 3.913 | 4.091 | 4.169 |
| B | - | - | 3.846 | 3.614 | 3.852 | 3.438 | 3.747 | 1.614 | 1.674 | 4.075 | 3.901 | 2.58 | 1.165 | 0.243 |
| C | - | - | - | 0.921 | 4.636 | 4.46 | -0.312 | 3.311 | 3.272 | 1.936 | -0.784 | 2.866 | 3.394 | 3.517 |
| D | - | - | - | - | 4.584 | 4.366 | -0.185 | 3.024 | 3.055 | 2.189 | 1.114 | 2.507 | 3.203 | 3.351 |
| E | - | - | - | - | - | 2.494 | 5.064 | 4.546 | 4.408 | 5.31 | 5.132 | 4.662 | 4.264 | 4.155 |
| F | - | - | - | - | - | - | 4.744 | 4.02 | 3.926 | 5.013 | 4.827 | 4.22 | 3.77 | 3.638 |
| G | - | - | - | - | - | - | - | 3.229 | 3.157 | 2.103 | 0.493 | 2.72 | 3.275 | 3.408 |
| H | - | - | - | - | - | - | - | - | -1.458 | 3.807 | 3.555 | 1.592 | -0.998 | 0.634 |
| I | - | - | - | - | - | - | - | - | - | 3.493 | 3.485 | 1.152 | -1.984 | 0.472 |
| J | - | - | - | - | - | - | - | - | - | - | 1.749 | 3.331 | 3.945 | 4.049 |
| K | - | - | - | - | - | - | - | - | - | - | - | 2.971 | 3.455 | 3.572 |
| L | - | - | - | - | - | - | - | - | - | - | - | - | 1.713 | 2.099 |
| M | - | - | - | - | - | - | - | - | - | - | - | - | - | 0.086 |
| N | - | - | - | - | - | - | - | - | - | - | - | - | - | - |

| Max | A | B | C | D | E | F | G | H | I | J | K | L | M | N |
|---|---|---|---|---|---|---|---|---|---|---|---|---|---|---|
| A | - | -1.874 | 3.716 | 4.096 | 3.791 | 1.901 | 3.163 | -1.216 | 3.906 | 6.966 | 3.574 | 0.177 | 2.166 | 2.849 |
| B | - | - | 3.68 | 4.079 | 3.815 | 1.828 | 3.108 | -2.831 | 3.931 | 6.999 | 3.535 | -0.205 | 2.259 | 2.904 |
| C | - | - | - | 1.857 | -0.307 | 3.042 | 1.892 | 3.491 | 4.64 | 6.637 | 0.172 | 3.408 | 3.965 | 4.17 |
| D | - | - | - | - | 1.282 | 3.514 | 2.822 | 3.84 | 4.78 | 6.558 | 2.174 | 3.78 | 4.214 | 4.38 |
| E | - | - | - | - | - | 3.186 | 2.174 | 3.596 | 4.685 | 6.627 | 0.986 | 3.543 | 4.051 | 4.224 |
| F | - | - | - | - | - | - | 2.3 | 1.718 | 4.186 | 6.89 | 2.958 | 1.387 | 3.015 | 3.409 |
| G | - | - | - | - | - | - | - | 2.968 | 4.46 | 6.751 | 1.43 | 2.837 | 3.626 | 3.892 |
| H | - | - | - | - | - | - | - | - | 3.944 | 6.987 | 3.513 | -0.501 | 2.298 | 2.927 |
| I | - | - | - | - | - | - | - | - | - | 7.41 | 4.984 | 4.248 | 3.626 | 3.197 |
| J | - | - | - | - | - | - | - | - | - | - | 5.678 | 5.809 | 5.871 | 5.891 |
| K | - | - | - | - | - | - | - | - | - | - | - | 3.263 | 3.873 | 4.094 |
| L | - | - | - | - | - | - | - | - | - | - | - | - | 2.485 | 3.051 |
| M | - | - | - | - | - | - | - | - | - | - | - | - | - | 1.507 |
| N | - | - | - | - | - | - | - | - | - | - | - | - | - | - |

| Colour | Legend |
|---|---|
| (red) | Larger than 10 |
| (orange) | Between 8 and 9,99 |
| (light orange) | Between 6 and 7,99 |
| (light green) | Between 4 and 5,99 |
| (green) | Between 2 and 3,99 |
| (dark green) | Between 0 and 1,99 |
| (grey) | NA (Calculation failed) |
