## Supplementary Data S2 for "Interlaboratory comparison of standardised metabolomics and lipidomics analyses in human and rodent blood using the MxP® Quant 500 kit": TG(17.1_38.6).pdf

**Amount of relative differences between laboratories**

| Min | A | B | C | D | E | F | G | H | I | J | K | L | M | N |
|---|---|---|---|---|---|---|---|---|---|---|---|---|---|---|
| A | - | 6.081 | 5.539 | 3.857 | NA | 6.245 | 2.231 | NA | 5.764 | 5.8 | 6.163 | 3.955 | 3.941 | 4.864 |
| B | - | - | 5.326 | 4.909 | NA | 3.602 | 5.488 | NA | 2.326 | 3.472 | 1.833 | 5.994 | 4.857 | 3.31 |
| C | - | - | - | 1.87 | NA | 5.454 | 2.766 | NA | 4.546 | 4.448 | 5.17 | 4.253 | 2.27 | 4.247 |
| D | - | - | - | - | NA | 5.942 | 3.415 | NA | 4.805 | 4.916 | 5.437 | 4.096 | -0.03 | 4.036 |
| E | - | - | - | - | - | NA | NA | NA | NA | NA | NA | NA | NA | NA |
| F | - | - | - | - | - | - | 6.161 | NA | 4.97 | 4.994 | 3.94 | 6.631 | 5.715 | 4.861 |
| G | - | - | - | - | - | - | - | NA | 5.574 | 5.696 | 6.071 | 2.322 | 3.462 | 4.675 |
| H | - | - | - | - | - | - | - | - | NA | NA | NA | NA | NA | NA |
| I | - | - | - | - | - | - | - | - | - | -3.7 | 3.83 | 5.894 | 4.975 | 3.533 |
| J | - | - | - | - | - | - | - | - | - | - | 3.578 | 5.47 | 3.427 | 2.512 |
| K | - | - | - | - | - | - | - | - | - | - | - | 6.266 | 5.331 | 4.079 |
| L | - | - | - | - | - | - | - | - | - | - | - | - | 3.506 | 4.688 |
| M | - | - | - | - | - | - | - | - | - | - | - | - | - | 3.931 |
| N | - | - | - | - | - | - | - | - | - | - | - | - | - | - |

| | |
|---|---|
| | Larger than 10 |
| | Between 8 and 9,99 |
| | Between 6 and 7,99 |
| | Between 4 and 5,99 |
| | Between 2 and 3,99 |
| | Between 0 and 1,99 |
| | NA (Calculation failed) |

| Median | A | B | C | D | E | F | G | H | I | J | K | L | M | N |
|---|---|---|---|---|---|---|---|---|---|---|---|---|---|---|
| A | - | 2.284 | 4.603 | 2.213 | NA | 3.997 | 1.406 | NA | 3.959 | 4.673 | 4.008 | 3.601 | 2.097 | 3.138 |
| B | - | - | 4.993 | 4.237 | NA | 3.298 | 3.898 | NA | 3.194 | 4.655 | 2.435 | 4.326 | 4.151 | -0.314 |
| C | - | - | - | 3.346 | NA | 5.031 | 3.726 | NA | 4.952 | 2.197 | 4.938 | 3.161 | 3.473 | 4.571 |
| D | - | - | - | - | NA | 4.688 | 1.629 | NA | 4.48 | 3.268 | 4.482 | 1.527 | -0.379 | 3.864 |
| E | - | - | - | - | - | NA | NA | NA | NA | NA | NA | NA | NA | NA |
| F | - | - | - | - | - | - | 4.94 | NA | 1.639 | 5.289 | 2.661 | 5.158 | 5.088 | 3.801 |
| G | - | - | - | - | - | - | - | NA | 4.235 | 4.225 | 4.319 | 3.03 | 0.867 | 3.525 |
| H | - | - | - | - | - | - | - | - | NA | NA | NA | NA | NA | NA |
| I | - | - | - | - | - | - | - | - | - | 5.324 | 1.515 | 5.011 | 4.812 | 3.35 |
| J | - | - | - | - | - | - | - | - | - | - | 4.493 | 1.428 | 1.822 | 4.159 |
| K | - | - | - | - | - | - | - | - | - | - | - | 4.865 | 4.786 | 2.846 |
| L | - | - | - | - | - | - | - | - | - | - | - | - | 2.278 | 4.151 |
| M | - | - | - | - | - | - | - | - | - | - | - | - | - | 3.786 |
| N | - | - | - | - | - | - | - | - | - | - | - | - | - | - |

| Max | A | B | C | D | E | F | G | H | I | J | K | L | M | N |
|---|---|---|---|---|---|---|---|---|---|---|---|---|---|---|
| A | - | 6.133 | 6.838 | 5.821 | NA | 6.061 | 3.909 | NA | 4.819 | 7.309 | 5.964 | 5.286 | 5.816 | 5.316 |
| B | - | - | 4.415 | 3.522 | NA | 2.735 | 5.154 | NA | 4.651 | 5.594 | 3.003 | 4.486 | 3.586 | 4.394 |
| C | - | - | - | 4.799 | NA | 4.114 | 5.554 | NA | 5.305 | 4.565 | 4.531 | 5.177 | 4.795 | 5.137 |
| D | - | - | - | - | NA | 4.56 | 4.715 | NA | 3.995 | 6.227 | 2.873 | 3.453 | -2.234 | 3.342 |
| E | - | - | - | - | - | NA | NA | NA | NA | NA | NA | NA | NA | NA |
| F | - | - | - | - | - | - | 5.254 | NA | 4.845 | 5.672 | 2.685 | 4.7 | 3.914 | 4.477 |
| G | - | - | - | - | - | - | - | NA | 3.772 | 7.07 | 5.551 | 4.391 | 5.165 | 4.465 |
| H | - | - | - | - | - | - | - | - | NA | NA | NA | NA | NA | NA |
| I | - | - | - | - | - | - | - | - | - | 6.725 | 4.755 | 2.522 | 4.261 | 2.785 |
| J | - | - | - | - | - | - | - | - | - | - | 5.227 | 5.598 | 5.373 | 5.554 |
| K | - | - | - | - | - | - | - | - | - | - | - | 4.119 | 2.575 | 4.007 |
| L | - | - | - | - | - | - | - | - | - | - | - | - | 3.634 | 0.516 |
| M | - | - | - | - | - | - | - | - | - | - | - | - | - | 3.368 |
| N | - | - | - | - | - | - | - | - | - | - | - | - | - | - |
