## Supplementary Data S2 for "Interlaboratory comparison of standardised metabolomics and lipidomics analyses in human and rodent blood using the MxP® Quant 500 kit": TG(17.1_38.7).pdf

**Amount of relative differences between laboratories**

| Min | A | B | C | D | E | F | G | H | I | J | K | L | M | N |
|---|---|---|---|---|---|---|---|---|---|---|---|---|---|---|
| A | - | 7.736 | NA | 4.217 | 5.301 | NA | 6.205 | NA | NA | NA | 9.808 | 4.988 | NA | 4.855 |
| B | - | - | NA | 7.415 | 7.966 | NA | 8.306 | NA | NA | NA | 8.132 | 7.628 | NA | 6.197 |
| C | - | - | - | NA | NA | NA | NA | NA | NA | NA | NA | NA | NA | NA |
| D | - | - | - | - | 3.91 | NA | 5.585 | NA | NA | NA | 9.716 | 3.666 | NA | 5.193 |
| E | - | - | - | - | - | NA | 5.243 | NA | NA | NA | 9.395 | 3.365 | NA | 5.516 |
| F | - | - | - | - | - | - | NA | NA | NA | NA | NA | NA | NA | NA |
| G | - | - | - | - | - | - | - | NA | NA | NA | 9.39 | 4.194 | NA | 5.767 |
| H | - | - | - | - | - | - | - | - | NA | NA | NA | NA | NA | NA |
| I | - | - | - | - | - | - | - | - | - | NA | NA | NA | NA | NA |
| J | - | - | - | - | - | - | - | - | - | - | NA | NA | NA | NA |
| K | - | - | - | - | - | - | - | - | - | - | - | 5.572 | NA | 3.174 |
| L | - | - | - | - | - | - | - | - | - | - | - | - | NA | 5.487 |
| M | - | - | - | - | - | - | - | - | - | - | - | - | - | NA |
| N | - | - | - | - | - | - | - | - | - | - | - | - | - | - |

| | |
|---|---|
| | Larger than 10 |
| | Between 8 and 9,99 |
| | Between 6 and 7,99 |
| | Between 4 and 5,99 |
| | Between 2 and 3,99 |
| | Between 0 and 1,99 |
| | NA (Calculation failed) |

| Median | A | B | C | D | E | F | G | H | I | J | K | L | M | N |
|---|---|---|---|---|---|---|---|---|---|---|---|---|---|---|
| A | - | 5.357 | NA | 4.177 | 3.553 | NA | 5.286 | NA | NA | NA | 5.46 | 2.742 | NA | 4.248 |
| B | - | - | NA | 6.984 | 6.859 | NA | 7.437 | NA | NA | NA | 6.673 | 6.78 | NA | 5.844 |
| C | - | - | - | NA | NA | NA | NA | NA | NA | NA | NA | NA | NA | NA |
| D | - | - | - | - | 2.71 | NA | 4.185 | NA | NA | NA | 3.416 | 3.164 | NA | 4.881 |
| E | - | - | - | - | - | NA | 4.726 | NA | NA | NA | 4.925 | 1.288 | NA | 4.712 |
| F | - | - | - | - | - | - | NA | NA | NA | NA | NA | NA | NA | NA |
| G | - | - | - | - | - | - | - | NA | NA | NA | 4.042 | 4.541 | NA | 5.245 |
| H | - | - | - | - | - | - | - | - | NA | NA | NA | NA | NA | NA |
| I | - | - | - | - | - | - | - | - | - | NA | NA | NA | NA | NA |
| J | - | - | - | - | - | - | - | - | - | - | NA | NA | NA | NA |
| K | - | - | - | - | - | - | - | - | - | - | - | 2.12 | NA | 4.245 |
| L | - | - | - | - | - | - | - | - | - | - | - | - | NA | 4.169 |
| M | - | - | - | - | - | - | - | - | - | - | - | - | - | NA |
| N | - | - | - | - | - | - | - | - | - | - | - | - | - | - |

| Max | A | B | C | D | E | F | G | H | I | J | K | L | M | N |
|---|---|---|---|---|---|---|---|---|---|---|---|---|---|---|
| A | - | 6.035 | NA | 4.06 | 4.249 | NA | 3.163 | NA | NA | NA | 8.914 | 5.269 | NA | 4.675 |
| B | - | - | NA | 2.717 | 5.023 | NA | 4.628 | NA | NA | NA | 4.24 | 5.611 | NA | 3.77 |
| C | - | - | - | NA | NA | NA | NA | NA | NA | NA | NA | NA | NA | NA |
| D | - | - | - | - | 4.931 | NA | 4.523 | NA | NA | NA | 8.671 | 5.587 | NA | 1.282 |
| E | - | - | - | - | - | NA | 3.419 | NA | NA | NA | 8.485 | 4.671 | NA | 5.938 |
| F | - | - | - | - | - | - | NA | NA | NA | NA | NA | NA | NA | NA |
| G | - | - | - | - | - | - | - | NA | NA | NA | 8.395 | 5.011 | NA | 5.174 |
| H | - | - | - | - | - | - | - | - | NA | NA | NA | NA | NA | NA |
| I | - | - | - | - | - | - | - | - | - | NA | NA | NA | NA | NA |
| J | - | - | - | - | - | - | - | - | - | - | NA | NA | NA | NA |
| K | - | - | - | - | - | - | - | - | - | - | - | 6.204 | NA | 5.986 |
| L | - | - | - | - | - | - | - | - | - | - | - | - | NA | 6.676 |
| M | - | - | - | - | - | - | - | - | - | - | - | - | - | NA |
| N | - | - | - | - | - | - | - | - | - | - | - | - | - | - |
