## Supplementary Data S2 for "Interlaboratory comparison of standardised metabolomics and lipidomics analyses in human and rodent blood using the MxP® Quant 500 kit": TG(17.2_34.3).pdf

**Amount of relative differences between laboratories**

| Min | A | B | C | D | E | F | G | H | I | J | K | L | M | N |
|---|---|---|---|---|---|---|---|---|---|---|---|---|---|---|
| A | - | 6.469 | 5.834 | 6.093 | 6.287 | NA | 5.295 | 5.615 | 5.919 | 6.536 | 5.798 | 5.02 | 5.819 | 6.013 |
| B | - | - | 4.069 | 5.893 | -0.255 | NA | 5.672 | 2.558 | 5.694 | 3.408 | 1.79 | 6.217 | 5.746 | 3.979 |
| C | - | - | - | 5.897 | 3.886 | NA | 6.303 | 4.488 | 6.273 | 4.632 | 4.698 | 6.727 | 6.092 | 2.147 |
| D | - | - | - | - | 5.214 | NA | 2.877 | 4.653 | -0.92 | 5.158 | 4.256 | 4.551 | 1.208 | 5.275 |
| E | - | - | - | - | - | NA | 6.297 | 4.206 | 5.828 | -2.038 | 1.339 | 6.735 | 6.019 | 1.531 |
| F | - | - | - | - | - | - | NA | NA | NA | NA | NA | NA | NA | NA |
| G | - | - | - | - | - | - | - | 4.349 | 5.952 | 7.123 | 6.488 | 3.065 | 5.626 | 5.405 |
| H | - | - | - | - | - | - | - | - | 5.003 | 0.351 | 4.666 | 6.145 | 5.374 | 4.37 |
| I | - | - | - | - | - | - | - | - | - | 5.858 | 4.803 | 5.451 | 2.604 | 4.891 |
| J | - | - | - | - | - | - | - | - | - | - | 1.95 | 5.812 | 5.243 | 4.551 |
| K | - | - | - | - | - | - | - | - | - | - | - | 6.184 | 5.458 | 4.384 |
| L | - | - | - | - | - | - | - | - | - | - | - | - | 5.839 | 5.517 |
| M | - | - | - | - | - | - | - | - | - | - | - | - | - | 5.238 |
| N | - | - | - | - | - | - | - | - | - | - | - | - | - | - |

| | |
|---|---|
| | Larger than 10 |
| | Between 8 and 9,99 |
| | Between 6 and 7,99 |
| | Between 4 and 5,99 |
| | Between 2 and 3,99 |
| | Between 0 and 1,99 |
| | NA (Calculation failed) |

| Median | A | B | C | D | E | F | G | H | I | J | K | L | M | N |
|---|---|---|---|---|---|---|---|---|---|---|---|---|---|---|
| A | - | 5.378 | 5.396 | 4.994 | 5.841 | NA | 5.591 | 5.484 | 5.516 | 5.405 | 4.823 | 5.537 | 5.044 | 5.947 |
| B | - | - | 3.914 | 5.095 | 4.534 | NA | 1.237 | 2.839 | 2.307 | 3.58 | 3.167 | 1.245 | 4.43 | 4.796 |
| C | - | - | - | 5.314 | 3.707 | NA | 3.862 | 4.282 | 3.233 | 4.667 | 4.852 | 3.8 | 5.086 | 4.192 |
| D | - | - | - | - | 5.378 | NA | 4.702 | 4.625 | 4.74 | 4.007 | 2.724 | 4.66 | 2.826 | 5.481 |
| E | - | - | - | - | - | NA | 4.708 | 4.412 | 4.651 | 5.44 | 5.281 | 4.821 | 5.876 | 2.39 |
| F | - | - | - | - | - | - | NA | NA | NA | NA | NA | NA | NA | NA |
| G | - | - | - | - | - | - | - | 2.549 | 2.946 | 2.873 | 3.205 | 2.312 | 4.986 | 4.805 |
| H | - | - | - | - | - | - | - | - | 2.524 | 4.108 | 1.771 | -0.952 | 4.541 | 4.857 |
| I | - | - | - | - | - | - | - | - | - | 3.136 | 3.901 | -2.256 | 4.645 | 4.876 |
| J | - | - | - | - | - | - | - | - | - | - | 2.132 | 3.69 | 3.069 | 5.117 |
| K | - | - | - | - | - | - | - | - | - | - | - | 2.937 | 3.076 | 5.078 |
| L | - | - | - | - | - | - | - | - | - | - | - | - | 4.659 | 4.96 |
| M | - | - | - | - | - | - | - | - | - | - | - | - | - | 5.375 |
| N | - | - | - | - | - | - | - | - | - | - | - | - | - | - |

| Max | A | B | C | D | E | F | G | H | I | J | K | L | M | N |
|---|---|---|---|---|---|---|---|---|---|---|---|---|---|---|
| A | - | 4.315 | 3.12 | 4.303 | 5.179 | NA | 5.88 | 1.34 | 5.129 | 5.16 | 5.625 | 5.857 | 1.392 | 5.742 |
| B | - | - | 3.213 | 0.746 | 5.573 | NA | 6.05 | 4.304 | 5.536 | 3.811 | 4.625 | 6.035 | 4.321 | 5.952 |
| C | - | - | - | 3.539 | 5.371 | NA | 5.969 | 3.372 | 5.332 | 4.721 | 5.215 | 5.951 | 3.373 | 5.849 |
| D | - | - | - | - | 5.529 | NA | 6.045 | 4.314 | 5.52 | 3.841 | 4.854 | 6.032 | 4.266 | 5.944 |
| E | - | - | - | - | - | NA | 5.2 | 5.631 | 0.963 | 6.785 | 7.291 | 5.177 | 5.665 | 4.825 |
| F | - | - | - | - | - | - | NA | NA | NA | NA | NA | NA | NA | NA |
| G | - | - | - | - | - | - | - | 7.116 | 5.873 | 7.874 | 8.182 | 1.113 | 7.093 | 3.697 |
| H | - | - | - | - | - | - | - | - | 5.062 | 5.521 | 5.883 | 5.842 | -1.861 | 5.718 |
| I | - | - | - | - | - | - | - | - | - | 6.81 | 7.095 | 5.157 | 5.63 | 4.835 |
| J | - | - | - | - | - | - | - | - | - | - | 2.552 | 6.125 | 4.894 | 6.061 |
| K | - | - | - | - | - | - | - | - | - | - | - | 6.207 | 5.229 | 6.139 |
| L | - | - | - | - | - | - | - | - | - | - | - | - | 7.044 | 3.406 |
| M | - | - | - | - | - | - | - | - | - | - | - | - | - | 5.708 |
| N | - | - | - | - | - | - | - | - | - | - | - | - | - | - |
