## Supplementary Data S2 for "Interlaboratory comparison of standardised metabolomics and lipidomics analyses in human and rodent blood using the MxP® Quant 500 kit": TG(17.2_36.2).pdf

**Amount of relative differences between laboratories**

| Min | A | B | C | D | E | F | G | H | I | J | K | L | M | N |
|---|---|---|---|---|---|---|---|---|---|---|---|---|---|---|
| A | - | 5.084 | 4.433 | 4.255 | 5.288 | 5.696 | 3.69 | 5.846 | 4.25 | 7.953 | 5.597 | 4.351 | 4.453 | 5.597 |
| B | - | - | 2.916 | 2.98 | 3.838 | 3.942 | 4.085 | 4.24 | 3.708 | 6.7 | 3.406 | 1.2 | 4.99 | 4.122 |
| C | - | - | - | -2.822 | 4.34 | 4.541 | 3.11 | 4.527 | 2.254 | 6.533 | 3.96 | 2.418 | 4.617 | 3.526 |
| D | - | - | - | - | 4.457 | 4.83 | 2.963 | 4.941 | 1.387 | 7.332 | 4.442 | 2.155 | 3.966 | 1.061 |
| E | - | - | - | - | - | 2.32 | 5.194 | 2.345 | 4.982 | 6.947 | 1.838 | 4.313 | 5.752 | 5.227 |
| F | - | - | - | - | - | - | 4.848 | 1.827 | 4.802 | 5.245 | 3.048 | 3.367 | 5.901 | 5.762 |
| G | - | - | - | - | - | - | - | 5.401 | 2.486 | 7.585 | 5.038 | 3.377 | -2.237 | 4.533 |
| H | - | - | - | - | - | - | - | - | 5.1 | 6.474 | 3.054 | 4.391 | 5.925 | 5.553 |
| I | - | - | - | - | - | - | - | - | - | 7.497 | 4.809 | 2.592 | 3.263 | 3.356 |
| J | - | - | - | - | - | - | - | - | - | - | 3.064 | 3.437 | 6.019 | 5.887 |
| K | - | - | - | - | - | - | - | - | - | - | - | 1.222 | 5.568 | 5.393 |
| L | - | - | - | - | - | - | - | - | - | - | - | - | 2.482 | 4.386 |
| M | - | - | - | - | - | - | - | - | - | - | - | - | - | 3.913 |
| N | - | - | - | - | - | - | - | - | - | - | - | - | - | - |

| | |
|---|---|
| (red) | Larger than 10 |
| (orange) | Between 8 and 9,99 |
| (yellow) | Between 6 and 7,99 |
| (light green) | Between 4 and 5,99 |
| (green) | Between 2 and 3,99 |
| (dark green) | Between 0 and 1,99 |
| (grey) | NA (Calculation failed) |

| Median | A | B | C | D | E | F | G | H | I | J | K | L | M | N |
|---|---|---|---|---|---|---|---|---|---|---|---|---|---|---|
| A | - | -0.203 | 3.158 | 1.794 | 4.142 | 2.743 | 1.973 | 3.546 | 0.858 | -0.254 | 3.704 | 3.959 | 3.969 | 2.16 |
| B | - | - | 3.304 | 0.535 | 3.851 | 3.081 | 0.241 | 3.519 | 0.284 | 2.98 | 3.9 | 3.806 | 4.256 | 2.282 |
| C | - | - | - | 2.882 | 4.456 | 1.524 | 2.925 | 4.279 | 3.353 | 0.419 | 2.242 | 4.438 | 3.124 | 3.758 |
| D | - | - | - | - | 3.955 | 2.712 | -1.099 | 3.539 | 0.696 | 2.063 | 3.842 | 3.856 | 4.17 | 2.259 |
| E | - | - | - | - | - | 4.602 | 4.191 | 1.828 | 3.944 | 4.593 | 5.094 | -0.613 | 5.295 | 3.494 |
| F | - | - | - | - | - | - | 1.334 | 4.042 | 2.83 | 1.199 | 2.956 | 4.255 | 3.755 | 3.548 |
| G | - | - | - | - | - | - | - | 3.28 | -1.148 | 2.631 | 4.021 | 3.723 | 4.28 | 1.385 |
| H | - | - | - | - | - | - | - | - | 3.532 | 4.435 | 4.904 | 1.268 | 5.141 | 2.871 |
| I | - | - | - | - | - | - | - | - | - | 2.365 | 3.95 | 3.728 | 4.28 | 1.662 |
| J | - | - | - | - | - | - | - | - | - | - | 1.628 | 4.458 | 3.223 | 3.98 |
| K | - | - | - | - | - | - | - | - | - | - | - | 4.64 | 2.266 | 4.197 |
| L | - | - | - | - | - | - | - | - | - | - | - | - | 5.214 | 3.653 |
| M | - | - | - | - | - | - | - | - | - | - | - | - | - | 4.285 |
| N | - | - | - | - | - | - | - | - | - | - | - | - | - | - |

| Max | A | B | C | D | E | F | G | H | I | J | K | L | M | N |
|---|---|---|---|---|---|---|---|---|---|---|---|---|---|---|
| A | - | 5.272 | 5.859 | 4.621 | 4.228 | 6.77 | 3.172 | 5.06 | 4.049 | 7.173 | 6.956 | 3.231 | 5.63 | 4.095 |
| B | - | - | 3.769 | 3.341 | 3.872 | 5.649 | 4.43 | 2.055 | 4.04 | 6.178 | 5.933 | 5.115 | 2.945 | 3.981 |
| C | - | - | - | 4.384 | 4.635 | 5.037 | 4.953 | 3.986 | 4.712 | 5.681 | 5.372 | 5.406 | 2.472 | 4.691 |
| D | - | - | - | - | 2.186 | 6.083 | 3.629 | 2.837 | 2.618 | 6.637 | 6.314 | 4.766 | 4.332 | 2.611 |
| E | - | - | - | - | - | 6.239 | 3.03 | 3.588 | 0.929 | 6.771 | 6.483 | 4.569 | 4.703 | 0.665 |
| F | - | - | - | - | - | - | 5.574 | 5.175 | 5.474 | 4.19 | 2.774 | 5.818 | 4.817 | 5.462 |
| G | - | - | - | - | - | - | - | 4.475 | 2.789 | 6.913 | 6.715 | 4.068 | 5.214 | 2.9 |
| H | - | - | - | - | - | - | - | - | 3.663 | 6.415 | 6.11 | 5.004 | 3.621 | 3.602 |
| I | - | - | - | - | - | - | - | - | - | 6.811 | 6.556 | 4.486 | 4.822 | -1.696 |
| J | - | - | - | - | - | - | - | - | - | - | 2.688 | 5.991 | 5.347 | 5.716 |
| K | - | - | - | - | - | - | - | - | - | - | - | 5.896 | 5.071 | 5.584 |
| L | - | - | - | - | - | - | - | - | - | - | - | - | 6.034 | 4.858 |
| M | - | - | - | - | - | - | - | - | - | - | - | - | - | 4.437 |
| N | - | - | - | - | - | - | - | - | - | - | - | - | - | - |
