## Supplementary Data S2 for "Interlaboratory comparison of standardised metabolomics and lipidomics analyses in human and rodent blood using the MxP® Quant 500 kit": TG(17.2_36.4).pdf

**Amount of relative differences between laboratories**

| Min | A | B | C | D | E | F | G | H | I | J | K | L | M | N |
|---|---|---|---|---|---|---|---|---|---|---|---|---|---|---|
| A | - | 6.266 | 6.283 | 5.724 | NA | 5.938 | 5.199 | 5.675 | 5.766 | 6.465 | 4.245 | 5.289 | 5.602 | 6.401 |
| B | - | - | 4.047 | 3.058 | NA | 5.073 | 3.625 | 3.123 | 5.847 | 4.026 | 5.248 | 4.507 | 4.337 | 1.481 |
| C | - | - | - | -0.259 | NA | 5.37 | 1.158 | 1.597 | 5.247 | 5.171 | 4.48 | 3.779 | 3.016 | 3.167 |
| D | - | - | - | - | NA | 5.131 | 3.004 | 0.995 | 5.232 | 5.052 | 5.003 | 3.938 | 3.59 | 3.834 |
| E | - | - | - | - | - | NA | NA | NA | NA | NA | NA | NA | NA | NA |
| F | - | - | - | - | - | - | 6.102 | 6.014 | 7.108 | 4.583 | 6.744 | 6.403 | 6.371 | 5.847 |
| G | - | - | - | - | - | - | - | 3.814 | 2.593 | 6.073 | 4.849 | 0.635 | 2.864 | 5.802 |
| H | - | - | - | - | - | - | - | - | 5.103 | 5.28 | 5.046 | 3.784 | 3.226 | 4.177 |
| I | - | - | - | - | - | - | - | - | - | 6.092 | 4.601 | 0.745 | 3.19 | 5.599 |
| J | - | - | - | - | - | - | - | - | - | - | 5.736 | 5.255 | 5.305 | 4.793 |
| K | - | - | - | - | - | - | - | - | - | - | - | 3.728 | 4.097 | 5.227 |
| L | - | - | - | - | - | - | - | - | - | - | - | - | 2.985 | 5.863 |
| M | - | - | - | - | - | - | - | - | - | - | - | - | - | 5.022 |
| N | - | - | - | - | - | - | - | - | - | - | - | - | - | - |

| Larger than 10 |
|---|
| Between 8 and 9,99 |
| Between 6 and 7,99 |
| Between 4 and 5,99 |
| Between 2 and 3,99 |
| Between 0 and 1,99 |
| NA (Calculation failed) |

| Median | A | B | C | D | E | F | G | H | I | J | K | L | M | N |
|---|---|---|---|---|---|---|---|---|---|---|---|---|---|---|
| A | - | 3.874 | 3.727 | 4.292 | NA | 5.564 | 5.048 | 4.597 | 4.759 | 4.955 | 1.271 | 5.087 | 4.498 | 5.468 |
| B | - | - | 3.461 | 2.981 | NA | 5.068 | 4.081 | 2.646 | 3.29 | 2.213 | 4.79 | 4.483 | 3.173 | 5.173 |
| C | - | - | - | 3.986 | NA | 5.328 | 4.599 | 3.903 | 4.197 | 4.082 | 3.89 | 4.851 | 4.101 | 5.385 |
| D | - | - | - | - | NA | 4.897 | 3.606 | -2.259 | 2.046 | 1.113 | 4.998 | 4.042 | 0.818 | 4.931 |
| E | - | - | - | - | - | NA | NA | NA | NA | NA | NA | NA | NA | NA |
| F | - | - | - | - | - | - | 4.633 | 5.365 | 5.16 | 5.417 | 6.52 | 4.063 | 5.212 | 1.895 |
| G | - | - | - | - | - | - | - | 3.855 | 3.34 | 2.578 | 5.575 | 1.052 | 3.986 | 4.093 |
| H | - | - | - | - | - | - | - | - | 1.921 | 1.713 | 5.048 | 4.005 | 0.475 | 4.914 |
| I | - | - | - | - | - | - | - | - | - | 2.289 | 5.169 | 3.507 | 2.095 | 4.671 |
| J | - | - | - | - | - | - | - | - | - | - | 5.106 | 4.371 | 2.766 | 5.129 |
| K | - | - | - | - | - | - | - | - | - | - | - | 5.2 | 4.69 | 5.569 |
| L | - | - | - | - | - | - | - | - | - | - | - | - | 4.17 | 3.95 |
| M | - | - | - | - | - | - | - | - | - | - | - | - | - | 4.852 |
| N | - | - | - | - | - | - | - | - | - | - | - | - | - | - |

| Max | A | B | C | D | E | F | G | H | I | J | K | L | M | N |
|---|---|---|---|---|---|---|---|---|---|---|---|---|---|---|
| A | - | 5.339 | 5.598 | 3.058 | NA | 2.562 | 4.863 | 2.82 | 3.9 | 5.891 | 5.39 | 4.947 | 1.697 | 4.908 |
| B | - | - | 2.522 | 4.427 | NA | 5.054 | 5.607 | 4.565 | 5.311 | 3.78 | -5.878 | 5.641 | 4.945 | 5.639 |
| C | - | - | - | 4.83 | NA | 5.195 | 5.727 | 4.85 | 5.439 | 3.716 | 2.399 | 5.778 | 5.169 | 5.739 |
| D | - | - | - | - | NA | 3.838 | 5.093 | 1.47 | 4.502 | 5.546 | 4.986 | 5.165 | 3.592 | 5.18 |
| E | - | - | - | - | - | NA | NA | NA | NA | NA | NA | NA | NA | NA |
| F | - | - | - | - | - | - | 4.515 | 3.802 | 3.273 | 6.176 | 5.659 | 4.63 | 1.569 | 4.667 |
| G | - | - | - | - | - | - | - | 5.575 | 4.113 | 6.925 | 6.618 | 1.258 | 5.048 | 1.756 |
| H | - | - | - | - | - | - | - | - | 4.375 | 5.723 | 5.053 | 5.099 | 3.234 | 5.105 |
| I | - | - | - | - | - | - | - | - | - | 6.569 | 6.074 | 4.111 | 3.834 | 4.116 |
| J | - | - | - | - | - | - | - | - | - | - | 4.009 | 5.824 | 5.37 | 5.838 |
| K | - | - | - | - | - | - | - | - | - | - | - | 5.676 | 5.014 | 5.673 |
| L | - | - | - | - | - | - | - | - | - | - | - | - | 5.222 | -1.565 |
| M | - | - | - | - | - | - | - | - | - | - | - | - | - | 4.786 |
| N | - | - | - | - | - | - | - | - | - | - | - | - | - | - |
