## Supplementary Data S2 for "Interlaboratory comparison of standardised metabolomics and lipidomics analyses in human and rodent blood using the MxP® Quant 500 kit": TG(17.2_38.5).pdf

**Amount of relative differences between laboratories**

| Min | A | B | C | D | E | F | G | H | I | J | K | L | M | N |
|---|---|---|---|---|---|---|---|---|---|---|---|---|---|---|
| A | - | 6.009 | NA | 5.486 | 5.66 | NA | 4.79 | 5.165 | 5.604 | 7.845 | 6.831 | 4.824 | 5.187 | 5.879 |
| B | - | - | NA | 5.547 | 5.157 | NA | 6.319 | 5.864 | 2.396 | 5.196 | 2.437 | 6.064 | 5.137 | 3.525 |
| C | - | - | - | NA | NA | NA | NA | NA | NA | NA | NA | NA | NA | NA |
| D | - | - | - | - | 3.169 | NA | 4.681 | 2.519 | 4.499 | 7.064 | 5.881 | 4.223 | 1.151 | 4.515 |
| E | - | - | - | - | - | NA | 5.296 | 4.192 | 3.755 | 7.393 | 6.009 | 5.004 | 3.172 | 3.945 |
| F | - | - | - | - | - | - | NA | NA | NA | NA | NA | NA | NA | NA |
| G | - | - | - | - | - | - | - | 3.854 | 5.087 | 7.869 | 6.619 | 1.648 | 4.117 | 5.337 |
| H | - | - | - | - | - | - | - | - | 4.859 | 6.722 | 5.83 | 3.287 | 3.152 | 4.736 |
| I | - | - | - | - | - | - | - | - | - | 8.11 | 6.363 | 5.572 | 4.661 | 3.14 |
| J | - | - | - | - | - | - | - | - | - | - | 4.212 | 5.173 | 3.103 | 2.878 |
| K | - | - | - | - | - | - | - | - | - | - | - | 6.116 | 5.209 | 3.778 |
| L | - | - | - | - | - | - | - | - | - | - | - | - | 3.84 | 5.299 |
| M | - | - | - | - | - | - | - | - | - | - | - | - | - | 4.359 |
| N | - | - | - | - | - | - | - | - | - | - | - | - | - | - |

| Median | A | B | C | D | E | F | G | H | I | J | K | L | M | N |
|---|---|---|---|---|---|---|---|---|---|---|---|---|---|---|
| A | - | 4.108 | NA | 3.807 | 4.668 | NA | 3.806 | 1.159 | 5.346 | 2.046 | 4.75 | 4.269 | 3.815 | 5.291 |
| B | - | - | NA | 3.993 | -0.347 | NA | 4.424 | 5 | 4.476 | 4.274 | 3.302 | 3.096 | 3.785 | 4.557 |
| C | - | - | - | NA | NA | NA | NA | NA | NA | NA | NA | NA | NA | NA |
| D | - | - | - | - | 3.816 | NA | 1.561 | 3.49 | 5.013 | 3.128 | 4.192 | 2.634 | 0.548 | 4.985 |
| E | - | - | - | - | - | NA | 4.268 | 4.92 | 4.407 | 4.693 | 2.252 | 3.184 | 3.867 | 4.334 |
| F | - | - | - | - | - | - | NA | NA | NA | NA | NA | NA | NA | NA |
| G | - | - | - | - | - | - | - | 3.463 | 5.019 | 2.407 | 4.181 | 2.86 | 0.915 | 4.955 |
| H | - | - | - | - | - | - | - | - | 5.317 | 0.713 | 4.81 | 4.04 | 3.548 | 5.282 |
| I | - | - | - | - | - | - | - | - | - | 5.824 | 4.463 | 5.224 | 5.499 | 0.864 |
| J | - | - | - | - | - | - | - | - | - | - | 4.866 | 3.937 | 3.565 | 5.337 |
| K | - | - | - | - | - | - | - | - | - | - | - | 3.943 | 4.37 | 4.139 |
| L | - | - | - | - | - | - | - | - | - | - | - | - | 2.54 | 4.661 |
| M | - | - | - | - | - | - | - | - | - | - | - | - | - | 4.938 |
| N | - | - | - | - | - | - | - | - | - | - | - | - | - | - |

| Max | A | B | C | D | E | F | G | H | I | J | K | L | M | N |
|---|---|---|---|---|---|---|---|---|---|---|---|---|---|---|
| A | - | 5.684 | NA | 4.517 | -0.144 | NA | -1.13 | 5.545 | 4.954 | 6.643 | 4.896 | 3.486 | 4.544 | 4.576 |
| B | - | - | NA | 4.392 | 5.183 | NA | 5.182 | 2.633 | 5.801 | 5.497 | 3.674 | 5.444 | 4.46 | 5.666 |
| C | - | - | - | NA | NA | NA | NA | NA | NA | NA | NA | NA | NA | NA |
| D | - | - | - | - | 4.251 | NA | 4.287 | 4.265 | 5.463 | 6.218 | 3.041 | 4.829 | -2.384 | 5.268 |
| E | - | - | - | - | - | NA | -0.887 | 5.577 | 4.936 | 6.849 | 5.093 | 3.517 | 4.58 | 4.578 |
| F | - | - | - | - | - | - | NA | NA | NA | NA | NA | NA | NA | NA |
| G | - | - | - | - | - | - | - | 5.588 | 4.932 | 6.852 | 5.094 | 3.452 | 4.585 | 4.567 |
| H | - | - | - | - | - | - | - | - | 5.729 | 5.532 | 3.208 | 5.317 | 4.036 | 5.598 |
| I | - | - | - | - | - | - | - | - | - | 7.715 | 6.494 | 4.821 | 6.296 | 3.334 |
| J | - | - | - | - | - | - | - | - | - | - | 5.173 | 5.936 | 5.524 | 6.047 |
| K | - | - | - | - | - | - | - | - | - | - | - | 5.036 | 2.733 | 5.391 |
| L | - | - | - | - | - | - | - | - | - | - | - | - | 5.304 | 3.846 |
| M | - | - | - | - | - | - | - | - | - | - | - | - | - | 5.268 |
| N | - | - | - | - | - | - | - | - | - | - | - | - | - | - |

| | |
|---|---|
| (red) | Larger than 10 |
| (orange) | Between 8 and 9,99 |
| (light orange) | Between 6 and 7,99 |
| (yellow-green) | Between 4 and 5,99 |
| (light green) | Between 2 and 3,99 |
| (dark green) | Between 0 and 1,99 |
| (light grey) | NA (Calculation failed) |
