## Supplementary Data S2 for "Interlaboratory comparison of standardised metabolomics and lipidomics analyses in human and rodent blood using the MxP® Quant 500 kit": TG(18.0_30.0).pdf

**Amount of relative differences between laboratories**

| Min | A | B | C | D | E | F | G | H | I | J | K | L | M | N |
|---|---|---|---|---|---|---|---|---|---|---|---|---|---|---|
| A | - | 6.52 | 6.097 | 8.08 | 11.741 | 6.056 | 7.913 | 8.222 | 5.728 | 7.773 | 6.234 | 7.959 | 7.552 | 6.961 |
| B | - | - | 3.727 | 7.741 | 11.675 | 4.07 | 3.686 | 2.396 | 5.937 | 6.613 | 1.317 | 8.158 | 6.846 | 1.718 |
| C | - | - | - | 7.876 | 11.536 | 4.514 | 0.685 | 4.349 | 5.529 | 7.006 | 3.084 | 8.03 | 6.871 | 1.966 |
| D | - | - | - | - | 8.705 | 5.184 | 8.955 | 9.177 | 7.752 | 7.356 | 6.797 | 10.259 | 5.904 | 7.789 |
| E | - | - | - | - | - | 5.921 | 9.277 | 9.506 | 8.124 | 7.851 | 7.278 | 10.517 | 6.854 | 8.112 |
| F | - | - | - | - | - | - | 6.556 | 4.546 | 7.173 | 7.707 | 5.706 | 9.176 | 6.365 | 6.109 |
| G | - | - | - | - | - | - | - | 3.836 | 4.757 | 6.463 | 4.246 | 7.684 | 6.373 | 4.222 |
| H | - | - | - | - | - | - | - | - | 4.787 | 6.363 | 4.193 | 7.732 | 6.179 | 4.482 |
| I | - | - | - | - | - | - | - | - | - | 7.496 | 5.323 | 5.331 | 7.322 | 6.201 |
| J | - | - | - | - | - | - | - | - | - | - | 1.89 | 8.923 | 4.481 | 5.476 |
| K | - | - | - | - | - | - | - | - | - | - | - | 8.36 | 6.558 | 3.561 |
| L | - | - | - | - | - | - | - | - | - | - | - | - | 6.98 | 6.179 |
| M | - | - | - | - | - | - | - | - | - | - | - | - | - | 7.834 |
| N | - | - | - | - | - | - | - | - | - | - | - | - | - | - |

| | |
|---|---|
| | Larger than 10 |
| | Between 8 and 9,99 |
| | Between 6 and 7,99 |
| | Between 4 and 5,99 |
| | Between 2 and 3,99 |
| | Between 0 and 1,99 |
| | NA (Calculation failed) |

| Median | A | B | C | D | E | F | G | H | I | J | K | L | M | N |
|---|---|---|---|---|---|---|---|---|---|---|---|---|---|---|
| A | - | 5.459 | 5.306 | 6.762 | 9.667 | 4.058 | 5.506 | 5.869 | 4.853 | 6.637 | 5.385 | 5.12 | 5.478 | 5.854 |
| B | - | - | 3.191 | 6.219 | 9.583 | -0.401 | 4.816 | 4.736 | 4.484 | 4.178 | 3.253 | 5.515 | 5.82 | 0.185 |
| C | - | - | - | 6.527 | 9.396 | 1.711 | 4.434 | 4.308 | 3.828 | 5.392 | 0.13 | 5.323 | 5.703 | 2.487 |
| D | - | - | - | - | 6.467 | 4.442 | 6.839 | 7.129 | 5.986 | 6.818 | 5.849 | 7.106 | 2.249 | 6.033 |
| E | - | - | - | - | - | 4.504 | 7.097 | 7.399 | 6.416 | 7.207 | 6.343 | 7.335 | 4.008 | 6.367 |
| F | - | - | - | - | - | - | 5.392 | 4.54 | 5.812 | 6.76 | 5.292 | 6.365 | 5.52 | 4.549 |
| G | - | - | - | - | - | - | - | 0.511 | 2.533 | 5.011 | 3.941 | 3.471 | 4.602 | 4.618 |
| H | - | - | - | - | - | - | - | - | 2.794 | 4.932 | 4.007 | 3.261 | 4.501 | 4.8 |
| I | - | - | - | - | - | - | - | - | - | 6.035 | 3.83 | 2.491 | 6.054 | 4.896 |
| J | - | - | - | - | - | - | - | - | - | - | 3.866 | 4.566 | 4.3 | 3.849 |
| K | - | - | - | - | - | - | - | - | - | - | - | 5.158 | 4.902 | 2.879 |
| L | - | - | - | - | - | - | - | - | - | - | - | - | 5.75 | 5.192 |
| M | - | - | - | - | - | - | - | - | - | - | - | - | - | 6.248 |
| N | - | - | - | - | - | - | - | - | - | - | - | - | - | - |

| Max | A | B | C | D | E | F | G | H | I | J | K | L | M | N |
|---|---|---|---|---|---|---|---|---|---|---|---|---|---|---|
| A | - | 2.877 | 3.273 | 8.134 | 10.049 | 4.553 | 5.495 | 5.685 | 2.776 | 7.709 | 5.124 | 2.724 | 7.266 | 0.865 |
| B | - | - | 1.09 | 8.026 | 10.007 | 3.895 | 5.14 | 5.366 | 3.726 | 7.559 | 4.68 | -0.603 | 7.101 | 2.348 |
| C | - | - | - | 7.972 | 9.928 | 3.641 | 5.016 | 5.255 | 3.912 | 7.511 | 4.524 | 1.474 | 7.041 | 2.833 |
| D | - | - | - | - | 7.205 | 6.097 | 5.962 | 5.925 | 6.246 | 4.269 | 6.023 | 6.18 | 5.137 | 6.202 |
| E | - | - | - | - | - | 6.487 | 6.442 | 6.434 | 6.521 | 6.086 | 6.461 | 6.501 | 6.288 | 6.506 |
| F | - | - | - | - | - | - | 4.16 | 4.537 | 4.615 | 7.251 | 3.225 | 3.755 | 6.732 | 4.117 |
| G | - | - | - | - | - | - | - | 2.156 | 5.164 | 6.818 | 2.828 | 4.73 | 6.224 | 4.898 |
| H | - | - | - | - | - | - | - | - | 5.268 | 6.696 | 3.463 | 4.89 | 6.075 | 5.034 |
| I | - | - | - | - | - | - | - | - | - | 7.86 | 5.485 | 3.857 | 7.435 | 3.227 |
| J | - | - | - | - | - | - | - | - | - | - | 5.818 | 6.034 | 4.206 | 6.067 |
| K | - | - | - | - | - | - | - | - | - | - | - | 4.387 | 6.465 | 4.613 |
| L | - | - | - | - | - | - | - | - | - | - | - | - | 7.115 | 2.156 |
| M | - | - | - | - | - | - | - | - | - | - | - | - | - | 5.905 |
| N | - | - | - | - | - | - | - | - | - | - | - | - | - | - |
