## Supplementary Data S2 for "Interlaboratory comparison of standardised metabolomics and lipidomics analyses in human and rodent blood using the MxP® Quant 500 kit": TG(18.0_30.1).pdf

**Amount of relative differences between laboratories**

| Min | A | B | C | D | E | F | G | H | I | J | K | L | M | N |
|---|---|---|---|---|---|---|---|---|---|---|---|---|---|---|
| A | - | 7.232 | 6.853 | 6.585 | 7.344 | 5.996 | 8.095 | 5.693 | 4.969 | 10.196 | 5.694 | 8.225 | 5.238 | 6.954 |
| B | - | - | 3.394 | 3.84 | 6.886 | 4.635 | 7.88 | 4.862 | 7.26 | 9.937 | 4.084 | 8.341 | 4.267 | 4.361 |
| C | - | - | - | 5.016 | 6.751 | 5.336 | 7.79 | 2.335 | 6.771 | 9.477 | 3.299 | 8.245 | 2.52 | 3.324 |
| D | - | - | - | - | 6.368 | 4.844 | 8.858 | 5.175 | 7.505 | 9.365 | 3.489 | 9.199 | 3.373 | 6.261 |
| E | - | - | - | - | - | 4.444 | 10.419 | 7.297 | 8.856 | 6.835 | 6.789 | 10.756 | 6.412 | 8.376 |
| F | - | - | - | - | - | - | 9.627 | 6.662 | 8.356 | 9.807 | 6.054 | 9.969 | 5.75 | 7.457 |
| G | - | - | - | - | - | - | - | 4.159 | 5.729 | 9.824 | 4.596 | 5.031 | 3.926 | 5.659 |
| H | - | - | - | - | - | - | - | - | 6.629 | 9.618 | 3.997 | 8.088 | 3.688 | 1.899 |
| I | - | - | - | - | - | - | - | - | - | 10.373 | 5.357 | 8.008 | 4.717 | 6.808 |
| J | - | - | - | - | - | - | - | - | - | - | 4.117 | 9.698 | 2.734 | 6.899 |
| K | - | - | - | - | - | - | - | - | - | - | - | 8.57 | 0.763 | 4.985 |
| L | - | - | - | - | - | - | - | - | - | - | - | - | 3.499 | 5.82 |
| M | - | - | - | - | - | - | - | - | - | - | - | - | - | 5.092 |
| N | - | - | - | - | - | - | - | - | - | - | - | - | - | - |

| | |
|---|---|
| | Larger than 10 |
| | Between 8 and 9,99 |
| | Between 6 and 7,99 |
| | Between 4 and 5,99 |
| | Between 2 and 3,99 |
| | Between 0 and 1,99 |
| | NA (Calculation failed) |

| Median | A | B | C | D | E | F | G | H | I | J | K | L | M | N |
|---|---|---|---|---|---|---|---|---|---|---|---|---|---|---|
| A | - | 4.948 | 4.111 | 4.202 | 5.945 | 5.396 | 2.075 | 4.126 | 3.041 | 3.961 | 3.396 | 3.513 | 4.528 | 4.789 |
| B | - | - | 4.738 | 4.519 | 5.137 | 3.679 | 5.212 | 5.068 | 5.015 | 5.119 | 4.762 | 4.988 | 3.81 | 3.093 |
| C | - | - | - | 2.712 | 5.604 | 5.019 | 2.856 | 2.281 | 1.458 | 2.996 | 1.874 | 1.838 | 3.544 | 3.864 |
| D | - | - | - | - | 5.428 | 4.778 | 3.726 | 4.162 | 3.186 | 3.34 | -2.441 | 3.402 | 2.51 | 2.937 |
| E | - | - | - | - | - | 3.165 | 6.293 | 6.509 | 6.047 | 6.528 | 5.668 | 6.347 | 5.358 | 5.657 |
| F | - | - | - | - | - | - | 5.9 | 6.099 | 5.783 | 3.274 | 5.393 | 5.854 | 4.885 | 4.97 |
| G | - | - | - | - | - | - | - | 2.236 | 1.259 | 2.407 | 2.548 | 2.369 | 3.863 | 4.464 |
| H | - | - | - | - | - | - | - | - | 0.972 | 3.782 | 3.164 | 0.835 | 4.12 | 4.219 |
| I | - | - | - | - | - | - | - | - | - | 5.068 | 1.777 | 2.634 | 3.91 | 4.519 |
| J | - | - | - | - | - | - | - | - | - | - | 4.47 | 3.771 | 4.521 | 4.817 |
| K | - | - | - | - | - | - | - | - | - | - | - | 2.668 | 3.307 | 3.486 |
| L | - | - | - | - | - | - | - | - | - | - | - | - | 3.235 | 4.16 |
| M | - | - | - | - | - | - | - | - | - | - | - | - | - | 2.991 |
| N | - | - | - | - | - | - | - | - | - | - | - | - | - | - |

| Max | A | B | C | D | E | F | G | H | I | J | K | L | M | N |
|---|---|---|---|---|---|---|---|---|---|---|---|---|---|---|
| A | - | 2.352 | 4.761 | 5.519 | 5.369 | 3.402 | 3.996 | 5.045 | 1.356 | 7.867 | 4.922 | 2.833 | -1.435 | -1.902 |
| B | - | - | 5.087 | 5.749 | 5.634 | 4.036 | 4.472 | 5.326 | 1.399 | 7.983 | 5.218 | 3.688 | 2.503 | 2.356 |
| C | - | - | - | 3.888 | 3.543 | 3.711 | 3.137 | 2.196 | 4.548 | 7.349 | 1.293 | 3.977 | 4.401 | 4.429 |
| D | - | - | - | - | 1.438 | 4.603 | 4.36 | 3.158 | 5.056 | 7.017 | 3.425 | 4.734 | 4.97 | 4.984 |
| E | - | - | - | - | - | 4.464 | 4.193 | 2.683 | 4.973 | 7.095 | 3.04 | 4.629 | 4.884 | 4.889 |
| F | - | - | - | - | - | - | 2.314 | 4.345 | 3.569 | 7.675 | 4.159 | 1.588 | 3.206 | 3.278 |
| G | - | - | - | - | - | - | - | 3.874 | 4.001 | 7.558 | 3.621 | 2.926 | 3.758 | 3.805 |
| H | - | - | - | - | - | - | - | - | 4.741 | 7.251 | 1.018 | 4.28 | 4.618 | 4.643 |
| I | - | - | - | - | - | - | - | - | - | 7.934 | 5.076 | 3.319 | 1.576 | 1.275 |
| J | - | - | - | - | - | - | - | - | - | - | 5.937 | 6.088 | 6.135 | 6.133 |
| K | - | - | - | - | - | - | - | - | - | - | - | 4.148 | 4.523 | 4.548 |
| L | - | - | - | - | - | - | - | - | - | - | - | - | 2.68 | 2.787 |
| M | - | - | - | - | - | - | - | - | - | - | - | - | - | -0.966 |
| N | - | - | - | - | - | - | - | - | - | - | - | - | - | - |
