## Supplementary Data S2 for "Interlaboratory comparison of standardised metabolomics and lipidomics analyses in human and rodent blood using the MxP® Quant 500 kit": TG(18.0_32.0).pdf

**Amount of relative differences between laboratories**

| Min | A | B | C | D | E | F | G | H | I | J | K | L | M | N |
|---|---|---|---|---|---|---|---|---|---|---|---|---|---|---|
| A | - | 6.616 | 6.079 | 7.564 | 11.985 | 5.878 | 8.221 | 6.812 | 6.467 | 9.227 | 6.984 | 6.238 | 9.744 | 6.594 |
| B | - | - | 5.274 | 7.255 | 11.734 | 3.712 | 6.715 | 0.007 | 4.696 | 8.34 | 4.682 | 5.993 | 9.406 | 2.732 |
| C | - | - | - | 7.441 | 11.986 | 4.991 | 7.62 | 5.467 | 4.156 | 8.887 | 5.167 | 3.502 | 9.6 | 5.376 |
| D | - | - | - | - | 9.449 | 5.926 | 9.442 | 8.952 | 9.048 | 9.533 | 9.421 | 9.409 | 9.029 | 8.599 |
| E | - | - | - | - | - | 3.937 | 8.072 | 7.394 | 7.503 | 8.195 | 7.996 | 7.958 | 7.713 | 6.886 |
| F | - | - | - | - | - | - | 8.515 | 3.385 | 5.07 | 10.004 | 5.69 | 6.679 | 11.085 | 3.304 |
| G | - | - | - | - | - | - | - | 4.617 | 5.35 | 6.571 | 5.928 | 6.292 | 8.04 | 2.831 |
| H | - | - | - | - | - | - | - | - | 4.519 | 8.173 | 4.65 | 5.888 | 9.202 | 3.092 |
| I | - | - | - | - | - | - | - | - | - | 8.542 | 2.959 | 4.995 | 9.379 | 4.749 |
| J | - | - | - | - | - | - | - | - | - | - | 6.312 | 6.465 | 6.923 | 3.938 |
| K | - | - | - | - | - | - | - | - | - | - | - | 4.928 | 9.063 | 4.681 |
| L | - | - | - | - | - | - | - | - | - | - | - | - | 9.677 | 5.581 |
| M | - | - | - | - | - | - | - | - | - | - | - | - | - | 6.415 |
| N | - | - | - | - | - | - | - | - | - | - | - | - | - | - |

| Median | A | B | C | D | E | F | G | H | I | J | K | L | M | N |
|---|---|---|---|---|---|---|---|---|---|---|---|---|---|---|
| A | - | 5.338 | 4.889 | 5.855 | 8.086 | 4.817 | 5.52 | 5.537 | 5.157 | 5.508 | 4.422 | 4.801 | 7.073 | 5.437 |
| B | - | - | 3.259 | 5.652 | 7.77 | 4.07 | 3.293 | 2.394 | 3.631 | 3.608 | 4.657 | 4.563 | 6.666 | 3.265 |
| C | - | - | - | 5.636 | 8.036 | 4.117 | 2.95 | 3.41 | 0.296 | 2.684 | 3.734 | 3.219 | 6.767 | 4.189 |
| D | - | - | - | - | 6.299 | 3.25 | 8.077 | 7.944 | 7.904 | 8.111 | 7.967 | 8.068 | 7.567 | 7.255 |
| E | - | - | - | - | - | 5.471 | 5.916 | 5.676 | 5.559 | 5.861 | 5.673 | 5.902 | 4.647 | 3.123 |
| F | - | - | - | - | - | - | 6.103 | 3.503 | 4.82 | 7.559 | 5.757 | 5.74 | 9.147 | -3.188 |
| G | - | - | - | - | - | - | - | 2.419 | 1.497 | 1.533 | 2.243 | 2.664 | 5.788 | 4.611 |
| H | - | - | - | - | - | - | - | - | 2.656 | 2.738 | 4.171 | 4.077 | 6.515 | 3.889 |
| I | - | - | - | - | - | - | - | - | - | 1.018 | 3.678 | 3.363 | 6.638 | 4.216 |
| J | - | - | - | - | - | - | - | - | - | - | 0.944 | 0.074 | 5.423 | 4.871 |
| K | - | - | - | - | - | - | - | - | - | - | - | 0.355 | 6.558 | 4.798 |
| L | - | - | - | - | - | - | - | - | - | - | - | - | 7.04 | 4.651 |
| M | - | - | - | - | - | - | - | - | - | - | - | - | - | 2.209 |
| N | - | - | - | - | - | - | - | - | - | - | - | - | - | - |

| Max | A | B | C | D | E | F | G | H | I | J | K | L | M | N |
|---|---|---|---|---|---|---|---|---|---|---|---|---|---|---|
| A | - | 4.385 | 1.391 | 7.639 | 10.299 | 1.519 | 7.005 | 5.066 | 4.245 | 8.019 | 5.674 | 3.391 | 7.933 | 2.579 |
| B | - | - | 3.921 | 7.232 | 10.067 | 4.311 | 6.478 | 3.381 | 0.705 | 7.624 | 4.639 | 3.111 | 7.649 | 3.628 |
| C | - | - | - | 7.593 | 10.315 | 2.435 | 6.941 | 4.914 | 3.996 | 7.97 | 5.562 | 2.943 | 7.912 | 1.724 |
| D | - | - | - | - | 8.62 | 6.081 | 4.579 | 5.802 | 5.919 | 4.383 | 5.642 | 5.987 | 3.927 | 6.018 |
| E | - | - | - | - | - | 6.55 | 6.388 | 6.506 | 6.521 | 6.154 | 6.488 | 6.532 | 6.166 | 6.534 |
| F | - | - | - | - | - | - | 7.094 | 5.239 | 4.505 | 8.094 | 5.801 | 3.8 | 8.102 | 3.225 |
| G | - | - | - | - | - | - | - | 5.379 | 5.584 | 5.829 | 5.086 | 5.692 | 5.749 | 5.745 |
| H | - | - | - | - | - | - | - | - | 3.447 | 7.403 | 3.714 | 4.109 | 7.403 | 4.366 |
| I | - | - | - | - | - | - | - | - | - | 7.659 | 4.754 | 2.833 | 7.624 | 3.448 |
| J | - | - | - | - | - | - | - | - | - | - | 5.858 | 6.115 | 0.847 | 6.14 |
| K | - | - | - | - | - | - | - | - | - | - | - | 4.747 | 7.096 | 4.899 |
| L | - | - | - | - | - | - | - | - | - | - | - | - | 7.836 | 2.025 |
| M | - | - | - | - | - | - | - | - | - | - | - | - | - | 6.13 |
| N | - | - | - | - | - | - | - | - | - | - | - | - | - | - |

| | |
|---|---|
| (red) | Larger than 10 |
| (orange) | Between 8 and 9,99 |
| (yellow) | Between 6 and 7,99 |
| (light green) | Between 4 and 5,99 |
| (green) | Between 2 and 3,99 |
| (dark green) | Between 0 and 1,99 |
| (grey) | NA (Calculation failed) |
