## Supplementary Data S2 for "Interlaboratory comparison of standardised metabolomics and lipidomics analyses in human and rodent blood using the MxP® Quant 500 kit": TG(18.0_32.1).pdf

**Amount of relative differences between laboratories**

| Min | A | B | C | D | E | F | G | H | I | J | K | L | M | N |
|---|---|---|---|---|---|---|---|---|---|---|---|---|---|---|
| A | - | 7.103 | 6.317 | 9.034 | 8.298 | 7.334 | 7.051 | 6.995 | 5.335 | 10.583 | 6.852 | 6.957 | 6.868 | 6.617 |
| B | - | - | 5.753 | 8.579 | 8.06 | 6.365 | 4.158 | 2.002 | 6.903 | 10.309 | 3.848 | 1.484 | 3.689 | 4.309 |
| C | - | - | - | 8.81 | 8.208 | 6.879 | 5.425 | 5.457 | 5.829 | 10.472 | 5.786 | 5.62 | 5.79 | 4.794 |
| D | - | - | - | - | 6.581 | 5.413 | 7.765 | 7.482 | 8.195 | 8.195 | 6.887 | 7.451 | 6.962 | 7.366 |
| E | - | - | - | - | - | 7.504 | 9.047 | 8.824 | 9.318 | 7.23 | 8.368 | 8.782 | 8.421 | 8.69 |
| F | - | - | - | - | - | - | 7.509 | 7.206 | 8.211 | 9.91 | 6.59 | 7.153 | 6.658 | 7.241 |
| G | - | - | - | - | - | - | - | 3.544 | 6.509 | 10.164 | 4.755 | 3.656 | 4.688 | 1.58 |
| H | - | - | - | - | - | - | - | - | 6.755 | 10.192 | 4.157 | 0.263 | 4.037 | 3.834 |
| I | - | - | - | - | - | - | - | - | - | 10.788 | 6.645 | 6.719 | 6.658 | 6.255 |
| J | - | - | - | - | - | - | - | - | - | - | 6.948 | 7.487 | 7.018 | 7.386 |
| K | - | - | - | - | - | - | - | - | - | - | - | 4.424 | 1.004 | 5.091 |
| L | - | - | - | - | - | - | - | - | - | - | - | - | 3.936 | 4.017 |
| M | - | - | - | - | - | - | - | - | - | - | - | - | - | 5.052 |
| N | - | - | - | - | - | - | - | - | - | - | - | - | - | - |

| Median | A | B | C | D | E | F | G | H | I | J | K | L | M | N |
|---|---|---|---|---|---|---|---|---|---|---|---|---|---|---|
| A | - | 4.626 | 3.35 | 5.441 | 6.339 | 4.97 | 3.963 | 4.35 | 4.277 | 5.173 | 4.722 | 4.651 | 4.584 | 4.972 |
| B | - | - | 4.122 | 3.975 | 5.955 | 3.626 | 3.687 | 2.853 | 2.696 | 1.896 | 2.35 | 2.066 | 0.76 | 2.496 |
| C | - | - | - | 5.296 | 6.343 | 4.757 | 2.404 | 3.183 | 3.149 | 5.351 | 3.74 | 3.778 | 3.876 | 4.394 |
| D | - | - | - | - | 5.302 | 4.084 | 3.708 | 3.081 | 2.335 | 3.856 | 3.79 | 3.378 | 1.23 | 1.168 |
| E | - | - | - | - | - | 4.239 | 6.228 | 6.105 | 5.981 | 6.232 | 6.364 | 6.214 | 5.877 | 5.76 |
| F | - | - | - | - | - | - | 5.167 | 4.898 | 4.847 | 3.418 | 5.028 | 4.87 | 4.505 | 4.092 |
| G | - | - | - | - | - | - | - | 2.328 | 2.483 | 3.835 | 2.818 | 2.965 | 3.307 | 4.042 |
| H | - | - | - | - | - | - | - | - | -0.482 | 3.268 | 0.879 | 1.46 | 2.37 | 3.579 |
| I | - | - | - | - | - | - | - | - | - | 4.367 | 2.873 | 2.628 | 2.463 | 3.835 |
| J | - | - | - | - | - | - | - | - | - | - | 0.303 | 2.484 | 3.751 | 4.022 |
| K | - | - | - | - | - | - | - | - | - | - | - | 0.413 | 2.144 | 3.44 |
| L | - | - | - | - | - | - | - | - | - | - | - | - | 1.282 | 3.238 |
| M | - | - | - | - | - | - | - | - | - | - | - | - | - | 2.781 |
| N | - | - | - | - | - | - | - | - | - | - | - | - | - | - |

| Max | A | B | C | D | E | F | G | H | I | J | K | L | M | N |
|---|---|---|---|---|---|---|---|---|---|---|---|---|---|---|
| A | - | 0.266 | 3.133 | 6.297 | 6.24 | 3.674 | 3.701 | 3.236 | 3.925 | 7.561 | 3.969 | 2.55 | 2.998 | 3.584 |
| B | - | - | 2.898 | 6.274 | 6.229 | 3.516 | 3.543 | 3.021 | 4.018 | 7.541 | 3.837 | 2.208 | 2.747 | 3.705 |
| C | - | - | - | 6.016 | 5.963 | 1.898 | 1.969 | -0.711 | 4.461 | 7.378 | 2.669 | 1.4 | -0.535 | 4.254 |
| D | - | - | - | - | 0.811 | 5.216 | 5.205 | 5.282 | 5.718 | 5.923 | 5.147 | 5.359 | 5.314 | 5.668 |
| E | - | - | - | - | - | 5.182 | 5.168 | 5.251 | 5.703 | 5.998 | 5.112 | 5.329 | 5.285 | 5.647 |
| F | - | - | - | - | - | - | -2.283 | 1.571 | 4.633 | 7.293 | 1.36 | 2.613 | 2.083 | 4.457 |
| G | - | - | - | - | - | - | - | 1.665 | 4.642 | 7.284 | 1.236 | 2.656 | 2.148 | 4.467 |
| H | - | - | - | - | - | - | - | - | 4.492 | 7.362 | 2.513 | 1.695 | 0.378 | 4.291 |
| I | - | - | - | - | - | - | - | - | - | 7.918 | 5.186 | 4.634 | 4.773 | 1.92 |
| J | - | - | - | - | - | - | - | - | - | - | 5.908 | 5.987 | 5.971 | 6.121 |
| K | - | - | - | - | - | - | - | - | - | - | - | 3.081 | 2.729 | 4.58 |
| L | - | - | - | - | - | - | - | - | - | - | - | - | 1.001 | 4.077 |
| M | - | - | - | - | - | - | - | - | - | - | - | - | - | 4.211 |
| N | - | - | - | - | - | - | - | - | - | - | - | - | - | - |

| | |
|---|---|
| | Larger than 10 |
| | Between 8 and 9,99 |
| | Between 6 and 7,99 |
| | Between 4 and 5,99 |
| | Between 2 and 3,99 |
| | Between 0 and 1,99 |
| | NA (Calculation failed) |
