## Supplementary Data S2 for "Interlaboratory comparison of standardised metabolomics and lipidomics analyses in human and rodent blood using the MxP® Quant 500 kit": TG(18.0_38.6).pdf

**Amount of relative differences between laboratories**

| Min | A | B | C | D | E | F | G | H | I | J | K | L | M | N |
|---|---|---|---|---|---|---|---|---|---|---|---|---|---|---|
| A | - | 3.061 | 5.2 | 4.286 | NA | 5.283 | 3.072 | 4.873 | 3.083 | 6.882 | 5.69 | 4.834 | 4.663 | 5.084 |
| B | - | - | 4.863 | 3.277 | NA | 5.058 | -3.492 | 4.457 | -0.854 | 7.011 | 5.509 | 4.403 | 4.055 | 4.745 |
| C | - | - | - | 4.206 | NA | 4.347 | 4.271 | 0.68 | 4.41 | 5.946 | 4.099 | 1.937 | 3.441 | 1.196 |
| D | - | - | - | - | NA | 4.956 | 1.661 | 4.012 | 2.176 | 6.518 | 4.995 | 4.117 | 2.738 | 4.152 |
| E | - | - | - | - | - | NA | NA | NA | NA | NA | NA | NA | NA | NA |
| F | - | - | - | - | - | - | 5.631 | 4.575 | 5.697 | 5.615 | 2.483 | 4.357 | 5.316 | 4.54 |
| G | - | - | - | - | - | - | - | 4.208 | 0.289 | 6.231 | 5.074 | 4.322 | 3.17 | 4.328 |
| H | - | - | - | - | - | - | - | - | 4.547 | 5.97 | 4.044 | 1.045 | 3.591 | 0.204 |
| I | - | - | - | - | - | - | - | - | - | 7.006 | 5.538 | 4.453 | 4.182 | 4.782 |
| J | - | - | - | - | - | - | - | - | - | - | 1.133 | 2.053 | 4.859 | 3.562 |
| K | - | - | - | - | - | - | - | - | - | - | - | 3.833 | 5.028 | 4.065 |
| L | - | - | - | - | - | - | - | - | - | - | - | - | 3.777 | -1.117 |
| M | - | - | - | - | - | - | - | - | - | - | - | - | - | 3.534 |
| N | - | - | - | - | - | - | - | - | - | - | - | - | - | - |

| Median | A | B | C | D | E | F | G | H | I | J | K | L | M | N |
|---|---|---|---|---|---|---|---|---|---|---|---|---|---|---|
| A | - | 2.535 | 1.628 | 2.362 | NA | 4.554 | 3.19 | 2.753 | 2.879 | 4.74 | 4.096 | 3.441 | -0.879 | 4.26 |
| B | - | - | 1.419 | 3.536 | NA | 4.226 | 3.985 | 0.192 | 0.788 | 5.096 | 3.613 | 2.448 | 2.744 | 3.836 |
| C | - | - | - | 3.249 | NA | 4.266 | 3.828 | 1.818 | 1.936 | 4.735 | 3.89 | 2.938 | 2.298 | 3.992 |
| D | - | - | - | - | NA | 4.728 | 2.049 | 3.556 | 3.598 | 4.107 | 4.456 | 3.96 | 2.087 | 4.536 |
| E | - | - | - | - | - | NA | NA | NA | NA | NA | NA | NA | NA | NA |
| F | - | - | - | - | - | - | 5.368 | 4.268 | 4.24 | 5.812 | 2.394 | 3.821 | 4.896 | 1.998 |
| G | - | - | - | - | - | - | - | 3.936 | 3.968 | 3.581 | 4.651 | 4.246 | 3.01 | 4.716 |
| H | - | - | - | - | - | - | - | - | -1.482 | 4.98 | 3.544 | 2.104 | 3.114 | 3.687 |
| I | - | - | - | - | - | - | - | - | - | 5.203 | 3.402 | 1.925 | 3.091 | 3.672 |
| J | - | - | - | - | - | - | - | - | - | - | 4.912 | 4.541 | 3.58 | 4.9 |
| K | - | - | - | - | - | - | - | - | - | - | - | 3.074 | 4.541 | 0.282 |
| L | - | - | - | - | - | - | - | - | - | - | - | - | 3.752 | 3.167 |
| M | - | - | - | - | - | - | - | - | - | - | - | - | - | 4.306 |
| N | - | - | - | - | - | - | - | - | - | - | - | - | - | - |

| Max | A | B | C | D | E | F | G | H | I | J | K | L | M | N |
|---|---|---|---|---|---|---|---|---|---|---|---|---|---|---|
| A | - | 1.696 | 4.323 | 4.571 | NA | 0.138 | 4.914 | 3.791 | 2.706 | 6.687 | 1.91 | 2.732 | 4.361 | 3.271 |
| B | - | - | 4.614 | 4.791 | NA | 1.286 | 5.117 | 4.158 | 1.775 | 6.91 | 2.981 | 3.336 | 4.629 | 2.634 |
| C | - | - | - | 1.459 | NA | 4.14 | 3.014 | 2.4 | 4.49 | 6.194 | 3.748 | 3.519 | -1.643 | 4.625 |
| D | - | - | - | - | NA | 4.317 | 2.407 | 2.957 | 4.613 | 6.222 | 3.994 | 3.786 | 1.248 | 4.738 |
| E | - | - | - | - | - | NA | NA | NA | NA | NA | NA | NA | NA | NA |
| F | - | - | - | - | - | - | 4.971 | 3.916 | 2.49 | 6.803 | 2.337 | 2.969 | 4.458 | 3.073 |
| G | - | - | - | - | - | - | - | 3.634 | 4.817 | 5.856 | 4.318 | 4.167 | 2.874 | 4.92 |
| H | - | - | - | - | - | - | - | - | 4.17 | 6.345 | 3.128 | 2.703 | 2.595 | 4.353 |
| I | - | - | - | - | - | - | - | - | - | 6.955 | 3.475 | 3.822 | 4.865 | 1.693 |
| J | - | - | - | - | - | - | - | - | - | - | 5.633 | 5.657 | 5.448 | 5.825 |
| K | - | - | - | - | - | - | - | - | - | - | - | 1.406 | 4.006 | 3.675 |
| L | - | - | - | - | - | - | - | - | - | - | - | - | 3.732 | 3.913 |
| M | - | - | - | - | - | - | - | - | - | - | - | - | - | 4.648 |
| N | - | - | - | - | - | - | - | - | - | - | - | - | - | - |

| | |
|---|---|
| (red) | Larger than 10 |
| (orange) | Between 8 and 9,99 |
| (light orange) | Between 6 and 7,99 |
| (light green) | Between 4 and 5,99 |
| (green) | Between 2 and 3,99 |
| (dark green) | Between 0 and 1,99 |
| (grey) | NA (Calculation failed) |
