## Supplementary Data S2 for "Interlaboratory comparison of standardised metabolomics and lipidomics analyses in human and rodent blood using the MxP® Quant 500 kit": TG(18.0_38.7).pdf

**Amount of relative differences between laboratories**

| Min | A | B | C | D | E | F | G | H | I | J | K | L | M | N |
|---|---|---|---|---|---|---|---|---|---|---|---|---|---|---|
| A | - | 3.854 | 5.146 | 5.053 | 6.019 | 5.904 | 5.342 | 5.763 | 5.526 | 6.777 | 6.131 | 5.138 | 3.907 | 5.229 |
| B | - | - | 3.411 | 3.784 | 5.485 | 5.17 | 3.497 | 5.118 | 4.927 | 6.336 | 5.211 | 3.286 | 2.44 | 4.665 |
| C | - | - | - | 2.832 | 4.128 | 4.631 | -1.714 | 4.721 | 4.632 | 5.881 | 4.166 | -0.118 | 3.594 | 4.441 |
| D | - | - | - | - | 5.234 | 4.262 | 2.405 | 4.368 | 4.27 | 5.868 | 3.752 | 2.453 | 4.357 | 3.943 |
| E | - | - | - | - | - | 4.912 | 4.362 | 4.198 | 4.002 | 5.643 | 3.778 | 4.054 | 4.082 | 4.251 |
| F | - | - | - | - | - | - | 5.068 | 2.133 | 2.528 | 1.27 | 4.432 | 5.04 | 5.224 | 2.676 |
| G | - | - | - | - | - | - | - | 4.732 | 4.685 | 6.111 | 4.179 | -3.415 | 2.976 | 4.568 |
| H | - | - | - | - | - | - | - | - | -4.823 | 4.725 | 4.176 | 5.173 | 5.597 | 1.39 |
| I | - | - | - | - | - | - | - | - | - | 6.471 | 4.464 | 4.534 | 5.684 | 3.311 |
| J | - | - | - | - | - | - | - | - | - | - | 4.846 | 5.383 | 5.356 | 2.261 |
| K | - | - | - | - | - | - | - | - | - | - | - | 4.232 | 4.863 | 3.511 |
| L | - | - | - | - | - | - | - | - | - | - | - | - | 2.814 | 4.638 |
| M | - | - | - | - | - | - | - | - | - | - | - | - | - | 4.941 |
| N | - | - | - | - | - | - | - | - | - | - | - | - | - | - |

| | |
|---|---|
| | Larger than 10 |
| | Between 8 and 9,99 |
| | Between 6 and 7,99 |
| | Between 4 and 5,99 |
| | Between 2 and 3,99 |
| | Between 0 and 1,99 |
| | NA (Calculation failed) |

| Median | A | B | C | D | E | F | G | H | I | J | K | L | M | N |
|---|---|---|---|---|---|---|---|---|---|---|---|---|---|---|
| A | - | 2.768 | 1.814 | 3.836 | -1.669 | 3.885 | 0.071 | 4.489 | 5.317 | 4.407 | 4.019 | 2.666 | 3.566 | 4.846 |
| B | - | - | 0.756 | 2.838 | 1.959 | 3.218 | 2.714 | 4.247 | 5.096 | 3.751 | 3.3 | 3.438 | 4.179 | 4.549 |
| C | - | - | - | 2.918 | 1.595 | 3.369 | 2.195 | 4.377 | 5.09 | 3.545 | 3.314 | 3.044 | 4.013 | 4.577 |
| D | - | - | - | - | 3.138 | 1.547 | 3.688 | 3.825 | 4.847 | 2.295 | 1.46 | 4.078 | 4.682 | 4.135 |
| E | - | - | - | - | - | 3.929 | 1.462 | 4.551 | 5.062 | 3.78 | 3.731 | 2.358 | 3.666 | 4.658 |
| F | - | - | - | - | - | - | 4.021 | 3.665 | 4.629 | 2.853 | 1.139 | 4.283 | 4.863 | 3.854 |
| G | - | - | - | - | - | - | - | 4.604 | 5.188 | 4.166 | 3.791 | 1.733 | 3.414 | 4.752 |
| H | - | - | - | - | - | - | - | - | 3.967 | 3.949 | 3.907 | 5.189 | 5.56 | 1.594 |
| I | - | - | - | - | - | - | - | - | - | 4.95 | 5.233 | 6.15 | 6.345 | 4.094 |
| J | - | - | - | - | - | - | - | - | - | - | 2.092 | 3.466 | 4.405 | 4.201 |
| K | - | - | - | - | - | - | - | - | - | - | - | 4.268 | 4.832 | 3.969 |
| L | - | - | - | - | - | - | - | - | - | - | - | - | 2.885 | 4.858 |
| M | - | - | - | - | - | - | - | - | - | - | - | - | - | 5.095 |
| N | - | - | - | - | - | - | - | - | - | - | - | - | - | - |

| Max | A | B | C | D | E | F | G | H | I | J | K | L | M | N |
|---|---|---|---|---|---|---|---|---|---|---|---|---|---|---|
| A | - | 3.905 | 5.035 | 4.463 | 5.603 | 5.549 | 5.38 | 3.752 | 4.937 | 6.404 | 4.911 | 5.705 | 6.01 | 2.631 |
| B | - | - | 3.974 | 2.48 | 5.321 | 4.763 | 4.583 | 0.305 | 5.314 | 6.068 | 3.757 | 5.068 | 5.425 | 4.202 |
| C | - | - | - | 3.069 | 4.187 | 3.232 | 3.129 | 3.855 | 5.579 | 5.488 | 1.617 | 4.029 | 4.591 | 4.879 |
| D | - | - | - | - | 5.434 | 4.27 | 4.343 | 2.782 | 5.421 | 5.885 | 2.691 | 4.881 | 5.162 | 4.537 |
| E | - | - | - | - | - | 2.341 | 3.869 | 4.866 | 5.887 | 4.889 | 3.69 | 2.912 | 2.129 | 5.366 |
| F | - | - | - | - | - | - | -0.62 | 4.462 | 5.724 | 4.229 | 3.854 | 2.22 | 3.689 | 5.14 |
| G | - | - | - | - | - | - | - | 4.446 | 5.711 | 5.436 | 3.347 | 2.783 | 4.011 | 5.142 |
| H | - | - | - | - | - | - | - | - | 5.281 | 6.035 | 3.628 | 5.208 | 5.485 | 4.135 |
| I | - | - | - | - | - | - | - | - | - | 7.444 | 6.474 | 6.951 | 7.128 | 5.15 |
| J | - | - | - | - | - | - | - | - | - | - | 5.029 | 4.51 | 4.133 | 5.74 |
| K | - | - | - | - | - | - | - | - | - | - | - | 4.307 | 4.769 | 4.736 |
| L | - | - | - | - | - | - | - | - | - | - | - | - | 3.015 | 5.307 |
| M | - | - | - | - | - | - | - | - | - | - | - | - | - | 5.429 |
| N | - | - | - | - | - | - | - | - | - | - | - | - | - | - |
