## Supplementary Data S2 for "Interlaboratory comparison of standardised metabolomics and lipidomics analyses in human and rodent blood using the MxP® Quant 500 kit": TG(18.1_26.0).pdf

**Amount of relative differences between laboratories**

| Min | A | B | C | D | E | F | G | H | I | J | K | L | M | N |
|---|---|---|---|---|---|---|---|---|---|---|---|---|---|---|
| A | - | 5.763 | 5.969 | 6.181 | 7.287 | 5.812 | 7.428 | 6.974 | 6.847 | 7.565 | 6.228 | 8.065 | 5.429 | 7.573 |
| B | - | - | 4.06 | 1.689 | 6.042 | 4.688 | 6.044 | 2.064 | 4.948 | 6.553 | 4.394 | 8.034 | 4.96 | 2.679 |
| C | - | - | - | 3.53 | 6.139 | 5.143 | 5.379 | 3.45 | 3.602 | 6.561 | 1.909 | 7.706 | 3.652 | 3.168 |
| D | - | - | - | - | 6.055 | 4.845 | 5.939 | 0.24 | 4.746 | 6.549 | 4.113 | 7.978 | 4.751 | 1.932 |
| E | - | - | - | - | - | 3.411 | 7.788 | 6.452 | 6.937 | 4.586 | 6.376 | 9.258 | 6.243 | 7.158 |
| F | - | - | - | - | - | - | 7.693 | 6.508 | 7 | 6.088 | 6.507 | 9.259 | 6.518 | 7.029 |
| G | - | - | - | - | - | - | - | 5.074 | 4.14 | 6.82 | 3.45 | 6.863 | 1.519 | 5.308 |
| H | - | - | - | - | - | - | - | - | 4.61 | 6.531 | 3.988 | 7.966 | 4.672 | 2.991 |
| I | - | - | - | - | - | - | - | - | - | 6.844 | -0.406 | 7.432 | 3.958 | 4.426 |
| J | - | - | - | - | - | - | - | - | - | - | 5.807 | 9.266 | 5.267 | 7.261 |
| K | - | - | - | - | - | - | - | - | - | - | - | 7.631 | 3.083 | 3.617 |
| L | - | - | - | - | - | - | - | - | - | - | - | - | -1.019 | 6.442 |
| M | - | - | - | - | - | - | - | - | - | - | - | - | - | 6.248 |
| N | - | - | - | - | - | - | - | - | - | - | - | - | - | - |

| Median | A | B | C | D | E | F | G | H | I | J | K | L | M | N |
|---|---|---|---|---|---|---|---|---|---|---|---|---|---|---|
| A | - | 4.437 | 2.985 | 4.079 | 5.248 | 5.221 | 3.859 | 4.495 | 4.524 | 2.819 | 2.074 | 4.441 | 3.626 | 5.073 |
| B | - | - | 4.031 | 1.824 | 3.624 | 4.329 | 4.221 | 2.276 | 2.607 | 5.326 | 4.376 | 3.374 | 3.222 | 2.376 |
| C | - | - | - | 3.459 | 4.615 | 4.939 | 0.949 | 3.307 | 3.136 | 4.371 | 1.934 | 2.374 | 2.582 | 4.208 |
| D | - | - | - | - | 3.931 | 4.534 | 3.87 | 0.364 | 1.302 | 5.142 | 4.057 | 2.733 | 2.485 | 3.065 |
| E | - | - | - | - | - | 3.887 | 5.003 | 3.937 | 4.123 | 5.726 | 4.886 | 4.586 | 4.048 | 2.984 |
| F | - | - | - | - | - | - | 5.704 | 5.158 | 5.226 | 6.036 | 5.711 | 5.495 | 5.24 | 4.631 |
| G | - | - | - | - | - | - | - | 3.509 | 3.56 | 4.31 | 1.852 | 2.848 | 2.573 | 4.285 |
| H | - | - | - | - | - | - | - | - | -0.136 | 5.088 | 3.921 | 2.446 | 2.103 | 3.096 |
| I | - | - | - | - | - | - | - | - | - | 5.04 | 3.959 | 2.433 | 2.272 | 3.103 |
| J | - | - | - | - | - | - | - | - | - | - | 4.097 | 3.88 | 4.693 | 4.683 |
| K | - | - | - | - | - | - | - | - | - | - | - | 3.113 | 3.24 | 4.426 |
| L | - | - | - | - | - | - | - | - | - | - | - | - | 2.175 | 3.754 |
| M | - | - | - | - | - | - | - | - | - | - | - | - | - | 4.12 |
| N | - | - | - | - | - | - | - | - | - | - | - | - | - | - |

| Max | A | B | C | D | E | F | G | H | I | J | K | L | M | N |
|---|---|---|---|---|---|---|---|---|---|---|---|---|---|---|
| A | - | 3.592 | 4.912 | 4.02 | 5.137 | 1.699 | 4.393 | 4.149 | 3.2 | 7.781 | 5.133 | -0.801 | 2.826 | 1.638 |
| B | - | - | 4.01 | 1.898 | 4.38 | 2.981 | 2.996 | 2.351 | 1.356 | 7.534 | 4.365 | 3.493 | 2.153 | 2.984 |
| C | - | - | - | 3.415 | 1.987 | 4.372 | 2.809 | 3.247 | 4.009 | 7.184 | 1.948 | 4.559 | 4.147 | 4.373 |
| D | - | - | - | - | 4.042 | 3.484 | 2.036 | 0.41 | 2.596 | 7.452 | 4.024 | 3.853 | 2.979 | 3.489 |
| E | - | - | - | - | - | 4.568 | 3.393 | 3.689 | 4.269 | 7.091 | -2.8 | 4.73 | 4.385 | 4.569 |
| F | - | - | - | - | - | - | 4.111 | 3.818 | 2.532 | 7.718 | 4.95 | 1.877 | 1.899 | -4.029 |
| G | - | - | - | - | - | - | - | 1.416 | 3.287 | 7.356 | 3.547 | 4.155 | 3.527 | 3.884 |
| H | - | - | - | - | - | - | - | - | 2.869 | 7.419 | 3.883 | 3.961 | 3.188 | 3.635 |
| I | - | - | - | - | - | - | - | - | - | 7.593 | 4.571 | 3.166 | 0.957 | 2.476 |
| J | - | - | - | - | - | - | - | - | - | - | 5.851 | 6.107 | 6.057 | 6.082 |
| K | - | - | - | - | - | - | - | - | - | - | - | 4.724 | 4.377 | 4.564 |
| L | - | - | - | - | - | - | - | - | - | - | - | - | 2.942 | 1.902 |
| M | - | - | - | - | - | - | - | - | - | - | - | - | - | 1.86 |
| N | - | - | - | - | - | - | - | - | - | - | - | - | - | - |

| | |
|---|---|
| (red) | Larger than 10 |
| (orange) | Between 8 and 9,99 |
| (yellow) | Between 6 and 7,99 |
| (light green) | Between 4 and 5,99 |
| (green) | Between 2 and 3,99 |
| (dark green) | Between 0 and 1,99 |
| (grey) | NA (Calculation failed) |
