## Supplementary Data S2 for "Interlaboratory comparison of standardised metabolomics and lipidomics analyses in human and rodent blood using the MxP® Quant 500 kit": TG(18.1_28.1).pdf

**Amount of relative differences between laboratories**

| Min | A | B | C | D | E | F | G | H | I | J | K | L | M | N |
|---|---|---|---|---|---|---|---|---|---|---|---|---|---|---|
| A | - | 6.346 | 7.518 | 5.685 | 6.631 | 4.492 | 7.487 | 6.486 | 6.678 | 8.065 | 5.953 | 7.038 | 6.251 | 6.928 |
| B | - | - | 2.793 | 3.397 | 6.105 | 1.666 | 4.137 | 4.768 | 2.494 | 7.578 | 4.415 | 6.609 | 5.356 | 2.914 |
| C | - | - | - | 4.095 | 6.088 | 1.833 | 3.156 | 3.303 | 4.124 | 7.424 | 2.87 | 6.219 | 4.387 | 2.18 |
| D | - | - | - | - | 5.805 | 2.952 | 6.746 | 6.22 | 5.175 | 7.148 | 5.637 | 7.602 | 6.458 | 6.11 |
| E | - | - | - | - | - | 6.05 | 9.005 | 8.279 | 7.821 | 5.337 | 7.73 | 9.378 | 8.361 | 8.353 |
| F | - | - | - | - | - | - | 4.132 | 5.267 | 3.228 | 7.772 | 5.134 | 6.79 | 5.874 | 4.891 |
| G | - | - | - | - | - | - | - | 2.766 | 4.352 | 7.501 | 2.27 | 6.053 | 4.201 | 3.206 |
| H | - | - | - | - | - | - | - | - | 4.681 | 7.514 | 1.662 | 5.788 | 3.451 | 3.948 |
| I | - | - | - | - | - | - | - | - | - | 7.406 | 4.135 | 6.865 | 5.399 | 4.22 |
| J | - | - | - | - | - | - | - | - | - | - | 7.123 | 9.025 | 7.875 | 7.975 |
| K | - | - | - | - | - | - | - | - | - | - | - | 5.941 | 4.078 | 2.776 |
| L | - | - | - | - | - | - | - | - | - | - | - | - | 3.463 | 5.561 |
| M | - | - | - | - | - | - | - | - | - | - | - | - | - | 5.068 |
| N | - | - | - | - | - | - | - | - | - | - | - | - | - | - |

| | |
|---|---|
| (red) | Larger than 10 |
| (orange) | Between 8 and 9,99 |
| (light orange) | Between 6 and 7,99 |
| (light green) | Between 4 and 5,99 |
| (green) | Between 2 and 3,99 |
| (dark green) | Between 0 and 1,99 |
| (grey) | NA (Calculation failed) |

| Median | A | B | C | D | E | F | G | H | I | J | K | L | M | N |
|---|---|---|---|---|---|---|---|---|---|---|---|---|---|---|
| A | - | 4.22 | 1.67 | 3.817 | 5.108 | 2.378 | 3.044 | 2.274 | 3.147 | 4.066 | 0.41 | 3.414 | 3.123 | 4.649 |
| B | - | - | 3.712 | 1.373 | 4.612 | 2.579 | 3.334 | 4.156 | 3.042 | 1.244 | 4.003 | 2.883 | 3.022 | 3.023 |
| C | - | - | - | 2.933 | 4.974 | 1.789 | 1.424 | 1.501 | 2.372 | 2.803 | 0.972 | 2.519 | 2.534 | 4.266 |
| D | - | - | - | - | 4.744 | 1.075 | 3.075 | 4.137 | 2.702 | 0.837 | 3.497 | 2.502 | 2.601 | 2.923 |
| E | - | - | - | - | - | 5.163 | 5.956 | 6.271 | 5.735 | 5.323 | 5.726 | 5.831 | 5.751 | 5.243 |
| F | - | - | - | - | - | - | 0.402 | 3.701 | 2.148 | 4.527 | 3.606 | -0.011 | 2.193 | 3.415 |
| G | - | - | - | - | - | - | - | 2.472 | 1.293 | 2.335 | 2.248 | 1.636 | 1.598 | 4.096 |
| H | - | - | - | - | - | - | - | - | 2.915 | 3.589 | 0.182 | 3.029 | 3.016 | 4.423 |
| I | - | - | - | - | - | - | - | - | - | 2.303 | 2.424 | -1.823 | -0.412 | 3.849 |
| J | - | - | - | - | - | - | - | - | - | - | -0.428 | 2.578 | 1.159 | 2.413 |
| K | - | - | - | - | - | - | - | - | - | - | - | 2.85 | 2.719 | 4.294 |
| L | - | - | - | - | - | - | - | - | - | - | - | - | -0.336 | 3.91 |
| M | - | - | - | - | - | - | - | - | - | - | - | - | - | 3.937 |
| N | - | - | - | - | - | - | - | - | - | - | - | - | - | - |

| Max | A | B | C | D | E | F | G | H | I | J | K | L | M | N |
|---|---|---|---|---|---|---|---|---|---|---|---|---|---|---|
| A | - | 3.42 | 4.715 | 4.961 | 5.297 | 4.477 | 4.352 | 4.731 | 4.666 | 7.52 | 4.709 | 2.454 | 3.284 | 1.263 |
| B | - | - | 3.819 | 4.208 | 4.71 | 3.382 | 3.142 | 3.842 | 3.735 | 7.291 | 3.803 | 2.227 | -0.217 | 3.561 |
| C | - | - | - | 1.928 | 3.369 | 1.717 | 2.21 | -2.153 | -0.538 | 6.959 | -2.888 | 4.04 | 3.712 | 4.504 |
| D | - | - | - | - | 2.684 | 2.772 | 3.024 | 1.802 | 2.119 | 6.862 | 1.924 | 4.288 | 4.029 | 4.676 |
| E | - | - | - | - | - | 3.643 | 3.763 | 3.205 | 3.331 | 6.703 | 3.255 | 4.61 | 4.419 | 4.905 |
| F | - | - | - | - | - | - | 0.479 | 1.833 | 1.392 | 7.054 | 1.67 | 3.772 | 3.354 | 4.331 |
| G | - | - | - | - | - | - | - | 2.347 | 2.045 | 7.081 | 2.239 | 3.632 | 3.152 | 4.244 |
| H | - | - | - | - | - | - | - | - | -0.138 | 6.956 | -1.565 | 4.057 | 3.733 | 4.515 |
| I | - | - | - | - | - | - | - | - | - | 6.982 | -0.783 | 3.991 | 3.645 | 4.47 |
| J | - | - | - | - | - | - | - | - | - | - | 5.799 | 5.974 | 5.94 | 6.036 |
| K | - | - | - | - | - | - | - | - | - | - | - | 4.034 | 3.701 | 4.499 |
| L | - | - | - | - | - | - | - | - | - | - | - | - | 2.01 | 2.902 |
| M | - | - | - | - | - | - | - | - | - | - | - | - | - | 3.465 |
| N | - | - | - | - | - | - | - | - | - | - | - | - | - | - |
