## Supplementary Data S2 for "Interlaboratory comparison of standardised metabolomics and lipidomics analyses in human and rodent blood using the MxP® Quant 500 kit": TG(18.1_30.0).pdf

**Amount of relative differences between laboratories**

| Min | A | B | C | D | E | F | G | H | I | J | K | L | M | N |
|---|---|---|---|---|---|---|---|---|---|---|---|---|---|---|
| A | - | 7.622 | 7.261 | 8.624 | 7.594 | 7.35 | 7.595 | 6.919 | 5.267 | 11.11 | 7.325 | 7.573 | 7.312 | 7.241 |
| B | - | - | 3.633 | 7.201 | 6.965 | 4.319 | 5.682 | 5.851 | 7.336 | 10.689 | 4.859 | 5.298 | 5.117 | 6.05 |
| C | - | - | - | 7.59 | 7.121 | 2.931 | 5.07 | 5.691 | 7.375 | 10.931 | 4.076 | 4.467 | 4.506 | 5.776 |
| D | - | - | - | - | 6.339 | 6.705 | 7.367 | 7.066 | 7.872 | 10.079 | 6.851 | 7.215 | 6.948 | 7.348 |
| E | - | - | - | - | - | 8.546 | 9.017 | 8.658 | 9.127 | 9.464 | 8.615 | 8.927 | 8.68 | 8.927 |
| F | - | - | - | - | - | - | 4.608 | 5.315 | 7.178 | 10.808 | 3.079 | 3.799 | 3.791 | 5.435 |
| G | - | - | - | - | - | - | - | 4.236 | 6.856 | 10.726 | 3.471 | 3.167 | 2.687 | 4.3 |
| H | - | - | - | - | - | - | - | - | 6.648 | 10.904 | 5.073 | 5.225 | 4.874 | 2.339 |
| I | - | - | - | - | - | - | - | - | - | 11.276 | 7.146 | 7.409 | 7.12 | 6.99 |
| J | - | - | - | - | - | - | - | - | - | - | 8.257 | 8.588 | 8.315 | 8.556 |
| K | - | - | - | - | - | - | - | - | - | - | - | 2.076 | 2.33 | 5.034 |
| L | - | - | - | - | - | - | - | - | - | - | - | - | 0.955 | 4.901 |
| M | - | - | - | - | - | - | - | - | - | - | - | - | - | 4.813 |
| N | - | - | - | - | - | - | - | - | - | - | - | - | - | - |

| Median | A | B | C | D | E | F | G | H | I | J | K | L | M | N |
|---|---|---|---|---|---|---|---|---|---|---|---|---|---|---|
| A | - | 4.24 | 4.124 | 4.269 | 5.17 | 4.321 | 3.821 | 3.994 | 3.915 | 3.424 | 4.061 | 4.235 | 4.204 | 4.521 |
| B | - | - | -0.527 | 1.145 | 4.546 | 0.092 | 2.904 | 1.639 | 2.013 | 3.081 | 2.118 | 0.827 | 0.648 | 2.474 |
| C | - | - | - | -3.265 | 4.642 | -1.402 | 3.003 | 2.199 | 2.731 | 2.042 | 2.337 | 1.147 | -0.698 | 2.191 |
| D | - | - | - | - | 4.735 | 2.185 | 2.093 | -0.115 | -2.655 | 2.386 | 0.321 | 0.02 | 2.6 | 3.211 |
| E | - | - | - | - | - | 5.203 | 5.648 | 5.373 | 5.446 | 5.276 | 5.524 | 5.436 | 5.021 | 4.878 |
| F | - | - | - | - | - | - | 3.088 | 2.112 | 2.404 | 3.24 | 2.409 | 1.443 | -1.986 | 2.158 |
| G | - | - | - | - | - | - | - | 1.9 | 1.488 | 0.97 | 1.552 | 2.421 | 2.976 | 3.563 |
| H | - | - | - | - | - | - | - | - | -0.055 | 2.341 | -0.862 | 0.975 | 2.033 | 3.071 |
| I | - | - | - | - | - | - | - | - | - | 1.737 | 0.693 | 2.041 | 2.067 | 3.221 |
| J | - | - | - | - | - | - | - | - | - | - | 0.246 | 1.689 | 3.559 | 3.749 |
| K | - | - | - | - | - | - | - | - | - | - | - | 1.314 | 2.387 | 3.209 |
| L | - | - | - | - | - | - | - | - | - | - | - | - | 1.442 | 2.781 |
| M | - | - | - | - | - | - | - | - | - | - | - | - | - | 2.108 |
| N | - | - | - | - | - | - | - | - | - | - | - | - | - | - |

| Max | A | B | C | D | E | F | G | H | I | J | K | L | M | N |
|---|---|---|---|---|---|---|---|---|---|---|---|---|---|---|
| A | - | 3.002 | 2.324 | 5.02 | 4.647 | 1.683 | 3.293 | 0.656 | 3.556 | 7.075 | 3.022 | 2.145 | -0.089 | 2.885 |
| B | - | - | 1.514 | 4.51 | 4.01 | 2.16 | 0.702 | 2.582 | 4.196 | 6.886 | -3.625 | 1.73 | 2.716 | 3.835 |
| C | - | - | - | 4.725 | 4.28 | 0.744 | 2.204 | 1.701 | 3.998 | 6.963 | 1.59 | -1.055 | 1.939 | 3.557 |
| D | - | - | - | - | 2.462 | 4.472 | 4.106 | 4.549 | 5.063 | 6.269 | 4.207 | 4.412 | 4.58 | 4.915 |
| E | - | - | - | - | - | 4.148 | 3.625 | 4.246 | 4.889 | 6.46 | 3.78 | 4.072 | 4.293 | 4.706 |
| F | - | - | - | - | - | - | 2.674 | 0.669 | 3.859 | 7.008 | 2.253 | 0.265 | 1.134 | 3.359 |
| G | - | - | - | - | - | - | - | 2.906 | 4.297 | 6.834 | 0.607 | 2.283 | 3.011 | 3.968 |
| H | - | - | - | - | - | - | - | - | 3.716 | 7.046 | 2.691 | 1.504 | -0.705 | 3.14 |
| I | - | - | - | - | - | - | - | - | - | 7.389 | 4.495 | 4.198 | 3.848 | 2.316 |
| J | - | - | - | - | - | - | - | - | - | - | 5.76 | 5.802 | 5.841 | 5.925 |
| K | - | - | - | - | - | - | - | - | - | - | - | 1.763 | 2.732 | 3.842 |
| L | - | - | - | - | - | - | - | - | - | - | - | - | 1.748 | 3.502 |
| M | - | - | - | - | - | - | - | - | - | - | - | - | - | 3.044 |
| N | - | - | - | - | - | - | - | - | - | - | - | - | - | - |

| Colour | Legend |
|---|---|
| (red) | Larger than 10 |
| (orange) | Between 8 and 9,99 |
| (light orange) | Between 6 and 7,99 |
| (light green) | Between 4 and 5,99 |
| (green) | Between 2 and 3,99 |
| (dark green) | Between 0 and 1,99 |
| (grey) | NA (Calculation failed) |
