## Supplementary Data S2 for "Interlaboratory comparison of standardised metabolomics and lipidomics analyses in human and rodent blood using the MxP® Quant 500 kit": TG(18.1_32.1).pdf

**Amount of relative differences between laboratories**

| Min | A | B | C | D | E | F | G | H | I | J | K | L | M | N |
|---|---|---|---|---|---|---|---|---|---|---|---|---|---|---|
| A | - | 6.903 | 6.861 | 7.037 | 6.989 | 7.245 | 6.488 | 6.186 | 4.568 | 10.308 | 6.522 | 6.566 | 6.577 | 6.086 |
| B | - | - | 3.384 | -0.579 | 5.422 | 5.374 | 4.943 | 4.774 | 6.421 | 9.99 | 4.746 | 4.449 | 4.32 | 5.43 |
| C | - | - | - | 3.441 | 5.613 | 5.607 | 4.292 | 4.108 | 6.184 | 9.915 | 3.999 | 3.503 | 3.258 | 4.99 |
| D | - | - | - | - | 5.386 | 5.265 | 4.789 | 4.527 | 6.256 | 9.818 | 4.569 | 4.282 | 4.132 | 5.247 |
| E | - | - | - | - | - | 3.686 | 6.598 | 6.401 | 7.259 | 9.955 | 6.518 | 6.425 | 6.376 | 6.737 |
| F | - | - | - | - | - | - | 6.253 | 6.036 | 7.002 | 9.828 | 6.163 | 6.051 | 5.997 | 6.415 |
| G | - | - | - | - | - | - | - | 0.399 | 5.814 | 10.019 | 1.747 | 2.975 | 3.234 | 3.836 |
| H | - | - | - | - | - | - | - | - | 5.87 | 10.175 | 2.783 | 3.517 | 3.705 | 3.691 |
| I | - | - | - | - | - | - | - | - | - | 10.509 | 6.27 | 6.335 | 6.35 | 5.661 |
| J | - | - | - | - | - | - | - | - | - | - | 7.819 | 7.782 | 7.753 | 7.763 |
| K | - | - | - | - | - | - | - | - | - | - | - | 2.18 | 2.591 | 4.134 |
| L | - | - | - | - | - | - | - | - | - | - | - | - | 0.734 | 4.514 |
| M | - | - | - | - | - | - | - | - | - | - | - | - | - | 4.616 |
| N | - | - | - | - | - | - | - | - | - | - | - | - | - | - |

| | |
|---|---|
| | Larger than 10 |
| | Between 8 and 9,99 |
| | Between 6 and 7,99 |
| | Between 4 and 5,99 |
| | Between 2 and 3,99 |
| | Between 0 and 1,99 |
| | NA (Calculation failed) |

| Median | A | B | C | D | E | F | G | H | I | J | K | L | M | N |
|---|---|---|---|---|---|---|---|---|---|---|---|---|---|---|
| A | - | 3.982 | 3.535 | 3.33 | 4.793 | 4.413 | 3.468 | 3.97 | 3.836 | 1.85 | 3.624 | 3.918 | 3.825 | 4.123 |
| B | - | - | 2.503 | 3.105 | 3.917 | 2.727 | 2.77 | -4.366 | 0.677 | 3.601 | 2.555 | 0.884 | -0.356 | 1.373 |
| C | - | - | - | 1.448 | 4.293 | 3.536 | 0.098 | 2.428 | 1.922 | 2.804 | -2.623 | 1.859 | 2.189 | 2.959 |
| D | - | - | - | - | 4.458 | 3.819 | 0.757 | 3.018 | 2.725 | 1.767 | 1.352 | 2.645 | 2.885 | 3.39 |
| E | - | - | - | - | - | 3.371 | 4.733 | 4.169 | 4.316 | 5.021 | 4.682 | 4.374 | 4.221 | 3.889 |
| F | - | - | - | - | - | - | 3.88 | 2.798 | 3.127 | 4.355 | 3.795 | 3.232 | 2.92 | 2.024 |
| G | - | - | - | - | - | - | - | 2.628 | 2.203 | 2.622 | -0.017 | 2.217 | 2.385 | 3.082 |
| H | - | - | - | - | - | - | - | - | 0.739 | 3.657 | 2.423 | 0.618 | 0.137 | 1.268 |
| I | - | - | - | - | - | - | - | - | - | 3.356 | 1.308 | -0.216 | -7.444 | 1.902 |
| J | - | - | - | - | - | - | - | - | - | - | 0.46 | 2.613 | 3.476 | 3.755 |
| K | - | - | - | - | - | - | - | - | - | - | - | 1.889 | 2.112 | 2.926 |
| L | - | - | - | - | - | - | - | - | - | - | - | - | -0.577 | 2.022 |
| M | - | - | - | - | - | - | - | - | - | - | - | - | - | 1.773 |
| N | - | - | - | - | - | - | - | - | - | - | - | - | - | - |

| Max | A | B | C | D | E | F | G | H | I | J | K | L | M | N |
|---|---|---|---|---|---|---|---|---|---|---|---|---|---|---|
| A | - | 2.17 | 3.234 | 3.95 | 2.201 | 2.264 | 2.627 | 1.176 | 3.619 | 6.924 | 2.512 | 0.966 | 0.564 | 2.914 |
| B | - | - | 2.236 | 3.4 | 3.11 | -1.499 | 0.662 | 2.696 | 4.008 | 6.831 | 0.172 | 1.287 | 1.542 | 3.528 |
| C | - | - | - | 2.482 | 3.674 | 2.06 | 1.578 | 3.418 | 4.314 | 6.688 | 1.776 | 2.772 | 2.864 | 3.956 |
| D | - | - | - | - | 4.118 | 3.207 | 3.021 | 3.945 | 4.59 | 6.54 | 3.089 | 3.556 | 3.607 | 4.32 |
| E | - | - | - | - | - | 3.298 | 3.486 | 1.25 | 3.025 | 7.065 | 3.421 | 2.758 | 2.646 | 1.662 |
| F | - | - | - | - | - | - | 0.308 | 2.767 | 4.036 | 6.812 | -0.358 | 1.484 | 1.707 | 3.565 |
| G | - | - | - | - | - | - | - | 2.989 | 4.122 | 6.773 | -1.148 | 1.985 | 2.144 | 3.692 |
| H | - | - | - | - | - | - | - | - | 3.362 | 7.003 | 3.025 | 2.11 | 1.934 | 2.435 |
| I | - | - | - | - | - | - | - | - | - | 7.28 | 4.36 | 4.023 | 3.973 | 2.441 |
| J | - | - | - | - | - | - | - | - | - | - | 5.724 | 5.771 | 5.776 | 5.875 |
| K | - | - | - | - | - | - | - | - | - | - | - | 1.818 | 1.999 | 3.647 |
| L | - | - | - | - | - | - | - | - | - | - | - | - | -1.061 | 3.222 |
| M | - | - | - | - | - | - | - | - | - | - | - | - | - | 3.152 |
| N | - | - | - | - | - | - | - | - | - | - | - | - | - | - |
