## Supplementary Data S2 for "Interlaboratory comparison of standardised metabolomics and lipidomics analyses in human and rodent blood using the MxP® Quant 500 kit": TG(18.1_33.2).pdf

**Amount of relative differences between laboratories**

| Min | A | B | C | D | E | F | G | H | I | J | K | L | M | N |
|---|---|---|---|---|---|---|---|---|---|---|---|---|---|---|
| A | - | 4.204 | 2.459 | 4.176 | 5.775 | 5.034 | 3.526 | 4.429 | 2.799 | 6.72 | 3.345 | 4.266 | 3.9 | 3.714 |
| B | - | - | 3.784 | 0.626 | 5.19 | 4.133 | 2.979 | 2.086 | 3.831 | 6.199 | 3.24 | 1.268 | 2.166 | 3.401 |
| C | - | - | - | 3.489 | 5.506 | 4.796 | 2.443 | 4.009 | -4.103 | 6.279 | 1.997 | 3.833 | 3.085 | 1.953 |
| D | - | - | - | - | 5.128 | 4.262 | 2.477 | 2.643 | 3.583 | 5.985 | 2.84 | 2.259 | 1.299 | 3.267 |
| E | - | - | - | - | - | 3.693 | 5.551 | 4.942 | 5.822 | 2.789 | 5.623 | 4.995 | 5.436 | 5.815 |
| F | - | - | - | - | - | - | 4.934 | 3.998 | 5.27 | 5.656 | 5.022 | 4.151 | 4.74 | 5.16 |
| G | - | - | - | - | - | - | - | 3.494 | 2.523 | 6.348 | 0.598 | 3.205 | 1.683 | 0.167 |
| H | - | - | - | - | - | - | - | - | 4.261 | 6.281 | 3.841 | 1.085 | 3.179 | 3.874 |
| I | - | - | - | - | - | - | - | - | - | 6.822 | 1.894 | 3.743 | 3.081 | 2.625 |
| J | - | - | - | - | - | - | - | - | - | - | 3.613 | -5.313 | 3.308 | 4.295 |
| K | - | - | - | - | - | - | - | - | - | - | - | 3.454 | 2.109 | 0.483 |
| L | - | - | - | - | - | - | - | - | - | - | - | - | 2.772 | 3.655 |
| M | - | - | - | - | - | - | - | - | - | - | - | - | - | 2.264 |
| N | - | - | - | - | - | - | - | - | - | - | - | - | - | - |

| | |
|---|---|
| | Larger than 10 |
| | Between 8 and 9,99 |
| | Between 6 and 7,99 |
| | Between 4 and 5,99 |
| | Between 2 and 3,99 |
| | Between 0 and 1,99 |
| | NA (Calculation failed) |

| Median | A | B | C | D | E | F | G | H | I | J | K | L | M | N |
|---|---|---|---|---|---|---|---|---|---|---|---|---|---|---|
| A | - | 3.033 | 0.454 | 0.881 | 4.313 | 4.403 | 2.221 | 3.825 | 2.565 | 2.368 | 0.903 | 3.52 | 2.872 | 1.275 |
| B | - | - | 3.543 | 2.713 | 3.812 | 3.79 | 2.072 | 2.633 | 0.647 | 2.962 | 3.481 | 2.013 | 0.095 | 2.456 |
| C | - | - | - | 2.086 | 4.493 | 4.504 | 2.704 | 3.987 | 3.107 | 1.449 | -2.197 | 3.78 | 3.212 | 2.358 |
| D | - | - | - | - | 4.3 | 4.245 | 0.98 | 3.556 | 2.301 | 1.516 | 2.183 | 3.325 | 2.327 | 0.273 |
| E | - | - | - | - | - | 1.405 | 4.596 | 3.564 | 4.244 | 5.371 | 5.014 | 3.715 | 4.325 | 4.56 |
| F | - | - | - | - | - | - | 4.389 | 3.15 | 4.111 | 4.806 | 4.856 | 3.46 | 4.103 | 4.451 |
| G | - | - | - | - | - | - | - | 3.32 | 1.081 | 1.621 | 2.755 | 2.933 | 1.533 | 0.669 |
| H | - | - | - | - | - | - | - | - | 3.095 | 3.832 | 4.213 | 1.311 | 2.966 | 3.668 |
| I | - | - | - | - | - | - | - | - | - | 0.641 | 3.055 | 2.557 | 0.466 | 1.91 |
| J | - | - | - | - | - | - | - | - | - | - | 3.244 | 4.51 | 4.128 | 4.05 |
| K | - | - | - | - | - | - | - | - | - | - | - | 3.784 | 3.18 | 2.401 |
| L | - | - | - | - | - | - | - | - | - | - | - | - | 2.385 | 3.322 |
| M | - | - | - | - | - | - | - | - | - | - | - | - | - | 2.213 |
| N | - | - | - | - | - | - | - | - | - | - | - | - | - | - |

| Max | A | B | C | D | E | F | G | H | I | J | K | L | M | N |
|---|---|---|---|---|---|---|---|---|---|---|---|---|---|---|
| A | - | 2.363 | 3.755 | 4.207 | 3.411 | 1.776 | 2.143 | 2.072 | 2.32 | 6.558 | 4.277 | 0.381 | 0.99 | 2.71 |
| B | - | - | 2.982 | 3.682 | 2.487 | 3.032 | -0.59 | 3.146 | 3.28 | 6.465 | 3.773 | 2.625 | 1.64 | 0.306 |
| C | - | - | - | 2.166 | 1.281 | 3.899 | 2.99 | 3.957 | 4.031 | 6.161 | 2.399 | 3.708 | 3.354 | 2.627 |
| D | - | - | - | - | 2.666 | 4.218 | 3.579 | 4.264 | 4.321 | 6.097 | -0.463 | 4.075 | 3.824 | 3.362 |
| E | - | - | - | - | - | 3.706 | 2.551 | 3.779 | 3.864 | 6.24 | 2.931 | 3.509 | 3.068 | 1.919 |
| F | - | - | - | - | - | - | 3.037 | -0.461 | 0.757 | 6.724 | 4.574 | 1.118 | 2.46 | 3.356 |
| G | - | - | - | - | - | - | - | 3.044 | 3.188 | 6.44 | 3.845 | 2.476 | 1.326 | 0.986 |
| H | - | - | - | - | - | - | - | - | -0.12 | 6.721 | 4.626 | 1.561 | 2.654 | 3.47 |
| I | - | - | - | - | - | - | - | - | - | 6.769 | 4.69 | 1.974 | 2.864 | 3.593 |
| J | - | - | - | - | - | - | - | - | - | - | 5.361 | 5.717 | 5.672 | 5.551 |
| K | - | - | - | - | - | - | - | - | - | - | - | 4.125 | 3.894 | 3.443 |
| L | - | - | - | - | - | - | - | - | - | - | - | - | 1.702 | 2.969 |
| M | - | - | - | - | - | - | - | - | - | - | - | - | - | 2.168 |
| N | - | - | - | - | - | - | - | - | - | - | - | - | - | - |
