## Supplementary Data S2 for "Interlaboratory comparison of standardised metabolomics and lipidomics analyses in human and rodent blood using the MxP® Quant 500 kit": TG(18.1_33.3).pdf

**Amount of relative differences between laboratories**

| Min | A | B | C | D | E | F | G | H | I | J | K | L | M | N |
|---|---|---|---|---|---|---|---|---|---|---|---|---|---|---|
| A | - | 4.098 | 3.072 | 3.842 | 5.376 | 4.519 | 3.97 | 1.765 | 2.376 | 4.785 | 3.386 | 3.808 | 1.584 | 3.416 |
| B | - | - | 1.529 | 0.1 | 4.902 | 3.735 | 1.439 | 3.205 | 2.399 | 4.3 | 2.403 | 1.499 | 3.095 | 1.775 |
| C | - | - | - | 1.936 | 4.975 | 3.972 | 2.281 | 2.695 | 0.888 | 4.521 | 1.948 | 2.499 | 2.497 | 0.099 |
| D | - | - | - | - | 4.917 | 3.642 | 0.392 | 3.6 | 2.983 | 5.366 | 2.671 | -1.173 | 3.591 | 2.314 |
| E | - | - | - | - | - | 4.129 | 5.148 | 5.605 | 5.328 | 3.35 | 5.642 | 4.983 | 5.535 | 5.45 |
| F | - | - | - | - | - | - | 3.771 | 4.689 | 4.387 | 0.293 | 4.595 | 3.626 | 4.669 | 4.361 |
| G | - | - | - | - | - | - | - | 3.764 | 3.32 | 4.737 | 2.858 | 0.008 | 3.792 | 2.606 |
| H | - | - | - | - | - | - | - | - | 2.54 | 3.618 | -0.388 | 3.557 | 0.773 | 1.697 |
| I | - | - | - | - | - | - | - | - | - | 5.894 | 3.788 | 3.334 | 1.58 | 3.324 |
| J | - | - | - | - | - | - | - | - | - | - | 3.937 | 1.186 | 3.709 | 3.429 |
| K | - | - | - | - | - | - | - | - | - | - | - | 3.535 | 0.529 | 1.9 |
| L | - | - | - | - | - | - | - | - | - | - | - | - | 3.655 | 2.435 |
| M | - | - | - | - | - | - | - | - | - | - | - | - | - | 2.892 |
| N | - | - | - | - | - | - | - | - | - | - | - | - | - | - |

| | |
|---|---|
| | Larger than 10 |
| | Between 8 and 9,99 |
| | Between 6 and 7,99 |
| | Between 4 and 5,99 |
| | Between 2 and 3,99 |
| | Between 0 and 1,99 |
| | NA (Calculation failed) |

| Median | A | B | C | D | E | F | G | H | I | J | K | L | M | N |
|---|---|---|---|---|---|---|---|---|---|---|---|---|---|---|
| A | - | 2.052 | -1.964 | 3.102 | 4.384 | 3.822 | 3.45 | 3.847 | 3.52 | 3.695 | 3.974 | 3.428 | 0.863 | 2.395 |
| B | - | - | 2.217 | 2.334 | 4.199 | 3.332 | 2.929 | 4.306 | 2.836 | 4.112 | 4.29 | 2.858 | 1.134 | 0.34 |
| C | - | - | - | 3.239 | 4.477 | 3.806 | 3.568 | 3.868 | 3.489 | 3.55 | 3.809 | 3.524 | 1.243 | 2.513 |
| D | - | - | - | - | 3.788 | 2.556 | 1.388 | 4.699 | 1.389 | 4.051 | 4.714 | 1.176 | 2.951 | 1.972 |
| E | - | - | - | - | - | 3.63 | 3.814 | 5.633 | 4.007 | 5.521 | 5.554 | 3.876 | 4.741 | 4.493 |
| F | - | - | - | - | - | - | 1.128 | 5.046 | 1.768 | 5.024 | 4.999 | 1.434 | 3.74 | 3.237 |
| G | - | - | - | - | - | - | - | 4.864 | -3.239 | 4.568 | 4.891 | -1.651 | 3.4 | 2.726 |
| H | - | - | - | - | - | - | - | - | 4.485 | 0.575 | 0.937 | 4.551 | 3.94 | 4.183 |
| I | - | - | - | - | - | - | - | - | - | 4.204 | 5.035 | 1.034 | 3.607 | 2.988 |
| J | - | - | - | - | - | - | - | - | - | - | 2.703 | 4.661 | 4.075 | 4.355 |
| K | - | - | - | - | - | - | - | - | - | - | - | 4.461 | 3.79 | 4.065 |
| L | - | - | - | - | - | - | - | - | - | - | - | - | 3.343 | 2.666 |
| M | - | - | - | - | - | - | - | - | - | - | - | - | - | 1.805 |
| N | - | - | - | - | - | - | - | - | - | - | - | - | - | - |

| Max | A | B | C | D | E | F | G | H | I | J | K | L | M | N |
|---|---|---|---|---|---|---|---|---|---|---|---|---|---|---|
| A | - | 2.097 | 3.496 | 2.009 | 2.683 | 0.833 | 2.913 | 5.371 | 4.271 | 6.193 | 5.162 | 2.927 | -0.925 | -0.164 |
| B | - | - | 2.786 | 3.033 | 3.366 | 2.461 | 3.477 | 5.163 | 4.501 | 6.175 | 4.932 | 3.529 | 2.254 | 2.336 |
| C | - | - | - | 3.796 | 3.985 | 3.535 | 4.076 | 4.751 | 4.783 | 5.982 | 4.448 | 4.102 | 3.48 | 3.483 |
| D | - | - | - | - | 1.219 | 1.77 | 1.829 | 5.554 | 3.977 | 6.679 | 5.375 | 2.008 | 1.581 | 1.622 |
| E | - | - | - | - | - | 2.377 | 0.334 | 5.673 | 3.805 | 6.629 | 5.483 | 0.819 | 2.381 | 2.436 |
| F | - | - | - | - | - | - | 2.593 | 5.461 | 4.206 | 6.305 | 5.248 | 2.735 | -3.231 | -1.16 |
| G | - | - | - | - | - | - | - | 5.719 | 3.667 | 6.623 | 5.551 | -1.52 | 2.793 | 2.816 |
| H | - | - | - | - | - | - | - | - | 5.429 | 4.455 | 1.99 | 5.12 | 4.922 | 4.916 |
| I | - | - | - | - | - | - | - | - | - | 7.067 | 6.1 | 3.789 | 4.477 | 4.486 |
| J | - | - | - | - | - | - | - | - | - | - | 4.542 | 5.702 | 5.635 | 5.513 |
| K | - | - | - | - | - | - | - | - | - | - | - | 5.007 | 4.778 | 4.767 |
| L | - | - | - | - | - | - | - | - | - | - | - | - | 2.899 | 2.829 |
| M | - | - | - | - | - | - | - | - | - | - | - | - | - | -1.628 |
| N | - | - | - | - | - | - | - | - | - | - | - | - | - | - |
