## Supplementary Data S2 for "Interlaboratory comparison of standardised metabolomics and lipidomics analyses in human and rodent blood using the MxP® Quant 500 kit": TG(18.1_34.1).pdf

**Amount of relative differences between laboratories**

| Min | A | B | C | D | E | F | G | H | I | J | K | L | M | N |
|---|---|---|---|---|---|---|---|---|---|---|---|---|---|---|
| A | - | 6.729 | 6.189 | 7.137 | 6.048 | 5.868 | 4.82 | 6.835 | 5.296 | 9.736 | 4.842 | 5.335 | 6.168 | 5.758 |
| B | - | - | 5.028 | 4.469 | 3.008 | 4.995 | 6.068 | 3.668 | 5.747 | 9.146 | 6.087 | 5.733 | 4.491 | 5.283 |
| C | - | - | - | 5.88 | 3.905 | 1.946 | 5.235 | 5.463 | 4.634 | 9.353 | 5.252 | 4.594 | 2.761 | 3.353 |
| D | - | - | - | - | 3.819 | 5.3 | 6.149 | 1.873 | 5.878 | 8.696 | 6.167 | 5.876 | 5.011 | 5.514 |
| E | - | - | - | - | - | 4.47 | 6.027 | 5.393 | 5.61 | 9.807 | 6.033 | 5.585 | 3.017 | 4.912 |
| F | - | - | - | - | - | - | 5.236 | 5.872 | 4.525 | 9.719 | 5.232 | 4.492 | 3.691 | 2.705 |
| G | - | - | - | - | - | - | - | 6.467 | 3.618 | 9.698 | -3.428 | 3.723 | 5.518 | 4.787 |
| H | - | - | - | - | - | - | - | - | 6.004 | 8.95 | 6.297 | 6.004 | 5.076 | 5.626 |
| I | - | - | - | - | - | - | - | - | - | 9.798 | 3.747 | -0.362 | 5.177 | 4.043 |
| J | - | - | - | - | - | - | - | - | - | - | 7.174 | 7.033 | 6.774 | 6.874 |
| K | - | - | - | - | - | - | - | - | - | - | - | 3.671 | 5.46 | 4.741 |
| L | - | - | - | - | - | - | - | - | - | - | - | - | 5.088 | 3.913 |
| M | - | - | - | - | - | - | - | - | - | - | - | - | - | 4.146 |
| N | - | - | - | - | - | - | - | - | - | - | - | - | - | - |

| Median | A | B | C | D | E | F | G | H | I | J | K | L | M | N |
|---|---|---|---|---|---|---|---|---|---|---|---|---|---|---|
| A | - | 3.392 | 2.958 | -1.188 | 4.747 | 4.077 | 2.775 | 3.495 | 3.807 | 3.135 | 1.866 | 3.496 | 3.572 | 3.971 |
| B | - | - | 2.132 | 3.415 | 4.224 | 2.785 | 2.506 | -2.295 | 1.99 | 3.118 | 3.088 | -2.377 | 0.469 | 2.859 |
| C | - | - | - | 2.701 | 4.44 | 3.399 | 0.285 | 2.093 | 2.935 | 1.269 | 2.046 | 2.074 | 2.39 | 3.382 |
| D | - | - | - | - | 4.731 | 3.976 | 2.042 | 3.254 | 3.723 | -0.086 | 0.817 | 3.261 | 3.428 | 4.045 |
| E | - | - | - | - | - | 3.824 | 4.859 | 4.463 | 4.162 | 4.781 | 5.036 | 4.473 | 4.39 | 3.861 |
| F | - | - | - | - | - | - | 3.691 | 2.763 | 1.707 | 3.367 | 4.025 | 2.808 | 2.573 | -1.181 |
| G | - | - | - | - | - | - | - | 2.345 | 3.019 | 1.107 | 1.687 | 2.323 | 2.537 | 3.362 |
| H | - | - | - | - | - | - | - | - | 1.957 | 3.474 | 3.121 | -4.698 | 0.316 | 2.88 |
| I | - | - | - | - | - | - | - | - | - | 2.585 | 3.63 | 1.782 | 1.336 | 1.282 |
| J | - | - | - | - | - | - | - | - | - | - | 3.219 | 4.053 | 4.198 | 4.66 |
| K | - | - | - | - | - | - | - | - | - | - | - | 2.994 | 3.129 | 3.719 |
| L | - | - | - | - | - | - | - | - | - | - | - | - | -0.134 | 2.528 |
| M | - | - | - | - | - | - | - | - | - | - | - | - | - | 2.463 |
| N | - | - | - | - | - | - | - | - | - | - | - | - | - | - |

| Max | A | B | C | D | E | F | G | H | I | J | K | L | M | N |
|---|---|---|---|---|---|---|---|---|---|---|---|---|---|---|
| A | - | 3.88 | 3.419 | 5.189 | 3.952 | 2.416 | -1.882 | 4.203 | 2.864 | 6.977 | 1.982 | 1.536 | 2.428 | 2.668 |
| B | - | - | 1.853 | 4.27 | 4.723 | 4.143 | 3.721 | 1.768 | 4.269 | 6.651 | 3.254 | 3.961 | 3.045 | 4.207 |
| C | - | - | - | 4.567 | 4.564 | 3.863 | 3.318 | 2.855 | 4.024 | 6.744 | 2.627 | 3.63 | 2.263 | 3.948 |
| D | - | - | - | - | 5.261 | 4.956 | 4.764 | 3.739 | 5.014 | 6.073 | 4.591 | 4.872 | 4.533 | 4.982 |
| E | - | - | - | - | - | 3.578 | 4.163 | 5.341 | 3.276 | 7.451 | 4.518 | 3.885 | 4.625 | 3.427 |
| F | - | - | - | - | - | - | 2.409 | 4.681 | 0.986 | 7.195 | 3.295 | 1.342 | 3.508 | 0.049 |
| G | - | - | - | - | - | - | - | 4.246 | 2.807 | 7.024 | 2.079 | 1.401 | 2.514 | 2.603 |
| H | - | - | - | - | - | - | - | - | 4.456 | 6.522 | 3.646 | 4.205 | 3.5 | 4.401 |
| I | - | - | - | - | - | - | - | - | - | 7.22 | 3.592 | 2.219 | 3.772 | -0.029 |
| J | - | - | - | - | - | - | - | - | - | - | 5.806 | 5.886 | 5.785 | 5.904 |
| K | - | - | - | - | - | - | - | - | - | - | - | 2.721 | 0.49 | 3.303 |
| L | - | - | - | - | - | - | - | - | - | - | - | - | 3.108 | 1.813 |
| M | - | - | - | - | - | - | - | - | - | - | - | - | - | 3.479 |
| N | - | - | - | - | - | - | - | - | - | - | - | - | - | - |

| | |
|---|---|
| | Larger than 10 |
| | Between 8 and 9,99 |
| | Between 6 and 7,99 |
| | Between 4 and 5,99 |
| | Between 2 and 3,99 |
| | Between 0 and 1,99 |
| | NA (Calculation failed) |
