## Supplementary Data S2 for "Interlaboratory comparison of standardised metabolomics and lipidomics analyses in human and rodent blood using the MxP® Quant 500 kit": TG(18.1_35.3).pdf

**Amount of relative differences between laboratories**

| Min | A | B | C | D | E | F | G | H | I | J | K | L | M | N |
|---|---|---|---|---|---|---|---|---|---|---|---|---|---|---|
| A | - | 3.599 | 1.102 | 3.702 | 5.537 | 4.471 | 3.7 | 3.057 | -0.096 | 7.671 | 3.245 | 4.827 | 3.889 | 4.147 |
| B | - | - | 3.586 | -0.296 | 5.221 | 3.649 | 0.893 | 0.289 | 4.287 | 7.3 | 2.043 | 4.127 | 2.074 | 2.247 |
| C | - | - | - | 3.51 | 5.459 | 4.442 | 3.607 | 3.263 | 2.813 | 7.193 | 2.757 | 4.713 | 3.85 | 2.576 |
| D | - | - | - | - | 5.186 | 3.574 | 0.149 | 0.353 | 4.452 | 7.121 | 2.329 | 4.048 | 1.86 | 3.19 |
| E | - | - | - | - | - | 5.096 | 5.672 | 5.706 | 6.533 | 6.05 | 5.961 | 4.878 | 5.55 | 6.333 |
| F | - | - | - | - | - | - | 3.631 | 3.811 | 5.376 | 6.509 | 4.38 | 2.116 | 3.227 | 4.919 |
| G | - | - | - | - | - | - | - | 1.683 | 4.417 | 7.18 | 2.542 | 4.021 | 1.301 | 2.616 |
| H | - | - | - | - | - | - | - | - | 4.132 | 7.299 | 1.287 | 4.22 | 2.436 | 1.396 |
| I | - | - | - | - | - | - | - | - | - | 7.594 | 3.489 | 4.887 | 4.053 | 4.071 |
| J | - | - | - | - | - | - | - | - | - | - | 3.639 | 3.253 | -3.406 | 5.21 |
| K | - | - | - | - | - | - | - | - | - | - | - | 4.373 | 3.058 | 1.433 |
| L | - | - | - | - | - | - | - | - | - | - | - | - | 4.024 | 4.75 |
| M | - | - | - | - | - | - | - | - | - | - | - | - | - | 2.378 |
| N | - | - | - | - | - | - | - | - | - | - | - | - | - | - |

| Median | A | B | C | D | E | F | G | H | I | J | K | L | M | N |
|---|---|---|---|---|---|---|---|---|---|---|---|---|---|---|
| A | - | -0.182 | 3.689 | 1.599 | 4.658 | 1.848 | 1.332 | -2.004 | 2.404 | 4.462 | 2.972 | 4.161 | 2.985 | 1.886 |
| B | - | - | 3.641 | 1.627 | 4.652 | 1.852 | 1.386 | -0.367 | 2.511 | 4.553 | 2.834 | 4.172 | 2.887 | 2.041 |
| C | - | - | - | 3.061 | 5.064 | 3.834 | 3.742 | 3.371 | 2.583 | 3.352 | 2.248 | 4.761 | 4.141 | 2.888 |
| D | - | - | - | - | 4.775 | 2.69 | 2.467 | 0.925 | 1.416 | 4.356 | 1.866 | 4.359 | 3.32 | 0.247 |
| E | - | - | - | - | - | 4.836 | 4.892 | 5.179 | 5.379 | 6.139 | 5.371 | 3.175 | 4.609 | 5.309 |
| F | - | - | - | - | - | - | -0.119 | 2.457 | 3.322 | 5.017 | 3.381 | 3.921 | 1.799 | 3.075 |
| G | - | - | - | - | - | - | - | 1.758 | 3.085 | 4.845 | 3.325 | 3.987 | 2.315 | 2.785 |
| H | - | - | - | - | - | - | - | - | 2.291 | 4.508 | 2.67 | 4.217 | 3.014 | 1.731 |
| I | - | - | - | - | - | - | - | - | - | 3.955 | 1.095 | 4.468 | 3.64 | 0.596 |
| J | - | - | - | - | - | - | - | - | - | - | 4.395 | 5.261 | 4.838 | 4.335 |
| K | - | - | - | - | - | - | - | - | - | - | - | 4.548 | 3.736 | 1.344 |
| L | - | - | - | - | - | - | - | - | - | - | - | - | 3.693 | 4.757 |
| M | - | - | - | - | - | - | - | - | - | - | - | - | - | 3.636 |
| N | - | - | - | - | - | - | - | - | - | - | - | - | - | - |

| Max | A | B | C | D | E | F | G | H | I | J | K | L | M | N |
|---|---|---|---|---|---|---|---|---|---|---|---|---|---|---|
| A | - | 3.68 | 4.786 | 4.41 | 0.748 | 4.519 | 3.143 | 3.681 | 3.058 | 7.261 | 4.578 | 3.014 | -2.145 | 4.073 |
| B | - | - | 3.712 | 2.911 | 3.274 | 3.193 | 1.78 | -4.251 | 1.977 | 7.009 | 3.31 | 4.212 | 3.48 | 1.899 |
| C | - | - | - | 2.289 | 4.326 | 1.842 | 3.876 | 3.529 | 3.913 | 6.663 | 1.529 | 4.804 | 4.423 | 3.05 |
| D | - | - | - | - | 3.991 | 0.419 | 3.367 | 2.816 | 3.423 | 6.825 | 1.246 | 4.6 | 4.128 | 1.881 |
| E | - | - | - | - | - | 4.358 | 2.726 | 3.376 | 2.609 | 7.229 | 4.462 | 3.328 | 1.046 | 3.862 |
| F | - | - | - | - | - | - | 3.524 | 3.061 | 3.581 | 6.703 | -0.355 | 4.659 | 4.22 | 2.311 |
| G | - | - | - | - | - | - | - | 1.872 | -1.05 | 7.021 | 3.798 | 3.955 | 2.99 | 2.88 |
| H | - | - | - | - | - | - | - | - | 1.971 | 6.996 | 3.321 | 4.215 | 3.489 | 1.904 |
| I | - | - | - | - | - | - | - | - | - | 7.097 | 3.859 | 3.92 | 2.925 | 2.976 |
| J | - | - | - | - | - | - | - | - | - | - | 5.702 | 6.029 | 5.96 | 5.788 |
| K | - | - | - | - | - | - | - | - | - | - | - | 4.695 | 4.265 | 2.524 |
| L | - | - | - | - | - | - | - | - | - | - | - | - | 3.17 | 4.761 |
| M | - | - | - | - | - | - | - | - | - | - | - | - | - | 4.076 |
| N | - | - | - | - | - | - | - | - | - | - | - | - | - | - |

| Colour | Legend |
|---|---|
| (red) | Larger than 10 |
| (orange) | Between 8 and 9,99 |
| (light orange) | Between 6 and 7,99 |
| (light yellow-green) | Between 4 and 5,99 |
| (light green) | Between 2 and 3,99 |
| (dark green) | Between 0 and 1,99 |
| (light grey) | NA (Calculation failed) |
