## Supplementary Data S2 for "Interlaboratory comparison of standardised metabolomics and lipidomics analyses in human and rodent blood using the MxP® Quant 500 kit": TG(18.1_36.0).pdf

**Amount of relative differences between laboratories**

| Min | A | B | C | D | E | F | G | H | I | J | K | L | M | N |
|---|---|---|---|---|---|---|---|---|---|---|---|---|---|---|
| A | - | 6.28 | 6.099 | 7.66 | 8.002 | 6.174 | 5.935 | 5.743 | 4.281 | 9.681 | 6.107 | 5.691 | 6.17 | 5.852 |
| B | - | - | 1.725 | 6.914 | 7.68 | 3.777 | 5.154 | 3.837 | 5.354 | 9.456 | 4.495 | 5.37 | 2.721 | 2.242 |
| C | - | - | - | 6.633 | 7.393 | 4.207 | 4.987 | 3.043 | 4.748 | 9.126 | 4.316 | 5.157 | 2.066 | 3.603 |
| D | - | - | - | - | 6.266 | 4.926 | 6.807 | 5.917 | 6.038 | 8.055 | 6.604 | 6.799 | 6.149 | 4.819 |
| E | - | - | - | - | - | 6.469 | 7.703 | 6.992 | 6.931 | 6.139 | 7.58 | 7.693 | 7.251 | 6.292 |
| F | - | - | - | - | - | - | 6.01 | 5.057 | 5.542 | 9.28 | 5.654 | 6.101 | 4.934 | 2.27 |
| G | - | - | - | - | - | - | - | 3.627 | 3.848 | 9.394 | 3.407 | 2.621 | 4.46 | 4.676 |
| H | - | - | - | - | - | - | - | - | 4.747 | 9.499 | 1.705 | 4.447 | 3.322 | 4.101 |
| I | - | - | - | - | - | - | - | - | - | 10.026 | 5.865 | 5.186 | 5.967 | 5.465 |
| J | - | - | - | - | - | - | - | - | - | - | 6.916 | 7.019 | 6.491 | 5.118 |
| K | - | - | - | - | - | - | - | - | - | - | - | 3.95 | 3.697 | 4.485 |
| L | - | - | - | - | - | - | - | - | - | - | - | - | 4.832 | 4.784 |
| M | - | - | - | - | - | - | - | - | - | - | - | - | - | 3.525 |
| N | - | - | - | - | - | - | - | - | - | - | - | - | - | - |

| | |
|---|---|
| | Larger than 10 |
| | Between 8 and 9,99 |
| | Between 6 and 7,99 |
| | Between 4 and 5,99 |
| | Between 2 and 3,99 |
| | Between 0 and 1,99 |
| | NA (Calculation failed) |

| Median | A | B | C | D | E | F | G | H | I | J | K | L | M | N |
|---|---|---|---|---|---|---|---|---|---|---|---|---|---|---|
| A | - | 4.432 | 3.462 | 4.048 | 4.837 | 3.891 | 3.307 | 4.203 | 3.82 | 1.691 | 2.499 | 3.825 | 3.86 | 4.455 |
| B | - | - | 3.882 | 3.037 | 3.31 | 2.772 | 4.074 | 2.3 | 3.316 | 4.695 | 4.465 | 3.522 | 2.993 | 0.977 |
| C | - | - | - | 2.44 | 4.356 | 2.995 | 0.942 | 3.105 | 2.319 | 3.492 | 2.694 | 1.443 | 2.646 | 3.961 |
| D | - | - | - | - | 4.066 | 2.549 | 2.86 | 2.196 | -0.127 | 4.352 | 3.571 | 1.434 | 1.883 | 3.766 |
| E | - | - | - | - | - | 3.398 | 4.843 | 3.814 | 4.16 | 5.539 | 5.112 | 4.586 | 3.828 | 0.237 |
| F | - | - | - | - | - | - | 3.602 | -0.064 | 1.89 | 2.217 | 4.084 | 2.89 | 1.276 | 3.043 |
| G | - | - | - | - | - | - | - | 3.326 | 2.426 | 3.032 | 2.184 | 2.208 | 2.819 | 3.936 |
| H | - | - | - | - | - | - | - | - | 2.123 | 4.327 | 4.021 | 2.605 | 1.682 | 2.728 |
| I | - | - | - | - | - | - | - | - | - | 3.93 | 2.96 | 0.384 | 1.326 | 3.238 |
| J | - | - | - | - | - | - | - | - | - | - | 2.965 | 3.917 | 4.501 | 4.943 |
| K | - | - | - | - | - | - | - | - | - | - | - | 3.118 | 3.53 | 4.306 |
| L | - | - | - | - | - | - | - | - | - | - | - | - | 1.095 | 3.412 |
| M | - | - | - | - | - | - | - | - | - | - | - | - | - | 3.229 |
| N | - | - | - | - | - | - | - | - | - | - | - | - | - | - |

| Max | A | B | C | D | E | F | G | H | I | J | K | L | M | N |
|---|---|---|---|---|---|---|---|---|---|---|---|---|---|---|
| A | - | -0.293 | 4.314 | 5.871 | 6.193 | 4.128 | 3.174 | -0.669 | 3.418 | 7.73 | 4.385 | -1.505 | 2.918 | 0.133 |
| B | - | - | 4.385 | 5.914 | 6.272 | 4.202 | 3.31 | -2.432 | 3.326 | 7.768 | 4.453 | -1.083 | 3.085 | -1.469 |
| C | - | - | - | 5.023 | 5.507 | 1.044 | 3.182 | 4.098 | 4.677 | 7.336 | -0.262 | 4.078 | 3.36 | 4.135 |
| D | - | - | - | - | 3.342 | 4.716 | 4.972 | 5.227 | 5.457 | 6.584 | 4.58 | 5.224 | 5.016 | 5.245 |
| E | - | - | - | - | - | 5.062 | 5.233 | 5.433 | 5.626 | 6.273 | 4.96 | 5.441 | 5.279 | 5.449 |
| F | - | - | - | - | - | - | 2.839 | 3.945 | 4.585 | 7.408 | 1.564 | 3.923 | 3.068 | 3.985 |
| G | - | - | - | - | - | - | - | 3.146 | 4.182 | 7.553 | 3.446 | 3.103 | 0.394 | 3.223 |
| H | - | - | - | - | - | - | - | - | 3.346 | 7.757 | 4.438 | -1.835 | 3.05 | -0.99 |
| I | - | - | - | - | - | - | - | - | - | 7.998 | 5.149 | 3.541 | 4.361 | 3.428 |
| J | - | - | - | - | - | - | - | - | - | - | 5.948 | 6.098 | 6.042 | 6.105 |
| K | - | - | - | - | - | - | - | - | - | - | - | 4.136 | 3.461 | 4.19 |
| L | - | - | - | - | - | - | - | - | - | - | - | - | 2.996 | -0.303 |
| M | - | - | - | - | - | - | - | - | - | - | - | - | - | 3.024 |
| N | - | - | - | - | - | - | - | - | - | - | - | - | - | - |
