## Supplementary Data S2 for "Interlaboratory comparison of standardised metabolomics and lipidomics analyses in human and rodent blood using the MxP® Quant 500 kit": TG(18.1_36.1).pdf

**Amount of relative differences between laboratories**

| Min | A | B | C | D | E | F | G | H | I | J | K | L | M | N |
|---|---|---|---|---|---|---|---|---|---|---|---|---|---|---|
| A | - | 6.182 | 6.372 | 6.559 | 6.973 | 6.828 | 6.213 | 5.487 | 2.413 | 10.42 | 5.807 | 6.224 | 5.947 | 5.966 |
| B | - | - | 3.679 | 4.337 | 6.218 | 5.72 | 1.019 | 3.743 | 5.958 | 10.266 | 2.826 | 2.353 | 2.72 | 2.511 |
| C | - | - | - | 2.424 | 5.931 | 5.273 | 3.355 | 4.307 | 6.055 | 10.02 | 3.97 | 2.755 | 4.073 | 3.989 |
| D | - | - | - | - | 5.815 | 5.071 | 3.888 | 4.473 | 6.091 | 9.964 | 4.234 | 3.482 | 4.366 | 4.307 |
| E | - | - | - | - | - | 5.124 | 6.644 | 6.577 | 7.294 | 9.83 | 6.592 | 6.558 | 6.695 | 6.674 |
| F | - | - | - | - | - | - | 5.807 | 5.831 | 6.775 | 9.866 | 5.802 | 5.676 | 5.914 | 5.887 |
| G | - | - | - | - | - | - | - | 3.706 | 5.91 | 10.113 | 2.947 | 1.653 | 2.947 | 2.792 |
| H | - | - | - | - | - | - | - | - | 5.6 | 10.416 | 3.065 | 4.552 | 3.409 | 3.543 |
| I | - | - | - | - | - | - | - | - | - | 10.781 | 5.914 | 6.395 | 6.087 | 6.114 |
| J | - | - | - | - | - | - | - | - | - | - | 7.019 | 7.172 | 7.176 | 7.151 |
| K | - | - | - | - | - | - | - | - | - | - | - | 3.4 | -3.452 | 0.006 |
| L | - | - | - | - | - | - | - | - | - | - | - | - | 3.505 | 3.388 |
| M | - | - | - | - | - | - | - | - | - | - | - | - | - | 0.032 |
| N | - | - | - | - | - | - | - | - | - | - | - | - | - | - |

| | |
|---|---|
| | Larger than 10 |
| | Between 8 and 9,99 |
| | Between 6 and 7,99 |
| | Between 4 and 5,99 |
| | Between 2 and 3,99 |
| | Between 0 and 1,99 |
| | NA (Calculation failed) |

| Median | A | B | C | D | E | F | G | H | I | J | K | L | M | N |
|---|---|---|---|---|---|---|---|---|---|---|---|---|---|---|
| A | - | 3.762 | 3.178 | 3.014 | 4.827 | 4.045 | 3.407 | 3.864 | 3.872 | 4.058 | 2.672 | 3.827 | 3.917 | 4.165 |
| B | - | - | 2.592 | 2.975 | 4.047 | 1.804 | 2.169 | 0.233 | 0.092 | -2.279 | 3.34 | -1.643 | 1.189 | 2.504 |
| C | - | - | - | 0.942 | 4.409 | 3.199 | 0.469 | 2.811 | 2.779 | 1.639 | 2.038 | 2.41 | 3.031 | 3.513 |
| D | - | - | - | - | 4.51 | 3.464 | 1.713 | 3.158 | 3.132 | 2.463 | 1.105 | 2.834 | 3.335 | 3.73 |
| E | - | - | - | - | - | 3.858 | 4.657 | 4.1 | 4.13 | 4.857 | 5.038 | 4.347 | 3.938 | 3.47 |
| F | - | - | - | - | - | - | 3.188 | 1.22 | 1.349 | 3.314 | 3.955 | 2.264 | -0.245 | 1.488 |
| G | - | - | - | - | - | - | - | 2.465 | 2.426 | 0.899 | 2.45 | 1.997 | 2.728 | 3.304 |
| H | - | - | - | - | - | - | - | - | -4.143 | 0.315 | 3.444 | 0.266 | -0.038 | 2.132 |
| I | - | - | - | - | - | - | - | - | - | 3.124 | 3.22 | -1.598 | -1.315 | 1.894 |
| J | - | - | - | - | - | - | - | - | - | - | 2.385 | 3.917 | 4.406 | 4.616 |
| K | - | - | - | - | - | - | - | - | - | - | - | 3.158 | 3.506 | 3.854 |
| L | - | - | - | - | - | - | - | - | - | - | - | - | 1.444 | 2.611 |
| M | - | - | - | - | - | - | - | - | - | - | - | - | - | 1.793 |
| N | - | - | - | - | - | - | - | - | - | - | - | - | - | - |

| Max | A | B | C | D | E | F | G | H | I | J | K | L | M | N |
|---|---|---|---|---|---|---|---|---|---|---|---|---|---|---|
| A | - | 1.27 | 3.883 | 4.296 | 2.749 | 3.926 | 3.184 | 1.996 | 4.114 | 7.511 | 3.812 | 2.145 | 1.265 | 2.682 |
| B | - | - | 3.597 | 4.078 | 2.134 | 3.654 | 2.705 | 2.646 | 4.269 | 7.498 | 3.507 | 0.992 | 2.232 | 3.106 |
| C | - | - | - | 2.092 | 2.782 | -1.176 | 2.317 | 4.034 | 4.807 | 7.22 | -0.615 | 3.174 | 3.906 | 4.207 |
| D | - | - | - | - | 3.416 | 1.873 | 3.15 | 4.307 | 4.95 | 7.134 | 2.234 | 3.673 | 4.207 | 4.447 |
| E | - | - | - | - | - | 2.973 | 1.044 | 3.35 | 4.501 | 7.384 | 2.74 | 1.229 | 3.129 | 3.635 |
| F | - | - | - | - | - | - | 2.433 | 4.065 | 4.82 | 7.215 | 0.091 | 3.238 | 3.942 | 4.233 |
| G | - | - | - | - | - | - | - | 3.585 | 4.599 | 7.328 | 2.183 | 2.089 | 3.397 | 3.829 |
| H | - | - | - | - | - | - | - | - | 3.796 | 7.64 | 4.233 | 3.135 | 0.729 | 1.325 |
| I | - | - | - | - | - | - | - | - | - | 7.958 | 5.246 | 4.717 | 4.173 | 3.723 |
| J | - | - | - | - | - | - | - | - | - | - | 5.925 | 6.002 | 6.054 | 6.078 |
| K | - | - | - | - | - | - | - | - | - | - | - | 3.075 | 3.852 | 4.165 |
| L | - | - | - | - | - | - | - | - | - | - | - | - | 2.713 | 3.378 |
| M | - | - | - | - | - | - | - | - | - | - | - | - | - | 2.034 |
| N | - | - | - | - | - | - | - | - | - | - | - | - | - | - |
