## Supplementary Data S2 for "Interlaboratory comparison of standardised metabolomics and lipidomics analyses in human and rodent blood using the MxP® Quant 500 kit": TG(18.1_36.6).pdf

**Amount of relative differences between laboratories**

| Min | A | B | C | D | E | F | G | H | I | J | K | L | M | N |
|---|---|---|---|---|---|---|---|---|---|---|---|---|---|---|
| A | - | 3.821 | 4.582 | 4.954 | NA | 5.615 | 2.976 | 3.953 | 4.025 | 6.058 | 5.364 | 4.864 | 4.779 | 4.752 |
| B | - | - | 2.851 | 4.008 | NA | 5.036 | 3.254 | 1.124 | 0.755 | 5.683 | 4.803 | 3.953 | 3.659 | 3.296 |
| C | - | - | - | 3.268 | NA | 4.682 | 4.079 | 3.297 | 3.226 | 5.298 | 4.496 | 3.246 | 2.62 | 1.386 |
| D | - | - | - | - | NA | 4.096 | 4.823 | 4.387 | 4.345 | 5.183 | 3.851 | -2.26 | 1.712 | 2.931 |
| E | - | - | - | - | - | NA | NA | NA | NA | NA | NA | NA | NA | NA |
| F | - | - | - | - | - | - | 5.465 | 5.228 | 5.235 | 1.934 | 0.573 | 3.653 | 4.092 | 4.581 |
| G | - | - | - | - | - | - | - | 2.828 | 2.927 | 5.74 | 5.132 | 4.503 | 4.346 | 4.196 |
| H | - | - | - | - | - | - | - | - | -2.617 | 5.285 | 4.85 | 4.153 | 3.875 | 3.422 |
| I | - | - | - | - | - | - | - | - | - | 6.07 | 4.925 | 3.934 | 3.771 | 3.679 |
| J | - | - | - | - | - | - | - | - | - | - | 2.875 | 2.138 | 3.132 | 4.083 |
| K | - | - | - | - | - | - | - | - | - | - | - | 4.022 | 4.34 | 4.698 |
| L | - | - | - | - | - | - | - | - | - | - | - | - | 1.661 | 2.407 |
| M | - | - | - | - | - | - | - | - | - | - | - | - | - | 1.757 |
| N | - | - | - | - | - | - | - | - | - | - | - | - | - | - |

| Median | A | B | C | D | E | F | G | H | I | J | K | L | M | N |
|---|---|---|---|---|---|---|---|---|---|---|---|---|---|---|
| A | - | 3.09 | 3.45 | 4.098 | NA | 4.183 | -0.176 | -3.856 | 3.853 | 3.466 | 4.69 | 4.582 | 4.234 | 4.122 |
| B | - | - | 1.846 | 3.351 | NA | 3.695 | 2.862 | 3.243 | 3.002 | 1.62 | 4.447 | 4.124 | 3.515 | 3.528 |
| C | - | - | - | 2.769 | NA | 3.346 | 3.482 | 3.804 | 2.27 | 1.657 | 4.302 | 3.832 | 2.974 | 3.105 |
| D | - | - | - | - | NA | 2.109 | 4.257 | 4.489 | 0.78 | 3.103 | 3.872 | 2.975 | 0.084 | 1.257 |
| E | - | - | - | - | - | NA | NA | NA | NA | NA | NA | NA | NA | NA |
| F | - | - | - | - | - | - | 4.683 | 4.925 | 2.809 | 4.494 | 3.549 | 1.279 | 2.347 | 1.187 |
| G | - | - | - | - | - | - | - | 0.83 | 3.82 | 3.03 | 4.741 | 4.523 | 4.132 | 4.116 |
| H | - | - | - | - | - | - | - | - | 4.092 | 3.049 | 4.936 | 4.639 | 4.267 | 4.358 |
| I | - | - | - | - | - | - | - | - | - | -0.537 | 3.777 | 3.436 | 2.303 | 1.855 |
| J | - | - | - | - | - | - | - | - | - | - | 5.043 | 4.392 | 3.872 | 4.317 |
| K | - | - | - | - | - | - | - | - | - | - | - | 2.947 | 3.967 | 3.845 |
| L | - | - | - | - | - | - | - | - | - | - | - | - | 2.838 | 2.883 |
| M | - | - | - | - | - | - | - | - | - | - | - | - | - | -1.217 |
| N | - | - | - | - | - | - | - | - | - | - | - | - | - | - |

| Max | A | B | C | D | E | F | G | H | I | J | K | L | M | N |
|---|---|---|---|---|---|---|---|---|---|---|---|---|---|---|
| A | - | 2.608 | 2.956 | 2.237 | NA | 3.828 | 3.35 | 4.996 | 3.655 | 6.83 | 3.413 | 3.977 | 1.928 | 2.984 |
| B | - | - | 0.675 | -0.079 | NA | 2.981 | 1.875 | 4.609 | 4.156 | 6.72 | 3.983 | 4.359 | 3.225 | 3.728 |
| C | - | - | - | 1.25 | NA | 2.608 | 1.065 | 4.489 | 4.261 | 6.623 | 4.082 | 4.445 | 3.436 | 3.866 |
| D | - | - | - | - | NA | 3.186 | 2.192 | 4.676 | 4.096 | 6.774 | 3.893 | 4.324 | 3.102 | 3.631 |
| E | - | - | - | - | - | NA | NA | NA | NA | NA | NA | NA | NA | NA |
| F | - | - | - | - | - | - | 1.863 | 3.986 | 4.586 | 6.352 | 4.444 | 4.716 | 3.972 | 4.273 |
| G | - | - | - | - | - | - | - | 4.319 | 4.372 | 6.547 | 4.214 | 4.555 | 3.663 | 4.031 |
| H | - | - | - | - | - | - | - | - | 5.094 | 5.923 | 5.017 | 5.179 | 4.765 | 4.92 |
| I | - | - | - | - | - | - | - | - | - | 7.258 | 1.437 | 1.905 | 3.349 | 2.473 |
| J | - | - | - | - | - | - | - | - | - | - | 5.863 | 5.939 | 5.789 | 5.826 |
| K | - | - | - | - | - | - | - | - | - | - | - | 2.546 | 2.933 | 1.493 |
| L | - | - | - | - | - | - | - | - | - | - | - | - | 3.815 | 3.24 |
| M | - | - | - | - | - | - | - | - | - | - | - | - | - | 2.082 |
| N | - | - | - | - | - | - | - | - | - | - | - | - | - | - |

| Colour | Legend |
|---|---|
| (red) | Larger than 10 |
| (orange) | Between 8 and 9,99 |
| (light orange) | Between 6 and 7,99 |
| (light green) | Between 4 and 5,99 |
| (green) | Between 2 and 3,99 |
| (dark green) | Between 0 and 1,99 |
| (grey) | NA (Calculation failed) |
