## Supplementary Data S2 for "Interlaboratory comparison of standardised metabolomics and lipidomics analyses in human and rodent blood using the MxP® Quant 500 kit": TG(18.1_38.7).pdf

Amount of relative differences between laboratories

| Min | A | B | C | D | E | F | G | H | I | J | K | L | M | N |
|---|---|---|---|---|---|---|---|---|---|---|---|---|---|---|
| A | - | 5.435 | 5.892 | 4.603 | 4.645 | 5.807 | 3.159 | 4.894 | 4.226 | 8.089 | 2.814 | 4.335 | 4.646 | 4.329 |
| B | - | - | 0.53 | 4.681 | 4.229 | 2.706 | 5.443 | 5.32 | 4.571 | 7.555 | 5.686 | 4.749 | 4.43 | 4.935 |
| C | - | - | - | 2.766 | 1.896 | 3.509 | 4.161 | 4.962 | -2.236 | 6.529 | 4.757 | 2.192 | 1.273 | 3.374 |
| D | - | - | - | - | 3.13 | 4.821 | 3.731 | 3.121 | -1.482 | 7.756 | 4.322 | -1.786 | 1.478 | 2.071 |
| E | - | - | - | - | - | 2.678 | 4.491 | 5.369 | 2.058 | 6.433 | 4.982 | 3.195 | 3.106 | 3.961 |
| F | - | - | - | - | - | - | 4.992 | 5.543 | 3.561 | 6.15 | 5.44 | 4.096 | 3.923 | 4.538 |
| G | - | - | - | - | - | - | - | 2.503 | 3.811 | 7.146 | 2.573 | 3.462 | 3.716 | 2.78 |
| H | - | - | - | - | - | - | - | - | 4.035 | 7.214 | 1.763 | 3.805 | 4.021 | 3.32 |
| I | - | - | - | - | - | - | - | - | - | 8.392 | 3.681 | 1.373 | 3.125 | 1.252 |
| J | - | - | - | - | - | - | - | - | - | - | 5.047 | 3.096 | 3.183 | 4.106 |
| K | - | - | - | - | - | - | - | - | - | - | - | 3.952 | 4.225 | 3.798 |
| L | - | - | - | - | - | - | - | - | - | - | - | - | 1.165 | 2.349 |
| M | - | - | - | - | - | - | - | - | - | - | - | - | - | 2.873 |
| N | - | - | - | - | - | - | - | - | - | - | - | - | - | - |

| | |
|---|---|
| | Larger than 10 |
| | Between 8 and 9,99 |
| | Between 6 and 7,99 |
| | Between 4 and 5,99 |
| | Between 2 and 3,99 |
| | Between 0 and 1,99 |
| | NA (Calculation failed) |

| Median | A | B | C | D | E | F | G | H | I | J | K | L | M | N |
|---|---|---|---|---|---|---|---|---|---|---|---|---|---|---|
| A | - | 4.643 | 2.396 | 3.19 | 0.535 | 2.476 | 3.823 | 3.87 | 4.329 | 4.473 | 2.882 | 0.706 | 2.883 | 3.293 |
| B | - | - | 5.277 | 4.452 | 5.08 | 4.612 | 5.713 | 5.734 | 2.883 | 5.952 | 5.554 | 5.142 | 4.621 | 4.342 |
| C | - | - | - | 3.446 | 0.726 | 3.387 | 3.459 | 3.552 | 4.415 | 4.06 | 3.203 | 0.82 | 3.101 | 3.638 |
| D | - | - | - | - | 3.088 | 1.235 | 4.636 | 4.686 | 3.577 | 5.019 | 4.322 | 3.453 | 1.148 | 0.284 |
| E | - | - | - | - | - | 3.242 | 3.652 | 3.759 | 4.318 | 4.156 | 3.462 | -1.453 | 2.768 | 3.482 |
| F | - | - | - | - | - | - | 4.581 | 4.628 | 3.599 | 4.943 | 4.479 | 3.278 | 1.137 | 1.185 |
| G | - | - | - | - | - | - | - | -0.559 | 4.87 | 2.401 | 1.494 | 3.498 | 4.151 | 4.393 |
| H | - | - | - | - | - | - | - | - | 4.898 | 2.305 | 1.836 | 3.569 | 4.205 | 4.427 |
| I | - | - | - | - | - | - | - | - | - | 5.732 | 5.086 | 4.773 | 3.991 | 3.706 |
| J | - | - | - | - | - | - | - | - | - | - | 0.937 | 3.87 | 4.261 | 4.565 |
| K | - | - | - | - | - | - | - | - | - | - | - | 2.885 | 3.884 | 4.151 |
| L | - | - | - | - | - | - | - | - | - | - | - | - | 2.864 | 3.446 |
| M | - | - | - | - | - | - | - | - | - | - | - | - | - | 1.855 |
| N | - | - | - | - | - | - | - | - | - | - | - | - | - | - |

| Max | A | B | C | D | E | F | G | H | I | J | K | L | M | N |
|---|---|---|---|---|---|---|---|---|---|---|---|---|---|---|
| A | - | 3.518 | 5.479 | 2.376 | 5.633 | 4.915 | 5.563 | 5.454 | 4.442 | 6.96 | 4.672 | 4.931 | 3.761 | 0.08 |
| B | - | - | 5.983 | 4.197 | 6.055 | 5.536 | 6.029 | 5.953 | 3.577 | 7.217 | 5.369 | 5.539 | 4.819 | 3.527 |
| C | - | - | - | 4.786 | 1.497 | 3.293 | 0.154 | 0.084 | 5.531 | 5.77 | 3.724 | 3.344 | 4.434 | 4.993 |
| D | - | - | - | - | 5.52 | 4.588 | 5.374 | 5.208 | 4.677 | 6.893 | 4.321 | 4.648 | 3.026 | 2.542 |
| E | - | - | - | - | - | 3.54 | 0.715 | 2.25 | 5.595 | 5.756 | 3.895 | 3.698 | 4.649 | 5.084 |
| F | - | - | - | - | - | - | 3.656 | 3.366 | 5.333 | 6.047 | 2.083 | -3.298 | 3.72 | 4.582 |
| G | - | - | - | - | - | - | - | 1.382 | 5.56 | 5.816 | 3.791 | 3.545 | 4.539 | 5.046 |
| H | - | - | - | - | - | - | - | - | 5.512 | 5.864 | 3.624 | 3.183 | 4.397 | 4.968 |
| I | - | - | - | - | - | - | - | - | - | 7.557 | 5.921 | 6.065 | 5.495 | 4.71 |
| J | - | - | - | - | - | - | - | - | - | - | 5.52 | 5.502 | 5.719 | 5.869 |
| K | - | - | - | - | - | - | - | - | - | - | - | 1.734 | 3.335 | 4.396 |
| L | - | - | - | - | - | - | - | - | - | - | - | - | 3.728 | 4.614 |
| M | - | - | - | - | - | - | - | - | - | - | - | - | - | 3.692 |
| N | - | - | - | - | - | - | - | - | - | - | - | - | - | - |
