## Supplementary Data S2 for "Interlaboratory comparison of standardised metabolomics and lipidomics analyses in human and rodent blood using the MxP® Quant 500 kit": TG(18.2_33.0).pdf

**Amount of relative differences between laboratories**

| Min | A | B | C | D | E | F | G | H | I | J | K | L | M | N |
|---|---|---|---|---|---|---|---|---|---|---|---|---|---|---|
| A | - | 4.516 | 5.092 | 5.905 | 6.71 | 5.537 | 3.809 | 4.77 | 3.305 | 6.611 | 4.26 | 4.441 | 4.785 | 5.283 |
| B | - | - | 3.449 | 4.956 | 6.235 | 4.76 | 3.161 | 1.782 | 4.7 | 5.862 | 2.179 | 1.719 | 0.91 | 3.538 |
| C | - | - | - | 3.827 | 5.871 | 4.173 | 4.282 | 2.925 | 5.427 | 5.131 | 3.939 | 3.884 | 3.339 | 0.439 |
| D | - | - | - | - | 5.388 | 3.383 | 4.74 | 4.116 | 5.878 | 4.003 | 4.583 | 4.666 | 4.344 | 3.645 |
| E | - | - | - | - | - | 4.889 | 6.332 | 6.095 | 7.035 | 5.338 | 6.267 | 6.304 | 6.215 | 6.031 |
| F | - | - | - | - | - | - | 5.53 | 4.975 | 6.218 | 4.551 | 5.36 | 5.332 | 5.105 | 4.547 |
| G | - | - | - | - | - | - | - | 3.601 | 3.805 | 6.034 | 2.144 | 2.585 | 3.454 | 4.367 |
| H | - | - | - | - | - | - | - | - | 4.962 | 5.638 | 2.982 | 2.772 | 0.899 | 2.953 |
| I | - | - | - | - | - | - | - | - | - | 6.61 | 3.456 | 3.777 | 4.32 | 5.004 |
| J | - | - | - | - | - | - | - | - | - | - | 4.191 | 4.264 | 4.031 | 3.557 |
| K | - | - | - | - | - | - | - | - | - | - | - | 0.405 | 2.66 | 3.994 |
| L | - | - | - | - | - | - | - | - | - | - | - | - | 2.371 | 3.933 |
| M | - | - | - | - | - | - | - | - | - | - | - | - | - | 3.257 |
| N | - | - | - | - | - | - | - | - | - | - | - | - | - | - |

| Median | A | B | C | D | E | F | G | H | I | J | K | L | M | N |
|---|---|---|---|---|---|---|---|---|---|---|---|---|---|---|
| A | - | 0.78 | 0.369 | 2.44 | 3.424 | 4.265 | 1.845 | -0.244 | 2.903 | 3.681 | 0.975 | 0.089 | -0.246 | 3.702 |
| B | - | - | 0.863 | 2.674 | 3.562 | 4.192 | 2.295 | -2.983 | 2.564 | 4.328 | 1.696 | -1.589 | -0.047 | 3.419 |
| C | - | - | - | 2.349 | 3.821 | 4.283 | 1.8 | 0.862 | 2.81 | 4.474 | 0.842 | 1.015 | 0.099 | 3.5 |
| D | - | - | - | - | 4.288 | 4.545 | 0.623 | 2.742 | 3.464 | 4.274 | 1.597 | 2.763 | 2.691 | 3.893 |
| E | - | - | - | - | - | 1.599 | 4.635 | 4.139 | 3.694 | 5.809 | 4.511 | 4.123 | 4.126 | 3.138 |
| F | - | - | - | - | - | - | 4.832 | 4.457 | 3.949 | 5.676 | 4.722 | 4.441 | 4.521 | 3.323 |
| G | - | - | - | - | - | - | - | 2.241 | 3.376 | 3.797 | 0.676 | 2.304 | 1.835 | 3.919 |
| H | - | - | - | - | - | - | - | - | 2.502 | 4.455 | 1.782 | -2.218 | 0.023 | 3.372 |
| I | - | - | - | - | - | - | - | - | - | 4.423 | 3.346 | 2.775 | 3.148 | 2.568 |
| J | - | - | - | - | - | - | - | - | - | - | 4.499 | 4.716 | 4.725 | 4.961 |
| K | - | - | - | - | - | - | - | - | - | - | - | 1.783 | 1.097 | 3.764 |
| L | - | - | - | - | - | - | - | - | - | - | - | - | 0.501 | 3.411 |
| M | - | - | - | - | - | - | - | - | - | - | - | - | - | 3.502 |
| N | - | - | - | - | - | - | - | - | - | - | - | - | - | - |

| Max | A | B | C | D | E | F | G | H | I | J | K | L | M | N |
|---|---|---|---|---|---|---|---|---|---|---|---|---|---|---|
| A | - | 4.073 | 4.99 | 5.861 | 5.692 | 2.147 | 4.212 | 4.298 | 2.683 | 7.415 | 4.28 | 3.752 | 4.341 | 2.362 |
| B | - | - | 3.677 | 5.192 | 4.926 | 3.408 | 0.565 | 1.288 | 4.319 | 7.041 | 1.123 | 1.528 | 1.543 | 3.33 |
| C | - | - | - | 4.368 | 3.941 | 4.372 | 3.332 | 3.189 | 4.861 | 6.7 | 3.233 | 3.79 | 3.124 | 4.336 |
| D | - | - | - | - | 2.226 | 5.111 | 4.674 | 4.662 | 5.385 | 6.126 | 4.663 | 4.88 | 4.629 | 5.088 |
| E | - | - | - | - | - | 4.974 | 4.487 | 4.442 | 5.29 | 6.292 | 4.461 | 4.696 | 4.418 | 4.946 |
| F | - | - | - | - | - | - | 3.748 | 3.868 | 3.399 | 7.299 | 3.834 | 3.114 | 3.918 | -0.644 |
| G | - | - | - | - | - | - | - | 0.028 | 4.4 | 6.991 | -0.445 | 2.067 | 0.565 | 3.494 |
| H | - | - | - | - | - | - | - | - | 4.452 | 6.957 | -2.02 | 2.371 | -1.199 | 3.607 |
| I | - | - | - | - | - | - | - | - | - | 7.581 | 4.789 | 4.414 | 4.836 | 3.63 |
| J | - | - | - | - | - | - | - | - | - | - | 5.818 | 5.865 | 5.799 | 5.935 |
| K | - | - | - | - | - | - | - | - | - | - | - | 2.302 | -0.475 | 3.578 |
| L | - | - | - | - | - | - | - | - | - | - | - | - | 2.564 | 2.887 |
| M | - | - | - | - | - | - | - | - | - | - | - | - | - | 3.652 |
| N | - | - | - | - | - | - | - | - | - | - | - | - | - | - |

| | |
|---|---|
| | Larger than 10 |
| | Between 8 and 9,99 |
| | Between 6 and 7,99 |
| | Between 4 and 5,99 |
| | Between 2 and 3,99 |
| | Between 0 and 1,99 |
| | NA (Calculation failed) |
