## Supplementary Data S2 for "Interlaboratory comparison of standardised metabolomics and lipidomics analyses in human and rodent blood using the MxP® Quant 500 kit": TG(18.2_33.1).pdf

**Amount of relative differences between laboratories**

| Min | A | B | C | D | E | F | G | H | I | J | K | L | M | N |
|---|---|---|---|---|---|---|---|---|---|---|---|---|---|---|
| A | - | 4.68 | 4.233 | 4.441 | 5.77 | 5.002 | 3.864 | 4.152 | 3.787 | 6.279 | 3.419 | 4.647 | 4.161 | 4.768 |
| B | - | - | 3.109 | 2.542 | 4.968 | 3.411 | 3.338 | 3.227 | 3.908 | 5.738 | 4.117 | -1.06 | 2.824 | 1.953 |
| C | - | - | - | 1.456 | 5.211 | 4.143 | 0.733 | -0.938 | 2.557 | 5.547 | 3.009 | 3.07 | 0.306 | 2.323 |
| D | - | - | - | - | 5.107 | 3.978 | 1.954 | 1.703 | 3.099 | 5.728 | 3.438 | 2.597 | 0.047 | 0.599 |
| E | - | - | - | - | - | 4.295 | 5.51 | 5.567 | 5.777 | 0.726 | 5.819 | 5.086 | 5.415 | 5.479 |
| F | - | - | - | - | - | - | 4.519 | 4.488 | 4.85 | 4.971 | 4.96 | 3.52 | 4.289 | 4.105 |
| G | - | - | - | - | - | - | - | 0.657 | 1.875 | 5.773 | 2.609 | 3.372 | 1.635 | 2.966 |
| H | - | - | - | - | - | - | - | - | 2.358 | 5.858 | 2.934 | 3.16 | 0.657 | 2.695 |
| I | - | - | - | - | - | - | - | - | - | 6.408 | 1.784 | 3.72 | 2.289 | 3.926 |
| J | - | - | - | - | - | - | - | - | - | - | 4.783 | 3.49 | 4.101 | 4.554 |
| K | - | - | - | - | - | - | - | - | - | - | - | 3.951 | 3.129 | 3.656 |
| L | - | - | - | - | - | - | - | - | - | - | - | - | 2.963 | 1.934 |
| M | - | - | - | - | - | - | - | - | - | - | - | - | - | 2.501 |
| N | - | - | - | - | - | - | - | - | - | - | - | - | - | - |

| | |
|---|---|
| | Larger than 10 |
| | Between 8 and 9,99 |
| | Between 6 and 7,99 |
| | Between 4 and 5,99 |
| | Between 2 and 3,99 |
| | Between 0 and 1,99 |
| | NA (Calculation failed) |

| Median | A | B | C | D | E | F | G | H | I | J | K | L | M | N |
|---|---|---|---|---|---|---|---|---|---|---|---|---|---|---|
| A | - | 2.96 | 1.29 | -0.289 | 4.35 | 4.14 | 1.942 | 2.338 | 3.602 | 3.872 | 2.231 | 3.301 | 2.906 | 3.351 |
| B | - | - | 2.755 | 2.877 | 3.874 | 3.088 | 2.622 | 2.117 | 2.225 | 1.008 | 3.808 | 1.383 | -0.098 | 1.357 |
| C | - | - | - | -0.625 | 4.312 | 3.85 | -0.637 | 1.234 | 3.411 | 1.961 | 2.769 | 3.116 | 2.44 | 3.117 |
| D | - | - | - | - | 4.36 | 3.871 | -0.097 | 1.431 | 3.484 | 2.909 | 2.651 | 3.215 | 2.562 | 3.203 |
| E | - | - | - | - | - | 3.315 | 4.752 | 4.609 | 3.642 | 4.906 | 5.123 | 3.87 | 4.299 | 3.904 |
| F | - | - | - | - | - | - | 3.993 | 3.788 | 1.922 | 3.741 | 4.596 | 2.58 | 3.318 | 2.603 |
| G | - | - | - | - | - | - | - | 0.725 | 3.294 | 2.558 | 2.957 | 2.96 | 2.221 | 2.969 |
| H | - | - | - | - | - | - | - | - | 3.057 | 2.573 | 3.254 | 2.639 | 1.615 | 2.653 |
| I | - | - | - | - | - | - | - | - | - | 2.891 | 4.263 | 1.354 | 2.421 | 1.123 |
| J | - | - | - | - | - | - | - | - | - | - | 2.508 | 4.341 | 3.979 | 4.313 |
| K | - | - | - | - | - | - | - | - | - | - | - | 3.857 | 3.526 | 3.858 |
| L | - | - | - | - | - | - | - | - | - | - | - | - | 1.817 | -5.642 |
| M | - | - | - | - | - | - | - | - | - | - | - | - | - | 1.715 |
| N | - | - | - | - | - | - | - | - | - | - | - | - | - | - |

| Max | A | B | C | D | E | F | G | H | I | J | K | L | M | N |
|---|---|---|---|---|---|---|---|---|---|---|---|---|---|---|
| A | - | 3.407 | 3.942 | 4.329 | 3.241 | 2.206 | 3.472 | 3.173 | 3.446 | 6.619 | 4.454 | 2.408 | 2.046 | 1.269 |
| B | - | - | 2.235 | 3.149 | -2.76 | 2.382 | -0.636 | 0.295 | 4.283 | 6.492 | 3.36 | 2.242 | 2.57 | 2.873 |
| C | - | - | - | 2.015 | 2.283 | 3.237 | 1.947 | 2.468 | 4.527 | 6.188 | 2.443 | 3.16 | 3.338 | 3.51 |
| D | - | - | - | - | 3.064 | 3.689 | 2.934 | 3.215 | 4.702 | 6.21 | 0.238 | 3.645 | 3.773 | 3.898 |
| E | - | - | - | - | - | 2.32 | -0.064 | 0.06 | 4.272 | 6.438 | 3.417 | 2.224 | 2.589 | 2.8 |
| F | - | - | - | - | - | - | 2.622 | 2.095 | 3.911 | 6.547 | 4.036 | -1.003 | -0.448 | 1.111 |
| G | - | - | - | - | - | - | - | 0.798 | 4.317 | 6.349 | 3.267 | 2.382 | 2.679 | 2.971 |
| H | - | - | - | - | - | - | - | - | 4.204 | 6.447 | 3.545 | 1.816 | 2.241 | 2.622 |
| I | - | - | - | - | - | - | - | - | - | 7.038 | 5.203 | 4.19 | 4.072 | 3.911 |
| J | - | - | - | - | - | - | - | - | - | - | 5.422 | 5.692 | 5.717 | 5.7 |
| K | - | - | - | - | - | - | - | - | - | - | - | 3.757 | 3.87 | 3.991 |
| L | - | - | - | - | - | - | - | - | - | - | - | - | 0.33 | 1.482 |
| M | - | - | - | - | - | - | - | - | - | - | - | - | - | 0.605 |
| N | - | - | - | - | - | - | - | - | - | - | - | - | - | - |
