## Supplementary Data S2 for "Interlaboratory comparison of standardised metabolomics and lipidomics analyses in human and rodent blood using the MxP® Quant 500 kit": TG(18.2_34.2).pdf

**Amount of relative differences between laboratories**

| Min | A | B | C | D | E | F | G | H | I | J | K | L | M | N |
|---|---|---|---|---|---|---|---|---|---|---|---|---|---|---|
| A | - | 4.307 | 3.937 | 2.554 | 4.895 | 4.967 | 3.485 | 3.997 | 3.821 | 5.149 | 3.715 | 3.501 | 4.137 | 3.749 |
| B | - | - | 2.473 | 4.1 | 3.83 | 3.499 | 3.413 | 2.331 | 3.463 | 3.889 | 2.742 | 3.131 | 1.593 | 3.398 |
| C | - | - | - | 3.453 | 4.237 | 3.887 | 2.249 | -1.543 | 2.709 | 4.17 | -0.78 | 1.426 | 1.226 | 2.429 |
| D | - | - | - | - | 4.757 | 4.658 | 2.454 | 3.332 | 2.569 | 4.753 | 3.096 | 2.796 | 3.583 | 2.664 |
| E | - | - | - | - | - | 1.311 | 4.852 | 4.556 | 4.844 | 1.009 | 4.651 | 4.795 | 4.309 | 4.808 |
| F | - | - | - | - | - | - | 4.462 | 4.019 | 4.664 | -2.911 | 4.091 | 4.287 | 3.79 | 4.568 |
| G | - | - | - | - | - | - | - | 2.326 | -0.809 | 4.522 | 1.859 | 0.801 | 2.781 | -4.794 |
| H | - | - | - | - | - | - | - | - | 2.346 | 4.325 | 0.682 | 1.842 | 0.94 | 2.237 |
| I | - | - | - | - | - | - | - | - | - | 5.039 | -1.291 | 1.328 | 2.116 | -4.516 |
| J | - | - | - | - | - | - | - | - | - | - | 2.333 | 2.835 | 2.314 | 4.06 |
| K | - | - | - | - | - | - | - | - | - | - | - | 0.84 | 1.994 | 1.268 |
| L | - | - | - | - | - | - | - | - | - | - | - | - | 2.658 | -1.578 |
| M | - | - | - | - | - | - | - | - | - | - | - | - | - | 2.814 |
| N | - | - | - | - | - | - | - | - | - | - | - | - | - | - |

| | |
|---|---|
| | Larger than 10 |
| | Between 8 and 9,99 |
| | Between 6 and 7,99 |
| | Between 4 and 5,99 |
| | Between 2 and 3,99 |
| | Between 0 and 1,99 |
| | NA (Calculation failed) |

| Median | A | B | C | D | E | F | G | H | I | J | K | L | M | N |
|---|---|---|---|---|---|---|---|---|---|---|---|---|---|---|
| A | - | 3.773 | 2.486 | 0.54 | 4.729 | 4.184 | 1.835 | 3.307 | 3.748 | 3.532 | 3.322 | 3.281 | 3.49 | 3.646 |
| B | - | - | 3.172 | 3.804 | 3.915 | 2.354 | 3.479 | 2.034 | -1.717 | 4.776 | 2.163 | 2.1 | 1.613 | 0.274 |
| C | - | - | - | 2.21 | 4.447 | 3.676 | 0.887 | 2.162 | 3.121 | 4.008 | 1.956 | 2.063 | 2.395 | 2.848 |
| D | - | - | - | - | 4.692 | 4.072 | 1.204 | 3.08 | 3.611 | 3.698 | 3.086 | 3.102 | 3.264 | 3.478 |
| E | - | - | - | - | - | 3.549 | 4.959 | 4.539 | 4.141 | 5.695 | 4.547 | 4.543 | 4.417 | 4.264 |
| F | - | - | - | - | - | - | 4.052 | 3.237 | 2.196 | 5.08 | 3.369 | 3.306 | 3.11 | 2.685 |
| G | - | - | - | - | - | - | - | 2.659 | 3.365 | 3.856 | 2.572 | 2.618 | 2.855 | 3.158 |
| H | - | - | - | - | - | - | - | - | 2.09 | 4.48 | -2.142 | -3.294 | 0.072 | 1.528 |
| I | - | - | - | - | - | - | - | - | - | 4.698 | 1.938 | 2.051 | 1.361 | -0.256 |
| J | - | - | - | - | - | - | - | - | - | - | 3.723 | 3.869 | 3.899 | 4.174 |
| K | - | - | - | - | - | - | - | - | - | - | - | -3.465 | 0.41 | 1.547 |
| L | - | - | - | - | - | - | - | - | - | - | - | - | 0.521 | 1.473 |
| M | - | - | - | - | - | - | - | - | - | - | - | - | - | 0.921 |
| N | - | - | - | - | - | - | - | - | - | - | - | - | - | - |

| Max | A | B | C | D | E | F | G | H | I | J | K | L | M | N |
|---|---|---|---|---|---|---|---|---|---|---|---|---|---|---|
| A | - | 2.051 | 3.239 | 2.275 | 4.369 | 0.828 | 2.847 | 0.614 | 3.689 | 6.447 | 1.997 | 2.83 | 0.312 | 3.537 |
| B | - | - | 3.845 | 3.238 | 4.058 | 1.401 | 3.567 | 1.487 | 3.184 | 6.574 | -2.041 | 1.574 | 1.644 | 2.976 |
| C | - | - | - | 2.13 | 4.763 | 3.36 | 1.169 | 3.347 | 4.359 | 6.149 | 3.637 | 3.938 | 3.3 | 4.281 |
| D | - | - | - | - | 4.568 | 2.647 | 1.124 | 2.64 | 4.095 | 6.271 | 3.067 | 3.492 | 2.554 | 3.984 |
| E | - | - | - | - | - | 4.539 | 5.097 | 4.515 | 3.179 | 7.043 | 4.325 | 4.008 | 4.572 | 3.414 |
| F | - | - | - | - | - | - | 3.155 | -3.692 | 3.527 | 6.459 | 1.336 | 2.526 | -1.086 | 3.375 |
| G | - | - | - | - | - | - | - | 3.019 | 4.227 | 6.224 | 3.377 | 3.731 | 2.964 | 4.141 |
| H | - | - | - | - | - | - | - | - | 3.523 | 6.499 | 1.272 | 2.473 | -1.689 | 3.361 |
| I | - | - | - | - | - | - | - | - | - | 6.914 | 3.377 | 2.766 | 3.745 | 0.761 |
| J | - | - | - | - | - | - | - | - | - | - | 5.59 | 5.642 | 5.529 | 5.696 |
| K | - | - | - | - | - | - | - | - | - | - | - | 1.652 | 1.505 | 3.019 |
| L | - | - | - | - | - | - | - | - | - | - | - | - | 2.638 | 2.365 |
| M | - | - | - | - | - | - | - | - | - | - | - | - | - | 3.403 |
| N | - | - | - | - | - | - | - | - | - | - | - | - | - | - |
