## Supplementary Data S2 for "Interlaboratory comparison of standardised metabolomics and lipidomics analyses in human and rodent blood using the MxP® Quant 500 kit": TG(18.2_34.3).pdf

**Amount of relative differences between laboratories**

| Min | A | B | C | D | E | F | G | H | I | J | K | L | M | N |
|---|---|---|---|---|---|---|---|---|---|---|---|---|---|---|
| A | - | 4.373 | 4.281 | 3.224 | 5.191 | 5.051 | 3.806 | 4.571 | 3.672 | 5.686 | 4.015 | 3.93 | 4.295 | 4.413 |
| B | - | - | 0.507 | 3.793 | 4.376 | 3.947 | 2.936 | 1.587 | 3.306 | 4.788 | 1.163 | 2.155 | 0.822 | -0.158 |
| C | - | - | - | 3.564 | 4.464 | 4.034 | 2.55 | 1.957 | 3.052 | 4.755 | -1.045 | 1.3 | -0.766 | -1.443 |
| D | - | - | - | - | 4.958 | 4.691 | 2.435 | 3.869 | 1.668 | 5.145 | 3.308 | 3.06 | 3.386 | 3.541 |
| E | - | - | - | - | - | 2.758 | 5.163 | 4.558 | 5.296 | 1.368 | 4.854 | 5.016 | 4.854 | 4.787 |
| F | - | - | - | - | - | - | 4.718 | 3.923 | 4.895 | 2.24 | 4.195 | 4.472 | 4.349 | 4.318 |
| G | - | - | - | - | - | - | - | 3.269 | 1.116 | 4.967 | 2.435 | 1.643 | 2.405 | 2.59 |
| H | - | - | - | - | - | - | - | - | 3.673 | 4.635 | 1.952 | 2.75 | 2.236 | 1.918 |
| I | - | - | - | - | - | - | - | - | - | 5.597 | 2.299 | 1.956 | 2.984 | 3.287 |
| J | - | - | - | - | - | - | - | - | - | - | 2.686 | 3.557 | 3.901 | 3.742 |
| K | - | - | - | - | - | - | - | - | - | - | - | 0.796 | 0.4 | 1.278 |
| L | - | - | - | - | - | - | - | - | - | - | - | - | 1.865 | 2.131 |
| M | - | - | - | - | - | - | - | - | - | - | - | - | - | 0.268 |
| N | - | - | - | - | - | - | - | - | - | - | - | - | - | - |

| Median | A | B | C | D | E | F | G | H | I | J | K | L | M | N |
|---|---|---|---|---|---|---|---|---|---|---|---|---|---|---|
| A | - | 3.566 | 2.584 | 0.254 | 4.835 | 4.237 | 1.708 | 3.452 | 3.427 | 4.167 | 3.583 | 3.529 | 3.33 | 3.961 |
| B | - | - | 2.655 | 3.558 | 4.259 | 3.059 | 3.229 | -0.19 | -0.577 | 5.003 | -1.268 | -2.452 | 1.185 | 2.074 |
| C | - | - | - | 2.346 | 4.579 | 3.785 | 1.527 | 2.322 | 2.439 | 4.528 | 2.359 | 2.475 | 1.82 | 3.307 |
| D | - | - | - | - | 4.802 | 4.168 | 1.1 | 3.272 | 3.293 | 4.241 | 3.374 | 3.395 | 3.087 | 3.842 |
| E | - | - | - | - | - | 3.729 | 5.139 | 4.644 | 4.616 | 5.991 | 4.622 | 4.598 | 4.737 | 4.205 |
| F | - | - | - | - | - | - | 4.277 | 3.365 | 3.285 | 5.436 | 3.397 | 3.289 | 3.601 | 2.25 |
| G | - | - | - | - | - | - | - | 2.951 | 2.999 | 4.359 | 3.011 | 3.063 | 2.679 | 3.631 |
| H | - | - | - | - | - | - | - | - | -1.318 | 4.896 | -2.012 | -0.615 | 0.593 | 2.355 |
| I | - | - | - | - | - | - | - | - | - | 4.943 | 0.385 | -0.01 | -0.15 | 2.426 |
| J | - | - | - | - | - | - | - | - | - | - | 3.984 | 4.2 | 3.997 | 4.522 |
| K | - | - | - | - | - | - | - | - | - | - | - | -3.664 | 0.984 | 2.145 |
| L | - | - | - | - | - | - | - | - | - | - | - | - | 0.957 | 2.088 |
| M | - | - | - | - | - | - | - | - | - | - | - | - | - | 2.691 |
| N | - | - | - | - | - | - | - | - | - | - | - | - | - | - |

| Max | A | B | C | D | E | F | G | H | I | J | K | L | M | N |
|---|---|---|---|---|---|---|---|---|---|---|---|---|---|---|
| A | - | 1.847 | 2.729 | 2.597 | 4.386 | 2.407 | 2.807 | -1.251 | 3.222 | 6.403 | 2.542 | 2.952 | 0.166 | 3.491 |
| B | - | - | 3.443 | 3.312 | 4.138 | 0.69 | 3.463 | 2.11 | 2.564 | 6.481 | 1.182 | 2.074 | 1.457 | 3.013 |
| C | - | - | - | -0.138 | 4.68 | 3.502 | -4.924 | 2.583 | 3.928 | 6.153 | 3.58 | 3.788 | 2.883 | 4.096 |
| D | - | - | - | - | 4.636 | 3.397 | -0.028 | 2.4 | 3.862 | 6.15 | 3.467 | 3.673 | 2.739 | 4.034 |
| E | - | - | - | - | - | 4.279 | 5.115 | 4.716 | 3.804 | 6.99 | 4.193 | 3.986 | 4.619 | 3.526 |
| F | - | - | - | - | - | - | 3.672 | 2.583 | 2.183 | 6.482 | -0.131 | 1.58 | 2.154 | 2.749 |
| G | - | - | - | - | - | - | - | 2.594 | 3.931 | 6.134 | 3.581 | 3.78 | 2.895 | 4.105 |
| H | - | - | - | - | - | - | - | - | 3.292 | 6.361 | 2.657 | 3.043 | 0.566 | 3.566 |
| I | - | - | - | - | - | - | - | - | - | 6.738 | 2.005 | 0.981 | 3.19 | 1.04 |
| J | - | - | - | - | - | - | - | - | - | - | 5.602 | 5.626 | 5.486 | 5.657 |
| K | - | - | - | - | - | - | - | - | - | - | - | 0.96 | 2.338 | 2.568 |
| L | - | - | - | - | - | - | - | - | - | - | - | - | 2.838 | 2.056 |
| M | - | - | - | - | - | - | - | - | - | - | - | - | - | 3.382 |
| N | - | - | - | - | - | - | - | - | - | - | - | - | - | - |

| | |
|---|---|
| (red) | Larger than 10 |
| (orange) | Between 8 and 9,99 |
| (light orange) | Between 6 and 7,99 |
| (light green) | Between 4 and 5,99 |
| (green) | Between 2 and 3,99 |
| (dark green) | Between 0 and 1,99 |
| (grey) | NA (Calculation failed) |
