## Supplementary Data S2 for "Interlaboratory comparison of standardised metabolomics and lipidomics analyses in human and rodent blood using the MxP® Quant 500 kit": TG(18.2_35.1).pdf

**Amount of relative differences between laboratories**

| Min | A | B | C | D | E | F | G | H | I | J | K | L | M | N |
|---|---|---|---|---|---|---|---|---|---|---|---|---|---|---|
| A | - | 4.095 | 4.135 | 4.285 | 5.585 | 5.208 | 3.931 | 4.168 | 2.161 | 7.336 | 2.388 | 4.618 | 4.122 | 4.732 |
| B | - | - | -0.243 | 0.877 | 5.011 | 4.431 | -0.098 | 1.013 | 4.339 | 6.973 | 3.896 | 2.793 | 0.444 | 1.013 |
| C | - | - | - | -0.16 | 4.97 | 4.368 | 0.828 | 0.287 | 4.431 | 6.853 | 3.995 | 2.587 | -0.79 | -0.577 |
| D | - | - | - | - | 4.834 | 4.257 | 0.464 | 0.453 | 4.605 | 6.629 | 4.053 | 2.214 | -0.547 | 2.045 |
| E | - | - | - | - | - | 3.364 | 5.105 | 4.957 | 6.134 | 5.541 | 5.815 | 4.867 | 4.993 | 5.546 |
| F | - | - | - | - | - | - | 4.637 | 4.427 | 5.761 | 6.208 | 5.465 | 4.148 | 4.477 | 4.871 |
| G | - | - | - | - | - | - | - | 1.525 | 4.305 | 6.791 | 3.814 | 2.907 | 1.176 | 0.946 |
| H | - | - | - | - | - | - | - | - | 4.489 | 6.939 | 4.105 | 2.363 | -0.554 | -0.694 |
| I | - | - | - | - | - | - | - | - | - | 7.529 | -2.092 | 4.382 | 3.679 | 4.575 |
| J | - | - | - | - | - | - | - | - | - | - | 4.755 | 2.554 | 2.106 | 4.767 |
| K | - | - | - | - | - | - | - | - | - | - | - | 4.261 | 3.798 | 4.001 |
| L | - | - | - | - | - | - | - | - | - | - | - | - | 2.393 | 2.792 |
| M | - | - | - | - | - | - | - | - | - | - | - | - | - | 0.287 |
| N | - | - | - | - | - | - | - | - | - | - | - | - | - | - |

| | |
|---|---|
| | Larger than 10 |
| | Between 8 and 9,99 |
| | Between 6 and 7,99 |
| | Between 4 and 5,99 |
| | Between 2 and 3,99 |
| | Between 0 and 1,99 |
| | NA (Calculation failed) |

| Median | A | B | C | D | E | F | G | H | I | J | K | L | M | N |
|---|---|---|---|---|---|---|---|---|---|---|---|---|---|---|
| A | - | 2.624 | 2.494 | -5.919 | 4.036 | 4.073 | 1.702 | 3.207 | 2.719 | 3.872 | 2.603 | 3.179 | 2.852 | 3.225 |
| B | - | - | -6.57 | 2.672 | 3.563 | 3.603 | 1.181 | 1.348 | 0.369 | 4.535 | 3.645 | 1.91 | 0.89 | 1.995 |
| C | - | - | - | 2.683 | 3.565 | 3.603 | 1.183 | 1.289 | 0.381 | 4.607 | 3.648 | 1.911 | 0.893 | 1.997 |
| D | - | - | - | - | 4.133 | 4.153 | 2.104 | 2.975 | 2.934 | 4.056 | 2.47 | 3.3 | 3.017 | 3.327 |
| E | - | - | - | - | - | -2.269 | 4.016 | 3.742 | 3.595 | 5.505 | 4.834 | 3.257 | 3.54 | 3.232 |
| F | - | - | - | - | - | - | 4.037 | 3.619 | 3.631 | 5.49 | 4.833 | 3.272 | 3.56 | 3.239 |
| G | - | - | - | - | - | - | - | 2.179 | 1.801 | 4.466 | 3.332 | 2.557 | 2.008 | 2.61 |
| H | - | - | - | - | - | - | - | - | 0.463 | 4.767 | 3.957 | 0.22 | -0.334 | 0.508 |
| I | - | - | - | - | - | - | - | - | - | 4.353 | 3.756 | 1.415 | -0.584 | 1.581 |
| J | - | - | - | - | - | - | - | - | - | - | 3.817 | 4.77 | 4.702 | 4.775 |
| K | - | - | - | - | - | - | - | - | - | - | - | 3.85 | 3.669 | 3.874 |
| L | - | - | - | - | - | - | - | - | - | - | - | - | 0.973 | -2.271 |
| M | - | - | - | - | - | - | - | - | - | - | - | - | - | 1.137 |
| N | - | - | - | - | - | - | - | - | - | - | - | - | - | - |

| Max | A | B | C | D | E | F | G | H | I | J | K | L | M | N |
|---|---|---|---|---|---|---|---|---|---|---|---|---|---|---|
| A | - | 2.754 | 2.942 | 4.597 | 4.021 | 1.926 | 3.43 | 1.659 | 2.963 | 7.207 | 4.135 | 2.564 | 2.071 | 0.2 |
| B | - | - | -0.119 | 4.052 | 3.217 | 1.327 | 1.936 | 1.72 | 3.768 | 7.084 | 3.335 | -0.519 | 1.265 | 2.375 |
| C | - | - | - | 3.956 | 3.058 | 1.781 | 1.51 | 2.063 | 3.849 | 7.024 | 3.182 | 0.67 | 1.718 | 2.589 |
| D | - | - | - | - | 2.63 | 4.037 | 3.452 | 4.094 | 4.699 | 6.7 | 2.51 | 3.895 | 4.035 | 4.223 |
| E | - | - | - | - | - | 3.455 | 2.365 | 3.549 | 4.408 | 6.845 | -1.877 | 3.229 | 3.461 | 3.739 |
| F | - | - | - | - | - | - | 2.705 | -0.31 | 3.535 | 7.075 | 3.652 | 0.907 | -4.459 | 1.404 |
| G | - | - | - | - | - | - | - | 2.78 | 4.071 | 6.936 | 2.586 | 2.115 | 2.585 | 3.113 |
| H | - | - | - | - | - | - | - | - | 3.419 | 7.135 | 3.79 | 1.42 | -0.121 | 0.935 |
| I | - | - | - | - | - | - | - | - | - | 7.455 | 4.78 | 3.896 | 3.694 | 3.28 |
| J | - | - | - | - | - | - | - | - | - | - | 5.804 | 5.889 | 5.907 | 5.923 |
| K | - | - | - | - | - | - | - | - | - | - | - | 3.299 | 3.509 | 3.791 |
| L | - | - | - | - | - | - | - | - | - | - | - | - | 0.784 | 2.188 |
| M | - | - | - | - | - | - | - | - | - | - | - | - | - | 1.484 |
| N | - | - | - | - | - | - | - | - | - | - | - | - | - | - |
