## Supplementary Data S2 for "Interlaboratory comparison of standardised metabolomics and lipidomics analyses in human and rodent blood using the MxP® Quant 500 kit": TG(18.2_36.2).pdf

**Amount of relative differences between laboratories**

| Min | A | B | C | D | E | F | G | H | I | J | K | L | M | N |
|---|---|---|---|---|---|---|---|---|---|---|---|---|---|---|
| A | - | 4.642 | 4.201 | 4.245 | 5.618 | 5.347 | 3.874 | 4.176 | 3.019 | 6.468 | 4.059 | 4.395 | 4.312 | 4.912 |
| B | - | - | 3.152 | 3.492 | 4.797 | 3.783 | 3.683 | 2.904 | 4.524 | 5.814 | 3.122 | 2.167 | 2.61 | -0.395 |
| C | - | - | - | 1.485 | 5.035 | 4.289 | 1.795 | 0.683 | 3.706 | 5.784 | -0.794 | 2.05 | 1.444 | 2.968 |
| D | - | - | - | - | 5.117 | 4.449 | -0.697 | 2.081 | 3.315 | 5.954 | 1.671 | 2.737 | 2.39 | 3.354 |
| E | - | - | - | - | - | 4.349 | 5.6 | 5.356 | 5.982 | 4.644 | 5.401 | 5.255 | 5.326 | 5.197 |
| F | - | - | - | - | - | - | 4.625 | 4.196 | 5.211 | 5.031 | 4.281 | 3.979 | 4.119 | 3.784 |
| G | - | - | - | - | - | - | - | 2.251 | 3.195 | 5.891 | 1.881 | 2.876 | 2.578 | 3.561 |
| H | - | - | - | - | - | - | - | - | 3.785 | 6.092 | 0.37 | 1.58 | 0.634 | 3.311 |
| I | - | - | - | - | - | - | - | - | - | 6.605 | 3.188 | 3.807 | 3.66 | 4.637 |
| J | - | - | - | - | - | - | - | - | - | - | 3.592 | 3.401 | 3.549 | 4.138 |
| K | - | - | - | - | - | - | - | - | - | - | - | 1.937 | 1.236 | 3.118 |
| L | - | - | - | - | - | - | - | - | - | - | - | - | 0.596 | 2.314 |
| M | - | - | - | - | - | - | - | - | - | - | - | - | - | 2.929 |
| N | - | - | - | - | - | - | - | - | - | - | - | - | - | - |

| Median | A | B | C | D | E | F | G | H | I | J | K | L | M | N |
|---|---|---|---|---|---|---|---|---|---|---|---|---|---|---|
| A | - | 3.637 | -4.853 | 0.99 | 4.646 | 3.591 | -0.926 | 3.167 | 3.265 | 4.277 | 2.155 | 2.966 | 3.041 | 3.796 |
| B | - | - | 3.667 | 3.949 | 3.925 | -0.585 | 3.747 | 1.439 | 1.023 | 5.186 | 3.158 | 2.095 | 1.816 | 1.358 |
| C | - | - | - | 1.357 | 4.623 | 3.561 | -0.561 | 3.143 | 3.236 | 4.42 | 1.629 | 2.885 | 3.022 | 3.75 |
| D | - | - | - | - | 4.73 | 3.809 | 0.885 | 3.473 | 3.547 | 4.117 | 2.463 | 3.277 | 3.378 | 3.963 |
| E | - | - | - | - | - | 4.073 | 5.054 | 4.37 | 4.314 | 5.924 | 4.863 | 4.513 | 4.434 | 3.885 |
| F | - | - | - | - | - | - | 3.815 | 1.73 | 1.426 | 5.278 | 3.372 | 2.348 | 2.038 | 0.841 |
| G | - | - | - | - | - | - | - | 3.239 | 3.326 | 4.359 | 1.974 | 3.009 | 3.126 | 3.815 |
| H | - | - | - | - | - | - | - | - | -0.574 | 5.002 | 2.512 | 0.537 | -0.326 | 2.393 |
| I | - | - | - | - | - | - | - | - | - | 4.933 | 2.487 | 0.978 | 0.628 | 2.268 |
| J | - | - | - | - | - | - | - | - | - | - | 4.328 | 4.671 | 4.743 | 4.926 |
| K | - | - | - | - | - | - | - | - | - | - | - | 2.097 | 2.314 | 3.402 |
| L | - | - | - | - | - | - | - | - | - | - | - | - | -0.47 | 2.709 |
| M | - | - | - | - | - | - | - | - | - | - | - | - | - | 2.575 |
| N | - | - | - | - | - | - | - | - | - | - | - | - | - | - |

| Max | A | B | C | D | E | F | G | H | I | J | K | L | M | N |
|---|---|---|---|---|---|---|---|---|---|---|---|---|---|---|
| A | - | 0.119 | 4.142 | 4.338 | 1.607 | 3.456 | 3.937 | -0.873 | 3.406 | 7.289 | 3.378 | 2.355 | 1.24 | 1.505 |
| B | - | - | 4.031 | 4.256 | 1.869 | 3.302 | 3.807 | 0.778 | 3.543 | 7.268 | 3.2 | 2.022 | 0.227 | 1.923 |
| C | - | - | - | 1.236 | 4.104 | 2.479 | 1.029 | 3.958 | 4.591 | 6.889 | 2.621 | 3.401 | 3.705 | 4.112 |
| D | - | - | - | - | 4.252 | 2.948 | 2.086 | 4.128 | 4.69 | 6.884 | 3.06 | 3.661 | 3.916 | 4.268 |
| E | - | - | - | - | - | 3.804 | 4.196 | 0.905 | 3.068 | 7.406 | 3.716 | 2.965 | 2.259 | -2.599 |
| F | - | - | - | - | - | - | 1.907 | 3.407 | 4.303 | 7.018 | -0.503 | 2.417 | 2.995 | 3.632 |
| G | - | - | - | - | - | - | - | 3.785 | 4.495 | 6.93 | 2.079 | 3.128 | 3.492 | 3.966 |
| H | - | - | - | - | - | - | - | - | 3.338 | 7.316 | 3.469 | 2.534 | 1.554 | 1.051 |
| I | - | - | - | - | - | - | - | - | - | 7.593 | 4.555 | 4.135 | 3.854 | 3.146 |
| J | - | - | - | - | - | - | - | - | - | - | 5.886 | 5.923 | 5.95 | 5.98 |
| K | - | - | - | - | - | - | - | - | - | - | - | 2.237 | 2.879 | 3.567 |
| L | - | - | - | - | - | - | - | - | - | - | - | - | 1.462 | 2.927 |
| M | - | - | - | - | - | - | - | - | - | - | - | - | - | 2.273 |
| N | - | - | - | - | - | - | - | - | - | - | - | - | - | - |

| | |
|---|---|
| | Larger than 10 |
| | Between 8 and 9,99 |
| | Between 6 and 7,99 |
| | Between 4 and 5,99 |
| | Between 2 and 3,99 |
| | Between 0 and 1,99 |
| | NA (Calculation failed) |
