## Supplementary Data S2 for "Interlaboratory comparison of standardised metabolomics and lipidomics analyses in human and rodent blood using the MxP® Quant 500 kit": TG(18.2_36.3).pdf

**Amount of relative differences between laboratories**

| Min | A | B | C | D | E | F | G | H | I | J | K | L | M | N |
|---|---|---|---|---|---|---|---|---|---|---|---|---|---|---|
| A | - | 4.049 | 3.913 | 1.74 | 4.993 | 4.793 | 3.242 | 4.462 | 3.139 | 6.151 | 3.311 | 2.822 | 4.161 | 3.668 |
| B | - | - | 1.109 | 3.885 | 4.283 | 3.766 | 3.009 | 2.711 | 3.01 | 5.624 | 2.701 | 3.164 | 0.156 | 1.903 |
| C | - | - | - | 3.539 | 4.468 | 3.959 | 2.476 | 3.083 | 2.374 | 5.58 | 1.847 | 2.562 | 1.578 | -0.591 |
| D | - | - | - | - | 4.913 | 4.657 | 2.61 | 4.235 | 2.427 | 5.931 | 2.849 | 2.272 | 3.869 | 3.252 |
| E | - | - | - | - | - | 2.944 | 5.124 | 4.112 | 5.165 | 5.315 | 5.058 | 5.217 | 4.473 | 4.827 |
| F | - | - | - | - | - | - | 4.602 | 3.034 | 4.526 | 4.662 | 4.421 | 4.562 | 3.88 | 4.182 |
| G | - | - | - | - | - | - | - | 3.74 | -3.062 | 5.729 | 0.75 | -0.773 | 3.067 | 2.075 |
| H | - | - | - | - | - | - | - | - | 3.885 | 5.398 | 3.702 | 3.952 | 2.494 | 3.359 |
| I | - | - | - | - | - | - | - | - | - | 6.205 | 0.763 | 0.551 | 3.401 | 2.161 |
| J | - | - | - | - | - | - | - | - | - | - | 3.112 | 3.051 | 2.714 | 2.377 |
| K | - | - | - | - | - | - | - | - | - | - | - | 1.486 | 3.033 | 1.476 |
| L | - | - | - | - | - | - | - | - | - | - | - | - | 3.593 | 2.428 |
| M | - | - | - | - | - | - | - | - | - | - | - | - | - | 2.264 |
| N | - | - | - | - | - | - | - | - | - | - | - | - | - | - |

| Median | A | B | C | D | E | F | G | H | I | J | K | L | M | N |
|---|---|---|---|---|---|---|---|---|---|---|---|---|---|---|
| A | - | 3.566 | 2.976 | 0.778 | 4.913 | 4.222 | 1.824 | 3.947 | 3.287 | 3.52 | 3.175 | 3.084 | 3.697 | 3.753 |
| B | - | - | 2.148 | 3.518 | 4.419 | 2.991 | 3.16 | 1.994 | 1.112 | 4.725 | 1.411 | 1.682 | 0.134 | 1.107 |
| C | - | - | - | 2.731 | 4.653 | 3.57 | 2.074 | 2.986 | 1.159 | 4.332 | 0.908 | 0.322 | 2.356 | 2.724 |
| D | - | - | - | - | 4.875 | 4.121 | 0.983 | 3.786 | 3.012 | 3.872 | 2.95 | 2.876 | 3.509 | 3.584 |
| E | - | - | - | - | - | 4.111 | 5.284 | 4.501 | 4.93 | 5.966 | 4.951 | 4.993 | 4.67 | 4.592 |
| F | - | - | - | - | - | - | 4.153 | 2.189 | 3.41 | 5.185 | 3.454 | 3.553 | 2.94 | 2.499 |
| G | - | - | - | - | - | - | - | 3.554 | 2.632 | 3.951 | 2.545 | 2.401 | 3.191 | 3.357 |
| H | - | - | - | - | - | - | - | - | 2.618 | 4.914 | 2.697 | 2.844 | 1.514 | 0.642 |
| I | - | - | - | - | - | - | - | - | - | 4.514 | -0.677 | 0.198 | 1.911 | 2.055 |
| J | - | - | - | - | - | - | - | - | - | - | 3.978 | 3.918 | 4.064 | 4.333 |
| K | - | - | - | - | - | - | - | - | - | - | - | -1.01 | 2.016 | 2.248 |
| L | - | - | - | - | - | - | - | - | - | - | - | - | 2.298 | 2.319 |
| M | - | - | - | - | - | - | - | - | - | - | - | - | - | 0.161 |
| N | - | - | - | - | - | - | - | - | - | - | - | - | - | - |

| Max | A | B | C | D | E | F | G | H | I | J | K | L | M | N |
|---|---|---|---|---|---|---|---|---|---|---|---|---|---|---|
| A | - | 2.14 | 1.622 | 0.122 | 4.79 | 1.393 | 2.431 | 2.014 | 3.495 | 6.668 | 2.935 | 3.446 | 2.452 | 3.867 |
| B | - | - | 2.981 | 2.521 | 4.594 | 1.034 | 3.362 | -1.215 | 2.845 | 6.771 | 1.724 | 2.735 | 0.099 | 3.424 |
| C | - | - | - | 0.973 | 4.883 | 2.431 | 1.118 | 2.802 | 3.807 | 6.561 | 3.374 | 3.757 | 3.065 | 4.103 |
| D | - | - | - | - | 4.819 | 1.827 | 2.089 | 2.359 | 3.624 | 6.605 | 3.093 | 3.542 | 2.694 | 3.961 |
| E | - | - | - | - | - | 5.097 | 5.496 | 4.994 | 4.457 | 7.446 | 4.767 | 4.504 | 4.926 | 4.162 |
| F | - | - | - | - | - | - | 3.001 | 0.754 | 3.179 | 6.698 | 2.396 | 3.11 | 1.616 | 3.646 |
| G | - | - | - | - | - | - | - | 3.164 | 3.985 | 6.497 | 3.618 | 3.939 | 3.363 | 4.249 |
| H | - | - | - | - | - | - | - | - | 2.916 | 6.771 | 1.861 | 2.815 | 0.461 | 3.471 |
| I | - | - | - | - | - | - | - | - | - | 7.006 | 2.08 | -0.731 | 2.723 | 1.885 |
| J | - | - | - | - | - | - | - | - | - | - | 5.744 | 5.788 | 5.706 | 5.834 |
| K | - | - | - | - | - | - | - | - | - | - | - | 1.789 | 1.177 | 2.929 |
| L | - | - | - | - | - | - | - | - | - | - | - | - | 2.579 | 2.145 |
| M | - | - | - | - | - | - | - | - | - | - | - | - | - | 3.283 |
| N | - | - | - | - | - | - | - | - | - | - | - | - | - | - |

| | |
|---|---|
| | Larger than 10 |
| | Between 8 and 9,99 |
| | Between 6 and 7,99 |
| | Between 4 and 5,99 |
| | Between 2 and 3,99 |
| | Between 0 and 1,99 |
| | NA (Calculation failed) |
