## Supplementary Data S2 for "Interlaboratory comparison of standardised metabolomics and lipidomics analyses in human and rodent blood using the MxP® Quant 500 kit": TG(18.2_36.4).pdf

**Amount of relative differences between laboratories**

| Min | A | B | C | D | E | F | G | H | I | J | K | L | M | N |
|---|---|---|---|---|---|---|---|---|---|---|---|---|---|---|
| A | - | 4.261 | 4.14 | 3.447 | 5.541 | 5.407 | 2.912 | 5.178 | 2.714 | 6.112 | 2.302 | -0.456 | 4.592 | 3.168 |
| B | - | - | 1.502 | 2.928 | 4.993 | 4.677 | 3.907 | 4.308 | 3.914 | 5.63 | 3.978 | 4.362 | 2.216 | 3.479 |
| C | - | - | - | 1.459 | 5.071 | 4.716 | 3.576 | 4.441 | 3.542 | 5.512 | 3.508 | 3.975 | 2.72 | 2.4 |
| D | - | - | - | - | 5.216 | 5.046 | 2.708 | 4.731 | 2.885 | 5.852 | 3.375 | 3.763 | 3.614 | 2.207 |
| E | - | - | - | - | - | 3.394 | 6.101 | 4.219 | 6.149 | 4.483 | 6.114 | 6.271 | 5.31 | 5.898 |
| F | - | - | - | - | - | - | 5.536 | 1.838 | 5.482 | 3.977 | 5.423 | 5.589 | 4.52 | 5.165 |
| G | - | - | - | - | - | - | - | 4.931 | -0.883 | 5.897 | 0.508 | 2.552 | 4.111 | 1.617 |
| H | - | - | - | - | - | - | - | - | 5.312 | 4.44 | 5.27 | 5.451 | 4.186 | 5.051 |
| I | - | - | - | - | - | - | - | - | - | 6.249 | 1.304 | 2.871 | 4.345 | 1.136 |
| J | - | - | - | - | - | - | - | - | - | - | 3.966 | 4.252 | 2.731 | 3.579 |
| K | - | - | - | - | - | - | - | - | - | - | - | 2.258 | 4.494 | 2.241 |
| L | - | - | - | - | - | - | - | - | - | - | - | - | 4.849 | 3.161 |
| M | - | - | - | - | - | - | - | - | - | - | - | - | - | 3.982 |
| N | - | - | - | - | - | - | - | - | - | - | - | - | - | - |

| Median | A | B | C | D | E | F | G | H | I | J | K | L | M | N |
|---|---|---|---|---|---|---|---|---|---|---|---|---|---|---|
| A | - | 4.002 | 3.477 | 3.093 | 5.312 | 4.863 | 2.179 | 4.809 | 3.285 | 1.142 | 2.967 | 2.611 | 4.279 | 3.723 |
| B | - | - | 2.451 | 2.906 | 4.911 | 3.98 | 3.678 | 3.819 | 2.701 | 4.026 | 3.16 | 3.415 | 1.847 | 1.183 |
| C | - | - | - | 0.355 | 5.118 | 4.324 | 2.78 | 4.195 | -0.079 | 3.35 | 1.668 | 2.199 | 2.994 | 2.136 |
| D | - | - | - | - | 5.135 | 4.519 | 2.251 | 4.381 | -0.686 | 2.5 | 1.724 | 2.002 | 3.445 | 1.996 |
| E | - | - | - | - | - | 4.445 | 5.971 | 4.629 | 5.785 | 6.07 | 5.848 | 5.922 | 5.295 | 5.522 |
| F | - | - | - | - | - | - | 5.032 | 1.403 | 4.655 | 5.214 | 4.821 | 4.891 | 3.96 | 4.221 |
| G | - | - | - | - | - | - | - | 4.593 | 2.453 | 1.611 | 1.788 | 0.797 | 3.894 | 3.243 |
| H | - | - | - | - | - | - | - | - | 4.459 | 5.057 | 4.651 | 4.732 | 3.611 | 4.062 |
| I | - | - | - | - | - | - | - | - | - | 2.954 | 0.892 | 2.007 | 3.558 | 1.962 |
| J | - | - | - | - | - | - | - | - | - | - | 2.535 | 2.492 | 3.629 | 3.729 |
| K | - | - | - | - | - | - | - | - | - | - | - | 1.117 | 3.747 | 2.501 |
| L | - | - | - | - | - | - | - | - | - | - | - | - | 4.048 | 2.865 |
| M | - | - | - | - | - | - | - | - | - | - | - | - | - | 2.509 |
| N | - | - | - | - | - | - | - | - | - | - | - | - | - | - |

| Max | A | B | C | D | E | F | G | H | I | J | K | L | M | N |
|---|---|---|---|---|---|---|---|---|---|---|---|---|---|---|
| A | - | 2.989 | 0.971 | 0.415 | 5.1 | 3.115 | -1.113 | 3.043 | 3.909 | 6.618 | 3.879 | 4.14 | 3.189 | 4.397 |
| B | - | - | 3.431 | 2.84 | 4.842 | -0.409 | 3.191 | -1.612 | 2.945 | 6.845 | 2.877 | 3.397 | 0.378 | 3.837 |
| C | - | - | - | 1.665 | 5.153 | 3.384 | 0.634 | 3.33 | 4.067 | 6.554 | 4.036 | 4.268 | 3.458 | 4.488 |
| D | - | - | - | - | 5.068 | 2.897 | 0.858 | 2.804 | 3.795 | 6.655 | 3.762 | 4.045 | 2.985 | 4.325 |
| E | - | - | - | - | - | 5.288 | 5.725 | 5.306 | 4.857 | 7.655 | 4.894 | 4.651 | 5.262 | 4.351 |
| F | - | - | - | - | - | - | 3.308 | -1.375 | 2.827 | 6.863 | 2.741 | 3.308 | -0.708 | 3.754 |
| G | - | - | - | - | - | - | - | 3.116 | 3.948 | 6.6 | 3.916 | 4.172 | 3.258 | 4.42 |
| H | - | - | - | - | - | - | - | - | 2.887 | 6.855 | 2.814 | 3.356 | -0.125 | 3.808 |
| I | - | - | - | - | - | - | - | - | - | 7.06 | -1.344 | 1.609 | 2.803 | 2.821 |
| J | - | - | - | - | - | - | - | - | - | - | 5.832 | 5.87 | 5.756 | 5.914 |
| K | - | - | - | - | - | - | - | - | - | - | - | 1.777 | 2.707 | 2.889 |
| L | - | - | - | - | - | - | - | - | - | - | - | - | 3.376 | 2.067 |
| M | - | - | - | - | - | - | - | - | - | - | - | - | - | 3.718 |
| N | - | - | - | - | - | - | - | - | - | - | - | - | - | - |

| Colour | Legend |
|---|---|
| (red) | Larger than 10 |
| (orange) | Between 8 and 9,99 |
| (light orange) | Between 6 and 7,99 |
| (light green) | Between 4 and 5,99 |
| (green) | Between 2 and 3,99 |
| (dark green) | Between 0 and 1,99 |
| (grey) | NA (Calculation failed) |
