## Supplementary Data S2 for "Interlaboratory comparison of standardised metabolomics and lipidomics analyses in human and rodent blood using the MxP® Quant 500 kit": TG(18.2_38.5).pdf

Amount of relative differences between laboratories

| Min | A | B | C | D | E | F | G | H | I | J | K | L | M | N |
|---|---|---|---|---|---|---|---|---|---|---|---|---|---|---|
| A | - | 4.648 | 4.104 | 1.541 | 4.905 | 5.506 | 3.871 | 4.527 | 3.011 | 6.26 | 3.435 | 3.484 | 4.294 | 4.195 |
| B | - | - | 4.102 | 4.575 | 3.323 | 4.437 | 3.681 | 1.263 | 4.38 | 5.14 | 4.159 | 3.767 | 2.585 | 2.17 |
| C | - | - | - | 2.876 | 4.476 | 5.057 | 1.446 | 3.57 | 1.969 | 5.613 | 0.677 | 0.455 | 3.14 | 3.257 |
| D | - | - | - | - | 4.845 | 5.349 | 3.45 | 4.304 | 2.009 | 6.013 | 2.795 | 3.129 | 4.023 | 3.962 |
| E | - | - | - | - | - | 4.041 | 4.595 | 3.44 | 5.061 | 5.576 | 4.899 | 4.817 | 4.095 | 4.109 |
| F | - | - | - | - | - | - | 5.336 | 4.802 | 5.653 | -0.092 | 5.543 | 5.316 | 4.971 | 4.82 |
| G | - | - | - | - | - | - | - | 3.238 | 2.77 | 5.653 | 2.139 | 0.071 | 2.636 | 2.831 |
| H | - | - | - | - | - | - | - | - | 4.122 | 5.255 | 3.907 | 3.475 | 1.729 | 0.603 |
| I | - | - | - | - | - | - | - | - | - | 6.359 | 1.834 | 1.937 | 3.657 | 3.482 |
| J | - | - | - | - | - | - | - | - | - | - | 4.565 | 4.089 | 3.607 | 2.108 |
| K | - | - | - | - | - | - | - | - | - | - | - | 1.231 | 3.327 | 3.321 |
| L | - | - | - | - | - | - | - | - | - | - | - | - | 3.135 | 2.884 |
| M | - | - | - | - | - | - | - | - | - | - | - | - | - | -0.388 |
| N | - | - | - | - | - | - | - | - | - | - | - | - | - | - |

| | |
|---|---|
| | Larger than 10 |
| | Between 8 and 9,99 |
| | Between 6 and 7,99 |
| | Between 4 and 5,99 |
| | Between 2 and 3,99 |
| | Between 0 and 1,99 |
| | NA (Calculation failed) |

| Median | A | B | C | D | E | F | G | H | I | J | K | L | M | N |
|---|---|---|---|---|---|---|---|---|---|---|---|---|---|---|
| A | - | 3.646 | -1.256 | -1.053 | 4.771 | 4.557 | 1.611 | 3.132 | 2.988 | 3.679 | 1.722 | 3.273 | 3.227 | 3.662 |
| B | - | - | 3.726 | 3.839 | 4.159 | 3.795 | 3.422 | 2.068 | 2.064 | 4.628 | 3.281 | 1.513 | 1.182 | -0.011 |
| C | - | - | - | -0.267 | 4.782 | 4.581 | 1.27 | 3.033 | 3.022 | 3.386 | 1.676 | 3.205 | 3.285 | 3.661 |
| D | - | - | - | - | 4.813 | 4.581 | 1.749 | 3.19 | 3.073 | 3.541 | 1.922 | 3.348 | 3.321 | 3.711 |
| E | - | - | - | - | - | 2.518 | 5.094 | 4.689 | 4.755 | 5.782 | 5.069 | 4.647 | 4.644 | 4.358 |
| F | - | - | - | - | - | - | 4.884 | 4.456 | 4.423 | 5.465 | 4.806 | 4.34 | 4.237 | 3.943 |
| G | - | - | - | - | - | - | - | 2.568 | 2.554 | 3.669 | -0.27 | 2.823 | 2.912 | 3.395 |
| H | - | - | - | - | - | - | - | - | -2.078 | 4.224 | 2.382 | 0.142 | 1.015 | 2.35 |
| I | - | - | - | - | - | - | - | - | - | 4.436 | 2.283 | 1.011 | 0.652 | 2.359 |
| J | - | - | - | - | - | - | - | - | - | - | 2.811 | 3.522 | 3.786 | 4.053 |
| K | - | - | - | - | - | - | - | - | - | - | - | 2.695 | 2.712 | 3.28 |
| L | - | - | - | - | - | - | - | - | - | - | - | - | -1.811 | 1.659 |
| M | - | - | - | - | - | - | - | - | - | - | - | - | - | 1.722 |
| N | - | - | - | - | - | - | - | - | - | - | - | - | - | - |

| Max | A | B | C | D | E | F | G | H | I | J | K | L | M | N |
|---|---|---|---|---|---|---|---|---|---|---|---|---|---|---|
| A | - | 0.036 | 3.523 | 2.125 | 4.631 | 2.8 | 3.228 | 2.79 | 2.97 | 5.958 | 1.759 | 3.004 | -1.865 | 2.627 |
| B | - | - | 3.428 | 1.772 | 4.659 | 3 | 3.079 | 2.623 | 3.114 | 5.854 | 1.304 | 3.132 | 0.056 | 2.865 |
| C | - | - | - | 2.727 | 5.022 | 4.075 | 1.053 | 2.048 | 4.122 | 5.434 | 2.899 | 4.146 | 3.402 | 4.018 |
| D | - | - | - | - | 4.782 | 3.486 | 2.252 | 1.341 | 3.561 | 5.747 | -0.018 | 3.552 | 2.148 | 3.358 |
| E | - | - | - | - | - | 4.493 | 5.488 | 5.399 | 4.465 | 6.804 | 5.205 | 4.438 | 5.001 | 4.599 |
| F | - | - | - | - | - | - | 4.16 | 3.948 | -0.257 | 6.088 | 3.507 | -0.16 | 2.958 | -0.637 |
| G | - | - | - | - | - | - | - | 1.06 | 3.969 | 5.57 | 2.456 | 3.987 | 3.119 | 3.85 |
| H | - | - | - | - | - | - | - | - | 3.792 | 5.646 | 1.805 | 3.838 | 2.756 | 3.665 |
| I | - | - | - | - | - | - | - | - | - | 6.301 | 3.629 | -4.131 | 3.04 | 0.842 |
| J | - | - | - | - | - | - | - | - | - | - | 5.101 | 5.402 | 5.203 | 5.342 |
| K | - | - | - | - | - | - | - | - | - | - | - | 3.451 | 1.792 | 3.231 |
| L | - | - | - | - | - | - | - | - | - | - | - | - | 3.058 | 0.753 |
| M | - | - | - | - | - | - | - | - | - | - | - | - | - | 2.633 |
| N | - | - | - | - | - | - | - | - | - | - | - | - | - | - |
