## Supplementary Data S2 for "Interlaboratory comparison of standardised metabolomics and lipidomics analyses in human and rodent blood using the MxP® Quant 500 kit": TG(18.3_36.2).pdf

**Amount of relative differences between laboratories**

| Min | A | B | C | D | E | F | G | H | I | J | K | L | M | N |
|---|---|---|---|---|---|---|---|---|---|---|---|---|---|---|
| A | - | 4.82 | 4.618 | 4.598 | 5.794 | 4.979 | 4.207 | 4.459 | 3.984 | 6.443 | 4.097 | 4.638 | 4.39 | 4.565 |
| B | - | - | 2.697 | 3.441 | 4.968 | 1.811 | 3.84 | 3.148 | 4.159 | 5.893 | 3.437 | 2.557 | 3.559 | 3.379 |
| C | - | - | - | 2.347 | 5.093 | 3.229 | 2.839 | 0.942 | 3.382 | 5.606 | 1.411 | -0.856 | 2.302 | 2.077 |
| D | - | - | - | - | 5.23 | 3.791 | 1.096 | 1.53 | 2.378 | 6.002 | 1.038 | 2.374 | -3.295 | -0.215 |
| E | - | - | - | - | - | 5.004 | 5.803 | 5.584 | 5.914 | 4.303 | 5.527 | 5.503 | 5.743 | 5.737 |
| F | - | - | - | - | - | - | 4.187 | 3.639 | 4.455 | 5.358 | 3.738 | 3.308 | 3.982 | 3.907 |
| G | - | - | - | - | - | - | - | 2.264 | 1.646 | 5.822 | 1.737 | 2.864 | 1.109 | 1.686 |
| H | - | - | - | - | - | - | - | - | 3.036 | 5.961 | 1.381 | 1.501 | 1.249 | -0.438 |
| I | - | - | - | - | - | - | - | - | - | 6.502 | 1.169 | 3.41 | 2.547 | 3.15 |
| J | - | - | - | - | - | - | - | - | - | - | 2.937 | 3.686 | 4.252 | 4.396 |
| K | - | - | - | - | - | - | - | - | - | - | - | 2.418 | -2.649 | 1.083 |
| L | - | - | - | - | - | - | - | - | - | - | - | - | 2.466 | 2.115 |
| M | - | - | - | - | - | - | - | - | - | - | - | - | - | 0.778 |
| N | - | - | - | - | - | - | - | - | - | - | - | - | - | - |

| | |
|---|---|
| | Larger than 10 |
| | Between 8 and 9,99 |
| | Between 6 and 7,99 |
| | Between 4 and 5,99 |
| | Between 2 and 3,99 |
| | Between 0 and 1,99 |
| | NA (Calculation failed) |

| Median | A | B | C | D | E | F | G | H | I | J | K | L | M | N |
|---|---|---|---|---|---|---|---|---|---|---|---|---|---|---|
| A | - | 3.872 | 2.265 | 0.05 | 4.734 | 3.85 | 0.075 | 3.371 | 3.766 | 3.349 | 2.938 | 3.143 | 2.956 | 3.63 |
| B | - | - | 3.582 | 4.052 | 3.875 | 0.445 | 3.997 | 2.433 | 0.234 | 5.019 | 2.915 | 2.736 | 3.008 | 1.529 |
| C | - | - | - | 2.064 | 4.59 | 3.214 | 1.784 | 2.559 | 3.288 | 4.391 | 2.088 | 2.292 | 1.867 | 3.031 |
| D | - | - | - | - | 4.759 | 3.697 | -0.557 | 3.273 | 3.741 | 3.709 | 3.009 | 3.117 | 2.902 | 3.567 |
| E | - | - | - | - | - | 4.383 | 5.206 | 4.633 | 4.274 | 5.931 | 4.713 | 4.688 | 4.782 | 4.437 |
| F | - | - | - | - | - | - | 3.808 | 1.898 | -0.875 | 5.084 | 2.469 | 2.274 | 2.632 | 0.275 |
| G | - | - | - | - | - | - | - | 3.184 | 3.673 | 4.001 | 2.869 | 3.005 | 2.773 | 3.494 |
| H | - | - | - | - | - | - | - | - | 1.975 | 4.704 | 1.157 | 0.387 | 1.366 | 1.243 |
| I | - | - | - | - | - | - | - | - | - | 4.69 | 2.764 | 2.464 | 2.762 | 0.536 |
| J | - | - | - | - | - | - | - | - | - | - | 4.863 | 4.843 | 4.785 | 4.934 |
| K | - | - | - | - | - | - | - | - | - | - | - | -0.143 | -1.522 | 2.179 |
| L | - | - | - | - | - | - | - | - | - | - | - | - | 0.37 | 1.819 |
| M | - | - | - | - | - | - | - | - | - | - | - | - | - | 2.273 |
| N | - | - | - | - | - | - | - | - | - | - | - | - | - | - |

| Max | A | B | C | D | E | F | G | H | I | J | K | L | M | N |
|---|---|---|---|---|---|---|---|---|---|---|---|---|---|---|
| A | - | 2.159 | 3.466 | 3.903 | 3.217 | -0.207 | 3.574 | 0.836 | 3.627 | 6.942 | -2.524 | 1.198 | -0.192 | 2.559 |
| B | - | - | 4.034 | 4.36 | 2.24 | 2.514 | 4.103 | 1.512 | 3.045 | 7.119 | 2.268 | 2.849 | 2.485 | 0.522 |
| C | - | - | - | 1.829 | 4.174 | 3.2 | -0.548 | 3.54 | 4.405 | 6.67 | 3.32 | 2.983 | 3.21 | 3.935 |
| D | - | - | - | - | 4.381 | 3.609 | 1.473 | 3.872 | 4.577 | 6.68 | 3.703 | 3.472 | 3.635 | 4.184 |
| E | - | - | - | - | - | 3.439 | 4.514 | 2.958 | 1.924 | 7.273 | 3.283 | 3.611 | 3.392 | 1.783 |
| F | - | - | - | - | - | - | 3.444 | 1.428 | 3.727 | 6.917 | -0.088 | 0.41 | -3.692 | 2.749 |
| G | - | - | - | - | - | - | - | 3.613 | 4.441 | 6.647 | 3.406 | 3.095 | 3.306 | 3.99 |
| H | - | - | - | - | - | - | - | - | 3.431 | 7.006 | 0.94 | 2.061 | 1.438 | 2.054 |
| I | - | - | - | - | - | - | - | - | - | 7.33 | 3.837 | 4.062 | 3.91 | 2.885 |
| J | - | - | - | - | - | - | - | - | - | - | 5.897 | 5.845 | 5.859 | 5.913 |
| K | - | - | - | - | - | - | - | - | - | - | - | 1.118 | -0.557 | 2.564 |
| L | - | - | - | - | - | - | - | - | - | - | - | - | 0.56 | 3.009 |
| M | - | - | - | - | - | - | - | - | - | - | - | - | - | 2.728 |
| N | - | - | - | - | - | - | - | - | - | - | - | - | - | - |
