## Supplementary Data S2 for "Interlaboratory comparison of standardised metabolomics and lipidomics analyses in human and rodent blood using the MxP® Quant 500 kit": TG(18.3_36.4).pdf

**Amount of relative differences between laboratories**

| Min | A | B | C | D | E | F | G | H | I | J | K | L | M | N |
|---|---|---|---|---|---|---|---|---|---|---|---|---|---|---|
| A | - | 5.017 | 4.078 | 3.4 | 5.556 | 5.183 | 3.396 | 4.894 | 2.654 | 5.954 | 4.043 | 1.977 | 4.536 | 2.622 |
| B | - | - | 4.111 | 4.644 | 4.397 | 2.349 | 4.8 | -2.273 | 4.939 | 5.031 | 4.064 | 5.031 | 3.405 | 4.952 |
| C | - | - | - | 2.689 | 5.045 | 4.221 | 3.257 | 3.833 | 3.532 | 5.385 | -1.055 | 3.724 | 2.35 | 3.425 |
| D | - | - | - | - | 5.293 | 4.703 | 0.294 | 4.378 | 2.232 | 5.735 | 2.732 | 2.714 | 3.685 | 2.329 |
| E | - | - | - | - | - | 4.469 | 6.021 | 4.907 | 6.114 | 4.02 | 5.64 | 6.173 | 5.384 | 6.031 |
| F | - | - | - | - | - | - | 4.848 | -4.619 | 4.774 | 4.183 | 3.808 | 4.815 | 3.629 | 4.698 |
| G | - | - | - | - | - | - | - | 4.436 | 1.824 | 5.742 | 2.936 | 2.475 | 3.785 | 1.766 |
| H | - | - | - | - | - | - | - | - | 4.879 | 4.936 | 3.957 | 4.946 | 3.444 | 4.889 |
| I | - | - | - | - | - | - | - | - | - | 6.107 | 3.468 | 1.531 | 4.332 | -1.829 |
| J | - | - | - | - | - | - | - | - | - | - | 3.687 | 4.653 | 4.014 | 4.655 |
| K | - | - | - | - | - | - | - | - | - | - | - | 3.903 | 3.026 | 3.625 |
| L | - | - | - | - | - | - | - | - | - | - | - | - | 4.628 | 1.398 |
| M | - | - | - | - | - | - | - | - | - | - | - | - | - | 4.295 |
| N | - | - | - | - | - | - | - | - | - | - | - | - | - | - |

| | |
|---|---|
| | Larger than 10 |
| | Between 8 and 9,99 |
| | Between 6 and 7,99 |
| | Between 4 and 5,99 |
| | Between 2 and 3,99 |
| | Between 0 and 1,99 |
| | NA (Calculation failed) |

| Median | A | B | C | D | E | F | G | H | I | J | K | L | M | N |
|---|---|---|---|---|---|---|---|---|---|---|---|---|---|---|
| A | - | 4.509 | 3.088 | 2.845 | 5.298 | 3.941 | 2.689 | 4.299 | 3.441 | 2.562 | 4.078 | 3.526 | 4.218 | 3.74 |
| B | - | - | 4.248 | 4.348 | 4.451 | 3.498 | 4.432 | 2.217 | 3.959 | 5.258 | 3.003 | 3.894 | 2.69 | 3.648 |
| C | - | - | - | 0.168 | 5.118 | 2.682 | 1.195 | 3.555 | 1.489 | 3.979 | 3.197 | 1.714 | 3.368 | 2.522 |
| D | - | - | - | - | 5.128 | 3.003 | -0.116 | 3.73 | 1.988 | 3.789 | 3.375 | 2.238 | 3.581 | 2.702 |
| E | - | - | - | - | - | 5.431 | 5.821 | 5.143 | 5.616 | 6.258 | 5.277 | 5.596 | 5.196 | 5.441 |
| F | - | - | - | - | - | - | 2.914 | 2.798 | 0.727 | 4.476 | 2.21 | 0.573 | 2.313 | 0.503 |
| G | - | - | - | - | - | - | - | 3.81 | 2.27 | 3.706 | 3.481 | 2.461 | 3.669 | 2.915 |
| H | - | - | - | - | - | - | - | - | 3.316 | 4.913 | 1.511 | 3.239 | 0.717 | 2.844 |
| I | - | - | - | - | - | - | - | - | - | 4.137 | 2.795 | -0.01 | 3.216 | 1.517 |
| J | - | - | - | - | - | - | - | - | - | - | 4.361 | 3.922 | 4.308 | 4.151 |
| K | - | - | - | - | - | - | - | - | - | - | - | 2.677 | 1.05 | 2.081 |
| L | - | - | - | - | - | - | - | - | - | - | - | - | 3.134 | 0.75 |
| M | - | - | - | - | - | - | - | - | - | - | - | - | - | 2.565 |
| N | - | - | - | - | - | - | - | - | - | - | - | - | - | - |

| Max | A | B | C | D | E | F | G | H | I | J | K | L | M | N |
|---|---|---|---|---|---|---|---|---|---|---|---|---|---|---|
| A | - | 3.995 | 0.846 | 1.723 | 5.116 | 3.4 | 1.423 | 2.778 | 3.938 | 6.566 | 4.116 | 4.306 | 3.809 | 4.346 |
| B | - | - | 4.4 | 3.911 | 4.483 | 4.981 | 3.982 | 3.439 | -0.448 | 7.035 | 0.728 | 2.183 | 1.207 | 2.399 |
| C | - | - | - | 2.327 | 5.161 | 3.123 | 2.145 | 3.1 | 4.077 | 6.509 | 4.234 | 4.409 | 3.96 | 4.439 |
| D | - | - | - | - | 5.019 | 3.836 | -0.638 | 1.873 | 3.637 | 6.667 | 3.859 | 4.088 | 3.468 | 4.142 |
| E | - | - | - | - | - | 6.11 | 5.607 | 5.397 | 4.872 | 7.629 | 4.724 | 4.489 | 4.98 | 4.471 |
| F | - | - | - | - | - | - | 3.596 | 3.989 | 4.555 | 6.232 | 4.662 | 4.784 | 4.481 | 4.802 |
| G | - | - | - | - | - | - | - | 2.103 | 3.701 | 6.646 | 3.913 | 4.134 | 3.543 | 4.18 |
| H | - | - | - | - | - | - | - | - | 3.186 | 6.776 | 3.488 | 3.793 | 2.928 | 3.862 |
| I | - | - | - | - | - | - | - | - | - | 7.032 | 1.242 | 2.384 | 0.658 | 2.561 |
| J | - | - | - | - | - | - | - | - | - | - | 5.85 | 5.885 | 5.803 | 5.89 |
| K | - | - | - | - | - | - | - | - | - | - | - | 1.553 | 2.004 | 1.852 |
| L | - | - | - | - | - | - | - | - | - | - | - | - | 2.845 | -0.397 |
| M | - | - | - | - | - | - | - | - | - | - | - | - | - | 2.878 |
| N | - | - | - | - | - | - | - | - | - | - | - | - | - | - |
