## Supplementary Data S2 for "Interlaboratory comparison of standardised metabolomics and lipidomics analyses in human and rodent blood using the MxP® Quant 500 kit": TG(18.3_38.5).pdf

**Amount of relative differences between laboratories**

| Min | A | B | C | D | E | F | G | H | I | J | K | L | M | N |
|---|---|---|---|---|---|---|---|---|---|---|---|---|---|---|
| A | - | 1.991 | -0.612 | 3.431 | 5.923 | 6.025 | 4.922 | 5.448 | 3.617 | 5.717 | 5.66 | 5.086 | 5.146 | 4.131 |
| B | - | - | 1.558 | 2.961 | 5.911 | 5.984 | 4.797 | 5.416 | 4.016 | 5.726 | 5.584 | 5.068 | 5.049 | 4.53 |
| C | - | - | - | 3.244 | 5.872 | 6.026 | 4.867 | 5.454 | 3.886 | 5.7 | 5.723 | 5.029 | 5.088 | 4.06 |
| D | - | - | - | - | 5.801 | 5.927 | 4.562 | 5.237 | 4.682 | 5.566 | 5.563 | 4.817 | 4.772 | 4.813 |
| E | - | - | - | - | - | 3.01 | 5.974 | 4.969 | 7.288 | 4.894 | 4.512 | 5.838 | 5.554 | 7.237 |
| F | - | - | - | - | - | - | 6.062 | 5.146 | 7.249 | 5.76 | 5.308 | 6.028 | 5.385 | 7.087 |
| G | - | - | - | - | - | - | - | 4.552 | 5.924 | 4.958 | 5.387 | 2.688 | 2.85 | 5.976 |
| H | - | - | - | - | - | - | - | - | 6.486 | 2.741 | -0.61 | 4.483 | 2.89 | 6.391 |
| I | - | - | - | - | - | - | - | - | - | 6.198 | 6.074 | 5.54 | 5.38 | 3.28 |
| J | - | - | - | - | - | - | - | - | - | - | 3.113 | 3.011 | 2.719 | 5.742 |
| K | - | - | - | - | - | - | - | - | - | - | - | 5.112 | 3.601 | 6.652 |
| L | - | - | - | - | - | - | - | - | - | - | - | - | 1.737 | 6.091 |
| M | - | - | - | - | - | - | - | - | - | - | - | - | - | 6.254 |
| N | - | - | - | - | - | - | - | - | - | - | - | - | - | - |

| Median | A | B | C | D | E | F | G | H | I | J | K | L | M | N |
|---|---|---|---|---|---|---|---|---|---|---|---|---|---|---|
| A | - | 2.482 | 0.527 | 3.378 | 5.594 | 5.381 | 4.683 | 4.491 | 1.731 | 0.704 | 4.828 | 4.791 | 5.05 | 3.777 |
| B | - | - | 2.761 | 2.503 | 5.499 | 5.243 | 4.392 | 4.167 | 1.214 | 2.098 | 4.561 | 4.543 | 4.874 | 4.338 |
| C | - | - | - | 3.522 | 5.624 | 5.433 | 4.786 | 4.626 | 2.314 | 2.175 | 4.955 | 4.892 | 5.09 | 3.52 |
| D | - | - | - | - | 5.406 | 5.128 | 4.136 | 3.798 | 2.949 | 3.227 | 4.381 | 4.31 | 4.662 | 4.648 |
| E | - | - | - | - | - | 4.166 | 5.375 | 5.6 | 6.478 | 6.471 | 5.177 | 5.179 | 5.02 | 6.977 |
| F | - | - | - | - | - | - | 3.602 | 4.26 | 5.783 | 5.694 | 3.169 | 2.663 | 3.255 | 6.534 |
| G | - | - | - | - | - | - | - | 1.716 | 4.971 | 4.996 | 2.713 | 1.708 | 3.105 | 5.77 |
| H | - | - | - | - | - | - | - | - | 4.829 | 4.766 | 2.995 | 3.201 | 3.298 | 5.843 |
| I | - | - | - | - | - | - | - | - | - | 0.909 | 4.68 | 4.541 | 4.968 | 3.874 |
| J | - | - | - | - | - | - | - | - | - | - | 4.809 | 4.896 | 4.788 | 4.708 |
| K | - | - | - | - | - | - | - | - | - | - | - | -0.272 | 1.312 | 6.19 |
| L | - | - | - | - | - | - | - | - | - | - | - | - | 2.474 | 5.888 |
| M | - | - | - | - | - | - | - | - | - | - | - | - | - | 6.103 |
| N | - | - | - | - | - | - | - | - | - | - | - | - | - | - |

| Max | A | B | C | D | E | F | G | H | I | J | K | L | M | N |
|---|---|---|---|---|---|---|---|---|---|---|---|---|---|---|
| A | - | 2.892 | 2.626 | 3.27 | 5.208 | 4.174 | 4.457 | 3.455 | 4.03 | 6.169 | 2.632 | 4.555 | 4.888 | 2.215 |
| B | - | - | 3.948 | 0.682 | 4.98 | 4.743 | 3.958 | 4.399 | 3.21 | 6.44 | -0.412 | 4.041 | 4.558 | 3.737 |
| C | - | - | - | 3.941 | 5.353 | 3.67 | 4.718 | 2.472 | 4.424 | 6 | 3.2 | 4.793 | 5.094 | -0.076 |
| D | - | - | - | - | 4.915 | 4.949 | 3.671 | 4.566 | 2.861 | 6.511 | 2.495 | 3.833 | 4.475 | 4.157 |
| E | - | - | - | - | - | 6.395 | 4.608 | 6.241 | 5.047 | 7.441 | 5.631 | 4.473 | 3.349 | 6.061 |
| F | - | - | - | - | - | - | 4.987 | 2.705 | 4.701 | 5.593 | 4.437 | 5.046 | 5.315 | 3.388 |
| G | - | - | - | - | - | - | - | 5.509 | 2.776 | 6.94 | 4.846 | 0.194 | 3.425 | 5.11 |
| H | - | - | - | - | - | - | - | - | 4.507 | 5.829 | 4.054 | 4.921 | 5.18 | 2.513 |
| I | - | - | - | - | - | - | - | - | - | 6.788 | 4.012 | 2.952 | 3.951 | 4.725 |
| J | - | - | - | - | - | - | - | - | - | - | 5.568 | 5.783 | 5.903 | 5.287 |
| K | - | - | - | - | - | - | - | - | - | - | - | 4.443 | 4.637 | 2.878 |
| L | - | - | - | - | - | - | - | - | - | - | - | - | 3.164 | 5.245 |
| M | - | - | - | - | - | - | - | - | - | - | - | - | - | 5.666 |
| N | - | - | - | - | - | - | - | - | - | - | - | - | - | - |

| | |
|---|---|
| | Larger than 10 |
| | Between 8 and 9,99 |
| | Between 6 and 7,99 |
| | Between 4 and 5,99 |
| | Between 2 and 3,99 |
| | Between 0 and 1,99 |
| | NA (Calculation failed) |
