## Supplementary Data S2 for "Interlaboratory comparison of standardised metabolomics and lipidomics analyses in human and rodent blood using the MxP® Quant 500 kit": TG(20.0_34.1).pdf

**Amount of relative differences between laboratories**

| Min | A | B | C | D | E | F | G | H | I | J | K | L | M | N |
|---|---|---|---|---|---|---|---|---|---|---|---|---|---|---|
| A | - | 6.142 | NA | 6.505 | 7.184 | 6.088 | 5.036 | 4.909 | 4.479 | 8.113 | 3.573 | 5.144 | 4.318 | 5.401 |
| B | - | - | NA | 5.304 | 3.95 | 3.902 | 6.056 | 6.438 | 7.486 | 7.004 | 7.009 | 6.494 | 3.121 | 6.376 |
| C | - | - | - | NA | NA | NA | NA | NA | NA | NA | NA | NA | NA | NA |
| D | - | - | - | - | 3.201 | 5.232 | 4.621 | 5.359 | 6.739 | 4.637 | 6.233 | 5.452 | 2.735 | 5.25 |
| E | - | - | - | - | - | 5.403 | 3.355 | 4.667 | 6.369 | 4.097 | 5.809 | 4.818 | 3.923 | 4.532 |
| F | - | - | - | - | - | - | 6.533 | 6.822 | 7.815 | 7.04 | 7.363 | 6.844 | 4.738 | 6.749 |
| G | - | - | - | - | - | - | - | 2.959 | 6.244 | 7.872 | 5.114 | 1.854 | -3.102 | 2.106 |
| H | - | - | - | - | - | - | - | - | 5.93 | 7.602 | 4.631 | 1.595 | 3.712 | 3.29 |
| I | - | - | - | - | - | - | - | - | - | 8.098 | 4.98 | 5.688 | 4.849 | 5.859 |
| J | - | - | - | - | - | - | - | - | - | - | 6.526 | 5.869 | 0.945 | 5.702 |
| K | - | - | - | - | - | - | - | - | - | - | - | 4.381 | 5.035 | 4.671 |
| L | - | - | - | - | - | - | - | - | - | - | - | - | 1.79 | 3.049 |
| M | - | - | - | - | - | - | - | - | - | - | - | - | - | 6.109 |
| N | - | - | - | - | - | - | - | - | - | - | - | - | - | - |

| | |
|---|---|
| | Larger than 10 |
| | Between 8 and 9,99 |
| | Between 6 and 7,99 |
| | Between 4 and 5,99 |
| | Between 2 and 3,99 |
| | Between 0 and 1,99 |
| | NA (Calculation failed) |

| Median | A | B | C | D | E | F | G | H | I | J | K | L | M | N |
|---|---|---|---|---|---|---|---|---|---|---|---|---|---|---|
| A | - | 4.719 | NA | 2.026 | 2.735 | 5.307 | 4.386 | 3.183 | 3.397 | 3.686 | 4.874 | 4.544 | 2.726 | 4.367 |
| B | - | - | NA | 5.336 | 5.656 | 4.088 | 4.527 | 5.119 | 5.883 | 1.999 | 6.317 | 4.03 | 3.805 | 3.551 |
| C | - | - | - | NA | NA | NA | NA | NA | NA | NA | NA | NA | NA | NA |
| D | - | - | - | - | 3.963 | 5.339 | 3.29 | -0.054 | 3.937 | 2.555 | 5.035 | 3.856 | 2.434 | 4.211 |
| E | - | - | - | - | - | 5.571 | 4.432 | 3.81 | -2.463 | 4.162 | 3.904 | 4.682 | 3.931 | 4.884 |
| F | - | - | - | - | - | - | 5.521 | 5.893 | 6.485 | 4.927 | 6.847 | 5.209 | 5.111 | 5.033 |
| G | - | - | - | - | - | - | - | 3.757 | 5.185 | 3.123 | 5.879 | 1.883 | 3.661 | 1.315 |
| H | - | - | - | - | - | - | - | - | 4.308 | 2.018 | 5.296 | 3.907 | 1.446 | 3.856 |
| I | - | - | - | - | - | - | - | - | - | 4.318 | 4.11 | 4.97 | 3.919 | 4.807 |
| J | - | - | - | - | - | - | - | - | - | - | 5.341 | 3.158 | 0.676 | 3.788 |
| K | - | - | - | - | - | - | - | - | - | - | - | 5.253 | 4.772 | 5.271 |
| L | - | - | - | - | - | - | - | - | - | - | - | - | 4.176 | 1.009 |
| M | - | - | - | - | - | - | - | - | - | - | - | - | - | 3.173 |
| N | - | - | - | - | - | - | - | - | - | - | - | - | - | - |

| Max | A | B | C | D | E | F | G | H | I | J | K | L | M | N |
|---|---|---|---|---|---|---|---|---|---|---|---|---|---|---|
| A | - | 4.669 | NA | 6.305 | 6.864 | -0.838 | 2.896 | 3.305 | 1.64 | 6.316 | 6.085 | 3.938 | 6.486 | 3.147 |
| B | - | - | NA | 5.366 | 6.241 | 4.309 | 4.71 | 3.615 | 4.062 | 5.655 | 5.081 | 5.032 | 5.704 | 4.763 |
| C | - | - | - | NA | NA | NA | NA | NA | NA | NA | NA | NA | NA | NA |
| D | - | - | - | - | 4.337 | 5.475 | 5.606 | 5.284 | 5.404 | -0.346 | 2.787 | 5.734 | 1.878 | 5.625 |
| E | - | - | - | - | - | 5.773 | 5.848 | 5.635 | 5.711 | 4.183 | 4.553 | 5.942 | 3.988 | 5.863 |
| F | - | - | - | - | - | - | 2.784 | 3.439 | 2.056 | 6.447 | 6.106 | 3.887 | 6.514 | 3.004 |
| G | - | - | - | - | - | - | - | 4.226 | 3.53 | 6.553 | 6.349 | 3.095 | 6.713 | 0.582 |
| H | - | - | - | - | - | - | - | - | 2.613 | 5.969 | 5.725 | 4.522 | 6.176 | 4.091 |
| I | - | - | - | - | - | - | - | - | - | 6.111 | 5.976 | 4.16 | 6.389 | 3.539 |
| J | - | - | - | - | - | - | - | - | - | - | 2.957 | 5.747 | -0.386 | 5.635 |
| K | - | - | - | - | - | - | - | - | - | - | - | 5.634 | 3.601 | 5.516 |
| L | - | - | - | - | - | - | - | - | - | - | - | - | 6.951 | 2.93 |
| M | - | - | - | - | - | - | - | - | - | - | - | - | - | 5.688 |
| N | - | - | - | - | - | - | - | - | - | - | - | - | - | - |
