## Supplementary Data S2 for "Interlaboratory comparison of standardised metabolomics and lipidomics analyses in human and rodent blood using the MxP® Quant 500 kit": TG(20.1_30.1).pdf

**Amount of relative differences between laboratories**

| Min | A | B | C | D | E | F | G | H | I | J | K | L | M | N |
|---|---|---|---|---|---|---|---|---|---|---|---|---|---|---|
| A | - | 5.668 | 5.585 | 5.203 | 4.664 | NA | 3.845 | 3.94 | 5.465 | NA | NA | 5.821 | NA | NA |
| B | - | - | 3.766 | 2.807 | 5.828 | NA | 4.545 | 4.068 | 4.686 | NA | NA | 4.923 | NA | NA |
| C | - | - | - | 4.146 | 4.294 | NA | 4.173 | 3.523 | 3.937 | NA | NA | 4.424 | NA | NA |
| D | - | - | - | - | 6.552 | NA | 5.155 | 5.019 | 3.603 | NA | NA | 0.208 | NA | NA |
| E | - | - | - | - | - | NA | 4.862 | 4.969 | 4.235 | NA | NA | 4.687 | NA | NA |
| F | - | - | - | - | - | - | NA | NA | NA | NA | NA | NA | NA | NA |
| G | - | - | - | - | - | - | - | 2.302 | 4.027 | NA | NA | 4.157 | NA | NA |
| H | - | - | - | - | - | - | - | - | -0.19 | NA | NA | -0.744 | NA | NA |
| I | - | - | - | - | - | - | - | - | - | NA | NA | 2.478 | NA | NA |
| J | - | - | - | - | - | - | - | - | - | - | NA | NA | NA | NA |
| K | - | - | - | - | - | - | - | - | - | - | - | NA | NA | NA |
| L | - | - | - | - | - | - | - | - | - | - | - | - | NA | NA |
| M | - | - | - | - | - | - | - | - | - | - | - | - | - | NA |
| N | - | - | - | - | - | - | - | - | - | - | - | - | - | - |

| | |
|---|---|
| | Larger than 10 |
| | Between 8 and 9,99 |
| | Between 6 and 7,99 |
| | Between 4 and 5,99 |
| | Between 2 and 3,99 |
| | Between 0 and 1,99 |
| | NA (Calculation failed) |

| Median | A | B | C | D | E | F | G | H | I | J | K | L | M | N |
|---|---|---|---|---|---|---|---|---|---|---|---|---|---|---|
| A | - | 4.785 | 4.867 | 4.901 | 5.502 | NA | 3.03 | 3.731 | 4.866 | NA | NA | 5.562 | NA | NA |
| B | - | - | 2.858 | 4.192 | 5.855 | NA | 3.877 | 3.997 | 1.604 | NA | NA | 1.686 | NA | NA |
| C | - | - | - | 4.486 | 3.126 | NA | 3.644 | 3.509 | -1.355 | NA | NA | -0.09 | NA | NA |
| D | - | - | - | - | 6.754 | NA | 5.167 | 5.07 | 3.621 | NA | NA | 2.148 | NA | NA |
| E | - | - | - | - | - | NA | 5.036 | 4.981 | 3.723 | NA | NA | 2.968 | NA | NA |
| F | - | - | - | - | - | - | NA | NA | NA | NA | NA | NA | NA | NA |
| G | - | - | - | - | - | - | - | 1.992 | 4.084 | NA | NA | 4.643 | NA | NA |
| H | - | - | - | - | - | - | - | - | 3.634 | NA | NA | 3.995 | NA | NA |
| I | - | - | - | - | - | - | - | - | - | NA | NA | 3.612 | NA | NA |
| J | - | - | - | - | - | - | - | - | - | - | NA | NA | NA | NA |
| K | - | - | - | - | - | - | - | - | - | - | - | NA | NA | NA |
| L | - | - | - | - | - | - | - | - | - | - | - | - | NA | NA |
| M | - | - | - | - | - | - | - | - | - | - | - | - | - | NA |
| N | - | - | - | - | - | - | - | - | - | - | - | - | - | - |

| Max | A | B | C | D | E | F | G | H | I | J | K | L | M | N |
|---|---|---|---|---|---|---|---|---|---|---|---|---|---|---|
| A | - | 4.262 | 3.518 | 4.361 | 6.496 | NA | 3.116 | 4.453 | 3.411 | NA | NA | 4.981 | NA | NA |
| B | - | - | 2.695 | 5.06 | 5.915 | NA | 3.137 | 1.207 | 4.645 | NA | NA | 5.415 | NA | NA |
| C | - | - | - | 4.844 | 6.182 | NA | 1.343 | 3.253 | 4.308 | NA | NA | 5.28 | NA | NA |
| D | - | - | - | - | 7.126 | NA | 5.2 | 5.74 | 3.643 | NA | NA | 3.795 | NA | NA |
| E | - | - | - | - | - | NA | 5.422 | 5.168 | 5.723 | NA | NA | 6.014 | NA | NA |
| F | - | - | - | - | - | - | NA | NA | NA | NA | NA | NA | NA | NA |
| G | - | - | - | - | - | - | - | 3.608 | 4.151 | NA | NA | 5.213 | NA | NA |
| H | - | - | - | - | - | - | - | - | 4.737 | NA | NA | 5.458 | NA | NA |
| I | - | - | - | - | - | - | - | - | - | NA | NA | 4.554 | NA | NA |
| J | - | - | - | - | - | - | - | - | - | - | NA | NA | NA | NA |
| K | - | - | - | - | - | - | - | - | - | - | - | NA | NA | NA |
| L | - | - | - | - | - | - | - | - | - | - | - | - | NA | NA |
| M | - | - | - | - | - | - | - | - | - | - | - | - | - | NA |
| N | - | - | - | - | - | - | - | - | - | - | - | - | - | - |
