## Supplementary Data S2 for "Interlaboratory comparison of standardised metabolomics and lipidomics analyses in human and rodent blood using the MxP® Quant 500 kit": TG(20.1_32.1).pdf

**Amount of relative differences between laboratories**

| Min | A | B | C | D | E | F | G | H | I | J | K | L | M | N |
|---|---|---|---|---|---|---|---|---|---|---|---|---|---|---|
| A | - | 6.225 | 5.039 | 5.919 | NA | 5.785 | 5.107 | 5.216 | 4.356 | 8.686 | 4.303 | 5.641 | 5.699 | 5.373 |
| B | - | - | 1.431 | 2.177 | NA | 4.755 | 2.053 | 4.725 | 5.951 | 8.228 | 4.713 | -3.429 | 3.983 | 4.145 |
| C | - | - | - | 3.455 | NA | 4.645 | 1.956 | 5.084 | 6.269 | 8.381 | 4.564 | 1.947 | 4.622 | 4.428 |
| D | - | - | - | - | NA | 4.899 | -2.058 | 4.443 | 5.802 | 8.25 | 4.639 | 1.861 | 3.454 | 3.75 |
| E | - | - | - | - | - | NA | NA | NA | NA | NA | NA | NA | NA | NA |
| F | - | - | - | - | - | - | 5.354 | 6.649 | 7.3 | 7.671 | 5.974 | 5.862 | 6.545 | 6.451 |
| G | - | - | - | - | - | - | - | 4.202 | 5.681 | 8.654 | 4.41 | 2.609 | 1.915 | 3.458 |
| H | - | - | - | - | - | - | - | - | 4.805 | 8.213 | 1.637 | 4.379 | 3.31 | 2.755 |
| I | - | - | - | - | - | - | - | - | - | 8.845 | 0.146 | 5.22 | 5.248 | 4.733 |
| J | - | - | - | - | - | - | - | - | - | - | 5.295 | 5.002 | 6.132 | 5.812 |
| K | - | - | - | - | - | - | - | - | - | - | - | 4.345 | 2.656 | 2.77 |
| L | - | - | - | - | - | - | - | - | - | - | - | - | 3.825 | 4.168 |
| M | - | - | - | - | - | - | - | - | - | - | - | - | - | 2.718 |
| N | - | - | - | - | - | - | - | - | - | - | - | - | - | - |

| Median | A | B | C | D | E | F | G | H | I | J | K | L | M | N |
|---|---|---|---|---|---|---|---|---|---|---|---|---|---|---|
| A | - | 4.596 | 4.241 | 4.617 | NA | 5.364 | 4.778 | 3.055 | 4.024 | 3.94 | 2.337 | 4.826 | 4.674 | 3.812 |
| B | - | - | 2.623 | 1.322 | NA | 4.302 | 0.243 | 4.316 | 3.484 | 3.673 | 4.81 | 1.913 | 1.959 | 3.554 |
| C | - | - | - | 1.178 | NA | 4.41 | 1.015 | 3.933 | 3.206 | 4.269 | 4.214 | 2.965 | 3.084 | 2.534 |
| D | - | - | - | - | NA | 4.47 | 0.984 | 4.101 | 3.1 | 3.314 | 4.743 | 2.606 | 2.598 | 3.21 |
| E | - | - | - | - | - | NA | NA | NA | NA | NA | NA | NA | NA | NA |
| F | - | - | - | - | - | - | 4.786 | 5.665 | 5.37 | 4.934 | 5.455 | 4.547 | 4.25 | 5.332 |
| G | - | - | - | - | - | - | - | 4.422 | 3.611 | 0.788 | 4.938 | 1.611 | 1.614 | 3.768 |
| H | - | - | - | - | - | - | - | - | 2.899 | 2.793 | 3.03 | 4.327 | 4.299 | 2.738 |
| I | - | - | - | - | - | - | - | - | - | -3.366 | 4.717 | 3.846 | 3.499 | 1.267 |
| J | - | - | - | - | - | - | - | - | - | - | -1.238 | 3.751 | 4.213 | 2.054 |
| K | - | - | - | - | - | - | - | - | - | - | - | 4.621 | 4.616 | 3.687 |
| L | - | - | - | - | - | - | - | - | - | - | - | - | -1.406 | 4.064 |
| M | - | - | - | - | - | - | - | - | - | - | - | - | - | 4.001 |
| N | - | - | - | - | - | - | - | - | - | - | - | - | - | - |

| Max | A | B | C | D | E | F | G | H | I | J | K | L | M | N |
|---|---|---|---|---|---|---|---|---|---|---|---|---|---|---|
| A | - | 1.55 | 4.269 | 1.952 | NA | 3.398 | 2.408 | 4.13 | 3.804 | 7.24 | 5.098 | 2.363 | 3.541 | 3.283 |
| B | - | - | 3.968 | -0.147 | NA | 2.857 | 3.028 | 3.813 | 4.044 | 7.17 | 4.916 | 2.986 | 3.832 | 2.719 |
| C | - | - | - | 3.668 | NA | 2.876 | 4.365 | 0.487 | 4.8 | 6.815 | 3.65 | 4.349 | 4.693 | 2.977 |
| D | - | - | - | - | NA | 2.657 | 3.164 | 3.706 | 4.108 | 7.15 | 4.857 | 3.127 | 3.906 | 2.487 |
| E | - | - | - | - | - | NA | NA | NA | NA | NA | NA | NA | NA | NA |
| F | - | - | - | - | - | - | 3.841 | 2.666 | 4.47 | 7.011 | 4.424 | 3.821 | 4.324 | -0.511 |
| G | - | - | - | - | - | - | - | 4.594 | 3.19 | 7.381 | 5.383 | -2.139 | 2.733 | 3.993 |
| H | - | - | - | - | - | - | - | - | 4.744 | 6.843 | 3.82 | 4.268 | 4.633 | 2.723 |
| I | - | - | - | - | - | - | - | - | - | 7.594 | 5.807 | 3.363 | 1.442 | 4.786 |
| J | - | - | - | - | - | - | - | - | - | - | 5.56 | 5.966 | 6.025 | 5.821 |
| K | - | - | - | - | - | - | - | - | - | - | - | 4.869 | 5.092 | 4.181 |
| L | - | - | - | - | - | - | - | - | - | - | - | - | 2.781 | 3.968 |
| M | - | - | - | - | - | - | - | - | - | - | - | - | - | 4.597 |
| N | - | - | - | - | - | - | - | - | - | - | - | - | - | - |

| Colour | Legend |
|---|---|
| (red) | Larger than 10 |
| (orange) | Between 8 and 9,99 |
| (light orange) | Between 6 and 7,99 |
| (light green) | Between 4 and 5,99 |
| (green) | Between 2 and 3,99 |
| (dark green) | Between 0 and 1,99 |
| (grey) | NA (Calculation failed) |
