## Supplementary Data S2 for "Interlaboratory comparison of standardised metabolomics and lipidomics analyses in human and rodent blood using the MxP® Quant 500 kit": TG(20.1_34.0).pdf

**Amount of relative differences between laboratories**

| Min | A | B | C | D | E | F | G | H | I | J | K | L | M | N |
|---|---|---|---|---|---|---|---|---|---|---|---|---|---|---|
| A | - | 5.469 | -0.516 | 6.71 | NA | 5.697 | 4.21 | 3.972 | 1.236 | 7.73 | 4.308 | 4.562 | 5.806 | 4.357 |
| B | - | - | 5.4 | 5.576 | NA | 2.424 | 5.53 | 5.256 | 6.298 | 7.037 | 6.73 | 5.414 | 3.65 | 5.444 |
| C | - | - | - | 5.899 | NA | 4.95 | 3.776 | 2.735 | 5.242 | 7.072 | 5.795 | 3.307 | 5.044 | 3.149 |
| D | - | - | - | - | NA | -0.34 | 5.52 | 5.12 | 6.226 | 3.129 | 6.519 | 5.524 | 2.575 | 5.458 |
| E | - | - | - | - | - | NA | NA | NA | NA | NA | NA | NA | NA | NA |
| F | - | - | - | - | - | - | 5.809 | 5.06 | 6.441 | 3.299 | 6.754 | 5.727 | -0.033 | 5.67 |
| G | - | - | - | - | - | - | - | 0.825 | 4.569 | 7.447 | 5.395 | 2.145 | 5.418 | 1.078 |
| H | - | - | - | - | - | - | - | - | 4.756 | 7.239 | 5.609 | -1.513 | 5.275 | 0.788 |
| I | - | - | - | - | - | - | - | - | - | 7.502 | 3.983 | 4.576 | 5.786 | 4.427 |
| J | - | - | - | - | - | - | - | - | - | - | 5.899 | 4.543 | 4.304 | 4.416 |
| K | - | - | - | - | - | - | - | - | - | - | - | 4.964 | 5.864 | 4.91 |
| L | - | - | - | - | - | - | - | - | - | - | - | - | 5.353 | 1.519 |
| M | - | - | - | - | - | - | - | - | - | - | - | - | - | 5.923 |
| N | - | - | - | - | - | - | - | - | - | - | - | - | - | - |

| Median | A | B | C | D | E | F | G | H | I | J | K | L | M | N |
|---|---|---|---|---|---|---|---|---|---|---|---|---|---|---|
| A | - | 4.606 | 3.405 | 4.817 | NA | 1.011 | 2.805 | -0.899 | 3.18 | 2.04 | 5.228 | 3.859 | 5.016 | 2.148 |
| B | - | - | 5.511 | 2.241 | NA | 4.865 | 4.621 | 5.153 | 5.562 | 5.246 | 6.48 | 3.716 | 3.082 | 5.173 |
| C | - | - | - | 4.954 | NA | 3.799 | 3.476 | 0.937 | 2.378 | 4.636 | 5.099 | 4.366 | 5.173 | 2.471 |
| D | - | - | - | - | NA | -5.174 | -0.331 | 3.725 | 4.288 | 4.657 | 5.673 | 3.219 | 4.589 | 2.873 |
| E | - | - | - | - | - | NA | NA | NA | NA | NA | NA | NA | NA | NA |
| F | - | - | - | - | - | - | 2.726 | 3.651 | 4.728 | 4.714 | 5.953 | 2.495 | 4.372 | 3.54 |
| G | - | - | - | - | - | - | - | 3.358 | 4.131 | 3.883 | 5.592 | 3.098 | 4.741 | 3.46 |
| H | - | - | - | - | - | - | - | - | 2.951 | 3.028 | 5.303 | 4.129 | 5.05 | -4.916 |
| I | - | - | - | - | - | - | - | - | - | 1.682 | 4.706 | 4.454 | 5.241 | 2.305 |
| J | - | - | - | - | - | - | - | - | - | - | 4.275 | 4.856 | 5.318 | 4.021 |
| K | - | - | - | - | - | - | - | - | - | - | - | 5.268 | 5.658 | 4.679 |
| L | - | - | - | - | - | - | - | - | - | - | - | - | 4.34 | 4.435 |
| M | - | - | - | - | - | - | - | - | - | - | - | - | - | 5.547 |
| N | - | - | - | - | - | - | - | - | - | - | - | - | - | - |

| Max | A | B | C | D | E | F | G | H | I | J | K | L | M | N |
|---|---|---|---|---|---|---|---|---|---|---|---|---|---|---|
| A | - | 2.752 | 6.262 | 6.885 | NA | 6.846 | 2.662 | 5.256 | 4.093 | 8.047 | 6.118 | 2.723 | 2.145 | 5.142 |
| B | - | - | 6.035 | 6.714 | NA | 6.665 | -1.454 | 4.883 | 3.278 | 7.916 | 5.876 | 3.642 | 1.079 | 4.744 |
| C | - | - | - | 4.575 | NA | 4.438 | 5.316 | 4.447 | 5.077 | 6.742 | 2.044 | 5.559 | 5.354 | 4.549 |
| D | - | - | - | - | NA | 0.651 | 5.682 | 5.202 | 5.537 | 6.047 | 4.493 | 5.843 | 5.708 | 5.251 |
| E | - | - | - | - | - | NA | NA | NA | NA | NA | NA | NA | NA | NA |
| F | - | - | - | - | - | - | 5.659 | 5.157 | 5.51 | 6.122 | 4.408 | 5.822 | 5.685 | 5.211 |
| G | - | - | - | - | - | - | - | 4.904 | 3.337 | 7.921 | 5.891 | 3.605 | 0.837 | 4.769 |
| H | - | - | - | - | - | - | - | - | 3.933 | 7.356 | 4.497 | 5.018 | 4.611 | 1.046 |
| I | - | - | - | - | - | - | - | - | - | 7.72 | 5.483 | 4.339 | 3.431 | 3.962 |
| J | - | - | - | - | - | - | - | - | - | - | 5.743 | 6.217 | 6.156 | 5.973 |
| K | - | - | - | - | - | - | - | - | - | - | - | 5.488 | 5.262 | 4.331 |
| L | - | - | - | - | - | - | - | - | - | - | - | - | 3.569 | 5.488 |
| M | - | - | - | - | - | - | - | - | - | - | - | - | - | 4.885 |
| N | - | - | - | - | - | - | - | - | - | - | - | - | - | - |

| Colour | Legend |
|---|---|
| Red | Larger than 10 |
| Orange | Between 8 and 9,99 |
| Light orange | Between 6 and 7,99 |
| Light green | Between 4 and 5,99 |
| Green | Between 2 and 3,99 |
| Dark green | Between 0 and 1,99 |
| Grey | NA (Calculation failed) |
