## Supplementary Data S2 for "Interlaboratory comparison of standardised metabolomics and lipidomics analyses in human and rodent blood using the MxP® Quant 500 kit": TG(20.2_34.2).pdf

Amount of relative differences between laboratories

| Min | A | B | C | D | E | F | G | H | I | J | K | L | M | N |
|---|---|---|---|---|---|---|---|---|---|---|---|---|---|---|
| A | - | 5.095 | 5.2 | 5.063 | 6.047 | 6.146 | 5.105 | 5.255 | 3.458 | 7.385 | 4.86 | 5.057 | 5.103 | 4.962 |
| B | - | - | 2.529 | -1.317 | 5.402 | 5.564 | 0.392 | 2.748 | 4.843 | 7.191 | 2.709 | 1.826 | 1.985 | 2.908 |
| C | - | - | - | 2.943 | 4.984 | 5.293 | 2.203 | -2.319 | 5.094 | 6.887 | 3.733 | 3.646 | -0.874 | 3.998 |
| D | - | - | - | - | 5.306 | 5.471 | 1.189 | 2.855 | 4.729 | 6.914 | 2.554 | 1.904 | 2.444 | 2.992 |
| E | - | - | - | - | - | 3.021 | 5.617 | 5.406 | 6.498 | 6.404 | 5.96 | 5.99 | 5.436 | 6.083 |
| F | - | - | - | - | - | - | 5.757 | 5.57 | 6.56 | 5.552 | 6.065 | 6.097 | 5.517 | 6.235 |
| G | - | - | - | - | - | - | - | 2.328 | 4.877 | 6.999 | 3.037 | 2.608 | 1.745 | 3.326 |
| H | - | - | - | - | - | - | - | - | 5.17 | 6.8 | 3.783 | 3.656 | 0.66 | 4.06 |
| I | - | - | - | - | - | - | - | - | - | 7.759 | 4.439 | 4.614 | 4.705 | 4.702 |
| J | - | - | - | - | - | - | - | - | - | - | 5.989 | 6.097 | 5.423 | 6.19 |
| K | - | - | - | - | - | - | - | - | - | - | - | 1.779 | 3.251 | -2.303 |
| L | - | - | - | - | - | - | - | - | - | - | - | - | 2.914 | 2.024 |
| M | - | - | - | - | - | - | - | - | - | - | - | - | - | 3.469 |
| N | - | - | - | - | - | - | - | - | - | - | - | - | - | - |

| | |
|---|---|
| | Larger than 10 |
| | Between 8 and 9,99 |
| | Between 6 and 7,99 |
| | Between 4 and 5,99 |
| | Between 2 and 3,99 |
| | Between 0 and 1,99 |
| | NA (Calculation failed) |

| Median | A | B | C | D | E | F | G | H | I | J | K | L | M | N |
|---|---|---|---|---|---|---|---|---|---|---|---|---|---|---|
| A | - | 3.902 | 2.842 | 2.431 | 4.89 | 4.238 | 3.167 | 2.664 | 4.006 | 4.668 | 3.622 | 3.72 | 4.197 | 3.868 |
| B | - | - | 3.16 | 3.482 | 4.086 | 2.129 | 2.834 | 3.195 | 0.841 | 5.449 | 1.757 | 1.04 | 1.862 | -0.583 |
| C | - | - | - | 1.171 | 4.515 | 3.578 | 0.643 | -1.839 | 3.334 | 4.893 | 2.266 | 2.617 | 3.494 | 2.886 |
| D | - | - | - | - | 4.647 | 3.788 | 1.871 | 1.163 | 3.563 | 4.816 | 2.782 | 3.044 | 3.719 | 3.228 |
| E | - | - | - | - | - | 3.846 | 4.776 | 4.828 | 4.018 | 6.064 | 4.553 | 4.44 | 3.929 | 4.328 |
| F | - | - | - | - | - | - | 3.507 | 3.47 | 0.175 | 5.386 | 3.012 | 2.56 | -0.067 | 2.332 |
| G | - | - | - | - | - | - | - | 0.636 | 3.084 | 5.042 | 1.757 | 2.246 | 3.313 | 2.58 |
| H | - | - | - | - | - | - | - | - | 3.267 | 4.939 | 2.172 | 2.559 | 3.441 | 2.821 |
| I | - | - | - | - | - | - | - | - | - | 5.578 | 2.073 | 1.88 | 1.489 | 0.439 |
| J | - | - | - | - | - | - | - | - | - | - | 4.234 | 4.384 | 4.558 | 4.426 |
| K | - | - | - | - | - | - | - | - | - | - | - | 0.357 | 2.762 | 1.413 |
| L | - | - | - | - | - | - | - | - | - | - | - | - | 2.513 | 0.418 |
| M | - | - | - | - | - | - | - | - | - | - | - | - | - | 2.11 |
| N | - | - | - | - | - | - | - | - | - | - | - | - | - | - |

| Max | A | B | C | D | E | F | G | H | I | J | K | L | M | N |
|---|---|---|---|---|---|---|---|---|---|---|---|---|---|---|
| A | - | 2.708 | 3.252 | 3.138 | 3.441 | 4.043 | 2.088 | 3.181 | 4.152 | 6.79 | 2.259 | 2.398 | 3.287 | 3.065 |
| B | - | - | 4.107 | 4.056 | 1.987 | 4.635 | 3.526 | 4.073 | 3.586 | 6.997 | 0.969 | 0.556 | 1.791 | 0.932 |
| C | - | - | - | -0.989 | 4.197 | 2.727 | 2.255 | -1.654 | 4.638 | 6.574 | 3.696 | 3.752 | 4.137 | 4.018 |
| D | - | - | - | - | 4.135 | 2.801 | 2.129 | -2.494 | 4.624 | 6.556 | 3.657 | 3.696 | 4.113 | 3.991 |
| E | - | - | - | - | - | 4.892 | 4.012 | 4.434 | 3.111 | 7.066 | 2.591 | 2.391 | -2.125 | 1.072 |
| F | - | - | - | - | - | - | 3.442 | 2.707 | 4.912 | 6.342 | 4.205 | 4.225 | 4.526 | 4.422 |
| G | - | - | - | - | - | - | - | 2.243 | 4.403 | 6.703 | 3.104 | 3.171 | 3.75 | 3.586 |
| H | - | - | - | - | - | - | - | - | 4.639 | 6.548 | 3.67 | 3.719 | 4.131 | 4.001 |
| I | - | - | - | - | - | - | - | - | - | 7.365 | 3.988 | 3.884 | 3.27 | 3.545 |
| J | - | - | - | - | - | - | - | - | - | - | 5.804 | 5.808 | 5.886 | 5.836 |
| K | - | - | - | - | - | - | - | - | - | - | - | -1.203 | 2.411 | 1.904 |
| L | - | - | - | - | - | - | - | - | - | - | - | - | 2.241 | 1.754 |
| M | - | - | - | - | - | - | - | - | - | - | - | - | - | 0.826 |
| N | - | - | - | - | - | - | - | - | - | - | - | - | - | - |
