## Supplementary Data S2 for "Interlaboratory comparison of standardised metabolomics and lipidomics analyses in human and rodent blood using the MxP® Quant 500 kit": TG(20.2_34.4).pdf

**Amount of relative differences between laboratories**

| Min | A | B | C | D | E | F | G | H | I | J | K | L | M | N |
|---|---|---|---|---|---|---|---|---|---|---|---|---|---|---|
| A | - | 6.109 | 5.305 | 5.166 | 5.034 | NA | 4.735 | NA | 4.462 | NA | 6.45 | 3.548 | 5.882 | 5.643 |
| B | - | - | 4.524 | 3.349 | 5.46 | NA | 6.223 | NA | 5.302 | NA | 5.046 | 7.046 | 3.688 | 5.613 |
| C | - | - | - | 1.591 | 3.855 | NA | 4.945 | NA | 4.658 | NA | 5.888 | 6.452 | 4.482 | 2.515 |
| D | - | - | - | - | 4.581 | NA | 2.407 | NA | 4.438 | NA | 5.966 | 4.48 | 4.975 | 4.759 |
| E | - | - | - | - | - | NA | 3.199 | NA | 4.087 | NA | 6.273 | 6.038 | 5.14 | 4.425 |
| F | - | - | - | - | - | - | NA | NA | NA | NA | NA | NA | NA | NA |
| G | - | - | - | - | - | - | - | NA | 3.275 | NA | 6.328 | 5.562 | 5.169 | 4.824 |
| H | - | - | - | - | - | - | - | - | NA | NA | NA | NA | NA | NA |
| I | - | - | - | - | - | - | - | - | - | NA | 6.445 | 5.011 | 5.805 | 5.69 |
| J | - | - | - | - | - | - | - | - | - | - | NA | NA | NA | NA |
| K | - | - | - | - | - | - | - | - | - | - | - | 8.077 | 5.896 | 6.895 |
| L | - | - | - | - | - | - | - | - | - | - | - | - | 5.833 | 5.963 |
| M | - | - | - | - | - | - | - | - | - | - | - | - | - | 5.549 |
| N | - | - | - | - | - | - | - | - | - | - | - | - | - | - |

| | |
|---|---|
| | Larger than 10 |
| | Between 8 and 9,99 |
| | Between 6 and 7,99 |
| | Between 4 and 5,99 |
| | Between 2 and 3,99 |
| | Between 0 and 1,99 |
| | NA (Calculation failed) |

| Median | A | B | C | D | E | F | G | H | I | J | K | L | M | N |
|---|---|---|---|---|---|---|---|---|---|---|---|---|---|---|
| A | - | 4.743 | 4.939 | 3.732 | 5.146 | NA | 4.612 | NA | 4.635 | NA | 5.804 | 3.69 | 5.161 | 4.714 |
| B | - | - | 2.921 | 3.656 | 0.606 | NA | 2.945 | NA | 3.112 | NA | 5.246 | 4.876 | 3.707 | 3.05 |
| C | - | - | - | 2.622 | 2.861 | NA | 2.082 | NA | 2.971 | NA | 5.258 | 4.575 | 2.962 | 2.135 |
| D | - | - | - | - | 4.844 | NA | 3.718 | NA | 1.828 | NA | 5.414 | 2.441 | 4.033 | 3.801 |
| E | - | - | - | - | - | NA | 4.186 | NA | 4.054 | NA | 4.881 | 5.121 | 1.599 | 4.061 |
| F | - | - | - | - | - | - | NA | NA | NA | NA | NA | NA | NA | NA |
| G | - | - | - | - | - | - | - | NA | 1.484 | NA | 5.369 | 3.862 | 3.321 | 0.723 |
| H | - | - | - | - | - | - | - | - | NA | NA | NA | NA | NA | NA |
| I | - | - | - | - | - | - | - | - | - | NA | 5.28 | 3.658 | 4.165 | 2.27 |
| J | - | - | - | - | - | - | - | - | - | - | NA | NA | NA | NA |
| K | - | - | - | - | - | - | - | - | - | - | - | 6.773 | 5.733 | 6.125 |
| L | - | - | - | - | - | - | - | - | - | - | - | - | 4.225 | 3.502 |
| M | - | - | - | - | - | - | - | - | - | - | - | - | - | 3.872 |
| N | - | - | - | - | - | - | - | - | - | - | - | - | - | - |

| Max | A | B | C | D | E | F | G | H | I | J | K | L | M | N |
|---|---|---|---|---|---|---|---|---|---|---|---|---|---|---|
| A | - | 4.701 | 3.367 | 1.62 | 5.258 | NA | 4.521 | NA | 4.891 | NA | 4.572 | 4.925 | 2.919 | 2.6 |
| B | - | - | 4.801 | 4.518 | 5.666 | NA | 5.248 | NA | 5.475 | NA | 5.424 | 5.446 | 3.741 | 4.717 |
| C | - | - | - | 2.921 | 4.943 | NA | 3.796 | NA | 4.47 | NA | 4.086 | 4.419 | 4.28 | 1.708 |
| D | - | - | - | - | 5.084 | NA | 4.211 | NA | 4.792 | NA | 4.487 | 4.651 | 3.56 | 1.471 |
| E | - | - | - | - | - | NA | 4.607 | NA | 3.996 | NA | 4.266 | 3.686 | 6.221 | 5.619 |
| F | - | - | - | - | - | - | NA | NA | NA | NA | NA | NA | NA | NA |
| G | - | - | - | - | - | - | - | NA | 3.181 | NA | 0.752 | 3.116 | 5.269 | 4.395 |
| H | - | - | - | - | - | - | - | - | NA | NA | NA | NA | NA | NA |
| I | - | - | - | - | - | - | - | - | - | NA | 3.354 | 0.385 | 5.729 | 5.128 |
| J | - | - | - | - | - | - | - | - | - | - | NA | NA | NA | NA |
| K | - | - | - | - | - | - | - | - | - | - | - | 2.216 | 5.369 | 4.806 |
| L | - | - | - | - | - | - | - | - | - | - | - | - | 5.724 | 5.004 |
| M | - | - | - | - | - | - | - | - | - | - | - | - | - | 3.669 |
| N | - | - | - | - | - | - | - | - | - | - | - | - | - | - |
