## Supplementary Data S2 for "Interlaboratory comparison of standardised metabolomics and lipidomics analyses in human and rodent blood using the MxP® Quant 500 kit": TG(20.2_36.5).pdf

**Amount of relative differences between laboratories**

| Min | A | B | C | D | E | F | G | H | I | J | K | L | M | N |
|---|---|---|---|---|---|---|---|---|---|---|---|---|---|---|
| A | - | 4.826 | 4.525 | 5.218 | NA | 6.864 | 5.989 | 5.001 | NA | 3.671 | 6.447 | 5.332 | 4.37 | 4.791 |
| B | - | - | 5.244 | 5.453 | NA | 6.153 | 5.151 | 5.002 | NA | 4.427 | 4.852 | 4.497 | 3.237 | 3.259 |
| C | - | - | - | 2.646 | NA | 6.481 | 5.849 | 5.548 | NA | 3.428 | 5.847 | 5.531 | 5.108 | 5.166 |
| D | - | - | - | - | NA | 6.639 | 6.1 | 5.654 | NA | 4.089 | 6.305 | 5.744 | 5.357 | 5.475 |
| E | - | - | - | - | - | NA | NA | NA | NA | NA | NA | NA | NA | NA |
| F | - | - | - | - | - | - | 4.561 | 3.245 | NA | 6.465 | 6.491 | 4.806 | 5.501 | 5.785 |
| G | - | - | - | - | - | - | - | 3.341 | NA | 5.874 | 5.532 | 2.614 | 4.483 | 4.809 |
| H | - | - | - | - | - | - | - | - | NA | 6.143 | 6.032 | 4.106 | 5.015 | 5.318 |
| I | - | - | - | - | - | - | - | - | - | NA | NA | NA | NA | NA |
| J | - | - | - | - | - | - | - | - | - | - | 4.284 | 5.049 | 4.636 | 4.445 |
| K | - | - | - | - | - | - | - | - | - | - | - | 4.014 | -0.494 | 0.495 |
| L | - | - | - | - | - | - | - | - | - | - | - | - | 3.867 | 4.274 |
| M | - | - | - | - | - | - | - | - | - | - | - | - | - | 1.314 |
| N | - | - | - | - | - | - | - | - | - | - | - | - | - | - |

| | |
|---|---|
| | Larger than 10 |
| | Between 8 and 9,99 |
| | Between 6 and 7,99 |
| | Between 4 and 5,99 |
| | Between 2 and 3,99 |
| | Between 0 and 1,99 |
| | NA (Calculation failed) |

| Median | A | B | C | D | E | F | G | H | I | J | K | L | M | N |
|---|---|---|---|---|---|---|---|---|---|---|---|---|---|---|
| A | - | 4.531 | 5.677 | 5.393 | NA | 4.603 | 5.224 | 4.025 | NA | 6.091 | 5.26 | 3.996 | 1.076 | 1.773 |
| B | - | - | 4.688 | 4.758 | NA | 2.13 | 2.059 | 3.275 | NA | 5.323 | 2.982 | 1.543 | 4.113 | 3.925 |
| C | - | - | - | 0.493 | NA | 4.415 | 3.846 | 4.653 | NA | 3.507 | 3.581 | 4.529 | 5.11 | 5.006 |
| D | - | - | - | - | NA | 3.72 | 2.632 | 4.332 | NA | 4.076 | 2.279 | 4.265 | 4.994 | 4.823 |
| E | - | - | - | - | - | NA | NA | NA | NA | NA | NA | NA | NA | NA |
| F | - | - | - | - | - | - | 2.421 | 1.729 | NA | 6.023 | 4.106 | 3.598 | 0.077 | 1.187 |
| G | - | - | - | - | - | - | - | 2.778 | NA | 5.632 | 3.093 | 1.08 | 3.385 | 3.435 |
| H | - | - | - | - | - | - | - | - | NA | 5.841 | 3.909 | 3.123 | 2.363 | 2.467 |
| I | - | - | - | - | - | - | - | - | - | NA | NA | NA | NA | NA |
| J | - | - | - | - | - | - | - | - | - | - | 4.719 | 4.957 | 5.352 | 5.332 |
| K | - | - | - | - | - | - | - | - | - | - | - | 3.052 | 4.326 | 4.282 |
| L | - | - | - | - | - | - | - | - | - | - | - | - | 3.705 | 3.686 |
| M | - | - | - | - | - | - | - | - | - | - | - | - | - | 1.846 |
| N | - | - | - | - | - | - | - | - | - | - | - | - | - | - |

| Max | A | B | C | D | E | F | G | H | I | J | K | L | M | N |
|---|---|---|---|---|---|---|---|---|---|---|---|---|---|---|
| A | - | 6.057 | 6.42 | 5.706 | NA | 7.651 | 7.289 | 7.294 | NA | 7.719 | 6.865 | 7.116 | 4.384 | 5.03 |
| B | - | - | 3.451 | 3.085 | NA | 6.312 | 5.673 | 5.793 | NA | 6.455 | 4.946 | 5.322 | 4.758 | 4.356 |
| C | - | - | - | 4.145 | NA | 6 | 5.195 | 5.24 | NA | 6.095 | 4.041 | 4.78 | 5.112 | 4.837 |
| D | - | - | - | - | NA | 6.576 | 6.075 | 6.083 | NA | 6.701 | 5.372 | 5.802 | 4.36 | 3.709 |
| E | - | - | - | - | - | NA | NA | NA | NA | NA | NA | NA | NA | NA |
| F | - | - | - | - | - | - | 4.015 | 3.872 | NA | 2.35 | 4.814 | 4.494 | 5.869 | 5.815 |
| G | - | - | - | - | - | - | - | 1.424 | NA | 4.713 | 3.907 | 2.73 | 5.674 | 5.565 |
| H | - | - | - | - | - | - | - | - | NA | 4.523 | 4.037 | 3.3 | 5.688 | 5.592 |
| I | - | - | - | - | - | - | - | - | - | NA | NA | NA | NA | NA |
| J | - | - | - | - | - | - | - | - | - | - | 4.937 | 4.733 | 5.922 | 5.85 |
| K | - | - | - | - | - | - | - | - | - | - | - | 3.148 | 5.46 | 5.288 |
| L | - | - | - | - | - | - | - | - | - | - | - | - | 5.562 | 5.432 |
| M | - | - | - | - | - | - | - | - | - | - | - | - | - | 3.219 |
| N | - | - | - | - | - | - | - | - | - | - | - | - | - | - |
