## Supplementary Data S2 for "Interlaboratory comparison of standardised metabolomics and lipidomics analyses in human and rodent blood using the MxP® Quant 500 kit": TG(20.3_34.2).pdf

**Amount of relative differences between laboratories**

| Min | A | B | C | D | E | F | G | H | I | J | K | L | M | N |
|---|---|---|---|---|---|---|---|---|---|---|---|---|---|---|
| A | - | 5.424 | 5.866 | 5.307 | 6.232 | 5.892 | 5.345 | 5.395 | 4.159 | 7.648 | 5.331 | 5.145 | 5.263 | 5.289 |
| B | - | - | 4.126 | 2.772 | 5.471 | 4.752 | 2.911 | -3.533 | 5.357 | 7.258 | 1.671 | 3.196 | 1.467 | 3.133 |
| C | - | - | - | 4.651 | 4.84 | 3.888 | 4.726 | 3.974 | 5.976 | 6.87 | 4.348 | 4.738 | 4.136 | 4.758 |
| D | - | - | - | - | 5.526 | 4.991 | -0.808 | 2.669 | 4.959 | 7.06 | 1.855 | 0.412 | 2.185 | 0.757 |
| E | - | - | - | - | - | 3.982 | 6.157 | 5.878 | 6.9 | 6.623 | 5.96 | 6.189 | 5.905 | 6.189 |
| F | - | - | - | - | - | - | 5.561 | 5.102 | 6.488 | 6.593 | 5.314 | 5.558 | 5.175 | 5.591 |
| G | - | - | - | - | - | - | - | 2.845 | 4.947 | 7.106 | 1.891 | 0.239 | 2.332 | 0.22 |
| H | - | - | - | - | - | - | - | - | 5.365 | 7.394 | 1.493 | 3.27 | 1.604 | 3.003 |
| I | - | - | - | - | - | - | - | - | - | 7.619 | 4.77 | 4.455 | 4.686 | 4.654 |
| J | - | - | - | - | - | - | - | - | - | - | 5.973 | 6.128 | 5.801 | 6.226 |
| K | - | - | - | - | - | - | - | - | - | - | - | 2.625 | -0.201 | 2.33 |
| L | - | - | - | - | - | - | - | - | - | - | - | - | 2.501 | -1.517 |
| M | - | - | - | - | - | - | - | - | - | - | - | - | - | 2.553 |
| N | - | - | - | - | - | - | - | - | - | - | - | - | - | - |

| | |
|---|---|
| | Larger than 10 |
| | Between 8 and 9,99 |
| | Between 6 and 7,99 |
| | Between 4 and 5,99 |
| | Between 2 and 3,99 |
| | Between 0 and 1,99 |
| | NA (Calculation failed) |

| Median | A | B | C | D | E | F | G | H | I | J | K | L | M | N |
|---|---|---|---|---|---|---|---|---|---|---|---|---|---|---|
| A | - | 3.95 | 3.633 | 2.068 | 5.05 | 4.241 | 2.495 | 4.162 | 3.139 | 3.653 | 3.665 | 3.617 | 3.546 | 3.918 |
| B | - | - | 1.659 | 3.798 | 4.36 | 2.126 | 3.621 | 1.588 | 2.84 | 4.953 | 2.086 | 2.019 | 2.012 | -1.35 |
| C | - | - | - | 3.474 | 4.476 | 2.874 | 3.253 | 2.536 | 1.898 | 4.678 | 1.026 | 0.57 | -0.232 | 1.32 |
| D | - | - | - | - | 4.915 | 4.021 | 0.536 | 3.897 | 2.629 | 3.867 | 3.069 | 3.127 | 3.144 | 3.584 |
| E | - | - | - | - | - | 4.223 | 5.338 | 4.393 | 5.028 | 5.928 | 4.949 | 4.928 | 4.874 | 4.643 |
| F | - | - | - | - | - | - | 4.194 | 0.688 | 3.559 | 5.131 | 3.379 | 3.274 | 3.125 | 2.38 |
| G | - | - | - | - | - | - | - | 3.774 | 2.235 | 4.039 | 2.809 | 2.878 | 2.903 | 3.415 |
| H | - | - | - | - | - | - | - | - | 3.399 | 5.113 | 2.866 | 2.864 | 2.864 | 1.77 |
| I | - | - | - | - | - | - | - | - | - | 4.528 | 1.97 | 1.853 | 1.635 | 2.767 |
| J | - | - | - | - | - | - | - | - | - | - | 3.669 | 3.834 | 4.031 | 4.218 |
| K | - | - | - | - | - | - | - | - | - | - | - | -2.861 | -1.704 | 1.922 |
| L | - | - | - | - | - | - | - | - | - | - | - | - | -3.314 | 1.761 |
| M | - | - | - | - | - | - | - | - | - | - | - | - | - | 1.811 |
| N | - | - | - | - | - | - | - | - | - | - | - | - | - | - |

| Max | A | B | C | D | E | F | G | H | I | J | K | L | M | N |
|---|---|---|---|---|---|---|---|---|---|---|---|---|---|---|
| A | - | 2.001 | 2.95 | 3.622 | 3.806 | 2.074 | 3.308 | 2.989 | 2.642 | 6.538 | -1.021 | 1.296 | -1.098 | 2.705 |
| B | - | - | 3.608 | 4.105 | 3.274 | 3.094 | 3.851 | 2.074 | 1.276 | 6.649 | 2.249 | 0.913 | 2.206 | 1.384 |
| C | - | - | - | 2.194 | 4.274 | 1.616 | 1.009 | 3.875 | 3.707 | 6.31 | 2.75 | 3.203 | 2.756 | 3.71 |
| D | - | - | - | - | 4.518 | 2.878 | 1.231 | 4.203 | 4.066 | 6.137 | 3.416 | 3.702 | 3.432 | 4.079 |
| E | - | - | - | - | - | 4.344 | 4.739 | 2.439 | 3.004 | 6.919 | 3.992 | 3.631 | 3.946 | 2.966 |
| F | - | - | - | - | - | - | 2.454 | 3.572 | 3.367 | 6.403 | 1.862 | 2.635 | 1.901 | 3.35 |
| G | - | - | - | - | - | - | - | 4.031 | 3.885 | 6.22 | 3.102 | 3.456 | 3.103 | 3.898 |
| H | - | - | - | - | - | - | - | - | 0.889 | 6.795 | 3.237 | 2.649 | 3.203 | 0.816 |
| I | - | - | - | - | - | - | - | - | - | 6.727 | 2.859 | 2.057 | 2.845 | -2.114 |
| J | - | - | - | - | - | - | - | - | - | - | 5.61 | 5.644 | 5.618 | 5.703 |
| K | - | - | - | - | - | - | - | - | - | - | - | 1.492 | -4.292 | 2.797 |
| L | - | - | - | - | - | - | - | - | - | - | - | - | 1.444 | 2.163 |
| M | - | - | - | - | - | - | - | - | - | - | - | - | - | 2.78 |
| N | - | - | - | - | - | - | - | - | - | - | - | - | - | - |
