## Supplementary Data S2 for "Interlaboratory comparison of standardised metabolomics and lipidomics analyses in human and rodent blood using the MxP® Quant 500 kit": TG(20.3_34.3).pdf

**Amount of relative differences between laboratories**

| Min | A | B | C | D | E | F | G | H | I | J | K | L | M | N |
|---|---|---|---|---|---|---|---|---|---|---|---|---|---|---|
| A | - | 5.128 | 5.222 | 4.734 | 5.685 | 5.303 | 4.5 | 4.165 | 5.312 | 7.145 | 5.143 | 5.074 | 5.615 | 5.737 |
| B | - | - | 1.111 | 3.953 | 4.44 | 3.886 | 4.307 | 3.391 | 1.68 | 6.937 | 1.656 | 2.013 | 3.806 | 3.45 |
| C | - | - | - | 4.11 | 4.453 | 4.469 | 4.367 | 1.971 | 0.996 | 6.484 | 1.579 | 2.278 | 2.744 | 2.313 |
| D | - | - | - | - | 5.131 | 4.835 | 2.012 | 0.127 | 3.918 | 6.859 | 3.978 | 3.399 | 4.623 | 4.338 |
| E | - | - | - | - | - | 0.924 | 5.809 | 5.158 | 4.787 | 6.545 | 4.519 | 5.192 | 4.092 | 5.029 |
| F | - | - | - | - | - | - | 6.113 | 4.663 | 5.524 | 5.487 | 4.973 | 5.675 | 5.295 | 6.046 |
| G | - | - | - | - | - | - | - | 2.24 | 4.271 | 6.988 | 4.135 | 3.828 | 4.88 | 4.791 |
| H | - | - | - | - | - | - | - | - | 2.733 | 6.716 | 2.691 | -1.018 | 4.042 | 3.432 |
| I | - | - | - | - | - | - | - | - | - | 7.151 | 1.678 | 3.096 | 3.747 | 3.825 |
| J | - | - | - | - | - | - | - | - | - | - | 4.893 | 5.692 | 5.64 | 6.446 |
| K | - | - | - | - | - | - | - | - | - | - | - | 2.955 | 2.751 | 1.075 |
| L | - | - | - | - | - | - | - | - | - | - | - | - | 4.021 | 3.403 |
| M | - | - | - | - | - | - | - | - | - | - | - | - | - | 3.651 |
| N | - | - | - | - | - | - | - | - | - | - | - | - | - | - |

| | |
|---|---|
| | Larger than 10 |
| | Between 8 and 9,99 |
| | Between 6 and 7,99 |
| | Between 4 and 5,99 |
| | Between 2 and 3,99 |
| | Between 0 and 1,99 |
| | NA (Calculation failed) |

| Median | A | B | C | D | E | F | G | H | I | J | K | L | M | N |
|---|---|---|---|---|---|---|---|---|---|---|---|---|---|---|
| A | - | 4.62 | 3.63 | 3.414 | 5.086 | 4.563 | 3.612 | 2.985 | 5.016 | -3.593 | 4.521 | 4.317 | 4.016 | 4.493 |
| B | - | - | 4.062 | 4.258 | 3.531 | -0.619 | 4.163 | 4.275 | 3.407 | 4.952 | 1.179 | 2.819 | 3.389 | 1.395 |
| C | - | - | - | 0.902 | 4.627 | 3.913 | 0.973 | 1.694 | 4.593 | 3.186 | 3.612 | 3.048 | 2.756 | 3.688 |
| D | - | - | - | - | 4.726 | 4.004 | -0.513 | 1.619 | 4.643 | 3.207 | 3.802 | 3.354 | 2.903 | 3.8 |
| E | - | - | - | - | - | 3.548 | 5.147 | 5.227 | -0.637 | 5.521 | 4.004 | 4.477 | 4.589 | 4.011 |
| F | - | - | - | - | - | - | 4.55 | 4.187 | 3.382 | 4.735 | 2.101 | 3.439 | 3.303 | 1.744 |
| G | - | - | - | - | - | - | - | 1.398 | 4.586 | 3.448 | 3.68 | 3.231 | 2.632 | 3.688 |
| H | - | - | - | - | - | - | - | - | 4.772 | 1.272 | 3.864 | 3.524 | 3.455 | 4.138 |
| I | - | - | - | - | - | - | - | - | - | 5.551 | 3.834 | 4.261 | 4.562 | 3.87 |
| J | - | - | - | - | - | - | - | - | - | - | 3.822 | 3.038 | 3.485 | 3.971 |
| K | - | - | - | - | - | - | - | - | - | - | - | 2.216 | 2.945 | -5.327 |
| L | - | - | - | - | - | - | - | - | - | - | - | - | 1.549 | 2.054 |
| M | - | - | - | - | - | - | - | - | - | - | - | - | - | 2.717 |
| N | - | - | - | - | - | - | - | - | - | - | - | - | - | - |

| Max | A | B | C | D | E | F | G | H | I | J | K | L | M | N |
|---|---|---|---|---|---|---|---|---|---|---|---|---|---|---|
| A | - | 4.213 | 2.786 | 0.169 | 4.472 | 2.102 | 2.311 | 3.239 | 4.883 | 5.991 | 3.758 | 3.484 | -1.395 | 3.692 |
| B | - | - | 4.967 | 4.413 | 2.24 | 4.801 | 4.054 | 5.117 | 3.749 | 6.645 | 2.677 | 3.166 | 4.476 | 2.776 |
| C | - | - | - | 2.875 | 4.734 | 1.362 | 3.47 | 1.036 | 5.094 | 5.742 | 4.269 | 4.055 | 2.762 | 4.206 |
| D | - | - | - | - | 4.369 | 2.588 | 2.003 | 3.362 | 4.847 | 6.037 | 3.651 | 3.325 | -0.365 | 3.581 |
| E | - | - | - | - | - | 5.157 | 4.395 | 5.338 | 3.278 | 6.764 | 3.431 | 3.721 | 4.778 | 3.455 |
| F | - | - | - | - | - | - | 3.096 | 2.553 | 5.004 | 5.878 | 4.036 | 3.884 | 2.099 | 4.016 |
| G | - | - | - | - | - | - | - | 3.943 | 4.69 | 6.19 | 3.198 | 2.661 | 2.32 | 3.051 |
| H | - | - | - | - | - | - | - | - | 5.14 | 5.668 | 4.375 | 4.21 | 3.055 | 4.303 |
| I | - | - | - | - | - | - | - | - | - | 7.044 | 4.505 | 4.696 | 5.373 | 4.559 |
| J | - | - | - | - | - | - | - | - | - | - | 5.573 | 5.534 | 5.343 | 5.576 |
| K | - | - | - | - | - | - | - | - | - | - | - | 1.416 | 3.873 | -0.683 |
| L | - | - | - | - | - | - | - | - | - | - | - | - | 3.603 | 1.107 |
| M | - | - | - | - | - | - | - | - | - | - | - | - | - | 3.647 |
| N | - | - | - | - | - | - | - | - | - | - | - | - | - | - |
