## Supplementary Data S2 for "Interlaboratory comparison of standardised metabolomics and lipidomics analyses in human and rodent blood using the MxP® Quant 500 kit": TG(20.3_36.3).pdf

**Amount of relative differences between laboratories**

| Min | A | B | C | D | E | F | G | H | I | J | K | L | M | N |
|---|---|---|---|---|---|---|---|---|---|---|---|---|---|---|
| A | - | 4.445 | 5.413 | 4.381 | 5.546 | 5.848 | 4.724 | 4.934 | 4.899 | 6.907 | 5.031 | 4.641 | 5.036 | 5.008 |
| B | - | - | 4.45 | 1.746 | 4.917 | 5.379 | 0.963 | 3.629 | 2.164 | 6.61 | 1.169 | 0.912 | 3.577 | 2.693 |
| C | - | - | - | 4.912 | 3.57 | 4.62 | 4.622 | 3.003 | 4.519 | 6.268 | 4.931 | 4.566 | 3.404 | 4.427 |
| D | - | - | - | - | 5.073 | 5.466 | 2.319 | 3.893 | 2.995 | 6.608 | 2.373 | 2.42 | 3.885 | 3.186 |
| E | - | - | - | - | - | 4.138 | 5.353 | 4.514 | 5.174 | 6.097 | 5.539 | 5.369 | 4.687 | 5.189 |
| F | - | - | - | - | - | - | 5.874 | 5.022 | 5.866 | 4.278 | 6.232 | 5.861 | 5.303 | 5.845 |
| G | - | - | - | - | - | - | - | 3.528 | 1.056 | 6.43 | 1.018 | -0.564 | 3.417 | 1.808 |
| H | - | - | - | - | - | - | - | - | 3.083 | 6.725 | 3.438 | 3.517 | -3.127 | 2.62 |
| I | - | - | - | - | - | - | - | - | - | 6.805 | -1.279 | 2.306 | 2.703 | 1.494 |
| J | - | - | - | - | - | - | - | - | - | - | 5.662 | 4.86 | 3.85 | 4.939 |
| K | - | - | - | - | - | - | - | - | - | - | - | 1.043 | 3.539 | 2.373 |
| L | - | - | - | - | - | - | - | - | - | - | - | - | 3.306 | 2.442 |
| M | - | - | - | - | - | - | - | - | - | - | - | - | - | 2.756 |
| N | - | - | - | - | - | - | - | - | - | - | - | - | - | - |

| | |
|---|---|
| | Larger than 10 |
| | Between 8 and 9,99 |
| | Between 6 and 7,99 |
| | Between 4 and 5,99 |
| | Between 2 and 3,99 |
| | Between 0 and 1,99 |
| | NA (Calculation failed) |

| Median | A | B | C | D | E | F | G | H | I | J | K | L | M | N |
|---|---|---|---|---|---|---|---|---|---|---|---|---|---|---|
| A | - | 3.492 | 4.381 | 3.052 | 5.005 | 4.797 | 3.167 | 4.499 | 4.375 | 4.748 | 3.013 | 4.207 | 4.439 | 4.445 |
| B | - | - | 3.606 | 0.952 | 4.621 | 4.295 | 0.61 | 3.69 | 3.65 | 5.231 | 0.778 | 3.189 | 3.713 | 3.732 |
| C | - | - | - | 4.026 | 3.812 | 3.041 | 4.007 | -2.73 | -0.392 | 5.759 | 3.974 | 1.959 | 0.071 | 0.3 |
| D | - | - | - | - | 4.71 | 4.408 | -1.344 | 3.87 | 3.824 | 5.137 | -3.184 | 3.447 | 3.882 | 3.892 |
| E | - | - | - | - | - | 2.665 | 5.125 | 4.015 | 3.948 | 6.318 | 5.117 | 4.385 | 3.922 | 3.897 |
| F | - | - | - | - | - | - | 4.725 | 3.358 | 2.744 | 5.945 | 4.638 | 3.747 | 2.838 | 2.761 |
| G | - | - | - | - | - | - | - | 3.813 | 3.814 | 5.106 | -3.941 | 3.399 | 3.862 | 3.874 |
| H | - | - | - | - | - | - | - | - | -0.982 | 5.869 | 4.079 | 2.124 | -2.519 | -1.469 |
| I | - | - | - | - | - | - | - | - | - | 5.873 | 4.014 | 1.816 | -0.16 | 0.087 |
| J | - | - | - | - | - | - | - | - | - | - | 4.251 | 4.65 | 4.904 | 4.935 |
| K | - | - | - | - | - | - | - | - | - | - | - | 3.416 | 3.866 | 3.881 |
| L | - | - | - | - | - | - | - | - | - | - | - | - | 2.042 | 2.071 |
| M | - | - | - | - | - | - | - | - | - | - | - | - | - | -3.004 |
| N | - | - | - | - | - | - | - | - | - | - | - | - | - | - |

| Max | A | B | C | D | E | F | G | H | I | J | K | L | M | N |
|---|---|---|---|---|---|---|---|---|---|---|---|---|---|---|
| A | - | 1.701 | 3.395 | 1.25 | 4.643 | 3.144 | -2.797 | 4.007 | 4.155 | 6.154 | -0.375 | 3.958 | 4.038 | 4.203 |
| B | - | - | 2.924 | 0.338 | 4.452 | 2.495 | 1.716 | 3.738 | 3.922 | 6.293 | 1.531 | 3.633 | 3.772 | 3.953 |
| C | - | - | - | 3.288 | 3.916 | 1.114 | 3.555 | 2.656 | 3.069 | 6.551 | 3.502 | 2.348 | 2.715 | 3.101 |
| D | - | - | - | - | 4.493 | 2.725 | 0.951 | 3.852 | 4.014 | 6.234 | 0.479 | 3.747 | 3.88 | 4.062 |
| E | - | - | - | - | - | 4.431 | 4.995 | 3.417 | 3.133 | 6.961 | 4.964 | 3.54 | 3.341 | 2.986 |
| F | - | - | - | - | - | - | 3.341 | 3 | 3.31 | 6.428 | 3.319 | 2.777 | 3.03 | 3.262 |
| G | - | - | - | - | - | - | - | 4.011 | 4.164 | 6.147 | -0.954 | 3.91 | 4.033 | 4.191 |
| H | - | - | - | - | - | - | - | - | 1.042 | 6.756 | 4.225 | 0.192 | -1.683 | 1.301 |
| I | - | - | - | - | - | - | - | - | - | 6.782 | 4.397 | 1.523 | 0.821 | -1.013 |
| J | - | - | - | - | - | - | - | - | - | - | 5.312 | 5.634 | 5.671 | 5.655 |
| K | - | - | - | - | - | - | - | - | - | - | - | 3.866 | 3.999 | 4.156 |
| L | - | - | - | - | - | - | - | - | - | - | - | - | 0.465 | 1.927 |
| M | - | - | - | - | - | - | - | - | - | - | - | - | - | 1.231 |
| N | - | - | - | - | - | - | - | - | - | - | - | - | - | - |
