## Supplementary Data S2 for "Interlaboratory comparison of standardised metabolomics and lipidomics analyses in human and rodent blood using the MxP® Quant 500 kit": TG(20.4_32.2).pdf

**Amount of relative differences between laboratories**

| Min | A | B | C | D | E | F | G | H | I | J | K | L | M | N |
|---|---|---|---|---|---|---|---|---|---|---|---|---|---|---|
| A | - | 5.47 | 5.066 | 5.395 | 6.053 | 6.297 | 5.023 | 5.526 | 4.694 | 6.554 | 5.862 | 5.337 | 5.411 | 5.484 |
| B | - | - | 4.28 | 3.534 | 4.708 | 5.503 | 4.477 | -3.42 | 5.053 | 5.486 | 4.217 | 2.944 | 1.825 | 2.338 |
| C | - | - | - | 2.778 | 5.267 | 5.768 | 1.251 | 4.033 | 3.526 | 5.764 | 4.991 | 3.327 | 4.26 | 4.35 |
| D | - | - | - | - | 5.065 | 5.648 | 3.223 | 3.436 | 4.23 | 5.587 | 4.758 | 1.982 | 3.839 | 3.947 |
| E | - | - | - | - | - | 4.616 | 5.902 | 4.947 | 6.206 | 4.062 | 2.974 | 5.366 | 4.628 | 4.52 |
| F | - | - | - | - | - | - | 6.645 | 6.032 | 6.874 | 4.347 | 5.233 | 6.251 | 5.722 | 5.697 |
| G | - | - | - | - | - | - | - | 4.203 | 3.155 | 5.84 | 5.073 | 3.606 | 4.373 | 4.461 |
| H | - | - | - | - | - | - | - | - | 5.04 | 5.417 | 4.194 | 2.982 | 2.096 | 2.48 |
| I | - | - | - | - | - | - | - | - | - | 6.114 | 5.344 | 4.293 | 4.706 | 4.794 |
| J | - | - | - | - | - | - | - | - | - | - | 0.417 | 4.597 | 2.125 | 2.224 |
| K | - | - | - | - | - | - | - | - | - | - | - | 4.869 | 3.631 | 3.509 |
| L | - | - | - | - | - | - | - | - | - | - | - | - | 3.27 | 3.473 |
| M | - | - | - | - | - | - | - | - | - | - | - | - | - | 0.731 |
| N | - | - | - | - | - | - | - | - | - | - | - | - | - | - |

| Median | A | B | C | D | E | F | G | H | I | J | K | L | M | N |
|---|---|---|---|---|---|---|---|---|---|---|---|---|---|---|
| A | - | 4.676 | 3.293 | 3.525 | 5.222 | 5.493 | 3.1 | 4.458 | 3.123 | 1.755 | 4.684 | 4.385 | 4.821 | 4.892 |
| B | - | - | 4.214 | 3.921 | 4.172 | 4.773 | 4.247 | 1.818 | 4.296 | 4.96 | 1.562 | 2.061 | 2.443 | 2.605 |
| C | - | - | - | 1.538 | 4.949 | 5.275 | -1.379 | 3.667 | -0.119 | 3.425 | 4.177 | 3.609 | 4.341 | 4.379 |
| D | - | - | - | - | 4.849 | 5.2 | 1.763 | 3.316 | 1.991 | 3.764 | 3.973 | 3.239 | 4.149 | 4.184 |
| E | - | - | - | - | - | 3.47 | 5.485 | 4.73 | 5.52 | 5.875 | 4.192 | 4.773 | 3.964 | 3.914 |
| F | - | - | - | - | - | - | 5.957 | 5.431 | 5.998 | 6.233 | 5.034 | 5.432 | 4.913 | 4.923 |
| G | - | - | - | - | - | - | - | 3.717 | -1.096 | 3.394 | 4.205 | 3.659 | 4.367 | 4.41 |
| H | - | - | - | - | - | - | - | - | 3.943 | 4.675 | 2.698 | -0.667 | 3.123 | 3.201 |
| I | - | - | - | - | - | - | - | - | - | 3.44 | 4.239 | 3.721 | 4.403 | 4.464 |
| J | - | - | - | - | - | - | - | - | - | - | 4.455 | 3.907 | 4.511 | 4.452 |
| K | - | - | - | - | - | - | - | - | - | - | - | 2.986 | 1.01 | 1.042 |
| L | - | - | - | - | - | - | - | - | - | - | - | - | 3.198 | 3.304 |
| M | - | - | - | - | - | - | - | - | - | - | - | - | - | -0.368 |
| N | - | - | - | - | - | - | - | - | - | - | - | - | - | - |

| Max | A | B | C | D | E | F | G | H | I | J | K | L | M | N |
|---|---|---|---|---|---|---|---|---|---|---|---|---|---|---|
| A | - | 1.623 | 3.818 | 3.847 | 3.903 | 3.636 | 3.654 | 2.107 | 2.425 | 6.71 | 3.503 | 1.304 | 3.403 | 3.354 |
| B | - | - | 4.146 | 4.182 | 3.56 | 3.296 | 4.007 | 2.902 | 3.094 | 6.784 | 3.898 | -0.243 | 2.96 | 2.89 |
| C | - | - | - | -1.162 | 4.645 | 4.548 | 0.464 | 3.12 | 2.958 | 6.295 | 1.286 | 3.833 | 4.435 | 4.416 |
| D | - | - | - | - | 4.658 | 4.57 | 0.893 | 3.173 | 3.028 | 6.281 | 1.454 | 3.87 | 4.459 | 4.441 |
| E | - | - | - | - | - | 0.975 | 4.979 | 4.45 | 4.521 | 7.1 | 4.915 | 3.827 | 2.113 | 2.296 |
| F | - | - | - | - | - | - | 4.839 | 4.234 | 4.329 | 7.048 | 4.766 | 3.59 | 1.055 | 1.368 |
| G | - | - | - | - | - | - | - | 2.889 | 2.69 | 6.341 | 0.091 | 3.707 | 4.36 | 4.34 |
| H | - | - | - | - | - | - | - | - | 0.005 | 6.594 | 2.808 | 2.64 | 3.829 | 3.792 |
| I | - | - | - | - | - | - | - | - | - | 6.558 | 2.566 | 2.834 | 3.912 | 3.882 |
| J | - | - | - | - | - | - | - | - | - | - | 5.506 | 5.701 | 5.818 | 5.811 |
| K | - | - | - | - | - | - | - | - | - | - | - | 3.607 | 4.31 | 4.277 |
| L | - | - | - | - | - | - | - | - | - | - | - | - | 3.103 | 3.051 |
| M | - | - | - | - | - | - | - | - | - | - | - | - | - | -1.45 |
| N | - | - | - | - | - | - | - | - | - | - | - | - | - | - |

| | |
|---|---|
| | Larger than 10 |
| | Between 8 and 9,99 |
| | Between 6 and 7,99 |
| | Between 4 and 5,99 |
| | Between 2 and 3,99 |
| | Between 0 and 1,99 |
| | NA (Calculation failed) |
