## Supplementary Data S2 for "Interlaboratory comparison of standardised metabolomics and lipidomics analyses in human and rodent blood using the MxP® Quant 500 kit": TG(20.4_34.3).pdf

**Amount of relative differences between laboratories**

| Min | A | B | C | D | E | F | G | H | I | J | K | L | M | N |
|---|---|---|---|---|---|---|---|---|---|---|---|---|---|---|
| A | - | 4.403 | 3.628 | 1.361 | 5.097 | 5.037 | 0.904 | 4.113 | 1.376 | 6.024 | 4.771 | 4.055 | 4.309 | 4.691 |
| B | - | - | 3.714 | 4.93 | 4.135 | 3.827 | 4.641 | 2.654 | 4.678 | 5.22 | 2.038 | 2.566 | 1.422 | 0.935 |
| C | - | - | - | 3.96 | 4.78 | 4.554 | 3.427 | 2.588 | 3.487 | 5.489 | 3.955 | 2.639 | 3.207 | 3.787 |
| D | - | - | - | - | 5.224 | 5.096 | 2.052 | 4.237 | 1.791 | 5.778 | 4.778 | 4.248 | 4.42 | 4.701 |
| E | - | - | - | - | - | 2.172 | 5.703 | 4.881 | 5.739 | 4.193 | 4.007 | 4.909 | 4.673 | 4.115 |
| F | - | - | - | - | - | - | 5.462 | 4.505 | 5.522 | 4.445 | 3.537 | 4.542 | 4.193 | 3.802 |
| G | - | - | - | - | - | - | - | 3.943 | -1.794 | 5.761 | 4.609 | 3.953 | 4.183 | 4.516 |
| H | - | - | - | - | - | - | - | - | 4.192 | 5.131 | 3.183 | -1.372 | 1.904 | 2.909 |
| I | - | - | - | - | - | - | - | - | - | 6.212 | 4.76 | 3.876 | 4.162 | 4.691 |
| J | - | - | - | - | - | - | - | - | - | - | 2.836 | 3.538 | 3.219 | 3.168 |
| K | - | - | - | - | - | - | - | - | - | - | - | 3.404 | 2.859 | 1.014 |
| L | - | - | - | - | - | - | - | - | - | - | - | - | 1.743 | 3.194 |
| M | - | - | - | - | - | - | - | - | - | - | - | - | - | 2.503 |
| N | - | - | - | - | - | - | - | - | - | - | - | - | - | - |

| | |
|---|---|
| | Larger than 10 |
| | Between 8 and 9,99 |
| | Between 6 and 7,99 |
| | Between 4 and 5,99 |
| | Between 2 and 3,99 |
| | Between 0 and 1,99 |
| | NA (Calculation failed) |

| Median | A | B | C | D | E | F | G | H | I | J | K | L | M | N |
|---|---|---|---|---|---|---|---|---|---|---|---|---|---|---|
| A | - | 3.632 | -1.482 | 3.27 | 4.798 | 4.392 | 2.414 | 0.144 | 2.032 | 3.997 | 4.016 | 3.493 | 3.56 | 4.114 |
| B | - | - | 3.513 | 4.618 | 4.176 | 3.437 | 4.213 | 3.512 | 2.915 | 4.79 | 1.922 | -5.038 | -0.69 | 2.699 |
| C | - | - | - | 3.537 | 4.746 | 4.31 | 2.671 | -3.318 | 1.793 | 3.906 | 3.801 | 3.366 | 3.311 | 4.025 |
| D | - | - | - | - | 5.112 | 4.815 | 2.139 | 3.355 | 3.697 | 2.126 | 4.526 | 4.285 | 4.272 | 4.632 |
| E | - | - | - | - | - | 3.181 | 5.491 | 5.182 | 4.987 | 5.843 | 4.12 | 4.505 | 4.537 | 3.783 |
| F | - | - | - | - | - | - | 5.016 | 4.607 | 4.331 | 5.385 | 3.009 | 3.603 | 3.662 | 2.166 |
| G | - | - | - | - | - | - | - | 2.61 | 3.179 | 3.177 | 4.29 | 3.981 | 3.971 | 4.416 |
| H | - | - | - | - | - | - | - | - | 1.866 | 3.769 | 3.705 | 3.333 | 3.223 | 4.012 |
| I | - | - | - | - | - | - | - | - | - | 4.293 | 3.744 | 2.939 | 3.055 | 3.846 |
| J | - | - | - | - | - | - | - | - | - | - | 4.077 | 4.094 | 3.843 | 4.476 |
| K | - | - | - | - | - | - | - | - | - | - | - | 1.998 | 2.222 | 1.552 |
| L | - | - | - | - | - | - | - | - | - | - | - | - | -5.012 | 2.742 |
| M | - | - | - | - | - | - | - | - | - | - | - | - | - | 2.834 |
| N | - | - | - | - | - | - | - | - | - | - | - | - | - | - |

| Max | A | B | C | D | E | F | G | H | I | J | K | L | M | N |
|---|---|---|---|---|---|---|---|---|---|---|---|---|---|---|
| A | - | 0.691 | 3.505 | 4.214 | 4.216 | 2.387 | 3.725 | 4.293 | 2.408 | 6.296 | 1.002 | 2.406 | -0.65 | 3.231 |
| B | - | - | 3.287 | 4.094 | 4.232 | 2.668 | 3.526 | 4.166 | 2.837 | 6.282 | 1.759 | 2.704 | 1.166 | 3.428 |
| C | - | - | - | 2.728 | 4.695 | 3.904 | 0.635 | 2.899 | 3.945 | 5.938 | 3.578 | 3.892 | 3.444 | 4.216 |
| D | - | - | - | - | 4.935 | 4.33 | 2.229 | -0.329 | 4.372 | 5.648 | 4.126 | 4.326 | 4.036 | 4.565 |
| E | - | - | - | - | - | 3.921 | 5.217 | 5.493 | 3.813 | 6.857 | 4.251 | 3.863 | 4.324 | 3.385 |
| F | - | - | - | - | - | - | 4.286 | 4.728 | -0.189 | 6.506 | 1.815 | -2.098 | 2.271 | 2.11 |
| G | - | - | - | - | - | - | - | 2.527 | 4.058 | 5.857 | 3.743 | 4.012 | 3.612 | 4.315 |
| H | - | - | - | - | - | - | - | - | 4.425 | 5.68 | 4.176 | 4.381 | 4.097 | 4.595 |
| I | - | - | - | - | - | - | - | - | - | 6.524 | 2.008 | -3.131 | 2.329 | 1.902 |
| J | - | - | - | - | - | - | - | - | - | - | 5.569 | 5.613 | 5.562 | 5.682 |
| K | - | - | - | - | - | - | - | - | - | - | - | 1.733 | 0.096 | 2.915 |
| L | - | - | - | - | - | - | - | - | - | - | - | - | 2.241 | 2.137 |
| M | - | - | - | - | - | - | - | - | - | - | - | - | - | 3.086 |
| N | - | - | - | - | - | - | - | - | - | - | - | - | - | - |
