## Supplementary Data S2 for "Interlaboratory comparison of standardised metabolomics and lipidomics analyses in human and rodent blood using the MxP® Quant 500 kit": TG(20.4_36.4).pdf

**Amount of relative differences between laboratories**

| Min | A | B | C | D | E | F | G | H | I | J | K | L | M | N |
|---|---|---|---|---|---|---|---|---|---|---|---|---|---|---|
| A | - | 4.182 | 3.311 | 0.316 | 4.3 | 5.378 | 2.351 | 4.96 | 3.953 | 5.048 | 4.099 | 0.515 | 4.262 | 4.564 |
| B | - | - | 3.32 | 4.425 | 2.366 | 4.782 | 3.995 | 3.973 | 1.285 | 2.931 | 0.709 | 4.106 | 0.116 | 2.914 |
| C | - | - | - | 3.473 | 3.797 | 5.093 | 2.518 | 4.53 | 2.733 | 4.255 | 2.915 | 2.883 | 3.265 | 4.041 |
| D | - | - | - | - | 4.399 | 5.404 | 2.685 | 5.008 | 4.062 | 5.071 | 4.198 | 1.512 | 4.335 | 4.626 |
| E | - | - | - | - | - | 4.987 | 3.697 | 4.177 | 1.14 | 5.047 | 0.378 | 4.481 | 1.786 | 3.366 |
| F | - | - | - | - | - | - | 5.886 | 3.994 | 5.288 | 5.161 | 5.287 | 5.853 | 5.151 | 4.652 |
| G | - | - | - | - | - | - | - | 4.777 | 3.516 | 4.629 | 3.667 | 1.21 | 3.879 | 4.323 |
| H | - | - | - | - | - | - | - | - | 4.343 | 3.93 | 4.326 | 5.181 | 4.101 | 3.09 |
| I | - | - | - | - | - | - | - | - | - | 4.238 | 0.591 | 4.006 | 1.863 | 3.37 |
| J | - | - | - | - | - | - | - | - | - | - | -0.539 | 3.476 | 1.815 | 3.471 |
| K | - | - | - | - | - | - | - | - | - | - | - | 4.075 | 1.18 | 3.198 |
| L | - | - | - | - | - | - | - | - | - | - | - | - | 4.272 | 4.555 |
| M | - | - | - | - | - | - | - | - | - | - | - | - | - | 2.725 |
| N | - | - | - | - | - | - | - | - | - | - | - | - | - | - |

| | |
|---|---|
| | Larger than 10 |
| | Between 8 and 9,99 |
| | Between 6 and 7,99 |
| | Between 4 and 5,99 |
| | Between 2 and 3,99 |
| | Between 0 and 1,99 |
| | NA (Calculation failed) |

| Median | A | B | C | D | E | F | G | H | I | J | K | L | M | N |
|---|---|---|---|---|---|---|---|---|---|---|---|---|---|---|
| A | - | 3.242 | 0.593 | -0.911 | 4.792 | 4.659 | -3.134 | 3.985 | 3.69 | 3.807 | 3.363 | 2.751 | 3.562 | 4.082 |
| B | - | - | 2.954 | 3.319 | 4.398 | 4.224 | 3.223 | 3.013 | 2.194 | 4.49 | 1.391 | 0.967 | 1.927 | 3.157 |
| C | - | - | - | 1.12 | 4.752 | 4.609 | 0.686 | 3.863 | 3.515 | 3.787 | 3.241 | 2.396 | 3.394 | 3.978 |
| D | - | - | - | - | 4.821 | 4.678 | -1.277 | 4.027 | 3.746 | 3.781 | 3.423 | 2.872 | 3.611 | 4.122 |
| E | - | - | - | - | - | 2.041 | 5.273 | 4.056 | 4.344 | 5.944 | 4.664 | 4.863 | 4.494 | 3.763 |
| F | - | - | - | - | - | - | 5.142 | 3.787 | 4.193 | 5.598 | 4.286 | 4.729 | 4.231 | 3.687 |
| G | - | - | - | - | - | - | - | 3.995 | 3.692 | 3.706 | 3.419 | 2.78 | 3.578 | 4.081 |
| H | - | - | - | - | - | - | - | - | 1.978 | 4.946 | 2.375 | 3.459 | 2.164 | -0.46 |
| I | - | - | - | - | - | - | - | - | - | 4.9 | 1.46 | 2.819 | 0.347 | 2.281 |
| J | - | - | - | - | - | - | - | - | - | - | 4.245 | 3.807 | 4.272 | 4.529 |
| K | - | - | - | - | - | - | - | - | - | - | - | 2.182 | 0.354 | 2.811 |
| L | - | - | - | - | - | - | - | - | - | - | - | - | 2.377 | 3.472 |
| M | - | - | - | - | - | - | - | - | - | - | - | - | - | 2.477 |
| N | - | - | - | - | - | - | - | - | - | - | - | - | - | - |

| Max | A | B | C | D | E | F | G | H | I | J | K | L | M | N |
|---|---|---|---|---|---|---|---|---|---|---|---|---|---|---|
| A | - | -1.706 | 2.623 | -3.112 | 5.119 | 3.314 | 2.12 | -1.488 | 3.401 | 5.318 | 2.402 | 3.526 | 2.499 | 3.331 |
| B | - | - | 2.515 | -1.073 | 5.133 | 3.379 | 1.989 | -8.141 | 3.45 | 5.286 | 2.519 | 3.562 | 2.602 | 3.369 |
| C | - | - | - | 2.569 | 5.265 | 3.933 | 0.796 | 2.461 | 3.979 | 4.995 | 3.439 | 4.058 | 3.493 | 3.911 |
| D | - | - | - | - | 5.111 | 3.302 | 2.162 | -0.989 | 3.382 | 5.315 | 2.366 | 3.505 | 2.478 | 3.315 |
| E | - | - | - | - | - | 5.239 | 5.896 | 5.738 | 5.199 | 6.828 | 5.486 | 5.138 | 5.466 | 5.249 |
| F | - | - | - | - | - | - | 4 | 3.53 | -0.859 | 5.791 | 2.389 | 0.769 | 2.262 | -1.61 |
| G | - | - | - | - | - | - | - | 1.928 | 3.835 | 5.081 | 3.206 | 3.921 | 3.267 | 3.782 |
| H | - | - | - | - | - | - | - | - | 3.444 | 5.3 | 2.506 | 3.56 | 2.61 | 3.348 |
| I | - | - | - | - | - | - | - | - | - | 5.825 | 2.552 | 0.101 | 2.425 | -0.413 |
| J | - | - | - | - | - | - | - | - | - | - | 5.014 | 5.196 | 5.023 | 5.144 |
| K | - | - | - | - | - | - | - | - | - | - | - | 2.715 | -1.141 | 2.266 |
| L | - | - | - | - | - | - | - | - | - | - | - | - | 2.721 | 0.687 |
| M | - | - | - | - | - | - | - | - | - | - | - | - | - | 2.169 |
| N | - | - | - | - | - | - | - | - | - | - | - | - | - | - |
