## Supplementary Data S2 for "Interlaboratory comparison of standardised metabolomics and lipidomics analyses in human and rodent blood using the MxP® Quant 500 kit": TG(20.4_36.5).pdf

**Amount of relative differences between laboratories**

| Min | A | B | C | D | E | F | G | H | I | J | K | L | M | N |
|---|---|---|---|---|---|---|---|---|---|---|---|---|---|---|
| A | - | 5.161 | 4.435 | 2.79 | 4.999 | 6.07 | 0.632 | 5.187 | 4.757 | 5.372 | 3.375 | 1.907 | 4.626 | 4.297 |
| B | - | - | 3.784 | 5.381 | 4.015 | 5.607 | 5.222 | 3.118 | 1.634 | 2.784 | 5.797 | 4.819 | 2.631 | 3.539 |
| C | - | - | - | 4.85 | 4.463 | 5.805 | 4.61 | 4.317 | 3.086 | 3.783 | 5.229 | 4.071 | 2.492 | -2.453 |
| D | - | - | - | - | 5.076 | 6.12 | 2.872 | 5.473 | 5.181 | 5.913 | 0.154 | 3.551 | 5.078 | 4.814 |
| E | - | - | - | - | - | 5.686 | 5.673 | 2.882 | 1.985 | 3.669 | 5.859 | 5.415 | 3.212 | 4.113 |
| F | - | - | - | - | - | - | 7.302 | 6.269 | 6.689 | 7.008 | 7.673 | 7.11 | 6.735 | 6.854 |
| G | - | - | - | - | - | - | - | 5.184 | 4.806 | 5.219 | 3.271 | 2.566 | 4.641 | 4.357 |
| H | - | - | - | - | - | - | - | - | 3.679 | 4.272 | 6.12 | 5.295 | 4.043 | 4.445 |
| I | - | - | - | - | - | - | - | - | - | 3.027 | 5.67 | 4.885 | 1.596 | 3.267 |
| J | - | - | - | - | - | - | - | - | - | - | 5.18 | 3.529 | 2.521 | 0.456 |
| K | - | - | - | - | - | - | - | - | - | - | - | 3.397 | 4.989 | 4.697 |
| L | - | - | - | - | - | - | - | - | - | - | - | - | 4.509 | 4.096 |
| M | - | - | - | - | - | - | - | - | - | - | - | - | - | 2.49 |
| N | - | - | - | - | - | - | - | - | - | - | - | - | - | - |

| Median | A | B | C | D | E | F | G | H | I | J | K | L | M | N |
|---|---|---|---|---|---|---|---|---|---|---|---|---|---|---|
| A | - | 3.372 | 2.844 | 1.766 | 5.014 | 5.496 | 2.631 | 4.152 | 4.26 | 3.938 | 1.406 | 2.701 | 3.839 | 3.647 |
| B | - | - | 1.903 | 4.177 | 4.568 | 5.189 | 4.378 | 2.48 | 3.339 | 4.658 | 3.786 | 2.598 | 1.216 | 0.693 |
| C | - | - | - | 3.713 | 4.738 | 5.305 | 3.998 | 3.337 | 3.718 | 4.447 | 3.309 | 0.971 | 2.7 | 2.407 |
| D | - | - | - | - | 5.057 | 5.543 | 0.919 | 4.462 | 4.505 | 3.204 | -0.144 | 3.346 | 4.215 | 4.019 |
| E | - | - | - | - | - | 4.261 | 5.852 | 4.417 | 4.236 | 6.155 | 5.676 | 5.275 | 4.717 | 4.87 |
| F | - | - | - | - | - | - | 6.547 | 5.693 | 5.362 | 6.441 | 6.283 | 6.119 | 5.832 | 5.812 |
| G | - | - | - | - | - | - | - | 4.486 | 4.614 | 2.873 | 1.999 | 3.6 | 4.268 | 4.163 |
| H | - | - | - | - | - | - | - | - | 2.404 | 4.963 | 4.301 | 3.608 | 1.804 | 1.946 |
| I | - | - | - | - | - | - | - | - | - | 5.375 | 4.744 | 4.086 | 2.814 | 3.14 |
| J | - | - | - | - | - | - | - | - | - | - | 3.064 | 3.741 | 4.323 | 4.315 |
| K | - | - | - | - | - | - | - | - | - | - | - | 3.209 | 4.098 | 3.889 |
| L | - | - | - | - | - | - | - | - | - | - | - | - | 3.082 | 2.649 |
| M | - | - | - | - | - | - | - | - | - | - | - | - | - | 0.344 |
| N | - | - | - | - | - | - | - | - | - | - | - | - | - | - |

| Max | A | B | C | D | E | F | G | H | I | J | K | L | M | N |
|---|---|---|---|---|---|---|---|---|---|---|---|---|---|---|
| A | - | 3.139 | 2.12 | -0.683 | 5.042 | 2.9 | 3.508 | 2.852 | 3.721 | 5.607 | 0.74 | 3.221 | 1.809 | 2.611 |
| B | - | - | 2.091 | 3.151 | 5.27 | 3.953 | 1.217 | 0.512 | 4.333 | 5.172 | 3.259 | 4.059 | 3.484 | 3.759 |
| C | - | - | - | 2.284 | 5.163 | 3.526 | 2.711 | 1.611 | 4.063 | 5.419 | 2.464 | 3.729 | 2.889 | 3.31 |
| D | - | - | - | - | 5.018 | 2.887 | 3.606 | 2.988 | 3.632 | 5.648 | -0.343 | 3.1 | 1.464 | 2.453 |
| E | - | - | - | - | - | 5.173 | 6.029 | 5.904 | 4.914 | 6.916 | 5.562 | 5.116 | 5.461 | 5.321 |
| F | - | - | - | - | - | - | 4.437 | 4.145 | 2.579 | 5.961 | 2.858 | -0.287 | 2.425 | 1.372 |
| G | - | - | - | - | - | - | - | 1.89 | 4.448 | 5.07 | 3.54 | 4.209 | 3.737 | 3.963 |
| H | - | - | - | - | - | - | - | - | 4.25 | 5.255 | 3.031 | 3.959 | 3.327 | 3.619 |
| I | - | - | - | - | - | - | - | - | - | 6.144 | 3.733 | 2.037 | 3.486 | 3.016 |
| J | - | - | - | - | - | - | - | - | - | - | 5.058 | 5.282 | 5.124 | 5.193 |
| K | - | - | - | - | - | - | - | - | - | - | - | 2.988 | 0.915 | 2.266 |
| L | - | - | - | - | - | - | - | - | - | - | - | - | 2.684 | 1.797 |
| M | - | - | - | - | - | - | - | - | - | - | - | - | - | 1.405 |
| N | - | - | - | - | - | - | - | - | - | - | - | - | - | - |

| | |
|---|---|
| | Larger than 10 |
| | Between 8 and 9,99 |
| | Between 6 and 7,99 |
| | Between 4 and 5,99 |
| | Between 2 and 3,99 |
| | Between 0 and 1,99 |
| | NA (Calculation failed) |
