## Supplementary Data S2 for "Interlaboratory comparison of standardised metabolomics and lipidomics analyses in human and rodent blood using the MxP® Quant 500 kit": TG(20.5_34.2).pdf

**Amount of relative differences between laboratories**

| Min | A | B | C | D | E | F | G | H | I | J | K | L | M | N |
|---|---|---|---|---|---|---|---|---|---|---|---|---|---|---|
| A | - | 5.55 | 5.091 | 4.564 | 5.656 | 6.057 | 4.673 | 3.624 | -3.487 | 7.606 | 2.357 | 5.045 | 4.694 | 4.542 |
| B | - | - | 4.971 | 5.554 | 0.516 | 3.52 | 5.47 | 5.924 | 6.349 | 7.488 | 6.165 | 4.929 | 5.309 | 5.415 |
| C | - | - | - | 3.492 | 4.506 | 4.897 | 3.217 | 4.36 | 5.161 | 7.144 | 4.837 | 0.152 | 2.394 | 2.887 |
| D | - | - | - | - | 5.007 | 5.299 | 0.716 | 3.216 | 4.505 | 7.199 | 4.059 | 3.531 | 2.304 | 1.805 |
| E | - | - | - | - | - | 3.195 | 5.272 | 5.675 | 6.084 | 6.955 | 5.929 | 4.781 | 5.065 | 5.032 |
| F | - | - | - | - | - | - | 5.482 | 5.831 | 6.256 | 6.77 | 6.07 | 4.996 | 5.261 | 5.337 |
| G | - | - | - | - | - | - | - | 3.453 | 4.649 | 7.168 | 4.209 | 3.295 | 1.784 | 0.626 |
| H | - | - | - | - | - | - | - | - | 3.691 | 7.154 | 2.781 | 4.248 | 3.709 | 3.47 |
| I | - | - | - | - | - | - | - | - | - | 7.768 | 2.436 | 5.094 | 4.732 | 4.572 |
| J | - | - | - | - | - | - | - | - | - | - | 6.226 | 5.654 | 5.687 | 5.659 |
| K | - | - | - | - | - | - | - | - | - | - | - | 4.61 | 4.232 | 4.091 |
| L | - | - | - | - | - | - | - | - | - | - | - | - | 2.942 | 3.294 |
| M | - | - | - | - | - | - | - | - | - | - | - | - | - | 0.946 |
| N | - | - | - | - | - | - | - | - | - | - | - | - | - | - |

| | |
|---|---|
| | Larger than 10 |
| | Between 8 and 9,99 |
| | Between 6 and 7,99 |
| | Between 4 and 5,99 |
| | Between 2 and 3,99 |
| | Between 0 and 1,99 |
| | NA (Calculation failed) |

| Median | A | B | C | D | E | F | G | H | I | J | K | L | M | N |
|---|---|---|---|---|---|---|---|---|---|---|---|---|---|---|
| A | - | 5.023 | 2.504 | -4.834 | 4.492 | 4.439 | -0.021 | 3.026 | 1.873 | 4.058 | 3.926 | 3.924 | 2.453 | 3.807 |
| B | - | - | 5.307 | 5.593 | 3.937 | 4.016 | 5.566 | 5.948 | 5.462 | 6.08 | 6.145 | 4.719 | 5.32 | 4.805 |
| C | - | - | - | 2.652 | 4.114 | 4.064 | 2.519 | 4.014 | 1.788 | 4.27 | 4.485 | 3.226 | -1.866 | 3.036 |
| D | - | - | - | - | 4.486 | 4.437 | -0.826 | 3.191 | 1.537 | 3.7 | 3.923 | 3.891 | 2.486 | 3.783 |
| E | - | - | - | - | - | -1.713 | 4.79 | 5.333 | 4.622 | 5.427 | 5.572 | 3.286 | 4.393 | 3.385 |
| F | - | - | - | - | - | - | 4.763 | 5.322 | 4.627 | 5.368 | 5.554 | 3.194 | 4.344 | 3.374 |
| G | - | - | - | - | - | - | - | 3.291 | 1.156 | 3.75 | 3.984 | 3.839 | 2.348 | 3.723 |
| H | - | - | - | - | - | - | - | - | 3.449 | 1.842 | 2.475 | 4.455 | 3.755 | 4.384 |
| I | - | - | - | - | - | - | - | - | - | 4.444 | 4.302 | 3.572 | 0.774 | 3.413 |
| J | - | - | - | - | - | - | - | - | - | - | 3.603 | 3.95 | 3.061 | 3.922 |
| K | - | - | - | - | - | - | - | - | - | - | - | 4.704 | 4.173 | 4.649 |
| L | - | - | - | - | - | - | - | - | - | - | - | - | 3.447 | 0.354 |
| M | - | - | - | - | - | - | - | - | - | - | - | - | - | 3.105 |
| N | - | - | - | - | - | - | - | - | - | - | - | - | - | - |

| Max | A | B | C | D | E | F | G | H | I | J | K | L | M | N |
|---|---|---|---|---|---|---|---|---|---|---|---|---|---|---|
| A | - | 4.614 | 3.065 | 3.589 | 3.154 | 2.111 | 3.634 | 4.495 | 2.477 | 6.173 | 4.819 | 2.692 | 2.495 | 3.201 |
| B | - | - | 5.448 | 5.613 | 4.465 | 4.748 | 5.617 | 5.965 | 4.647 | 7.05 | 6.132 | 4.604 | 5.327 | 4.347 |
| C | - | - | - | 1.907 | 3.877 | 3.546 | 1.919 | 3.709 | 3.695 | 5.942 | 4.203 | 3.743 | 1.286 | 4.024 |
| D | - | - | - | - | 4.142 | 3.888 | -5.317 | 3.173 | 3.991 | 5.781 | 3.828 | 4.038 | 2.576 | 4.254 |
| E | - | - | - | - | - | 1.987 | 4.439 | 5.033 | 1.437 | 6.442 | 5.286 | 0.111 | 3.83 | 0.102 |
| F | - | - | - | - | - | - | 4.135 | 4.811 | 0.615 | 6.336 | 5.088 | 0.976 | 3.391 | 2.373 |
| G | - | - | - | - | - | - | - | 3.164 | 4.009 | 5.775 | 3.819 | 4.045 | 2.582 | 4.266 |
| H | - | - | - | - | - | - | - | - | 4.525 | 5.389 | 2.238 | 4.547 | 3.779 | 4.698 |
| I | - | - | - | - | - | - | - | - | - | 6.404 | 5.164 | 0.014 | 3.565 | 1.947 |
| J | - | - | - | - | - | - | - | - | - | - | 4.811 | 5.573 | 5.355 | 5.619 |
| K | - | - | - | - | - | - | - | - | - | - | - | 4.748 | 4.14 | 4.87 |
| L | - | - | - | - | - | - | - | - | - | - | - | - | 3.662 | 1.648 |
| M | - | - | - | - | - | - | - | - | - | - | - | - | - | 3.813 |
| N | - | - | - | - | - | - | - | - | - | - | - | - | - | - |
