## Supplementary Data S2 for "Interlaboratory comparison of standardised metabolomics and lipidomics analyses in human and rodent blood using the MxP® Quant 500 kit": TG(20.5_36.2).pdf

**Amount of relative differences between laboratories**

| Min | A | B | C | D | E | F | G | H | I | J | K | L | M | N |
|---|---|---|---|---|---|---|---|---|---|---|---|---|---|---|
| A | - | 4.307 | 2.836 | 4.517 | 5.899 | 5.3 | 4.661 | -2.746 | 2.001 | 7.006 | 4.265 | 4.864 | 4.743 | 4.911 |
| B | - | - | 4.353 | 1.366 | 5.255 | 4.513 | 0.302 | 4.679 | 4.004 | 6.733 | 2.811 | 3.085 | 2.512 | 3.662 |
| C | - | - | - | 3.891 | 5.58 | 5.164 | 4.15 | 1.922 | 2.267 | 6.42 | 3.445 | 4.599 | 4.446 | 4.803 |
| D | - | - | - | - | 5.308 | 4.622 | 1.78 | 4.436 | 3.519 | 6.807 | 2.19 | 3.454 | 3.059 | 3.974 |
| E | - | - | - | - | - | 2.856 | 5.372 | 6.154 | 5.685 | 5.183 | 5.719 | 4.832 | 4.996 | 4.149 |
| F | - | - | - | - | - | - | 5.008 | 5.914 | 5.434 | 4.075 | 5.427 | 4.283 | 4.51 | 3.513 |
| G | - | - | - | - | - | - | - | 4.681 | 3.88 | 6.383 | 3.047 | 2.993 | 2.407 | 3.711 |
| H | - | - | - | - | - | - | - | - | 2.632 | 7.053 | 4.04 | 4.805 | 4.68 | 4.923 |
| I | - | - | - | - | - | - | - | - | - | 7.467 | 4.62 | 4.969 | 4.831 | 4.902 |
| J | - | - | - | - | - | - | - | - | - | - | 4.863 | 3.815 | 4.161 | 1.971 |
| K | - | - | - | - | - | - | - | - | - | - | - | 3.867 | 3.574 | 4.162 |
| L | - | - | - | - | - | - | - | - | - | - | - | - | 1.61 | 2.059 |
| M | - | - | - | - | - | - | - | - | - | - | - | - | - | 2.847 |
| N | - | - | - | - | - | - | - | - | - | - | - | - | - | - |

| Median | A | B | C | D | E | F | G | H | I | J | K | L | M | N |
|---|---|---|---|---|---|---|---|---|---|---|---|---|---|---|
| A | - | 2.512 | 4.133 | -2.582 | 4.176 | 3.64 | 0.024 | 3.353 | 3.887 | 5.071 | 0.349 | 3.904 | 3.797 | 4.524 |
| B | - | - | 4.632 | 2.602 | 3.675 | 2.663 | 2.301 | 4.145 | 3.145 | 5.321 | 2.482 | 3.269 | 3.055 | 4.087 |
| C | - | - | - | 3.907 | 4.911 | 4.633 | 3.996 | 2.483 | 4.747 | 3.463 | 3.933 | 4.786 | 4.727 | 5.057 |
| D | - | - | - | - | 4.151 | 3.529 | 0.151 | 3.467 | 3.805 | 5.007 | -1.588 | 3.888 | 3.757 | 4.431 |
| E | - | - | - | - | - | 3.009 | 4.291 | 5.143 | 2.316 | 5.786 | 4.436 | 1.72 | 2.353 | 1.819 |
| F | - | - | - | - | - | - | 3.407 | 4.659 | 1.421 | 5.4 | 3.656 | 2.155 | 1.466 | 3.389 |
| G | - | - | - | - | - | - | - | 3.635 | 3.688 | 5.004 | -0.159 | 3.789 | 3.643 | 4.359 |
| H | - | - | - | - | - | - | - | - | 4.499 | 4.384 | 3.371 | 4.528 | 4.46 | 4.895 |
| I | - | - | - | - | - | - | - | - | - | 5.924 | 3.993 | -1.335 | 0.572 | 3.355 |
| J | - | - | - | - | - | - | - | - | - | - | 4.141 | 4.935 | 4.857 | 5.078 |
| K | - | - | - | - | - | - | - | - | - | - | - | 3.847 | 3.716 | 4.42 |
| L | - | - | - | - | - | - | - | - | - | - | - | - | 0.547 | 3.037 |
| M | - | - | - | - | - | - | - | - | - | - | - | - | - | 3.275 |
| N | - | - | - | - | - | - | - | - | - | - | - | - | - | - |

| Max | A | B | C | D | E | F | G | H | I | J | K | L | M | N |
|---|---|---|---|---|---|---|---|---|---|---|---|---|---|---|
| A | - | 0.64 | 4.835 | 3.204 | 1.43 | 2.86 | 3.167 | 3.959 | 4.343 | 5.95 | 2.72 | 3.136 | 3.006 | 4.214 |
| B | - | - | 4.741 | 2.91 | 1.757 | 2.525 | 2.901 | 3.767 | 4.428 | 6.032 | 2.31 | 3.343 | 3.247 | 4.302 |
| C | - | - | - | 3.914 | 4.556 | 4.064 | 3.927 | 3.38 | 5.251 | 4.799 | 4.104 | 4.861 | 4.839 | 5.196 |
| D | - | - | - | - | 3.277 | 0.962 | -2.702 | 2.488 | 4.752 | 5.839 | 1.308 | 4.043 | 3.989 | 4.658 |
| E | - | - | - | - | - | 3.112 | 3.404 | 4.075 | 4.225 | 6.152 | 3.011 | 2.945 | 2.829 | 4.077 |
| F | - | - | - | - | - | - | 0.789 | 2.858 | 4.704 | 5.714 | -0.49 | 3.897 | 3.848 | 4.598 |
| G | - | - | - | - | - | - | - | 2.49 | 4.753 | 5.705 | 1.221 | 4.034 | 3.979 | 4.654 |
| H | - | - | - | - | - | - | - | - | 4.953 | 5.573 | 2.94 | 4.388 | 4.347 | 4.877 |
| I | - | - | - | - | - | - | - | - | - | 6.673 | 5.073 | 3.832 | 3.919 | 1.08 |
| J | - | - | - | - | - | - | - | - | - | - | 5.252 | 5.646 | 5.653 | 5.715 |
| K | - | - | - | - | - | - | - | - | - | - | - | 3.838 | 3.776 | 4.561 |
| L | - | - | - | - | - | - | - | - | - | - | - | - | -0.442 | 3.426 |
| M | - | - | - | - | - | - | - | - | - | - | - | - | - | 3.49 |
| N | - | - | - | - | - | - | - | - | - | - | - | - | - | - |

| |
|---|
| Larger than 10 |
| Between 8 and 9,99 |
| Between 6 and 7,99 |
| Between 4 and 5,99 |
| Between 2 and 3,99 |
| Between 0 and 1,99 |
| NA (Calculation failed) |
