## Supplementary Data S2 for "Interlaboratory comparison of standardised metabolomics and lipidomics analyses in human and rodent blood using the MxP® Quant 500 kit": TG(20.5_36.3).pdf

**Amount of relative differences between laboratories**

| Min | A | B | C | D | E | F | G | H | I | J | K | L | M | N |
|---|---|---|---|---|---|---|---|---|---|---|---|---|---|---|
| A | - | 4.54 | 5.14 | 4.797 | 5.47 | 6.423 | 5.444 | 5.655 | 1.971 | 6.547 | 2.728 | 4.848 | 4.511 | 3.8 |
| B | - | - | 3.497 | 2.825 | 5.092 | 5.928 | 4.42 | 4.884 | 4.178 | 6.07 | 3.25 | 3.364 | -1.958 | -1.377 |
| C | - | - | - | 2.478 | 4.636 | 5.79 | 3.357 | 4.32 | 5.009 | 4.858 | 4.52 | 0.226 | 3.64 | 3.575 |
| D | - | - | - | - | 4.905 | 5.876 | 3.943 | 4.604 | 4.701 | 6.69 | 4.183 | 1.762 | 2.937 | 3.218 |
| E | - | - | - | - | - | 5.127 | 4.466 | 2.875 | 6.151 | 6.068 | 6.035 | 5.279 | 5.749 | 5.573 |
| F | - | - | - | - | - | - | 6.002 | 5.63 | 6.828 | 6.594 | 6.609 | 6.252 | 6.608 | 6.371 |
| G | - | - | - | - | - | - | - | 3.403 | 5.33 | 5.913 | 4.985 | 3.57 | 4.621 | 4.341 |
| H | - | - | - | - | - | - | - | - | 5.651 | 5.23 | 5.348 | 4.515 | 5.212 | 4.94 |
| I | - | - | - | - | - | - | - | - | - | 7.103 | 3.654 | 4.909 | 4.634 | 4.086 |
| J | - | - | - | - | - | - | - | - | - | - | 4.766 | 5.189 | 5.565 | 2.251 |
| K | - | - | - | - | - | - | - | - | - | - | - | 4.416 | 3.262 | 3.205 |
| L | - | - | - | - | - | - | - | - | - | - | - | - | 3.418 | 3.824 |
| M | - | - | - | - | - | - | - | - | - | - | - | - | - | -0.131 |
| N | - | - | - | - | - | - | - | - | - | - | - | - | - | - |

| | |
|---|---|
| | Larger than 10 |
| | Between 8 and 9,99 |
| | Between 6 and 7,99 |
| | Between 4 and 5,99 |
| | Between 2 and 3,99 |
| | Between 0 and 1,99 |
| | NA (Calculation failed) |

| Median | A | B | C | D | E | F | G | H | I | J | K | L | M | N |
|---|---|---|---|---|---|---|---|---|---|---|---|---|---|---|
| A | - | 3.645 | 1.969 | 1.497 | 4.432 | 2.739 | 1.567 | 2.632 | 2.064 | 5.259 | 3.363 | 2.414 | 3.054 | 0.771 |
| B | - | - | 2.923 | 3.111 | 4.919 | 4.155 | 3.144 | 2.487 | 2.662 | 4.516 | 0.854 | 3.979 | 1.889 | 3.1 |
| C | - | - | - | 0.236 | 4.618 | 3.475 | 0.532 | 1.09 | 0.597 | 5 | 2.644 | 3.147 | 2.066 | -0.125 |
| D | - | - | - | - | 4.562 | 3.322 | -2.24 | 1.729 | 1.28 | 5.154 | 2.901 | 2.965 | 2.43 | -4.506 |
| E | - | - | - | - | - | 4.133 | 4.933 | 5.144 | 5.108 | 6.263 | 5.35 | 4.375 | 5.249 | 4.969 |
| F | - | - | - | - | - | - | 3.521 | 4.028 | 4.177 | 5.602 | 4.418 | 1.984 | 4.226 | 3.926 |
| G | - | - | - | - | - | - | - | 1.905 | 1.769 | 5.141 | 3.001 | 2.888 | 2.543 | -0.066 |
| H | - | - | - | - | - | - | - | - | -1.916 | 4.911 | 2.067 | 3.422 | 1.009 | 1.068 |
| I | - | - | - | - | - | - | - | - | - | 5.089 | 2.559 | 3.196 | 1.975 | 1.247 |
| J | - | - | - | - | - | - | - | - | - | - | 4.246 | 4.954 | 4.401 | 4.556 |
| K | - | - | - | - | - | - | - | - | - | - | - | 3.825 | 0.94 | 2.865 |
| L | - | - | - | - | - | - | - | - | - | - | - | - | 3.841 | 3.06 |
| M | - | - | - | - | - | - | - | - | - | - | - | - | - | 2.369 |
| N | - | - | - | - | - | - | - | - | - | - | - | - | - | - |

| Max | A | B | C | D | E | F | G | H | I | J | K | L | M | N |
|---|---|---|---|---|---|---|---|---|---|---|---|---|---|---|
| A | - | 4.168 | 3.342 | 2.876 | 4.162 | 2.361 | 3.245 | 4.124 | 2.086 | 5.672 | 3.722 | 1.025 | 3.701 | 2.461 |
| B | - | - | 2.807 | 3.149 | 4.888 | 3.508 | 2.824 | -0.226 | 3.602 | 4.98 | 1.908 | 4.047 | 2.08 | 3.401 |
| C | - | - | - | 1.12 | 4.615 | 2.175 | -2.24 | 2.704 | 2.533 | 5.243 | 1.758 | 3.359 | 1.553 | 2.013 |
| D | - | - | - | - | 4.498 | 1.445 | 0.936 | 3.138 | 1.668 | 5.615 | 2.493 | 3.075 | 2.335 | 1.142 |
| E | - | - | - | - | - | 4.666 | 4.988 | 5.355 | 4.688 | 6.503 | 5.192 | 4.186 | 5.155 | 4.775 |
| F | - | - | - | - | - | - | 2.037 | 3.281 | 2.206 | 5.457 | 3.052 | 2.63 | 2.763 | 0.285 |
| G | - | - | - | - | - | - | - | 2.72 | 2.419 | 5.476 | 1.978 | 3.337 | 1.579 | 1.902 |
| H | - | - | - | - | - | - | - | - | 3.538 | 5.194 | 1.059 | 3.948 | 1.718 | 3.331 |
| I | - | - | - | - | - | - | - | - | - | 5.661 | 3.18 | 2.64 | 3.05 | 0.237 |
| J | - | - | - | - | - | - | - | - | - | - | 4.75 | 5.339 | 4.977 | 4.926 |
| K | - | - | - | - | - | - | - | - | - | - | - | 3.718 | -0.4 | 2.768 |
| L | - | - | - | - | - | - | - | - | - | - | - | - | 3.896 | 2.792 |
| M | - | - | - | - | - | - | - | - | - | - | - | - | - | 2.732 |
| N | - | - | - | - | - | - | - | - | - | - | - | - | - | - |
