## Supplementary Data S2 for "Interlaboratory comparison of standardised metabolomics and lipidomics analyses in human and rodent blood using the MxP® Quant 500 kit": TG(22.0_32.4).pdf

**Amount of relative differences between laboratories**

| Min | A | B | C | D | E | F | G | H | I | J | K | L | M | N |
|---|---|---|---|---|---|---|---|---|---|---|---|---|---|---|
| A | - | 5.374 | 5.681 | 4.85 | NA | 7.722 | 5.229 | 6.384 | 4.446 | 6.132 | 4.956 | 5.583 | NA | 1.956 |
| B | - | - | 4.306 | 7.062 | NA | 8.567 | 7.141 | 6.248 | 6.753 | 5.951 | 2.653 | 3.529 | NA | 6.086 |
| C | - | - | - | 6.788 | NA | 8.013 | 7.212 | 5.803 | 6.651 | 5.102 | 5.065 | 5.399 | NA | 6.386 |
| D | - | - | - | - | NA | 8.358 | 0.884 | 7.156 | 2.874 | 6.144 | 5.404 | 6.17 | NA | 4.769 |
| E | - | - | - | - | - | NA | NA | NA | NA | NA | NA | NA | NA | NA |
| F | - | - | - | - | - | - | 6.569 | 4.57 | 5.746 | 5.053 | 1.418 | 5.412 | NA | 5.456 |
| G | - | - | - | - | - | - | - | 6.774 | 3.391 | 6.043 | 5.623 | 5.969 | NA | 5.056 |
| H | - | - | - | - | - | - | - | - | 6.722 | 6.295 | 3.607 | 6.399 | NA | 6.602 |
| I | - | - | - | - | - | - | - | - | - | 6.023 | 5.473 | 5.671 | NA | 4.493 |
| J | - | - | - | - | - | - | - | - | - | - | 4.054 | -4.04 | NA | 2.889 |
| K | - | - | - | - | - | - | - | - | - | - | - | 5.785 | NA | 5.573 |
| L | - | - | - | - | - | - | - | - | - | - | - | - | NA | 5.214 |
| M | - | - | - | - | - | - | - | - | - | - | - | - | - | NA |
| N | - | - | - | - | - | - | - | - | - | - | - | - | - | - |

| | |
|---|---|
| | Larger than 10 |
| | Between 8 and 9,99 |
| | Between 6 and 7,99 |
| | Between 4 and 5,99 |
| | Between 2 and 3,99 |
| | Between 0 and 1,99 |
| | NA (Calculation failed) |

| Median | A | B | C | D | E | F | G | H | I | J | K | L | M | N |
|---|---|---|---|---|---|---|---|---|---|---|---|---|---|---|
| A | - | 5.504 | 5.056 | 4.02 | NA | 3.414 | 4.286 | 3.855 | 3.693 | 3.55 | 3.759 | 4.962 | NA | 2.976 |
| B | - | - | 4.657 | 6.863 | NA | 6.85 | 6.915 | 5.818 | 6.701 | 6.87 | 5.294 | 5.114 | NA | 5.981 |
| C | - | - | - | 6.14 | NA | 5.826 | 6.338 | 4.051 | 5.996 | 6.377 | 5.205 | 0.923 | NA | 5.344 |
| D | - | - | - | - | NA | 2.583 | 1.521 | 4.789 | 1.665 | 3.112 | 4.354 | 5.231 | NA | 4.528 |
| E | - | - | - | - | - | NA | NA | NA | NA | NA | NA | NA | NA | NA |
| F | - | - | - | - | - | - | 4.457 | 4.828 | 3.214 | 4.216 | 3.873 | 5.106 | NA | 1.697 |
| G | - | - | - | - | - | - | - | 4.607 | 2.735 | 2.918 | 4.735 | 5.239 | NA | 4.791 |
| H | - | - | - | - | - | - | - | - | 5.628 | 5.987 | 2.336 | 3.434 | NA | 5.086 |
| I | - | - | - | - | - | - | - | - | - | 3.532 | 4.529 | 5.082 | NA | 4.416 |
| J | - | - | - | - | - | - | - | - | - | - | 4.614 | 5.536 | NA | 4.539 |
| K | - | - | - | - | - | - | - | - | - | - | - | 3.624 | NA | 3.412 |
| L | - | - | - | - | - | - | - | - | - | - | - | - | NA | 4.624 |
| M | - | - | - | - | - | - | - | - | - | - | - | - | - | NA |
| N | - | - | - | - | - | - | - | - | - | - | - | - | - | - |

| Max | A | B | C | D | E | F | G | H | I | J | K | L | M | N |
|---|---|---|---|---|---|---|---|---|---|---|---|---|---|---|
| A | - | 5.624 | 3.566 | 2.168 | NA | 7.113 | -0.35 | 6.113 | -4.144 | 6.823 | 4.944 | 4.608 | NA | 4.477 |
| B | - | - | 6.298 | 6.463 | NA | 8.645 | 6.636 | 7.761 | 6.621 | 8.404 | 7.272 | 5.459 | NA | 5.84 |
| C | - | - | - | 2.306 | NA | 7.896 | 3.433 | 6.705 | 3.336 | 7.166 | 5.509 | 3.85 | NA | 3.592 |
| D | - | - | - | - | NA | 7.489 | 1.976 | 6.803 | 1.546 | 6.742 | 5.129 | 4.563 | NA | 4.163 |
| E | - | - | - | - | - | NA | NA | NA | NA | NA | NA | NA | NA | NA |
| F | - | - | - | - | - | - | 5.755 | 5.089 | 5.8 | -1.067 | 5.492 | 6.036 | NA | 5.931 |
| G | - | - | - | - | - | - | - | 6.299 | 0.053 | 6.625 | 4.945 | 4.79 | NA | 4.381 |
| H | - | - | - | - | - | - | - | - | 5.033 | 5.481 | 3.704 | 5.625 | NA | 5.441 |
| I | - | - | - | - | - | - | - | - | - | 6.735 | 4.951 | 4.752 | NA | 4.314 |
| J | - | - | - | - | - | - | - | - | - | - | 5.473 | 6.027 | NA | 5.926 |
| K | - | - | - | - | - | - | - | - | - | - | - | 5.31 | NA | 5.116 |
| L | - | - | - | - | - | - | - | - | - | - | - | - | NA | 3.231 |
| M | - | - | - | - | - | - | - | - | - | - | - | - | - | NA |
| N | - | - | - | - | - | - | - | - | - | - | - | - | - | - |
