## Supplementary Data S2 for "Interlaboratory comparison of standardised metabolomics and lipidomics analyses in human and rodent blood using the MxP® Quant 500 kit": TG(22.4_32.0).pdf

### Amount of relative differences between laboratories

| Min | A | B | C | D | E | F | G | H | I | J | K | L | M | N |
|---|---|---|---|---|---|---|---|---|---|---|---|---|---|---|
| A | - | 1.515 | 0.372 | 6.575 | 7.014 | 5.878 | -1.034 | 5.708 | 5.051 | 7.374 | 5.13 | -0.355 | 4.89 | 4.886 |
| B | - | - | 2.931 | 6.09 | 6.598 | 5.851 | 1.517 | 5.665 | 4.814 | 7.01 | 5.294 | 2.203 | 4.967 | 4.446 |
| C | - | - | - | 5.878 | 6.533 | 5.678 | 3.873 | 5.495 | 3.969 | 6.961 | 4.926 | 3.991 | 4.593 | 3.592 |
| D | - | - | - | - | 5.068 | 5.105 | 5.83 | 4.587 | 4.004 | 5.747 | 4.174 | 5.902 | 1.175 | 4.963 |
| E | - | - | - | - | - | 3.354 | 6.838 | 2.035 | 5.894 | 4.58 | 3.417 | 6.9 | 5.04 | 6.342 |
| F | - | - | - | - | - | - | 7.09 | 3.937 | 6.288 | 3.296 | 4.218 | 7.143 | 5.464 | 6.699 |
| G | - | - | - | - | - | - | - | 5.705 | 4.94 | 7.031 | 5.318 | -0.461 | 5.038 | 4.641 |
| H | - | - | - | - | - | - | - | - | 5.717 | 5.386 | 2.952 | 6.784 | 4.831 | 6.183 |
| I | - | - | - | - | - | - | - | - | - | 7.347 | 3.32 | 5.56 | -1.246 | 2.891 |
| J | - | - | - | - | - | - | - | - | - | - | 4.281 | 6.137 | -1.981 | 5.547 |
| K | - | - | - | - | - | - | - | - | - | - | - | 6.588 | 4.282 | 5.978 |
| L | - | - | - | - | - | - | - | - | - | - | - | - | 4.968 | 4.861 |
| M | - | - | - | - | - | - | - | - | - | - | - | - | - | 4.205 |
| N | - | - | - | - | - | - | - | - | - | - | - | - | - | - |

| Median | A | B | C | D | E | F | G | H | I | J | K | L | M | N |
|---|---|---|---|---|---|---|---|---|---|---|---|---|---|---|
| A | - | 4.191 | 4.164 | 2.359 | 4.155 | 4.473 | 4.132 | 4.337 | 4.567 | 4.999 | 4.139 | 2.519 | 3.884 | -1.906 |
| B | - | - | 0.809 | 3.413 | 4.941 | 5.085 | -1.872 | 5.009 | 5.146 | 3.294 | 4.845 | 3.403 | 4.801 | 3.927 |
| C | - | - | - | 3.418 | 5.015 | 4.974 | 1.031 | 4.939 | 5.048 | 2.645 | 4.523 | 3.172 | 4.697 | 3.795 |
| D | - | - | - | - | 4.588 | 4.674 | 3.549 | 4.573 | 4.81 | 4.417 | 4.053 | 0.19 | 4.318 | 2.218 |
| E | - | - | - | - | - | 0.785 | 5.563 | -0.799 | 2.383 | 5.904 | 3.417 | 4.922 | 2.463 | 4.691 |
| F | - | - | - | - | - | - | 5.615 | 1.017 | 2.077 | 5.911 | 3.783 | 4.961 | 2.687 | 4.787 |
| G | - | - | - | - | - | - | - | 5.001 | 5.138 | 3.488 | 4.842 | 3.387 | 4.792 | 3.909 |
| H | - | - | - | - | - | - | - | - | 2.494 | 5.867 | 3.247 | 4.891 | 2.325 | 4.649 |
| I | - | - | - | - | - | - | - | - | - | 6.166 | 3.219 | 5.262 | 3.542 | 4.954 |
| J | - | - | - | - | - | - | - | - | - | - | 4.741 | 4.141 | 5.046 | 4.269 |
| K | - | - | - | - | - | - | - | - | - | - | - | 4.111 | 2.29 | 3.759 |
| L | - | - | - | - | - | - | - | - | - | - | - | - | 4.278 | 2.489 |
| M | - | - | - | - | - | - | - | - | - | - | - | - | - | 4.163 |
| N | - | - | - | - | - | - | - | - | - | - | - | - | - | - |

| Max | A | B | C | D | E | F | G | H | I | J | K | L | M | N |
|---|---|---|---|---|---|---|---|---|---|---|---|---|---|---|
| A | - | 5.024 | 5.274 | 5.929 | 4.83 | 5.529 | 4.876 | 4.527 | 4.252 | 7.6 | 5.389 | 3.054 | -2.056 | 4.158 |
| B | - | - | 2.198 | 4.409 | 1.711 | 3.385 | 1.292 | 2.843 | 5.284 | 6.947 | 2.842 | 4.207 | 4.613 | 3.477 |
| C | - | - | - | 4.002 | 2.871 | 2.448 | 2.746 | 3.49 | 5.38 | 6.825 | 1.214 | 4.457 | 4.795 | 3.908 |
| D | - | - | - | - | 4.329 | 3.156 | 4.296 | 4.554 | 5.635 | 6.373 | 3.541 | 5.036 | 5.239 | 4.744 |
| E | - | - | - | - | - | 3.823 | -0.531 | 2.064 | 5.208 | 7.006 | 3.415 | 3.998 | 4.465 | 3.058 |
| F | - | - | - | - | - | - | 3.527 | 3.993 | 5.481 | 6.664 | 1.491 | 4.706 | 4.98 | 4.289 |
| G | - | - | - | - | - | - | - | 2.28 | 5.226 | 6.995 | 3.283 | 4.025 | 4.493 | 3.152 |
| H | - | - | - | - | - | - | - | - | 5.097 | 7.133 | 3.953 | 3.606 | 4.219 | 2.106 |
| I | - | - | - | - | - | - | - | - | - | 8.044 | 6.236 | 5.078 | 4.574 | 5.51 |
| J | - | - | - | - | - | - | - | - | - | - | 5.692 | 5.986 | 6.046 | 5.909 |
| K | - | - | - | - | - | - | - | - | - | - | - | 4.57 | 4.879 | 4.092 |
| L | - | - | - | - | - | - | - | - | - | - | - | - | 2.89 | 3.15 |
| M | - | - | - | - | - | - | - | - | - | - | - | - | - | 4.137 |
| N | - | - | - | - | - | - | - | - | - | - | - | - | - | - |

| | |
|---|---|
| | Larger than 10 |
| | Between 8 and 9,99 |
| | Between 6 and 7,99 |
| | Between 4 and 5,99 |
| | Between 2 and 3,99 |
| | Between 0 and 1,99 |
| | NA (Calculation failed) |
