## Supplementary Data S2 for "Interlaboratory comparison of standardised metabolomics and lipidomics analyses in human and rodent blood using the MxP® Quant 500 kit": TG(22.5_32.1).pdf

**Amount of relative differences between laboratories**

| Min | A | B | C | D | E | F | G | H | I | J | K | L | M | N |
|---|---|---|---|---|---|---|---|---|---|---|---|---|---|---|
| A | - | 4.838 | 4.732 | 4.443 | NA | 5.729 | 4.082 | 3.72 | 4.221 | 6.266 | 5.06 | 4.208 | 4.317 | 5.262 |
| B | - | - | 3.072 | 3.227 | NA | 4.036 | 4.015 | 3.197 | 4.839 | 5.317 | 1.801 | 4.004 | 2.923 | 1.607 |
| C | - | - | - | -1.323 | NA | 4.439 | 2.736 | -0.522 | 4.276 | 5.414 | 3.443 | 2.741 | 0.609 | 3.32 |
| D | - | - | - | - | NA | 4.528 | 2.651 | -1.039 | 4.175 | 5.485 | 3.541 | 2.629 | 0.733 | 3.448 |
| E | - | - | - | - | - | NA | NA | NA | NA | NA | NA | NA | NA | NA |
| F | - | - | - | - | - | - | 4.79 | 3.898 | 5.582 | 4.353 | 3.04 | 4.836 | 4.049 | 3.499 |
| G | - | - | - | - | - | - | - | 2.246 | 3.219 | 5.65 | 4.088 | -1.71 | 2.785 | 4.13 |
| H | - | - | - | - | - | - | - | - | 3.466 | 5.733 | 3.881 | 1.501 | 1.693 | 4.022 |
| I | - | - | - | - | - | - | - | - | - | 5.981 | 4.434 | 2.268 | 3.177 | 4.608 |
| J | - | - | - | - | - | - | - | - | - | - | 3.435 | 4.895 | 3.728 | 4.222 |
| K | - | - | - | - | - | - | - | - | - | - | - | 4.363 | 3.681 | 0.049 |
| L | - | - | - | - | - | - | - | - | - | - | - | - | 2.558 | 4.201 |
| M | - | - | - | - | - | - | - | - | - | - | - | - | - | 3.422 |
| N | - | - | - | - | - | - | - | - | - | - | - | - | - | - |

| | |
|---|---|
| | Larger than 10 |
| | Between 8 and 9,99 |
| | Between 6 and 7,99 |
| | Between 4 and 5,99 |
| | Between 2 and 3,99 |
| | Between 0 and 1,99 |
| | NA (Calculation failed) |

| Median | A | B | C | D | E | F | G | H | I | J | K | L | M | N |
|---|---|---|---|---|---|---|---|---|---|---|---|---|---|---|
| A | - | 3.484 | 0.473 | 1.908 | NA | 3.604 | -1.962 | 2.658 | 2.168 | 4.089 | 3.977 | 1.973 | 2.828 | 4.066 |
| B | - | - | 3.486 | 3.309 | NA | -1.755 | 3.695 | 2.515 | 2.713 | 4.803 | 3.008 | 3.152 | 2.659 | 2.765 |
| C | - | - | - | 0.048 | NA | 3.356 | 0.641 | 2.296 | 2.095 | 3.869 | 4.133 | 1.031 | 2.08 | 4.02 |
| D | - | - | - | - | NA | 3.212 | 1.459 | 1.956 | 1.624 | 4.013 | 4.039 | -0.086 | 1.697 | 3.925 |
| E | - | - | - | - | - | NA | NA | NA | NA | NA | NA | NA | NA | NA |
| F | - | - | - | - | - | - | 3.661 | 2.477 | 2.532 | 4.701 | 3.272 | 3.168 | 2.802 | 2.869 |
| G | - | - | - | - | - | - | - | 2.68 | 2.462 | 3.806 | 4.182 | 1.88 | 2.599 | 4.119 |
| H | - | - | - | - | - | - | - | - | 0.228 | 4.47 | 3.593 | 1.528 | -2.805 | 3.538 |
| I | - | - | - | - | - | - | - | - | - | 4.411 | 3.655 | 0.335 | 0.762 | 3.666 |
| J | - | - | - | - | - | - | - | - | - | - | 4.831 | 3.275 | 3.362 | 4.607 |
| K | - | - | - | - | - | - | - | - | - | - | - | 4.136 | 3.869 | -0.337 |
| L | - | - | - | - | - | - | - | - | - | - | - | - | 1.407 | 3.813 |
| M | - | - | - | - | - | - | - | - | - | - | - | - | - | 3.604 |
| N | - | - | - | - | - | - | - | - | - | - | - | - | - | - |

| Max | A | B | C | D | E | F | G | H | I | J | K | L | M | N |
|---|---|---|---|---|---|---|---|---|---|---|---|---|---|---|
| A | - | 3.47 | 4.757 | 4.522 | NA | 4.819 | 4.401 | 3.156 | 1.949 | 7.11 | 0.999 | 3.607 | 3.961 | -2.273 |
| B | - | - | 3.844 | 3.423 | NA | 3.964 | 3.178 | 0.949 | 2.695 | 6.849 | 3.536 | -0.056 | 2.025 | 3.284 |
| C | - | - | - | 1.662 | NA | 0.101 | 2.213 | 3.833 | 4.188 | 6.467 | 4.508 | 3.549 | 3.171 | 4.399 |
| D | - | - | - | - | NA | 2.127 | 0.619 | 3.511 | 3.961 | 6.565 | 4.335 | 3.143 | 2.593 | 4.213 |
| E | - | - | - | - | - | NA | NA | NA | NA | NA | NA | NA | NA | NA |
| F | - | - | - | - | - | - | 2.482 | 3.919 | 4.254 | 6.424 | 4.561 | 3.663 | 3.316 | 4.452 |
| G | - | - | - | - | - | - | - | 3.335 | 3.834 | 6.596 | 4.252 | 2.89 | 2.199 | 4.111 |
| H | - | - | - | - | - | - | - | - | 2.221 | 6.899 | 3.319 | 1.558 | 2.611 | 2.997 |
| I | - | - | - | - | - | - | - | - | - | 7.024 | 2.476 | 2.998 | 3.497 | 1.817 |
| J | - | - | - | - | - | - | - | - | - | - | 5.886 | 5.737 | 5.696 | 5.86 |
| K | - | - | - | - | - | - | - | - | - | - | - | 3.835 | 4.15 | 1.152 |
| L | - | - | - | - | - | - | - | - | - | - | - | - | 1.616 | 3.414 |
| M | - | - | - | - | - | - | - | - | - | - | - | - | - | 3.732 |
| N | - | - | - | - | - | - | - | - | - | - | - | - | - | - |
