## Supplementary Data S2 for "Interlaboratory comparison of standardised metabolomics and lipidomics analyses in human and rodent blood using the MxP® Quant 500 kit": TG(22.5_34.1).pdf

**Amount of relative differences between laboratories**

| Min | A | B | C | D | E | F | G | H | I | J | K | L | M | N |
|---|---|---|---|---|---|---|---|---|---|---|---|---|---|---|
| A | - | 4.093 | 3.715 | 3.593 | 5.177 | 4.26 | 2.925 | 2.453 | 2.546 | 6.729 | 3.787 | 4.202 | 3.817 | 4.922 |
| B | - | - | 2.713 | 2.858 | 4.358 | -2.307 | 3.79 | 3.991 | 4.293 | 6.246 | 2.987 | 0.943 | 2.031 | 2.728 |
| C | - | - | - | -0.898 | 4.656 | 2.529 | 2.792 | 3.131 | 3.695 | 6.138 | 0.843 | 2.024 | 1.352 | 3.448 |
| D | - | - | - | - | 4.661 | 2.582 | 2.716 | 3.053 | 3.73 | 6.061 | 0.755 | 2.042 | 1.662 | 3.284 |
| E | - | - | - | - | - | 4.674 | 5.381 | 5.445 | 5.743 | 4.843 | 5.2 | 4.835 | 4.773 | 4.718 |
| F | - | - | - | - | - | - | 3.681 | 3.841 | 4.48 | 5.354 | 3.181 | 1.274 | 0.456 | -0.543 |
| G | - | - | - | - | - | - | - | 0.767 | 2.538 | 6.128 | 2.21 | 3.352 | 3.156 | 4.028 |
| H | - | - | - | - | - | - | - | - | 1.718 | 6.263 | 2.761 | 3.587 | 3.354 | 4.264 |
| I | - | - | - | - | - | - | - | - | - | 6.834 | 3.055 | 3.699 | 2.985 | 4.75 |
| J | - | - | - | - | - | - | - | - | - | - | 3.3 | 1.07 | 1.579 | 3.07 |
| K | - | - | - | - | - | - | - | - | - | - | - | 2.502 | 1.993 | 3.666 |
| L | - | - | - | - | - | - | - | - | - | - | - | - | 0.875 | 2.931 |
| M | - | - | - | - | - | - | - | - | - | - | - | - | - | 3.241 |
| N | - | - | - | - | - | - | - | - | - | - | - | - | - | - |

| | |
|---|---|
| | Larger than 10 |
| | Between 8 and 9,99 |
| | Between 6 and 7,99 |
| | Between 4 and 5,99 |
| | Between 2 and 3,99 |
| | Between 0 and 1,99 |
| | NA (Calculation failed) |

| Median | A | B | C | D | E | F | G | H | I | J | K | L | M | N |
|---|---|---|---|---|---|---|---|---|---|---|---|---|---|---|
| A | - | 3.674 | 2.64 | 2.388 | 4.635 | 1.825 | -2.107 | 0.072 | 2.903 | 1.97 | 2.192 | 3.448 | 3.166 | 3.997 |
| B | - | - | 2.969 | 3.256 | 3.751 | 3.326 | 3.961 | 4.007 | 2.74 | 4.342 | 3.249 | 1.458 | 2.22 | 1.812 |
| C | - | - | - | 0.76 | 4.303 | 0.92 | 2.856 | 2.932 | 0.004 | 3.59 | 0.652 | 2.222 | 1.561 | 3.393 |
| D | - | - | - | - | 4.402 | -3.167 | 2.432 | 2.508 | 1.435 | 3.286 | -1.656 | 2.656 | 2.221 | 3.599 |
| E | - | - | - | - | - | 4.688 | 5.078 | 5.078 | 4.557 | 5.325 | 4.708 | 4.243 | 4.375 | 3.495 |
| F | - | - | - | - | - | - | 2.675 | 2.674 | 0.829 | 3.557 | -1.54 | 2.434 | 1.993 | 3.527 |
| G | - | - | - | - | - | - | - | -1.533 | 2.941 | 2.245 | 2.398 | 3.472 | 3.249 | 4.048 |
| H | - | - | - | - | - | - | - | - | 3.015 | 1.999 | 2.496 | 3.52 | 3.298 | 4.075 |
| I | - | - | - | - | - | - | - | - | - | 3.551 | 1.673 | 1.881 | 0.629 | 3.192 |
| J | - | - | - | - | - | - | - | - | - | - | 3.546 | 4.024 | 3.955 | 4.477 |
| K | - | - | - | - | - | - | - | - | - | - | - | 2.613 | 2.122 | 3.574 |
| L | - | - | - | - | - | - | - | - | - | - | - | - | 0.876 | 2.607 |
| M | - | - | - | - | - | - | - | - | - | - | - | - | - | 2.969 |
| N | - | - | - | - | - | - | - | - | - | - | - | - | - | - |

| Max | A | B | C | D | E | F | G | H | I | J | K | L | M | N |
|---|---|---|---|---|---|---|---|---|---|---|---|---|---|---|
| A | - | -5.506 | 3.459 | 4.096 | 2.408 | 5.042 | 4.316 | 4.042 | 3.253 | 6.955 | 3.613 | 2.171 | 2.676 | -2.026 |
| B | - | - | 3.46 | 4.093 | 2.401 | 5.044 | 4.317 | 4.044 | 3.251 | 6.969 | 3.631 | 2.152 | 2.673 | -3.342 |
| C | - | - | - | 2.452 | 2.359 | 4.307 | 3.007 | 2.309 | 4.209 | 6.674 | 0.221 | 2.561 | 2.052 | 3.33 |
| D | - | - | - | - | 3.33 | 3.768 | 1.288 | -1.065 | 4.506 | 6.544 | 1.989 | 3.432 | 3.195 | 3.879 |
| E | - | - | - | - | - | 4.719 | 3.799 | 3.413 | 3.818 | 6.848 | 2.764 | -0.281 | 0.003 | 2.377 |
| F | - | - | - | - | - | - | 3.293 | 3.632 | 4.999 | 6.115 | 3.958 | 4.427 | 4.324 | 4.638 |
| G | - | - | - | - | - | - | - | 1.515 | 4.619 | 6.432 | 2.687 | 3.692 | 3.498 | 4.069 |
| H | - | - | - | - | - | - | - | - | 4.481 | 6.528 | 1.835 | 3.372 | 3.118 | 3.836 |
| I | - | - | - | - | - | - | - | - | - | 7.221 | 4.593 | 3.951 | 4.135 | 3.368 |
| J | - | - | - | - | - | - | - | - | - | - | 5.669 | 5.757 | 5.73 | 5.803 |
| K | - | - | - | - | - | - | - | - | - | - | - | 2.818 | 2.451 | 3.466 |
| L | - | - | - | - | - | - | - | - | - | - | - | - | 0.881 | 2.143 |
| M | - | - | - | - | - | - | - | - | - | - | - | - | - | 2.597 |
| N | - | - | - | - | - | - | - | - | - | - | - | - | - | - |
