## Supplementary Data S2 for "Interlaboratory comparison of standardised metabolomics and lipidomics analyses in human and rodent blood using the MxP® Quant 500 kit": TG(22.6_34.2).pdf

**Amount of relative differences between laboratories**

| Min | A | B | C | D | E | F | G | H | I | J | K | L | M | N |
|---|---|---|---|---|---|---|---|---|---|---|---|---|---|---|
| A | - | 4.942 | 5.124 | 3.114 | 5.227 | 5.379 | 4.607 | 4.949 | 3.665 | 6.136 | 4.26 | 3.672 | 5.146 | 4.226 |
| B | - | - | 0.724 | 4.717 | 3.585 | 3.384 | 3.863 | 2.065 | 4.143 | 5.015 | 4.038 | 4.228 | -3.902 | 3.564 |
| C | - | - | - | 4.315 | 3.989 | 3.638 | 3.57 | 0.565 | 3.435 | 4.908 | 3.596 | 3.681 | 0.962 | 2.448 |
| D | - | - | - | - | 5.001 | 5.045 | 3.776 | 4.42 | 2.425 | 5.86 | 3.259 | 2.402 | 4.675 | 3.475 |
| E | - | - | - | - | - | 0.55 | 4.822 | 3.859 | 5.145 | 5.274 | 4.995 | 5.263 | 3.442 | 4.806 |
| F | - | - | - | - | - | - | 4.78 | 3.955 | 4.599 | 3.958 | 4.747 | 4.722 | 3.587 | 4.229 |
| G | - | - | - | - | - | - | - | 3.196 | 0.962 | 5.445 | 0.437 | 1.718 | 3.674 | 1.712 |
| H | - | - | - | - | - | - | - | - | 3.464 | 5.043 | 3.496 | 3.688 | 1.885 | 2.475 |
| I | - | - | - | - | - | - | - | - | - | 5.994 | 2.254 | -0.163 | 4.452 | 2.674 |
| J | - | - | - | - | - | - | - | - | - | - | 5.104 | 4.838 | 4.4 | 3.925 |
| K | - | - | - | - | - | - | - | - | - | - | - | 2.458 | 4.161 | 0.719 |
| L | - | - | - | - | - | - | - | - | - | - | - | - | 4.64 | 2.592 |
| M | - | - | - | - | - | - | - | - | - | - | - | - | - | 3.514 |
| N | - | - | - | - | - | - | - | - | - | - | - | - | - | - |

| Median | A | B | C | D | E | F | G | H | I | J | K | L | M | N |
|---|---|---|---|---|---|---|---|---|---|---|---|---|---|---|
| A | - | 3.561 | -2.887 | 0.061 | 4.855 | 3.15 | 1.358 | 1.984 | 2.862 | 3.366 | 3.492 | 3.437 | 3.614 | 3.949 |
| B | - | - | 3.716 | 3.699 | 4.264 | 1.994 | 4.03 | 3.2 | 2.427 | 4.642 | -1.338 | 0.086 | -4.569 | 1.916 |
| C | - | - | - | -1.794 | 4.828 | 2.952 | 1.58 | 1.703 | 2.733 | 3.384 | 3.487 | 3.395 | 3.506 | 3.883 |
| D | - | - | - | - | 4.828 | 2.989 | 1.754 | 1.64 | 2.691 | 3.477 | 3.409 | 3.346 | 3.511 | 3.874 |
| E | - | - | - | - | - | 4.833 | 5.468 | 5.133 | 4.927 | 5.772 | 4.62 | 4.667 | 4.568 | 4.251 |
| F | - | - | - | - | - | - | 3.519 | 2.25 | 0.132 | 4.29 | 1.988 | 1.669 | 1.992 | 2.935 |
| G | - | - | - | - | - | - | - | 2.617 | 3.236 | 2.849 | 3.778 | 3.713 | 3.813 | 4.113 |
| H | - | - | - | - | - | - | - | - | 1.806 | 3.839 | 3.023 | 2.897 | 3.073 | 3.571 |
| I | - | - | - | - | - | - | - | - | - | 4.205 | 2.144 | 1.956 | 2.402 | 3.136 |
| J | - | - | - | - | - | - | - | - | - | - | 4.28 | 4.212 | 4.239 | 4.491 |
| K | - | - | - | - | - | - | - | - | - | - | - | -0.885 | -0.16 | 2.187 |
| L | - | - | - | - | - | - | - | - | - | - | - | - | 0.632 | 2.341 |
| M | - | - | - | - | - | - | - | - | - | - | - | - | - | 1.917 |
| N | - | - | - | - | - | - | - | - | - | - | - | - | - | - |

| Max | A | B | C | D | E | F | G | H | I | J | K | L | M | N |
|---|---|---|---|---|---|---|---|---|---|---|---|---|---|---|
| A | - | 2.105 | 3.552 | 1.256 | 4.676 | 2.475 | 3.563 | 2.636 | 2.248 | 5.37 | 3.132 | 3.332 | 2.157 | 3.814 |
| B | - | - | 4.083 | 2.806 | 4.464 | 3.37 | 4.083 | 3.469 | -1.583 | 5.556 | 2.21 | 2.573 | -4.887 | 3.372 |
| C | - | - | - | 3.097 | 5.061 | 2.501 | -3.986 | 2.335 | 3.884 | 4.702 | 4.211 | 4.305 | 3.861 | 4.541 |
| D | - | - | - | - | 4.753 | 1.642 | 3.208 | 1.922 | 2.799 | 5.24 | 3.453 | 3.597 | 2.733 | 4.013 |
| E | - | - | - | - | - | 5.383 | 5.636 | 5.421 | 4.813 | 6.502 | 4.479 | 4.34 | 4.81 | 3.933 |
| F | - | - | - | - | - | - | 2.576 | -0.524 | 3.273 | 5.048 | 3.768 | 3.897 | 3.225 | 4.231 |
| G | - | - | - | - | - | - | - | 2.32 | 3.879 | 4.742 | 4.205 | 4.294 | 3.857 | 4.538 |
| H | - | - | - | - | - | - | - | - | 3.358 | 5.023 | 3.829 | 3.955 | 3.322 | 4.266 |
| I | - | - | - | - | - | - | - | - | - | 5.647 | 2.108 | 2.529 | -1.788 | 3.294 |
| J | - | - | - | - | - | - | - | - | - | - | 5.181 | 5.218 | 5.015 | 5.263 |
| K | - | - | - | - | - | - | - | - | - | - | - | 0.508 | 2.267 | 2.531 |
| L | - | - | - | - | - | - | - | - | - | - | - | - | 2.657 | 2.221 |
| M | - | - | - | - | - | - | - | - | - | - | - | - | - | 3.349 |
| N | - | - | - | - | - | - | - | - | - | - | - | - | - | - |

| | |
|---|---|
| (red) | Larger than 10 |
| (orange) | Between 8 and 9,99 |
| (light orange) | Between 6 and 7,99 |
| (light green) | Between 4 and 5,99 |
| (green) | Between 2 and 3,99 |
| (dark green) | Between 0 and 1,99 |
| (grey) | NA (Calculation failed) |
