## Supplementary Data S2 for "Interlaboratory comparison of standardised metabolomics and lipidomics analyses in human and rodent blood using the MxP® Quant 500 kit": TG(22.6_34.3).pdf

**Amount of relative differences between laboratories**

| Min | A | B | C | D | E | F | G | H | I | J | K | L | M | N |
|---|---|---|---|---|---|---|---|---|---|---|---|---|---|---|
| A | - | 4.728 | 4.762 | 4.495 | 5.495 | 5.261 | 5.327 | 3.574 | 4.471 | 5.034 | 3.66 | 4.311 | 5.04 | 4.284 |
| B | - | - | 1.578 | 2.197 | 4.676 | 4.31 | 3.858 | 3.288 | 2.383 | 2.047 | 4.167 | 0.392 | 1.984 | 2.467 |
| C | - | - | - | 2.127 | 4.775 | 4.256 | 4.136 | 3.566 | 2.31 | 3.92 | 4.162 | 1.392 | 2.404 | 2.612 |
| D | - | - | - | - | 4.937 | 4.444 | 4.442 | 3.187 | -0.618 | 4.28 | 3.777 | 0.263 | 3.382 | 1.35 |
| E | - | - | - | - | - | 3.074 | 3.848 | 5.728 | 5.444 | 3.174 | 5.924 | 5.348 | 4.956 | 5.532 |
| F | - | - | - | - | - | - | 3.293 | 5.004 | 5.037 | 5.265 | 5.754 | 4.539 | 4.669 | 5.004 |
| G | - | - | - | - | - | - | - | 4.931 | 4.645 | 0.66 | 5.428 | 4.26 | 3.914 | 4.674 |
| H | - | - | - | - | - | - | - | - | 2.494 | 3.651 | 2.912 | 3.05 | 3.87 | 2.151 |
| I | - | - | - | - | - | - | - | - | - | 3.854 | 3.677 | -1.244 | 3.514 | 1.1 |
| J | - | - | - | - | - | - | - | - | - | - | 3.398 | 2.303 | 2.712 | 2.301 |
| K | - | - | - | - | - | - | - | - | - | - | - | 4.021 | 4.302 | 3.552 |
| L | - | - | - | - | - | - | - | - | - | - | - | - | 4.141 | 0.455 |
| M | - | - | - | - | - | - | - | - | - | - | - | - | - | 3.306 |
| N | - | - | - | - | - | - | - | - | - | - | - | - | - | - |

| | |
|---|---|
| | Larger than 10 |
| | Between 8 and 9,99 |
| | Between 6 and 7,99 |
| | Between 4 and 5,99 |
| | Between 2 and 3,99 |
| | Between 0 and 1,99 |
| | NA (Calculation failed) |

| Median | A | B | C | D | E | F | G | H | I | J | K | L | M | N |
|---|---|---|---|---|---|---|---|---|---|---|---|---|---|---|
| A | - | 2.186 | 3.318 | 3.476 | 5.098 | 2.168 | 4.047 | 1.301 | 3.65 | 3.775 | 4.229 | 4.501 | 3.211 | 3.741 |
| B | - | - | 2.569 | 2.75 | 4.968 | -0.885 | 3.719 | 2.954 | 3.128 | 4.188 | 4.548 | 4.255 | 2.281 | 3.335 |
| C | - | - | - | -0.193 | 4.754 | 2.57 | 2.92 | 3.856 | 1.518 | 4.666 | 4.978 | 3.801 | 0.193 | 2.054 |
| D | - | - | - | - | 4.716 | 2.82 | 2.715 | 3.951 | 0.915 | 4.746 | 5.043 | 3.72 | 1.033 | 1.618 |
| E | - | - | - | - | - | 5.488 | 4.734 | 5.826 | 5.044 | 6.158 | 6.322 | 4.158 | 5.27 | 4.967 |
| F | - | - | - | - | - | - | 3.699 | 3.26 | 3.022 | 4.2 | 4.576 | 4.19 | 1.909 | 3.32 |
| G | - | - | - | - | - | - | - | 4.616 | 2.389 | 5.146 | 5.404 | 2.744 | 3.293 | 1.857 |
| H | - | - | - | - | - | - | - | - | 3.932 | 3.273 | 3.88 | 4.638 | 3.553 | 4.038 |
| I | - | - | - | - | - | - | - | - | - | 4.886 | 5.153 | 3.512 | 1.988 | 0.208 |
| J | - | - | - | - | - | - | - | - | - | - | 2.139 | 4.93 | 4.184 | 4.568 |
| K | - | - | - | - | - | - | - | - | - | - | - | 5.106 | 4.516 | 4.765 |
| L | - | - | - | - | - | - | - | - | - | - | - | - | 4.114 | 3.607 |
| M | - | - | - | - | - | - | - | - | - | - | - | - | - | 2.452 |
| N | - | - | - | - | - | - | - | - | - | - | - | - | - | - |

| Max | A | B | C | D | E | F | G | H | I | J | K | L | M | N |
|---|---|---|---|---|---|---|---|---|---|---|---|---|---|---|
| A | - | 2.231 | 2.157 | 2.772 | 4.91 | 3.99 | 3.057 | 3.482 | 3.239 | 4.886 | 4.929 | 4.572 | -0.178 | 3.504 |
| B | - | - | 3.138 | 3.457 | 5.066 | 3.47 | 3.668 | 2.706 | 3.728 | 4.526 | 4.668 | 4.774 | 2.373 | 3.934 |
| C | - | - | - | 1.51 | 4.746 | 4.394 | 2.032 | 4.006 | 2.432 | 5.294 | 5.193 | 4.368 | 1.775 | 2.919 |
| D | - | - | - | - | 4.621 | 4.61 | 0.374 | 4.272 | 1.436 | 5.414 | 5.328 | 4.172 | 2.693 | 2.304 |
| E | - | - | - | - | - | 6.063 | 4.948 | 5.882 | 4.887 | 6.535 | 6.447 | 3.102 | 5.366 | 4.716 |
| F | - | - | - | - | - | - | 4.386 | 2.023 | 4.418 | 3.525 | 3.656 | 5.132 | 3.817 | 4.583 |
| G | - | - | - | - | - | - | - | 4.385 | 0.217 | 5.554 | 5.395 | 4.102 | 2.988 | 1.87 |
| H | - | - | - | - | - | - | - | - | 4.2 | 4.11 | 4.122 | 4.983 | 3.421 | 4.352 |
| I | - | - | - | - | - | - | - | - | - | 5.48 | 5.463 | 4.018 | 3.235 | 1.171 |
| J | - | - | - | - | - | - | - | - | - | - | 1.08 | 5.487 | 4.752 | 5.055 |
| K | - | - | - | - | - | - | - | - | - | - | - | 5.386 | 4.585 | 5.014 |
| L | - | - | - | - | - | - | - | - | - | - | - | - | 4.893 | 3.98 |
| M | - | - | - | - | - | - | - | - | - | - | - | - | - | 3.403 |
| N | - | - | - | - | - | - | - | - | - | - | - | - | - | - |
