## Supplementary Data S2 for "Interlaboratory comparison of standardised metabolomics and lipidomics analyses in human and rodent blood using the MxP® Quant 500 kit": Trigonelline.pdf

**Amount of relative differences between laboratories**

| Min | A | B | C | D | E | F | G | H | I | J | K | L | M | N |
|---|---|---|---|---|---|---|---|---|---|---|---|---|---|---|
| A | - | 5.157 | 4.341 | 1.832 | 3.216 | 4.713 | 4.04 | 3.455 | 1.196 | 3.306 | 3.033 | 3.014 | 4.43 | 5.161 |
| B | - | - | 4.703 | 5.155 | 5.865 | 4.197 | 4.665 | 4.76 | 5.224 | 5.561 | 4.666 | 5.643 | 4.516 | 4 |
| C | - | - | - | 3.728 | 4.817 | 2.842 | 0.585 | 4.833 | 4.365 | 4.73 | 2.959 | 4.534 | 1.548 | 3.864 |
| D | - | - | - | - | 3.786 | 4.523 | 3.607 | 2.936 | 1.879 | 3.708 | 2.29 | 3.482 | 4.161 | 4.862 |
| E | - | - | - | - | - | 5.053 | 4.52 | 4.18 | 2.931 | 2.232 | 3.524 | 0.038 | 4.807 | 5.458 |
| F | - | - | - | - | - | - | 2.964 | 4.484 | 4.518 | 4.975 | 3.564 | 4.983 | 2.12 | 2.695 |
| G | - | - | - | - | - | - | - | 3.746 | 4.108 | 4.721 | 2.015 | 4.748 | 1.958 | 3.949 |
| H | - | - | - | - | - | - | - | - | 6.216 | 6.238 | 7.28 | 7.015 | 7.888 | 8.194 |
| I | - | - | - | - | - | - | - | - | - | 0.686 | 4.792 | 3.753 | 5.713 | 6.072 |
| J | - | - | - | - | - | - | - | - | - | - | 4.176 | 1.645 | 5.092 | 5.649 |
| K | - | - | - | - | - | - | - | - | - | - | - | 4.175 | 3.381 | 4.546 |
| L | - | - | - | - | - | - | - | - | - | - | - | - | 4.952 | 5.53 |
| M | - | - | - | - | - | - | - | - | - | - | - | - | - | 3.45 |
| N | - | - | - | - | - | - | - | - | - | - | - | - | - | - |

| Median | A | B | C | D | E | F | G | H | I | J | K | L | M | N |
|---|---|---|---|---|---|---|---|---|---|---|---|---|---|---|
| A | - | 3.81 | 3.743 | -0.071 | 0.937 | 3.709 | 1.189 | 3.098 | 2.54 | 2.6 | 1.979 | 2.662 | 3.1 | 4.024 |
| B | - | - | 1.2 | 3.678 | 3.644 | 0.781 | 3.624 | 4.295 | -1.981 | 4.328 | 3.4 | 4.343 | 1.911 | 1.002 |
| C | - | - | - | 3.427 | 3.377 | -1.557 | 3.254 | 4.564 | 2.497 | 4.145 | 3.018 | 4.091 | 1.741 | 1.729 |
| D | - | - | - | - | -0.098 | 3.623 | 0.112 | 2.544 | 2.661 | 2.804 | 1.478 | 2.824 | 2.981 | 3.888 |
| E | - | - | - | - | - | 3.457 | -0.453 | 3.979 | 1.038 | 2.966 | 0.88 | 2.851 | 2.518 | 3.768 |
| F | - | - | - | - | - | - | 3.319 | 4.177 | 1.63 | 4.158 | 3.041 | 4.174 | 1.684 | 1.74 |
| G | - | - | - | - | - | - | - | 3.054 | 3.057 | 3.147 | 0.911 | 3.175 | 2.979 | 3.916 |
| H | - | - | - | - | - | - | - | - | 4.83 | 5.054 | 6.323 | 5.64 | 6.439 | 6.746 |
| I | - | - | - | - | - | - | - | - | - | 2.388 | 3.312 | 1.172 | 3.84 | 4.435 |
| J | - | - | - | - | - | - | - | - | - | - | 3.385 | -2.841 | 3.904 | 4.542 |
| K | - | - | - | - | - | - | - | - | - | - | - | 3.337 | 2.31 | 3.619 |
| L | - | - | - | - | - | - | - | - | - | - | - | - | 3.955 | 4.567 |
| M | - | - | - | - | - | - | - | - | - | - | - | - | - | 2.742 |
| N | - | - | - | - | - | - | - | - | - | - | - | - | - | - |

| Max | A | B | C | D | E | F | G | H | I | J | K | L | M | N |
|---|---|---|---|---|---|---|---|---|---|---|---|---|---|---|
| A | - | 2.679 | 3.011 | 0.077 | 3.164 | 1.724 | 3.127 | 1.174 | 3.29 | 1.41 | -0.212 | 2.337 | -0.516 | 2.296 |
| B | - | - | 4.009 | 2.248 | 4.027 | 3.412 | 1.139 | 3.812 | 4.221 | 2.002 | 2.637 | 0.404 | 2.97 | 3.641 |
| C | - | - | - | 3.145 | -0.312 | 2.245 | 3.981 | 0.865 | 0.742 | 3.3 | 3.083 | 3.654 | 2.836 | 1.691 |
| D | - | - | - | - | 3.351 | 2.134 | 2.917 | 1.243 | 3.487 | 0.654 | -1.298 | 2.035 | 0.876 | 2.703 |
| E | - | - | - | - | - | 2.472 | 4.048 | 2.366 | -0.575 | 3.403 | 3.185 | 3.725 | 2.959 | 2.003 |
| F | - | - | - | - | - | - | 3.546 | 0.247 | 2.722 | 2.526 | 2.054 | 3.026 | 1.337 | 0.6 |
| G | - | - | - | - | - | - | - | 3.88 | 4.38 | 2.745 | 3.101 | 1.988 | 3.366 | 3.897 |
| H | - | - | - | - | - | - | - | - | 2.585 | 2.114 | 2.373 | 3.007 | 2.093 | -0.621 |
| I | - | - | - | - | - | - | - | - | - | 3.5 | 3.346 | 3.778 | 3.137 | 2.295 |
| J | - | - | - | - | - | - | - | - | - | - | 0.856 | 1.334 | 1.752 | 2.939 |
| K | - | - | - | - | - | - | - | - | - | - | - | 2.105 | 0.722 | 2.546 |
| L | - | - | - | - | - | - | - | - | - | - | - | - | 2.618 | 3.406 |
| M | - | - | - | - | - | - | - | - | - | - | - | - | - | 2.078 |
| N | - | - | - | - | - | - | - | - | - | - | - | - | - | - |

| | |
|---|---|
| (red) | Larger than 10 |
| (orange) | Between 8 and 9,99 |
| (light orange) | Between 6 and 7,99 |
| (light green) | Between 4 and 5,99 |
| (green) | Between 2 and 3,99 |
| (dark green) | Between 0 and 1,99 |
| (grey) | NA (Calculation failed) |
