## Supplementary Data S2 for "Interlaboratory comparison of standardised metabolomics and lipidomics analyses in human and rodent blood using the MxP® Quant 500 kit": Trp.pdf

**Amount of relative differences between laboratories**

| Min | A | B | C | D | E | F | G | H | I | J | K | L | M | N |
|---|---|---|---|---|---|---|---|---|---|---|---|---|---|---|
| A | - | 3.433 | 1.952 | 0.43 | 3.404 | 3.04 | 2.628 | 3.1 | 3.13 | 2.589 | 0.368 | 1.704 | 1.961 | 1.949 |
| B | - | - | 2.981 | 3.43 | -0.657 | 1.621 | 4.267 | 1.288 | 1.288 | 2.501 | 3.801 | 3.109 | 2.962 | 2.979 |
| C | - | - | - | 1.384 | 2.799 | 2.192 | 3.395 | 2.316 | 2.427 | 1.139 | 2.44 | -0.71 | -3.477 | -4.228 |
| D | - | - | - | - | 3.208 | 2.81 | 2.924 | 2.892 | 2.985 | 2.228 | 1.47 | 0.951 | 1.442 | 1.383 |
| E | - | - | - | - | - | 1.666 | 4.281 | 1.475 | 0.772 | 2.437 | 3.763 | 3.086 | 2.967 | 2.951 |
| F | - | - | - | - | - | - | 3.974 | -1.149 | 0.34 | 1.269 | 3.38 | 2.381 | 2.222 | 2.248 |
| G | - | - | - | - | - | - | - | 3.798 | 3.864 | 3.49 | 2.08 | 3.115 | 3.245 | 3.203 |
| H | - | - | - | - | - | - | - | - | -2.314 | 1.512 | 3.445 | 2.548 | 2.361 | 2.375 |
| I | - | - | - | - | - | - | - | - | - | 1.571 | 3.433 | 2.447 | 2.326 | 2.235 |
| J | - | - | - | - | - | - | - | - | - | - | 2.977 | 1.531 | 1.055 | 1.206 |
| K | - | - | - | - | - | - | - | - | - | - | - | 2.201 | 2.374 | 2.365 |
| L | - | - | - | - | - | - | - | - | - | - | - | - | -0.807 | -0.835 |
| M | - | - | - | - | - | - | - | - | - | - | - | - | - | -3.824 |
| N | - | - | - | - | - | - | - | - | - | - | - | - | - | - |

| | |
|---|---|
| | Larger than 10 |
| | Between 8 and 9,99 |
| | Between 6 and 7,99 |
| | Between 4 and 5,99 |
| | Between 2 and 3,99 |
| | Between 0 and 1,99 |
| | NA (Calculation failed) |

| Median | A | B | C | D | E | F | G | H | I | J | K | L | M | N |
|---|---|---|---|---|---|---|---|---|---|---|---|---|---|---|
| A | - | 3.441 | 1.616 | 0.492 | 3.085 | 2.941 | 2.032 | 3.005 | 1.776 | 2.883 | 0.665 | 1.688 | 1.754 | 1.819 |
| B | - | - | 3.131 | 3.41 | 1.464 | 1.844 | 4.07 | 1.665 | 3.064 | 1.98 | 3.808 | 3.103 | 3.071 | 3.042 |
| C | - | - | - | 0.835 | 2.442 | 2.242 | 2.899 | 2.356 | -1.401 | 2.141 | 2.277 | -2.85 | -1.893 | -1.209 |
| D | - | - | - | - | 2.838 | 2.671 | 2.477 | 2.749 | 1.064 | 2.596 | 1.609 | 0.877 | 1.009 | 1.112 |
| E | - | - | - | - | - | -0.909 | 3.753 | -4.258 | 2.328 | -0.287 | 3.405 | 2.408 | 2.4 | 2.345 |
| F | - | - | - | - | - | - | 3.675 | -1.312 | 2.177 | -1.604 | 3.328 | 2.263 | 2.225 | 2.173 |
| G | - | - | - | - | - | - | - | 3.527 | 2.845 | 3.448 | 1.198 | 2.788 | 2.838 | 2.861 |
| H | - | - | - | - | - | - | - | - | 2.327 | -0.447 | 3.391 | 2.39 | 2.351 | 2.302 |
| I | - | - | - | - | - | - | - | - | - | 2.018 | 2.392 | -2.884 | -2.136 | -7.328 |
| J | - | - | - | - | - | - | - | - | - | - | 3.274 | 2.168 | 2.087 | 2.039 |
| K | - | - | - | - | - | - | - | - | - | - | - | 2.247 | 2.288 | 2.333 |
| L | - | - | - | - | - | - | - | - | - | - | - | - | -2.813 | -1.697 |
| M | - | - | - | - | - | - | - | - | - | - | - | - | - | -2.657 |
| N | - | - | - | - | - | - | - | - | - | - | - | - | - | - |

| Max | A | B | C | D | E | F | G | H | I | J | K | L | M | N |
|---|---|---|---|---|---|---|---|---|---|---|---|---|---|---|
| A | - | 3.448 | 1.262 | 0.552 | 2.424 | 2.839 | 1.187 | 2.922 | 0.479 | 3.106 | 0.912 | 1.673 | 1.523 | 1.673 |
| B | - | - | 3.244 | 3.39 | 2.561 | 2.03 | 3.876 | 1.918 | 3.777 | 1.256 | 3.816 | 3.097 | 3.168 | 3.104 |
| C | - | - | - | -0.1 | 1.669 | 2.289 | 2.265 | 2.388 | 2.049 | 2.682 | 2.093 | -0.308 | -1.181 | -0.281 |
| D | - | - | - | - | 2.02 | 2.522 | 1.932 | 2.619 | 1.666 | 2.864 | 1.735 | 0.808 | 0.422 | 0.767 |
| E | - | - | - | - | - | 1.008 | 3.057 | 1.147 | 3.003 | 1.744 | 2.928 | 1.304 | 1.509 | 1.233 |
| F | - | - | - | - | - | - | 3.355 | -1.463 | 3.286 | 0.641 | 3.275 | 2.149 | 2.228 | 2.093 |
| G | - | - | - | - | - | - | - | 3.255 | -0.924 | 3.409 | -1.47 | 2.416 | 2.299 | 2.396 |
| H | - | - | - | - | - | - | - | - | 3.283 | 0.249 | 3.334 | 2.233 | 2.341 | 2.222 |
| I | - | - | - | - | - | - | - | - | - | 3.382 | -1.767 | 2.204 | 2.12 | 2.274 |
| J | - | - | - | - | - | - | - | - | - | - | 3.521 | 2.567 | 2.661 | 2.577 |
| K | - | - | - | - | - | - | - | - | - | - | - | 2.288 | 2.201 | 2.3 |
| L | - | - | - | - | - | - | - | - | - | - | - | - | -1.899 | -4.367 |
| M | - | - | - | - | - | - | - | - | - | - | - | - | - | -1.342 |
| N | - | - | - | - | - | - | - | - | - | - | - | - | - | - |
