## Supplementary Data S2 for "Interlaboratory comparison of standardised metabolomics and lipidomics analyses in human and rodent blood using the MxP® Quant 500 kit": TrpBetaine.pdf

**Amount of relative differences between laboratories**

| Min | A | B | C | D | E | F | G | H | I | J | K | L | M | N |
|---|---|---|---|---|---|---|---|---|---|---|---|---|---|---|
| A | - | 7.159 | 3.162 | 5.468 | 8.346 | 4.914 | 2.527 | 6.251 | 5.969 | 8.008 | 4.732 | 6.347 | 2.836 | 1.577 |
| B | - | - | 4.186 | 4.446 | 8.536 | 6.093 | 7.776 | 4.988 | 4.948 | 8.261 | 3.367 | 6.433 | 5.114 | 4.345 |
| C | - | - | - | 5.228 | 8.486 | 5.424 | 4.958 | 6.093 | 5.769 | 8.2 | 4.037 | 6.319 | 4.111 | 2.587 |
| D | - | - | - | - | 8.775 | 6.559 | 9.091 | 5.726 | 4.185 | 9.094 | 5.356 | 5.609 | 5.798 | 5.361 |
| E | - | - | - | - | - | 7.135 | 10.327 | 8.571 | 7.364 | 8.863 | 7.502 | 7.058 | 6.961 | 6.965 |
| F | - | - | - | - | - | - | 8.907 | 7.533 | 6.548 | 6.638 | 6.139 | 6.537 | 3.801 | 4.53 |
| G | - | - | - | - | - | - | - | 6.219 | 5.934 | 7.477 | 4.794 | 6.61 | 1.265 | 2.912 |
| H | - | - | - | - | - | - | - | - | 5.205 | 8.638 | 4.209 | 6.044 | 5.363 | 4.793 |
| I | - | - | - | - | - | - | - | - | - | 9.561 | 6.158 | 4.078 | 6.347 | 5.995 |
| J | - | - | - | - | - | - | - | - | - | - | 6.464 | 6.789 | 4.949 | 5.287 |
| K | - | - | - | - | - | - | - | - | - | - | - | 6.21 | 4.43 | 3.267 |
| L | - | - | - | - | - | - | - | - | - | - | - | - | 7.172 | 6.928 |
| M | - | - | - | - | - | - | - | - | - | - | - | - | - | 3.35 |
| N | - | - | - | - | - | - | - | - | - | - | - | - | - | - |

| Median | A | B | C | D | E | F | G | H | I | J | K | L | M | N |
|---|---|---|---|---|---|---|---|---|---|---|---|---|---|---|
| A | - | 4.404 | 3.028 | 4.712 | 7.674 | 3.164 | 2.351 | 4.442 | 5.252 | 4.275 | 3.23 | 5.064 | 0.575 | 1.361 |
| B | - | - | 3.592 | 3.27 | 7.899 | 5.019 | 4.82 | 2.325 | 3.943 | 5.102 | 3.661 | 4.808 | 4.175 | 4.03 |
| C | - | - | - | 4.274 | 7.833 | 4.218 | 3.87 | 3.875 | 4.928 | 4.88 | -0.606 | 4.82 | 2.823 | 2.55 |
| D | - | - | - | - | 8.084 | 5.44 | 5.404 | 3.032 | 3.152 | 5.818 | 4.562 | 3.925 | 4.818 | 4.819 |
| E | - | - | - | - | - | 6.49 | 6.636 | 6.719 | 6.817 | 6.443 | 6.592 | 6.291 | 6.461 | 6.537 |
| F | - | - | - | - | - | - | 3.025 | 5.056 | 5.687 | 3.589 | 4.369 | 5.006 | 3.057 | 3.466 |
| G | - | - | - | - | - | - | - | 4.668 | 5.303 | 2.724 | 3.719 | 5.454 | 2.644 | 2.885 |
| H | - | - | - | - | - | - | - | - | 4.237 | 5.414 | 3.956 | 4.41 | 4.406 | 4.363 |
| I | - | - | - | - | - | - | - | - | - | 6.204 | 5.069 | 2.515 | 5.325 | 5.344 |
| J | - | - | - | - | - | - | - | - | - | - | 4.879 | 5.374 | 4.194 | 4.443 |
| K | - | - | - | - | - | - | - | - | - | - | - | 4.982 | 2.403 | 2.151 |
| L | - | - | - | - | - | - | - | - | - | - | - | - | 6.168 | 6.229 |
| M | - | - | - | - | - | - | - | - | - | - | - | - | - | 1.171 |
| N | - | - | - | - | - | - | - | - | - | - | - | - | - | - |

| Max | A | B | C | D | E | F | G | H | I | J | K | L | M | N |
|---|---|---|---|---|---|---|---|---|---|---|---|---|---|---|
| A | - | 3.292 | 2.811 | 2.889 | 5.85 | 3.403 | 2.514 | 3.218 | 1.448 | 3.176 | 1.561 | 2.472 | 3.578 | 0.888 |
| B | - | - | 1.707 | 1.595 | 6.293 | -0.618 | 4.136 | 0.125 | 2.801 | 0.883 | 3.84 | 4.283 | 0.997 | 3.239 |
| C | - | - | - | -1.174 | 6.126 | 1.936 | 3.776 | 1.282 | 2.103 | 1.082 | 3.413 | 3.776 | 2.401 | 2.484 |
| D | - | - | - | - | 6.124 | 1.815 | 3.829 | 1.028 | 2.176 | 1.029 | 3.481 | 3.6 | 2.322 | 2.579 |
| E | - | - | - | - | - | 5.432 | 5.052 | 5.409 | 5.242 | 5.386 | 5.153 | 5.072 | 5.457 | 5.228 |
| F | - | - | - | - | - | - | 4.189 | 0.524 | 3.195 | 0.668 | 3.914 | 4.105 | 0.614 | 3.288 |
| G | - | - | - | - | - | - | - | 3.83 | 3.046 | 3.793 | 1.437 | -5.284 | 4.064 | 2.838 |
| H | - | - | - | - | - | - | - | - | 2.836 | -2.291 | 3.743 | 3.915 | 1.567 | 3.041 |
| I | - | - | - | - | - | - | - | - | - | 2.917 | 2.675 | 1.75 | 3.308 | -0.579 |
| J | - | - | - | - | - | - | - | - | - | - | 3.682 | 3.878 | 1.617 | 3.013 |
| K | - | - | - | - | - | - | - | - | - | - | - | 1.511 | 3.849 | 2.173 |
| L | - | - | - | - | - | - | - | - | - | - | - | - | 3.968 | 2.501 |
| M | - | - | - | - | - | - | - | - | - | - | - | - | - | 3.521 |
| N | - | - | - | - | - | - | - | - | - | - | - | - | - | - |

| | |
|---|---|
| (red) | Larger than 10 |
| (orange) | Between 8 and 9,99 |
| (light orange) | Between 6 and 7,99 |
| (light green) | Between 4 and 5,99 |
| (green) | Between 2 and 3,99 |
| (dark green) | Between 0 and 1,99 |
| (grey) | NA (Calculation failed) |
